# Molecular and neural mechanisms of behavioural integration in the extended-amygdala

**DOI:** 10.1101/2024.04.29.591588

**Authors:** Simon Chang, Federica Fermani, Lianyun Huang, Steffen Schneider, Mackenzie W. Mathis, Jan M. Deussing, Na Cai

**Author notes:** These authors contributed equally to this work (Simon Chang, Federica Fermani). These authors co-supervised this work (Jan M. Deussing, Na Cai). Corresponding Authors: Helmholtz Pioneer Campus, Helmholtz Munich, Neuherberg, Germany, (S.C); (N.C). Current address: University of Regensburg, Cellular Neurobiology Group, Regensburg, Germany (Simon Chang, Federica Fermani).

## Abstract

Integration of diverse stimuli is crucial for organisms to adapt and communicate effectively, enabling overall homeostasis and survival. Studies have been performed on identifying specific neuronal encoding of individual behaviours, but how neurons integrate diverse behaviours across contexts remains elusive. Here we use Ca^2+^ imaging in freely moving mice to identify neural ensembles in the extended amygdala encoding behaviours across six distinct contexts. We found extensive flexibility in these ensemble encodings that may act as reserves for behavioural integration, with those encoding aversive stimuli showing greater specificity. Finally, we identified differential gene expression profiles between ensembles that are enriched in associations with human psychiatric and neurodegenerative disorders. Overall, our results demonstrate the molecular mechanisms behind behavioural integration, and their potential implications in health and disease.

## Main Text

Changes in the brain that regulate behavioural states such as anxiety, preference and motivation are crucial for an individual’s well-being. Rapid processing of external stimuli for the generation of appropriate behavioural responses requires a large neural network that involves multiple sensory-related circuits in the cortex as well as subcortical structures. Among these structures, the extended amygdala, which includes the central amygdala (CeA) and the bed nucleus of stria terminalis (BNST), has been suggested to play a prominent role in regulating emotion (*1–3*). Only recently, the interstitial nucleus of the posterior limb of the anterior commissure (IPAC), also part of the extended amygdala (*4*), has emerged as another region that is involved in these processes.

IPAC activity has been suggested to be associated with a range of behaviours. IPAC neurons have been shown to be involved in both innate and learned disgust showing increased expression of immediate early genes such as Fos and Arc (*5*), activated following social defeat stress and regulated by direct projections from the medial prefrontal cortex (mPFC) (*6*). In addition, neurotensin (Nts) neurons in the IPAC encode dietary preference for unhealthy energy-dense food (*7*). We previously found that corticotropin-releasing hormone (CRH) neurons in the lateral part of IPAC (IPACL) mediate locomotor activation and avoidance behaviour via basal ganglia circuits. IPACL^CRH^ neurons promote this by stimulating presynaptic CRH receptor 1 (CRHR1) localised to terminals of projections originating from the external globus pallidus (*8*). In addition, a recent study demonstrated that the IPAC integrates rewarding and environmental memory information by receiving projections from ventral tegmental area (VTA) and nucleus accumbens shell region (NAcSh) (*9*). Together, these findings demonstrate the heterogeneous and diverse role of the IPAC, suggesting that it may constitute a crucial region for integrating behavioural stimuli.

However, we do not yet have a comprehensive understanding of the neural and molecular mechanisms through which IPAC and IPACL neurons process behavioural stimuli across different contexts. Existing research on IPAC and IPACL has predominantly focused on specific sub-populations of neurons or has confined experimental subjects to a single, isolated context. These approaches were tailored towards identifying specific neural populations engaged in particular tasks, and cannot fully capture how neurons process external stimuli. This has hindered our ability to grasp the broader functionality of IPACL neurons across a spectrum of diverse scenarios and contexts.

In this paper, we adopted a more comprehensive approach, longitudinally recording neural activities in freely moving animals across six different behavioural contexts. Here we applied state-of-the-art computational pipelines for time-series calcium activity data to identify neural ensembles in the IPACL mediating mouse behavioural features in diverse contexts, then used optogenetic inhibition of identified neural ensembles to verify their specificity and relevance to the behaviours. Additionally, with viral tracing we identified the neurocircuitry these neural ensembles are a part of. For the first time, our results show that IPACL neural ensembles integrate different behavioural stimuli across contexts, undertaking divergent behavioural responses while being also dedicated for specific tasks. Finally, using single-cell transcriptomics, we found that the genes up- and down-regulated in GABA and glutamatergic neurons in the IPACL, specifically in relation to particular behavioural contexts, had been previously associated with pathological human conditions. This demonstrates how neural ensembles involved in different behaviours may have pathological consequences when dysregulated.

Taken together, our work illustrates how IPACL neural ensembles mediate emotional, preference, and social behaviours through both shared and unique mechanisms, opening up opportunities for future explorations into the precise functionality of the IPACL and its circuits in behaviour in the extended amygdala and beyond.

### Ca^2+^ imaging in IPACL neurons in behavioural contexts

To investigate the relationship between individual neural activities and behavioural responses to contexts of emotion, preference and social interaction, we used a head mounted miniature microscope to track the relative changes in Ca^2+^ activity in populations of IPACL neurons in freely moving animals (n = 16 mice, n = 1,777 neurons, mean = 117 ± 6.5 neurons per animal) (**Fig 1, A and B**). In choosing the behavioural contexts, we ensured that they encompass a diverse spectrum of behaviours. These contexts are not only widely employed for standard behavioural screening but have also been extensively used for investigating disease-related phenotypes (*10–14*). To trigger innate fear responses in mice, we used a short, sudden air-puff (Puff) on the back of the animal to mimic predator approaches. To assess how IPACL neurons respond to high or low anxiety contexts, we subjected mice to open field (OF), dark-light box (DLB) and elevated plus maze (EPM), which provide contrast in regions that reflect anxiety level of the mice. To investigate the role of IPACL neurons in preference selection, we conducted conditioned place preference (CPP) tests with one chamber paired with quinine (aversive) and the other with condensed milk (rewarding). To investigate whether IPACL has a role in social contexts like CeA and BNST (*15*, *16*), mice were subjected to a three-chamber social interaction test (SOC) with novel conspecifics. Across all six behavioural contexts, mice showed observable behaviours that were consistent with previous findings (**fig. S1, A to G**).

**Figure 1.**
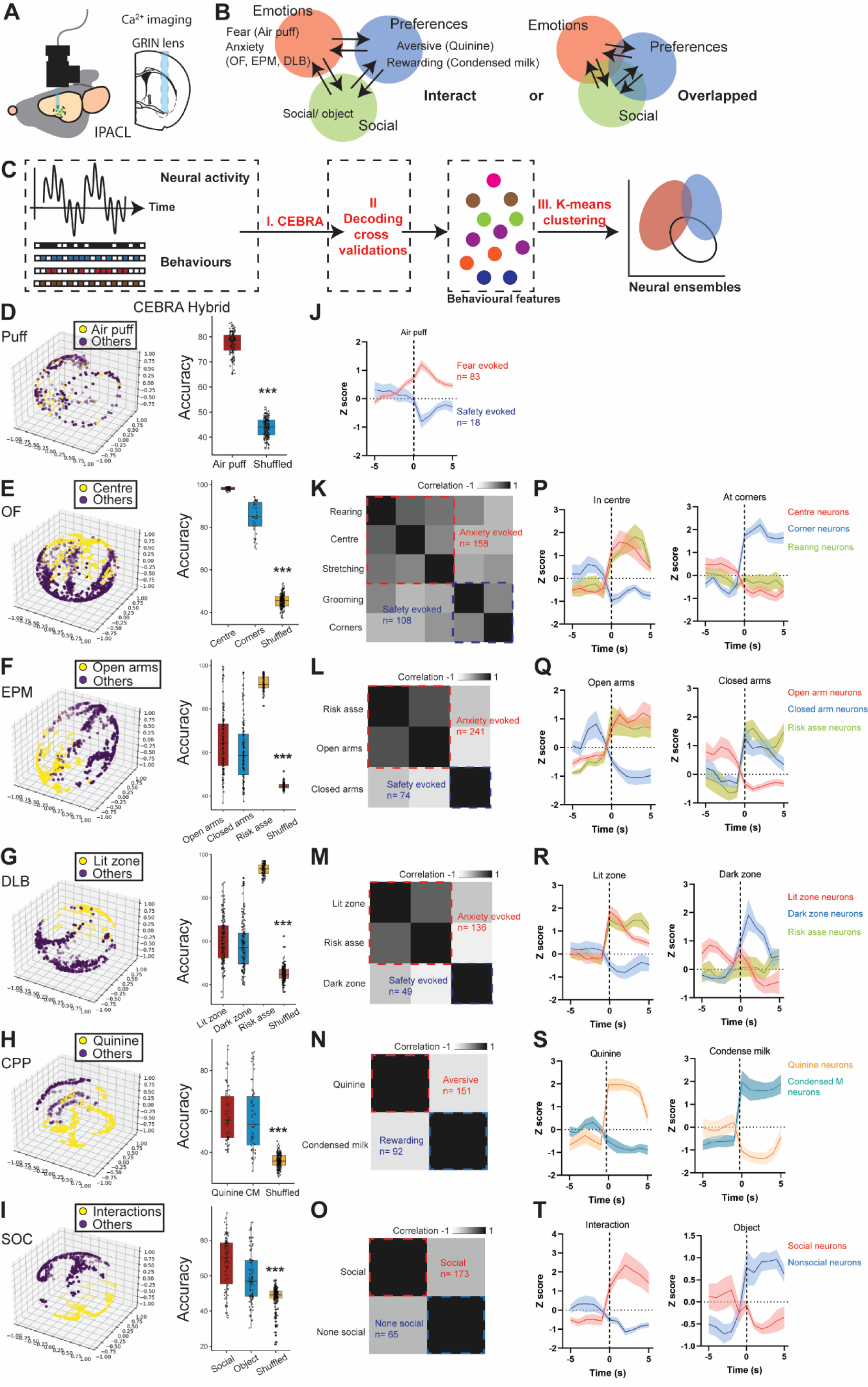
Deep brain imaging of the IPACL neurons in freely moving mice reveals encoding of different behavioural contexts. A. Scheme of *in vivo* Ca^2+^ imaging in freely moving mice in the IPACL showing virus injection and GRIN lens placement. B. We investigated the following models of behavioural integration in the IPACL: the **interaction** model, where neural ensembles in IPACL each encode for a specific behavioural context and interact only through signaling; and the **overlapped** model, where neural ensembles in the IPACL multitask and encode for more than one behavioural context. C. Workflow using computational modelling to identify neural ensembles for individual behavioural contexts: I. identification of features that contribute to the changes of neural activity in the IPACL in each behavioural context by generation and validation of CEBRA embeddings through comparison with shuffled data; II. cross-validation of CEBRA embeddings through prediction of behaviours across animals; III. K-means clustering of neural activity based on CEBRA identified and validated features into neural ensembles. D. CEBRA identifies that air puff contributes to the changes in neural activity and predicts air puff accurately compared to shuffled behavioural labels across animals (*p* < 0.0001, t = 69.44). E. CEBRA embeddings demonstrate that entering the centre zone of the OF contributes to changes of neural activity in the IPACL. CEBRA-generated embeddings predict both centre zone (98.2 ±0.12) and corners (84.8 ±1.13) entries accurately compared to shuffled (45.5 ± 0.21) behavioural labels across mice (One-way ANOVA, F_2,282_ = 4626, *p* < 0.0001. Bonferroni’s post-hoc test, centre/shuffled, *p* < 0.0001, t = 83.46; corners/shuffled, *p* < 0.0001, t = 61.71). F. CEBRA embeddings predict entry to open arms (64.9± 1.11) or closed arms (60.8± 1.13) and risk assessment (90.8± 0.38) compared to shuffled data (44.7± 0.09) (One-way ANOVA, F_3,573_ = 382.5, *p* < 0.0001. Bonferron’s post-hoc test: open arms/shuffled, *p* < 0.0001, t = 18.6; closed arms/shuffled, *p* < 0.0001, t = 14.6; risk assessment/shuffled, *p* < 0.0001, t = 32.9). G. CEBRA embeddings predict lit zone (60.8± 0.79) or dark zone (57.9± 0.85) entry, risk assessment (93.2± 0.25) compared to shuffled data (44.9± 0.23) (One-way ANOVA, F_3,637_ = 902.6, *p* < 0.0001. Bonferron’s post-hoc test: lit zone/shuffled, *p* < 0.0001, t = 19.3; dark zone/shuffled, *p* < 0.0001, t = 15.7; risk assessment/shuffled, *p* < 0.0001, t = 51.8). H. CEBRA embeddings predict aversive (58± 1.59) or rewarding (56.5± 1.99) chamber entries, compared to shuffled data (35.8± 0.23) (One-way ANOVA, F_2,326_ = 207.5, *p* < 0.0001. Bonferron’s post-hoc test: aversive/shuffled, *p* < 0.0001, t = 16.7; rewarding/shuffled, *p* < 0.0001, t = 15.4). I. CEBRA embeddings predict social (67.6± 1.42), nonsocial (58.9± 1.41) compared to shuffled data (47.9± 0.42) (One-way ANOVA, F_2,398_ = 115.4, *p* < 0.0001. Bonferroni’s post-hoc test: social/shuffled, *p* < 0.0001, t = 14.8; nonsocial/shuffled, *p* < 0.0001, t = 8.2). J. Traces showing the average Z-score of neurons during the administration of air puff, with a distinction between fear and safety responses. K. Neurons, recorded during OF, are clustered into anxiety-responding (n = 158) or safety-responding (n = 108) ensembles. L. Neurons, recorded during EPM, are clustered into anxiety (n = 241) or safety-evoked (n = 74) ensembles. M. EPM active neurons are clustered into anxiety (n = 136) and safety (n = 49) ensembles. N. Conditioned place preference neurons clustered into aversive (n = 151) or rewarding (n = 92) ensembles. O. Neurons, during social interaction, are clustered into social (n = 173) or nonsocial (n = 65) ensembles. Traces of ensembles while mice enter the centre zone or corners of the OF (P), different compartments of the EPM (Q) or the lit or dark zone of DLB (R). S. Traces of neural activities when animals stay in quinine or condensed milk conditioned chambers. T. Activities of neurons when in contact with a novel object or social partner. Values = *median ±min/max*, \*\*\**p* < 0.0001.

We simultaneously recorded spontaneous behaviours and Ca^2+^ activity, while the animals explored six behavioural contexts in random sequences to avoid generalisation of context-specific activity. We recorded Ca^2+^ activity from a comparable number of neurons across different exposures (air puff, 13% of neurons; OF, 18%; EPM, 20%; DLB, 14%; CPP, 19%; SOC, 16%) and observed a majority (76%) of these neurons showing context-dependent (not random) changes of neural activity (**fig. S2, A to F**).

### Identifying IPACL neural ensembles encoding diverse behavioural features

To investigate whether behaviours in diverse contexts contribute to changes of neural activity, we obtained embeddings for neural Ca^2+^ recordings using a recently developed machine learning algorithm that extracts neural population dynamics informed by behaviour: **C**onsistent **E**m**B**eddings of high-dimensional **R**ecordings using **A**uxiliary variables (CEBRA) (*17*) on Ca^2+^ recordings (**Materials and Methods, Fig. 1C**). We explored different CEBRA models to capture the relationship between neural activity, time and behavioural contexts (Time: making unsupervised use of only the time information in the neural recordings; Behaviour: making use of user-defined supervised behavioural labels in isolation; Hybrid: a combination of both). Since we wanted to assess if behavioural features significantly contribute to changes in neural activity over time while also considering unsupervised signals, we applied the CEBRA-Hybrid model throughout this study.

First, we found that behavioural features reflecting inner states of a mouse, such as responses to air puffs in Puff, generated quality Hybrid embeddings (**Fig. 1D, fig S2G**), with InfoNCE loss scores performing significantly better than that of a shuffled model (**fig. S2H**). In contrast, behavioural features such as speed and movement did not generate quality Hybrid embeddings, and showed similar InfoNCE loss scores as shuffled data, ruling out locomotor activity as the main driver of Ca^2+^ activity (**fig. S2, I and J**). These features generated embeddings that replicated well between mice in the same contexts (mean R^2^ = 66.3, SEM = 0.012, **fig. S2K**).

The same held true in other contexts. In OF, EPM and DLB, we found that locations and behaviours (exploratory, self-soothing and risk assessment) of mice generated quality embeddings, but not the speed of their movements (**Fig. 1, E to G, fig. S3, A to I**). In the conditioning phase of CPP (**fig S4A**) and the subsequent condition memory test, behavioural responses to the administration and memory of aversive (quinine) or rewarding (condensed milk) substances accounted for changes in neural activity, and generated quality Hybrid embeddings (**Fig. 1H, fig S4, B and C**). In SOC, both social interactions and inspection of novel objects resulted in increased neural activities (**fig. S4D**), and generated good Hybrid embeddings (**Fig. 1I, Fig. S4, E and F**).

To link neural activities to specific behavioural features, we compared how well the Hybrid-model embeddings predict specific behavioural features (cross-validated among mice) against embeddings obtained using shuffled feature labels (**Materials and Methods**). We found that the behavioural features that generate quality CEBRA embeddings were well-predicted in each context, validating their contribution to neural activity. Overall, Hybrid-model embeddings were able to predict specific behavioural features at 56-98% accuracy across all contexts, significantly better than shuffled dataset (35-48%, **Fig. 1, D to I**).

We next asked if the behavioural features associated with the Hybrid embeddings could be attributed to specific ensembles of neurons. We classified neurons into ensembles using k-Means clustering on their average neural Ca^2+^ activity in timeframes where mice demonstrated the specific behavioural features responsible to Ca^2+^ activity in each context (**Fig. 1C, fig. S5, A to J, Materials and Methods**). In general, we found that neurons cluster into two ensembles with anti-correlated neural activities in these timeframes, each encoding opposite behavioural states corresponding to the contexts, such as fear/safety and anxiety-evoked/safety-evoked (for Puff, EPM, DLB and OF), aversive/rewarding (for CPP) and social vs nonsocial (for SOC) (**Fig. 1, J to O, fig. S6, A and B**). These ensembles either exhibited distinct activation patterns associated with specific behavioural features, or demonstrated simultaneous activation across different behaviours (**Fig. 1, P to T**).

Finally, as our data showed both aversive and rewarding memories associated with neural ensembles in the IPACL, we further asked if they were programmed during the conditioning phase prior to the condition memory test. Using neural activity data from only the conditioning phase of CPP, we found that the aversive neurons identified in CPP show higher neural activity when mice receive quinine compared to condensed milk, but the rewarding neurons did not show such differences (**fig. S6, C and D**). In addition, we revealed that the non-social neurons have the highest activity when in contact with a novel object, opposite to the social neurons (f**ig. S6, E and F**). In other words, these neurons are programmed to respond to certain events such as aversive substance or social conspecifics.

### Flexibility of neural ensembles in the IPACL

Next, we asked if IPACL neurons associated with different behaviours are preserved across contexts, using longitudinal registration on neurons across all behavioural contexts. The majority (66%) of recorded neurons showed activities across behavioural contexts, while the rest showed activities unique to a single context (**Fig. 2, A and B**). We first asked if there were differences in spatial distribution of neural ensembles. Through visualising IPACL neurons under the GRIN lens in different regions of interests (ROIs) corresponding to the different contexts (**fig. S7, A to F**), we were able to quantify the distribution of neurons in each ensemble across the radius of the GRIN lens in each ROI (**fig S7, G to K**). We found no difference in spatial distribution of these neurons (**fig. S8, A to H**).

**Figure 2.**
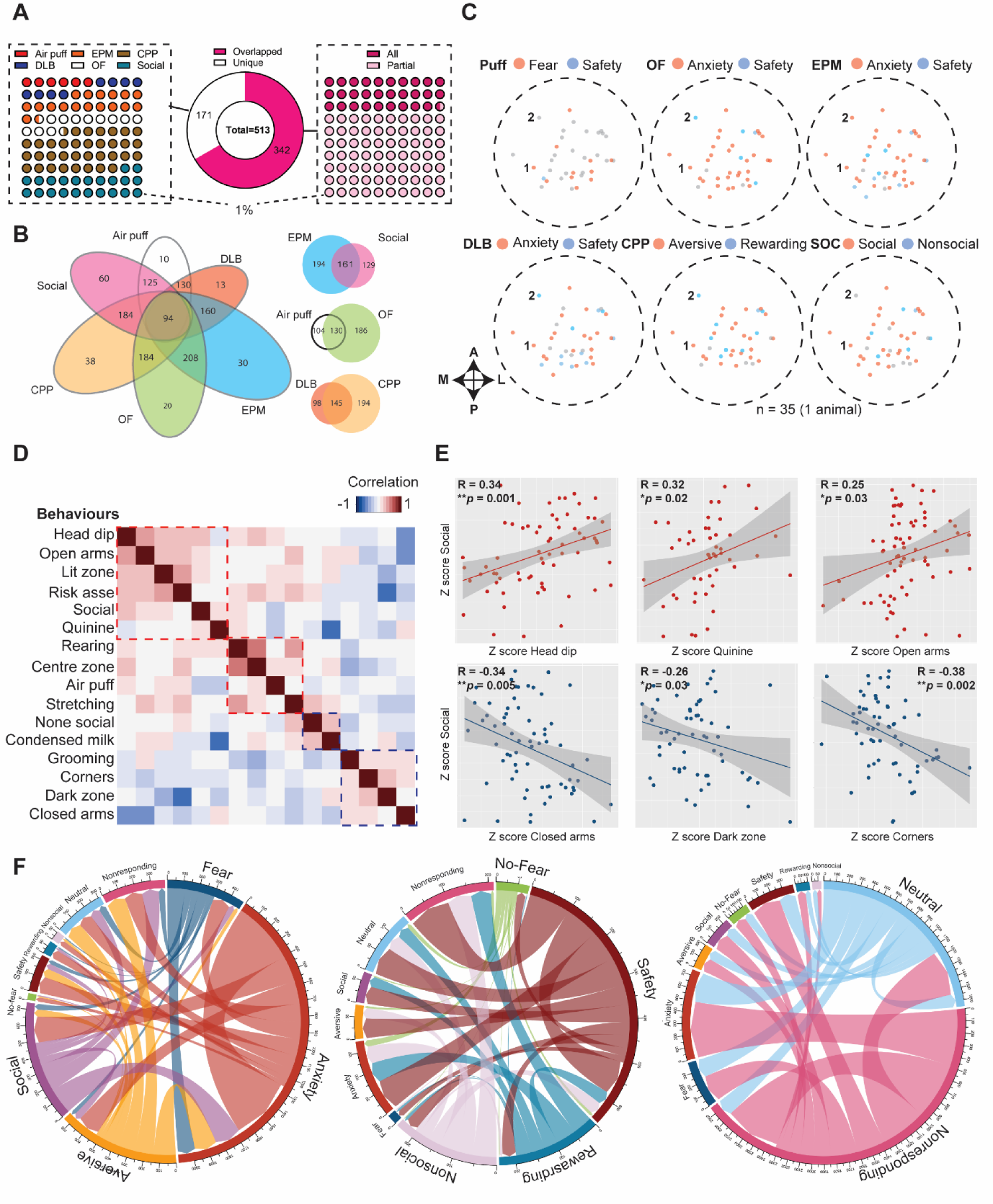
Flexible encoding of IPACL neural ensembles. A. Visual depictions of the outcomes from the longitudinal registration on neural ensembles across six behaviours. Left: percentages of unique neurons dedicated to each behaviour (each dot represents 1%). Middle: pie chart illustrating the number of neurons (342/513, 66%) that overlap in at least two tasks (magenta) and the number of neurons responding uniquely to one of the tasks (171/513) (white). Right: percentages of neurons that overlap in all six behaviours (magenta) or only partially between behaviours (pink) (each dot represents 1%). B. Numbers of neurons overlapping across all behaviours. Out of 342 neurons whose activities overlapped across tasks, 94 neurons responded to all six behaviours. C. Illustration of neurons recorded within the same ROI across various behaviours. We show that IPACL neural ensembles encode behaviours in two distinct patterns: in a flexible and interchangeable manner, as depicted by neuron 2, and in a stable way, as indicated by neuron 1. In this representation, red denotes fear, anxiety, aversive, or social neurons, blue represents safety, rewarding, nonsocial neurons, and grey signifies neutral neurons. D. Heatmap shows the correlation between activities of neural ensembles. Red boxes and blue boxes mark the ensembles encoding negative or positive experiences respectively. E. Scatter plots show that activities of neurons between social, head dip (R = 0.34, *p* = 0.001), quinine (R = 0.32, *p* = 0.02) and open arms (R = 0.25, *p* = 0.03) are positively correlated. In contrast, activities between social, closed arms (R = −0.34, *p* = 0.005), dark zone (R = −0.26, *p* = 0.03) and corners (R = −0.38, *p* = 0.002) are negatively correlated. F. Chord diagrams showing neural ensembles in the IPACL interchanging between tasks. Ensembles encoding negative experiences are more stable compared to positive or neutral experiences. \**p* < 0.05, \*\**p* < 0.01.

We further asked if neural ensembles are preserved (stable) across contexts. To do this, we inspected the ensembles in the ROIs at the single neuron level across all behavioural contexts. First, we found that some neurons preserve their ensemble identity across contexts (e.g., neuron 1 is aversive in all contexts, and therefore stable, **Fig. 2C**) while some neurons switched in between contexts (e.g., neuron 2 switches between safety and anxiety). In addition, we observed that neural activities triggered by negative experiences such as fear, anxiety and aversive substances were highly correlated. Further, although social interaction was usually indicated as a positive context, we observed that activity associated with social interaction was better correlated with negative experiences (**Fig. 2, D and E**). Finally, we quantified the neurons that were stable or switched in their ensemble identities between contexts, and found more stable neurons encoding negative experiences (52.1% remain stable) compared to the others (positive, 26.1%; neutral, 15.9% and N/A, 34.8% remain stable) (**Fig. 2F, fig. S9, A to F**). These results demonstrated that instead of completely remaining stable, many neural ensembles in the IPACL are flexible and interchangeable between contexts.

### Activity-labelled ensembles verify flexibility of neural activity across contexts

To validate the flexibility observed in IPACL ensembles, we performed activity-dependent labelling in FOSTRAP2 mice (*18*) (**Materials and Methods**) which were simultaneously monitored using *in vivo* Ca^2+^ imaging in the IPACL under the following selected contexts: elevated platform (EP) for anxiety ^14^, quinine administration (Q) for aversion, and SOC for social interaction with novel partner (**Materials and Methods**). Each of these “reduced” contexts would induce a specific behavioural feature which we previously identified as driving neural activity in wild-type (WT) mice in EPM, CPP and SOC. To validate the use of activity-dependent labelling using FOSTRAP2 mice as a means to identify neurons with increased activity, we tested if the exposure to reduced behavioural contexts triggers endogenous cFos expression in the IPACL which would be a prerequisite for activity-dependent labelling. We conducted cFos staining in the IPACL region 90 minutes after exposing test mice to the behavioural contexts, as well as control mice that remained in their home cage for the same duration (Ctrl). We found that test mice in all three contexts showed more cFos^+^ cells as compared to Ctrl mice (**fig. S10, A and B**), validating our approach. We first injected Cre-dependent GCaMP6s virus in the IPACL of FOSTRAP2 mice. After recovery, mice (n = 3-4) were subjected to EP, Q or SOC following tamoxifen injection (TAM, 100 mg/kg, 3.5 h) to label cells in an activity-dependent manner (X-TRAPed, X = context, **Fig. 3A**). We then exposed mice to the same contexts they were TRAPed in for behavioural recording and *in vivo* Ca^2+^ imaging of the TRAPed neurons 2 weeks later.

**Figure 3.**
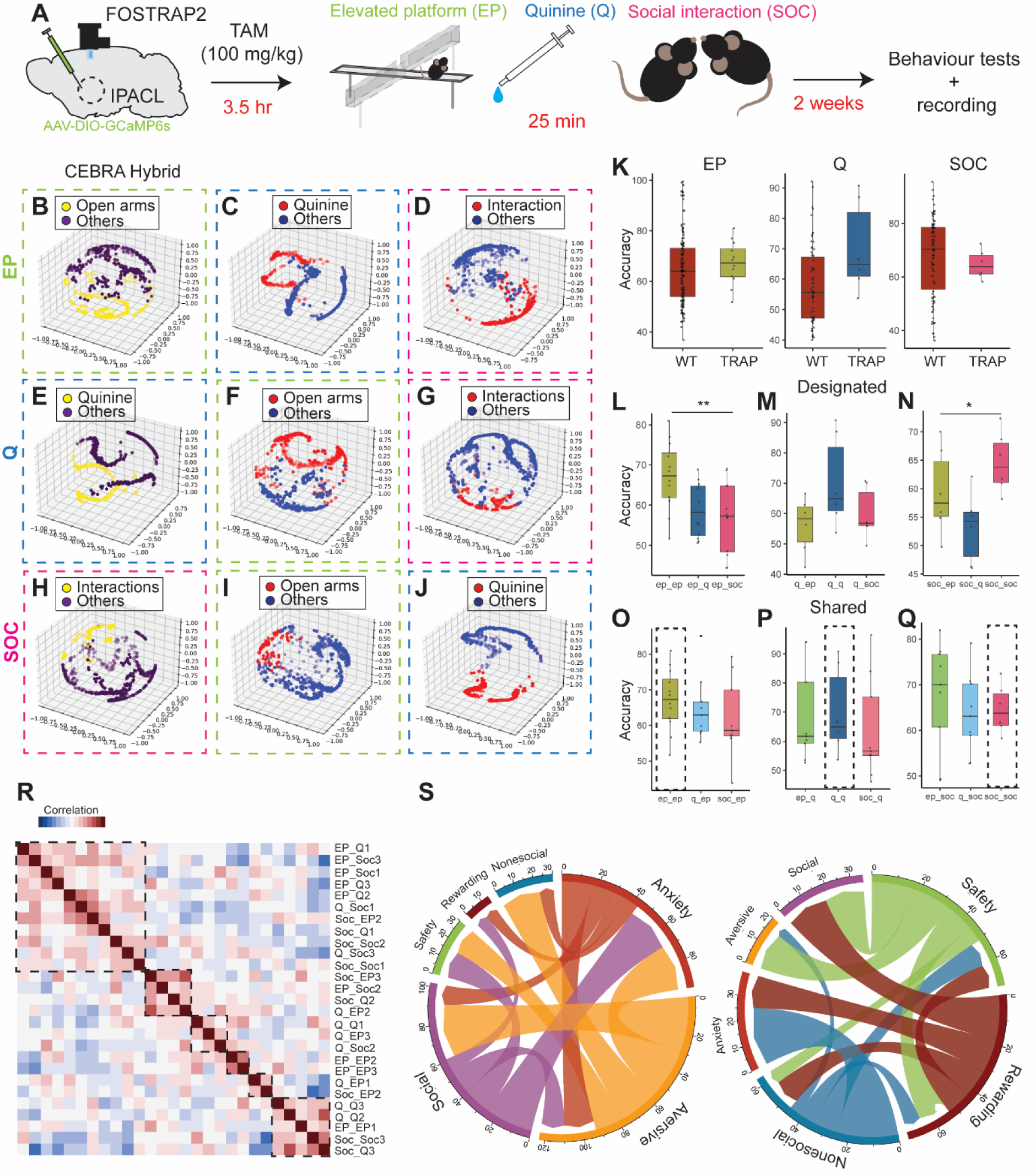
Designated ensembles show flexibility across distinct tasks. A. Scheme of experimental procedures of *in vivo* Ca^2+^ imaging on activity-dependent labelled cells. B-J. Examples of CEBRA embeddings of both activity-dependent and context-dependent ensembles along with example neural traces. B-D. CEBRA embeddings of EP-TRAPed mice in EP (dashed green line), receiving Q (dashed blue line) or in social contexts (dashed magenta line). E-G. CEBRA embeddings of Q-TRAPed animals receiving Q (dashed blue line), in EP (dashed green line) or in social contexts (dashed magenta line). H-J. CEBRA embeddings of SOC-TRAPed mice in social contexts (dashed magenta line), EP (dashed green line) or receiving Q (dashed blue line). K. Inspection of decoding accuracy between CEBRA ensembles in IPACL-wide recordings of WT and TRAPed (activity-dependent) animals. No significant difference between the two groups on decoding accuracy (EP, *p* = 0.59, t = 0.55; Q, *p* = 0.05, t = 1.96; SOC, *p* = 0.62, t = 0.49). L-Q. Behavioural decoding using either designated ensembles or shared ensembles. L-N. Designated ensembles predict specific tasks better than alternative tasks (One-way ANOVA, EP, F_2,33_ = 5.34, *p* = 0.0098; Q, F_2,15_ = 2.46, *p* = 0.12; SOC, F_2,15_ = 4.71, *p* = 0.025). O-Q. Shared ensembles decode behaviours similar to activity-dependent ensembles (One-way ANOVA. EP, F_2,27_ = 1.51, *p* = 0.23; Q, F_2,21_ = 0.39, *p* = 0.68; SOC, F_2,21_ = 0.39, *p* = 0.68). R. Heatmap of the correlation between activities of different TRAPed neural ensembles. S. Chord diagrams showing the stability of TRAPed ensembles in IPACL interchanging between tasks. Values represent *median ±min/max*, *p < 0.05, \*\**p* < 0.01.

We obtained CEBRA Hybrid embeddings on the Ca^2+^ recordings in the EP-, Q- and SOC-TRAPed neurons respectively. These embeddings represent, therefore, the neural activities of the activity-validated subset of the context-specific ensembles we previously identified using IPACL-wide Ca^2+^ recordings in WT mice, and we refer to them as designated ensembles (**Fig. 3, B, E and H**). Finally, we exposed the mice TRAPed in each context to the other two contexts for behavioural and *in vivo* Ca^2+^ recordings, to ask if the designated neural ensembles show activity across contexts. We found that Ca^2+^ activity in a subset of the TRAPed neurons, identified through clustering their CEBRA embeddings, accounted for neural activity in other contexts as well. We therefore performed longitudinal registration of activity-dependent labelled neurons across behaviours, and identified those with shared activity among contexts as shared ensembles (**EP: Fig. 3, C and D, Q: Fig. 3, F and G, SOC: Fig. 3, I and J**).

We then investigated how well designated and shared ensembles predict behaviours within or across contexts. We found that the CEBRA embeddings of the designated ensembles predicted their respective behavioural responses with high accuracy. For example, EP-based embeddings in EP-TRAPed neurons (ep_ep) predicted EP (cross-validated among mice) with 56 to 81% accuracy, similar to the prediction of mice being in open arms using IPACL-wide Ca^2+^ recordings in WT mice derived from the EPM context (**Fig. 3K**). In addition, they showed better accuracy at predicting EP compared to shared ensembles, such as Q-or SOC-based embeddings obtained in EP-TRAPed neurons (ep_q, 50-65%; ep_soc, 44-69%, **Fig. 3L**). We found similar results for Q-TRAPed and SOC-TRAPed neurons (**Fig. 3, M and N**).

Interestingly, we found that shared ensembles predicted behavioural contexts in which the Ca^2+^ signals were obtained for embeddings, regardless of whether they were TRAPed in the same contexts. For example, Q-based embeddings obtained in EP-TRAPed neurons (ep_q) predicted Q at 59 to 94% of accuracy, similar to the results of Q-based embeddings obtained in Q-TRAPed neurons (designated ensemble in q_q, 60-90%). This held true for EP- and SOC-based embeddings (**Fig. 3, O to Q**). This was consistent with our finding that neural activity between designated and shared ensembles were correlated demonstrating the neural ensembles were conserved across contexts (**Fig. 3R**). In addition, we also observed consistent similarity between these ensembles (**fig. S10C**).

Finally, we quantified the percentage of neurons showing unique or shared activities across contexts. We found that 44%, 42% and 42% of neurons respond uniquely to EP, Q or SOC respectively. There were more neurons encoding negative experiences (64%) including anxiety, aversive and social interaction compared to the positive ones (36%, **fig S10, D to F**). Consistent with our previous findings using IPACL-wide Ca^2+^ recordings in WT mice, we found that a greater percentage of the activity-dependent neural ensembles we identified as encoding negative experiences were stable (64.8%), as compared to neurons encoding positive experiences (37.3%, **Fig. 3S, fig. S10G**). We further ensured that our results were not due to non-specific labelling of cells (**fig. S11, A and B**). These results verified our previous findings that while IPACL neural ensembles showed behaviour specific activities, they also exhibited a high level of flexibility when exposed to different contexts.

### Specific modulation of behaviours by designated neural ensembles

To conclusively show that the neural ensembles which we identified through the Ca^2+^ recordings contribute directly to behavioural outcomes, we optogenetically inhibited context-TRAPed neurons (595 nm, constant light for 5 min) in FOSTRAP2 mice (n = 6-10) injected with a Cre-dependent eNpHR3.0 AAV in the IPACL, and compared their behaviours in different contexts to those of EYFP injected mice (Ctrl) (**Fig. 4A, Materials and Methods**). To make sure differences in the behaviours of test (eNpHR3.0) and Ctrl mice were not due to viral constructs and efficiency of virus distribution, we verified that the numbers of YFP^+^ cells were similar between eNpHR3.0 and Ctrl animals (**fig. S12A**). We subjected eNpHR3.0 and Ctrl animals to DLB and OF to assess anxiety, to real-time place preference (RTPP) and CPP for preference, and to SOC for sociability (**Fig. 4A**).

**Figure 4.**
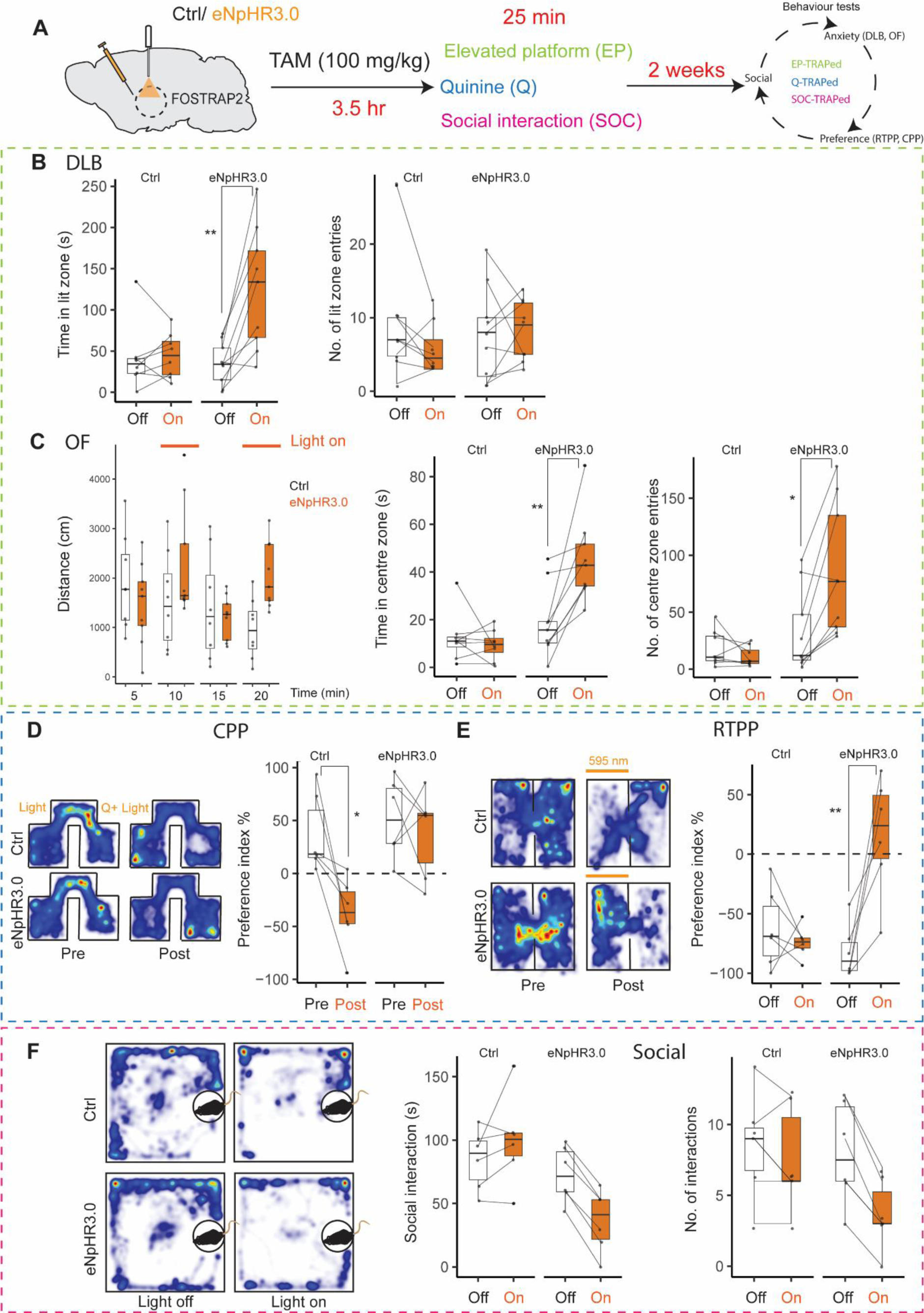
Contribution of activity-dependent ensembles to behaviours. A. Scheme showing the strategy of virus injection and behavioural timeline. Either AAV-hSyn-DIO-EYFP (Ctrl) or AAV-hSyn-DIO-eNpHR3.0 (eNpHR3.0) viruses were injected bilaterally in the IPACL of FOSTRAP2 mice. Two weeks after virus injection and optic fibre placement mice were exposed to different contexts following TAM (100 mg/kg) injection. Mice were subjected to different behavioural tests two weeks after context exposure. B. eNpHR3.0 EP-TRAPed mice (n = 10) show anxiolytic behaviour, spending more time in the lit zone during the light-on period (Two-way ANOVA, F_1,30_= 5.35, *p* = 0.027. Bonferroni’s post-hoc test, *p* = 0.0016, t = 4.12) of DLB, without significant difference in the number of entries into the lit zone (Two-way ANOVA, F_1,30_= 0.12, *p* = 0.73) compared to Ctrl mice (n = 9) when light is on. C. No significant differences were observed in locomotor activity between Ctrl and eNpHR3.0 EP-TRAPed mice (Two-way RM ANOVA, F_1,15_= 0.86, *p* = 0.37) during the open field test. eNpHR3.0 mice show lower anxiety levels as evidenced by increased time in the centre zone (Two-way ANOVA, F_1,30_= 21.6, *p* < 0.0001. Bonferroni post-hoc test, *p* = 0.001, t = 4.2) and by more entries into the centre zone (Two-way ANOVA, F_1,30_= 12.9, *p* = 0.001. Bonferroni’s post-hoc test, *p* = 0.024, t = 3.11) during the open field test compared to ctrl mice. D. Heatmap and boxplots showing that inhibition of Q-TRAPed ensemble in the IPACL results in place preference of the eNpHR3.0 mice in RTPP compared to Ctrl mice (Two-way ANOVA, F_1,20_= 10.77, *p* = 0.004. Bonferroni’s post-hoc test, eNpHR3.0, *p* = 0.0002, t = 5.24). E. Heatmap and box plots illustrate that inhibition of Q-TRAPed neurons in the IPACL reduces the aversive response to quinine in eNpHR3.0 mice, whereas Ctrl mice continue to avoid the chamber paired with quinine (Two-way ANOVA, F_1,20_= 8.8, *p* = 0.007. Bonferroni’s post-hoc test, Ctrl, *p* = 0.015, t = 3.5; eNPHR3.0, *p* > 0.99, t = 0.63). F. Heatmap and boxplots revealing that inhibition of SOC-TRAPed neurons leads to a modest reduction in sociability among eNPHR3.0 mice, manifested by a decrease in both the duration of social interaction and the number of interactions, however, there is no significant difference compared to Ctrl mice (social interaction, Two-way ANOVA, F_1,20_= 0.96, *p* = 0.33; No. of interactions, Two-way ANOVA, F_1,20_= 3.64, *p* = 0.07). Values represent *median ±min/max*, \**p* < 0.05, \*\**p* < 0.01.

Inhibition of EP-TRAPed neurons resulted in eNpHR3.0 mice staying longer in the lit zone of the DLB, though not entering the lit zone more often, as compared to Ctrl mice (**Fig. 4B**). Similarly in the OF, eNpHR3.0 mice entered the centre zone more frequently and spent longer time in that zone compared to Ctrl mice, while there was no difference in the total distance travelled between the two groups (**Fig. 4C**). eNpHR3.0 mice did not differ from Ctrls with regards to the time they spent on social interaction, or the number of interactions they had in SOC (**fig. S12B**). They also did not show any difference in behaviour in the RTPP or their avoidance of quinine in the CPP compared to Ctrl mice (**fig. S12, C and D**). These results suggest that the inhibition of EP-TRAPed neurons in the IPACL was specifically anxiolytic, and had no effects on locomotor function, social interaction, or aversive conditioning.

We found similar specificity when inhibiting other neural ensembles. For example, inhibition of Q-TRAPed ensembles which were trapped in conjunction with quinine administration resulted in the elimination of quinine avoidance in eNpHR3.0 animals, while Ctrl animals remained avoidant in the CPP (**Fig. 4D, Material and Methods**). The same eNpHR3.0 animals showed a strong place preference for the side where light was applied in RTPP, while Ctrl animals did not exhibit any preference (**Fig. 4E**). In contrast, inhibition of quinine-responding ensembles did not affect anxiety nor sociability (**fig. S13, A to C**). This suggests that the Q-TRAPed ensembles induced a generally aversive response and avoidance-related behavioural patterns.

Finally, we assessed the role of SOC-TRAPed ensembles in anxiety, preference, and sociability. Inhibiting SOC-TRAPed cells reduced the duration of interaction and the overall number of interactions in SOC, as compared to Ctrl animals (**Fig. 4F**). In addition, we did not observe any change in anxiety levels or preference when inhibiting SOC-TRAPed cells (**fig. S14, A to D**). This suggests that SOC-TRAPed ensembles might specifically affect social but not other behaviours. Overall, these findings conclusively demonstrated that we had correctly identified the designated ensembles for specific behaviours. Further, the specificity of our findings suggests that the flexibility of neural ensembles shown in previous experiments primarily served as reserves in neural networks that allowed for behavioural integration.

We therefore sought to understand how neurons of different IPACL ensembles were connected with other brain regions by mapping their connections. To map the inputs into IPACL, we injected a Cre-dependent retrograde virus into FOSTRAP2 mice. We then subjected the mice to various behavioural contexts (**Fig 5A, Materials and Methods**). In addition to behaviour-specific inputs from regions involved in appetitive behaviours such as PBN to quinine ensembles, we found that neurons from distinct neural ensembles shared common inputs originating from regions recognized for behavioural integration, including the basolateral amygdala (BLA), prefrontal cortex (PFC), anterior insular cortex (AIC), and entorhinal cortex (ECT, **Fig 5, B to E**). While we found common regions innervating these neural ensembles, the strength of inputs from shared brain regions varied among different neural ensembles. This variability in input strength could potentially indicate the diverse roles played by these neurons in responding to different stimuli.

**Figure 5.**
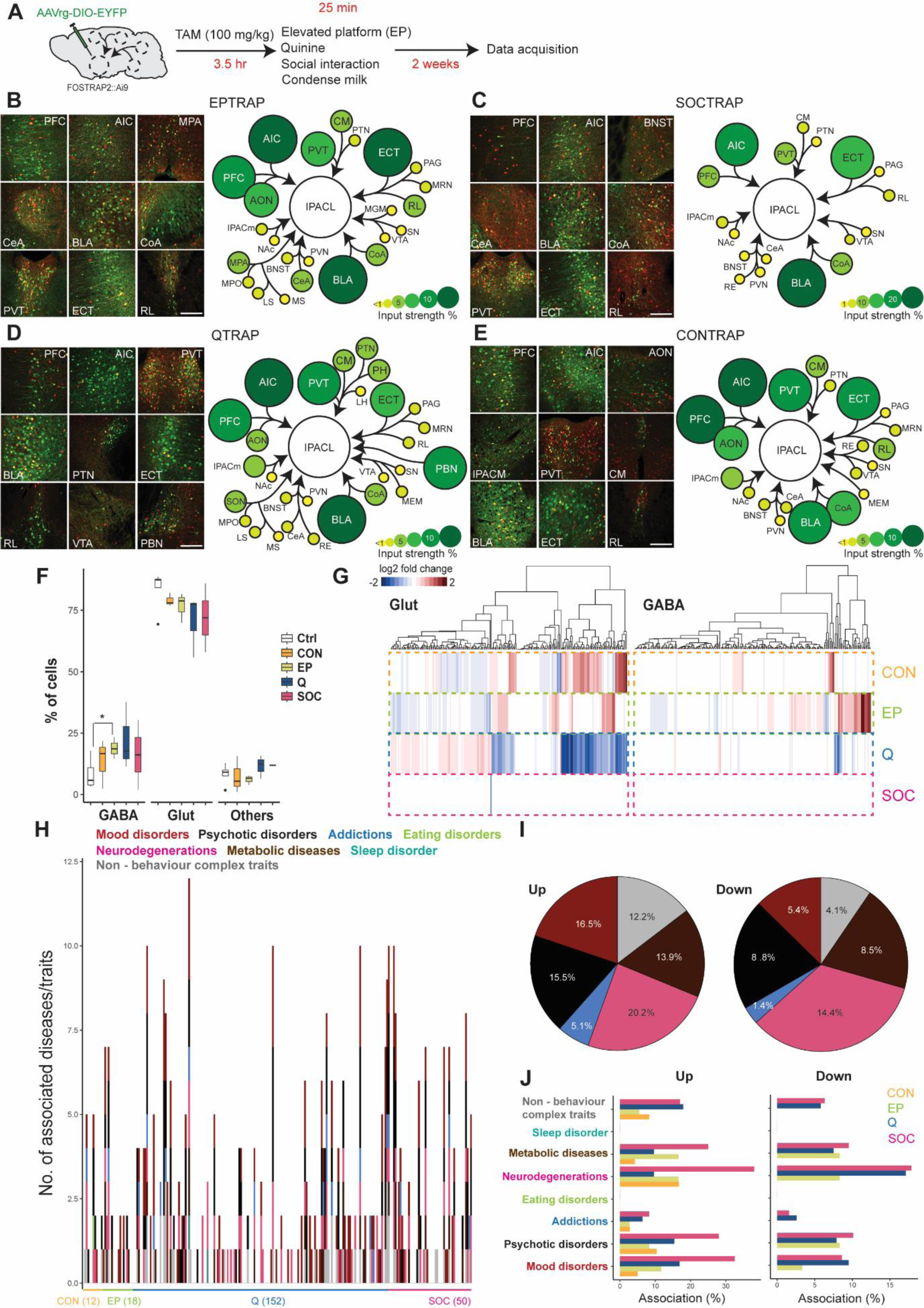
Networks and molecular mechanisms of behavioural integration. A. Scheme of the approach used for retrograde tracing on activity-dependent ensembles. B-E. Representative images of brain regions sending inputs into IPACL-TRAPed ensembles. Circle plots show the percentage of cells quantified in different brain regions, size and colours of the circles demonstrate the strength of input. Scale bars = 200 µm. F. Box plots demonstrate the distribution of different subtypes of cells. There is no difference in percentage of Glut (One-way ANOVA *p* = 0.44) and other cell types between Ctrl and context exposures. However, there are significantly more GABAergic neurons in EP compared to Ctrl (*p* = 0.043, t = 2.58). G. Heatmap demonstrates the level of differentially expressed genes in 4 conditions H. Number of associations with human diseases/complex traits in Glut neurons are shown with stacked bars. I. Average percentages of up-or down-regulated genes associated with different categories of traits and disease in Glut neurons. J. Bar graphs showing the percentages of up and down regulated genes associated with diseases in all four context exposures in Glut neurons. Abbreviation of brain regions: AIC, Anterior insula cortex; AON, anterior olfactory nucleus; BLA, basolateral amygdala; BNST, bed nucleus of the stria terminalis; CeA, central nucleus of amygdala; CM, Central medial thalamic nucleus; CoA, cortical amygdalar area; ECT, ectorhinal area; IPACm, interstitial nucleus of posterior limb of anterior commissure, medial; MEM, median eminence; MGM, magnocellular nucleus of medial geniculate body; MPA, medial preoptic nucleus; MPO, medial preoptic nucleus; MS, medial septal nucleus; MRN, median raphe; NAc, nucleus accumbens; LH, lateral hypothalamus; LS, lateral septal nucleus; PAG, periaqueductal gray; PBN, parabrachial nucleus; PFC, prefrontal cortex; PH, posterior hypothalamic area; PTN, parathalamic nucleus; PVN, periventricular nucleus; PVT, paraventricular thalamic nucleus; RE, reuniens thalamic nucleus; RL, rostral linear nucleus of the raphe; SN, substantia nigra; SON, supraoptic nucleus; VTA, ventral tegmental area. Scale bar = 200µm. Values represent *median ±min/max*, \**p* < 0.05.

### Molecular mechanisms of behavioural integration in the IPACL

Finally, we investigated the molecular mechanisms driving both the specific activities and those underlying their roles in behavioural integration. To this end, we performed single-cell RNA sequencing (scRNA-Seq) on FACs-sorted TRAPed neurons identified in FOSTRAP2 animals subjected to EP, SOC, Q (for aversion) and condensed milk CPP (CON, for reward, **Materials and Methods, fig. S15, A and B**). In order to assess whether c-Fos expressing cells are representative of the cell population in IPACL, we also performed scRNA-Seq on a combination of all non-activated cells (Ctrl) across all four behavioural contexts collected from FACs (**Materials and Methods**).

We first assessed the cellular compositions of the activated cells in each context, as well as all non-activated cells. Mapping each of our samples to the mouse brain reference atlases in MapMyCells (RRID:SCR_024672), we obtained an estimate of the percentage of cell types present in each sample, and compared our findings across samples obtained in different behavioural contexts. We found no difference in percentages of glutamatergic (Glut) cells between the Ctrl and activated samples, and the percentages of GABAergic (GABA) cells in EP were significantly higher than in Ctrls. We did not, however, observe significant differences between cellular compositions between all four behavioural contexts (**Fig. 5F**). This was consistent with our previous finding that neural activities are flexible, and the neural ensembles that were activated in different behavioural contexts overlap.

We then asked if there were differentially expressed genes among the Glut and GABA neurons that were activated in different behavioural contexts. We performed differential gene expression (DGE) analysis between each pair of contexts, using pseudo-bulk gene expression per sample per cell type (**Materials and Methods**). We identified 86 significantly up- and 293 down-regulated genes among behaviours in total (**table S1-S6**), and found that 48.9% and 73% of those were significant in more than one test. In other words, genes that were up-or down-regulated in one behaviour as compared to another, were also likely to be similarly up-or down-regulated in a third behaviour that shared neural ensembles and activities. To identify the uniquely up-or down-regulated genes per behavioural context, we further performed DGE analysis between pseudo-bulk gene expressions among Glut and GABA cells activated in each context, against those in all other contexts (**Materials and Methods**). We found 1-11 up-regulated genes and 1-33 down-regulated genes unique to each behavioural context across both cell types (**Fig. 5G, fig. S15, C and D and fig. S16, A to D, table S7-S10**).

Finally, we asked whether the genes up-or down-regulated in these neural ensembles had previously been found to be associated with human complex traits and diseases. We selected 21 human conditions which we grouped into 8 categories: mood disorders, psychotic disorders, addictions, eating disorders, neurodegeneration, metabolic diseases, sleeping disorders and non-behavioural traits. We obtained the previously found genetic associations between the genes we identified in DEG analysis (as up-or down-regulated in each context, as opposed to one or all other contexts) with these diseases and complex traits using the Harmonizome(*19*) database (**Materials and Methods**), and summarised our findings in **Fig. 5H and fig. S17A**. We found that the significant up-or down-regulated genes were associated with disease categories in a context-dependent manner. For instance, genes up-regulated in both EP- and Q-TRAPed Glut and GABA neurons were associated more with stress-related disorders (Glut: Cat I, 14.4%; others, 7.1%; GABA: Cat I, 27.8%; others, 20.1%) (**Fig. 5, I and J, fig. S17, B and C**).

## Discussion

Rapid processing of diverse environmental signals that pose threat or indicate precious resources is essential for survival. In our present study, we used a combination of computational, behavioural, and molecular approaches to unravel how neurons within the IPACL integrate and process behaviours across different contexts. Our results showed that these neurons encode a range of behaviours, including innate fear, anxiety, aversion and reward responses and associative memories, as well as responses to novelty and social interactions. While these functions are well-aligned with the roles of other regions in the amygdala (*20–22*) and extended amygdala (*23–28*), we showed, for the first time, a side-by-side comparison of IPACL neural function between contexts. Additionally, we assessed these behaviours in freely moving animals, providing a more ecologically valid understanding of how the brain responds to diverse stimuli.

We observed a greater number of neurons that encode negative experiences such as fear and anxiety, supporting the view that IPACL neurons are more sensitive to stressful stimuli (*6*). We further identified a bidirectional regulation of preferences in the IPACL, which is consistent with previous findings suggesting that IPAC^Nts^ neurons play a role in both dietary preference for unhealthy, energy-dense food (*7*) and reactions of innate and learned disgust (*5*, *7*). Finally, we found that the activity of social neurons is more similar to those that encode fear and anxiety, highlighting the potential role of IPACL in negative social valence (*29*) rather than reward (*30*). These results contributed to our understanding of social behaviour and emotional processing within this brain region.

With our multi-context setup, we were able to investigate and demonstrate, for the first time, the sharing and flexibility of neural encodings in a large portion of IPACL neurons across contexts. Of note, we found that this flexibility was not common to all neural activity - ensembles associated with negative memories consistently exhibited stable imprinting and displayed stronger activation while exposed to aversive contexts, while ensembles linked to rewarding memories tended to be more generalised in their responses. This was first corroborated by our results from activity-dependent labelled neurons in FOSTRAP2 mice, which showed that while IPACL neurons display behaviour-specific activities, they mediated other behaviours too. We then identified a further nuance through optogenetically inhibiting designated ensembles in the IPACL, which pointed to the reserve nature of this flexibility.

Through viral tracing, we found that IPACL neurons receive inputs from regions that are involved in a wide range of processes (*23*, *31–37*, *40*, *41*). While previous studies found the output neural circuits are functionally biased (*7*, *34*, *38–41*), we demonstrated that the strengths of inputs into IPACL ensembles are both region- and context-dependent. This diversity underscores the nuanced and context-dependent nature of IPACL functioning, shedding light on how the same subpopulation of neurons can play a role in facilitating opposite or contrasting behaviours. This complexity and adaptability of neural circuits challenges traditional notions of fixed neural functions.

Overall, our results indicated that IPACL neurons differ from those observed in other brain structures, highlighting the potential importance of IPACL as a hub for behavioural integration. For instance, our observed pattern of flexible encoding is not comparable to the CeA, where neurons display a higher degree of specificity (*38*). It further distinguishes the IPACL from recent findings related to the activities of serotonergic neurons in the hindbrain, which are preserved in various contexts, and do not exhibit flexibility across opposite valances (*39*).

We noticed that in our scRNA-seq data we captured a large amount of glutamatergic cells, which might be due to neurons being labelled as a consequence of cFOS expression (*42*). Nevertheless, our scRNA-seq analysis identified behaviour-relevant genes that were associated with human diseases such as major depression, ASD, addiction and AD. This finding suggests a potential link between mechanisms of behavioural regulation and human diseases, indicating that dysregulation of genes involved in fundamental behaviours might contribute to the development of a spectrum of psychiatric, metabolic, neurodevelopmental, and neurodegenerative pathologies.

Our study contributes to advancing the knowledge of the complex integration of neural signal and behavioural encoding that occurs in the IPACL. Nevertheless, an important aspect that remains unexplored pertains to the contribution of input circuits to the formation of these neural ensembles and their subsequent impact on behaviours. Future studies will need to disentangle how different IPACL circuits are recruited and coordinate to participate in behavioural integration.

## Acknowledgements

We would like to thank A. Varga, head of the animal facility, and staff for the dedicated support with animal care; S. Unkmeir, S. Bauer, and the scientific core unit Genetically Engineered Mouse Models for genotyping support from Max Planck Institute of Psychiatry. We would also like to thank Dr. Inti de la Rosa Velazquez and G. Eckstein for performing the NovaSeq sequencing at the Bioinformatics Core Facility of Helmholtz Munich and T. Walzthöni for bioinformatics support. We would like to thank Prof. Dr. Jonathan Flint, Dr. Rohit Menon for their critical comments which improved this manuscript.

## Fundings

This research is supported by Vallee Foundation to M.W.M and Google PhD Fellowship to S.S.

## Author contributions

Conceptualization: S.C., F.F., and N.C. Methodology: S.S., and M.W.M., and N.C. Investigation: S.C., F.F., L.H., S.S., M.W.M., and N.C. Data analysis: S.C., F.F., L.H., and N.C. Visualization: S.C., F.F., L.H., and N.C. Project administration: S.C., and N.C. Supervision: M.W.M, J.M.D., and N.C. Writing (original draft): S.C., F.F., L.H., and N.C. Writing (review and editing): S.C., F.F., L.H., S.S., M.W.M., J.M.D., and N.C.

## Conflict of interests

All authors declared there is no conflict of interest.

## Data and Materials availability

Custom code for single cell analysis is available at: https://github.com/caina89/calciumIPACL. Behaviour videos, calcium recordings and additional histology images are available from the corresponding authors upon reasonable request. They are not deposited to a public database owing to their large size and size limitation of online depositories.

## Supplementary Materials

Materials and Methods Figs. S1 to S18

Tables S1 to S10

References (43–51)

## Notes

### Competing Interest Statement

The authors have declared no competing interest.

