## Supplementary materials for "Molecular and neural mechanisms of behavioural integration in the extended-amygdala"

**The PDF file includes:**

Materials and Methods

Figs. S1 to S18

Tables S1 to S10

References (43-51)

**Materials and Methods**

Mice

All animal experiments are conducted with the approval of and in accordance with the Guide of the Care and Use of Laboratory Animals of the Government of Upper Bavaria, Germany. Mice are group-housed under standard laboratory conditions (22° ± 1°C, 55 ± 5% humidity) and maintained under a 12-hour light/12-hour dark cycle with food and water ad libitum. Experiments are conducted with adult male mice or equally mixed gender groups of mice (age, 2 to 5 months). The following mouse lines are used: C57/BL6NRj, $Fos^{2A\text{-}iCreER}$ (FOSTRAP2) [$Fos^{tm2.1(icre/ERT2)Luo}$/J, the Jackson Laboratory, stock no. 030323], Ai9 [$Gt(ROSA)26Sor^{tm9(CAG\text{-}tdTomato)Hze}$, the Jackson Laboratory, stock no. 007909]. The FOSTRAP2::Ai9 double transgenic line was generated by cross-breeding of single transgenic lines. Primer sequences and protocols for genotyping are available upon request.

**Stereotactic surgeries**

Mice are anesthetized for surgery with isoflurane (Floren, Abbott), 2% (v/v) in $O_2$, and placed in a stereotaxic frame (Kopf Instruments). Body temperature is maintained with a heating pad. Metacam (5 mg/kg of body weight) is administered as a systemic analgesic. Viruses are delivered using a 33-gauge microinjection needle with a 2.5-µl Hamilton syringe (Hamilton) coupled to an automated microinjection pump (World Precision Instruments Inc.) at 100 nl/min. Post surgery recovery involves Metacam supplementation (5 mg/kg, subcutaneous injection) for 3 days after surgery, with daily monitoring of food intake. For in vivo calcium imaging recordings 0.5 ul of virus $AAV_5$-hSyn-GCaMP6s or $AAV_5$-hSyn-DIO-GCaMP6s (~$10^{12}$ vg/ml; UPenn Vector Core) are injected unilaterally in the IPACL (AP: 0.26, ML: 2, DV: −5). For optogenetic experiments $AAV_5$-Ef1a-DIO-eNpHR3.0-EYFP, or $AAV_5$-Ef1a-DIO-EYFP virus (~$10^{12}$ vg/ml; UNC Vector Core) are injected bilaterally in IPACL. For retrograde tracing AAVrg-hSyn-DIO-EYFP (~$10^{13}$ vg/ml; Addgene) is injected unilaterally in the IPACL.

**GRIN lens implantation, baseplate fixation and optic fibre implantation**

The gradient index (GRIN) lens is implanted 1 week after viral injection ($AAV_5$-hSyn-GCaMP6s/$AAV_5$-hSyn-DIO-GCaMP6s) in the IPACL. Mice are anaesthetised as described above. Debris is removed from the hole and a customised blunted 20G needle (1 mm in diameter) slowly lowered down into the brain (AP: 0.26, ML: 2, DV: -4.8 mm). After retraction of the needle, a GRIN lens (ProView lens; diameter, 0.6 mm; length, ~7.4 mm, Inscopix) mounted on a GRIN lens-holder is slowly implanted above the IPACL. The exposed skull is covered with dental acrylic (Paladur, Heraeus). Two weeks after GRIN lens implantation, mice are anaesthetised and placed in the stereotaxic frame. A baseplate (BPL-2, Inscopix) attached to a miniature microscope (Inscopix) is positioned above the GRIN lens. The focal plane is adjusted until neural structures and GCaMP6s dynamics are observed. The baseplate is fixed using C&B-Metabond (Parkell).

In optogenetic experiments, an optic fibre (200 µm core, 0.39 NA, 1.25 mm ferrule, Thorlabs) is implanted bilaterally above the IPACL (AP: 0.26, ML: 2, DV: -4.8 mm) two weeks after virus injection. Implants are secured with cyanoacrylic glue (Braun), and the exposed skull is covered with dental acrylic (Paladur, Heraeus). The animals are left for two weeks for recovery before subjecting them to behavioural tests.

**Description of behavioural contexts and procedures for calcium imaging and optogenetic experiments**

Mice were single-housed for a week and entered the respective test apparatus in random order. Mice rested for 1 day between each behavioural test, except in the conditioned place preference test. All experiments were analysed using the automated video tracking Ethovision (Noldus, Wageningen).

Air puff (Puff): mice were placed in an open arena [40 cm (*l*) × 40 cm (*w*) × 60 cm (*h*)] for 5 min. Every 30 sec an air puff was delivered on the back of the mice.

Open field (OF): the test was conducted in an evenly illuminated (<15 lx) square apparatus [40 cm (*l*) × 40 cm (*w*) × 60 cm (*h*)]. The test duration was 20 min. Mice received a constant 595 nm (Thorlabs, M595F2) with 10 mW photoinhibition. The photoinhibition followed a pattern of 5 minutes off/on sequence starting with light off.

Dark-light box (DLB): the test was performed for 10 min in an apparatus consisting of a secure black compartment [10 cm (*l*) × 20 cm (*w*) × 35 cm (*h*), <5 lx] and an aversive, brightly illuminated white compartment [10 cm (*l*) × 20 cm (*w*) × 35 cm (*h*), 400 lx] connected by a door. The photoinhibition occurred during the last 5 minutes.

Elevated Plus Maze (EPM): mice were placed in the centre area of a plus-shaped platform with four intersecting arms, elevated 37 cm above the floor. Two opposing open (30 × 5 cm) and closed (30 × 5 × 15 cm) arms were connected by a central zone (5 × 5 cm). Mice were free to explore the maze for 15 min.

Real time place preference (RTPP): in the RTPP, mice were subjected to a two-chamber arena (40 × 40 × 25(h) with openings between the two chambers (20 cm each) for 10 min. One counterbalanced side of the chamber was assigned as the stimulation side. The light-emitting diode (595 nm) was triggered on the basis of the location of the animal using primmax software and controllers. In the photostimulated side of the arena, which was assigned on the basis of the initial preference, mice received a constant 595 nm with 10 mW of power at the fibre tips.

Conditioned Place Preference (CPP): the CPP apparatus [T shape, 40 cm (*l*) × 40 cm (*w*) × 35 cm (*h*)] consisted of a starting box and two conditioning boxes made of black acrylic boards. The conditioning compartments [18 cm (*l*) × 20 cm (*w*) × 35 cm (*h*)] were separated by a smaller compartment [10 cm (*l*) × 20 cm (*w*) × 35 cm (*h*)] in the middle. The left compartment had mosaic walls and smooth flooring, while the right compartment had black walls and thin-grid flooring. The experiment consisted of three phases: (i) preconditioning, (ii) conditioning, and (iii) postconditioning phase. During the preconditioning phase (pretest trial) (day 1), each mouse was subjected to the middle compartment with free access to all compartments for 10 min. The conditioning phase (days 2 to 4) consisted of six 10-min training sessions that were carried out during morning and afternoon training. Condensed milk was orally administered in the least preferred compartment at the start of the paradigm and at min 5, in the morning session; quinine (3mM) was always paired with the most preferred compartment at the same intervals, in the afternoon session. The morning and afternoon training sessions were repeated for 3 days in total from days 2 to 4. In the postconditioning phase (test trial, day 5), each mouse was subjected to the middle compartment with free access to the other compartments in the absence of treatments for 10 min. For the optogenetic experiment, during the conditioning phase the mice were sequestered in the least preferred chamber with constant 595 nm, 10 mW photoinhibition in the morning session . In the afternoon sessions, the mice were confined to the most preferred chamber and received photoinhibition and two oral administrations of quinine (3 mM). The test was conducted the following day after 3 days of conditioning for 10 min in absence of photoinhibition.

Social interaction (SOC): the test occurred in three sessions within a three-chambered arena (60 × 40 × 25(h) cm) with openings between the chambers (20 cm each). After 5 min habituation to the empty box, the mice were presented with 2 novel objects, one on each side of two chambers for 10 min. During the last 10 min one object in one chamber was replaced by an adult mouse inside a wire containment cup. For Optogenetic experiments, the mice are presented in the social apparatus with an adult mouse inside a wire containment cup for 10 min. The photoinhibition occurred during the last 5 minutes. The interactions with the novel mouse were manually scored with Ethovision.

**Analysis of calcium data**

Calcium imaging recordings are processed by Inscopix software into traces of calcium activity. We use z scores to represent the dynamic $Ca^{2+}$ activities in each neuron per trial, where trial means. To obtain the temporal z scores for a neuron, we first obtain the mean activity trace for the neuron by averaging the fluorescence signals (ΔF) at each time point across all trials and then computed the z scores as (F(t) – Fm)/s.d., where F(t) is the ΔF value at time t, and Fm and s.d. are the mean and standard deviation, respectively. For the trial-by-trial analyses, the z scores are computed for each trial using the same method (but without averaging across trials).

**CEBRA embeddings of calcium activity**

We apply the CEBRA pipeline (v0.2.0)(*17*) for generating calcium activity embeddings and feature selection, using the following parameters across Unsupervised (Time), Supervised (Behaviour) and Hybrid models: model_architecture = 'offset5-model'; distance = 'cosine'; batch_size = 600-1200; learning_rate = 1e-4; max_iterations = 8000. We use the additional parameters conditional = "time" for both the Time and Behavioural model, and conditional = "time-delta", hybrid = True for the Hybrid model.

We use the InfoNCE loss curve to inspect the model learning and compare actual data with shuffled data sets (1000 permutations along the time axis). To assess the embedding quality, we further apply the KNN decoder with cross animal validation (predicting every other animal besides itself). We validate our model by fitting a KNN decoder with randomly shuffled data sets. For the clustering analysis, we concatenate the feature-averaged responses (z scores) of individual neurons to each of the behavioural features. We perform principal-component analysis (PCA) on the z scores and use 3 PCs that capture more than 80% of the variance in the data for K-means clustering among behaviour features. We use the

Elbow method, which calculates the within-cluster sum of squares (WCSS), to determine the optimal number of clusters.

**Immunohistochemistry**

Mice are euthanized with an overdose of isoflurane (Floren, Abbott) and transcardially perfused with 20 ml phosphate-buffered saline (PBS), followed by 20 ml of 4% paraformaldehyde (PFA). Dissected brains are postfixed in 4% PFA overnight at 4°C, and 50-μm-thick sections are prepared using a vibratome (Microm HM 650 V, Thermo Fisher Scientific). Sections are rinsed in PBS and incubated overnight at 4°C with the primary antibody in PBS with 0.5% Triton X-100. On the next day, sections are washed, incubated with the secondary antibody coupled to suitable fluorescence (1:250 Alexa Fluor; Invitrogen, Thermo Fisher Scientific), and afterwards washed with PBS. Sections are mounted using Fluoromount-G mounting medium (SouthernBiotech) and either left to air-dry or stored at −20°C for image acquisition. The primary antibodies used were as follows: chicken anti-GFP antibody (1:500; Aves), rabbit anti-cFOS antibody (1:1000; Abcam). For quantification, all slides are randomised and coded before quantitative analysis. cFOS and GFP-labelled cells are counted on every sixth section through the entire rostrocaudal extent of regions of interest (ROIs; six sections per animal). Images are captured with either a Zeiss Axioplan2 fluorescence microscope and AxioVision 4.5 software or a Zeiss inverted laser scanning confocal microscope and Zen software. For confocal imaging, a z-stack of pictures of areas of interest is obtained with 0.4- to 1.2-μm step size and 2048-pixel by 2048-pixel picture size. Images are analysed with ImageJ (https://imagej.nih.gov/ij/).

**Activity-dependent labelling in FOSTRAP2 mice**

To label cells that are activated by specific contexts, we utilise FOSTRAP2 mice. For *in vivo* $Ca^{2+}$ imaging, we inject $AAV_5$-hSyn-DIO-GCaMP6s unilaterally in the IPACL. Two weeks after viral injection, we follow a similar procedure for GRIN lens implantation and baseplating. After recovery, mice are subjected to activity-dependent cell labelling. In brief, we injected 100 mg/kg tamoxifen (TAM) 3.5 hr prior to context exposures. Mice are exposed to either elevated platform, quinine or novel social conspecifics. 25 min after the exposure, mice are transferred back to their home cage. Two weeks after cell labelling, mice are subjected to *in vivo* $Ca^{2+}$ recordings. For retrograde tracing, mice are sacrificed two weeks after context exposures.

**FACs-sorting of TRAPed neurons**

Mice subjected to EP, SOC, Q and CON contexts (n = 3-4 per group) were euthanized at 2 months of age after context exposure by cervical dislocation. Brains were collected in ice-cold PBS and processed for tissue dissection. The target region (extended amygdala) was isolated by subjecting brains to a brain matrix (Fine Science Tools). Brains were cut into 1-mm thick slices covering the entire ROI. The dorsal part including the cortex was removed, and the remaining tissue was dissociated into single cells using the Papain Dissociation System (#LK0031500, Worthington Biochemical). Then, samples were processed to fluorescence-activated cell sorting (FACS) using a FACSMelody (Becton Dickinson) with a nozzle diameter of 100 µm and FACSFlow medium. Debris and aggregated nuclei were gated out by forward and sideward scatter. Single nuclei were gated out by FSC-width/FSC-area. Gating for fluorophores was done using isolated cells possessing GFP in the nuclear membrane.

FACs-sorted TRAPed neurons among mice subjected to the same context were pooled and redistributed into three pooled samples for scRNA-seq; non-FACs-sorted neurons (not activity-TRAPed) among all mice, regardless of the contexts they were subjected to, were pooled and redistributed into five pooled samples for scRNA-seq. The latter were used as control (Ctrl) samples containing non-activity labelled neurons. Of note, as these neurons were pooled across contexts, some non-TRAPed neurons in one context may in fact be TRAPed in another. As such, Ctrl samples are enriched for non-TRAPed neurons across all contexts, and increasingly depleted for neurons with greater overlapping activity across contexts (and therefore TRAPed in an increasing number of contexts).

**Single cell RNA sequencing (scRNA-seq)**

scRNA-seq was performed using a 10× genomics kit (10x Chromium 3.0) following the instructions of the manufacturer. Briefly, cells were mixed with solutions containing barcoded beads and partitioning oil. To achieve single-nucleus resolution, nuclei were delivered at a limiting dilution. Next, gel beads were dissolved, primers were released, and any co-partitioned cells were lysed. Primers were mixed with the cell lysate and with a master mix containing reverse transcription reagents. This produced barcoded, full-length cDNA from polyadenylated mRNA. After incubation, products were broken up and pooled fractions recovered. Silane magnetic beads were used to purify the first-strand cDNA from the post–reverse transcription reaction mixture. Barcoded, full-length cDNA was amplified via polymerase chain reaction to generate sufficient mass for library construction. Enzymatic fragmentation and size selection were used to optimise the cDNA amplicon size. A Chromium Single-Cell 3′ Gene Expression library comprises standard Illumina paired-end constructs beginning and ending with P5 and P7. The 16-bp 10× barcode and 12-bp unique molecular identifier (UMI) were encoded in Read 1, while Read 2 was used to sequence the cDNA fragments. Sample index sequences were incorporated as the i7 index read. TruSeq Read 1 and TruSeq Read 2 are standard Illumina sequencing primer sites used in paired-end sequencing.

**Preprocessing and quality control of snRNA-seq data**

Single-cell RNAseq data were analysed following the recommended pipeline suggested in the best practice for single-cell data analysis (*43*, *44*). We first used the cell calling algorithm in CellRanger (10x Genomics Cell Ranger 7.1.0) to call the raw gene expression matrix, taking into account the number of UMIs and the number of barcodes jointly. We used the "-- force" flag to assign 4500 cells per sample to meet the biological expectations. We then used the python package Scanpy (*45*) to analyse the raw gene expression matrices.

We carried out quality control (QC) steps on each sample individually: cells with >5 median absolute deviations (MADs) in log10 total counts, log10 total features, or cumulative percentage counts for 20 most expressed genes in a cell were considered outliers and removed; cells with > 3 MADs in mitochondrial gene fraction or >20% mitochondrial genes were further removed as outliers (*43*, *46*). We then used the R package SoupX to identify ambient RNA and corrected the count matrix for each sample (*47*) (**fig. S18 A to E**). Genes that were detected in less than 5 cells in the corrected matrix were then filtered out. Finally, we used the python package Scrublet to filter out doublets (*48*). Samples were then grouped by treatment for batch effects detection and correction using the python package Scanorama (*49*) , normalisation, log-transformation and dimensionality reduction using principal components analysis (PCA) and leiden algorithm clustering in Scanpy (*45*).

To identify the cell types present in each sample, we inputted our post-Scanorama matrix in the online tool MapMyCells (RRID:SCR_024672) provided by Allen institute, which maps our data onto high-quality, high-resolution cell type taxonomies available in mouse brain reference atlases. The identified cell types were visualised using the uniform manifold approximation and Projection algorithm (UMAP) implemented in Scanpy (*45*).

**Differential gene expression analysis (DGE)**

Pseudo-bulk gene expression levels are calculated for genes in each sample, in glutamatergic and GABAergic cells present in TRAPed cells from each context using the python package decoupleR(*50*). DEG analyses are performed between TRAPed glutamatergic and GABAergic neurons from pairs of contexts, as well as between the two cell types in each context against all other contexts. We perform the DEG analysis using the python package DEseq2 (*51*), and perform multiple-testing correction on our results accounting for all tests per cell type. Genetic associations of significant DEG genes with human diseases and complex traits are performed using the Harmonizome(*19*) database.

### Supplementary Figures

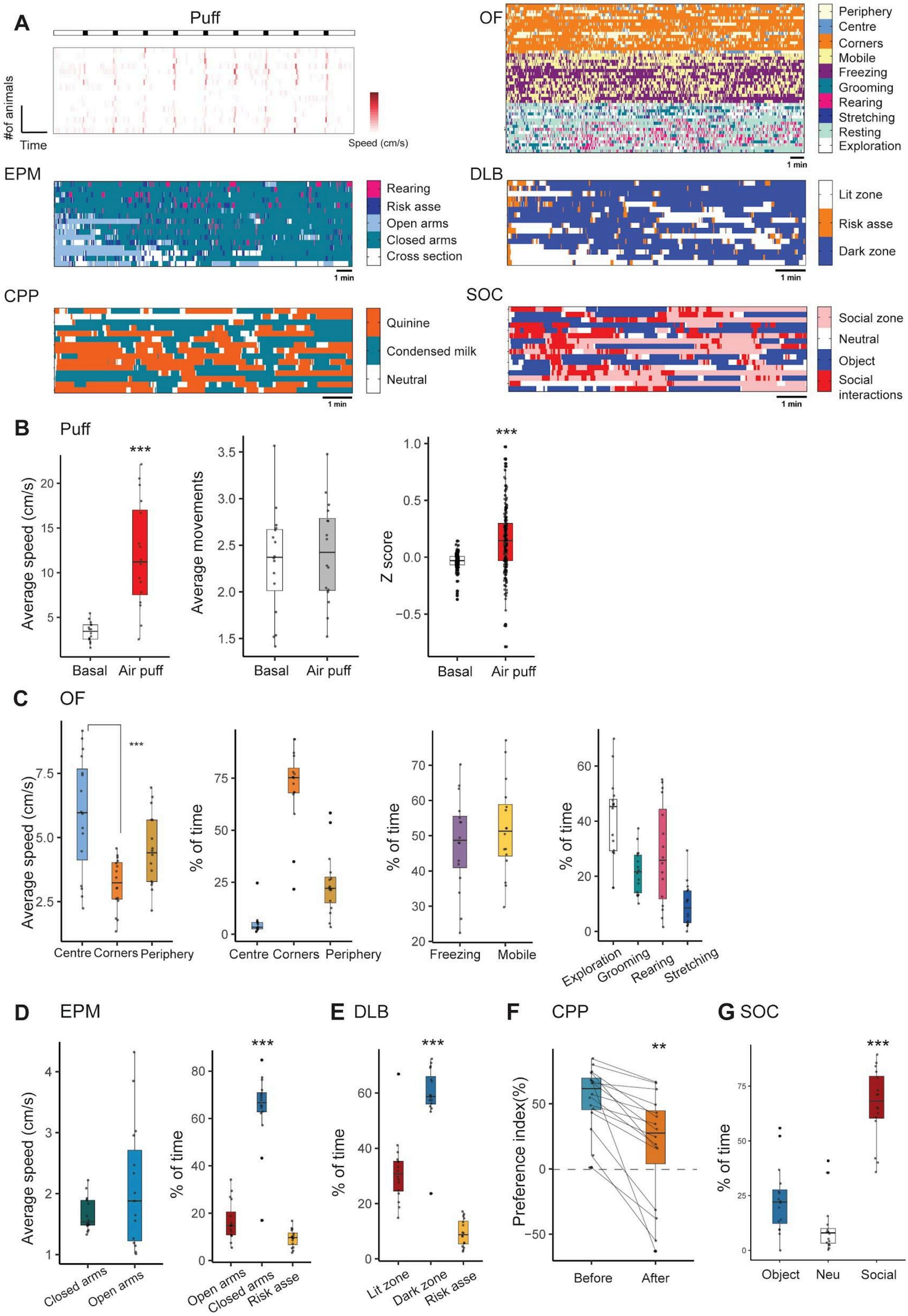

**Fig S1. Mouse behavioural responses to behavioural contexts**

A. Heatmaps demonstrating the spontaneous behaviours of freely moving mice recorded in each behavioural context. B. Air puff (Puff): The presentation of air puff induces flight behaviours in mice, as evidenced by a noticeable increase in their movement speed compared to baseline ($p$ < 0.0001, t = 8.21); Boxplot showing that air puff does not change average movements of mice ($p$ = 0.64, t = 0.47). IPACL neurons have higher average activity when receiving air puff ($p$ < 0.0001, t = 8.21). C. Open field (OF): Mice exhibit greater speed in the centre zone of the OF compared to the corners (One-way ANOVA, $F_{2,45}$ = 10.61, $p$ = 0.0002. Bonferroni's post-hoc test: centre/corners, $p$ < 0.0001, t = 4.6); percentages of time mice spent in corner (70.7% $\pm$ 4.8), centre zone (5.1% $\pm$ 1.4) or periphery (24.2% $\pm$ 3.8) during the OF test. Percentages of time dedicated to different behaviours in the OF (exploring, 40.71% $\pm$ 3.81; grooming, 21.69% $\pm$ 2 ; stretching, 9.58% $\pm$ 1.97 ; and rearing, 28.02% $\pm$ 4.63). D. Elevated plus maze (EPM): during EPM mice stay for the longest time majorly in the closed arms (64.1% $\pm$ 3.9) and occasionally come out to the open arms (16.6% $\pm$ 2.1), concurrently performing risk assessment (9.4% $\pm$ 0.9) (One-way ANOVA, $F_{2,45}$ = 130.4, p < 0.0001); boxplot demonstrating no difference in average speed during EPM when mice are in either open or closed arms (p = 0.1, t = 1.68). E. Dark-light box (DLB): Mice spend more time in the dark zone (59.1% $\pm$ 2.8) of the DLB compared with the lit zone (31.6% $\pm$ 3) where they show risk assessment (9.4% $\pm$ 1.22) behaviour (One-way ANOVA, $F_{2,45}$ = 100.8, p < 0.0001). F. Conditioned place preference (CPP): After CPP conditioning, mice show higher preference toward the chamber paired with condensed milk (p = 0.0002, t = 4.792). G. Social interaction test (SOC): Mice spend more time in social interactions as well as in the chamber where the social counterpart is present compared to the chamber containing an object (One-way ANOVA, $F_{2,45}$ = 65.42, p < 0.0001). Value = median $\pm$ $min/max$, **$p$ < 0.01, ***$p$ < 0.0001.

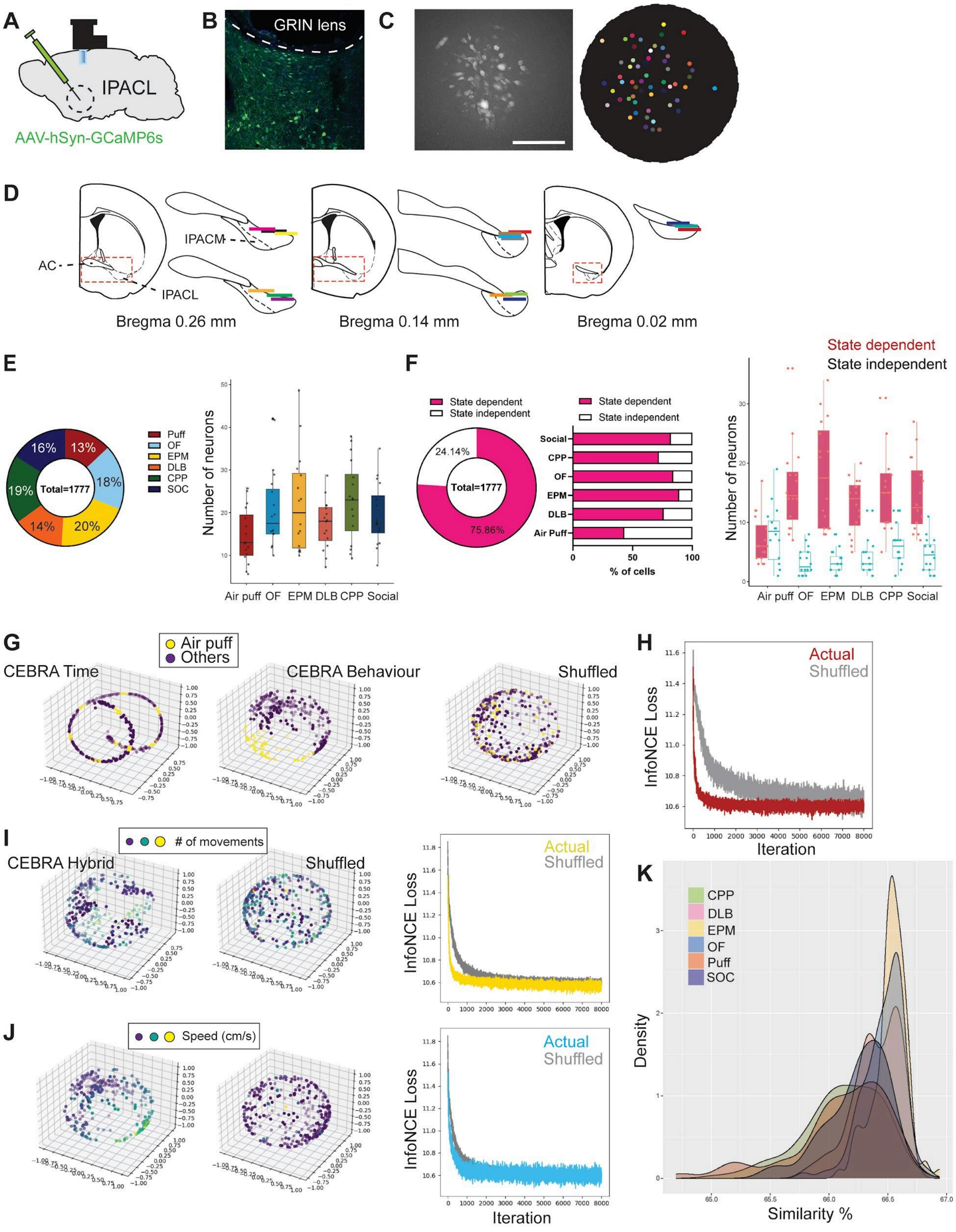

### Fig S2. Deep brain $Ca^{2+}$ imaging in the IPACL

A. Scheme of GRIN lens implantation and viral injection. B. Representative image of viral expression in IPACL and field of view during recording (C). D. Cartoon depicting the rostro-caudal extension. E. Pie chart and box plot showing respectively the percentage and number of neurons recorded across 6 different behavioural contexts.

F. Pie chart of percentages of state dependent (76%, magenta) and state independent (24%, blue) neurons across all behavioural contexts; bar plot of percentages per behavioural context; boxplots of the corresponding numbers of neurons per behavioural context. G. CEBRA models (trained with the Time or Behaviour models) show how the air puff contributes to changes in neural activity in the IPACL, resulting in separated embeddings, compared to the embedding generated from shuffled data. H. The InfoNCE loss curve shows the model learning through 8000 iterations in the CEBRA Hybrid model. The red curve indicates the actual data used in the study and the grey curve shows the result from shuffled data. I,J. CEBRA embeddings and the InfoNCE loss curve demonstrate no contribution of either movement or speed to changes of neural activity during air puff. K. Density plot showing distribution of similarity of CEBRA ensembles across six behavioural tasks. Value = median $\pm min/max$. Scale bar = 200 µm.

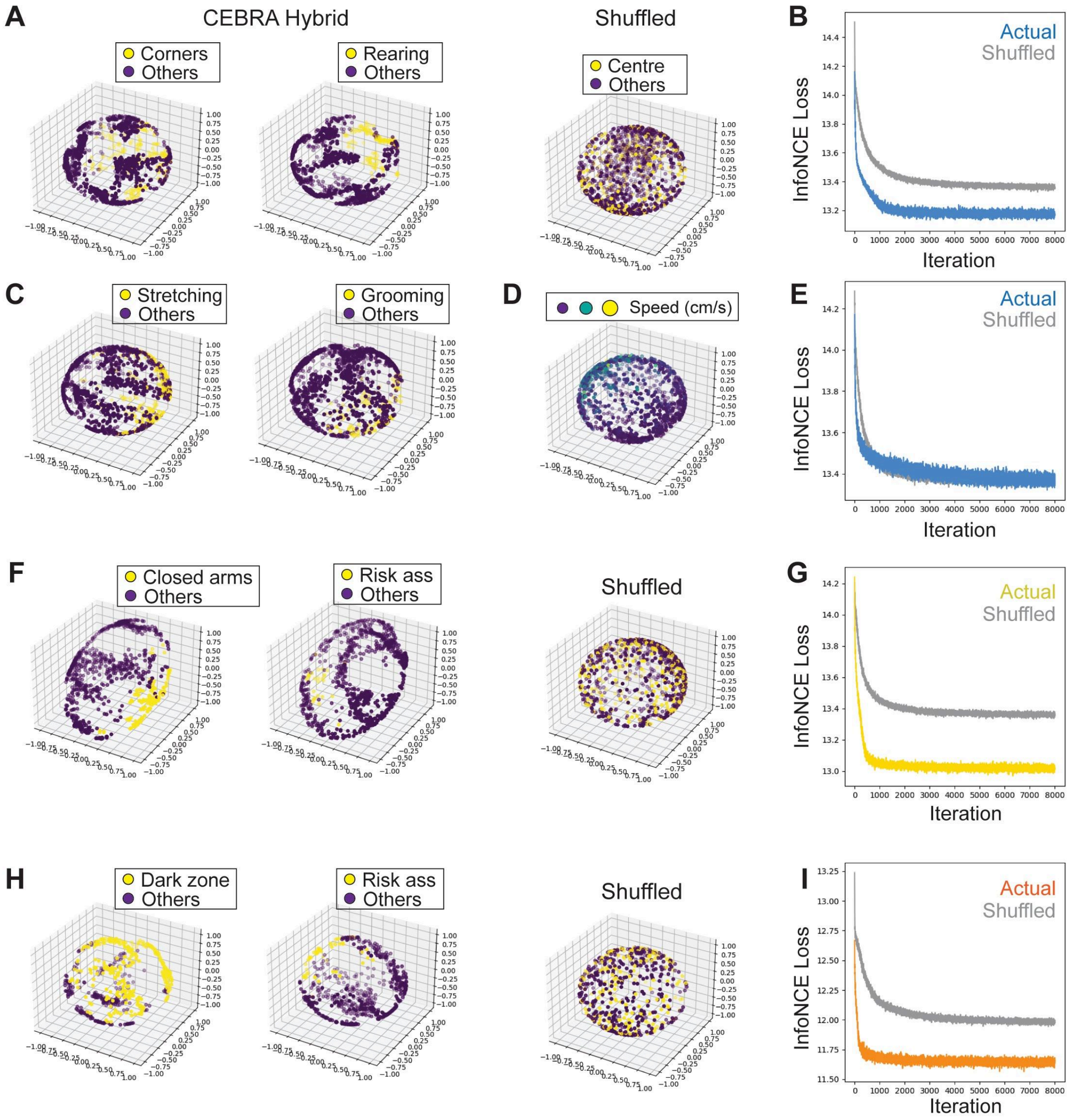

**Fig S3. IPACL neurons encode different behavioural contexts**

A-E. Examples of CEBRA embeddings and the InfoNCE loss curve. Behaviours such as corner entry and rearing (j) as well as stretching and grooming (l) contribute to changes of neural activity in OF. m,n. Speed of mice in the OF does not contribute to changes of neural activity. F. CEBRA embeddings reveal that both closed arms entry and risk assessment contribute to changes in neural activity in the IPACL during EPM. G. The InfoNCE loss curve of the CEBRA Hybrid model through 8000 iterations in EPM shows better learning in actual (yellow) data compared to shuffled (grey) data. H. CEBRA embeddings illustrate that changes in neuronal activity within the IPACL during DLB are influenced by the entry into the dark zone and risk assessment. I. The InfoNCE loss curve of the CEBRA-Hybrid model through 8000 iterations in DLB illustrates the difference between actual (orange) and shuffled (grey) data. Value = median $\pm min/max$.

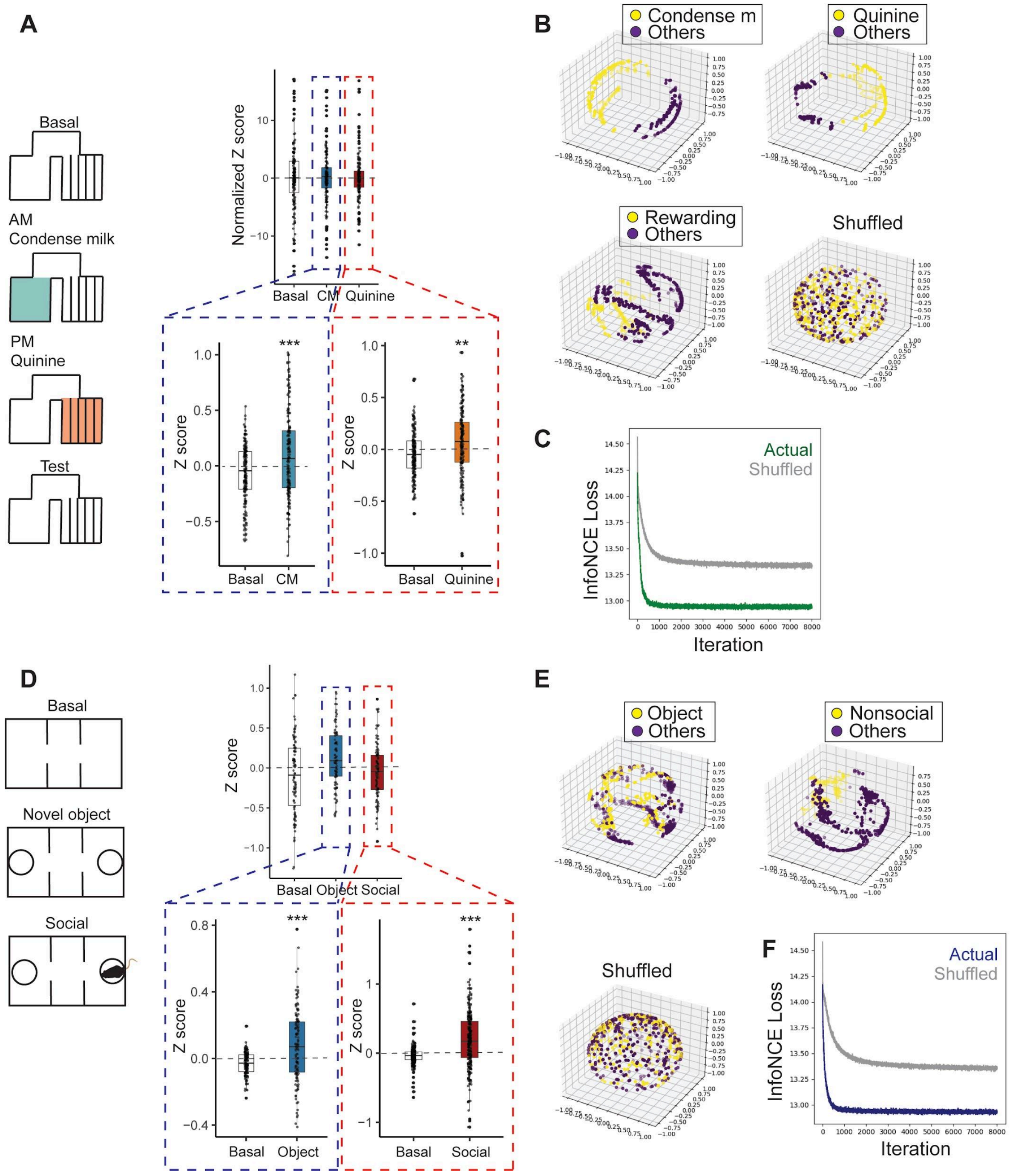

**Fig S4. Neural activity dynamics in CPP and three-chamber social interaction test**

A. Scheme showing the sequence of conditioning in the CPP. No differences in normalised activity under basal conditions, for both CM or Q encoded neurons. However, during conditioning, the administration of both condensed milk and quinine leads to significant increase of activity of IPACL neurons (CM, $p < 0.0001$, t = 3.41; Q, $p = 0.0041$, t = 2.91). B. CEBRA embeddings show that administration of condensed milk and quinine and entry into the reward conditioned chamber contribute to changes in neural activity in the IPACL. C. The InfoNCE loss curve of the CEBRA Hybrid model through 8000 iterations in the CPP illustrates successful learning of actual (green) data compared to shuffled (grey) data. D. Scheme of sequence in three-chamber social interaction test. No difference in activity between basal, nonsocial or social neurons. Activity of neurons increases when in contact with novel objects or social partners (object, $p < 0.0001$, t = 4.31; social, $p < 0.0001$, t = 8.34). E. CEBRA embeddings highlight that novel objects as well as nonsocial zones affect neural activity in the IPACL. F. The InfoNCE loss curve of the CEBRA Hybrid model through 8000 iterations in social interaction shows the difference between actual (dark blue) and shuffled (grey) data. Value = median $\pm$ $min/max$, $^{**}p < 0.01$, $^{***}p < 0.0001$.

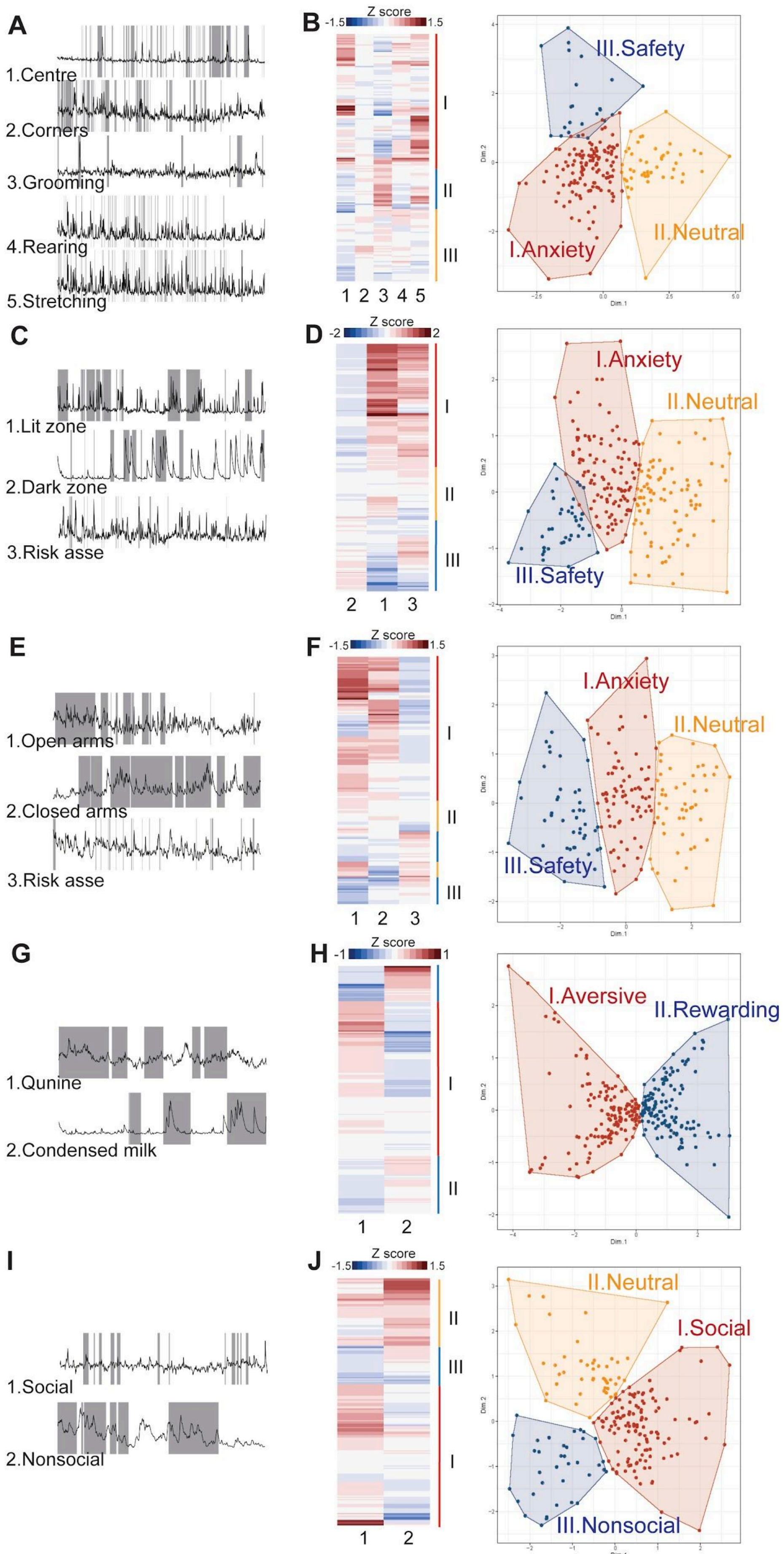

A
1.Centre
2.Corners
3.Grooming
4.Rearing
5.Stretching
B
Z score
-1.5
1.5
I
II
III
1 2 3 4 5
III.Safety
I.Anxiety
II.Neutral
Dim.1
Dim.2
C
1.Lit zone
2.Dark zone
3.Risk asse
D
Z score
-2
2
I
II
III
2 1 3
I.Anxiety
II.Neutral
III.Safety
E
1.Open arms
2.Closed arms
3.Risk asse
F
Z score
-1.5
1.5
I
II
III
1 2 3
I.Anxiety
II.Neutral
III.Safety
G
1.Qunine
2.Condensed milk
H
Z score
-1
1
I
II
1 2
I.Aversive
II.Rewarding
I
1.Social
2.Nonsocial
J
Z score
-1.5
1.5
II
III
I
1 2
II.Neutral
I.Social
III.Nonsocial

**Fig S5. Neuronal activity clustering during diverse behavioural contexts**

A. Example traces of neurons in the IPACL that account for changes in neural activity during different behaviours in the OF (in grey). B. Heatmap shows the average activities (Z-score) of behavioural features (time locked) used for clustering neurons into different ensembles during OF. Scatterplot shows that neurons can be clustered into 3 different clusters encoding I. anxiety, II. neutral or III. safety conditions. C,D. Example traces of neurons in the DLB. We used these behavioural features to cluster neurons into anxiety, neutral or safety clusters. E,F. Example traces of neurons in the EPM. Neurons are clustered into anxiety, neutral or safety. G,H. Example traces of neurons in the CPP while mice are either in the condensed milk or quinine chamber. Neurons are clustered into aversive or rewarding. I,J. Example traces of neurons during the social interaction behaviour. Neurons are clustered into social, nonsocial or neutral.

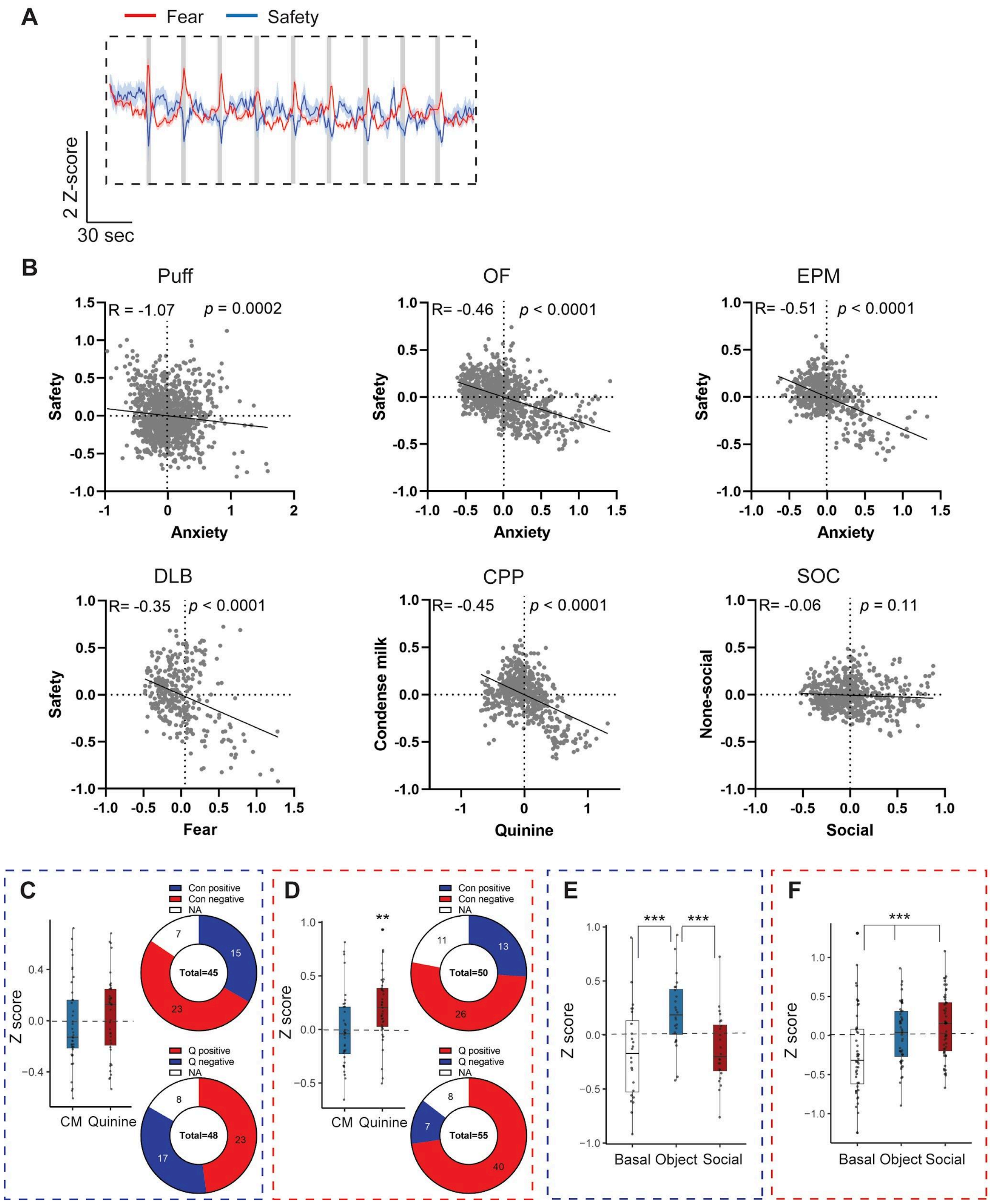

**Fig S6. Correlation between different IPACL ensembles**

A. Neuronal traces show the average activities of fear (red) and safety (blue) ensembles during Puff. B. Activities of fear and safety ensembles are negatively correlated with each other in all behavioural contexts except for SOC. C. Neurons encoding reward during the CPP test exhibit comparable activity levels during both condensed milk and quinine administration ($p = 0.34$, $t = 0.96$). Pie charts illustrating the number of neurons involved in encoding reward memory that respond to condensed milk or quinine during the conditioning phase, with a greater number of neurons displaying responsiveness to quinine. D. Neurons that encode aversion during CPP test show higher activity when animals receive quinine compared to condensed milk ($p = 0.0045$, t = 2.92). As expected, more aversive neurons are responsive to quinine during the conditioning phase. E. Neurons encoding nonsocial states show highest activity when mice interact with novel objects (Mixed effect model (REML), $p < 0.0001$. Bonferroni's post-hoc test: object/basal, $p < 0.0001$, t = 28.9; object/social, $p < 0.0001$, t = 28.9). F. Neurons encoding social state show highest activity when mice interact with social counterparts (Mixed effect model (REML), Bonferroni's post-hoc test: social/basal, $p < 0.0001$, t = 21.4; social/object, $p < 0.0001$, t = 10.7). Value = mean $\pm min/max$, $^{**}p < 0.001$, $^{***}p < 0.0001$.

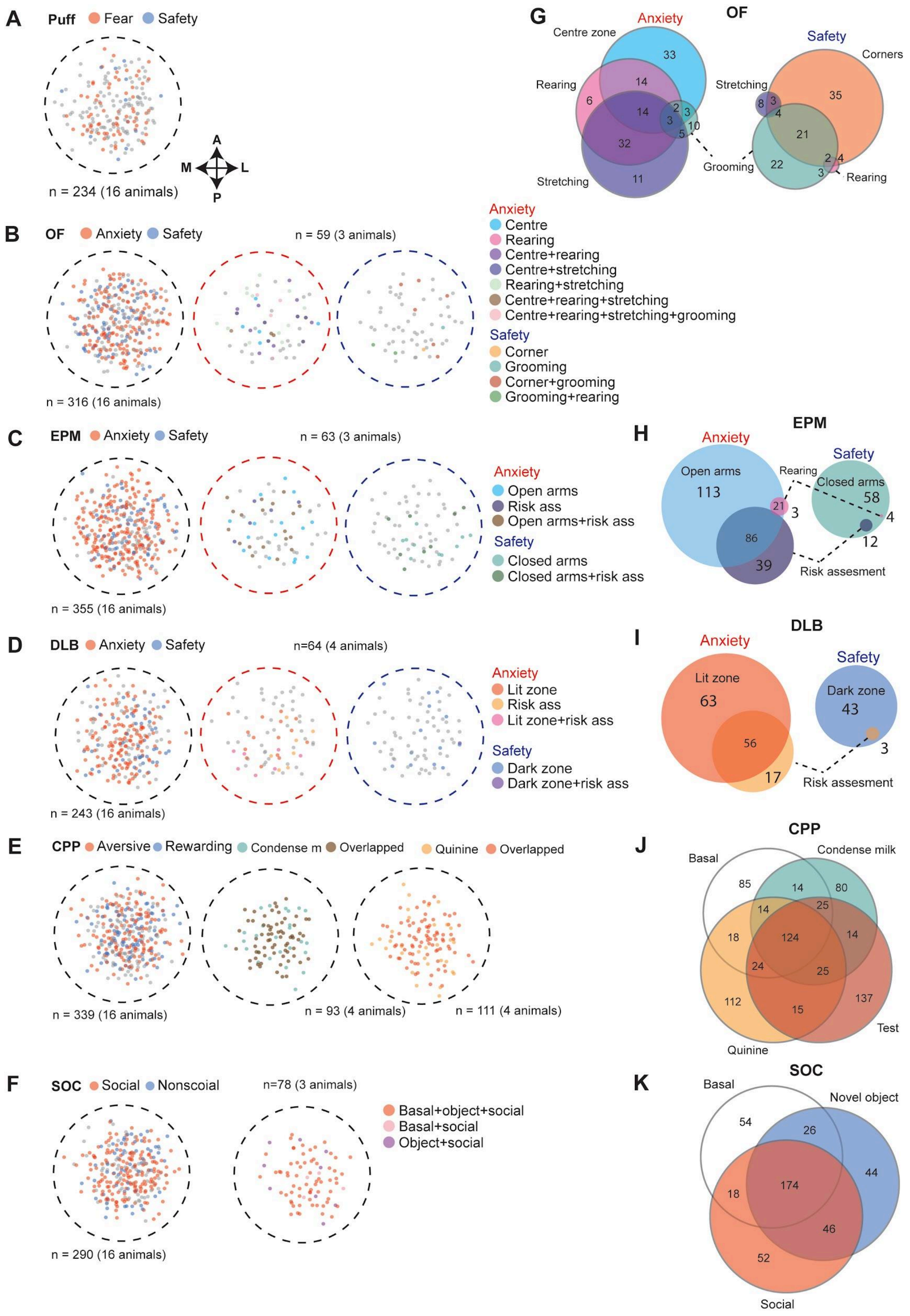

A
Puff Fear Safety
A
M
L
P
n = 234 (16 animals)
B
OF Anxiety Safety
n = 59 (3 animals)
n = 316 (16 animals)
Anxiety
Centre
Rearing
Centre+rearing
Centre+stretching
Rearing+stretching
Centre+rearing+stretching
Centre+rearing+stretching+grooming
Safety
Corner
Grooming
Corner+grooming
Grooming+rearing
C
EPM Anxiety Safety
n = 63 (3 animals)
n = 355 (16 animals)
Anxiety
Open arms
Risk ass
Open arms+risk ass
Safety
Closed arms
Closed arms+risk ass
D
DLB Anxiety Safety
n=64 (4 animals)
n = 243 (16 animals)
Anxiety
Lit zone
Risk ass
Lit zone+risk ass
Safety
Dark zone
Dark zone+risk ass
E
CPP Aversive Rewarding Condense m Overlapped Quinine Overlapped
n = 339 (16 animals)
n = 93 (4 animals)
n = 111 (4 animals)
F
SOC Social Nonscoial
n=78 (3 animals)
n = 290 (16 animals)
Basal+object+social
Basal+social
Object+social
G
OF
Anxiety
Centre zone
33
Rearing
14
6
14
2
3
3
3
10
5
32
11
Stretching
Grooming
Safety
Corners
Stretching
35
8
3
4
21
22
2
4
3
Rearing
H
EPM
Anxiety
Open arms
113
Rearing
Safety
Closed arms
58
21
3
4
86
12
39
Risk assesment
I
DLB
Anxiety
Lit zone
63
Safety
Dark zone
43
56
3
17
Risk assesment
J
CPP
Basal
Condense milk
85
14
80
14
25
18
124
14
24
25
112
15
137
Test
Quinine
K
SOC
Basal
Novel object
54
26
44
174
18
46
52
Social

**Fig S7. Distribution of neural ensembles in the IPACL**

A. Image of ensembles in the region of interest (ROI) encoding fear (red), safety (blue) or neutral (grey) states during Puff. B. Images of neurons distributed in the ROI with unique or concurrent activity in OF. C. Images of neurons recorded under ROI exhibiting unique or concurrent activity in the EPM. D. Images of neurons recorded in the DLB having unique or concurrent activity between states. E. Images of cells under ROI with concurrent activity across the CPP test. F. Neurons recorded in the social interaction test having unique or concurrent activity across the testing phase. G-K. Graphical representation of the number of neurons responsive to different contexts with unique or concurrent activities across contexts.

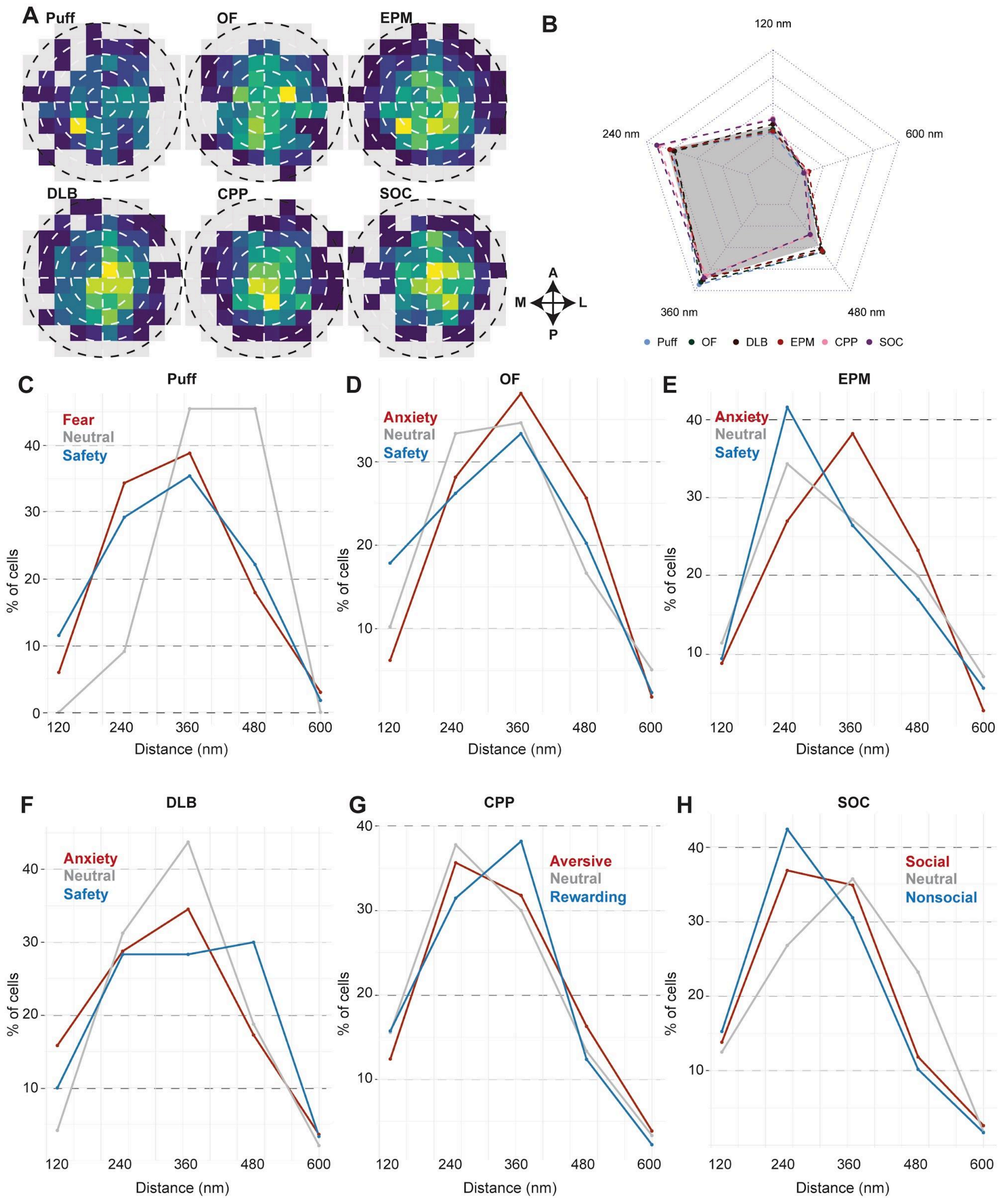

**Fig S8. Distribution analysis of cells and neural ensembles across behavioural contexts**

A. Heatmaps demonstrate the distribution of cells (percentage) recorded within ROI. Each white ring represents 20% of the surface under the ROI, distance covered by the ring is calculated from the centre of the ROI. B. Radar plot summarised the percentage of cells distributed within the ROI, each colour dot line represents one context and the grey polygon shows the average distribution of all conditions. There is no difference in distribution of cells between different behaviour exposures. C-H. Distribution of neural ensembles under ROIs in different behavioural contexts.

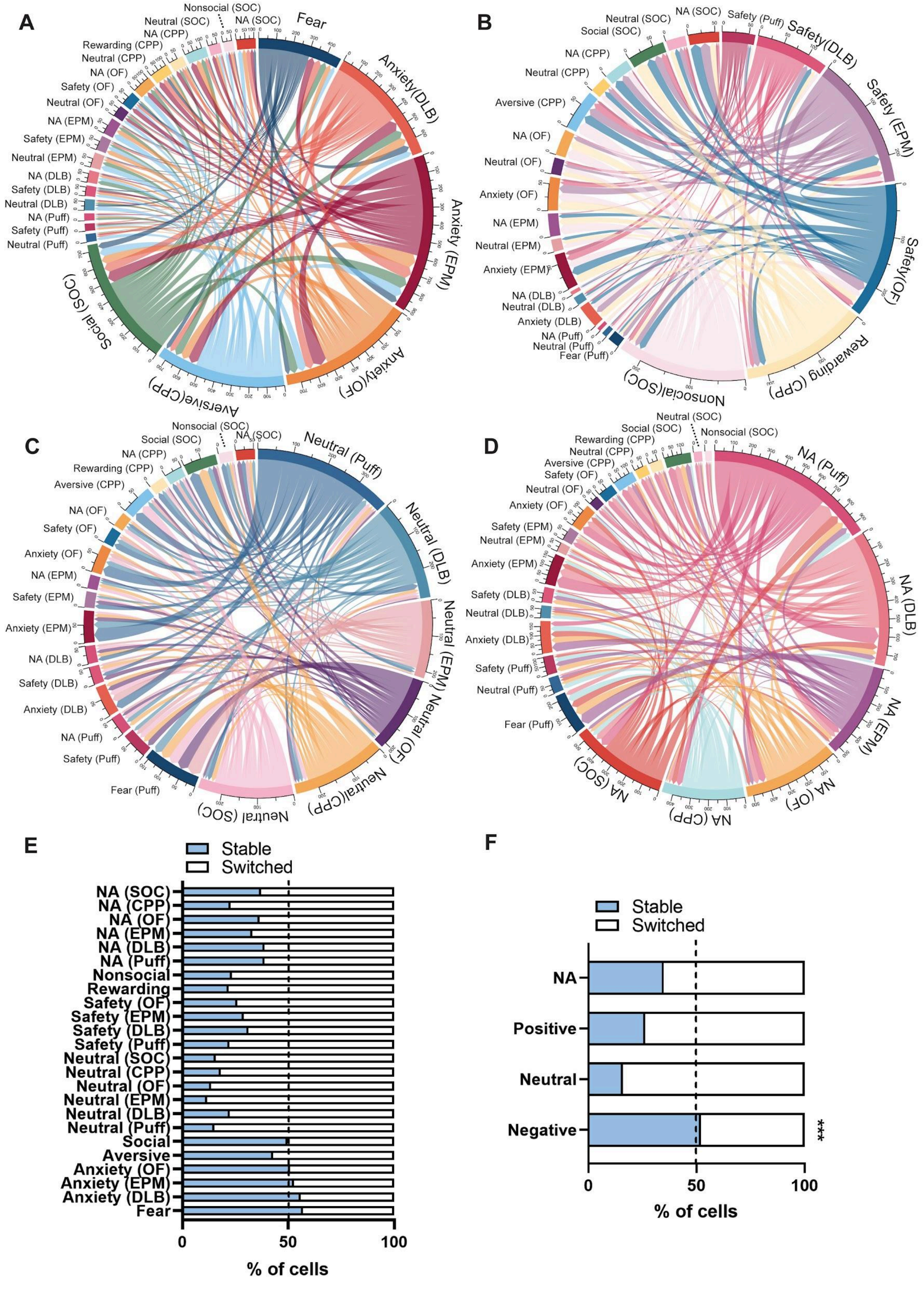

**Fig S9. Flexibility between neural ensembles in the IPACL**

A-D. Chord diagrams showing the relationship of neural ensembles and their encoding states across behavioural tasks. E. Detailed and summarised percentage of cells remaining stable in encoding state or switching between encoding across behavioural tasks. F. Ensembles encoding negative experiences have higher stability compared to the others (Two-way ANOVA, $F_{1,40} = 461$, $p < 0.0001$). Value = mean $\pm min/max$, ***$p < 0.0001$.

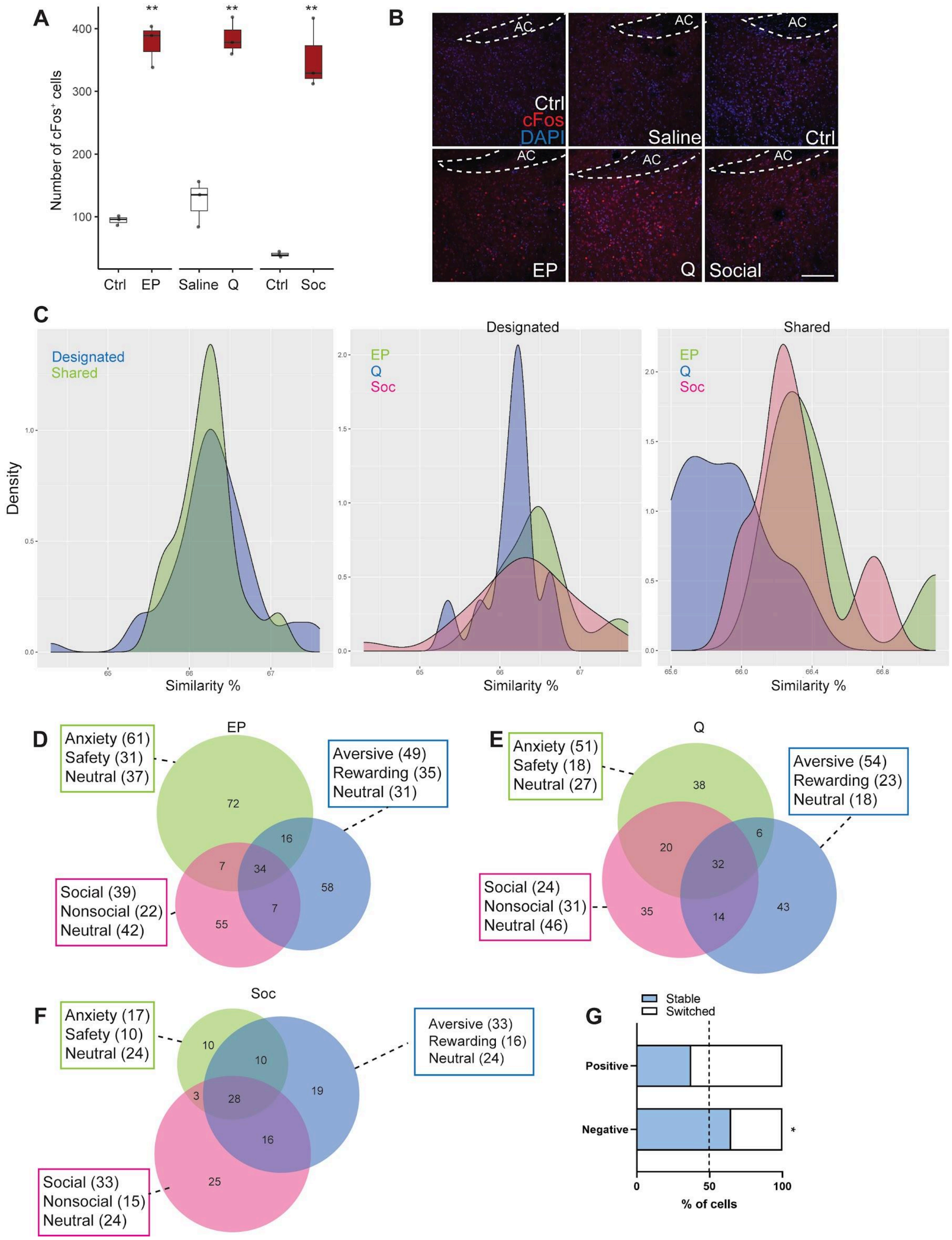
A
Number of cFos+ cells
Ctrl
EP
Saline
Q
Ctrl
Soc
B
AC
Ctrl
cFos
DAPI
Saline
Ctrl
EP
Q
Social
C
Designated
Shared
Density
Similarity %
Designated
EP
Q
Soc
Shared
D
EP
Anxiety (61)
Safety (31)
Neutral (37)
Aversive (49)
Rewarding (35)
Neutral (31)
Social (39)
Nonsocial (22)
Neutral (42)
E
Q
Anxiety (51)
Safety (18)
Neutral (27)
Aversive (54)
Rewarding (23)
Neutral (18)
Social (24)
Nonsocial (31)
Neutral (46)
F
Soc
Anxiety (17)
Safety (10)
Neutral (24)
Aversive (33)
Rewarding (16)
Neutral (24)
Social (33)
Nonsocial (15)
Neutral (24)
G
Stable
Switched
Positive
Negative
% of cells

**Fig S10. cFOS expression patterns and ensemble stability in mice exposed to diverse behavioural contexts**

A. cFOS expression in mice exposed to various contexts is higher compared to the Ctrl group (EP, $p$ = 0.0002, t = 13.1; Q, $p$ = 0.0007, t = 9.5; SOC, $p$ = 0.0007, t = 9.5). B. Representative images of IPACL cFOS expression in different behaviours. C. Density plots of the similarity between ensembles. Designated and shared ensembles show high similarity. D-F. Numbers of neurons demonstrating unique or conserved activity across three behavioural paradigms. g. Ensembles encoding negative experiences show higher stability (Two-way ANOVA, Bonferroni's post-hoc test, $p$ = 0.023, t = 3.99). Scale bar = 200 μm. Value = median $\pm\, min/max$, *$p < 0.05$, **$p < 0.001$.

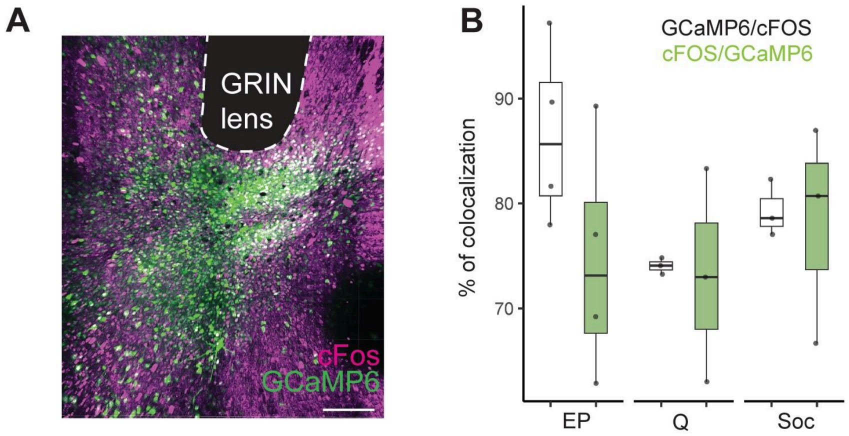

**Fig S11. Quantification of colocalization of $cFOS^+$ and $GCaMP6^+$ neurons in the IPACL**

A. Representative image showing colocalization (white) of $cFOS^+$ neurons (magenta) and $GCaMP6^+$ neurons (green) in relation to the position of the GRIN lens in the IPACL. B. Percentage of cells with colocalization of $GCaMP6^+$ and $cFOS^+$ neurons. More than 70% colocalization of $GCaMP^+$ and $cFOS^+$ neurons , the results are ruled out as being due to non-specific labelling of cells. Value = median $\pm\, min/max$. Scale bar = 200μm.

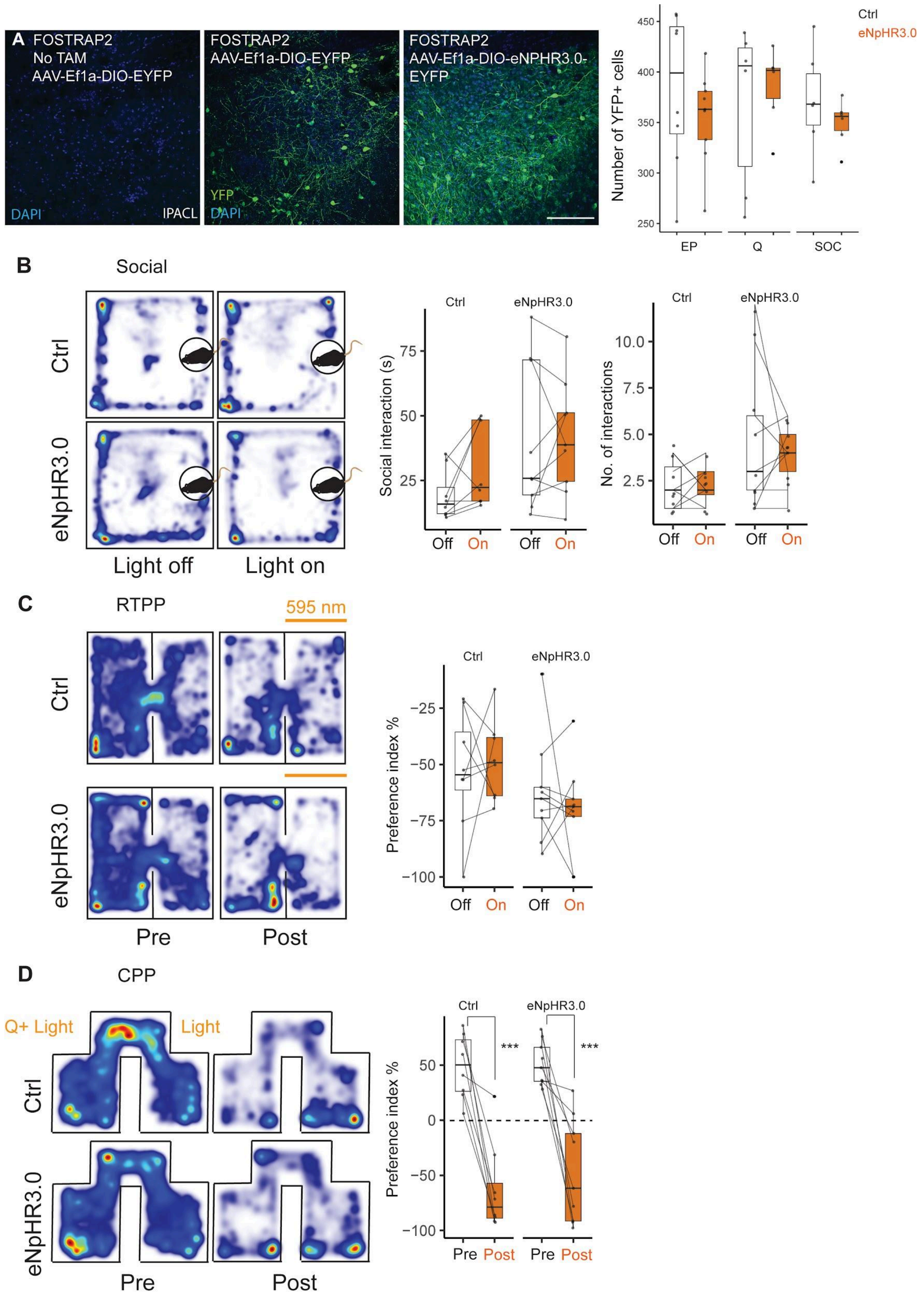
A
FOSTRAP2
No TAM
AAV-Ef1a-DIO-EYFP
DAPI
IPACL
FOSTRAP2
AAV-Ef1a-DIO-EYFP
YFP
DAPI
FOSTRAP2
AAV-Ef1a-DIO-eNPHR3.0-EYFP
Number of YFP+ cells
Ctrl
eNpHR3.0
EP
Q
SOC
B
Social
Ctrl
eNpHR3.0
Light off
Light on
Social interaction (s)
No. of interactions
Off
On
C
RTPP
595 nm
Pre
Post
Preference index %
D
CPP
Q+ Light
Light
Pre
Post
***

**Fig S12. Anxiety ensemble has no effect on sociability, place preference and aversive avoidance**

A. Images and bar plots show similar expression of GFP (Ctrl) and eNphR3.0 in the IPACL. There is no expression of virus without Tamoxifen (TAM, 100 mg/kg) injection. No difference between Ctrl or eNpHR3.0 animals in the number of $YFP^+$ cells in the IPACL after TAM administration (EP, $p = 0.37$, t = 0.92; Q, $p = 0.41$, t = 0.86; SOC, $p = 0.64$. t = 0.48). Scale bar = 200 µm. B. Heatmap illustrating the spatial positioning of the animal in relation to the social stimulus. Inhibition of anxiety responding ensembles does not affect sociability of mice (interaction time, Two-way ANOVA, effect of light, $F_{1,30} = 0.72$, $p = 0.4$; No. of interactions, Two-way ANOVA, effect of light, $F_{1,30} = 0.23$, $p = 0.63$). C. Heatmap showing the position of the animal during RTPP (pre: no light; post: with light). Inhibition of anxiety responding ensembles does not generate real time place preference. (Two-way ANOVA, effect of light, $F_{1,30} = 0.05$, $p = 0.82$). D. Heatmap of the mouse location during CPP (pre: before conditioning; post: after conditioning). Inhibition of the anxiety ensemble does not affect quinine induced place aversion (Two-way ANOVA, effect of opsin, $F_{1,30} = 0.57$, $p = 0.45$; effect of quinine, $F_{1,30} = 73.5$, $p < 0.0001$. Bonferroni's post-hoc test, Ctrl pre/post, $p < 0.0001$, t = 6.3; eNpHR3.0 pre/post, $p < 0.0001$, t = 5.8). Value = median $\pm min/max$, ***$p < 0.0001$. Scale bar = 200µm.

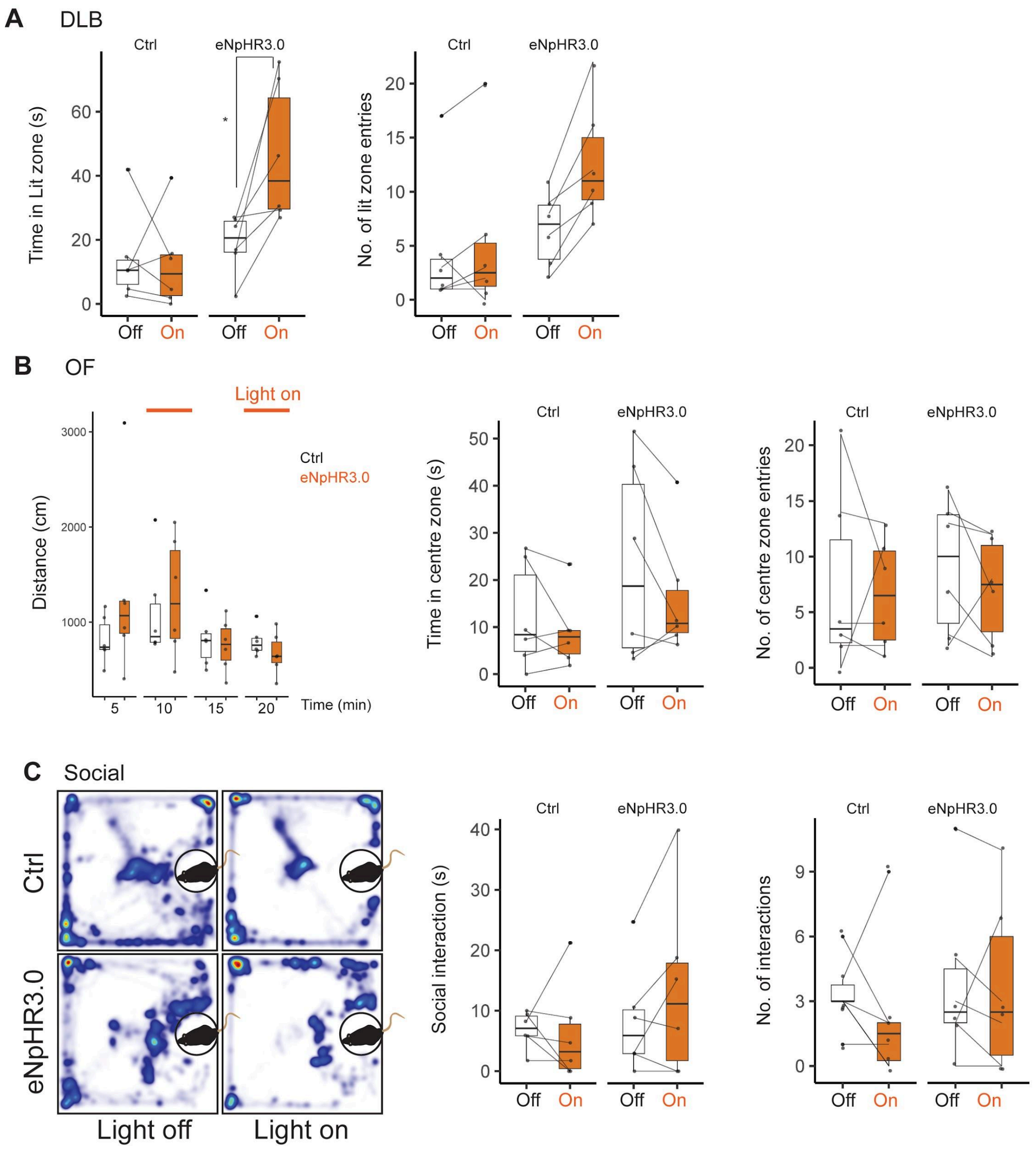

**Fig S13. Inhibition of Q-TRAPed neurons has no effect on anxiety and sociability**

A. Inhibition of Q-TRAPed ensembles in the IPACL during DLB results in longer time spent in the lit zone of eNpHR3.0 compared to Ctrl animals (Two-way ANOVA, effect of light, $F_{1,20} = 9.16$, $p = 0.006$; Bonferroni's post-hoc test, $p = 0.036$, $t = 3.07$). There is no difference between the two groups in the number of entries into the lit zone (Two-way ANOVA, effect of light, $F_{1,20} = 2.14$, $p = 0.16$). B. In OF, there is no difference between eNpHR3.0 and Ctrl animals with respect to distance travelled (Two-way RM ANOVA, effect of light, $F_{1,10} = 0.38$, $p = 0.54$), time in centre zone (Two-way ANOVA, effect of light, $F_{1,20} = 0.83$, $p = 0.37$) or number of centre zone entries (Two-way ANOVA, effect of light, $F_{1,20} = 0.32$, $p = 0.57$) when Q-TRAPed ensembles are inhibited. C. Heatmap and boxplots show no difference between eNpHR3.0 and Ctrl mice in sociability when inhibiting Q-TRAPed ensembles (interaction time, Two-way ANOVA, effect of light, $F_{1,20} = 0.3$, $p = 0.59$; No. of interaction, Two-way ANOVA, effect of light, $F_{1,20} = 0.18$, $p = 0.68$). Value = median $\pm min/max$, $*p < 0.05$.

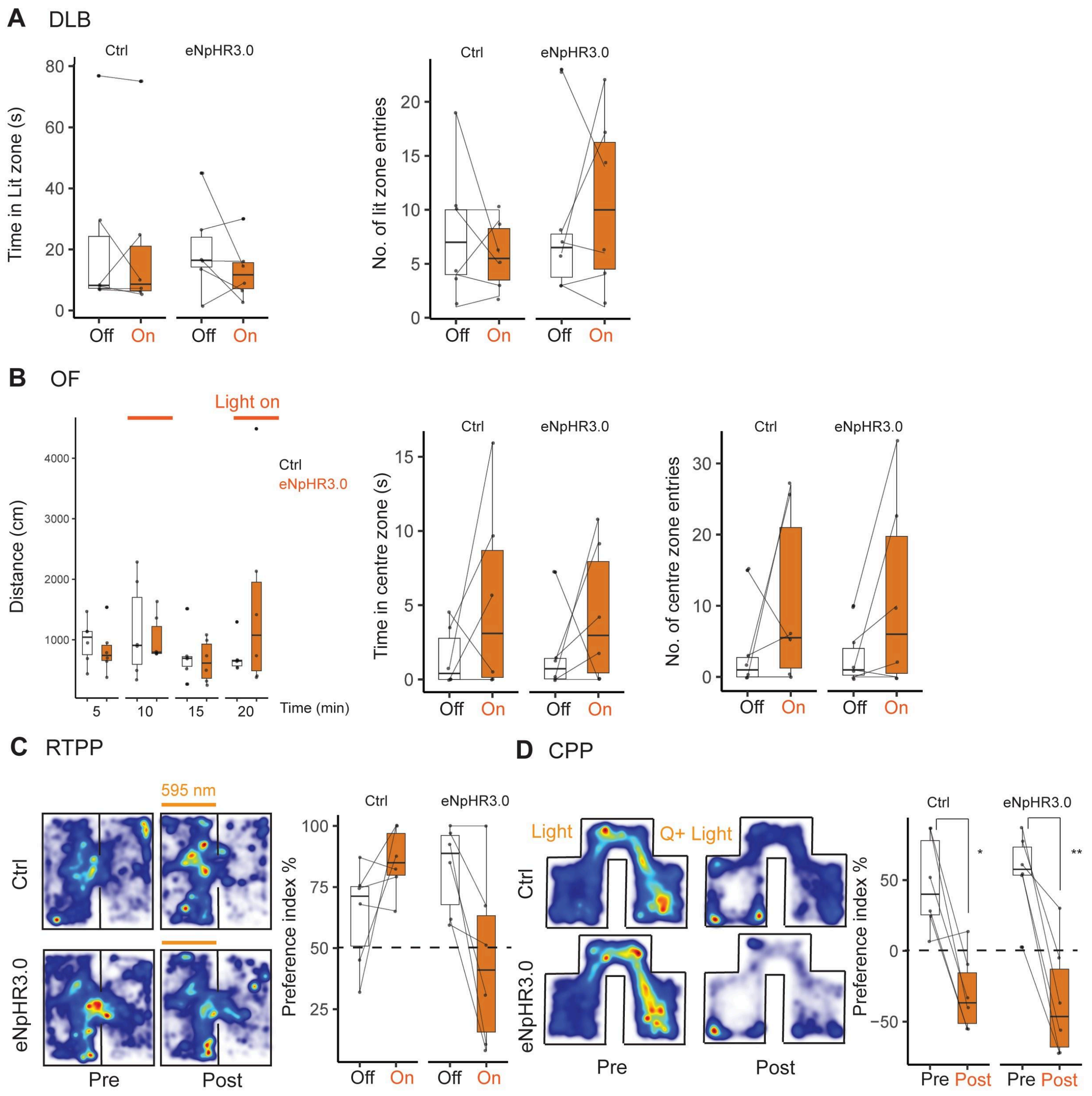

**Fig S14. Inhibition of SOC-TRAPed neurons has no influence on anxiety and place preference**

A. Inhibition of SOC-TRAPed ensembles in the IPACL does not affect anxiety in the DLB. There is no difference in time spent in the lit zone (Two-way ANOVA, effect of light, $F_{1,20}$ =

0.21, $p = 0.64$) and the number of lit zone entries (Two-way ANOVA, effect of light, $F_{1,20} = 0.001$, $p = 0.97$) between eNpHR3.0 and Ctrl animals. B. No difference in distance travelled during the OF (Two-way RM ANOVA, effect of light, $F_{1,10} = 0.29$, $p = 0.60$), time spent in centre zone (Two-way ANOVA, effect of light, $F_{1,20} = 3.32$, $p = 0.08$) or the number of centre zone entries (Two-way ANOVA, effect of light, $F_{1,20} = 3.78$, $p = 0.06$) when we inhibit SOC-TRAPed ensembles in the IPACL. C. Heatmap and box plots demonstrate there is no difference in place preference when we inhibit SOC-TRAPed ensembles (Two-way ANOVA, effect of light, $F_{1,20} = 0.71$, $p = 0.41$). D. Heatmap and box plots show there is no effect in quinine induced place aversion by inhibition of SOC-TRAPed ensembles during CPP (Two-way ANOVA, effect of opsin, $F_{1,20} = 0.01$, $p = 0.91$; effect of quinine, $F_{1,20} = 38.7$, $p < 0.0001$; Bonferroni's post-hoc test, Ctrl pre/post, $p = 0.004$, t = 4.03; eNpHR3.0 pre/post, $p = 0.0007$, t = 4.77). Value = median $\pm min/max$, $*p < 0.05$, $**p < 0.01$.

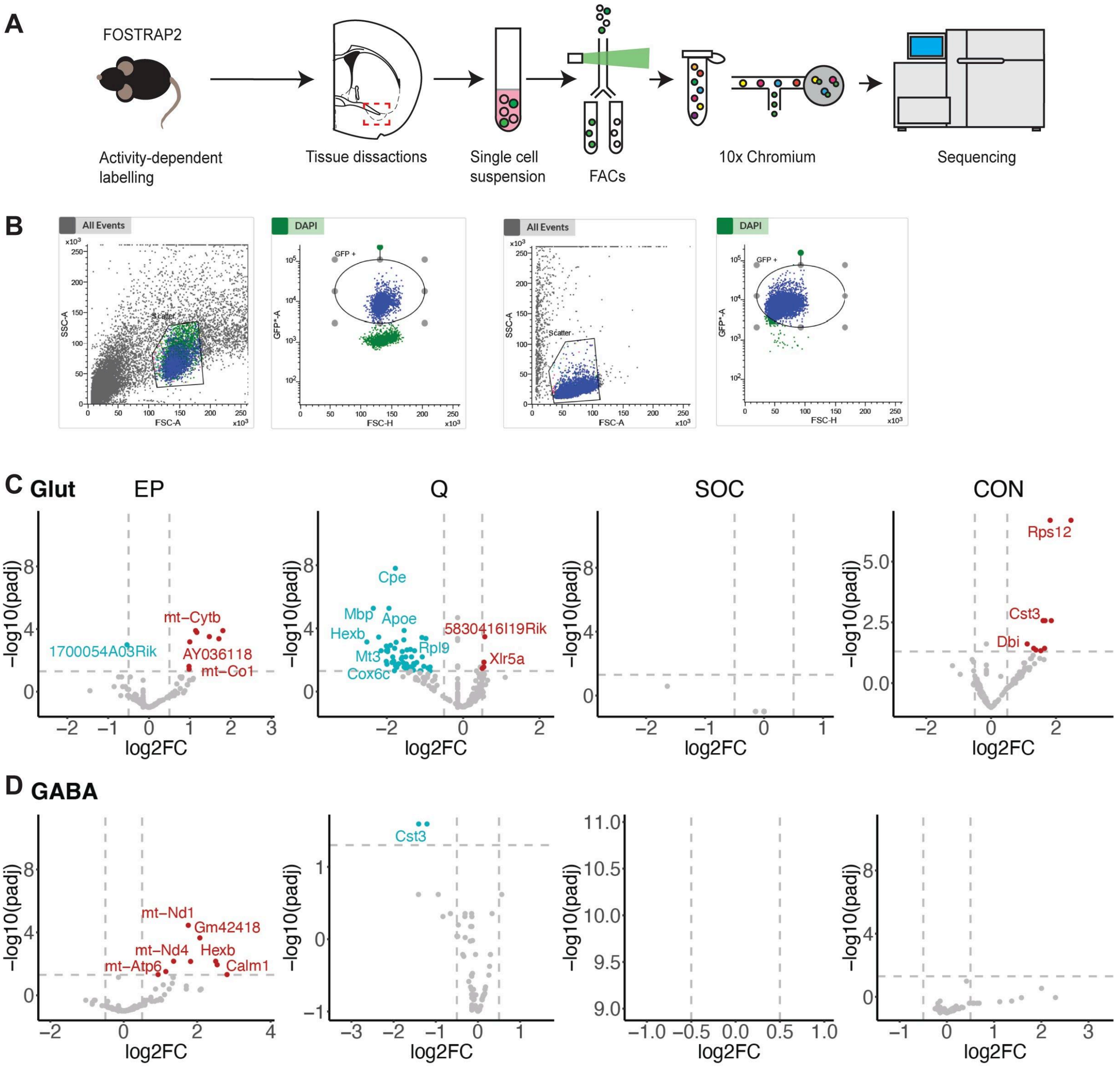

**Fig S15. Single-cell RNA sequencing in IPACL neuronal ensembles.**

A. Cartoon showing the different steps from activity-dependent cell labelling to FACs sorting and sequencing. B. Scatter plots demonstrating the selection of fluorescent positive cells during FACs sorting. C,D. Volcano plots indicating differentially expressed genes (DEGs) in GABA and Glut cells comparing one treatment with three other treatments. X-axis shows log2-fold change and y-axis shows -log10 p values. Genes survived multiple testing corrections and global corrections (10 times) are marked with colours, with blue indicating down-regulated genes and red indicating up-regulated genes.

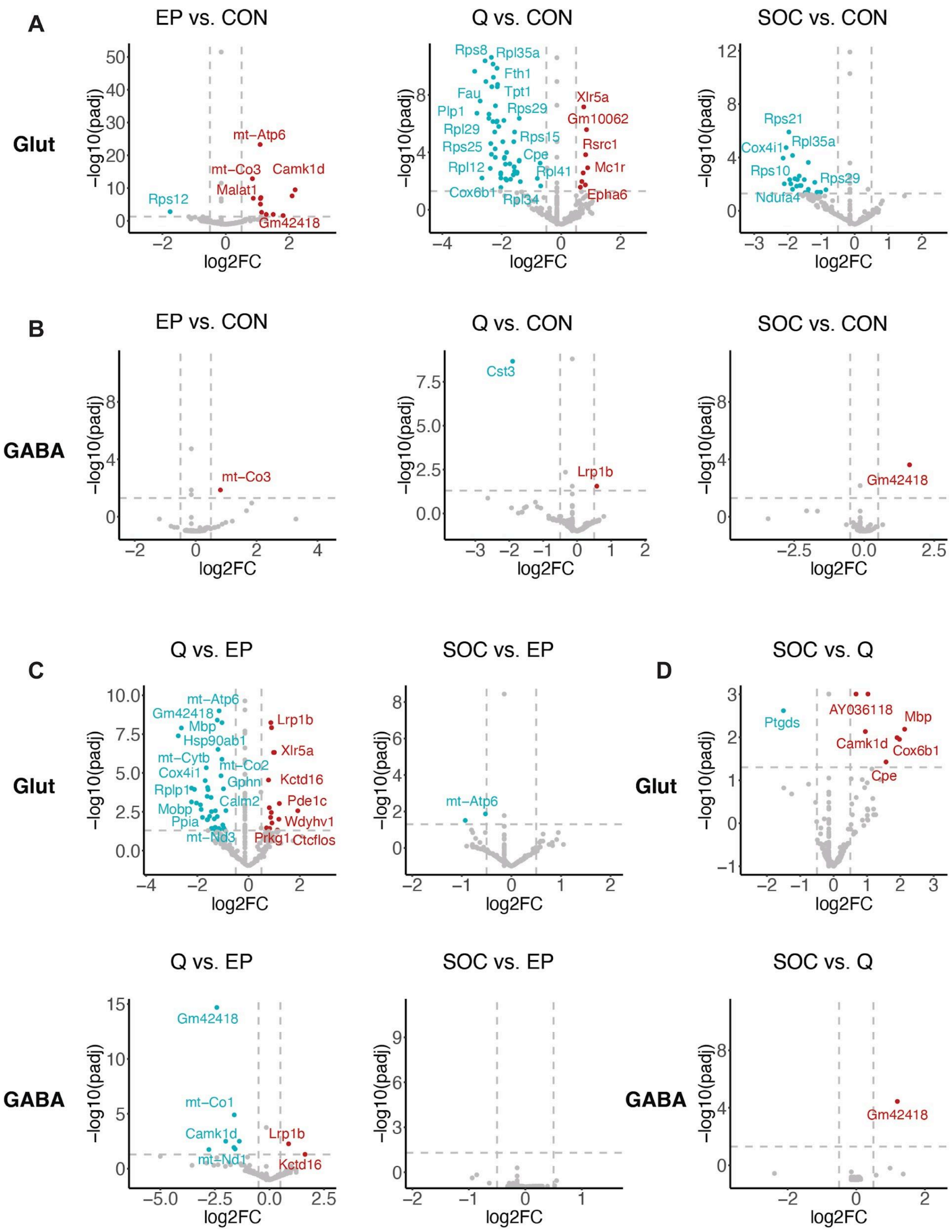

**Fig S16. Differential gene expression between two treatments**

A-D. Volcano plots indicating differentially expressed genes (DEGs) in GABA and Glut cells comparing one treatment with other treatments. X-axis shows log2-fold change and y-axis shows -log10 p values. Genes survived multiple testing corrections and global corrections (10 times) are marked with colours, with blue indicating down-regulated genes and red indicating up-regulated genes.

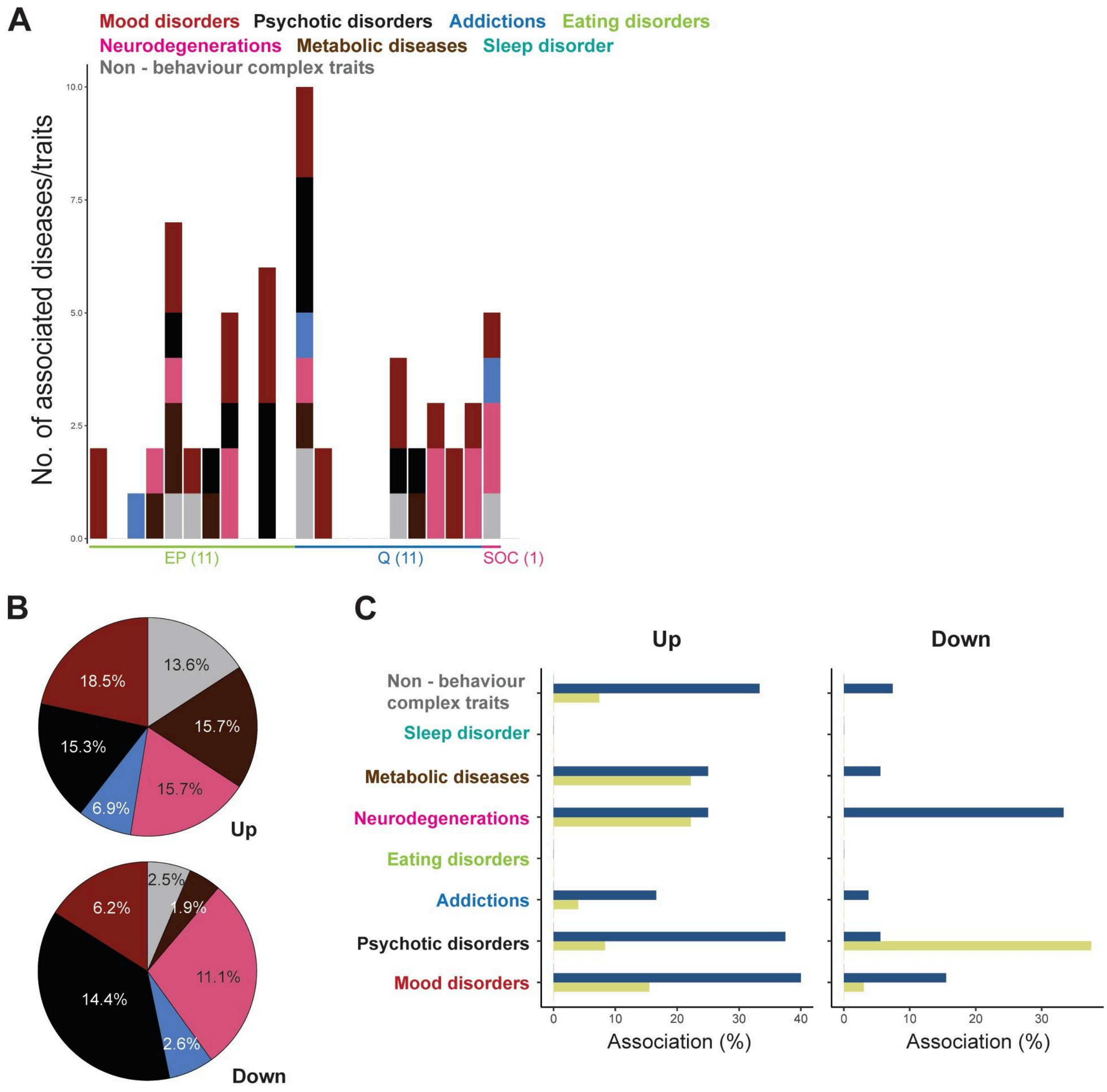

**Fig S17. Genes association with disease related traits in GABAergic neurons**

A. Stacked bars illustrate the association of genes with different human complex traits and diseases in GABA neurons. B. Mean percentages of up or down regulated genes linked to various categories of traits and diseases in GABA neurons. C. Bar graphs presenting the percentages of up and down regulated genes associated with diseases in all four context exposures.

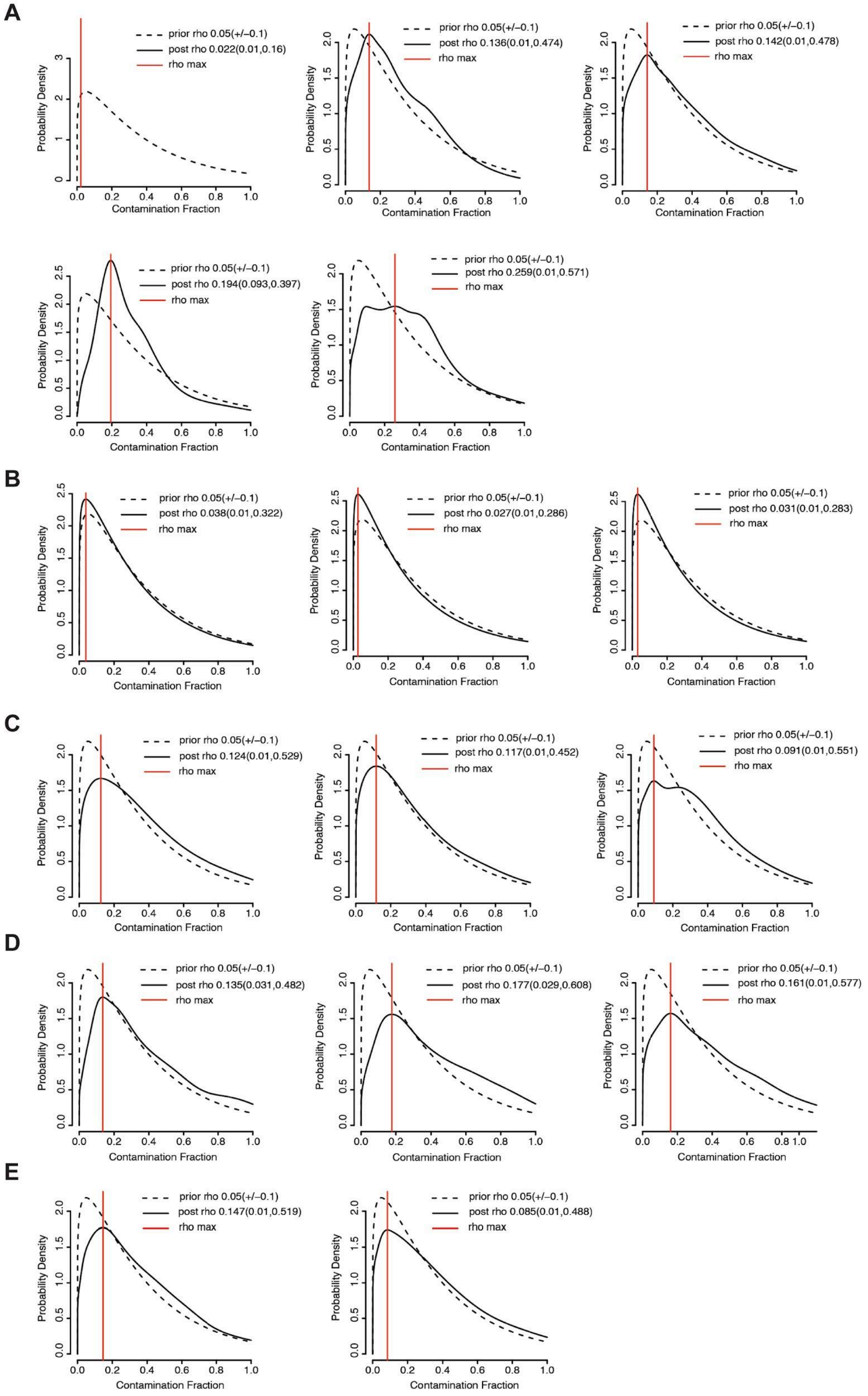
A
prior rho 0.05(+/−0.1)
post rho 0.022(0.01,0.16)
rho max
prior rho 0.05(+/−0.1)
post rho 0.136(0.01,0.474)
rho max
prior rho 0.05(+/−0.1)
post rho 0.142(0.01,0.478)
rho max
prior rho 0.05(+/−0.1)
post rho 0.194(0.093,0.397)
rho max
prior rho 0.05(+/−0.1)
post rho 0.259(0.01,0.571)
rho max
B
prior rho 0.05(+/−0.1)
post rho 0.038(0.01,0.322)
rho max
prior rho 0.05(+/−0.1)
post rho 0.027(0.01,0.286)
rho max
prior rho 0.05(+/−0.1)
post rho 0.031(0.01,0.283)
rho max
C
prior rho 0.05(+/−0.1)
post rho 0.124(0.01,0.529)
rho max
prior rho 0.05(+/−0.1)
post rho 0.117(0.01,0.452)
rho max
prior rho 0.05(+/−0.1)
post rho 0.091(0.01,0.551)
rho max
D
prior rho 0.05(+/−0.1)
post rho 0.135(0.031,0.482)
rho max
prior rho 0.05(+/−0.1)
post rho 0.177(0.029,0.608)
rho max
prior rho 0.05(+/−0.1)
post rho 0.161(0.01,0.577)
rho max
E
prior rho 0.05(+/−0.1)
post rho 0.147(0.01,0.519)
rho max
prior rho 0.05(+/−0.1)
post rho 0.085(0.01,0.488)
rho max
Probability Density
Contamination Fraction

**Fig S18. Correction of ambient RNA contamination fractions using SoupX.**

A-E. Graphical representations illustrate the estimated ambient RNA contamination fractions (parameterized as rho) for each sample through the application of SoupX, where rho = 0 means no contamination and rho = 1 means 100% of UMIs in a droplet are background RNA. The dashed curve shows prior rho values while the solid curve shows posterior rho values. The red lines indicate the peak rho values in each sample. A. controls; B. condensed milk; C. elevated platform; D.quinine; E. social.

**table S1: Differential gene expression list elevated platform compared to condensed milk in GABA and Glut neurons**

| **GABA** | baseMean | log2FoldChange | lfcSE | stat | pvalue | padj | pglobadj |
|---|---|---|---|---|---|---|---|
| Gm42418 | 63.825733 | 2.490023118 | 0.30885566 | 8.313449228 | 9.30E-17 | 2.36E-14 | 2.36E-13 |
| mt-Atp6 | 34.616814 | -0.144269504 | 0.533818667 | 5.667491383 | 1.45E-08 | 1.84E-06 | 1.84E-05 |
| mt-Co3 | 41.746395 | 0.810794634 | 0.207384986 | 4.246795332 | 2.17E-05 | 0.001376997 | 0.013769967 |
| Grik1 | 167.79897 | -0.144269504 | 0.533818461 | -4.305440953 | 1.67E-05 | 0.001376997 | 0.013769967 |
| mt-Co1 | 20.15368 | -0.144269504 | 0.53381867 | 4.023556335 | 5.73E-05 | 0.002912154 | 0.029121539 |
| AY036118 | 9.416064 | 1.838117441 | 0.606044425 | 3.647178454 | 0.000265136 | 0.011224082 | 0.112240818 |
| mt-Nd1 | 9.918921 | 1.667294864 | 0.662332181 | 3.27315204 | 0.001063553 | 0.038591763 | 0.385917627 |
| Hexb | 2.6150105 | -0.144269504 | 0.533818688 | 2.921216981 | 0.003486669 | 0.110701742 | 1.107017416 |
| Kctd16 | 9.053456 | -1.190333232 | 0.617155705 | -2.762385717 | 0.005738065 | 0.14574684 | 1.457468404 |
| Calm1 | 3.0364132 | 3.28937777 | 1.361120962 | 2.777428335 | 0.005479092 | 0.14574684 | 1.457468404 |
| Camk1d | 6.7925277 | 1.201390384 | 0.724956104 | 2.616575463 | 0.008881673 | 0.205085903 | 2.050859028 |
| mt-Nd4 | 10.174128 | 0.988646853 | 0.606080644 | 2.517500235 | 0.011819089 | 0.250170718 | 2.501707183 |
| 4930511M | 3.852436 | -0.144269504 | 0.533818668 | -2.41785241 | 0.015612408 | 0.30504243 | 3.050424303 |
| mt-Nd3 | 7.8861423 | 0.87137072 | 0.604798749 | 2.365963436 | 0.01798322 | 0.326266998 | 3.262669982 |
| Gria4 | 4.358638 | -0.809892788 | 1.180179421 | -2.212633035 | 0.026922956 | 0.455895382 | 4.558953816 |
| mt-Nd2 | 11.382935 | 0.740772557 | 0.62631066 | 2.163290577 | 0.030518837 | 0.484486544 | 4.84486544 |
| D430042O | 8.211102 | -0.703927442 | 0.774923213 | -2.087041712 | 0.036884364 | 0.551095794 | 5.51095794 |
| Gphn | 2.2153957 | 0.37418247 | 0.905976605 | 2.018391907 | 0.043550462 | 0.556494512 | 5.564945116 |
| mt-Cytb | 10.061957 | 0.635046631 | 0.568061436 | 2.015823211 | 0.043818465 | 0.556494512 | 5.564945116 |
| Rps15 | 3.859108 | -0.605173697 | 0.88491805 | -2.021297544 | 0.043248973 | 0.556494512 | 5.564945116 |
| Tpt1 | 3.266245 | 0.140427664 | 2.083030314 | 1.982382066 | 0.047436495 | 0.573755704 | 5.737557044 |
| Ckb | 2.7758057 | 0.438729997 | 0.851398523 | 1.942282459 | 0.052102915 | 0.575541591 | 5.755415914 |
| Epha6 | 7.4647217 | 0.596757487 | 0.589963887 | 1.942174567 | 0.052115971 | 0.575541591 | 5.755415914 |
| Rps20 | 2.0829651 | 0.310596938 | 0.863409493 | 1.894783672 | 0.058121067 | 0.615114629 | 6.151146285 |
| mt-Co2 | 20.432314 | 0.487156534 | 0.39699814 | 1.818759921 | 0.068948065 | 0.700512345 | 7.005123449 |
| Cpe | 1.806445 | 0.286941077 | 0.807175593 | 1.768730548 | 0.076938852 | 0.723795125 | 7.237951252 |
| Rpl23 | 3.024376 | 0.165974801 | 1.111257338 | 1.781628013 | 0.074809913 | 0.723795125 | 7.237951252 |
| Cox6b1 | 2.4086268 | 0.336556121 | 0.777589969 | 1.737801833 | 0.082245738 | 0.746086333 | 7.460863333 |
| Fyb2 | 2.214498 | -0.305171794 | 0.631788099 | -1.660758986 | 0.096761861 | 0.847500442 | 8.475004421 |
| Cox5b | 1.8724982 | -0.100431436 | 0.477770134 | -1.374461596 | 0.169298425 | 0.914530483 | 9.145304832 |
| Cox7c | 3.3538988 | 0.27948499 | 0.765364827 | 1.600505081 | 0.109486581 | 0.914530483 | 9.145304832 |
| Sil1 | 3.97458 | -0.144269504 | 0.533818674 | -1.390079999 | 0.164504586 | 0.914530483 | 9.145304832 |
| Sparcl1 | 1.0810266 | 0.186424603 | 1.049550019 | 1.578932579 | 0.114351522 | 0.914530483 | 9.145304832 |
| Gpx4 | 2.040009 | -0.076847607 | 0.466512283 | -1.289844304 | 0.197104722 | 0.914530483 | 9.145304832 |
| Cox4i1 | 3.030142 | 0.222117238 | 0.702217016 | 1.37451452 | 0.169282006 | 0.914530483 | 9.145304832 |
| Rpl39 | 1.735824 | -0.170797002 | 0.54698026 | -1.257000749 | 0.208753364 | 0.914530483 | 9.145304832 |
| Rps10 | 1.1794552 | -0.197971982 | 0.549816107 | -1.441975642 | 0.149309245 | 0.914530483 | 9.145304832 |
| Rps12 | 3.3723876 | -0.131111299 | 0.513466472 | -1.334258522 | 0.182119146 | 0.914530483 | 9.145304832 |
| Rps7 | 1.0332989 | 0.088826085 | 1.387758125 | 1.245545519 | 0.212931294 | 0.914530483 | 9.145304832 |
| Fth1 | 9.076847 | -0.258598728 | 0.565167378 | -1.327827904 | 0.184234966 | 0.914530483 | 9.145304832 |
| Csmd3 | 4.105139 | -0.144269504 | 0.533818667 | -1.386032691 | 0.165736922 | 0.914530483 | 9.145304832 |
| Cntnap2 | 9.239207 | 0.289185396 | 0.550463933 | 1.260596564 | 0.207454237 | 0.914530483 | 9.145304832 |
| Rpl30 | 3.7069292 | -0.207310615 | 0.561537005 | -1.566165881 | 0.117309788 | 0.914530483 | 9.145304832 |
| Tmsb4x | 3.28174 | -0.243458436 | 0.556002355 | -1.255174638 | 0.20941537 | 0.914530483 | 9.145304832 |
| Mbp | 2.6833713 | 0.242450013 | 0.755443603 | 1.501984508 | 0.13310111 | 0.914530483 | 9.145304832 |
| Mapk14 | 5.426243 | -0.144269504 | 0.533818664 | -1.282040134 | 0.199828568 | 0.914530483 | 9.145304832 |
| Xlr5a | 14.715869 | -0.348573062 | 0.41969787 | -1.373299726 | 0.169659188 | 0.914530483 | 9.145304832 |
| Wdr72 | 3.3485434 | -0.144269504 | 0.533818686 | 1.423981724 | 0.154451763 | 0.914530483 | 9.145304832 |
| 4930441J1 | 2.2203438 | -0.247908285 | 0.591482277 | -1.536414125 | 0.124436845 | 0.914530483 | 9.145304832 |
| Tmeff2 | 2.6209083 | 0.222749754 | 0.684126173 | 1.372395968 | 0.169940205 | 0.914530483 | 9.145304832 |
| Lrp1b | 26.611061 | -0.292212813 | 0.252804937 | -1.411379573 | 0.158132723 | 0.914530483 | 9.145304832 |
| Atp5k | 1.603309 | 0.116958908 | 0.87191644 | 1.237151555 | 0.216030823 | 0.914530483 | 9.145304832 |
| Mrc2 | 4.0515614 | -0.328741224 | 0.585305145 | -1.462082148 | 0.1437187 | 0.914530483 | 9.145304832 |
| Atp9b | 3.812995 | -0.297632748 | 0.568290817 | -1.372412544 | 0.169935047 | 0.914530483 | 9.145304832 |
| Rpl3 | 0.9445738 | 0.136709937 | 0.723471021 | 1.260783245 | 0.207386952 | 0.914530483 | 9.145304832 |
| Cdk8 | 2.7248116 | 0.260653961 | 0.704230709 | 1.491930219 | 0.13571744 | 0.914530483 | 9.145304832 |
| Mt1 | 1.5199275 | 0.170589659 | 0.754077005 | 1.392841373 | 0.163667764 | 0.914530483 | 9.145304832 |
| Cep128 | 1.3127552 | -0.171734188 | 0.54789223 | -1.302841475 | 0.192628888 | 0.914530483 | 9.145304832 |
| Igf2bp3 | 5.202918 | -0.292899776 | 0.51412113 | -1.258546426 | 0.208194208 | 0.914530483 | 9.145304832 |

| | | | | | | | |
|---|---|---|---|---|---|---|---|
| Uqcrh | 1.2902656 | -0.144269504 | 0.53381868 | 1.301097435 | 0.193225106 | 0.914530483 | 9.145304832 |
| Rpl35 | 1.1383914 | 0.105412354 | 0.894181582 | 1.220329127 | 0.222340132 | 0.925809731 | 9.258097311 |
| Rab29 | 2.2901654 | -0.15234968 | 0.537152473 | -1.172637644 | 0.240941147 | 0.9601486 | 9.601486004 |
| Camk2a | 2.641178 | -0.211099176 | 0.547886276 | -1.160839149 | 0.245707319 | 0.9601486 | 9.601486004 |
| Sec61a1 | 32.313854 | -0.286722053 | 0.403439636 | -1.178025401 | 0.23878648 | 0.9601486 | 9.601486004 |
| Rps28 | 2.9921 | -0.186894869 | 0.545345133 | -1.162357152 | 0.24509042 | 0.9601486 | 9.601486004 |
| Rpl12 | 1.3417516 | 0.118076132 | 0.749496137 | 1.140680821 | 0.254002771 | 0.964448377 | 9.644483771 |
| Gm49497 | 1.5720488 | -0.171928707 | 0.550333957 | -1.139722963 | 0.254401737 | 0.964448377 | 9.644483771 |
| Ptprd | 1.4235657 | 0.104344724 | 0.808577169 | 1.111270936 | 0.266451746 | 0.974188459 | 9.741884591 |
| Lyrm1 | 0.71984035 | 0.107308951 | 0.693286711 | 1.015748331 | 0.309749246 | 0.974188459 | 9.741884591 |
| Hdac9 | 0.7884868 | 0.098566385 | 0.687761987 | 1.022490527 | 0.306548794 | 0.974188459 | 9.741884591 |
| Rps14 | 2.1608822 | 0.059645778 | 1.154819973 | 1.053552831 | 0.292087694 | 0.974188459 | 9.741884591 |
| 1700054A( | 30.060472 | -0.250974093 | 0.476230781 | -1.087713073 | 0.276721795 | 0.974188459 | 9.741884591 |
| Nkain2 | 1.7783517 | 0.139331523 | 0.672870368 | 1.098151341 | 0.27213841 | 0.974188459 | 9.741884591 |
| Ctnna3 | 4.2239866 | -0.219892757 | 0.506491138 | -1.037913605 | 0.299310279 | 0.974188459 | 9.741884591 |
| Ttc28 | 2.4535105 | -0.158800852 | 0.537414379 | -1.017333967 | 0.308994583 | 0.974188459 | 9.741884591 |
| Uba52 | 1.3606701 | 0.108197519 | 0.703788081 | 1.013824718 | 0.310666398 | 0.974188459 | 9.741884591 |
| Chsy3 | 4.910581 | -0.214298495 | 0.503521307 | -1.020979503 | 0.30726415 | 0.974188459 | 9.741884591 |
| Fau | 1.8035932 | 0.118277851 | 0.737136404 | 1.070571836 | 0.284361993 | 0.974188459 | 9.741884591 |
| Vit | 0.7298555 | 0.105771222 | 0.69166722 | 1.017194888 | 0.309060727 | 0.974188459 | 9.741884591 |
| Ctnna2 | 3.4546864 | -0.187667341 | 0.542046147 | -1.071522469 | 0.283934573 | 0.974188459 | 9.741884591 |
| Shb | 8.229129 | -0.239012592 | 0.469129907 | -1.045237098 | 0.295913402 | 0.974188459 | 9.741884591 |
| Tshz2 | 5.412102 | 0.214362436 | 0.494046098 | 0.967018062 | 0.333535005 | 0.97584087 | 9.758408697 |
| Rpl11 | 3.7124317 | -0.100506579 | 0.523226041 | -0.942825069 | 0.345770387 | 0.97584087 | 9.758408697 |
| Rpl13a | 1.0170885 | 0.104318731 | 0.672240853 | 0.944987995 | 0.344665005 | 0.97584087 | 9.758408697 |
| Alpl | 1.3248256 | 0.132024704 | 0.620702306 | 0.955103437 | 0.339525391 | 0.97584087 | 9.758408697 |
| Ccdc60 | 1.6695045 | -0.144269504 | 0.533818675 | -0.988876229 | 0.322723702 | 0.97584087 | 9.758408697 |
| Gm11337 | 2.8759263 | -0.167219146 | 0.528566687 | -0.946254205 | 0.344018945 | 0.97584087 | 9.758408697 |
| 4930558K( | 3.2298863 | 0.171615217 | 0.576727731 | 0.96558422 | 0.334252275 | 0.97584087 | 9.758408697 |
| Pcsk1n | 2.002329 | -0.148791628 | 0.536014775 | -0.996423929 | 0.319044211 | 0.97584087 | 9.758408697 |
| 4930402H2 | 2.5481591 | -0.149385613 | 0.534803749 | -0.952727061 | 0.340728379 | 0.97584087 | 9.758408697 |
| Rps29 | 9.16326 | 0.148447656 | 0.640355317 | 0.931284262 | 0.351706541 | 0.981686388 | 9.816863884 |
| Apoe | 2.9698837 | -0.150263772 | 0.527535703 | -0.911180811 | 0.362200113 | 0.989234718 | 9.892347176 |
| Tacc1 | 0.78005695 | 0.078866627 | 0.772883031 | 0.917116456 | 0.359081619 | 0.989234718 | 9.892347176 |
| Pcdh15 | 6.3310084 | 0.115757691 | 0.486307316 | 0.576321231 | 0.564398075 | 0.991797939 | 9.917979392 |
| Pde6a | 9.454883 | -0.107425478 | 0.473891824 | -0.534774405 | 0.592805864 | 0.991797939 | 9.917979392 |
| Ppp1r16a | 3.5661983 | -0.143067924 | 0.52653696 | -0.845247194 | 0.397972837 | 0.991797939 | 9.917979392 |
| Zfhx3 | 2.2990522 | -0.063077972 | 0.534878604 | -0.516918426 | 0.605213105 | 0.991797939 | 9.917979392 |
| Vwc2l | 2.1770017 | 0.103899783 | 0.610031183 | 0.747854665 | 0.454547825 | 0.991797939 | 9.917979392 |
| Ptpn9 | 3.7900143 | -0.11525841 | 0.504815135 | -0.644538063 | 0.519226584 | 0.991797939 | 9.917979392 |
| Ppp1r9a | 2.6486123 | -0.073621499 | 0.524938238 | -0.523028243 | 0.600954599 | 0.991797939 | 9.917979392 |
| Plp1 | 1.951137 | 0.030088165 | 1.173338015 | 0.724844187 | 0.468547625 | 0.991797939 | 9.917979392 |
| Ppia | 1.5618752 | 0.066427617 | 0.658406198 | 0.63719586 | 0.523997274 | 0.991797939 | 9.917979392 |
| Ubb | 0.71324944 | -0.040915157 | 0.534484077 | -0.496133594 | 0.619800159 | 0.991797939 | 9.917979392 |
| Rps2 | 1.735171 | -0.097674302 | 0.532830694 | -0.890817237 | 0.373027227 | 0.991797939 | 9.917979392 |
| Trim2 | 5.161488 | 0.148246708 | 0.530816518 | 0.764584949 | 0.444518734 | 0.991797939 | 9.917979392 |
| Rpl27a | 3.7344685 | 0.077986846 | 0.605551044 | 0.555848533 | 0.578314404 | 0.991797939 | 9.917979392 |
| Rps19 | 1.6213824 | -0.048421887 | 0.487984147 | -0.868512705 | 0.38511372 | 0.991797939 | 9.917979392 |
| Rps18 | 0.55254894 | 0.061082947 | 1.243680717 | 0.864714913 | 0.387195282 | 0.991797939 | 9.917979392 |
| Rps17 | 1.3156979 | 0.042039933 | 0.897875378 | 0.707898165 | 0.4790085 | 0.991797939 | 9.917979392 |
| Rps24 | 2.3036187 | -0.124273805 | 0.526617307 | -0.81403675 | 0.415623898 | 0.991797939 | 9.917979392 |
| Ntm | 6.072977 | 0.092859256 | 0.503285937 | 0.47710546 | 0.633287027 | 0.991797939 | 9.917979392 |
| Rsrc1 | 7.085821 | -0.115697027 | 0.468444007 | -0.575234382 | 0.565132793 | 0.991797939 | 9.917979392 |
| Scamp1 | 2.2210886 | -0.139599458 | 0.52350302 | -0.838805269 | 0.401578594 | 0.991797939 | 9.917979392 |
| Rplp2 | 0.64398235 | 0.029374432 | 0.787761907 | 0.561464548 | 0.574480892 | 0.991797939 | 9.917979392 |
| Rpl27 | 0.5654194 | -0.057261298 | 0.478796288 | -0.70346356 | 0.481766908 | 0.991797939 | 9.917979392 |
| Rplp1 | 5.602014 | -0.073545231 | 0.521505874 | -0.477452294 | 0.633040084 | 0.991797939 | 9.917979392 |
| Rpl38 | 5.5418315 | 0.082277441 | 0.588342687 | 0.540657535 | 0.588743653 | 0.991797939 | 9.917979392 |
| Rpl37 | 3.905646 | 0.102303306 | 0.555390395 | 0.614425185 | 0.538934398 | 0.991797939 | 9.917979392 |
| Slc25a4 | 0.91304207 | -0.087466848 | 0.533108408 | -0.82220456 | 0.410960484 | 0.991797939 | 9.917979392 |
| Srgn | 1.6207895 | 0.075244441 | 0.599781289 | 0.559893707 | 0.575551941 | 0.991797939 | 9.917979392 |
| Rpl36 | 0.4683596 | -0.032796344 | 0.468431387 | -0.489619755 | 0.624402995 | 0.991797939 | 9.917979392 |

| | | | | | | | |
|---|---|---|---|---|---|---|---|
| Syt1 | 1.0789711 | -0.052397438 | 0.536779663 | -0.482644605 | 0.629348102 | 0.991797939 | 9.917979392 |
| Tns3 | 1.3923346 | -0.144269504 | 0.53381868 | -0.591155962 | 0.554415927 | 0.991797939 | 9.917979392 |
| Rps21 | 5.4247017 | -0.144269504 | 0.533818672 | -0.593154798 | 0.55307756 | 0.991797939 | 9.917979392 |
| Rpl41 | 3.506254 | -0.141912728 | 0.527827129 | -0.889688956 | 0.373632926 | 0.991797939 | 9.917979392 |
| Rpsa | 3.0339682 | -0.085208661 | 0.528038637 | -0.656659835 | 0.511399666 | 0.991797939 | 9.917979392 |
| Ninj2 | 2.7502337 | -0.161396015 | 0.515477288 | -0.880683341 | 0.378489236 | 0.991797939 | 9.917979392 |
| Nell1 | 2.1040456 | -0.08770312 | 0.533878122 | -0.642196999 | 0.520745281 | 0.991797939 | 9.917979392 |
| Cdh23 | 3.9961262 | 0.113927715 | 0.52776642 | 0.611807277 | 0.540665274 | 0.991797939 | 9.917979392 |
| Cfap74 | 1.6633576 | -0.078211063 | 0.531042157 | -0.646248436 | 0.518118476 | 0.991797939 | 9.917979392 |
| Kat7 | 1.8998642 | -0.076433243 | 0.51533783 | -0.485136257 | 0.627579695 | 0.991797939 | 9.917979392 |
| Chl1 | 2.9479158 | -0.145371675 | 0.518755962 | -0.834791149 | 0.40383531 | 0.991797939 | 9.917979392 |
| Ehbp1 | 5.512093 | -0.096788643 | 0.502809311 | -0.542338502 | 0.587585337 | 0.991797939 | 9.917979392 |
| Cmss1 | 3.2268436 | 0.129520764 | 0.524288697 | 0.692135384 | 0.488852312 | 0.991797939 | 9.917979392 |
| Hdac8 | 2.131761 | -0.080095553 | 0.528001217 | -0.564527307 | 0.572395321 | 0.991797939 | 9.917979392 |
| Cox8a | 5.6540856 | -0.065558653 | 0.533969738 | -0.518948395 | 0.603796728 | 0.991797939 | 9.917979392 |
| Gria1 | 0.8403544 | -0.089211526 | 0.533122748 | -0.799798347 | 0.423827641 | 0.991797939 | 9.917979392 |
| Gpm6b | 0.8889708 | -0.059734259 | 0.526017642 | -0.695162377 | 0.486953547 | 0.991797939 | 9.917979392 |
| Cspp1 | 1.3673939 | -0.043694633 | 0.533329505 | -0.489817239 | 0.624263231 | 0.991797939 | 9.917979392 |
| Kcnh7 | 0.8303394 | -0.082141766 | 0.531475877 | -0.767862798 | 0.442568701 | 0.991797939 | 9.917979392 |
| Cst3 | 15.013968 | -0.144269504 | 0.533818649 | -0.756373786 | 0.44942512 | 0.991797939 | 9.917979392 |
| Gm50306 | 3.4517357 | 0.103090627 | 0.615768347 | 0.697969338 | 0.48519637 | 0.991797939 | 9.917979392 |
| Gm38973 | 2.7419732 | 0.094212006 | 0.609417461 | 0.669034068 | 0.503473746 | 0.991797939 | 9.917979392 |
| Gm14582 | 2.8478622 | -0.100104564 | 0.499246867 | -0.561075701 | 0.574745932 | 0.991797939 | 9.917979392 |
| Gm12216 | 14.620876 | -0.166003237 | 0.361896509 | -0.723436 | 0.469412061 | 0.991797939 | 9.917979392 |
| Gm10062 | 5.3520646 | -0.157802763 | 0.430098578 | -0.708246569 | 0.478792151 | 0.991797939 | 9.917979392 |
| Dctd | 1.2889811 | 0.065416147 | 0.625835579 | 0.558046951 | 0.576812324 | 0.991797939 | 9.917979392 |
| Gapdh | 0.6887045 | -0.144269504 | 0.533818683 | -0.794605834 | 0.426842824 | 0.991797939 | 9.917979392 |
| mt-Nd4l | 1.4160243 | 0.046155893 | 0.942339274 | 0.783033019 | 0.433607723 | 0.991797939 | 9.917979392 |
| Dlg2 | 1.0649148 | -0.068178861 | 0.533992943 | -0.564333047 | 0.572527495 | 0.991797939 | 9.917979392 |
| Dpyd | 2.2141922 | 0.090691675 | 0.546055019 | 0.554381481 | 0.579317798 | 0.991797939 | 9.917979392 |
| Eef1a1 | 2.482557 | -0.106437947 | 0.528733726 | -0.764621689 | 0.44449685 | 0.991797939 | 9.917979392 |
| Gm765 | 1.2246474 | -0.084589818 | 0.532232977 | -0.694923564 | 0.487103204 | 0.991797939 | 9.917979392 |
| Bc1 | 1.6728477 | -0.055876584 | 0.536222478 | -0.466525909 | 0.640839107 | 0.991797939 | 9.917979392 |
| Cmpk1 | 7.641893 | -0.144269504 | 0.53381867 | -0.461608359 | 0.644362199 | 0.991797939 | 9.917979392 |
| Kcnip4 | 1.5396427 | -0.062131264 | 0.529870523 | -0.456287982 | 0.648182905 | 0.991797939 | 9.917979392 |
| Adk | 2.3225515 | -0.144269504 | 0.533818686 | 0.821970116 | 0.411093905 | 0.991797939 | 9.917979392 |
| AI314278 | 2.6000702 | 0.088171633 | 0.528722523 | 0.510222974 | 0.609895259 | 0.991797939 | 9.917979392 |
| Lrrc17 | 2.6446164 | -0.111514125 | 0.487336364 | -0.598803535 | 0.549303905 | 0.991797939 | 9.917979392 |
| Mt3 | 3.5244105 | 0.075030972 | 0.547763476 | 0.461710227 | 0.644289136 | 0.991797939 | 9.917979392 |
| 6030443J0 | 3.9329336 | -0.134952407 | 0.49215843 | -0.703890521 | 0.481500955 | 0.991797939 | 9.917979392 |
| Malat1 | 4.9240985 | 0.044343918 | 0.73489692 | 0.526426658 | 0.598591798 | 0.991797939 | 9.917979392 |
| Atp5j2 | 1.2947026 | -0.037446423 | 0.502748501 | -0.701547184 | 0.482961599 | 0.991797939 | 9.917979392 |
| Nckap5 | 10.468728 | -0.15849632 | 0.414114941 | -0.703545532 | 0.481715842 | 0.991797939 | 9.917979392 |
| 1700126G | 5.0824046 | -0.144269504 | 0.533818666 | -0.687254861 | 0.491922138 | 0.991797939 | 9.917979392 |
| Arvcf | 0.3312396 | -0.060308461 | 0.497705482 | -0.529539869 | 0.596430995 | 0.991797939 | 9.917979392 |
| Selenow | 1.1922412 | 0.00453741 | 0.56347535 | 0.040646509 | 0.967577706 | 0.998757862 | 9.987578619 |
| Sema3c | 12.56537 | -0.144269504 | 0.53381865 | -0.428742627 | 0.668110536 | 0.998757862 | 9.987578619 |
| Slx1b | 2.8473423 | -0.061887045 | 0.516274444 | -0.400498217 | 0.688789596 | 0.998757862 | 9.987578619 |
| 8430422H | 4.2783537 | 0.018700167 | 0.500621781 | 0.102023958 | 0.918737659 | 0.998757862 | 9.987578619 |
| Dop1b | 1.3326868 | 0.023630755 | 0.559385991 | 0.180327571 | 0.856895412 | 0.998757862 | 9.987578619 |
| 5830416I1 | 11.9921875 | -0.080020522 | 0.356633062 | -0.358575446 | 0.719912718 | 0.998757862 | 9.987578619 |
| Acoxl | 1.8561587 | -0.006498881 | 0.532268736 | -0.045363082 | 0.963817907 | 0.998757862 | 9.987578619 |
| Adamtsl1 | 4.5373645 | 0.047827364 | 0.46456604 | 0.240369556 | 0.810043777 | 0.998757862 | 9.987578619 |
| Dbi | 1.9905039 | -0.004094997 | 0.591251214 | -0.048636393 | 0.961209067 | 0.998757862 | 9.987578619 |
| Ctcflos | 2.801382 | -0.006662115 | 0.543309016 | -0.047276652 | 0.962292736 | 0.998757862 | 9.987578619 |
| Sgip1 | 1.0956551 | -0.144269504 | 0.533818679 | 0.198594897 | 0.842579646 | 0.998757862 | 9.987578619 |
| Zmat4 | 3.9567978 | 0.01676833 | 0.493136819 | 0.091516698 | 0.927082038 | 0.998757862 | 9.987578619 |
| Unc5d | 7.3008747 | -0.021805229 | 0.436386978 | -0.107535762 | 0.914363956 | 0.998757862 | 9.987578619 |
| Cadm2 | 0.1170899 | -0.021584505 | 0.498617419 | -0.075556423 | 0.939772007 | 0.998757862 | 9.987578619 |
| Car12 | 2.7007744 | 0.035872248 | 0.611853349 | 0.314379919 | 0.753232506 | 0.998757862 | 9.987578619 |
| Uqcr11 | 0.1170899 | -0.021584505 | 0.498617419 | -0.075556423 | 0.939772007 | 0.998757862 | 9.987578619 |
| Atxn7l1 | 1.0454535 | 0.011091764 | 0.561214084 | 0.09182296 | 0.926838702 | 0.998757862 | 9.987578619 |

| | | | | | | | |
|---|---|---|---|---|---|---|---|
| Anks1b | 0.9038186 | 0.023821994 | 0.633267508 | 0.280532606 | 0.779068914 | 0.998757862 | 9.987578619 |
| Wdyhv1 | 3.1423538 | -0.049029509 | 0.507488579 | -0.296661193 | 0.766725187 | 0.998757862 | 9.987578619 |
| Tmem108 | 1.8043896 | -0.035650711 | 0.525345412 | -0.249651148 | 0.80285714 | 0.998757862 | 9.987578619 |
| Cntn3 | 1.8330709 | 0.039981661 | 0.557515624 | 0.290192856 | 0.771668701 | 0.998757862 | 9.987578619 |
| Ankrd12 | 4.783603 | -0.059716464 | 0.497906515 | -0.3341868 | 0.738238604 | 0.998757862 | 9.987578619 |
| Cox6c | 2.4186468 | -0.055458188 | 0.51355012 | -0.357865926 | 0.720443651 | 0.998757862 | 9.987578619 |
| Svil | 1.5211977 | -0.043789344 | 0.531276628 | -0.337895507 | 0.735441934 | 0.998757862 | 9.987578619 |
| Zbtb7a | 3.031286 | 0.042978678 | 0.486743939 | 0.233030133 | 0.815738007 | 0.998757862 | 9.987578619 |
| Sult2a2 | 1.6488713 | 0.002130822 | 0.557635963 | 0.016342052 | 0.986961509 | 0.998757862 | 9.987578619 |
| Aldoc | 0.918843 | -0.019379497 | 0.512446705 | -0.424768047 | 0.671005773 | 0.998757862 | 9.987578619 |
| St18 | 1.1581548 | 0.003081371 | 0.554314573 | 0.025696187 | 0.979499665 | 0.998757862 | 9.987578619 |
| Zfp994 | 1.9295621 | -0.03467183 | 0.535640795 | -0.269555127 | 0.787502526 | 0.998757862 | 9.987578619 |
| Spock1 | 2.6526794 | 0.003952696 | 0.524912011 | 0.023589921 | 0.981179712 | 0.998757862 | 9.987578619 |
| Ahdc1 | 1.6529126 | -0.144269504 | 0.533818674 | 0.05251204 | 0.958120702 | 0.998757862 | 9.987578619 |
| Rps5 | 2.5000947 | 0.016981011 | 0.631173595 | 0.188105978 | 0.850793575 | 0.998757862 | 9.987578619 |
| Fbxl7 | 2.1408079 | 0.049180103 | 0.537166684 | 0.315954909 | 0.752036734 | 0.998757862 | 9.987578619 |
| Rps3a1 | 5.4120173 | 0.042875234 | 0.679837023 | 0.441371496 | 0.658944074 | 0.998757862 | 9.987578619 |
| Rpl24 | 0.43888316 | -0.016644063 | 0.568002801 | -0.12558895 | 0.90005731 | 0.998757862 | 9.987578619 |
| Rpl23a | 0.7610722 | 0.029194892 | 0.664613996 | 0.361808539 | 0.717495111 | 0.998757862 | 9.987578619 |
| Rpl21 | 2.4972112 | 0.043045874 | 0.589767472 | 0.33064671 | 0.74091136 | 0.998757862 | 9.987578619 |
| Rpl17 | 1.9896317 | 0.055646963 | 0.576361919 | 0.406172061 | 0.684616181 | 0.998757862 | 9.987578619 |
| Rpl13 | 4.1371584 | -0.04520575 | 0.533719226 | -0.315239706 | 0.752579661 | 0.998757862 | 9.987578619 |
| Kcnq5 | 1.2124442 | 0.012835099 | 0.56661147 | 0.111816827 | 0.910968645 | 0.998757862 | 9.987578619 |
| Klhl14 | 14.023906 | 0.028370516 | 0.358242314 | 0.12640236 | 0.899413434 | 0.998757862 | 9.987578619 |
| Rapgef4 | 7.5327554 | 0.024556757 | 0.430983605 | 0.115415656 | 0.90811567 | 0.998757862 | 9.987578619 |
| Ralyl | 1.865873 | 0.011569519 | 0.539720887 | 0.07936157 | 0.936745035 | 0.998757862 | 9.987578619 |
| Ptprg | 1.2726609 | 0.017890653 | 0.569496052 | 0.158928174 | 0.873725463 | 0.998757862 | 9.987578619 |
| Lrrc4c | 0.6306417 | 0.022928927 | 0.626631529 | 0.28313194 | 0.777075698 | 0.998757862 | 9.987578619 |
| Prkg1 | 3.3030834 | -0.045170007 | 0.507816253 | -0.270541043 | 0.786744046 | 0.998757862 | 9.987578619 |
| Prkag2 | 2.3407285 | -0.144269504 | 0.533818682 | -0.138389666 | 0.889932464 | 0.998757862 | 9.987578619 |
| Ppp3ca | 0.7254184 | 0.018773374 | 0.628904198 | 0.246954188 | 0.804943679 | 0.998757862 | 9.987578619 |
| Lsamp | 2.6785412 | 0.038917078 | 0.577271177 | 0.280612335 | 0.779007754 | 0.998757862 | 9.987578619 |
| Mc1r | 4.462505 | -0.012481096 | 0.478661493 | -0.067365072 | 0.946291074 | 0.998757862 | 9.987578619 |
| Meg3 | 1.4406489 | 0.01973452 | 0.634951323 | 0.2360471 | 0.813396122 | 0.998757862 | 9.987578619 |
| Phkg1 | 2.029296 | 0.002728104 | 0.540397666 | 0.0180485 | 0.985600162 | 0.998757862 | 9.987578619 |
| Ms4a1 | 2.864072 | -0.010217726 | 0.497605747 | -0.060338562 | 0.95188599 | 0.998757862 | 9.987578619 |
| Pde1c | 3.7180388 | 0.068380513 | 0.505239734 | 0.371148834 | 0.71052668 | 0.998757862 | 9.987578619 |
| Pcgf6 | 2.046505 | 0.037347998 | 0.547028895 | 0.249890533 | 0.802672004 | 0.998757862 | 9.987578619 |
| Pcca | 2.115995 | -0.030342523 | 0.52909521 | -0.213458973 | 0.830968987 | 0.998757862 | 9.987578619 |
| Ndufa4 | 1.8538784 | -0.033460529 | 0.543378434 | -0.398987927 | 0.6899021 | 0.998757862 | 9.987578619 |
| Rpl26 | 4.832376 | 0.033701136 | 0.55408582 | 0.221456417 | 0.824737069 | 0.998757862 | 9.987578619 |
| Rps4x | 2.1002257 | -0.020742103 | 0.538394946 | -0.149401742 | 0.881236638 | 0.998757862 | 9.987578619 |
| Kcnd2 | 1.7278643 | 0.054044546 | 0.579813017 | 0.394771235 | 0.693011725 | 0.998757862 | 9.987578619 |
| Il27 | 4.365445 | 0.003855057 | 0.477082433 | 0.018510979 | 0.985231219 | 0.998757862 | 9.987578619 |
| Rps3 | 1.5196611 | 0.013762535 | 0.620448937 | 0.164573062 | 0.869280035 | 0.998757862 | 9.987578619 |
| Esf1 | 1.0956551 | -0.144269504 | 0.533818679 | 0.198594897 | 0.842579646 | 0.998757862 | 9.987578619 |
| Fam169a | 2.507719 | -0.01033059 | 0.510008808 | -0.064931332 | 0.948228674 | 0.998757862 | 9.987578619 |
| Rps27a | 1.1677741 | -0.038642271 | 0.541482215 | -0.394983496 | 0.692855068 | 0.998757862 | 9.987578619 |
| Rps27 | 1.768813 | 0.018222294 | 0.586208134 | 0.158536205 | 0.874034294 | 0.998757862 | 9.987578619 |
| Rps25 | 0.21414971 | -0.027153724 | 0.482437982 | -0.240863286 | 0.809661078 | 0.998757862 | 9.987578619 |
| Rps23 | 0.9090493 | 0.00159684 | 0.607329615 | 0.026650241 | 0.978738701 | 0.998757862 | 9.987578619 |
| Galnt13 | 2.4272995 | -0.009310096 | 0.553344592 | -0.075881291 | 0.939513542 | 0.998757862 | 9.987578619 |
| Gas5 | 0.50752956 | -0.014945817 | 0.55599714 | -0.180326825 | 0.856895998 | 0.998757862 | 9.987578619 |
| Rps16 | 1.7898598 | -0.009794296 | 0.568635956 | -0.100987375 | 0.919560482 | 0.998757862 | 9.987578619 |
| Gm29325 | 2.2622397 | 0.007032309 | 0.515374709 | 0.042150018 | 0.966379107 | 0.998757862 | 9.987578619 |
| Rpl28 | 0.8195307 | 0.00908821 | 0.650610726 | 0.144471499 | 0.885128161 | 0.998757862 | 9.987578619 |
| Gm49767 | 1.7097859 | -0.022025932 | 0.532305452 | -0.158054418 | 0.874413919 | 0.998757862 | 9.987578619 |
| Rpl9 | 2.5286186 | 0.023366574 | 0.565695649 | 0.172885161 | 0.862741696 | 0.998757862 | 9.987578619 |
| Gnao1 | 0.94241995 | 0.032616584 | 0.601103896 | 0.314024652 | 0.753502316 | 0.998757862 | 9.987578619 |
| Gnaz | 6.549734 | 0.008924375 | 0.440300448 | 0.042054722 | 0.966455075 | 0.998757862 | 9.987578619 |
| Rpl37a | 3.1772478 | 0.04412804 | 0.607808117 | 0.370241591 | 0.71120249 | 0.998757862 | 9.987578619 |
| Rpl36a | 0.4488982 | -0.017659929 | 0.567259025 | -0.140704736 | 0.888103204 | 0.998757862 | 9.987578619 |

| | | | | | | | |
|---|---|---|---|---|---|---|---|
| Hapln1 | 1.8792313 | 0.046668511 | 0.548140332 | 0.324478801 | 0.745575569 | 0.998757862 | 9.987578619 |
| Rpl35a | 4.3182583 | -0.038235235 | 0.54737724 | -0.344327215 | 0.73060022 | 0.998757862 | 9.987578619 |
| Rpl34 | 1.5077101 | 0.014516173 | 0.588973837 | 0.1356586 | 0.892091177 | 0.998757862 | 9.987578619 |
| Il1rapl2 | 2.03188 | -0.034925477 | 0.520679282 | -0.227712884 | 0.819869456 | 0.998757862 | 9.987578619 |
| Gm960 | 0.5916446 | -0.011367657 | 0.55841263 | -0.217895954 | 0.82751018 | 0.998757862 | 9.987578619 |
| mt-Nd5 | 3.9185421 | 0.030394005 | 0.524421051 | 0.18286988 | 0.854900116 | 0.998757862 | 9.987578619 |
| Klkb1 | 0.80636305 | -0.000762263 | 0.573212028 | -0.007214447 | 0.994243754 | 0.999836329 | 9.998363292 |
| Rpl32 | 2.7427218 | 0.000589261 | 0.52293417 | 0.002399996 | 0.998085082 | 0.999836329 | 9.998363292 |
| Rps8 | 3.3647108 | 0.000328127 | 0.540777666 | 0.000205131 | 0.999836329 | 0.999836329 | 9.998363292 |

| **Glut** | baseMean | log2FoldChange | lfcSE | stat | pvalue | padj | pglobadj |
|---|---|---|---|---|---|---|---|
| Cst3 | 153.12349 | -0.144269504 | 0.513828888 | -15.71100388 | 1.27E-55 | 3.04E-53 | 3.04E-52 |
| mt-Atp6 | 236.50526 | 1.083960576 | 0.102048985 | 10.77975942 | 4.29E-27 | 5.13E-25 | 5.13E-24 |
| mt-Co3 | 295.9007 | 0.841302439 | 0.104311121 | 8.243868063 | 1.67E-16 | 1.33E-14 | 1.33E-13 |
| mt-Co2 | 163.4085 | -0.144269504 | 0.513829101 | 7.824931675 | 5.08E-15 | 3.03E-13 | 3.03E-12 |
| Camk1d | 71.690636 | 2.189615478 | 0.318930664 | 7.182032171 | 6.87E-13 | 3.28E-11 | 3.28E-10 |
| AY036118 | 96.97641 | 2.093149069 | 0.338010711 | 6.545542982 | 5.93E-11 | 2.36E-09 | 2.36E-08 |
| Malat1 | 56.60613 | 1.109004601 | 0.182690843 | 6.342155646 | 2.27E-10 | 7.74E-09 | 7.74E-08 |
| mt-Co1 | 159.95605 | 0.868853717 | 0.144855557 | 6.243767972 | 4.27E-10 | 1.28E-08 | 1.28E-07 |
| mt-Nd4 | 88.771065 | 1.09349557 | 0.185414493 | 6.19256802 | 5.92E-10 | 1.57E-08 | 1.57E-07 |
| Rpl13 | 53.985012 | -0.144269504 | 0.513829101 | -6.16450814 | 7.07E-10 | 1.69E-08 | 1.69E-07 |
| Cdk8 | 14.423764 | -0.144269504 | 0.513829183 | 5.864631161 | 4.50E-09 | 9.78E-08 | 9.78E-07 |
| mt-Cytb | 96.45872 | 1.094604526 | 0.212207248 | 5.495362676 | 3.90E-08 | 7.77E-07 | 7.77E-06 |
| Rps12 | 40.518112 | -1.756189803 | 0.448036825 | -4.457245744 | 8.30E-06 | 0.000152628 | 0.00152628 |
| Gm42418 | 454.46402 | 1.1304279 | 0.290734921 | 4.336999901 | 1.44E-05 | 0.000246581 | 0.002465809 |
| Gphn | 15.382317 | 1.497329951 | 0.437287805 | 3.965105634 | 7.34E-05 | 0.001095867 | 0.010958665 |
| Cmss1 | 51.62958 | 1.278037528 | 0.372224082 | 3.972406746 | 7.12E-05 | 0.001095867 | 0.010958665 |
| Meg3 | 20.876986 | 1.810646903 | 0.580118042 | 3.770218553 | 0.000163105 | 0.002293059 | 0.022930595 |
| Rps3 | 20.277365 | -1.165232392 | 0.391116801 | -3.547535495 | 0.000388853 | 0.005163108 | 0.051631084 |
| mt-Nd5 | 23.097977 | 1.106771087 | 0.372990863 | 3.529226136 | 0.000416777 | 0.005242614 | 0.052426138 |
| Rplp1 | 80.822174 | -0.72121287 | 0.231854693 | -3.507516862 | 0.00045231 | 0.005405101 | 0.054051008 |
| Rps8 | 46.15485 | -0.970778145 | 0.342163258 | -3.401708682 | 0.00066966 | 0.007621366 | 0.076213663 |
| Calm1 | 22.319403 | -0.144269504 | 0.513829155 | 3.285073346 | 0.001019558 | 0.01107611 | 0.110761103 |
| Rps20 | 31.994286 | -0.754223075 | 0.272407108 | -3.213472304 | 0.001311404 | 0.013627203 | 0.136272028 |
| Rpl35a | 41.280197 | -0.751965974 | 0.283129289 | -3.132686765 | 0.001732142 | 0.017249246 | 0.172492462 |
| Rps10 | 29.202696 | -0.833790392 | 0.340454514 | -3.012695868 | 0.002589383 | 0.024754499 | 0.247544993 |
| Apoe | 79.24645 | -0.589804273 | 0.229396743 | -2.951415223 | 0.003163214 | 0.027000288 | 0.270002882 |
| Rpl37 | 63.020767 | -0.144269504 | 0.513829108 | -2.973614753 | 0.002943143 | 0.027000288 | 0.270002882 |
| Rpl17 | 27.19542 | -1.095693047 | 0.502607484 | -2.9536389 | 0.003140512 | 0.027000288 | 0.270002882 |
| mt-Nd2 | 93.58352 | 0.658331511 | 0.266385474 | 2.92125464 | 0.003486248 | 0.028731489 | 0.287314886 |
| mt-Nd3 | 49.44484 | 0.609214025 | 0.252448177 | 2.841943575 | 0.004483943 | 0.034569752 | 0.345697516 |
| Mbp | 67.935005 | 0.896700007 | 0.415558247 | 2.846157635 | 0.004425028 | 0.034569752 | 0.345697516 |
| mt-Nd1 | 59.90675 | 0.755000758 | 0.333839698 | 2.830330449 | 0.004649995 | 0.03472965 | 0.347296501 |
| Rps18 | 21.838402 | -0.752536428 | 0.332018086 | -2.805672933 | 0.005021163 | 0.03619951 | 0.361995095 |
| Dlg2 | 13.382599 | 1.268689875 | 0.691720232 | 2.79751604 | 0.005149721 | 0.03619951 | 0.361995095 |
| Ppp3ca | 11.008586 | 1.007914844 | 0.505548687 | 2.77360794 | 0.005543845 | 0.037856539 | 0.378565386 |
| mt-Nd4l | 10.703118 | -0.144269504 | 0.513829178 | 2.712552352 | 0.006676723 | 0.043428609 | 0.434286095 |
| Syt1 | 10.909065 | 1.110941259 | 0.613348908 | 2.710249683 | 0.006723258 | 0.043428609 | 0.434286095 |
| Ralyl | 11.710411 | 1.075598642 | 0.607818615 | 2.68120096 | 0.007335845 | 0.044955561 | 0.449555613 |
| Cox6c | 37.26712 | -0.79024982 | 0.392475486 | -2.683623335 | 0.007282911 | 0.044955561 | 0.449555613 |
| Nkain2 | 14.873723 | -0.144269504 | 0.513829175 | 2.643088757 | 0.00821535 | 0.049086714 | 0.490867141 |
| Tmsb4x | 41.383972 | -0.69223787 | 0.366212234 | -2.513021928 | 0.011970189 | 0.069777446 | 0.697774457 |
| Rps19 | 30.389437 | -0.480175577 | 0.221223984 | -2.48552595 | 0.012936016 | 0.073612088 | 0.736120884 |
| Rpl3 | 13.666936 | -0.144269504 | 0.513829171 | 2.450377944 | 0.014270634 | 0.079318173 | 0.793181728 |
| Ntm | 40.96863 | 0.560863694 | 0.295882394 | 2.38283587 | 0.017179852 | 0.093317833 | 0.933178328 |
| Rps5 | 20.95526 | -0.778555445 | 0.514327335 | -2.366742589 | 0.017945409 | 0.095310063 | 0.953100625 |
| Cfap74 | 12.265499 | -0.713817722 | 0.433845889 | -2.34126787 | 0.019218373 | 0.09772747 | 0.977274703 |
| Cox4i1 | 62.974926 | -0.56736754 | 0.310030237 | -2.348787216 | 0.018834667 | 0.09772747 | 0.977274703 |
| Trim2 | 29.10721 | 0.530852317 | 0.284548746 | 2.325624589 | 0.020038587 | 0.099775463 | 0.997754627 |
| Rps27 | 22.94887 | -0.144269504 | 0.513829149 | 2.313353001 | 0.020703239 | 0.100981104 | 1.009811039 |
| Gm50306 | 22.39982 | -0.144269504 | 0.513829151 | 2.287508756 | 0.022166145 | 0.105954173 | 1.059541727 |

| | | | | | | | |
|---|---|---|---|---|---|---|---|
| Lrrc4c | 16.326674 | 0.778351608 | 0.609648275 | 2.279644948 | 0.022628755 | 0.106044557 | 1.060445571 |
| Galnt13 | 19.314667 | 0.610967044 | 0.371083253 | 2.269593442 | 0.023232262 | 0.10677905 | 1.067790499 |
| Mt3 | 37.2819 | -0.554705797 | 0.32518146 | -2.235882073 | 0.025359497 | 0.112239255 | 1.122392547 |
| Rpsa | 28.174269 | -0.745464497 | 0.626966134 | -2.240007184 | 0.025090457 | 0.112239255 | 1.122392547 |
| Rps24 | 56.464184 | -0.646034352 | 0.4589946 | -2.193258172 | 0.028288783 | 0.122927623 | 1.229276225 |
| Gm38973 | 10.315429 | -0.679971776 | 0.53000731 | -2.143341964 | 0.03208565 | 0.136936969 | 1.369369687 |
| Rpl26 | 36.338345 | -0.581159768 | 0.396307599 | -2.131031019 | 0.033086587 | 0.138731478 | 1.387314781 |
| Rps3a1 | 26.112642 | -0.595654234 | 0.437202 | -2.086740039 | 0.03691164 | 0.149523423 | 1.495234227 |
| Pcca | 7.60866 | -0.639913503 | 0.465392596 | -2.090986755 | 0.036529251 | 0.149523423 | 1.495234227 |
| Dbi | 18.70937 | -0.45396692 | 0.278493691 | -2.04304536 | 0.041047948 | 0.16350766 | 1.635076598 |
| Rps2 | 17.396173 | -0.59312499 | 0.510072269 | -2.010172672 | 0.044412917 | 0.174011264 | 1.740112639 |
| Sec61a1 | 131.5049 | -0.25483314 | 0.137434168 | -1.989869413 | 0.046605323 | 0.179656002 | 1.796560016 |
| Ppia | 17.1309 | 0.436783787 | 0.29107504 | 1.941802784 | 0.05216098 | 0.19102075 | 1.910207505 |
| Rpl28 | 21.463943 | -0.555464506 | 0.487914028 | -1.948808556 | 0.051318292 | 0.19102075 | 1.910207505 |
| 1700054A( | 96.21874 | -0.35037733 | 0.209907775 | -1.943947133 | 0.051901827 | 0.19102075 | 1.910207505 |
| Fam169a | 17.663467 | -0.144269504 | 0.513829145 | -1.936957816 | 0.0527505 | 0.19102075 | 1.910207505 |
| Cpe | 17.7603 | 0.421823923 | 0.283513882 | 1.906812197 | 0.056544911 | 0.201704982 | 2.017049823 |
| Uba52 | 9.519464 | 0.524257529 | 0.480774805 | 1.885862504 | 0.059313485 | 0.208469456 | 2.084694562 |
| Fau | 41.727573 | -0.502022297 | 0.59367275 | -1.853544796 | 0.063804312 | 0.22100334 | 2.210033403 |
| Cadm2 | 10.564265 | 0.504419183 | 0.486540345 | 1.836654302 | 0.066260946 | 0.2262338 | 2.262337997 |
| Rpl12 | 12.941884 | -0.496717529 | 0.43042118 | -1.830050323 | 0.067242414 | 0.226351226 | 2.263512256 |
| Rpl30 | 35.64468 | -0.399853654 | 0.300437378 | -1.758997496 | 0.078577934 | 0.257262003 | 2.572620028 |
| Rps14 | 40.009132 | -0.444047645 | 0.377502297 | -1.760873364 | 0.078259839 | 0.257262003 | 2.572620028 |
| Ccdc60 | 11.066734 | 0.446412918 | 0.465557499 | 1.699812808 | 0.089166142 | 0.287982539 | 2.879825386 |
| Rpl38 | 36.959415 | -0.390348561 | 0.33794118 | -1.642493621 | 0.100487747 | 0.320220954 | 3.202209537 |
| Rpl27a | 35.931152 | -0.395987991 | 0.35829275 | -1.628422586 | 0.103435312 | 0.323810437 | 3.238104374 |
| Rps7 | 22.277212 | -0.413256489 | 0.506186701 | -1.624243432 | 0.104323865 | 0.323810437 | 3.238104374 |
| Cntnap2 | 39.725906 | 0.310773792 | 0.242120459 | 1.580315422 | 0.11403465 | 0.328364835 | 3.28364835 |
| Phkg1 | 9.036313 | -0.144269504 | 0.513829156 | -1.585924642 | 0.112756405 | 0.328364835 | 3.28364835 |
| Rps16 | 25.17542 | -0.401925109 | 0.458483623 | -1.583929905 | 0.113209671 | 0.328364835 | 3.28364835 |
| AI314278 | 7.108044 | -0.417122199 | 0.472002587 | -1.589024612 | 0.112054836 | 0.328364835 | 3.28364835 |
| Rpl24 | 24.18354 | -0.408993961 | 0.435212599 | -1.603270488 | 0.108874949 | 0.328364835 | 3.28364835 |
| Gas5 | 23.488619 | 0.364968546 | 0.631666936 | 1.607906833 | 0.107855585 | 0.328364835 | 3.28364835 |
| Anks1b | 8.142463 | 0.378856144 | 0.468249837 | 1.521001813 | 0.128259382 | 0.364928481 | 3.649284809 |
| Rpl23 | 28.991348 | -0.364482982 | 0.395840913 | -1.489687293 | 0.136306477 | 0.383261742 | 3.832617421 |
| Gnaz | 27.21197 | -0.31320742 | 0.274617374 | -1.471059981 | 0.141274896 | 0.383993293 | 3.839932925 |
| Atxn7l1 | 8.474053 | -0.144269504 | 0.513829173 | 1.476809636 | 0.139726673 | 0.383993293 | 3.839932925 |
| Lsamp | 19.663153 | 0.342285433 | 0.338993046 | 1.470646833 | 0.141386652 | 0.383993293 | 3.839932925 |
| Cox5b | 15.267654 | -0.351027409 | 0.469166874 | -1.445259586 | 0.148384985 | 0.398472039 | 3.984720386 |
| Lrrc17 | 17.73422 | -0.349726544 | 0.408941335 | -1.430616447 | 0.152540171 | 0.401019935 | 4.010199349 |
| Rps28 | 24.591118 | -0.347447382 | 0.399527343 | -1.430095545 | 0.152689599 | 0.401019935 | 4.010199349 |
| Rps23 | 26.174232 | -0.270438714 | 0.232419942 | -1.400734875 | 0.16129337 | 0.415345341 | 4.153453412 |
| Gpm6b | 10.295604 | 0.334480223 | 0.482716262 | 1.39964472 | 0.161619735 | 0.415345341 | 4.153453412 |
| Rpl41 | 39.928654 | -0.305241015 | 0.296288235 | -1.39229108 | 0.163834272 | 0.41655735 | 4.1655735 |
| Rps21 | 74.39566 | -0.326042207 | 0.369935682 | -1.379667232 | 0.167689127 | 0.42187054 | 4.218705396 |
| Uqcr11 | 11.809479 | -0.329936383 | 0.402227315 | -1.360648068 | 0.173624933 | 0.43225374 | 4.322537404 |
| Ptpn9 | 10.586755 | -0.321730271 | 0.389237887 | -1.3327549 | 0.18261225 | 0.448948345 | 4.489483451 |
| Gm49767 | 8.390749 | -0.317800522 | 0.447708614 | -1.328273999 | 0.184087606 | 0.448948345 | 4.489483451 |
| Hexb | 14.844687 | 0.306901247 | 0.459671593 | 1.302123522 | 0.192874163 | 0.465625505 | 4.656255054 |
| Gapdh | 15.659986 | -0.304250717 | 0.375149119 | -1.288556608 | 0.197552277 | 0.472149942 | 4.721499418 |
| Tmem108 | 12.377369 | 0.298628362 | 0.367464027 | 1.279720885 | 0.200643317 | 0.474789631 | 4.747896306 |
| Rpl9 | 28.012674 | -0.277847262 | 0.491265436 | -1.265033967 | 0.205859152 | 0.482356248 | 4.823562478 |
| Gm12216 | 59.987026 | -0.174730635 | 0.159925984 | -1.200641834 | 0.229890166 | 0.49921839 | 4.992183898 |
| Tacc1 | 8.593654 | -0.144269504 | 0.513829166 | -1.235079858 | 0.216800785 | 0.49921839 | 4.992183898 |
| Sil1 | 16.818111 | 0.263883996 | 0.310623163 | 1.197151299 | 0.231247588 | 0.49921839 | 4.992183898 |
| Mt1 | 35.104218 | -0.242858962 | 0.260519433 | -1.194762997 | 0.232179641 | 0.49921839 | 4.992183898 |
| Rpl37a | 43.691177 | -0.26717274 | 0.30374115 | -1.221178878 | 0.2220183 | 0.49921839 | 4.992183898 |
| Hdac9 | 6.796373 | 0.285382515 | 0.438466961 | 1.229244596 | 0.218980113 | 0.49921839 | 4.992183898 |
| Pcsk1n | 18.374506 | -0.280206498 | 0.407606172 | -1.204359297 | 0.228450737 | 0.49921839 | 4.992183898 |
| Rps27a | 36.51063 | -0.27980249 | 0.354920251 | -1.21973236 | 0.22256635 | 0.49921839 | 4.992183898 |
| Cmpk1 | 31.528717 | 0.256692777 | 0.280307742 | 1.213872122 | 0.22479656 | 0.49921839 | 4.992183898 |
| Ctnna2 | 8.165573 | -0.144269504 | 0.51382918 | 1.190262218 | 0.233943346 | 0.49921839 | 4.992183898 |

| | | | | | | | |
|---|---|---|---|---|---|---|---|
| Pde1c | 28.128672 | -0.261567482 | 0.321334531 | -1.174797 | 0.240075941 | 0.507369192 | 5.073691922 |
| Car12 | 15.380535 | 0.265573986 | 0.348740398 | 1.169201356 | 0.242322515 | 0.507369192 | 5.073691922 |
| Adk | 10.355698 | 0.26893172 | 0.397705798 | 1.164721805 | 0.24413162 | 0.507369192 | 5.073691922 |
| Il1rapl2 | 7.5780206 | 0.263827093 | 0.424224467 | 1.150178749 | 0.250070257 | 0.510827277 | 5.108272772 |
| Camk2a | 15.41069 | 0.257593039 | 0.328228838 | 1.152274089 | 0.249208461 | 0.510827277 | 5.108272772 |
| Wdyhv1 | 8.780831 | -0.263201728 | 0.39714818 | -1.132031701 | 0.257621107 | 0.517407097 | 5.174070968 |
| 4930402H2 | 8.12458 | 0.259705291 | 0.407811307 | 1.132561741 | 0.257398343 | 0.517407097 | 5.174070968 |
| Plp1 | 12.918778 | -0.253913348 | 0.419023889 | -1.118604016 | 0.263309109 | 0.520089894 | 5.200898938 |
| Kcnq5 | 9.718189 | 0.254480333 | 0.437468059 | 1.120455781 | 0.262519586 | 0.520089894 | 5.200898938 |
| Atp9b | 13.626531 | 0.252543046 | 0.418311978 | 1.105465972 | 0.268957731 | 0.523458074 | 5.234580736 |
| Acoxl | 12.417441 | -0.144269504 | 0.513829168 | 1.104457475 | 0.269394741 | 0.523458074 | 5.234580736 |
| Zmat4 | 14.470482 | -0.144269504 | 0.513829161 | -1.099313162 | 0.271631493 | 0.523547798 | 5.235477975 |
| Rpl32 | 27.14873 | -0.234367634 | 0.293131294 | -1.09367179 | 0.274098951 | 0.524077194 | 5.240771939 |
| Ctnna3 | 17.311068 | -0.242818856 | 0.330282155 | -1.085520043 | 0.277691391 | 0.526732084 | 5.267320837 |
| Cox8a | 61.68057 | -0.243583588 | 0.380120348 | -1.075978978 | 0.281936658 | 0.530306458 | 5.303064578 |
| Aldoc | 16.639307 | -0.243493992 | 0.359779478 | -1.071346853 | 0.2840135 | 0.530306458 | 5.303064578 |
| Gm10062 | 35.610672 | 0.218205789 | 0.268138728 | 1.063424828 | 0.287589338 | 0.532820557 | 5.328205573 |
| Selenow | 18.020937 | -0.235758001 | 0.422019451 | -1.054108248 | 0.29183336 | 0.53361849 | 5.336184897 |
| Gria1 | 9.3569975 | -0.144269504 | 0.513829171 | 1.052684863 | 0.292485448 | 0.53361849 | 5.336184897 |
| Epha6 | 24.288336 | 0.200994768 | 0.237495623 | 1.044461772 | 0.296271797 | 0.536431511 | 5.36431511 |
| Cep128 | 7.9962883 | 0.233699484 | 0.360888064 | 1.009458248 | 0.312754916 | 0.562018232 | 5.620182319 |
| Gm29325 | 10.006942 | -0.219285979 | 0.370064577 | -0.973661652 | 0.330224564 | 0.580321108 | 5.803211082 |
| Mrc2 | 10.112897 | -0.214804633 | 0.421478545 | -0.974296653 | 0.329909266 | 0.580321108 | 5.803211082 |
| Vit | 8.487055 | -0.222277237 | 0.385868803 | -0.982529527 | 0.325839044 | 0.580321108 | 5.803211082 |
| Rpl21 | 40.990406 | -0.200025888 | 0.27974184 | -0.960735073 | 0.336685391 | 0.583100062 | 5.831000618 |
| D430042O | 42.442684 | 0.156714647 | 0.185757238 | 0.962135172 | 0.335981708 | 0.583100062 | 5.831000618 |
| Spock1 | 7.8019853 | -0.144269504 | 0.513829168 | -0.943325498 | 0.345514438 | 0.594085976 | 5.940859756 |
| Gnao1 | 8.048166 | 0.200059128 | 0.431575772 | 0.924903343 | 0.355016187 | 0.601765027 | 6.017650268 |
| Adamtsl1 | 20.072302 | -0.144269504 | 0.51382915 | -0.928373982 | 0.35321361 | 0.601765027 | 6.017650268 |
| Ckb | 25.570559 | 0.202154155 | 0.340588854 | 0.91385722 | 0.360791875 | 0.607248297 | 6.072482969 |
| Rps29 | 90.593666 | 0.198465655 | 0.368475042 | 0.891858922 | 0.372468555 | 0.614111769 | 6.14111769 |
| Atp5j2 | 13.27106 | -0.195761702 | 0.358466964 | -0.882110082 | 0.377717281 | 0.614111769 | 6.14111769 |
| Dpyd | 9.891637 | -0.144269504 | 0.51382917 | 0.882134526 | 0.377704063 | 0.614111769 | 6.14111769 |
| Cox6b1 | 16.960936 | -0.180971551 | 0.450830926 | -0.882648341 | 0.377426301 | 0.614111769 | 6.14111769 |
| Ehbp1 | 24.006865 | -0.192896375 | 0.301094686 | -0.900475749 | 0.367867125 | 0.614111769 | 6.14111769 |
| Rps17 | 12.333552 | 0.192499099 | 0.378204788 | 0.867764138 | 0.385523466 | 0.6225683 | 6.225683002 |
| Zbtb7a | 14.866871 | 0.184387187 | 0.353527802 | 0.835474187 | 0.403450775 | 0.634305974 | 6.343059739 |
| Rpl27 | 12.368381 | 0.187200218 | 0.364767886 | 0.848829629 | 0.395976101 | 0.634305974 | 6.343059739 |
| Bc1 | 14.420689 | -0.170414115 | 0.442045368 | -0.830843609 | 0.406061983 | 0.634305974 | 6.343059739 |
| Gria4 | 17.834356 | 0.181222043 | 0.313787145 | 0.84006311 | 0.400873002 | 0.634305974 | 6.343059739 |
| Ndufa4 | 17.974972 | -0.181013038 | 0.314777812 | -0.835792678 | 0.403271546 | 0.634305974 | 6.343059739 |
| Rab29 | 12.269017 | -0.175795068 | 0.373696532 | -0.802151563 | 0.422465291 | 0.655644186 | 6.556441856 |
| Rsrc1 | 25.437769 | -0.144269504 | 0.513829154 | 0.765741567 | 0.443830089 | 0.684357363 | 6.843573625 |
| Alpl | 7.422941 | 0.162727664 | 0.402391449 | 0.756837751 | 0.449147073 | 0.688116349 | 6.881163488 |
| Zfp994 | 10.542933 | -0.164143831 | 0.366496834 | -0.750575388 | 0.452908238 | 0.689459037 | 6.894590375 |
| Mc1r | 22.108246 | -0.144269504 | 0.51382916 | 0.74218859 | 0.457973081 | 0.692756749 | 6.927567487 |
| Sult2a2 | 7.131176 | -0.144269504 | 0.513829168 | 0.734818761 | 0.46244988 | 0.695129064 | 6.951290643 |
| Ppp1r16a | 9.812871 | -0.144269504 | 0.513829154 | -0.72986647 | 0.46547181 | 0.695298516 | 6.952985155 |
| Ctcflos | 9.990439 | -0.147876223 | 0.429622923 | -0.723350189 | 0.469464766 | 0.696907323 | 6.969073229 |
| Rps4x | 11.603481 | 0.133676571 | 0.494272801 | 0.713444855 | 0.475570525 | 0.701613306 | 7.016133058 |
| 8430422H0 | 24.16215 | -0.144269504 | 0.513829156 | -0.69936057 | 0.484326723 | 0.706222669 | 7.062226694 |
| Rpl34 | 22.231255 | -0.146088719 | 0.41236143 | -0.698915761 | 0.484604677 | 0.706222669 | 7.062226694 |
| Kcnip4 | 12.674187 | 0.143132832 | 0.386005838 | 0.666849828 | 0.504868061 | 0.731293737 | 7.31293737 |
| Rpl36 | 21.884354 | -0.131471451 | 0.384586562 | -0.619505563 | 0.535583359 | 0.758493433 | 7.584934334 |
| Ptprg | 7.291076 | 0.133210602 | 0.397699658 | 0.630065957 | 0.528651432 | 0.758493433 | 7.584934334 |
| Ppp1r9a | 11.041557 | -0.132823179 | 0.324370263 | -0.618356228 | 0.536340545 | 0.758493433 | 7.584934334 |
| Gm765 | 9.099705 | 0.132991764 | 0.374477313 | 0.619171856 | 0.53580315 | 0.758493433 | 7.584934334 |
| Nell1 | 15.036522 | -0.144269504 | 0.513829172 | 0.608383051 | 0.54293345 | 0.763300556 | 7.633005565 |
| Atp5k | 13.95847 | 0.121460373 | 0.303741719 | 0.575693232 | 0.564822551 | 0.789430348 | 7.894303484 |
| Cspp1 | 10.810677 | -0.119246856 | 0.322453263 | -0.557794197 | 0.576984926 | 0.793125407 | 7.931254074 |
| 6030443J0 | 21.46823 | -0.144269504 | 0.513829143 | 0.557154543 | 0.577421845 | 0.793125407 | 7.931254074 |
| Ankrd12 | 14.6444235 | -0.116168964 | 0.286273269 | -0.558624977 | 0.57641769 | 0.793125407 | 7.931254074 |

| | | | | | | | |
|---|---|---|---|---|---|---|---|
| Csmd3 | 18.023438 | -0.118280775 | 0.330618673 | -0.551680319 | 0.581167397 | 0.793708616 | 7.93708616 |
| Scamp1 | 10.45517 | 0.113200063 | 0.363583258 | 0.528430394 | 0.597200647 | 0.806389574 | 8.063895741 |
| Ttc28 | 18.252264 | -0.112749353 | 0.309899158 | -0.53139226 | 0.595146984 | 0.806389574 | 8.063895741 |
| Cdh23 | 21.341753 | -0.105193342 | 0.302341635 | -0.500948693 | 0.616407232 | 0.818451825 | 8.184518253 |
| Gm49497 | 8.222299 | 0.106711317 | 0.390290947 | 0.508271258 | 0.611263124 | 0.818451825 | 8.184518253 |
| Fth1 | 168.7455 | -0.104137496 | 0.294664088 | -0.503708966 | 0.614465907 | 0.818451825 | 8.184518253 |
| Igf2bp3 | 29.640501 | 0.102531254 | 0.371060929 | 0.478248464 | 0.632473372 | 0.825452844 | 8.254528437 |
| Sema3c | 40.454563 | 0.099395894 | 0.29366783 | 0.476556822 | 0.633677736 | 0.825452844 | 8.254528437 |
| Nckap5 | 57.710537 | 0.074663936 | 0.180753998 | 0.475211669 | 0.63463611 | 0.825452844 | 8.254528437 |
| Ptprd | 9.4254675 | 0.100782022 | 0.36858761 | 0.474006775 | 0.635495076 | 0.825452844 | 8.254528437 |
| Rpl23a | 12.074002 | 0.094666999 | 0.39569632 | 0.453653922 | 0.650077937 | 0.835315198 | 8.353151982 |
| Ninj2 | 12.011555 | -0.097932118 | 0.342750178 | -0.458332944 | 0.646713259 | 0.835315198 | 8.353151982 |
| Dop1b | 13.804519 | 0.093914219 | 0.361065076 | 0.438919786 | 0.66071966 | 0.844449191 | 8.444491907 |
| Rpl35 | 11.906886 | 0.090770159 | 0.375419282 | 0.429414305 | 0.667621746 | 0.8487319 | 8.487319001 |
| Hapln1 | 7.49247 | 0.089342152 | 0.37533299 | 0.424178284 | 0.671435797 | 0.849064315 | 8.490643146 |
| Tpt1 | 30.659975 | -0.085003198 | 0.308206079 | -0.406491644 | 0.684381395 | 0.851912257 | 8.519122574 |
| Eef1a1 | 22.805122 | 0.086706484 | 0.313260252 | 0.411045049 | 0.681039503 | 0.851912257 | 8.519122574 |
| Chsy3 | 15.136107 | -0.086899834 | 0.323614805 | -0.410441047 | 0.681482441 | 0.851912257 | 8.519122574 |
| 1700126G( | 23.795546 | -0.082769561 | 0.315771093 | -0.392907894 | 0.694387511 | 0.859889197 | 8.598891971 |
| Uqcrh | 20.020086 | -0.079867032 | 0.313258558 | -0.379871565 | 0.704040756 | 0.863501558 | 8.635015577 |
| Gm14582 | 7.9209514 | 0.07757983 | 0.404476463 | 0.37921184 | 0.70453056 | 0.863501558 | 8.635015577 |
| Rpl13a | 25.250313 | -0.078307556 | 0.357005263 | -0.370510565 | 0.711002106 | 0.866987262 | 8.669872621 |
| Gpx4 | 11.707394 | -0.065237053 | 0.457862469 | -0.363752097 | 0.716043137 | 0.86870208 | 8.687020797 |
| Rpl39 | 32.30936 | -0.075079165 | 0.331967762 | -0.356508504 | 0.721459785 | 0.870852972 | 8.708529723 |
| Klkb1 | 7.68376 | -0.071238282 | 0.375498262 | -0.342278952 | 0.732140974 | 0.872237079 | 8.72237079 |
| Xlr5a | 56.852993 | 0.065329479 | 0.23982968 | 0.345564443 | 0.729670072 | 0.872237079 | 8.72237079 |
| Prkg1 | 20.728361 | -0.144269504 | 0.513829168 | 0.336584722 | 0.736429975 | 0.872237079 | 8.72237079 |
| Slx1b | 16.138987 | 0.069244786 | 0.277313052 | 0.335557523 | 0.737204561 | 0.872237079 | 8.72237079 |
| 5830416I1 | 68.23154 | -0.144269504 | 0.513829111 | 0.320943152 | 0.748253472 | 0.880948669 | 8.809486688 |
| Rps15 | 20.13585 | -0.063520598 | 0.330795111 | -0.301615785 | 0.762944973 | 0.893842395 | 8.938423953 |
| Rplp2 | 15.350688 | 0.06116188 | 0.354518156 | 0.28704913 | 0.774074702 | 0.8950189 | 8.950189001 |
| Pde6a | 41.91164 | 0.051906457 | 0.208719255 | 0.294795503 | 0.768150102 | 0.8950189 | 8.950189001 |
| 4930558K( | 11.438983 | -0.144269504 | 0.513829165 | 0.284496835 | 0.776029659 | 0.8950189 | 8.950189001 |
| Cntn3 | 8.736117 | -0.058658986 | 0.3583983 | -0.280715548 | 0.778928582 | 0.8950189 | 8.950189001 |
| Chl1 | 9.660644 | -0.057375964 | 0.317617368 | -0.273034757 | 0.784826498 | 0.897481019 | 8.974810195 |
| Pcgf6 | 9.177818 | 0.055708367 | 0.334453281 | 0.262163592 | 0.793195321 | 0.902731818 | 9.027318182 |
| Prkag2 | 11.633308 | -0.144269504 | 0.513829172 | -0.232847811 | 0.815879585 | 0.90590211 | 9.059021103 |
| Svil | 6.522386 | -0.046772699 | 0.38678069 | -0.229187425 | 0.818723246 | 0.90590211 | 9.059021103 |
| Gm11337 | 7.686161 | 0.047982459 | 0.377501398 | 0.23000863 | 0.818085064 | 0.90590211 | 9.059021103 |
| Tmeff2 | 7.9769416 | 0.050028026 | 0.380691311 | 0.239914297 | 0.810396697 | 0.90590211 | 9.059021103 |
| Wdr72 | 13.736755 | 0.051800221 | 0.332333733 | 0.242309367 | 0.808540456 | 0.90590211 | 9.059021103 |
| Slc25a4 | 13.340878 | 0.046843059 | 0.416040172 | 0.231492553 | 0.81693217 | 0.90590211 | 9.059021103 |
| Ubb | 13.164841 | 0.046485788 | 0.35079306 | 0.218915795 | 0.826715642 | 0.907039092 | 9.070390916 |
| Sparcl1 | 15.654034 | -0.044646515 | 0.3673967 | -0.215241255 | 0.829579228 | 0.907039092 | 9.070390916 |
| Shb | 39.72005 | -0.04301523 | 0.278297688 | -0.213244528 | 0.831136239 | 0.907039092 | 9.070390916 |
| Pcdh15 | 19.016542 | -0.144269504 | 0.51382915 | 0.192638686 | 0.847241945 | 0.908807568 | 9.088075684 |
| Cox7c | 26.405092 | -0.144269504 | 0.513829156 | 0.200623906 | 0.840992664 | 0.908807568 | 9.088075684 |
| Lrp1b | 128.38896 | 0.033833102 | 0.212643238 | 0.189683103 | 0.849557462 | 0.908807568 | 9.088075684 |
| Ms4a1 | 16.984304 | -0.144269504 | 0.513829164 | 0.186861232 | 0.851769436 | 0.908807568 | 9.088075684 |
| Kat7 | 10.341788 | 0.04153951 | 0.378172453 | 0.198140897 | 0.842934828 | 0.908807568 | 9.088075684 |
| Il27 | 19.969423 | -0.036708447 | 0.279158964 | -0.181195512 | 0.856214117 | 0.909025431 | 9.090254307 |
| Gm960 | 6.655112 | 0.036296251 | 0.390130789 | 0.176908155 | 0.859580533 | 0.909025431 | 9.090254307 |
| Fbxl7 | 8.989198 | 0.034328455 | 0.352107306 | 0.161863951 | 0.871412992 | 0.917478877 | 9.174788765 |
| Ahdc1 | 8.223656 | -0.031258226 | 0.363216291 | -0.150543601 | 0.880335754 | 0.922808093 | 9.228080934 |
| Mapk14 | 26.49383 | -0.02563504 | 0.291961754 | -0.12741433 | 0.898612475 | 0.937853194 | 9.37853194 |
| Rpl11 | 37.099155 | 0.022555949 | 0.4458339 | 0.115118426 | 0.908351255 | 0.943710911 | 9.437109113 |
| Kctd16 | 50.94356 | -0.144269504 | 0.513829105 | -0.110362034 | 0.912122262 | 0.943710911 | 9.437109113 |
| Rapgef4 | 29.665274 | 0.018301571 | 0.275463601 | 0.089432211 | 0.928738426 | 0.956760706 | 9.567607058 |
| 4930441J1 | 8.310448 | -0.01494886 | 0.352282341 | -0.072090402 | 0.942529964 | 0.95985066 | 9.598506603 |
| Fyb2 | 15.566055 | 0.016512323 | 0.345059706 | 0.077248076 | 0.938426196 | 0.95985066 | 9.598506603 |
| Hdac8 | 7.574986 | -0.144269504 | 0.513829168 | 0.070511919 | 0.943786214 | 0.95985066 | 9.598506603 |
| Rpl36a | 10.0171585 | 0.011922019 | 0.344071929 | 0.055229252 | 0.955955825 | 0.959972446 | 9.599724458 |

| | | | | | | | |
|---|---|---|---|---|---|---|---|
| Zfhx3 | 9.152152 | 0.011945721 | 0.357699935 | 0.055666034 | 0.955607858 | 0.959972446 | 9.599724458 |
| Rps25 | 19.357801 | -0.012877695 | 0.307988228 | -0.063199905 | 0.949607321 | 0.959972446 | 9.599724458 |
| 4930511M | 23.912638 | -0.00521604 | 0.290127242 | -0.026374826 | 0.978958373 | 0.978958373 | 9.789583732 |
| Arvcf | 6.5191417 | -0.144269504 | 0.513829175 | 0.61217265 | 0.540423535 | | |
| Dctd | 4.7964993 | -0.001008548 | 0.418420614 | -0.00656554 | 0.994761495 | | |
| Esf1 | 6.320379 | 0.071650547 | 0.395952179 | 0.348326764 | 0.727594796 | | |
| Grik1 | 2.652864 | -0.524882052 | 0.712019127 | -1.811387795 | 0.070080844 | | |
| Kcnd2 | 6.3454533 | -0.144269504 | 0.513829182 | 2.150279594 | 0.031533106 | | |
| Kcnh7 | 5.200062 | 0.631298776 | 0.601876242 | 2.042341703 | 0.041117648 | | |
| Klhl14 | 1.3022913 | -0.144269504 | 0.513829188 | 1.690499601 | 0.090932414 | | |
| Lyrm1 | 5.873159 | -0.144269504 | 0.513829176 | 1.303737591 | 0.192323068 | | |
| Sgip1 | 5.4363647 | -0.144269504 | 0.513829183 | 1.408789099 | 0.158897539 | | |
| Srgn | 4.6724954 | 0.100520445 | 0.423935825 | 0.505391716 | 0.613283736 | | |
| St18 | 3.2631721 | -0.144269504 | 0.51382917 | -2.345749875 | 0.018988846 | | |
| Tns3 | 6.517016 | 0.051110122 | 0.390182421 | 0.248855103 | 0.803472866 | | |
| Tshz2 | 1.4463643 | -0.144269504 | 0.513829283 | 1.38150351 | 0.167124199 | | |
| Unc5d | 0.7641921 | -0.144269504 | 0.51382918 | 0.101456737 | 0.919187899 | | |
| Vwc2l | 5.621639 | -0.0344424 | 0.391883465 | -0.171026497 | 0.864202931 | | |

**table S2: Differential gene expression list quinine compared to condensed milk in GABA and Glut neurons**

| **GABA** | baseMean | log2FoldChange | lfcSE | stat | pvalue | padj | pglobadj |
|---|---|---|---|---|---|---|---|
| Rps12 | 12.221588 | -0.144269504 | 0.728707886 | -7.210270611 | 5.58E-13 | 1.57E-10 | 1.57E-09 |
| Cst3 | 35.174557 | -1.90118434 | 0.282247799 | -7.072732744 | 1.52E-12 | 2.14E-10 | 2.14E-09 |
| Grik1 | 363.9453 | -0.343570652 | 0.076864215 | -4.57668064 | 4.72E-06 | 0.000444067 | 0.004440671 |
| Pde1c | 16.1248 | -0.144269504 | 0.728707972 | 4.060422804 | 4.90E-05 | 0.002787127 | 0.027871271 |
| Lrp1b | 117.401024 | 0.579243125 | 0.151642036 | 4.058366609 | 4.94E-05 | 0.002787127 | 0.027871271 |
| Fau | 9.303338 | -0.144269504 | 0.728707895 | -3.766058705 | 0.000165845 | 0.007794706 | 0.077947062 |
| Rpl35a | 13.723231 | -2.631754122 | 0.962834588 | -3.588445307 | 0.000332656 | 0.013401275 | 0.134012748 |
| Rps21 | 14.005599 | -1.229051144 | 0.451151268 | -3.412039861 | 0.000644787 | 0.022728736 | 0.227287361 |
| Ntm | 28.322523 | -0.144269504 | 0.728708133 | 3.227959212 | 0.001246767 | 0.035158843 | 0.351588434 |
| Rps20 | 3.0477123 | -0.144269504 | 0.728707909 | -3.257159424 | 0.001125332 | 0.035158843 | 0.351588434 |
| Rps16 | 10.1992035 | -1.495594791 | 0.656292176 | -3.131474634 | 0.001739308 | 0.041208564 | 0.412085643 |
| Rps28 | 12.25369 | -1.171926149 | 0.498326465 | -3.115682558 | 0.001835197 | 0.041208564 | 0.412085643 |
| Prkg1 | 14.764182 | -0.144269504 | 0.728707963 | 3.105482971 | 0.001899686 | 0.041208564 | 0.412085643 |
| Rps27a | 9.805704 | -1.52568375 | 0.716191606 | -3.061797037 | 0.002200126 | 0.044316819 | 0.443168194 |
| Aldoc | 6.235399 | -1.558097109 | 0.760769287 | -3.012775202 | 0.002588706 | 0.047689518 | 0.476895177 |
| Cox8a | 15.239409 | -1.934143803 | 1.059439845 | -2.999324649 | 0.002705788 | 0.047689518 | 0.476895177 |
| Grm3 | 10.096467 | -1.072600203 | 0.490548293 | -2.972909951 | 0.00294991 | 0.048933798 | 0.489337982 |
| Rpl9 | 9.58222 | -1.619616937 | 1.169856698 | -2.749299011 | 0.005972288 | 0.084209257 | 0.84209257 |
| Nckap5 | 44.42943 | -0.144269504 | 0.728708061 | 2.760738756 | 0.005767079 | 0.084209257 | 0.84209257 |
| Rps3a1 | 7.1695037 | -1.630533546 | 1.09137246 | -2.781116588 | 0.005417228 | 0.084209257 | 0.84209257 |
| Uqcrh | 4.750526 | -1.732968705 | 1.468260253 | -2.692111671 | 0.007100117 | 0.095344425 | 0.953444245 |
| Cntnap2 | 24.043127 | -0.144269504 | 0.728707993 | 2.645089737 | 0.008166926 | 0.104685136 | 1.046851363 |
| Apoe | 15.710725 | -0.837977137 | 0.488659202 | -2.538093111 | 0.011145833 | 0.1366576 | 1.366575998 |
| Rps3 | 3.8628423 | -0.481051197 | 0.970589406 | -2.471067808 | 0.013471027 | 0.146601059 | 1.46601059 |
| Rpl8 | 8.226033 | -0.519537444 | 1.040516748 | -2.495990173 | 0.012560608 | 0.146601059 | 1.46601059 |
| Fam169a | 12.418809 | 0.784188324 | 0.413266465 | 2.469864876 | 0.01351641 | 0.146601059 | 1.46601059 |
| Il1rapl2 | 5.0466504 | -0.144269504 | 0.728707917 | -2.335747775 | 0.019504391 | 0.192210706 | 1.922107055 |
| Tnni3k | 5.5913777 | -0.832326404 | 0.817509932 | -2.331350915 | 0.019734864 | 0.192210706 | 1.922107055 |
| Ankrd12 | 13.006836 | -0.699417692 | 0.460036138 | -2.330753731 | 0.019766349 | 0.192210706 | 1.922107055 |
| Fth1 | 24.661276 | -0.75789965 | 0.927969922 | -2.302361229 | 0.02131481 | 0.200359209 | 2.003592094 |
| Ckb | 6.0336604 | -0.605859977 | 0.887692386 | -2.213477401 | 0.02686475 | 0.231915453 | 2.319154525 |
| Rpl41 | 7.1125774 | -0.71504672 | 0.777500241 | -2.211651378 | 0.026990763 | 0.231915453 | 2.319154525 |
| Tshz2 | 15.929192 | 0.647767893 | 0.420644737 | 2.209512092 | 0.027139042 | 0.231915453 | 2.319154525 |
| Car12 | 4.998469 | -0.640991123 | 0.848346698 | -2.192833791 | 0.028319356 | 0.234884074 | 2.348840736 |
| Hapln1 | 7.078533 | -0.144269504 | 0.728707953 | 2.16771335 | 0.030180503 | 0.234902205 | 2.349022052 |
| Rps15a | 4.942472 | -0.228314644 | 0.543442684 | -2.171813665 | 0.029869721 | 0.234902205 | 2.349022052 |
| Trim2 | 15.258869 | -0.144269504 | 0.728707956 | 2.15938253 | 0.030820502 | 0.234902205 | 2.349022052 |
| Rps29 | 16.899284 | -0.608037806 | 0.463662509 | -2.113014841 | 0.034599498 | 0.251432571 | 2.514325706 |
| Stxbp5l | 6.1767383 | -0.144269504 | 0.728707978 | 2.110996769 | 0.03477259 | 0.251432571 | 2.514325706 |
| Rps10 | 4.1539774 | -0.530694792 | 0.808802705 | -2.086996204 | 0.036888478 | 0.253641284 | 2.536412842 |
| Rps24 | 11.304478 | -0.401595383 | 0.728790008 | -2.0625057 | 0.039159615 | 0.253641284 | 2.536412842 |
| Rpl24 | 3.5317266 | -0.287757429 | 0.629153851 | -2.058155218 | 0.039575236 | 0.253641284 | 2.536412842 |
| Mc1r | 18.88242 | 0.586641047 | 0.40946232 | 2.09174719 | 0.036461137 | 0.253641284 | 2.536412842 |
| mt-Nd5 | 7.1929193 | -0.602111775 | 0.641954842 | -2.06111148 | 0.039292406 | 0.253641284 | 2.536412842 |
| Zbtb7a | 13.372302 | -0.144269504 | 0.728707984 | 1.989806981 | 0.046612202 | 0.264986386 | 2.649863856 |
| Tmem108 | 8.813258 | 0.638130711 | 0.566989712 | 2.01381191 | 0.044029285 | 0.264986386 | 2.649863856 |
| Klhl14 | 8.033774 | -0.144269504 | 0.728707962 | 2.011908491 | 0.044229584 | 0.264986386 | 2.649863856 |
| mt-Co1 | 37.010067 | -0.482954777 | 0.341730296 | -1.9864498 | 0.046983402 | 0.264986386 | 2.649863856 |
| Cox7c | 6.142248 | -0.217428208 | 0.537225504 | -2.012740272 | 0.04414196 | 0.264986386 | 2.649863856 |
| Rpl39 | 4.639669 | -0.436211739 | 0.754677261 | -1.992985257 | 0.046263061 | 0.264986386 | 2.649863856 |
| Riok3 | 3.166993 | -0.304231903 | 0.663264398 | -1.973121293 | 0.048481745 | 0.267440271 | 2.674402711 |
| Cox4i1 | 7.3111577 | -0.378144499 | 0.700655621 | -1.96585617 | 0.049315227 | 0.267440271 | 2.674402711 |
| Fbxl7 | 7.4968495 | 0.578911929 | 0.637851635 | 1.930466211 | 0.053549098 | 0.284921615 | 2.849216152 |
| Sh3glb1 | 3.9933329 | -0.425306001 | 0.723933866 | -1.908299269 | 0.056352547 | 0.28893488 | 2.889348796 |
| Ralyl | 5.797991 | -0.144269504 | 0.728707939 | 1.913199344 | 0.055722537 | 0.28893488 | 2.889348796 |
| Cmss1 | 9.21466 | 0.558005032 | 0.537841572 | 1.891581648 | 0.058546743 | 0.289652307 | 2.896523072 |
| Wdr72 | 11.093801 | 0.546348909 | 0.526462409 | 1.891870621 | 0.058508221 | 0.289652307 | 2.896523072 |
| Rpl32 | 3.9884064 | -0.208440772 | 0.538837685 | -1.854139025 | 0.063719273 | 0.289819918 | 2.898199183 |
| Rps15 | 9.761291 | -0.279743408 | 0.598274068 | -1.857284906 | 0.063270631 | 0.289819918 | 2.898199183 |
| Cox6b1 | 5.308594 | -0.399014395 | 0.692975741 | -1.854953576 | 0.063602856 | 0.289819918 | 2.898199183 |
| Igf2bp3 | 24.474657 | 0.454354986 | 0.33456511 | 1.881068899 | 0.059962547 | 0.289819918 | 2.898199183 |
| Lrrc4c | 5.5929084 | 0.526043194 | 0.69465674 | 1.858312664 | 0.063124626 | 0.289819918 | 2.898199183 |
| Slx1b | 12.773972 | -0.144269504 | 0.728707979 | 1.825460999 | 0.067931492 | 0.299323135 | 2.993231345 |
| Tmsb4x | 8.168612 | -0.353412693 | 0.656044932 | -1.829653065 | 0.067301833 | 0.299323135 | 2.993231345 |
| Rpl12 | 4.9237013 | -0.34987798 | 0.66448028 | -1.80117663 | 0.071675045 | 0.310959424 | 3.109594242 |
| Rplp1 | 19.55245 | -0.39977054 | 0.628948316 | -1.765595207 | 0.077463785 | 0.330981625 | 3.309816255 |

| | | | | | | | |
|---|---|---|---|---|---|---|---|
| 8430422H( | 17.26683 | 0.451065467 | 0.431715488 | 1.726101164 | 0.084329215 | 0.343875004 | 3.438750037 |
| Rpl11 | 8.120596 | -0.228991071 | 0.556706328 | -1.723247135 | 0.084843843 | 0.343875004 | 3.438750037 |
| Mt3 | 5.9679656 | -0.342810217 | 0.638324356 | -1.720403928 | 0.085359043 | 0.343875004 | 3.438750037 |
| Nkain2 | 6.4264207 | -0.144269504 | 0.728707932 | 1.721066963 | 0.085238673 | 0.343875004 | 3.438750037 |
| Hspa14 | 3.1303577 | 0.320079452 | 0.80941095 | 1.702888874 | 0.088588866 | 0.351860005 | 3.518600048 |
| Xlr5a | 50.572094 | 0.284086037 | 0.194769274 | 1.691155615 | 0.090807086 | 0.354022441 | 3.540224414 |
| Galnt13 | 12.247922 | 0.445493146 | 0.462729403 | 1.686719359 | 0.091657328 | 0.354022441 | 3.540224414 |
| mt-Co3 | 83.12322 | -0.376429272 | 0.310600669 | -1.680297198 | 0.092899506 | 0.354022441 | 3.540224414 |
| Stard7 | 4.528309 | 0.362958326 | 0.726511275 | 1.663688365 | 0.096174723 | 0.359311081 | 3.593110811 |
| Rpl29 | 4.449764 | -0.24516367 | 0.582775578 | -1.647847758 | 0.099383916 | 0.359311081 | 3.593110811 |
| 4930402H1 | 6.483537 | -0.365482369 | 0.606903258 | -1.652549938 | 0.098422494 | 0.359311081 | 3.593110811 |
| Eef1a1 | 7.3203444 | -0.144269504 | 0.728707922 | -1.657884407 | 0.0973408 | 0.359311081 | 3.593110811 |
| Mapk14 | 19.08004 | -0.144269504 | 0.728707971 | 1.62051375 | 0.105121963 | 0.37055492 | 3.705549199 |
| Klkb1 | 4.70573 | -0.144269504 | 0.728707962 | 1.621307664 | 0.104951672 | 0.37055492 | 3.705549199 |
| Rpl38 | 10.764174 | -0.272280224 | 0.581438628 | -1.585349548 | 0.112886937 | 0.37476705 | 3.747670498 |
| Rps11 | 3.451559 | -0.207684334 | 0.560376216 | -1.601223108 | 0.109327513 | 0.37476705 | 3.747670498 |
| Gm50306 | 11.34864 | 0.406046452 | 0.48832902 | 1.5850204 | 0.112961699 | 0.37476705 | 3.747670498 |
| Dop1b | 10.740474 | -0.144269504 | 0.728707976 | 1.590943134 | 0.111622374 | 0.37476705 | 3.747670498 |
| Nrg3 | 5.083226 | -0.378752648 | 0.559985185 | -1.607084613 | 0.108035808 | 0.37476705 | 3.747670498 |
| Rpl21 | 6.20196 | -0.186265215 | 0.525960517 | -1.556034965 | 0.119699778 | 0.390504178 | 3.905041779 |
| Mt1 | 3.0413835 | 0.234036698 | 0.788139811 | 1.5470157 | 0.12185946 | 0.390504178 | 3.905041779 |
| Atxn7l1 | 3.7801106 | 0.306389878 | 0.718066348 | 1.54906742 | 0.121365514 | 0.390504178 | 3.905041779 |
| Gm42791 | 3.118952 | -0.250590219 | 0.637417004 | -1.537132865 | 0.124260776 | 0.393725156 | 3.937251563 |
| Rpl17 | 3.2443411 | -0.200082364 | 0.574178894 | -1.483503162 | 0.137940777 | 0.418272034 | 4.182720337 |
| Acoxl | 9.650014 | 0.369155096 | 0.514459475 | 1.491775604 | 0.135757982 | 0.418272034 | 4.182720337 |
| Gm10062 | 29.320692 | 0.353718648 | 0.384726519 | 1.483907451 | 0.137833475 | 0.418272034 | 4.182720337 |
| Usp3 | 4.782866 | 0.275155129 | 0.6960937 | 1.483648495 | 0.137902197 | 0.418272034 | 4.182720337 |
| Cldn16 | 3.4228976 | 0.283791558 | 0.701566248 | 1.460617888 | 0.144120335 | 0.432361004 | 4.323610037 |
| Bc1 | 5.695118 | 0.324729457 | 0.612482802 | 1.453326684 | 0.146133072 | 0.433784487 | 4.337844868 |
| Alpl | 4.200237 | 0.272044261 | 0.658971185 | 1.382165876 | 0.166920775 | 0.480323047 | 4.803230469 |
| Rps27 | 3.8602371 | -0.257310335 | 0.598901616 | -1.383807241 | 0.166417485 | 0.480323047 | 4.803230469 |
| Hdac9 | 4.1805596 | 0.27444825 | 0.660370397 | 1.387925511 | 0.165159729 | 0.480323047 | 4.803230469 |
| Kcnq5 | 6.4754353 | -0.144269504 | 0.728707933 | 1.358070644 | 0.174441269 | 0.496893312 | 4.968933123 |
| Reps2 | 4.068739 | -0.241276026 | 0.58480249 | -1.347424101 | 0.177843683 | 0.497708386 | 4.97708386 |
| mt-Co2 | 45.869038 | -0.290233452 | 0.305302289 | -1.346140347 | 0.178257259 | 0.497708386 | 4.97708386 |
| Col5a2 | 4.2028365 | 0.242790875 | 0.660759872 | 1.320694523 | 0.186603241 | 0.515903078 | 5.159030782 |
| Ralgps1 | 3.16889 | -0.221347866 | 0.612643894 | -1.314163242 | 0.188791276 | 0.516596873 | 5.165968734 |
| Epha6 | 19.994076 | 0.299724304 | 0.369704609 | 1.303658926 | 0.1923499 | 0.516596873 | 5.165968734 |
| A630012P( | 8.247962 | -0.292555602 | 0.478113966 | -1.308083616 | 0.19084495 | 0.516596873 | 5.165968734 |
| Dtnb | 3.8421476 | 0.253470933 | 0.643871462 | 1.2950425 | 0.195305569 | 0.519586513 | 5.195865133 |
| 4930441J1 | 7.928579 | -0.267864869 | 0.518927449 | -1.279741329 | 0.200636124 | 0.528779318 | 5.287793181 |
| Rps26 | 4.0899043 | -0.196815421 | 0.564144145 | -1.26960956 | 0.204223742 | 0.532513492 | 5.325134917 |
| 5830416I1 | 50.722412 | 0.240841091 | 0.240915275 | 1.265116182 | 0.205829683 | 0.532513492 | 5.325134917 |
| Dpyd | 9.490453 | -0.144269504 | 0.728707943 | 1.252273247 | 0.210470314 | 0.53956935 | 5.395693502 |
| Gm26901 | 3.7745736 | -0.144269504 | 0.728707929 | -1.23742958 | 0.215927643 | 0.547471981 | 5.474719808 |
| Asxl3 | 6.784757 | 0.26395452 | 0.550947987 | 1.22818622 | 0.21937707 | 0.547471981 | 5.474719808 |
| Mrc2 | 13.059499 | -0.277019301 | 0.406655188 | -1.229658094 | 0.218825166 | 0.547471981 | 5.474719808 |
| Rpl13 | 10.277906 | -0.232545601 | 0.521015952 | -1.206957169 | 0.227448639 | 0.554846398 | 5.548463979 |
| Sult2a2 | 5.4808154 | 0.238906434 | 0.596749014 | 1.209400358 | 0.226509071 | 0.554846398 | 5.548463979 |
| Gm42889 | 5.6544228 | -0.144269504 | 0.728707907 | -1.204918701 | 0.228234688 | 0.554846398 | 5.548463979 |
| Tacc1 | 3.4373827 | -0.144269504 | 0.728707914 | 1.196888757 | 0.231349917 | 0.55761262 | 5.576126202 |
| Gm49767 | 5.579478 | -0.222683939 | 0.547644035 | -1.185914682 | 0.235655995 | 0.563177885 | 5.631778853 |
| Lrrc17 | 9.684402 | 0.259101481 | 0.482956874 | 1.166025347 | 0.243604198 | 0.572469864 | 5.724698643 |
| Pde6a | 33.298298 | 0.245803294 | 0.294912328 | 1.168318936 | 0.24267814 | 0.572469864 | 5.724698643 |
| Sec61a1 | 96.05451 | -0.183887333 | 0.186367857 | -1.152271639 | 0.249209467 | 0.580802229 | 5.808022294 |
| Gm12216 | 47.633396 | -0.144269504 | 0.728707994 | 1.139628258 | 0.254441208 | 0.583353013 | 5.833530125 |
| Cntnap5b | 3.511289 | -0.183802155 | 0.573463134 | -1.141530172 | 0.253649364 | 0.583353013 | 5.833530125 |
| Rps25 | 4.7889094 | -0.16684379 | 0.539120214 | -1.13203392 | 0.257620174 | 0.585878137 | 5.858781375 |
| Rpl30 | 12.249324 | -0.189160852 | 0.522007051 | -1.12683614 | 0.259811767 | 0.586135347 | 5.861353472 |
| Ptprd | 5.949268 | 0.189471681 | 0.626691788 | 1.120318184 | 0.262578196 | 0.58767501 | 5.8767501 |
| Rps8 | 8.564133 | -0.142483398 | 0.511303245 | -1.105580645 | 0.268908071 | 0.59710296 | 5.971029605 |
| Cox7a2 | 4.9629493 | -0.146552625 | 0.533920986 | -1.088690157 | 0.276290544 | 0.60398398 | 6.039839799 |
| Sema3c | 85.11226 | -0.144269504 | 0.728708018 | -1.089964945 | 0.275728586 | 0.60398398 | 6.039839799 |
| St18 | 30.757889 | -0.217592675 | 0.284992102 | -1.06805582 | 0.285495333 | 0.619305261 | 6.193052614 |
| Rab29 | 10.738876 | -0.227486072 | 0.446279153 | -1.057737336 | 0.290175208 | 0.624651975 | 6.246519753 |
| Fam189a2 | 4.7822156 | -0.144269504 | 0.728707913 | -1.045607813 | 0.295742141 | 0.627062283 | 6.270622832 |
| Ms4a1 | 13.051231 | 0.234026956 | 0.414903412 | 1.04669871 | 0.295238559 | 0.627062283 | 6.270622832 |
| 4930558K( | 11.312633 | 0.228282869 | 0.448220449 | 1.037237382 | 0.29962524 | 0.63055461 | 6.3055461 |
| Rapgef4 | 26.139738 | -0.144269504 | 0.728707959 | 1.022743304 | 0.306429231 | 0.635390023 | 6.353900229 |

| | | | | | | | |
|---|---|---|---|---|---|---|---|
| Gnaz | 22.523691 | 0.225205857 | 0.35312447 | 1.023872226 | 0.30589563 | 0.635390023 | 6.353900229 |
| Dctd | 3.2623832 | 0.165386921 | 0.621824636 | 0.970404123 | 0.331845094 | 0.673239687 | 6.732396874 |
| Dppa2 | 5.75971 | -0.172062896 | 0.524127412 | -0.970444203 | 0.331825124 | 0.673239687 | 6.732396874 |
| AI314278 | 5.2436852 | -0.19684422 | 0.498023399 | -0.970684901 | 0.331705213 | 0.673239687 | 6.732396874 |
| Rubcnl | 6.2419114 | -0.191283257 | 0.49384827 | -0.955737285 | 0.33920498 | 0.683255746 | 6.832557459 |
| Unc5d | 15.126615 | -0.198224785 | 0.394763307 | -0.926925537 | 0.353965197 | 0.707930394 | 7.079303945 |
| Rpl18a | 5.469551 | -0.116309084 | 0.515353972 | -0.903832907 | 0.36608401 | 0.721927907 | 7.21927907 |
| Zfpm2 | 4.781524 | -0.163824791 | 0.529999833 | -0.904365358 | 0.365801701 | 0.721927907 | 7.21927907 |
| Vmn2r103 | 5.408102 | -0.15156615 | 0.529860695 | -0.893448527 | 0.371617025 | 0.727750008 | 7.277500082 |
| Grid2 | 3.9817505 | 0.154214333 | 0.591751228 | 0.883385157 | 0.377028208 | 0.728232566 | 7.282325655 |
| Kcnh7 | 3.3050241 | 0.150727962 | 0.610022886 | 0.886183358 | 0.37551873 | 0.728232566 | 7.282325655 |
| Ppp1r9a | 10.468803 | -0.185958961 | 0.418625853 | -0.874954206 | 0.381598823 | 0.732046722 | 7.320467219 |
| Ccdc60 | 7.7561297 | 0.181873304 | 0.479858691 | 0.867353391 | 0.385748411 | 0.735007108 | 7.350071076 |
| Gm38973 | 6.152351 | 0.167561167 | 0.530782114 | 0.861990776 | 0.388692594 | 0.735646385 | 7.35646385 |
| Srgn | 3.6218865 | 0.174346464 | 0.54693308 | 0.856236921 | 0.391866744 | 0.735663811 | 7.356638112 |
| Phkg1 | 6.902163 | 0.169589587 | 0.512568895 | 0.852531299 | 0.393919275 | 0.735663811 | 7.356638112 |
| Prkag2 | 7.5835156 | 0.172889771 | 0.48420393 | 0.834876085 | 0.403787481 | 0.74423575 | 7.442357496 |
| Pcsk2 | 3.1961663 | 0.124893439 | 0.628308912 | 0.836327034 | 0.402970949 | 0.74423575 | 7.442357496 |
| Cmpk1 | 27.24363 | 0.174979359 | 0.315751048 | 0.829922129 | 0.406582812 | 0.744521773 | 7.445217731 |
| Klf10 | 4.765948 | -0.135480891 | 0.538658161 | -0.807747294 | 0.419236072 | 0.762739176 | 7.627391758 |
| Sgip1 | 3.156119 | 0.119869572 | 0.619938069 | 0.794954889 | 0.426639744 | 0.771233383 | 7.712333827 |
| Chl1 | 7.8284264 | -0.138306148 | 0.505344377 | -0.786496242 | 0.43157682 | 0.775188937 | 7.751889375 |
| Ppp1r16a | 9.846262 | -0.158766917 | 0.390636198 | -0.757271453 | 0.44888725 | 0.791742832 | 7.917428322 |
| mt-Nd4 | 20.98154 | -0.144269504 | 0.72870795 | 0.728780844 | 0.466135731 | 0.791742832 | 7.917428322 |
| 1700126G( | 17.642927 | 0.155944962 | 0.350970867 | 0.729753556 | 0.465540838 | 0.791742832 | 7.917428322 |
| Pard3b | 3.7414958 | -0.144269504 | 0.72870791 | -0.760800065 | 0.446776495 | 0.791742832 | 7.917428322 |
| Gm765 | 6.7321286 | -0.144269504 | 0.728707925 | 0.72114445 | 0.470820645 | 0.791742832 | 7.917428322 |
| Rpl37 | 8.0153265 | -0.101387051 | 0.512797979 | -0.73333369 | 0.463354932 | 0.791742832 | 7.917428322 |
| Elf1 | 3.119602 | 0.117688431 | 0.60410685 | 0.729371941 | 0.465774176 | 0.791742832 | 7.917428322 |
| Svil | 6.252604 | 0.139782144 | 0.520399871 | 0.732944321 | 0.463592389 | 0.791742832 | 7.917428322 |
| Cdh20 | 4.890388 | -0.126603957 | 0.519871592 | -0.716603202 | 0.473618968 | 0.791742832 | 7.917428322 |
| Shb | 29.469902 | 0.149958363 | 0.276955993 | 0.749801041 | 0.453374542 | 0.791742832 | 7.917428322 |
| Kat7 | 7.819134 | 0.148544206 | 0.470446018 | 0.72456198 | 0.468720791 | 0.791742832 | 7.917428322 |
| Adamtsl1 | 55.07283 | -0.144269504 | 0.728707995 | -0.715202087 | 0.47448418 | 0.791742832 | 7.917428322 |
| Gm20342 | 5.603552 | -0.134515368 | 0.49737134 | -0.710147335 | 0.477612775 | 0.792275309 | 7.922753095 |
| Cntn3 | 6.040966 | 0.133798738 | 0.520313317 | 0.704907293 | 0.480867931 | 0.793010272 | 7.930102724 |
| 1700054A( | 97.21484 | -0.120570102 | 0.20951411 | -0.700238196 | 0.483778561 | 0.793171827 | 7.931718272 |
| Ctnna2 | 10.168796 | -0.13472659 | 0.436296569 | -0.666938969 | 0.504811118 | 0.807097397 | 8.070973971 |
| Tatdn1 | 3.045824 | 0.107779774 | 0.60016044 | 0.670771905 | 0.502365848 | 0.807097397 | 8.070973971 |
| Ssh2 | 5.0847206 | -0.115966171 | 0.518358056 | -0.665316641 | 0.505847989 | 0.807097397 | 8.070973971 |
| Sorcs2 | 7.121006 | 0.128701803 | 0.497525793 | 0.664168599 | 0.506582409 | 0.807097397 | 8.070973971 |
| D430042O | 34.25977 | -0.136554264 | 0.304755411 | -0.668539126 | 0.503789516 | 0.807097397 | 8.070973971 |
| Gm42418 | 75.15459 | 0.100899705 | 0.181588013 | 0.648426216 | 0.516709319 | 0.813584731 | 8.135847313 |
| mt-Cytb | 21.743904 | 0.136313933 | 0.341164747 | 0.644410307 | 0.519309403 | 0.813584731 | 8.135847313 |
| Rpl27a | 7.7989106 | -0.094018568 | 0.514867599 | -0.651351399 | 0.514819676 | 0.813584731 | 8.135847313 |
| Cep128 | 5.2810345 | -0.144269504 | 0.728707947 | 0.630130104 | 0.528609465 | 0.823361955 | 8.233619546 |
| Gm960 | 3.3566253 | -0.098626608 | 0.549731737 | -0.625886134 | 0.53138963 | 0.823361955 | 8.233619546 |
| Csmd3 | 14.582181 | -0.127926036 | 0.372146799 | -0.61077549 | 0.541348222 | 0.824993199 | 8.24993199 |
| Kcnip4 | 6.6033077 | -0.098121445 | 0.495115167 | -0.549973681 | 0.582337425 | 0.824993199 | 8.24993199 |
| Adk | 6.9709363 | 0.104870202 | 0.514790864 | 0.574146828 | 0.565868447 | 0.824993199 | 8.24993199 |
| Naa11 | 5.386561 | -0.106563667 | 0.510670301 | -0.601441545 | 0.547545935 | 0.824993199 | 8.24993199 |
| mt-Atp6 | 66.57476 | -0.11018385 | 0.291314835 | -0.545948178 | 0.58510156 | 0.824993199 | 8.24993199 |
| Ano2 | 3.4092267 | 0.08393782 | 0.587695186 | 0.550515144 | 0.581966092 | 0.824993199 | 8.24993199 |
| Mbp | 4.5371413 | -0.089555719 | 0.52731716 | -0.558266219 | 0.576662609 | 0.824993199 | 8.24993199 |
| Meg3 | 7.231478 | 0.094201111 | 0.544781941 | 0.572763347 | 0.566804939 | 0.824993199 | 8.24993199 |
| Wdyhv1 | 9.009689 | -0.144269504 | 0.728707941 | 0.586772524 | 0.557356493 | 0.824993199 | 8.24993199 |
| Fbxo4 | 3.462966 | -0.095428963 | 0.546225032 | -0.577464606 | 0.563625642 | 0.824993199 | 8.24993199 |
| 6030443J0 | 14.052521 | -0.144269504 | 0.728707961 | 0.611364221 | 0.540958483 | 0.824993199 | 8.24993199 |
| Nell1 | 11.76224 | 0.105262594 | 0.472489444 | 0.549497144 | 0.582664324 | 0.824993199 | 8.24993199 |
| Dbi | 3.1805382 | -0.069306142 | 0.53954207 | -0.564998834 | 0.572074557 | 0.824993199 | 8.24993199 |
| Pinlyp | 3.7137024 | -0.093084929 | 0.549350407 | -0.59514541 | 0.551746277 | 0.824993199 | 8.24993199 |
| Rpl26 | 9.565828 | -0.078666799 | 0.507227411 | -0.565451432 | 0.57176675 | 0.824993199 | 8.24993199 |
| Tpd52l1 | 3.6817331 | -0.097068217 | 0.536921122 | -0.608045588 | 0.543157238 | 0.824993199 | 8.24993199 |
| Olfr57 | 3.1732054 | -0.081040137 | 0.556881369 | -0.605028246 | 0.545160229 | 0.824993199 | 8.24993199 |
| mt-Nd3 | 11.661037 | 0.122055907 | 0.373667614 | 0.574666572 | 0.56551682 | 0.824993199 | 8.24993199 |
| Lingo2 | 4.323964 | 0.077612871 | 0.575502228 | 0.506851665 | 0.612258902 | 0.826110098 | 8.261100976 |
| Gm11337 | 8.366889 | -0.144269504 | 0.728707938 | -0.511159121 | 0.60923964 | 0.826110098 | 8.261100976 |
| Rpl37a | 6.858608 | -0.091796655 | 0.488783411 | -0.507767332 | 0.611616522 | 0.826110098 | 8.261100976 |
| Gnao1 | 4.0632005 | 0.088931781 | 0.554665754 | 0.519151938 | 0.603654792 | 0.826110098 | 8.261100976 |

| | | | | | | | |
|---|---|---|---|---|---|---|---|
| Ninj2 | 11.427499 | -0.108104396 | 0.398985372 | -0.5266948 | 0.598405548 | 0.826110098 | 8.261100976 |
| Gria1 | 3.6526048 | -0.085643197 | 0.543946737 | -0.519442532 | 0.603452178 | 0.826110098 | 8.261100976 |
| Esf1 | 4.1028438 | 0.086205522 | 0.554285879 | 0.50786828 | 0.611545721 | 0.826110098 | 8.261100976 |
| Ttc28 | 14.362916 | 0.107692704 | 0.332803745 | 0.515525249 | 0.60618603 | 0.826110098 | 8.261100976 |
| Tns3 | 5.9620595 | 0.09517622 | 0.527926536 | 0.538266592 | 0.590393012 | 0.826110098 | 8.261100976 |
| Rgs6 | 15.358516 | 0.102568724 | 0.414038855 | 0.497854508 | 0.618586594 | 0.830673426 | 8.30673426 |
| Ahdc1 | 7.62892 | -0.144269504 | 0.728707942 | 0.487451803 | 0.625938191 | 0.834218732 | 8.342187321 |
| Lin28b | 3.585234 | 0.078204748 | 0.574967302 | 0.485751789 | 0.62714316 | 0.834218732 | 8.342187321 |
| Cspp1 | 6.8032126 | -0.144269504 | 0.728707933 | 0.466699501 | 0.640714887 | 0.844711051 | 8.447110514 |
| Spock1 | 7.3607025 | -0.080903331 | 0.493519767 | -0.466270542 | 0.641021862 | 0.844711051 | 8.447110514 |
| Nrp2 | 3.2675326 | 0.068878202 | 0.575602002 | 0.441096004 | 0.659143496 | 0.860548453 | 8.605484535 |
| A330008L1 | 4.552196 | 0.08053153 | 0.527110862 | 0.445053701 | 0.656280947 | 0.860548453 | 8.605484535 |
| Ptpn9 | 11.600575 | -0.088460627 | 0.401896778 | -0.433759123 | 0.664463368 | 0.863496174 | 8.634961742 |
| Gm28729 | 3.088824 | -0.060578873 | 0.556018305 | -0.393955256 | 0.693614074 | 0.867540864 | 8.675408637 |
| 4930511M | 17.627966 | -0.084913431 | 0.356601694 | -0.412780223 | 0.679767643 | 0.867540864 | 8.675408637 |
| Cdh23 | 13.755928 | -0.144269504 | 0.728707961 | 0.412131968 | 0.680242699 | 0.867540864 | 8.675408637 |
| Chsy3 | 12.854756 | -0.082135118 | 0.378782263 | -0.400653528 | 0.68867523 | 0.867540864 | 8.675408637 |
| Camk2a | 10.730389 | 0.08116599 | 0.450550993 | 0.410810659 | 0.681211377 | 0.867540864 | 8.675408637 |
| Opcml | 3.8994443 | 0.06565882 | 0.556284155 | 0.398379105 | 0.690350758 | 0.867540864 | 8.675408637 |
| Anks1b | 3.1884053 | -0.144269504 | 0.728707911 | -0.394888596 | 0.692925106 | 0.867540864 | 8.675408637 |
| Il23a | 3.9379215 | 0.066558424 | 0.547539196 | 0.391722524 | 0.695263245 | 0.867540864 | 8.675408637 |
| Hdac8 | 7.3132796 | 0.081584849 | 0.471983364 | 0.418466631 | 0.675605979 | 0.867540864 | 8.675408637 |
| Gria4 | 15.817715 | -0.144269504 | 0.728707954 | -0.386394374 | 0.699204616 | 0.868615426 | 8.686154264 |
| Rpl19 | 3.2033045 | 0.052714131 | 0.587464224 | 0.37756414 | 0.705754408 | 0.872906768 | 8.72906768 |
| Ehbp1 | 18.458115 | 0.067571304 | 0.367521017 | 0.324577741 | 0.745500676 | 0.898423892 | 8.984238917 |
| Pcca | 6.43877 | -0.05731577 | 0.49853757 | -0.344759946 | 0.730274847 | 0.898423892 | 8.984238917 |
| Slco2a1 | 3.8558457 | 0.055291907 | 0.552323231 | 0.335828987 | 0.73699983 | 0.898423892 | 8.984238917 |
| Gm14582 | 6.949116 | -0.144269504 | 0.728707924 | 0.327144791 | 0.743558376 | 0.898423892 | 8.984238917 |
| Pzp | 4.3720546 | -0.057665634 | 0.518947133 | -0.333883995 | 0.738467098 | 0.898423892 | 8.984238917 |
| AY036118 | 15.008479 | 0.062735797 | 0.454205404 | 0.329328205 | 0.741907627 | 0.898423892 | 8.984238917 |
| mt-Nd1 | 15.216 | -0.144269504 | 0.728707944 | -0.30357234 | 0.761453727 | 0.909700336 | 9.097003363 |
| Syt1 | 4.2310753 | 0.047649471 | 0.539006928 | 0.286868446 | 0.774213052 | 0.909700336 | 9.097003363 |
| Anapc15 | 4.5367656 | -0.048062078 | 0.523714094 | -0.289129234 | 0.772482483 | 0.909700336 | 9.097003363 |
| Zmat4 | 13.51533 | -0.061670797 | 0.353503028 | -0.303029748 | 0.761867191 | 0.909700336 | 9.097003363 |
| Cdh12 | 3.938876 | 0.048976038 | 0.549950676 | 0.294938468 | 0.768040885 | 0.909700336 | 9.097003363 |
| Fyb2 | 10.5768385 | 0.059160488 | 0.427854395 | 0.293008709 | 0.769515495 | 0.909700336 | 9.097003363 |
| Lsamp | 10.066482 | 0.057162084 | 0.422609647 | 0.280604475 | 0.779013783 | 0.911543099 | 9.11543099 |
| Trim33 | 3.9992864 | 0.04037111 | 0.561073567 | 0.267551702 | 0.789044412 | 0.919464976 | 9.194649757 |
| Cox6c | 6.096412 | -0.049064368 | 0.459785867 | -0.254543993 | 0.799075322 | 0.927321978 | 9.273219784 |
| Pkp2 | 6.4001365 | -0.044482755 | 0.476192518 | -0.240761816 | 0.809739725 | 0.928238222 | 9.282382218 |
| Cadm2 | 3.3506944 | 0.036271041 | 0.575256176 | 0.245386 | 0.806157568 | 0.928238222 | 9.282382218 |
| Vwc2l | 15.776025 | -0.049225948 | 0.351308447 | -0.241673626 | 0.809033067 | 0.928238222 | 9.282382218 |
| Dmd | 4.9338746 | -0.039004988 | 0.513031654 | -0.231361916 | 0.817033649 | 0.932807648 | 9.32807648 |
| Chchd6 | 3.772844 | -0.036512262 | 0.534844754 | -0.224355962 | 0.822480335 | 0.935239735 | 9.352397354 |
| Pcgf6 | 7.014855 | 0.039908528 | 0.472783697 | 0.208593808 | 0.834765345 | 0.945396896 | 9.453968963 |
| Rpl13a | 4.0575385 | -0.02783083 | 0.536351192 | -0.197790085 | 0.843209304 | 0.946100381 | 9.461003807 |
| Cpe | 3.726711 | 0.024504786 | 0.58301131 | 0.187428814 | 0.851324432 | 0.946100381 | 9.461003807 |
| Cdk8 | 3.9378386 | 0.027065839 | 0.566867268 | 0.194377047 | 0.845880661 | 0.946100381 | 9.461003807 |
| Rsrc1 | 22.222565 | -0.035426587 | 0.380816131 | -0.177811179 | 0.858871268 | 0.946100381 | 9.461003807 |
| Gm2895 | 3.8460655 | -0.029260971 | 0.532821697 | -0.179692414 | 0.857394049 | 0.946100381 | 9.461003807 |
| Gm29325 | 8.049865 | 0.039267891 | 0.439922242 | 0.195530816 | 0.844977415 | 0.946100381 | 9.461003807 |
| Ctcflos | 8.459374 | 0.035165512 | 0.458737928 | 0.182626504 | 0.855091087 | 0.946100381 | 9.461003807 |
| Cfap74 | 6.4889827 | 0.032903548 | 0.472744517 | 0.171466211 | 0.863857197 | 0.947889998 | 9.478899983 |
| Zfp994 | 7.226358 | 0.025327734 | 0.51337468 | 0.152047414 | 0.879149539 | 0.957220734 | 9.572207338 |
| A930016O | 5.371809 | -0.144269504 | 0.728707922 | -0.154359763 | 0.877326078 | 0.957220734 | 9.572207338 |
| Sdccag8 | 4.707819 | -0.144269504 | 0.72870793 | 0.147449409 | 0.882777309 | 0.957473851 | 9.574738509 |
| Pcdh15 | 14.132702 | -0.027573438 | 0.334090502 | -0.138359468 | 0.889956329 | 0.96156201 | 9.615620103 |
| mt-Nd2 | 21.215528 | -0.02555715 | 0.343545994 | -0.126372286 | 0.899437239 | 0.964415594 | 9.644155944 |
| Malat1 | 12.407729 | -0.014819235 | 0.52669274 | -0.12786894 | 0.898252691 | 0.964415594 | 9.644155944 |
| Atp9b | 10.61692 | 0.025091214 | 0.402905541 | 0.120949067 | 0.903731378 | 0.965349426 | 9.65349426 |
| Kctd16 | 47.13608 | 0.022798772 | 0.311400897 | 0.109852593 | 0.91252628 | 0.971065702 | 9.710657016 |
| Asic2 | 4.113033 | -0.012102681 | 0.531348923 | -0.075138698 | 0.940104358 | 0.980089172 | 9.800891723 |
| Rpl34 | 3.1515095 | 0.010521477 | 0.56896715 | 0.068458517 | 0.945420641 | 0.980089172 | 9.800891723 |
| Comt | 3.7562973 | -0.014206121 | 0.535465675 | -0.09072551 | 0.9277107 | 0.980089172 | 9.800891723 |
| Ctnna3 | 12.739343 | 0.015932361 | 0.363355399 | 0.075073953 | 0.940155872 | 0.980089172 | 9.800891723 |
| Scamp1 | 9.79033 | -0.012376983 | 0.416266342 | -0.064201566 | 0.948809731 | 0.980089172 | 9.800891723 |
| Sil1 | 15.781804 | 0.018567181 | 0.31281797 | 0.08794882 | 0.929917354 | 0.980089172 | 9.800891723 |
| Tcea1 | 5.23988 | 0.011484232 | 0.516165533 | 0.066216057 | 0.947205813 | 0.980089172 | 9.800891723 |
| Tmeff2 | 4.646029 | 0.011809233 | 0.513568747 | 0.06496399 | 0.948202671 | 0.980089172 | 9.800891723 |

| | | | | | | | |
|---|---|---|---|---|---|---|---|
| Ube2cbp | 4.3902645 | 0.007405999 | 0.531339567 | 0.042122632 | 0.966400938 | 0.988820715 | 9.888207147 |
| Il27 | 15.843124 | -0.007718893 | 0.37400224 | -0.040390284 | 0.967781976 | 0.988820715 | 9.888207147 |
| Camk1d | 9.065154 | -0.008953866 | 0.429303385 | -0.047061933 | 0.962463867 | 0.988820715 | 9.888207147 |
| Zfhx3 | 5.9161816 | 0.006591817 | 0.48819756 | 0.033733403 | 0.973089743 | 0.99065454 | 9.906545396 |
| Magi2 | 3.4612923 | 0.003153653 | 0.560974526 | 0.020581224 | 0.983579718 | 0.994155844 | 9.94155844 |
| Gm49497 | 7.449143 | 0.004661032 | 0.485049944 | 0.023096767 | 0.981573084 | 0.994155844 | 9.94155844 |
| Kcnd2 | 4.2012353 | 0.001424508 | 0.536289564 | 0.006417456 | 0.994879646 | 0.998420143 | 9.98420143 |
| Ptprg | 4.347489 | -0.001040085 | 0.52902264 | -0.008507571 | 0.993212022 | 0.998420143 | 9.98420143 |
| Arvcf | 4.196541 | 0.000507898 | 0.533918826 | 0.000668592 | 0.999466541 | 0.999466541 | 9.994665406 |
| 4931429P1 | 2.310384 | 0.085439905 | 0.64105118 | 0.653053476 | 0.513721804 | | |
| 9130015L2 | 2.9100034 | -0.121556746 | 0.573940173 | -0.809686761 | 0.418120229 | | |
| Alk | 2.1714432 | -0.144269504 | 0.728707943 | 1.375213184 | 0.169065362 | | |
| Ap3s1 | 2.7727785 | 0.148995166 | 0.670273382 | 1.020878894 | 0.30731182 | | |
| Atp5e | 2.6827114 | -0.153406493 | 0.526952916 | -1.488026485 | 0.136743909 | | |
| Atp5md | 2.276934 | 0.194278684 | 0.854193114 | 1.506059434 | 0.132051917 | | |
| Brip1os | 2.772364 | -0.025161452 | 0.56870312 | -0.171111615 | 0.864136003 | | |
| C330002G | 2.487561 | 0.135744284 | 0.644005585 | 0.881485049 | 0.378055343 | | |
| Cadps2 | 2.149689 | -0.150636907 | 0.608902463 | -1.046522069 | 0.295320061 | | |
| Calb1 | 1.6046982 | 0.068475864 | 0.665609922 | 0.596572523 | 0.550792821 | | |
| Calm1 | 2.9218295 | 0.021269721 | 0.578778152 | 0.161063116 | 0.87204369 | | |
| Celf2 | 2.863083 | 0.056463024 | 0.629670897 | 0.493606256 | 0.621584282 | | |
| Chchd2 | 2.0120494 | -0.031451556 | 0.587650224 | -0.236884642 | 0.812746285 | | |
| Cox11 | 2.6252534 | 0.036217822 | 0.591818031 | 0.253783531 | 0.799662798 | | |
| Crhr2 | 2.7763135 | -0.049889355 | 0.574297373 | -0.336965398 | 0.736142985 | | |
| Dlg2 | 2.9164665 | 0.084027996 | 0.618822551 | 0.616006994 | 0.537889904 | | |
| Dlgap1 | 2.2567644 | 0.087667912 | 0.634865419 | 0.642017369 | 0.520861905 | | |
| Fam155a | 2.3172529 | 0.080562985 | 0.627975084 | 0.572506794 | 0.566978684 | | |
| Fgf12 | 1.8536628 | 0.111932057 | 0.682197167 | 0.86648398 | 0.386224808 | | |
| Gas5 | 2.5405552 | -0.096595597 | 0.586301641 | -0.668297597 | 0.503943647 | | |
| Gm13219 | 1.5715543 | -0.039480447 | 0.615715748 | -0.333009851 | 0.739126847 | | |
| Gm16046 | 2.3358028 | -0.04773909 | 0.562145711 | -0.38559885 | 0.699793785 | | |
| Gm17529 | 2.7098987 | 0.176462392 | 0.654474313 | 1.063705096 | 0.287462315 | | |
| Gphn | 2.6941593 | -0.030194562 | 0.567663389 | -0.224182195 | 0.822615537 | | |
| Hexb | 2.463273 | 0.010609167 | 0.588859769 | 0.074784597 | 0.940386097 | | |
| Hipk2 | 2.3546884 | -0.259418051 | 0.665106113 | -1.663766957 | 0.09615901 | | |
| Kif5a | 2.1948159 | -0.04848062 | 0.582755751 | -0.404994611 | 0.685481472 | | |
| Lrrc51 | 2.2366729 | -0.125481195 | 0.594857978 | -0.880580349 | 0.378544999 | | |
| Ly75 | 2.5411646 | -0.088663402 | 0.580260534 | -0.636438755 | 0.524490488 | | |
| Lyrm1 | 2.925868 | 0.309139967 | 0.78955798 | 1.642182194 | 0.100552252 | | |
| Micu1 | 1.8262444 | -0.02403355 | 0.613946487 | -0.204996957 | 0.837574503 | | |
| Nup210l | 2.5739462 | -0.022577487 | 0.566117798 | -0.174311448 | 0.861620703 | | |
| Pecam1 | 2.0981572 | -0.019052604 | 0.588860003 | -0.139887456 | 0.888748912 | | |
| Plp1 | 1.9981508 | -0.067514806 | 0.595190229 | -0.514033709 | 0.607228423 | | |
| Ppia | 3.0083334 | -0.242431843 | 0.593352048 | -1.803066371 | 0.071377792 | | |
| Ppp3ca | 2.159522 | -0.032523685 | 0.587148095 | -0.251534097 | 0.801401204 | | |
| Rpl23 | 2.6464455 | 0.06553 | 0.595639369 | 0.440615859 | 0.659491119 | | |
| Rpl31 | 2.1410306 | 0.017364131 | 0.618285995 | 0.159254346 | 0.873468488 | | |
| Rpl36 | 2.0195076 | -0.093093282 | 0.559081508 | -0.775057536 | 0.438305662 | | |
| Rps13 | 1.6739355 | 0.176753534 | 0.782899725 | 1.282147027 | 0.199791075 | | |
| Rps14 | 2.2897723 | 0.192139504 | 0.713633758 | 1.242097107 | 0.214200736 | | |
| Rps17 | 2.0987666 | -0.0166028 | 0.586842461 | -0.126009278 | 0.89972458 | | |
| Rps23 | 2.5489166 | -0.064750109 | 0.538976256 | -0.580343019 | 0.561683323 | | |
| Rps7 | 0.8259451 | 0.06405067 | 1.074505021 | 0.777570166 | 0.436822455 | | |
| Rtkn | 2.9219913 | -0.071477012 | 0.546955063 | -0.5971176 | 0.550428868 | | |
| Selenow | 1.5719209 | 0.037323838 | 0.671432681 | 0.443165071 | 0.657646342 | | |
| Sgcz | 2.4237251 | 0.070912457 | 0.628200883 | 0.515384136 | 0.606284616 | | |
| Sh3bgrl2 | 2.953968 | 0.00374881 | 0.566615271 | 0.022572141 | 0.981991567 | | |
| Slc24a3 | 2.551067 | 0.210453258 | 0.740004429 | 1.368009848 | 0.171308993 | | |
| Srbd1 | 2.1976688 | 0.216368869 | 0.685992957 | 1.184795835 | 0.23609818 | | |
| Topaz1 | 1.939161 | -0.144269504 | 0.728707929 | 1.13396266 | 0.256810211 | | |
| Tpt1 | 0.76028436 | 0.051699672 | 1.133939583 | 0.682221282 | 0.49509904 | | |
| Tsacc | 2.6871746 | 0.026604923 | 0.59218142 | 0.183687476 | 0.854258634 | | |
| Tshr | 2.7266922 | 0.099121664 | 0.592468993 | 0.594624737 | 0.552094341 | | |
| Uba52 | 1.3251296 | 0.011897874 | 0.650003559 | 0.109150247 | 0.913083321 | | |
| Ubb | 2.8230526 | -0.135012798 | 0.548291884 | -1.062852955 | 0.28784864 | | |
| Vit | 0.39625272 | 0.036543088 | 1.030352308 | 0.365810021 | 0.714506842 | | |
| Wbscr25 | 1.1314259 | 0.060842774 | 0.719061136 | 0.634832713 | 0.525537526 | | |
| mt-Atp8 | 2.1064222 | 0.042112664 | 0.619435637 | 0.321643412 | 0.747722852 | | |
| mt-Nd4l | 1.9953672 | -0.032722464 | 0.586909895 | -0.250675428 | 0.80206506 | | |

| **Glut** | baseMean | log2FoldChange | lfcSE | stat | pvalue | padj | pglobadj |
|---|---|---|---|---|---|---|---|
| Rplp1 | 95.34244 | -2.418573756 | 0.174090882 | -14.01614222 | 1.24E-44 | 4.25E-42 | 4.25E-41 |
| Cox4i1 | 70.2725 | -2.733422984 | 0.201574956 | -13.67024315 | 1.53E-42 | 2.61E-40 | 2.61E-39 |
| Rpl37 | 69.7033 | -2.165090482 | 0.187728091 | -11.67259321 | 1.76E-31 | 2.01E-29 | 2.01E-28 |
| Cst3 | 223.14856 | -2.847546895 | 0.273775806 | -10.60771638 | 2.74E-26 | 2.35E-24 | 2.35E-23 |
| Rpl13 | 64.833275 | -2.890463149 | 0.328413462 | -9.039983778 | 1.57E-19 | 1.07E-17 | 1.07E-16 |
| Cox8a | 62.06467 | -3.089346022 | 0.358958703 | -8.838296423 | 9.72E-19 | 5.54E-17 | 5.54E-16 |
| Rps24 | 64.62592 | -2.829954907 | 0.352119581 | -8.293383059 | 1.10E-16 | 5.38E-15 | 5.38E-14 |
| Apoe | 94.01519 | -1.965681147 | 0.246333739 | -8.202299285 | 2.36E-16 | 1.01E-14 | 1.01E-13 |
| Rps20 | 39.05046 | -2.331450396 | 0.295693891 | -8.103527788 | 5.34E-16 | 2.03E-14 | 2.03E-13 |
| Rps8 | 60.03716 | -2.349384361 | 0.325445196 | -7.487184995 | 7.04E-14 | 2.41E-12 | 2.41E-11 |
| 5830416I1 | 133.81793 | -0.144269504 | 0.849531651 | 7.467634981 | 8.16E-14 | 2.54E-12 | 2.54E-11 |
| Rpl17 | 37.72632 | -2.561528706 | 0.359662593 | -7.390284469 | 1.47E-13 | 4.18E-12 | 4.18E-11 |
| Rps10 | 37.998 | -2.294345695 | 0.325431677 | -7.30910791 | 2.69E-13 | 7.07E-12 | 7.07E-11 |
| Rpl35a | 47.411045 | -2.160368241 | 0.310842062 | -7.209662754 | 5.61E-13 | 1.37E-11 | 1.37E-10 |
| Rps12 | 56.39298 | -2.91198652 | 0.426893226 | -7.132270026 | 9.87E-13 | 2.25E-11 | 2.25E-10 |
| Fth1 | 173.70662 | -2.281342461 | 0.340441076 | -6.990093516 | 2.75E-12 | 5.87E-11 | 5.87E-10 |
| Cox6c | 48.043636 | -2.533680219 | 0.385266905 | -6.880586765 | 5.96E-12 | 1.16E-10 | 1.16E-09 |
| Cox7c | 25.93898 | -0.144269504 | 0.849531729 | -6.876789974 | 6.12E-12 | 1.16E-10 | 1.16E-09 |
| Tpt1 | 32.081738 | -2.139470445 | 0.327307058 | -6.801571575 | 1.03E-11 | 1.86E-10 | 1.86E-09 |
| Rpl27a | 41.454174 | -2.334028177 | 0.361990031 | -6.737446026 | 1.61E-11 | 2.68E-10 | 2.68E-09 |
| Rpl19 | 32.01882 | -2.149473623 | 0.332021651 | -6.734582749 | 1.64E-11 | 2.68E-10 | 2.68E-09 |
| Fau | 50.04449 | -2.727079835 | 0.451406353 | -6.386761858 | 1.69E-10 | 2.63E-09 | 2.63E-08 |
| Dbi | 24.794054 | -0.144269504 | 0.849531681 | -6.274232103 | 3.51E-10 | 5.22E-09 | 5.22E-08 |
| Rpl26 | 45.40846 | -2.211559373 | 0.372242843 | -6.254578223 | 3.99E-10 | 5.68E-09 | 5.68E-08 |
| Xlr5a | 113.85682 | 0.759258226 | 0.124615791 | 6.219567299 | 4.99E-10 | 6.82E-09 | 6.82E-08 |
| Plp1 | 14.069005 | -2.835119685 | 0.490485899 | -6.052456856 | 1.43E-09 | 1.88E-08 | 1.88E-07 |
| Uqcrh | 18.11986 | -2.429634213 | 0.424088498 | -6.022445827 | 1.72E-09 | 2.18E-08 | 2.18E-07 |
| Rpl39 | 32.810528 | -2.435814658 | 0.438647763 | -5.904078339 | 3.55E-09 | 4.26E-08 | 4.26E-07 |
| Rps29 | 84.72292 | -1.411250323 | 0.249304903 | -5.9010404 | 3.61E-09 | 4.26E-08 | 4.26E-07 |
| Rps3 | 28.096464 | -1.928526411 | 0.348067217 | -5.839848932 | 5.22E-09 | 5.96E-08 | 5.96E-07 |
| Tmsb4x | 52.412006 | -2.157465215 | 0.393852986 | -5.818305968 | 5.94E-09 | 6.56E-08 | 6.56E-07 |
| Rps27a | 37.066845 | -2.29740611 | 0.421553514 | -5.799768178 | 6.64E-09 | 7.10E-08 | 7.10E-07 |
| Rpl30 | 40.908356 | -2.174013106 | 0.399740632 | -5.782206996 | 7.37E-09 | 7.64E-08 | 7.64E-07 |
| Rpl29 | 18.33695 | -2.101686681 | 0.392874129 | -5.654108157 | 1.57E-08 | 1.58E-07 | 1.58E-06 |
| Gm10062 | 67.03946 | 0.853516567 | 0.156660219 | 5.562101989 | 2.67E-08 | 2.60E-07 | 2.60E-06 |
| Rps14 | 48.404217 | -2.236924796 | 0.436421194 | -5.490106376 | 4.02E-08 | 3.82E-07 | 3.82E-06 |
| Rpl32 | 32.849392 | -1.568497473 | 0.301130199 | -5.484546835 | 4.15E-08 | 3.83E-07 | 3.83E-06 |
| Rps15 | 21.40912 | -1.595383855 | 0.324908946 | -5.199026568 | 2.00E-07 | 1.80E-06 | 1.80E-05 |
| Rpl36 | 24.249557 | -1.963852271 | 0.405724429 | -5.193105509 | 2.07E-07 | 1.80E-06 | 1.80E-05 |
| mt-Nd4 | 68.65068 | -0.144269504 | 0.849531598 | -5.189716973 | 2.11E-07 | 1.80E-06 | 1.80E-05 |
| Mt3 | 44.26073 | -2.371228006 | 0.503219071 | -5.129913857 | 2.90E-07 | 2.42E-06 | 2.42E-05 |
| Rpl23 | 34.328648 | -2.222589572 | 0.48759659 | -4.962297205 | 6.97E-07 | 5.67E-06 | 5.67E-05 |
| Rps25 | 18.493528 | -1.83548764 | 0.408944712 | -4.845125051 | 1.27E-06 | 1.01E-05 | 0.000100637 |
| Rps26 | 26.159721 | -2.347482007 | 0.539736362 | -4.7806203 | 1.75E-06 | 1.36E-05 | 0.000135832 |
| Rsrc1 | 51.52804 | 0.822696597 | 0.177456773 | 4.761645012 | 1.92E-06 | 1.46E-05 | 0.000145936 |
| Galnt13 | 32.602253 | -0.144269504 | 0.849531754 | 4.669691176 | 3.02E-06 | 2.20E-05 | 0.00021984 |
| Rps11 | 30.818487 | -1.903509903 | 0.444968631 | -4.669373325 | 3.02E-06 | 2.20E-05 | 0.00021984 |
| Rpl18a | 28.090158 | -2.216743545 | 0.524866098 | -4.661061545 | 3.15E-06 | 2.24E-05 | 0.00022414 |
| Car12 | 12.034065 | -1.914117644 | 0.448952727 | -4.624937504 | 3.75E-06 | 2.62E-05 | 0.000261533 |
| Cpe | 15.205574 | -1.410858313 | 0.330790834 | -4.556611737 | 5.20E-06 | 3.56E-05 | 0.00035558 |
| Atp5e | 16.749338 | -1.414593385 | 0.335033269 | -4.522457959 | 6.11E-06 | 4.10E-05 | 0.000409901 |
| mt-Co3 | 260.50076 | -0.708598258 | 0.163723752 | -4.451154106 | 8.54E-06 | 5.62E-05 | 0.000561735 |
| Rps3a1 | 35.08009 | -1.729887324 | 0.429522135 | -4.423230072 | 9.72E-06 | 6.27E-05 | 0.000627448 |
| Eef1a1 | 21.101805 | -1.877010492 | 0.470098918 | -4.408444727 | 1.04E-05 | 6.59E-05 | 0.000659399 |
| Rps21 | 92.48 | -1.585293011 | 0.403681998 | -4.311910236 | 1.62E-05 | 0.000100641 | 0.001006414 |
| Rps23 | 30.271101 | -1.581671332 | 0.403033626 | -4.304340594 | 1.67E-05 | 0.000102285 | 0.001022848 |
| Mc1r | 43.78233 | 0.889585995 | 0.216615803 | 4.266736732 | 1.98E-05 | 0.000119012 | 0.001190118 |
| mt-Atp6 | 196.57178 | -0.474270063 | 0.11325205 | -4.25551972 | 2.09E-05 | 0.000122981 | 0.001229808 |
| Rpl12 | 13.173615 | -2.386184734 | 0.636143362 | -4.237809572 | 2.26E-05 | 0.000130836 | 0.001308359 |
| Rpl11 | 38.71404 | -1.569351019 | 0.41981618 | -4.132768389 | 3.58E-05 | 0.000204299 | 0.002042993 |
| Rps16 | 30.492613 | -1.482768965 | 0.398901284 | -4.091833876 | 4.28E-05 | 0.000239947 | 0.002399468 |
| Mbp | 41.059708 | -2.032580335 | 0.571661668 | -4.062415246 | 4.86E-05 | 0.000266181 | 0.002661809 |
| Rps13 | 22.769087 | -1.512504066 | 0.411952515 | -4.060187489 | 4.90E-05 | 0.000266181 | 0.002661809 |
| Cmpk1 | 54.735962 | 0.738783283 | 0.188634768 | 4.054421774 | 5.03E-05 | 0.000268569 | 0.002685687 |
| Rpl41 | 50.65451 | -1.488532196 | 0.408277087 | -4.032382637 | 5.52E-05 | 0.000290512 | 0.002905116 |
| Rps7 | 29.148432 | -2.035637906 | 0.584822215 | -3.993433222 | 6.51E-05 | 0.000337458 | 0.003374577 |
| Rpl38 | 44.777405 | -1.610557287 | 0.454791285 | -3.970244347 | 7.18E-05 | 0.000366496 | 0.003664962 |

| | | | | | | | |
|---|---|---|---|---|---|---|---|
| Cox6b1 | 17.059458 | -2.665946381 | 0.8298069 | -3.840384034 | 0.000122842 | 0.000617823 | 0.00617823 |
| Rpl37a | 48.63827 | -2.030413475 | 0.618279044 | -3.83632121 | 0.000124891 | 0.000619025 | 0.006190252 |
| mt-Co1 | 134.72978 | -0.803769617 | 0.221115292 | -3.819091031 | 0.000133944 | 0.000654414 | 0.006544137 |
| Rpl21 | 43.89273 | -1.868326021 | 0.570770251 | -3.796473807 | 0.000146769 | 0.000706971 | 0.007069711 |
| Selenow | 21.611269 | -1.702435674 | 0.515250725 | -3.782686962 | 0.000155144 | 0.000736936 | 0.007369363 |
| Aldoc | 16.357277 | -1.83802411 | 0.567423712 | -3.739971104 | 0.000184041 | 0.000862221 | 0.008622214 |
| Epha6 | 42.884907 | 0.69503836 | 0.194441052 | 3.680827787 | 0.000232478 | 0.001074425 | 0.010744253 |
| Rpl34 | 28.263384 | -1.374613858 | 0.419769877 | -3.672979644 | 0.000239739 | 0.001093208 | 0.01093208 |
| Atp9b | 26.321203 | -0.144269504 | 0.849531767 | 3.617811643 | 0.000297104 | 0.00133697 | 0.013369702 |
| Sil1 | 28.911364 | 0.812336143 | 0.242369642 | 3.53007475 | 0.000415442 | 0.001845211 | 0.018452111 |
| mt-Co2 | 153.16696 | -0.6899307 | 0.207555934 | -3.482618469 | 0.000496535 | 0.002177117 | 0.02177117 |
| Nckap5 | 111.1208 | 0.644876075 | 0.19639548 | 3.425407551 | 0.000613878 | 0.002657547 | 0.02657547 |
| Rpl9 | 30.104624 | -2.031350497 | 0.73735968 | -3.408169867 | 0.000654002 | 0.002795857 | 0.027958572 |
| Hapln1 | 16.739304 | -0.144269504 | 0.849531793 | 3.392522492 | 0.000692522 | 0.002888325 | 0.028883254 |
| Scamp1 | 20.171698 | -0.144269504 | 0.84953178 | 3.394107593 | 0.000688526 | 0.002888325 | 0.028883254 |
| 4930511M | 45.435577 | -0.144269504 | 0.849531705 | 3.374701783 | 0.000738957 | 0.00304486 | 0.030448598 |
| Acoxl | 22.274984 | -0.144269504 | 0.849531742 | 3.370295673 | 0.000750876 | 0.003057137 | 0.030571367 |
| Zbtb7a | 26.736818 | -0.144269504 | 0.849531762 | 3.22554375 | 0.001257336 | 0.005048522 | 0.050485224 |
| Rpl8 | 19.731731 | -2.048418721 | 0.8176993 | -3.222784221 | 0.001269511 | 0.005048522 | 0.050485224 |
| Trim2 | 47.792538 | 0.684477228 | 0.225467986 | 3.197269206 | 0.001387354 | 0.005453736 | 0.054537357 |
| Rps15a | 16.681158 | -1.324731611 | 0.48561248 | -3.178370997 | 0.001481051 | 0.005755903 | 0.057559031 |
| Dpyd | 17.710781 | -0.144269504 | 0.84953176 | 3.174897151 | 0.001498896 | 0.005759803 | 0.057598033 |
| Mt1 | 41.963573 | -1.490444346 | 0.577640298 | -3.133204317 | 0.00172909 | 0.006570543 | 0.065705433 |
| Vit | 27.375063 | 0.857319642 | 0.311300441 | 3.00344037 | 0.002669459 | 0.010032471 | 0.100324706 |
| Lrp1b | 234.25195 | 0.506582019 | 0.17557482 | 2.988265147 | 0.00280566 | 0.010429737 | 0.104297374 |
| Atp5md | 17.48183 | -1.137313418 | 0.441148034 | -2.983622114 | 0.002848583 | 0.010475436 | 0.104754358 |
| Rpl24 | 27.844671 | -1.901799438 | 0.869742762 | -2.977753939 | 0.00290369 | 0.010564489 | 0.10564489 |
| Hdac8 | 16.366652 | 0.872542669 | 0.318937454 | 2.96887141 | 0.002988956 | 0.010760243 | 0.107602433 |
| Ubb | 13.961383 | -1.230261293 | 0.491115915 | -2.963097929 | 0.003045597 | 0.010849938 | 0.108499378 |
| mt-Nd4l | 7.8132463 | -0.144269504 | 0.849531757 | -2.935842422 | 0.003326433 | 0.011728247 | 0.117282466 |
| Grik1 | 7.980826 | 1.552217242 | 0.62546964 | 2.931258995 | 0.003375912 | 0.011781244 | 0.117812439 |
| Ntm | 65.63441 | 0.786980592 | 0.295635676 | 2.898739549 | 0.00374666 | 0.012943007 | 0.129430066 |
| Rubcnl | 14.1022215 | -0.144269504 | 0.849531797 | 2.871714938 | 0.00408251 | 0.013962186 | 0.139621858 |
| mt-Nd2 | 89.92444 | -0.583787923 | 0.214683052 | -2.867715237 | 0.004134475 | 0.013999905 | 0.139999045 |
| Prkg1 | 40.345478 | -0.144269504 | 0.849531737 | 2.848914059 | 0.004386873 | 0.014708927 | 0.147089267 |
| Ms4a1 | 33.088974 | 0.74324921 | 0.286112695 | 2.819622748 | 0.004808014 | 0.015935101 | 0.159351005 |
| Cntnap2 | 61.76419 | -0.144269504 | 0.849531692 | 2.817112058 | 0.004845762 | 0.015935101 | 0.159351005 |
| Sec61a1 | 209.65936 | -0.144269504 | 0.84953137 | -2.790653146 | 0.005260181 | 0.017133161 | 0.171331606 |
| Gm12216 | 111.9227 | -0.144269504 | 0.849531643 | 2.770579288 | 0.005595667 | 0.018053946 | 0.18053946 |
| 4930441J1 | 16.806175 | -0.144269504 | 0.849531775 | 2.750461915 | 0.005951131 | 0.019021372 | 0.190213717 |
| Cfap74 | 19.824886 | -0.844595298 | 0.345978234 | -2.735426771 | 0.006229945 | 0.019728158 | 0.19728158 |
| Gm50306 | 36.185795 | -0.144269504 | 0.849531709 | 2.687699717 | 0.007194606 | 0.022573903 | 0.225739025 |
| Hexb | 11.284686 | -1.488174962 | 0.751077883 | -2.683850043 | 0.007277974 | 0.022627883 | 0.226278835 |
| Rapgef4 | 58.898792 | 0.619428704 | 0.248296168 | 2.667176724 | 0.007649144 | 0.023357207 | 0.23357207 |
| Rps17 | 12.254832 | -1.00344809 | 0.44201592 | -2.66718316 | 0.007648997 | 0.023357207 | 0.23357207 |
| Calb1 | 9.242661 | -0.144269504 | 0.849531751 | -2.641287475 | 0.008259161 | 0.024996752 | 0.249967516 |
| Ppia | 15.149974 | -0.908533184 | 0.396654754 | -2.635977035 | 0.008389541 | 0.025168623 | 0.251686228 |
| Meg3 | 17.646185 | 1.054435336 | 0.47736932 | 2.629606719 | 0.00854837 | 0.025422108 | 0.25422108 |
| 8430422H( | 51.415527 | 0.572567018 | 0.233439813 | 2.608180795 | 0.009102486 | 0.026607268 | 0.266072678 |
| Igf2bp3 | 55.370945 | 0.591061066 | 0.241357325 | 2.610127549 | 0.009050847 | 0.026607268 | 0.266072678 |
| 4930558K( | 20.802944 | -0.144269504 | 0.849531777 | 2.56654646 | 0.010271686 | 0.029770481 | 0.297704809 |
| Rpl13a | 25.23023 | -1.489193205 | 0.87819328 | -2.534649789 | 0.011255979 | 0.032349117 | 0.323491171 |
| Usp3 | 11.741427 | 0.966361425 | 0.452843819 | 2.50056788 | 0.012399437 | 0.035046342 | 0.350463422 |
| Nkain2 | 21.204378 | -0.144269504 | 0.84953177 | 2.503093413 | 0.012311305 | 0.035046342 | 0.350463422 |
| Wdr72 | 27.201456 | 0.614531279 | 0.265776302 | 2.492615166 | 0.01268062 | 0.035547312 | 0.355473118 |
| Pcsk2 | 11.427464 | 1.021300839 | 0.490198912 | 2.488750548 | 0.012819288 | 0.035643873 | 0.356438728 |
| Gria4 | 32.433834 | 0.638343717 | 0.282406426 | 2.458522215 | 0.013951014 | 0.038290676 | 0.382906763 |
| Cdk8 | 11.849927 | -0.144269504 | 0.84953179 | 2.457388359 | 0.01399513 | 0.038290676 | 0.382906763 |
| Lsamp | 29.747162 | 0.528402582 | 0.228882689 | 2.44835281 | 0.014351105 | 0.038953 | 0.38953 |
| Svil | 13.7415695 | -0.144269504 | 0.849531745 | 2.436203328 | 0.014842339 | 0.039969133 | 0.399691331 |
| Gm49497 | 15.725553 | 0.834520978 | 0.393754419 | 2.430752579 | 0.0150675 | 0.040258476 | 0.402584764 |
| Tcea1 | 16.892315 | -0.144269504 | 0.849531761 | -2.408858323 | 0.016002509 | 0.042425256 | 0.424252558 |
| Nell1 | 26.662558 | 0.695590358 | 0.323794828 | 2.389572657 | 0.016867987 | 0.044375781 | 0.443757813 |
| Kctd16 | 87.87623 | 0.498290581 | 0.221328166 | 2.380426765 | 0.017292598 | 0.045145561 | 0.451455611 |
| Chchd2 | 13.155441 | -1.085660102 | 0.593593613 | -2.377003836 | 0.017453907 | 0.045221485 | 0.452214853 |
| Opcml | 9.554732 | -0.144269504 | 0.849531789 | 2.358636802 | 0.018342198 | 0.047165651 | 0.471656514 |
| Dop1b | 24.85656 | 0.687364375 | 0.327242241 | 2.342001395 | 0.019180643 | 0.048953582 | 0.48953582 |
| Tns3 | 13.053523 | 0.923454679 | 0.479854604 | 2.326421393 | 0.019996081 | 0.05065674 | 0.506567397 |
| Ctnna2 | 15.505933 | 0.844078398 | 0.443563211 | 2.265163933 | 0.023502622 | 0.059102181 | 0.591021805 |

| | | | | | | | |
|---|---|---|---|---|---|---|---|
| mt-Nd3 | 47.00168 | -0.696586088 | 0.369171147 | -2.174268957 | 0.029684942 | 0.074104017 | 0.741040165 |
| Adk | 16.03901 | -0.144269504 | 0.849531786 | 2.161509718 | 0.030655987 | 0.075973533 | 0.759735333 |
| Tmem108 | 20.846542 | 0.631728698 | 0.328872867 | 2.150659359 | 0.031503096 | 0.076957564 | 0.76957564 |
| Gas5 | 16.921923 | -0.833617123 | 0.477512949 | -2.150791177 | 0.031492686 | 0.076957564 | 0.76957564 |
| Gm42418 | 376.12793 | -0.392360256 | 0.191056138 | -2.147396849 | 0.031761701 | 0.07703902 | 0.770390202 |
| Naa11 | 7.6849 | 0.908415712 | 0.53173167 | 2.128997019 | 0.033254507 | 0.080091841 | 0.80091841 |
| Rpl31 | 14.029416 | -0.981554419 | 0.643967325 | -2.11175814 | 0.034707199 | 0.083006029 | 0.830060292 |
| Pzp | 18.460627 | -0.650341353 | 0.353132403 | -2.104425727 | 0.035341327 | 0.083935652 | 0.839356521 |
| Stard7 | 12.632127 | 0.718339523 | 0.411449596 | 2.051575528 | 0.040210932 | 0.094842337 | 0.948423373 |
| mt-Nd5 | 15.932015 | -0.683708171 | 0.392294189 | -2.042287481 | 0.041123023 | 0.096329273 | 0.963292734 |
| Cox7a2 | 21.137936 | -0.917026351 | 0.631839409 | -2.020032373 | 0.04338003 | 0.100924967 | 1.00924967 |
| Ctcflos | 21.047422 | -0.144269504 | 0.849531762 | 2.004380244 | 0.045029344 | 0.104054295 | 1.040542949 |
| Prkag2 | 21.88738 | -0.144269504 | 0.849531737 | 1.992087636 | 0.046361443 | 0.106413514 | 1.064135142 |
| Rps28 | 32.55888 | -0.771093845 | 0.521454025 | -1.912229361 | 0.055846782 | 0.127330662 | 1.273306622 |
| mt-Cytb | 80.49536 | -0.391301056 | 0.219088584 | -1.896550624 | 0.057887273 | 0.131108922 | 1.311089219 |
| Hspa14 | 9.755078 | -0.144269504 | 0.849531762 | 1.878854415 | 0.060264375 | 0.135594843 | 1.355948428 |
| Gm26901 | 7.0144696 | -0.144269504 | 0.849531774 | -1.854877946 | 0.063613658 | 0.142195236 | 1.421952356 |
| Gm14582 | 15.299492 | 0.599964804 | 0.378743926 | 1.843528422 | 0.065251896 | 0.144910055 | 1.449100546 |
| Slx1b | 31.099352 | 0.501077598 | 0.303413019 | 1.83484156 | 0.066529169 | 0.145852408 | 1.458524083 |
| Sema3c | 3.6835606 | 0.981787902 | 0.878091375 | 1.835467643 | 0.066436429 | 0.145852408 | 1.458524083 |
| Adamtsl1 | 1.6549803 | -0.144269504 | 0.849531787 | 1.827676299 | 0.067598147 | 0.147252015 | 1.472520149 |
| Pinlyp | 6.176205 | -0.696486631 | 0.483711068 | -1.818367117 | 0.069008041 | 0.147504687 | 1.47504687 |
| Il27 | 36.25184 | 0.500230954 | 0.305635261 | 1.820377776 | 0.068701494 | 0.147504687 | 1.47504687 |
| Fbxl7 | 19.16149 | 0.697751408 | 0.481463528 | 1.823498446 | 0.068227932 | 0.147504687 | 1.47504687 |
| 6030443J0 | 38.516613 | 0.466972184 | 0.28253827 | 1.811979996 | 0.069989287 | 0.148672896 | 1.486728963 |
| Gm11337 | 15.23903 | 0.604394454 | 0.39450304 | 1.803481713 | 0.071312595 | 0.150548811 | 1.505488113 |
| Alpl | 12.030566 | -0.144269504 | 0.84953178 | 1.799022848 | 0.072015067 | 0.151099097 | 1.510990967 |
| Trim33 | 10.494772 | -0.144269504 | 0.849531736 | -1.785560773 | 0.074170403 | 0.154672426 | 1.546724259 |
| Calm1 | 19.41618 | -0.671438343 | 0.476559064 | -1.774745701 | 0.075939886 | 0.157402672 | 1.574026721 |
| Gm13219 | 7.1886063 | -0.677116553 | 0.493671347 | -1.751547015 | 0.07985173 | 0.164513806 | 1.645138057 |
| Kcnh7 | 7.5752106 | 0.71512764 | 0.53444845 | 1.7398116 | 0.081892105 | 0.167707185 | 1.67707185 |
| Uba52 | 7.7806873 | -0.721917406 | 0.567748094 | -1.73010501 | 0.083611515 | 0.169202002 | 1.692020015 |
| Gnaz | 52.612423 | 0.343360762 | 0.209153037 | 1.732669693 | 0.083154394 | 0.169202002 | 1.692020015 |
| mt-Atp8 | 7.16448 | -0.144269504 | 0.849531753 | -1.723837475 | 0.084737187 | 0.170471283 | 1.704712827 |
| 4931429P1 | 6.027571 | -0.144269504 | 0.84953177 | -1.717283267 | 0.085927428 | 0.171854856 | 1.718548556 |
| Ninj2 | 21.596132 | -0.144269504 | 0.849531742 | 1.714041797 | 0.08652105 | 0.172036041 | 1.720360411 |
| Pde1c | 58.276245 | 0.350500215 | 0.218481632 | 1.701477579 | 0.088853344 | 0.175652276 | 1.756522763 |
| Vwc2l | 1.6346 | 1.039143835 | 1.192605325 | 1.686457025 | 0.091707806 | 0.180253273 | 1.802532734 |
| Pcdh15 | 32.498318 | -0.144269504 | 0.849531666 | 1.682719942 | 0.092429319 | 0.180633299 | 1.806332986 |
| Wdyhv1 | 20.146286 | -0.144269504 | 0.849531722 | 1.667859991 | 0.095343524 | 0.185269801 | 1.852698014 |
| Srbd1 | 7.4579577 | -0.144269504 | 0.849531751 | 1.643010584 | 0.100380744 | 0.193956013 | 1.939560129 |
| Cmss1 | 38.949615 | -0.406165804 | 0.271828395 | -1.636714564 | 0.101690124 | 0.195382149 | 1.953821491 |
| Asxl3 | 11.312686 | -0.144269504 | 0.84953177 | 1.596046608 | 0.110478388 | 0.211081613 | 2.110816127 |
| Chsy3 | 27.832537 | -0.144269504 | 0.849531758 | 1.586661042 | 0.112589434 | 0.213919925 | 2.139199252 |
| Pkp2 | 12.1512985 | -0.144269504 | 0.849531789 | 1.575661575 | 0.115103817 | 0.217488981 | 2.174889807 |
| Rps27 | 23.897285 | -0.529109175 | 0.405584354 | -1.566605521 | 0.117206927 | 0.220245983 | 2.202459829 |
| Sdccag8 | 9.023283 | 0.624836618 | 0.529971108 | 1.559978731 | 0.118764908 | 0.220747817 | 2.207478173 |
| Ckb | 24.59765 | -0.636413746 | 0.556153604 | -1.561799023 | 0.118335342 | 0.220747817 | 2.207478173 |
| Hdac9 | 10.124961 | 0.55426585 | 0.441728973 | 1.543848742 | 0.12262498 | 0.225471738 | 2.254717376 |
| Arvcf | 6.9049354 | -0.144269504 | 0.849531782 | -1.544283787 | 0.122519599 | 0.225471738 | 2.254717376 |
| mt-Nd1 | 55.00545 | -0.389745163 | 0.279225034 | -1.537695369 | 0.124123116 | 0.227005914 | 2.270059135 |
| Mapk14 | 50.016357 | 0.328374251 | 0.230855459 | 1.520831128 | 0.128302221 | 0.232984039 | 2.329840394 |
| Kcnip4 | 20.59791 | 0.530493681 | 0.427258456 | 1.519032412 | 0.128754338 | 0.232984039 | 2.329840394 |
| Zfhx3 | 17.124788 | 0.448449594 | 0.34914624 | 1.478498644 | 0.13927436 | 0.250693847 | 2.506938474 |
| Atxn7l1 | 13.80564 | 0.511765123 | 0.428470048 | 1.463255157 | 0.143397573 | 0.255487562 | 2.554875622 |
| Lyrm1 | 8.578427 | 0.557782072 | 0.489373645 | 1.463130718 | 0.143431614 | 0.255487562 | 2.554875622 |
| Spock1 | 16.135859 | 0.44815869 | 0.357624159 | 1.451672513 | 0.146592686 | 0.258426281 | 2.584262811 |
| Chchd6 | 9.167513 | 0.5535332 | 0.489609329 | 1.453482337 | 0.14608988 | 0.258426281 | 2.584262811 |
| Slco2a1 | 10.997745 | -0.144269504 | 0.849531758 | 1.446632734 | 0.147999813 | 0.259568903 | 2.595689032 |
| Fgf12 | 10.426932 | 0.506658056 | 0.444635426 | 1.412811156 | 0.157711257 | 0.275190051 | 2.751900507 |
| Gm2895 | 7.383854 | 0.543155237 | 0.51856786 | 1.384461049 | 0.166217327 | 0.28856003 | 2.885600296 |
| Pde6a | 68.24931 | 0.290176179 | 0.226231122 | 1.367075878 | 0.171601523 | 0.296402631 | 2.96402631 |
| Cldn16 | 12.688222 | 0.455614372 | 0.40478461 | 1.356130086 | 0.175057782 | 0.300853073 | 3.008530725 |
| Ehbp1 | 44.990513 | 0.290173825 | 0.229401683 | 1.351150661 | 0.176647176 | 0.302066671 | 3.020666708 |
| Ap3s1 | 13.637611 | -0.144269504 | 0.84953177 | 1.337026495 | 0.181213989 | 0.308334249 | 3.083342492 |
| Ccdc60 | 15.207973 | 0.439587261 | 0.398884522 | 1.324720163 | 0.185263983 | 0.313664763 | 3.136647634 |
| Ptprd | 15.91118 | 0.451462482 | 0.42081575 | 1.312322622 | 0.189411302 | 0.319106726 | 3.191067264 |
| Rgs6 | 1.1746267 | 0.60470146 | 1.104306744 | 1.301951622 | 0.192932924 | 0.323446373 | 3.234463727 |
| Stxbp5l | 17.244005 | 0.397965498 | 0.361173519 | 1.284923424 | 0.198819044 | 0.331688357 | 3.316883571 |

| | | | | | | | |
|---|---|---|---|---|---|---|---|
| Kif5a | 6.6975093 | -0.459294545 | 0.456215968 | -1.281952593 | 0.199859277 | 0.331805207 | 3.318052072 |
| Grid2 | 12.530739 | -0.144269504 | 0.849531764 | 1.273026506 | 0.203008636 | 0.335405573 | 3.354055728 |
| Zmat4 | 27.439436 | 0.329000971 | 0.289796427 | 1.259862959 | 0.207718803 | 0.339903496 | 3.399034956 |
| Klhl14 | 2.0752926 | -0.144269504 | 0.849531817 | 1.260837015 | 0.207367575 | 0.339903496 | 3.399034956 |
| Lrrc17 | 30.266716 | -0.354114982 | 0.321704944 | -1.250037416 | 0.21128588 | 0.344094147 | 3.440941471 |
| Gm28729 | 5.518057 | -0.144269504 | 0.849531767 | 1.241039214 | 0.214591263 | 0.346730745 | 3.467307451 |
| Gm16046 | 6.2665734 | -0.474441056 | 0.515728051 | -1.239325842 | 0.215224852 | 0.346730745 | 3.467307451 |
| Nrg3 | 10.689915 | 0.438369303 | 0.443491861 | 1.237379189 | 0.215946341 | 0.346730745 | 3.467307451 |
| Dctd | 9.343067 | -0.144269504 | 0.849531782 | 1.232984477 | 0.217581556 | 0.347723795 | 3.477237946 |
| Klkb1 | 14.826636 | 0.343202527 | 0.315452002 | 1.228442691 | 0.21928083 | 0.348809507 | 3.488095069 |
| Csmd3 | 32.346813 | 0.308046357 | 0.27793136 | 1.219381557 | 0.222699407 | 0.352607394 | 3.526073939 |
| Tshz2 | 3.178899 | 0.530667328 | 0.740735639 | 1.210015001 | 0.226273137 | 0.355168105 | 3.551681045 |
| Gm765 | 14.447982 | 0.395101478 | 0.392817716 | 1.207149892 | 0.227374424 | 0.355168105 | 3.551681045 |
| Gm20342 | 10.802338 | 0.459376034 | 0.504373156 | 1.206999836 | 0.227432207 | 0.355168105 | 3.551681045 |
| Srgn | 6.8719287 | -0.144269504 | 0.849531797 | 1.192870383 | 0.23292014 | 0.361560896 | 3.615608963 |
| C330002G( | 6.980444 | 0.439563151 | 0.473980796 | 1.191034024 | 0.233640228 | 0.361560896 | 3.615608963 |
| Micu1 | 6.7083936 | -0.43932142 | 0.49774982 | -1.173426884 | 0.240624662 | 0.370346687 | 3.703466865 |
| Shb | 68.3498 | 0.220769258 | 0.19797767 | 1.171287172 | 0.241483366 | 0.370346687 | 3.703466865 |
| Tacc1 | 13.956649 | -0.312794829 | 0.305703048 | -1.159584794 | 0.246217895 | 0.370953833 | 3.709538329 |
| Topaz1 | 8.223777 | -0.144269504 | 0.849531761 | 1.162468967 | 0.245045023 | 0.370953833 | 3.709538329 |
| Unc5d | 1.4896435 | 0.520997789 | 0.896371002 | 1.164886481 | 0.244064946 | 0.370953833 | 3.709538329 |
| Klf10 | 8.785145 | 0.400556464 | 0.425292672 | 1.161259791 | 0.245536266 | 0.370953833 | 3.709538329 |
| Cdh23 | 37.642277 | 0.254841995 | 0.235785051 | 1.155075374 | 0.248059562 | 0.372089343 | 3.720893426 |
| Asic2 | 18.594269 | -0.324557437 | 0.322399295 | -1.148186916 | 0.250891409 | 0.3733405 | 3.733405004 |
| Cspp1 | 20.2637 | 0.343712979 | 0.346178223 | 1.147737502 | 0.251076945 | 0.3733405 | 3.733405004 |
| 4930402H2 | 11.330419 | -0.144269504 | 0.849531772 | 1.128801513 | 0.258981574 | 0.383427266 | 3.834272661 |
| Il1rapl2 | 10.690964 | 0.366357444 | 0.394915512 | 1.112842726 | 0.265775982 | 0.391790456 | 3.91790456 |
| Sorcs2 | 11.712841 | 0.377156463 | 0.422490569 | 1.101289011 | 0.270770892 | 0.395742073 | 3.957420735 |
| AY036118 | 49.08356 | -0.226977667 | 0.219369089 | -1.102485037 | 0.27025086 | 0.395742073 | 3.957420735 |
| Bc1 | 22.944368 | -0.375938234 | 0.428693188 | -1.091601455 | 0.275008316 | 0.400224869 | 4.002248689 |
| 1700054A( | 174.88585 | 0.112521831 | 0.108678635 | 1.054732421 | 0.29154772 | 0.422497119 | 4.224971192 |
| Ttc28 | 32.3879 | 0.272194545 | 0.288209155 | 1.047156625 | 0.295027347 | 0.425735665 | 4.25735665 |
| Celf2 | 12.806203 | 0.418205538 | 0.597659321 | 1.040440961 | 0.29813508 | 0.428412594 | 4.284125936 |
| Esf1 | 10.965066 | 0.378828389 | 0.474911639 | 1.037512048 | 0.299497284 | 0.428569335 | 4.28569335 |
| Kcnd2 | 7.7279754 | 0.37330459 | 0.469861047 | 1.026129959 | 0.30483033 | 0.432580801 | 4.325808009 |
| Pard3b | 7.5184894 | 0.391798619 | 0.520089876 | 1.026528928 | 0.304642335 | 0.432580801 | 4.325808009 |
| Malat1 | 63.65539 | 0.324667335 | 0.383441232 | 1.008982361 | 0.312983094 | 0.442314951 | 4.423149507 |
| Pcgf6 | 14.770329 | -0.144269504 | 0.849531774 | 0.983300412 | 0.325459607 | 0.458054262 | 4.58054262 |
| Nrp2 | 8.941567 | -0.144269504 | 0.849531746 | 0.975691885 | 0.329217175 | 0.459634572 | 4.596345719 |
| Slc24a3 | 12.337384 | 0.325342296 | 0.406872447 | 0.975584557 | 0.32927038 | 0.459634572 | 4.596345719 |
| Lin28b | 10.255604 | -0.144269504 | 0.849531765 | 0.971780085 | 0.331159965 | 0.460393122 | 4.603931223 |
| Gm29325 | 19.200335 | -0.144269504 | 0.849531764 | 0.949724286 | 0.342252366 | 0.470081563 | 4.70081563 |
| D430042O | 64.132515 | 0.218782382 | 0.248354359 | 0.95134404 | 0.341429758 | 0.470081563 | 4.70081563 |
| Ppp1r16a | 19.354738 | 0.276519954 | 0.332401497 | 0.955059129 | 0.339547796 | 0.470081563 | 4.70081563 |
| Ptprg | 10.517569 | 0.322059034 | 0.432337748 | 0.932019537 | 0.351326429 | 0.480614555 | 4.80614555 |
| Cadps2 | 5.6570077 | -0.144269504 | 0.849531761 | 0.910030344 | 0.362806507 | 0.49238026 | 4.923802601 |
| Ppp1r9a | 20.427477 | 0.317642949 | 0.439526835 | 0.910324648 | 0.362651323 | 0.49238026 | 4.923802601 |
| Rab29 | 21.145254 | 0.22160641 | 0.274434984 | 0.889956158 | 0.373489428 | 0.501837534 | 5.018375344 |
| Tsacc | 4.575702 | -0.35267301 | 0.609643799 | -0.889528838 | 0.373718932 | 0.501837534 | 5.018375344 |
| Cox11 | 7.8023853 | -0.144269504 | 0.849531787 | 0.888676231 | 0.374177109 | 0.501837534 | 5.018375344 |
| Nup210l | 8.675225 | -0.293534318 | 0.440155215 | -0.848997363 | 0.39588276 | 0.528874625 | 5.288746245 |
| Gm960 | 10.207831 | 0.283302693 | 0.420881798 | 0.835086038 | 0.403669267 | 0.537178558 | 5.371785578 |
| Cntnap5b | 4.7811284 | -0.144269504 | 0.849531753 | -0.825549284 | 0.409059815 | 0.54224208 | 5.422420799 |
| Tpd52l1 | 10.871926 | -0.144269504 | 0.849531783 | 0.810088302 | 0.417889427 | 0.551807661 | 5.518076606 |
| 9130015L2 | 5.0294023 | -0.144269504 | 0.849531781 | 0.805132797 | 0.420743052 | 0.553438937 | 5.534389375 |
| Tatdn1 | 9.639689 | -0.144269504 | 0.849531756 | -0.798122964 | 0.424799137 | 0.556633351 | 5.566333513 |
| Fam189a2 | 7.773994 | -0.144269504 | 0.849531777 | 0.793784093 | 0.427321134 | 0.55780087 | 5.578008701 |
| Lrrc4c | 15.353004 | -0.144269504 | 0.849531782 | 0.780396923 | 0.435157279 | 0.563726475 | 5.637264745 |
| Ctnna3 | 32.19501 | -0.144269504 | 0.849531677 | 0.780995316 | 0.43480525 | 0.563726475 | 5.637264745 |
| Ano2 | 13.100007 | 0.254332701 | 0.400876403 | 0.774811531 | 0.438451035 | 0.565850014 | 5.658500144 |
| Camk1d | 35.845703 | -0.193484871 | 0.278412692 | -0.768499828 | 0.442190293 | 0.56640105 | 5.664010496 |
| Brip1os | 7.3789635 | -0.298224033 | 0.574702266 | -0.770910643 | 0.440759898 | 0.56640105 | 5.664010496 |
| Sult2a2 | 11.174783 | -0.144269504 | 0.849531755 | 0.732593845 | 0.463806186 | 0.591872074 | 5.918720736 |
| Elf1 | 10.401972 | -0.144269504 | 0.849531777 | 0.722648916 | 0.469895607 | 0.597413746 | 5.974137463 |
| Zfpm2 | 1.6973265 | 0.275928288 | 0.835593387 | 0.705872372 | 0.480267511 | 0.608338847 | 6.083388466 |
| Gm38973 | 20.448889 | -0.173057229 | 0.274080974 | -0.699906711 | 0.483985566 | 0.610786213 | 6.107862127 |
| A630012P( | 0.3618243 | 0.314545014 | 1.325412169 | 0.687623034 | 0.491690199 | 0.615963546 | 6.159635465 |
| Gm42791 | 4.867136 | -0.144269504 | 0.8495318 | 0.687675593 | 0.491657093 | 0.615963546 | 6.159635465 |
| Phkg1 | 17.80475 | -0.144269504 | 0.849531718 | 0.681947033 | 0.495272444 | 0.618186773 | 6.181867732 |

| | | | | | | | |
|---|---|---|---|---|---|---|---|
| Ralyl | 10.461764 | 0.228576696 | 0.42418319 | 0.673406651 | 0.500688618 | 0.622674573 | 6.226745726 |
| Sgip1 | 5.9846826 | -0.255115402 | 0.573975854 | -0.66408511 | 0.50663584 | 0.626147664 | 6.261476642 |
| 1700126G( | 40.101994 | 0.181682618 | 0.310640994 | 0.66043638 | 0.508973832 | 0.626147664 | 6.261476642 |
| Lingo2 | 7.67656 | 0.239107176 | 0.478467633 | 0.661450625 | 0.508323369 | 0.626147664 | 6.261476642 |
| Anks1b | 7.6343727 | -0.238729217 | 0.491453433 | -0.656723161 | 0.511358939 | 0.626827087 | 6.26827087 |
| Gm17529 | 10.398543 | 0.21569988 | 0.437467821 | 0.624592667 | 0.532238434 | 0.65009123 | 6.500912299 |
| A330008L1 | 12.58962 | 0.202403879 | 0.402771114 | 0.615216747 | 0.53841159 | 0.655290974 | 6.55290974 |
| Kat7 | 16.726942 | 0.200884936 | 0.403451082 | 0.608589954 | 0.542796265 | 0.658284832 | 6.582848319 |
| Il23a | 8.529622 | 0.21349615 | 0.460032074 | 0.60268291 | 0.546719653 | 0.660700075 | 6.607000753 |
| Ppp3ca | 8.880681 | -0.209640815 | 0.487685681 | -0.579590179 | 0.562191018 | 0.677004676 | 6.77004676 |
| Ptpn9 | 20.518375 | 0.159806327 | 0.31601228 | 0.576389253 | 0.564352107 | 0.677222528 | 6.772225279 |
| Olfr57 | 7.610871 | 0.200571977 | 0.526208225 | 0.533994488 | 0.593345347 | 0.707305991 | 7.073059907 |
| Zfp994 | 19.507566 | 0.148401811 | 0.313936049 | 0.533687222 | 0.593557951 | 0.707305991 | 7.073059907 |
| A930016O | 8.856556 | 0.182052433 | 0.449747922 | 0.520954595 | 0.602398402 | 0.715348102 | 7.153481018 |
| Tmeff2 | 11.775853 | -0.144269504 | 0.849531778 | 0.502594841 | 0.615249158 | 0.728080318 | 7.280803179 |
| Gphn | 10.085578 | -0.144269504 | 0.849531763 | -0.498592696 | 0.618066352 | 0.728892043 | 7.288920428 |
| Sh3bgrl2 | 6.459438 | -0.144269504 | 0.849531775 | 0.478769159 | 0.63210286 | 0.742883773 | 7.428837731 |
| Rtkn | 5.6617584 | 0.178215105 | 0.538345691 | 0.472134214 | 0.636830992 | 0.743331739 | 7.433317386 |
| Gnao1 | 10.356858 | -0.144269504 | 0.849531765 | 0.472710522 | 0.636419719 | 0.743331739 | 7.433317386 |
| Col5a2 | 9.190492 | -0.144269504 | 0.84953177 | -0.466036813 | 0.64118915 | 0.745873093 | 7.458730933 |
| Ahdc1 | 12.658006 | 0.126699203 | 0.329566773 | 0.435755572 | 0.663014081 | 0.768646833 | 7.686468331 |
| Alk | 7.22739 | -0.144269504 | 0.84953177 | -0.432277824 | 0.665539502 | 0.768967939 | 7.689679386 |
| Camk2a | 20.487011 | 0.120633611 | 0.355519384 | 0.396893756 | 0.691445809 | 0.796210325 | 7.962103253 |
| Pecam1 | 6.8732324 | 0.133441522 | 0.435965031 | 0.376236384 | 0.706741168 | 0.811092213 | 8.110922133 |
| Tshr | 9.432204 | -0.144269504 | 0.849531771 | 0.360065081 | 0.718798465 | 0.81400356 | 8.140035602 |
| Chl1 | 17.029634 | 0.114693301 | 0.382003962 | 0.360411653 | 0.718539313 | 0.81400356 | 8.140035602 |
| Cdh20 | 0.57248324 | 0.120546578 | 1.044114227 | 0.364745094 | 0.715301694 | 0.81400356 | 8.140035602 |
| Dmd | 10.356573 | 0.124035639 | 0.430716526 | 0.364145268 | 0.715749535 | 0.81400356 | 8.140035602 |
| Reps2 | 8.42129 | -0.121254103 | 0.444355837 | -0.353464288 | 0.723740397 | 0.814207947 | 8.142079466 |
| St18 | 2.4092991 | -0.098187945 | 0.903495563 | -0.355993972 | 0.72184508 | 0.814207947 | 8.142079466 |
| Dppa2 | 13.351748 | 0.110916475 | 0.417130017 | 0.330295575 | 0.741176636 | 0.831089867 | 8.310898671 |
| Lrrc51 | 5.020204 | -0.144269504 | 0.84953177 | 0.326909761 | 0.743736138 | 0.831234507 | 8.312345073 |
| Anapc15 | 8.500967 | 0.10486617 | 0.446680938 | 0.302205632 | 0.762495312 | 0.849424744 | 8.494247444 |
| Kcnq5 | 12.853368 | 0.103893312 | 0.505761215 | 0.280412982 | 0.779160679 | 0.865171922 | 8.651719223 |
| Gm42889 | 12.189364 | -0.089889616 | 0.411614633 | -0.273212667 | 0.784689743 | 0.86849156 | 8.684915603 |
| Riok3 | 9.316052 | -0.144269504 | 0.849531766 | 0.266919391 | 0.789531226 | 0.871031223 | 8.710312235 |
| Gm49767 | 14.771431 | -0.073314267 | 0.326063738 | -0.258749847 | 0.795828256 | 0.872350204 | 8.723502041 |
| Sh3glb1 | 6.793886 | 0.101692253 | 0.665739858 | 0.261671359 | 0.793574824 | 0.872350204 | 8.723502041 |
| Hipk2 | 6.885319 | -0.091958545 | 0.525658361 | -0.250389337 | 0.802286275 | 0.876619508 | 8.766195084 |
| Fbxo4 | 5.3313203 | -0.088773996 | 0.498977308 | -0.246772576 | 0.805084235 | 0.876875186 | 8.768751863 |
| Fyb2 | 24.773638 | 0.052845943 | 0.243259398 | 0.222865361 | 0.823640296 | 0.894238036 | 8.942380356 |
| Gria1 | 12.101741 | -0.144269504 | 0.849531734 | -0.20446334 | 0.837991437 | 0.90407909 | 9.040790899 |
| Wbscr25 | 7.445358 | -0.144269504 | 0.849531755 | -0.206619898 | 0.836306719 | 0.90407909 | 9.040790899 |
| Cadm2 | 10.189448 | 0.068941569 | 0.453019234 | 0.195344786 | 0.845123039 | 0.908905909 | 9.089059094 |
| Magi2 | 10.049273 | 0.058157983 | 0.435597663 | 0.166818043 | 0.867513221 | 0.925608973 | 9.25608973 |
| Dlg2 | 10.546779 | 0.06448642 | 0.638634331 | 0.16521681 | 0.868773334 | 0.925608973 | 9.25608973 |
| Fam155a | 7.520903 | 0.061425415 | 0.485175203 | 0.168415365 | 0.866256521 | 0.925608973 | 9.25608973 |
| Ube2cbp | 11.793842 | -0.144269504 | 0.849531764 | -0.140587277 | 0.888196 | 0.941179063 | 9.411790629 |
| AI314278 | 14.276881 | -0.043891829 | 0.385539662 | -0.1397072 | 0.888891337 | 0.941179063 | 9.411790629 |
| Tnni3k | 6.6259294 | 0.048777801 | 0.505075774 | 0.131516216 | 0.895366959 | 0.945109568 | 9.451095682 |
| Vmn2r103 | 8.293338 | 0.040917341 | 0.445622842 | 0.117478519 | 0.906480865 | 0.953896788 | 9.538967876 |
| Fam169a | 33.938984 | -0.144269504 | 0.84953167 | 0.099398101 | 0.920822191 | 0.966015918 | 9.66015918 |
| Sgcz | 7.1447697 | -0.032568546 | 0.622255056 | -0.087511557 | 0.9302649 | 0.967023087 | 9.670230872 |
| Mrc2 | 18.78206 | 0.028076411 | 0.373911217 | 0.087861067 | 0.929987101 | 0.967023087 | 9.670230872 |
| Comt | 21.540283 | -0.144269504 | 0.849531742 | -0.088923423 | 0.929142768 | 0.967023087 | 9.670230872 |
| Ly75 | 5.598034 | 0.0298386 | 0.541674995 | 0.077687244 | 0.938076841 | 0.9692516 | 9.692516001 |
| Dtnb | 7.362386 | 0.028221451 | 0.462875726 | 0.078642302 | 0.937317139 | 0.9692516 | 9.692516001 |
| Dlgap1 | 8.007651 | -0.144269504 | 0.849531761 | -0.07069566 | 0.943639975 | 0.972062866 | 9.72062866 |
| Ssh2 | 12.196491 | -0.006996846 | 0.495454513 | -0.022082845 | 0.982381871 | 0.995972566 | 9.95972566 |
| Ankrd12 | 21.407003 | -0.144269504 | 0.849531734 | 0.023682296 | 0.981106028 | 0.995972566 | 9.95972566 |
| Pcca | 15.593055 | -0.010144423 | 0.363448154 | -0.034817945 | 0.972224911 | 0.995972566 | 9.95972566 |
| Ralgps1 | 7.1980953 | -0.008520318 | 0.527371668 | -0.025623267 | 0.979557828 | 0.995972566 | 9.95972566 |
| Cntn3 | 14.418162 | -0.00546663 | 0.355907839 | -0.019648512 | 0.984323764 | 0.995972566 | 9.95972566 |
| Grm3 | 0.11452106 | 0.057920704 | 1.645476209 | 0.030710886 | 0.975500109 | 0.995972566 | 9.95972566 |
| Syt1 | 8.639386 | -0.00194023 | 0.52069903 | -0.008339789 | 0.993345889 | 0.9991891 | 9.991890997 |
| Cdh12 | 7.2734222 | -0.003268455 | 0.47683792 | -0.011673545 | 0.990686071 | 0.9991891 | 9.991890997 |
| Cep128 | 11.039948 | 0.0001289 | 0.401707047 | -0.001480219 | 0.998818956 | 0.999298278 | 9.992982781 |
| Crhr2 | 5.539928 | 0.000999476 | 0.520685099 | 0.000879478 | 0.999298278 | 0.999298278 | 9.992982781 |

**table S3: Differential gene expression list social compared to condensed milk in GABA and Glut neurons**

| **GABA** | baseMean | log2FoldChange | lfcSE | stat | pvalue | padj | pglobadj |
|---|---|---|---|---|---|---|---|
| Gm42418 | 85.60561 | 1.61868265 | 0.325259006 | 5.393816412 | 6.90E-08 | 2.43E-05 | 0.000242798 |
| Rps12 | 10.623244 | -0.144269504 | 0.471485981 | -4.616876071 | 3.90E-06 | 0.000685625 | 0.006856252 |
| Rplp1 | 12.864304 | -2.056919948 | 0.734469266 | -3.586395043 | 0.000335281 | 0.039339628 | 0.39339628 |
| Rps15 | 7.9549103 | -1.669660585 | 0.599085319 | -3.441035247 | 0.000579493 | 0.04079631 | 0.4079631 |
| mt-Atp6 | 61.179726 | -0.144269504 | 0.471486066 | 3.488995128 | 0.00048484 | 0.04079631 | 0.4079631 |
| Rps21 | 10.414843 | -0.144269504 | 0.471485993 | -3.028600624 | 0.002456892 | 0.118764336 | 1.187643362 |
| Wdr72 | 7.5489936 | -0.144269504 | 0.471486033 | 3.000068444 | 0.002699189 | 0.118764336 | 1.187643362 |
| mt-Co3 | 73.710365 | -0.144269504 | 0.471486059 | 3.049408005 | 0.002292928 | 0.118764336 | 1.187643362 |
| mt-Cytb | 19.946127 | -0.144269504 | 0.471486033 | 2.954706487 | 0.003129666 | 0.122404731 | 1.224047311 |
| Fau | 5.7841673 | -3.430766306 | 1.534664539 | -2.883140532 | 0.003937317 | 0.138593573 | 1.385935733 |
| Pde1c | 10.640474 | -0.144269504 | 0.471486034 | 2.711215533 | 0.006703703 | 0.214518512 | 2.14518512 |
| Rpl18 | 3.3089302 | -0.144269504 | 0.471486002 | -2.670112766 | 0.007582578 | 0.222422278 | 2.224222784 |
| Rpl30 | 7.003047 | -0.13707031 | 0.364045205 | -2.602600802 | 0.009251961 | 0.250514643 | 2.505146432 |
| Rpl35a | 8.314245 | -0.39256694 | 0.818250337 | -2.573208715 | 0.010076042 | 0.253340493 | 2.533404932 |
| Apoe | 13.775223 | -0.636960776 | 0.496054333 | -2.359832223 | 0.018283202 | 0.359602216 | 3.596022163 |
| Fth1 | 18.389235 | -0.203483395 | 0.433958059 | -2.340585048 | 0.019253553 | 0.359602216 | 3.596022163 |
| Hspa14 | 3.020616 | 0.223608174 | 0.73789178 | 2.326993794 | 0.019965596 | 0.359602216 | 3.596022163 |
| AY036118 | 14.730557 | 0.660506584 | 0.75604538 | 2.381306666 | 0.017251344 | 0.359602216 | 3.596022163 |
| Cox8a | 13.251005 | -0.319346329 | 0.604053089 | -2.382270156 | 0.017206269 | 0.359602216 | 3.596022163 |
| Fbxl7 | 5.554635 | -0.144269504 | 0.471486024 | 2.318319855 | 0.020431944 | 0.359602216 | 3.596022163 |
| Car12 | 9.558533 | -0.192015672 | 0.424921339 | -2.221394996 | 0.02632422 | 0.441244062 | 4.412440625 |
| Sec61a1 | 88.48557 | -0.384070469 | 0.25705238 | -2.092272022 | 0.036414189 | 0.500182029 | 5.001820288 |
| Nckap5 | 30.14854 | 0.415860265 | 0.289952132 | 2.086368418 | 0.036945263 | 0.500182029 | 5.001820288 |
| Mt1 | 4.0376062 | 0.137954884 | 0.6834085 | 2.112156262 | 0.034673049 | 0.500182029 | 5.001820288 |
| Tshz2 | 8.797974 | -0.144269504 | 0.47148601 | 2.089000439 | 0.036707684 | 0.500182029 | 5.001820288 |
| Trim2 | 12.478292 | 0.472201526 | 0.370363857 | 2.095644644 | 0.036113726 | 0.500182029 | 5.001820288 |
| Tnni3k | 3.2754028 | -0.176621352 | 0.424610957 | -2.07023649 | 0.038430204 | 0.501015989 | 5.010159887 |
| Galnt13 | 8.34316 | -0.144269504 | 0.471486026 | 1.980949409 | 0.047596947 | 0.582636034 | 5.82636034 |
| Calb1 | 9.7875395 | -0.260212834 | 0.490569946 | -1.977357236 | 0.048001264 | 0.582636034 | 5.82636034 |
| Rps16 | 6.573574 | -0.189606587 | 0.427413167 | -1.885008443 | 0.059428698 | 0.64472414 | 6.447241401 |
| Rps24 | 7.3618546 | -0.127027599 | 0.369461167 | -1.88524118 | 0.059397283 | 0.64472414 | 6.447241401 |
| Rpl11 | 6.8216047 | -0.208500567 | 0.447475471 | -1.880288674 | 0.060068746 | 0.64472414 | 6.447241401 |
| Atp5l | 2.4516213 | 0.168712973 | 0.552066268 | 1.877549 | 0.060442888 | 0.64472414 | 6.447241401 |
| Pcsk1n | 4.028841 | -0.132615501 | 0.381397359 | -1.85383723 | 0.06376245 | 0.660128899 | 6.601288985 |
| Mapk14 | 13.659798 | 0.35176482 | 0.484581445 | 1.840582506 | 0.065682763 | 0.660580934 | 6.605809344 |
| Camk1d | 8.601393 | -0.144269504 | 0.471486022 | 1.818323741 | 0.069014666 | 0.667177712 | 6.671777119 |
| Hapln1 | 3.5744648 | -0.144269504 | 0.471486013 | 1.797272783 | 0.072292325 | 0.667177712 | 6.671777119 |
| Rpsa | 6.2807426 | -0.097590941 | 0.350671221 | -1.790535043 | 0.073367939 | 0.667177712 | 6.671777119 |
| Rpl8 | 4.5539684 | -0.070357829 | 0.332812989 | -1.787106619 | 0.073920258 | 0.667177712 | 6.671777119 |
| Bc1 | 2.9660645 | -0.232871746 | 0.499616822 | -1.748128453 | 0.080441787 | 0.707887727 | 7.07887727 |
| Tpt1 | 1.8971672 | 0.110094022 | 0.58981089 | 1.713676733 | 0.086588112 | 0.743390622 | 7.433906225 |
| Grik1 | 278.6926 | -0.17430112 | 0.115888468 | -1.65983666 | 0.096947317 | 0.793615243 | 7.936152431 |
| Rps28 | 6.5464745 | -0.181982009 | 0.420946551 | -1.670237296 | 0.094872424 | 0.793615243 | 7.936152431 |
| Slc24a3 | 2.0621026 | -0.144269504 | 0.471486008 | 1.645989001 | 0.099766024 | 0.798128188 | 7.981281883 |
| Chsy3 | 9.549289 | -0.190568522 | 0.40677647 | -1.442876493 | 0.149055266 | 0.825247444 | 8.252474439 |
| Ms4a1 | 9.609934 | 0.247693868 | 0.375235799 | 1.425842749 | 0.153913738 | 0.825247444 | 8.252474439 |
| Rpl23a | 1.9417061 | -0.079744096 | 0.351092766 | -1.553868855 | 0.120215702 | 0.825247444 | 8.252474439 |
| Rpl13 | 8.0091 | -0.122156453 | 0.37676524 | -1.50174343 | 0.133163383 | 0.825247444 | 8.252474439 |
| Mc1r | 13.939334 | 0.26866586 | 0.394850998 | 1.518516067 | 0.128884352 | 0.825247444 | 8.252474439 |
| Cfap74 | 3.0673738 | -0.119350772 | 0.38132682 | -1.617551308 | 0.10575933 | 0.825247444 | 8.252474439 |
| Dctd | 2.9917564 | 0.1260717 | 0.499115657 | 1.484822689 | 0.1375908 | 0.825247444 | 8.252474439 |
| Gpx4 | 6.0329256 | -0.071578579 | 0.342649844 | -1.483726802 | 0.137881413 | 0.825247444 | 8.252474439 |
| Ndufa4 | 4.8546896 | -0.130335922 | 0.38543525 | -1.535873806 | 0.124569334 | 0.825247444 | 8.252474439 |
| Hdac9 | 2.0675054 | -0.144269504 | 0.471486009 | 1.467243057 | 0.142309948 | 0.825247444 | 8.252474439 |
| Rpl32 | 3.1907234 | -0.088411659 | 0.357833965 | -1.477014218 | 0.139671826 | 0.825247444 | 8.252474439 |
| Gm17529 | 2.4453607 | 0.10523836 | 0.512513892 | 1.426008893 | 0.153865775 | 0.825247444 | 8.252474439 |
| Ap3s1 | 1.9563315 | -0.144269504 | 0.471486007 | 1.467243044 | 0.142309951 | 0.825247444 | 8.252474439 |
| Usp3 | 3.9223971 | 0.107493243 | 0.532246395 | 1.495298924 | 0.134836446 | 0.825247444 | 8.252474439 |
| Rplp2 | 2.0136142 | 0.082820849 | 0.592876279 | 1.589913138 | 0.111854386 | 0.825247444 | 8.252474439 |
| Rps5 | 4.1130757 | -0.059974805 | 0.333801143 | -1.46918025 | 0.141783906 | 0.825247444 | 8.252474439 |
| Rps4x | 3.3408737 | -0.092122229 | 0.361159227 | -1.546691326 | 0.121937696 | 0.825247444 | 8.252474439 |
| Atp5md | 1.8126179 | 0.069003 | 0.608129571 | 1.434127676 | 0.151535831 | 0.825247444 | 8.252474439 |
| Alk | 1.9941908 | 0.097839465 | 0.516947608 | 1.461289104 | 0.143936119 | 0.825247444 | 8.252474439 |
| Rps27a | 8.137126 | -0.251666594 | 0.431108204 | -1.577718158 | 0.114630373 | 0.825247444 | 8.252474439 |
| Rps3 | 3.2402453 | -0.111493969 | 0.384706677 | -1.423007794 | 0.154733896 | 0.825247444 | 8.252474439 |
| 4930402H[illegible] | 5.2102904 | -0.151950809 | 0.404076961 | -1.58200382 | 0.113648699 | 0.825247444 | 8.252474439 |
| Dop1b | 7.854299 | -0.144269504 | 0.471486015 | 1.401330614 | 0.161115231 | 0.846456137 | 8.46456137 |
| Ctnna2 | 6.250261 | -0.138283557 | 0.393223894 | -1.368220463 | 0.171243078 | 0.854390974 | 8.543909745 |

| | | | | | | | |
|---|---|---|---|---|---|---|---|
| Ankrd12 | 10.434359 | -0.229680541 | 0.369589088 | -1.377979258 | 0.168209692 | 0.854390974 | 8.543909745 |
| Gm10062 | 18.717892 | 0.214676546 | 0.398953245 | 1.359338314 | 0.174039408 | 0.854390974 | 8.543909745 |
| Cox7b | 2.176436 | -0.048309725 | 0.333668449 | -1.357061122 | 0.17476179 | 0.854390974 | 8.543909745 |
| Mt3 | 3.2125957 | -0.09643541 | 0.370952253 | -1.359181162 | 0.174089189 | 0.854390974 | 8.543909745 |
| Gm50306 | 9.414768 | 0.173796929 | 0.434927477 | 1.3374758 | 0.181067377 | 0.873092008 | 8.730920076 |
| Lrrc17 | 4.438938 | -0.137951004 | 0.403248324 | -1.304928433 | 0.191917218 | 0.888325058 | 8.883250581 |
| Zfpm2 | 14.670027 | -0.19251376 | 0.377579381 | -1.302683102 | 0.192682973 | 0.888325058 | 8.883250581 |
| Epha6 | 14.623177 | 0.227945333 | 0.321818941 | 1.318694321 | 0.187271326 | 0.888325058 | 8.883250581 |
| Gnaz | 16.687979 | -0.144269504 | 0.471486021 | 1.297901727 | 0.194321106 | 0.888325058 | 8.883250581 |
| Fgf12 | 1.8593079 | 0.077158488 | 0.517066946 | 1.287946494 | 0.197764589 | 0.892476095 | 8.924760952 |
| Gm49497 | 4.498791 | -0.113746596 | 0.385699329 | -1.267914259 | 0.204828571 | 0.912653888 | 9.126538879 |
| Atxn7l1 | 3.313615 | 0.110762963 | 0.464887871 | 1.253454019 | 0.210040523 | 0.924178301 | 9.241783012 |
| Dmd | 2.9341674 | -0.093239302 | 0.379102864 | -1.217148246 | 0.223547817 | 0.927502959 | 9.275029593 |
| 8430422H( | 9.372745 | -0.144269504 | 0.471486016 | 1.18767763 | 0.23496044 | 0.927502959 | 9.275029593 |
| Kcnq5 | 4.7545915 | 0.109077431 | 0.438377821 | 1.128215878 | 0.25922876 | 0.927502959 | 9.275029593 |
| Gria4 | 12.067919 | -0.167469789 | 0.369781575 | -1.186933043 | 0.235254033 | 0.927502959 | 9.275029593 |
| Zbtb7a | 7.6496596 | -0.144269504 | 0.471486018 | 1.105161443 | 0.269089641 | 0.927502959 | 9.275029593 |
| Fam169a | 7.7225924 | 0.152321291 | 0.426725119 | 1.227989974 | 0.219450732 | 0.927502959 | 9.275029593 |
| Uqcrh | 3.5035825 | -0.091530061 | 0.377444955 | -1.113214998 | 0.265616103 | 0.927502959 | 9.275029593 |
| Lrp1b | 71.251175 | 0.189828341 | 0.269541901 | 1.131863921 | 0.257691649 | 0.927502959 | 9.275029593 |
| Rpl38 | 7.8183594 | -0.098241677 | 0.371250931 | -1.093817642 | 0.274034965 | 0.927502959 | 9.275029593 |
| St18 | 13.973656 | -0.146533539 | 0.366387938 | -1.098706602 | 0.271896062 | 0.927502959 | 9.275029593 |
| Ccdc60 | 5.917782 | 0.142058108 | 0.434737336 | 1.22591795 | 0.220229553 | 0.927502959 | 9.275029593 |
| Rpl36a | 1.6902087 | 0.068158404 | 0.494203489 | 1.127174187 | 0.259668842 | 0.927502959 | 9.275029593 |
| Cmpk1 | 18.649454 | 0.19047214 | 0.364691707 | 1.201127472 | 0.229701758 | 0.927502959 | 9.275029593 |
| Cmss1 | 4.58798 | 0.117585969 | 0.43894916 | 1.164742531 | 0.244123227 | 0.927502959 | 9.275029593 |
| Rps3a1 | 4.6815195 | -0.075775932 | 0.362747873 | -1.131220868 | 0.257962141 | 0.927502959 | 9.275029593 |
| Rpl3 | 1.3156979 | 0.063789223 | 0.528409203 | 1.240511584 | 0.214786232 | 0.927502959 | 9.275029593 |
| Alpl | 3.2864318 | -0.144269504 | 0.471486015 | 1.145393463 | 0.252046205 | 0.927502959 | 9.275029593 |
| mt-Nd3 | 10.215883 | 0.150511907 | 0.401412464 | 1.126073902 | 0.26013424 | 0.927502959 | 9.275029593 |
| Gm11337 | 5.139593 | -0.116941926 | 0.385452241 | -1.20429113 | 0.228477074 | 0.927502959 | 9.275029593 |
| Rps7 | 0.8670147 | 0.04648042 | 0.753618769 | 1.093862849 | 0.274015135 | 0.927502959 | 9.275029593 |
| Ppia | 2.3876035 | -0.059006827 | 0.353252857 | -1.118191647 | 0.263485151 | 0.927502959 | 9.275029593 |
| Scamp1 | 6.3706956 | -0.127712502 | 0.382484407 | -1.145031307 | 0.252196189 | 0.927502959 | 9.275029593 |
| Sh3glb1 | 2.1542354 | -0.056542811 | 0.358237873 | -1.09915212 | 0.271701719 | 0.927502959 | 9.275029593 |
| Prkg1 | 7.718428 | 0.167896309 | 0.373878306 | 1.109177693 | 0.267353534 | 0.927502959 | 9.275029593 |
| Gphn | 1.5765476 | 0.060375383 | 0.48317334 | 1.023221974 | 0.306202906 | 0.958613119 | 9.586131187 |
| mt-Nd4 | 16.179276 | 0.155206844 | 0.345609474 | 1.014257432 | 0.31045993 | 0.958613119 | 9.586131187 |
| Cdk8 | 3.0650709 | 0.083095506 | 0.451385263 | 1.024469739 | 0.305613457 | 0.958613119 | 9.586131187 |
| mt-Co1 | 36.11727 | 0.166816709 | 0.278141857 | 1.019651364 | 0.307893836 | 0.958613119 | 9.586131187 |
| Elf1 | 2.1457946 | 0.096879106 | 0.441042747 | 1.054266538 | 0.291760904 | 0.958613119 | 9.586131187 |
| Ninj2 | 6.848321 | -0.121167579 | 0.377710221 | -1.064261334 | 0.287210329 | 0.958613119 | 9.586131187 |
| Plp1 | 2.0220082 | -0.063177149 | 0.369665695 | -1.024808042 | 0.305453771 | 0.958613119 | 9.586131187 |
| Cox7c | 7.258831 | -0.067555054 | 0.355138744 | -1.034791969 | 0.300766074 | 0.958613119 | 9.586131187 |
| Atp9b | 6.884336 | -0.114066161 | 0.375280013 | -1.053486976 | 0.29211786 | 0.958613119 | 9.586131187 |
| Gria1 | 2.101285 | -0.06961501 | 0.376662709 | -1.037947764 | 0.299294375 | 0.958613119 | 9.586131187 |
| Vit | 2.0060596 | -0.06479362 | 0.373961686 | -0.986998083 | 0.323643578 | 0.960652306 | 9.606523058 |
| Rps29 | 11.359849 | -0.115544315 | 0.362587726 | -0.976089662 | 0.329020034 | 0.960652306 | 9.606523058 |
| Meg3 | 3.1593094 | 0.090513469 | 0.425126325 | 0.973023031 | 0.330541856 | 0.960652306 | 9.606523058 |
| mt-Nd5 | 6.3741317 | -0.097495619 | 0.37295234 | -0.98895224 | 0.32268651 | 0.960652306 | 9.606523058 |
| Asic2 | 3.3307571 | -0.144269504 | 0.471486005 | -0.968182254 | 0.332953356 | 0.960652306 | 9.606523058 |
| Ehbp1 | 14.291593 | 0.132631061 | 0.372882627 | 0.974043419 | 0.330034981 | 0.960652306 | 9.606523058 |
| 5830416I1 | 39.48457 | 0.162096839 | 0.287246994 | 0.99158314 | 0.321400917 | 0.960652306 | 9.606523058 |
| Atp5h | 5.8244596 | -0.057419638 | 0.350495324 | -0.989572847 | 0.322382947 | 0.960652306 | 9.606523058 |
| Pard3b | 2.687468 | -0.061420419 | 0.37236319 | -0.922566094 | 0.356233372 | 0.964046064 | 9.640460645 |
| Cdh23 | 9.650531 | 0.098139668 | 0.368141801 | 0.764543564 | 0.444543386 | 0.964046064 | 9.640460645 |
| Nup210l | 2.3698108 | 0.085977637 | 0.410521235 | 0.817973095 | 0.41337255 | 0.964046064 | 9.640460645 |
| Cst3 | 33.722935 | -0.109286428 | 0.322097729 | -0.765833342 | 0.443775473 | 0.964046064 | 9.640460645 |
| Ctnna3 | 9.395289 | -0.144269504 | 0.471486006 | -0.708283271 | 0.478769364 | 0.964046064 | 9.640460645 |
| Vwc2l | 0.95087564 | 0.02404605 | 0.540130695 | 0.695434077 | 0.486783311 | 0.964046064 | 9.640460645 |
| 1700054A( | 65.85053 | -0.100260765 | 0.192695724 | -0.688722437 | 0.490997954 | 0.964046064 | 9.640460645 |
| Rpl27a | 5.3361354 | -0.056329842 | 0.363069922 | -0.816215419 | 0.41437694 | 0.964046064 | 9.640460645 |
| Rps2 | 5.409779 | -0.020199124 | 0.330770988 | -0.767361603 | 0.442866551 | 0.964046064 | 9.640460645 |
| Dlgap1 | 1.291002 | 0.029997451 | 0.467291962 | 0.674518827 | 0.49998152 | 0.964046064 | 9.640460645 |
| A330008L1 | 3.729712 | 0.067721534 | 0.419000365 | 0.796387766 | 0.425806687 | 0.964046064 | 9.640460645 |
| Srgn | 2.0774462 | -0.056992043 | 0.370901031 | -0.902548494 | 0.366765573 | 0.964046064 | 9.640460645 |
| Calm1 | 2.599751 | 0.04198544 | 0.432176559 | 0.663558166 | 0.50697314 | 0.964046064 | 9.640460645 |
| Lyrm1 | 2.080669 | 0.051470945 | 0.462151403 | 0.883340945 | 0.377052088 | 0.964046064 | 9.640460645 |
| Rps26 | 3.4884536 | -0.036841467 | 0.35961816 | -0.659755562 | 0.509410705 | 0.964046064 | 9.640460645 |
| Acoxl | 7.196729 | 0.111908195 | 0.397586185 | 0.937580711 | 0.348459928 | 0.964046064 | 9.640460645 |
| Celf2 | 3.1833162 | 0.063820873 | 0.422350445 | 0.768438552 | 0.442226684 | 0.964046064 | 9.640460645 |

| | | | | | | | |
|---|---|---|---|---|---|---|---|
| Pcsk2 | 2.0356083 | 0.051103385 | 0.476974545 | 0.915305488 | 0.360031279 | 0.964046064 | 9.640460645 |
| Csmd3 | 11.044287 | -0.085487581 | 0.347304321 | -0.689984553 | 0.490203902 | 0.964046064 | 9.640460645 |
| Aldoc | 6.1002874 | -0.060586094 | 0.361044722 | -0.846229499 | 0.397424728 | 0.964046064 | 9.640460645 |
| Ahdc1 | 4.428569 | 0.07567611 | 0.406902049 | 0.77514099 | 0.438256353 | 0.964046064 | 9.640460645 |
| Rpl19 | 2.1935105 | 0.039680504 | 0.456001883 | 0.705182693 | 0.48069655 | 0.964046064 | 9.640460645 |
| Rpl18a | 2.4631073 | 0.047256209 | 0.480723459 | 0.850041866 | 0.39530181 | 0.964046064 | 9.640460645 |
| Col5a2 | 2.911921 | 0.048837266 | 0.427158304 | 0.68487743 | 0.493421269 | 0.964046064 | 9.640460645 |
| Rpl12 | 3.0547311 | -0.034391657 | 0.35543467 | -0.679988796 | 0.496511555 | 0.964046064 | 9.640460645 |
| Cox6b1 | 4.9958706 | -0.059366833 | 0.361319026 | -0.839175305 | 0.401370943 | 0.964046064 | 9.640460645 |
| Syt1 | 3.9825034 | 0.061684566 | 0.414183956 | 0.727003289 | 0.467223941 | 0.964046064 | 9.640460645 |
| Lrrc51 | 1.4710289 | -0.047302051 | 0.370639588 | -0.867552785 | 0.385639203 | 0.964046064 | 9.640460645 |
| Ralgps1 | 3.8249373 | 0.098093832 | 0.412610687 | 0.938773056 | 0.347847276 | 0.964046064 | 9.640460645 |
| Rpl37a | 5.236495 | -0.05698257 | 0.368832339 | -0.697043287 | 0.485775704 | 0.964046064 | 9.640460645 |
| Ptpn9 | 8.256581 | -0.090594173 | 0.360637915 | -0.804439077 | 0.421143441 | 0.964046064 | 9.640460645 |
| Rpl39 | 3.8647625 | -0.069656242 | 0.365977123 | -0.948914519 | 0.34266409 | 0.964046064 | 9.640460645 |
| Ppp1r9a | 6.954094 | -0.063812969 | 0.362198362 | -0.753966395 | 0.450869401 | 0.964046064 | 9.640460645 |
| Rpl41 | 6.378624 | -0.060345892 | 0.362482232 | -0.711882196 | 0.476537728 | 0.964046064 | 9.640460645 |
| Rpl7 | 1.628763 | -0.031940725 | 0.360431984 | -0.668673866 | 0.503703542 | 0.964046064 | 9.640460645 |
| Crhr2 | 1.8192136 | -0.047152425 | 0.368237996 | -0.83879945 | 0.401581859 | 0.964046064 | 9.640460645 |
| Pde6a | 23.63924 | 0.102433236 | 0.31030119 | 0.681256713 | 0.495709068 | 0.964046064 | 9.640460645 |
| Rab29 | 6.66237 | -0.106275545 | 0.350594347 | -0.775969466 | 0.437767015 | 0.964046064 | 9.640460645 |
| Lrrc4c | 3.1831484 | 0.062951476 | 0.426230702 | 0.79160138 | 0.428593142 | 0.964046064 | 9.640460645 |
| Tmem108 | 5.4817033 | 0.064550615 | 0.403251768 | 0.696334532 | 0.486219352 | 0.964046064 | 9.640460645 |
| Zmat4 | 8.376503 | -0.07965327 | 0.353736293 | -0.666449782 | 0.505123652 | 0.964046064 | 9.640460645 |
| Asxl3 | 3.21226 | 0.051389217 | 0.42672693 | 0.71689328 | 0.473439949 | 0.964046064 | 9.640460645 |
| 4930511M | 11.236544 | -0.144269504 | 0.471486015 | -0.85239883 | 0.393992769 | 0.964046064 | 9.640460645 |
| mt-Nd1 | 12.260099 | 0.106212837 | 0.351777475 | 0.770273679 | 0.441137566 | 0.964046064 | 9.640460645 |
| mt-Nd2 | 16.444113 | 0.135660518 | 0.363643325 | 0.955852631 | 0.339146694 | 0.964046064 | 9.640460645 |
| Slx1b | 9.106492 | 0.08296196 | 0.397761559 | 0.779792666 | 0.435512924 | 0.964046064 | 9.640460645 |
| Hexb | 2.7214704 | 0.058045331 | 0.429337252 | 0.780082433 | 0.435342356 | 0.964046064 | 9.640460645 |
| Atp5e | 2.2757869 | -0.043190189 | 0.361229757 | -0.826426571 | 0.408562156 | 0.964046064 | 9.640460645 |
| Igf2bp3 | 15.687628 | 0.103142653 | 0.341274292 | 0.735915922 | 0.461781867 | 0.964046064 | 9.640460645 |
| Il1rapl2 | 2.6027884 | -0.052186118 | 0.376699588 | -0.707288939 | 0.479386938 | 0.964046064 | 9.640460645 |
| Gm13219 | 1.6879736 | -0.035157326 | 0.370152302 | -0.684501801 | 0.493658353 | 0.964046064 | 9.640460645 |
| Gas5 | 1.5927483 | -0.036071494 | 0.373441294 | -0.664165732 | 0.506584244 | 0.964046064 | 9.640460645 |
| Sema3c | 75.611465 | 0.10497105 | 0.174701692 | 0.757255808 | 0.448896621 | 0.964046064 | 9.640460645 |
| 4931429P1 | 1.174723 | 0.05714242 | 0.45463605 | 0.879723856 | 0.379008923 | 0.964046064 | 9.640460645 |
| Gapdh | 3.903394 | -0.047928588 | 0.35594146 | -0.820131357 | 0.412141228 | 0.964046064 | 9.640460645 |
| Gm38973 | 4.996316 | 0.092606977 | 0.384536584 | 0.769101526 | 0.441833042 | 0.964046064 | 9.640460645 |
| Lin28b | 2.8430219 | 0.073306301 | 0.435190779 | 0.91785334 | 0.358695653 | 0.964046064 | 9.640460645 |
| Spock1 | 4.8161044 | -0.095494752 | 0.379878297 | -0.918895603 | 0.35815018 | 0.964046064 | 9.640460645 |
| Cadps2 | 2.1911073 | -0.047313291 | 0.375366606 | -0.69801263 | 0.485169296 | 0.964046064 | 9.640460645 |
| Kcnd2 | 1.7874845 | -0.05048702 | 0.375243719 | -0.804620275 | 0.421038838 | 0.964046064 | 9.640460645 |
| Kat7 | 5.802742 | -0.144269504 | 0.471486011 | 0.719679014 | 0.47172265 | 0.964046064 | 9.640460645 |
| mt-Atp8 | 1.6735712 | 0.044239862 | 0.438922595 | 0.683545799 | 0.494262021 | 0.964046064 | 9.640460645 |
| Klkb1 | 2.6711369 | 0.062251884 | 0.437203703 | 0.81001367 | 0.417932319 | 0.964046064 | 9.640460645 |
| Brip1os | 2.1485803 | 0.048018535 | 0.427813357 | 0.681428548 | 0.495600364 | 0.964046064 | 9.640460645 |
| Sgcz | 1.2276738 | 0.03331705 | 0.448311866 | 0.629593363 | 0.528960667 | 0.964736553 | 9.647365528 |
| Pcgf6 | 4.4977202 | -0.076174604 | 0.37010972 | -0.642167578 | 0.520764382 | 0.964736553 | 9.647365528 |
| Gm12216 | 34.58866 | 0.100501175 | 0.250370785 | 0.640463245 | 0.521871477 | 0.964736553 | 9.647365528 |
| Atp5j2 | 2.0205083 | -0.009495101 | 0.32389673 | -0.635723685 | 0.524956536 | 0.964736553 | 9.647365528 |
| Fbxo4 | 2.5364227 | 0.046207372 | 0.420132388 | 0.631280901 | 0.527856873 | 0.964736553 | 9.647365528 |
| Cdh20 | 9.640322 | -0.0771895 | 0.349579186 | -0.643262651 | 0.520053686 | 0.964736553 | 9.647365528 |
| Camk2a | 7.321739 | -0.064616143 | 0.360731272 | -0.631038137 | 0.528015589 | 0.964736553 | 9.647365528 |
| Dbi | 2.5629625 | -0.035621735 | 0.366283814 | -0.623174917 | 0.533169583 | 0.966426865 | 9.664268653 |
| Chl1 | 5.0338683 | -0.046194637 | 0.368584595 | -0.619816629 | 0.535378519 | 0.966426865 | 9.664268653 |
| 1700126G( | 12.808928 | -0.065599392 | 0.344895438 | -0.541839563 | 0.587929034 | 0.968544016 | 9.685440156 |
| Cldn16 | 2.0136976 | 0.030459311 | 0.428222052 | 0.495078086 | 0.620545002 | 0.968544016 | 9.685440156 |
| Rapgef4 | 16.668 | -0.144269504 | 0.471486014 | 0.599195931 | 0.549042236 | 0.968544016 | 9.685440156 |
| Gm49767 | 4.072876 | -0.144269504 | 0.471486009 | -0.493808979 | 0.621441092 | 0.968544016 | 9.685440156 |
| Sgip1 | 2.3842134 | -0.035318278 | 0.377823994 | -0.51435371 | 0.607004717 | 0.968544016 | 9.685440156 |
| Slc25a4 | 1.5848581 | -0.026247257 | 0.364460014 | -0.58228078 | 0.560377569 | 0.968544016 | 9.685440156 |
| Cntnap2 | 14.622923 | 0.074461127 | 0.33168191 | 0.53514603 | 0.592548883 | 0.968544016 | 9.685440156 |
| Rpl17 | 3.130916 | -0.031030046 | 0.36401711 | -0.568032581 | 0.570012845 | 0.968544016 | 9.685440156 |
| Rpl37 | 10.704559 | -0.036124598 | 0.359482514 | -0.543889513 | 0.586517506 | 0.968544016 | 9.685440156 |
| Tcea1 | 2.8919303 | -0.028972083 | 0.357738891 | -0.593856118 | 0.552608351 | 0.968544016 | 9.685440156 |
| Fam189a2 | 2.58684 | -0.041616234 | 0.373677991 | -0.591200744 | 0.554385924 | 0.968544016 | 9.685440156 |
| Sh3bgrl2 | 1.7434117 | -0.03247619 | 0.377863205 | -0.563720847 | 0.572944128 | 0.968544016 | 9.685440156 |
| Lsamp | 7.7228446 | 0.062221956 | 0.373080977 | 0.54998298 | 0.582331047 | 0.968544016 | 9.685440156 |
| Ssh2 | 3.001138 | -0.038304844 | 0.372100419 | -0.573207697 | 0.566504073 | 0.968544016 | 9.685440156 |
| Stard7 | 2.6808255 | 0.035721853 | 0.418333581 | 0.504789245 | 0.613706871 | 0.968544016 | 9.685440156 |

| | | | | | | | |
|---|---|---|---|---|---|---|---|
| Naa11 | 3.672039 | -0.042684427 | 0.371824395 | -0.584746809 | 0.558717974 | 0.968544016 | 9.685440156 |
| Vmn2r103 | 3.7036 | -0.031804887 | 0.368627046 | -0.508783224 | 0.61090418 | 0.968544016 | 9.685440156 |
| Rps15a | 4.9160523 | -0.021719497 | 0.352054072 | -0.525642364 | 0.599136717 | 0.968544016 | 9.685440156 |
| Sorcs2 | 4.7162952 | -0.046588274 | 0.368854376 | -0.518223767 | 0.604302154 | 0.968544016 | 9.685440156 |
| AI314278 | 3.2355316 | -0.043095948 | 0.372339364 | -0.554697438 | 0.579101629 | 0.968544016 | 9.685440156 |
| Ubb | 2.457146 | 0.034418325 | 0.422630628 | 0.526562049 | 0.598497752 | 0.968544016 | 9.685440156 |
| Nrp2 | 2.5498376 | -0.03467945 | 0.375354695 | -0.499367321 | 0.617520637 | 0.968544016 | 9.685440156 |
| Rps10 | 4.1450734 | -0.043224369 | 0.369131951 | -0.535201887 | 0.592510261 | 0.968544016 | 9.685440156 |
| Slco2a1 | 2.270384 | -0.036269089 | 0.371819133 | -0.581191257 | 0.56111156 | 0.968544016 | 9.685440156 |
| Rps27 | 4.1086383 | 0.051080059 | 0.399135803 | 0.548973264 | 0.583023798 | 0.968544016 | 9.685440156 |
| Cep128 | 3.80298 | -0.144269504 | 0.471486003 | -0.493808983 | 0.621441089 | 0.968544016 | 9.685440156 |
| Phkg1 | 5.782508 | 0.048455257 | 0.388611919 | 0.493231132 | 0.621849283 | 0.968544016 | 9.685440156 |
| Rpl9 | 4.4358544 | -0.044515536 | 0.367852775 | -0.539903833 | 0.589263355 | 0.968544016 | 9.685440156 |
| Grid2 | 3.5685031 | 0.051027643 | 0.408475371 | 0.603758715 | 0.546004071 | 0.968544016 | 9.685440156 |
| Svil | 4.1908226 | -0.144269504 | 0.471486008 | -0.497005823 | 0.619184945 | 0.968544016 | 9.685440156 |
| Rps19 | 6.743202 | -0.035324376 | 0.360003241 | -0.563985701 | 0.572763864 | 0.968544016 | 9.685440156 |
| Trim33 | 2.8988419 | -0.034683467 | 0.376528635 | -0.484286763 | 0.628182369 | 0.970740565 | 9.707405646 |
| Fam155a | 1.9344208 | 0.031552457 | 0.425841984 | 0.483451571 | 0.628775138 | 0.970740565 | 9.707405646 |
| Tmeff2 | 4.129691 | 0.03477516 | 0.410665391 | 0.456609297 | 0.647951896 | 0.975435278 | 9.75435278 |
| Sil1 | 10.131317 | -0.055770043 | 0.342800245 | -0.455854695 | 0.64849447 | 0.975435278 | 9.75435278 |
| A630012P( | 6.8366203 | -0.144269504 | 0.471486013 | 0.456492095 | 0.648036154 | 0.975435278 | 9.75435278 |
| Gm29325 | 5.441394 | 0.045992777 | 0.389140177 | 0.47158993 | 0.637219516 | 0.975435278 | 9.75435278 |
| Gm14582 | 5.5686007 | -0.043085557 | 0.367713833 | -0.451067671 | 0.651940778 | 0.975435278 | 9.75435278 |
| Esf1 | 2.7179956 | 0.034992545 | 0.410186762 | 0.462400317 | 0.643794272 | 0.975435278 | 9.75435278 |
| Adamtsl1 | 38.544395 | -0.068730372 | 0.249963158 | -0.448233046 | 0.653985016 | 0.975435278 | 9.75435278 |
| 4930441J1 | 6.0807805 | -0.045618223 | 0.365189719 | -0.470713962 | 0.637845013 | 0.975435278 | 9.75435278 |
| Pecam1 | 1.7546463 | -0.144269504 | 0.47148601 | 0.441542781 | 0.658820097 | 0.977806364 | 9.77806364 |
| Gm42791 | 3.289301 | -0.030950806 | 0.376049917 | -0.438352962 | 0.661130439 | 0.977806364 | 9.77806364 |
| Rpl35 | 2.7068913 | -0.017879429 | 0.363323886 | -0.425792851 | 0.670258796 | 0.983046235 | 9.830462346 |
| Cox4i1 | 6.8070807 | -0.034460548 | 0.366328904 | -0.429452213 | 0.667594163 | 0.983046235 | 9.830462346 |
| Klhl14 | 15.925067 | 0.054479694 | 0.335280382 | 0.409864969 | 0.681905004 | 0.984422791 | 9.844227915 |
| Malat1 | 8.823586 | -0.025125021 | 0.363695142 | -0.347619145 | 0.728126226 | 0.984422791 | 9.844227915 |
| Zfp994 | 4.97191 | -0.032988182 | 0.368185574 | -0.375470207 | 0.7073108 | 0.984422791 | 9.844227915 |
| Lingo2 | 1.8491825 | -0.021455424 | 0.385159052 | -0.345365453 | 0.729819646 | 0.984422791 | 9.844227915 |
| C330002G( | 2.0789924 | -0.144269504 | 0.471486006 | 0.368927408 | 0.712181833 | 0.984422791 | 9.844227915 |
| Xlr5a | 35.891243 | 0.045689684 | 0.288260161 | 0.305153233 | 0.760249449 | 0.984422791 | 9.844227915 |
| Mbp | 3.966034 | 0.02970951 | 0.391397612 | 0.3493985 | 0.726790159 | 0.984422791 | 9.844227915 |
| Wdyhv1 | 6.2250147 | -0.041967104 | 0.363307615 | -0.415263697 | 0.677948874 | 0.984422791 | 9.844227915 |
| Kcnip4 | 5.081118 | -0.144269504 | 0.471486007 | -0.307089792 | 0.758775027 | 0.984422791 | 9.844227915 |
| Itm2b | 2.3965573 | -0.016346767 | 0.365898002 | -0.361980129 | 0.71736688 | 0.984422791 | 9.844227915 |
| Tmsb4x | 7.7842736 | -0.035209145 | 0.353266316 | -0.319113559 | 0.749640402 | 0.984422791 | 9.844227915 |
| Il23a | 2.6201148 | 0.027569659 | 0.403033907 | 0.355656085 | 0.722098137 | 0.984422791 | 9.844227915 |
| Rtkn | 2.542767 | -0.024523253 | 0.376643808 | -0.377295791 | 0.7059538 | 0.984422791 | 9.844227915 |
| Fyb2 | 7.328687 | -0.041294316 | 0.33640488 | -0.315943847 | 0.75204513 | 0.984422791 | 9.844227915 |
| Gm42889 | 4.6184525 | -0.029687439 | 0.370798108 | -0.34378169 | 0.731010471 | 0.984422791 | 9.844227915 |
| Eef1a1 | 6.669673 | 0.034206493 | 0.374900569 | 0.337588145 | 0.735673577 | 0.984422791 | 9.844227915 |
| Rps8 | 6.691774 | -0.01549548 | 0.356873701 | -0.392168981 | 0.694933361 | 0.984422791 | 9.844227915 |
| Ppp3ca | 1.454139 | 0.018353177 | 0.420640251 | 0.331537135 | 0.740238796 | 0.984422791 | 9.844227915 |
| Cpe | 2.779824 | 0.018352776 | 0.416400493 | 0.32957723 | 0.74171943 | 0.984422791 | 9.844227915 |
| Ppp1r16a | 7.9613924 | -0.144269504 | 0.471486009 | -0.390304294 | 0.696311549 | 0.984422791 | 9.844227915 |
| Ralyl | 3.5146954 | 0.026934494 | 0.392496763 | 0.31547169 | 0.752403542 | 0.984422791 | 9.844227915 |
| Chchd6 | 2.8301105 | -0.027428164 | 0.37971199 | -0.356136827 | 0.721738099 | 0.984422791 | 9.844227915 |
| Cox6c | 3.7238717 | -0.023240947 | 0.37349574 | -0.337863037 | 0.735466403 | 0.984422791 | 9.844227915 |
| Olfr57 | 2.1893091 | -0.023481829 | 0.383302584 | -0.349938913 | 0.726384542 | 0.984422791 | 9.844227915 |
| Cntn3 | 3.344907 | -0.032782561 | 0.378017736 | -0.415200426 | 0.677995187 | 0.984422791 | 9.844227915 |
| Riok3 | 3.5079143 | -0.031648075 | 0.373646895 | -0.396533177 | 0.691711738 | 0.984422791 | 9.844227915 |
| Adk | 4.783787 | 0.02856442 | 0.383785259 | 0.304574375 | 0.760690339 | 0.984422791 | 9.844227915 |
| Nkain2 | 3.8744133 | 0.033487695 | 0.397558516 | 0.399569173 | 0.689473866 | 0.984422791 | 9.844227915 |
| A930016O | 3.4797902 | -0.023114459 | 0.377368303 | -0.304884253 | 0.76045431 | 0.984422791 | 9.844227915 |
| Cox11 | 1.366115 | -0.021633824 | 0.386219867 | -0.38351993 | 0.701334296 | 0.984422791 | 9.844227915 |
| Ntm | 14.425692 | -0.144269504 | 0.471486017 | 0.35695709 | 0.72112393 | 0.984422791 | 9.844227915 |
| Uba52 | 1.2092379 | 0.014026518 | 0.434520318 | 0.327520132 | 0.743274519 | 0.984422791 | 9.844227915 |
| Sdccag8 | 4.56701 | -0.144269504 | 0.471486018 | 0.29959824 | 0.764483627 | 0.985707826 | 9.857078262 |
| Ano2 | 2.04199 | 0.016535082 | 0.413007157 | 0.28054799 | 0.779057112 | 0.986110479 | 9.861104794 |
| Tacc1 | 2.6360633 | 0.020905854 | 0.402141168 | 0.273372556 | 0.784566846 | 0.986110479 | 9.861104794 |
| Tshr | 2.0314443 | 0.010626103 | 0.402949261 | 0.156306989 | 0.875791048 | 0.986110479 | 9.861104794 |
| 4930558K( | 6.648917 | -0.144269504 | 0.471486011 | 0.151482942 | 0.879594766 | 0.986110479 | 9.861104794 |
| Wbscr25 | 0.9819156 | -0.001182262 | 0.395325506 | -0.039134252 | 0.968783353 | 0.986110479 | 9.861104794 |
| Anks1b | 2.0393348 | 0.009510467 | 0.402947039 | 0.144306724 | 0.885258269 | 0.986110479 | 9.861104794 |
| Sparcl1 | 2.7145214 | 0.007062511 | 0.394321174 | 0.122082848 | 0.902833406 | 0.986110479 | 9.861104794 |
| Ube2cbp | 4.4138594 | -0.00269406 | 0.377523449 | -0.031816282 | 0.974618562 | 0.986110479 | 9.861104794 |

| | | | | | | | |
|---|---|---|---|---|---|---|---|
| Unc5d | 10.593002 | -0.013510499 | 0.352251285 | -0.124298935 | 0.90107859 | 0.986110479 | 9.861104794 |
| Uqcr11 | 1.3580568 | -0.006184923 | 0.380895722 | -0.156078513 | 0.875971135 | 0.986110479 | 9.861104794 |
| Srbd1 | 2.9975796 | 0.00638508 | 0.394356007 | 0.091879698 | 0.926793622 | 0.986110479 | 9.861104794 |
| 6030443J0 | 10.639899 | -0.023825243 | 0.338908894 | -0.196260145 | 0.844406554 | 0.986110479 | 9.861104794 |
| Zfhx3 | 5.1628814 | -0.144269504 | 0.471486006 | -0.120318763 | 0.904230641 | 0.986110479 | 9.861104794 |
| Sult2a2 | 3.6857228 | 0.002680168 | 0.385471126 | 0.031360653 | 0.97498192 | 0.986110479 | 9.861104794 |
| Stxbp5l | 2.7594132 | 0.009775969 | 0.392818651 | 0.125022715 | 0.900505567 | 0.986110479 | 9.861104794 |
| mt-Co2 | 39.68119 | 0.020965585 | 0.229796614 | 0.136664141 | 0.891296275 | 0.986110479 | 9.861104794 |
| Tns3 | 3.4865246 | -0.022950144 | 0.375350896 | -0.285807709 | 0.775025404 | 0.986110479 | 9.861104794 |
| Gm26901 | 3.472421 | 0.019007325 | 0.390373239 | 0.234540412 | 0.814565464 | 0.986110479 | 9.861104794 |
| Arvcf | 2.5824237 | -0.008183937 | 0.386155633 | -0.139145065 | 0.889335518 | 0.986110479 | 9.861104794 |
| Pkp2 | 4.7125683 | 0.020893723 | 0.381613831 | 0.219182849 | 0.826507616 | 0.986110479 | 9.861104794 |
| Cspp1 | 4.9962697 | -0.002021492 | 0.369686058 | -0.022899352 | 0.981730557 | 0.986110479 | 9.861104794 |
| Pcdh15 | 10.841716 | -0.008757697 | 0.336216502 | -0.072583665 | 0.942137426 | 0.986110479 | 9.861104794 |
| Pcca | 4.715829 | -0.002179043 | 0.37800634 | -0.031479006 | 0.974887535 | 0.986110479 | 9.861104794 |
| Nrg3 | 4.154472 | -0.002714864 | 0.378584516 | -0.036995437 | 0.970488644 | 0.986110479 | 9.861104794 |
| Nell1 | 8.847961 | 0.013318544 | 0.364974165 | 0.12733478 | 0.898675434 | 0.986110479 | 9.861104794 |
| Ctcflos | 7.2694855 | 0.027044236 | 0.336874476 | 0.204575223 | 0.837904015 | 0.986110479 | 9.861104794 |
| D430042O | 23.611097 | -0.029307429 | 0.317267904 | -0.220984166 | 0.825104762 | 0.986110479 | 9.861104794 |
| mt-Nd4l | 1.2082504 | 0.009123914 | 0.419227225 | 0.200143126 | 0.841368646 | 0.986110479 | 9.861104794 |
| Mrc2 | 9.3257475 | -0.020754315 | 0.344487085 | -0.17669602 | 0.859747167 | 0.986110479 | 9.861104794 |
| Micu1 | 1.7600491 | 0.003187834 | 0.401732585 | 0.05463148 | 0.956432059 | 0.986110479 | 9.861104794 |
| Dlg2 | 2.456121 | 0.011673312 | 0.40079636 | 0.199729023 | 0.841692514 | 0.986110479 | 9.861104794 |
| Dppa2 | 5.314692 | -0.019525162 | 0.369446655 | -0.231416382 | 0.81699134 | 0.986110479 | 9.861104794 |
| Magi2 | 3.1816022 | -0.003621462 | 0.383495831 | -0.052051876 | 0.958487359 | 0.986110479 | 9.861104794 |
| Ly75 | 2.1478915 | 0.005858258 | 0.397330576 | 0.085380323 | 0.931959036 | 0.986110479 | 9.861104794 |
| Dpyd | 5.1002426 | 0.029523597 | 0.377217575 | 0.29213847 | 0.770180754 | 0.986110479 | 9.861104794 |
| Dtnb | 2.131943 | 0.011705617 | 0.398445132 | 0.175347811 | 0.860806346 | 0.986110479 | 9.861104794 |
| Klf10 | 3.4066052 | -0.008254282 | 0.385102399 | -0.114813021 | 0.908593328 | 0.986110479 | 9.861104794 |
| Kctd16 | 32.01136 | -0.029845336 | 0.294697203 | -0.20913039 | 0.834346451 | 0.986110479 | 9.861104794 |
| Il27 | 12.893095 | -0.144269504 | 0.471486006 | 0.214229104 | 0.830368401 | 0.986110479 | 9.861104794 |
| Hipk2 | 2.2350583 | 0.007182389 | 0.399394591 | 0.121272998 | 0.903474807 | 0.986110479 | 9.861104794 |
| Hdac8 | 5.1240816 | 0.022114538 | 0.377680666 | 0.22373449 | 0.822963907 | 0.986110479 | 9.861104794 |
| Ftl1 | 2.7191048 | -0.012611249 | 0.377763429 | -0.223807709 | 0.822906931 | 0.986110479 | 9.861104794 |
| Gnao1 | 3.2582612 | 0.008859102 | 0.388048683 | 0.105556461 | 0.915934271 | 0.986110479 | 9.861104794 |
| Gm960 | 3.1102157 | -0.007795104 | 0.382545642 | -0.106535583 | 0.915157425 | 0.986110479 | 9.861104794 |
| Gm765 | 4.3821306 | 0.003448256 | 0.378191063 | 0.036262516 | 0.971073038 | 0.986110479 | 9.861104794 |
| Gm2895 | 2.6113677 | -0.007837079 | 0.384816755 | -0.107538402 | 0.914361861 | 0.986110479 | 9.861104794 |
| Prkag2 | 5.966652 | -0.020799067 | 0.361973093 | -0.203748588 | 0.838549971 | 0.986110479 | 9.861104794 |
| Shb | 20.2632 | -0.014364788 | 0.304646383 | -0.103389221 | 0.917654067 | 0.986110479 | 9.861104794 |
| Ptprd | 3.086377 | -0.00140036 | 0.383598006 | -0.020920556 | 0.983309029 | 0.986110479 | 9.861104794 |
| Ptprg | 3.3906567 | -0.002618649 | 0.385078004 | -0.03500206 | 0.972078099 | 0.986110479 | 9.861104794 |
| Serf2 | 1.2866701 | -0.010802367 | 0.377802617 | -0.253379045 | 0.799975319 | 0.986110479 | 9.861104794 |
| Selenow | 2.374226 | 0.014647625 | 0.406079258 | 0.23226313 | 0.816333647 | 0.986110479 | 9.861104794 |
| Rubcnl | 5.6925087 | 0.004846943 | 0.377371338 | 0.053022668 | 0.957713846 | 0.986110479 | 9.861104794 |
| Rsrc1 | 15.524399 | 0.026142739 | 0.350982219 | 0.223335958 | 0.823274043 | 0.986110479 | 9.861104794 |
| Rps9 | 2.2207394 | 0.001090326 | 0.397498699 | 0.02646599 | 0.97888566 | 0.986110479 | 9.861104794 |
| Atp5k | 0.7781336 | 0.002730198 | 0.413423977 | 0.082639209 | 0.934138423 | 0.986110479 | 9.861104794 |
| C1qa | 2.0312762 | 0.009099403 | 0.40823377 | 0.180063258 | 0.857102907 | 0.986110479 | 9.861104794 |
| Cadm2 | 2.0051184 | -0.002060864 | 0.396354514 | -0.032913399 | 0.973743648 | 0.986110479 | 9.861104794 |
| Rps23 | 3.127832 | -0.003727103 | 0.385047024 | -0.060414418 | 0.951825575 | 0.986110479 | 9.861104794 |
| Rps20 | 1.559221 | -0.009605335 | 0.391426721 | -0.179935001 | 0.857203597 | 0.986110479 | 9.861104794 |
| Rps18 | 1.9495506 | 0.007446922 | 0.403679582 | 0.149137801 | 0.881444898 | 0.986110479 | 9.861104794 |
| Rps17 | 1.4559374 | -0.001638943 | 0.394024296 | -0.030225347 | 0.975887334 | 0.986110479 | 9.861104794 |
| Cdh12 | 2.0843952 | 0.001780274 | 0.396959025 | 0.024987905 | 0.980064611 | 0.986110479 | 9.861104794 |
| Rps14 | 2.248389 | 0.003251849 | 0.403491874 | 0.065531699 | 0.947750668 | 0.986110479 | 9.861104794 |
| Rps13 | 0.11366117 | 0.009858803 | 0.637053472 | 0.126792979 | 0.899104253 | 0.986110479 | 9.861104794 |
| Ckb | 4.6122003 | -0.008129889 | 0.380035941 | -0.10474959 | 0.916574511 | 0.986110479 | 9.861104794 |
| Rpl36 | 1.611016 | -0.015250771 | 0.38033912 | -0.285990819 | 0.774885154 | 0.986110479 | 9.861104794 |
| Gm20342 | 4.540684 | 0.01090833 | 0.376367706 | 0.113169197 | 0.909896416 | 0.986110479 | 9.861104794 |
| Rpl28 | 3.187862 | 0.006084939 | 0.394435509 | 0.089302297 | 0.928841669 | 0.986110479 | 9.861104794 |
| Rpl27 | 2.1159945 | 0.014475692 | 0.40535387 | 0.255479197 | 0.798353009 | 0.986110479 | 9.861104794 |
| Rpl26 | 7.323041 | -0.013892206 | 0.37134063 | -0.201594581 | 0.840233682 | 0.986110479 | 9.861104794 |
| Comt | 5.622409 | -0.015446401 | 0.371649602 | -0.203216381 | 0.838965909 | 0.986110479 | 9.861104794 |
| Rpl23 | 2.1917963 | -0.001726419 | 0.388593656 | -0.028828145 | 0.977001654 | 0.986110479 | 9.861104794 |
| Rpl21 | 6.037702 | -0.001559742 | 0.37953319 | -0.024104577 | 0.980769192 | 0.986110479 | 9.861104794 |
| Rpl13a | 2.3098729 | -0.005825562 | 0.379598753 | -0.145424772 | 0.884375507 | 0.986110479 | 9.861104794 |
| Rgs6 | 0.43800712 | 0.005360717 | 0.439953679 | 0.133732945 | 0.893613753 | 0.986110479 | 9.861104794 |
| Reps2 | 3.6321213 | -0.005721114 | 0.380215609 | -0.080368611 | 0.935944091 | 0.986110479 | 9.861104794 |
| Rpl34 | 2.3196077 | 0.01727847 | 0.410093954 | 0.270490967 | 0.786782565 | 0.986110479 | 9.861104794 |
| Ttc28 | 9.37296 | 0.000215749 | 0.351552056 | -0.000174326 | 0.999860908 | 0.999860908 | 9.998609084 |

| **Glut** | baseMean | log2FoldChange | lfcSE | stat | pvalue | padj | pglobadj |
|---|---|---|---|---|---|---|---|
| Rpl37 | 65.28122 | -0.144269504 | 0.620659798 | -8.105043149 | 5.27E-16 | 1.24E-13 | 1.24E-12 |
| Rpl41 | 41.200634 | -0.144269504 | 0.620659824 | -7.549447468 | 4.37E-14 | 5.16E-12 | 5.16E-11 |
| Rps21 | 78.27365 | -1.964177646 | 0.34755854 | -6.037030179 | 1.57E-09 | 1.23E-07 | 1.23E-06 |
| Rpl35a | 45.59281 | -2.047165769 | 0.398991693 | -5.53743592 | 3.07E-08 | 1.81E-06 | 1.81E-05 |
| Cox4i1 | 68.31515 | -1.855212632 | 0.386346264 | -5.252707818 | 1.50E-07 | 7.07E-06 | 7.07E-05 |
| Rps12 | 53.979713 | -2.136242141 | 0.459542916 | -5.134512001 | 2.83E-07 | 1.11E-05 | 0.000111265 |
| Rpl21 | 42.341732 | -1.384884329 | 0.301359835 | -4.962901432 | 6.94E-07 | 2.34E-05 | 0.000234138 |
| Rpl30 | 42.08132 | -0.144269504 | 0.620659863 | -4.9168576 | 8.79E-07 | 2.59E-05 | 0.000259436 |
| Rpl34 | 24.784636 | -0.144269504 | 0.620659874 | -4.422655833 | 9.75E-06 | 0.000245191 | 0.002451909 |
| Rps10 | 35.73779 | -1.656051779 | 0.423191024 | -4.408905241 | 1.04E-05 | 0.000245191 | 0.002451909 |
| Nckap5 | 82.60741 | -0.144269504 | 0.620659877 | 4.310314863 | 1.63E-05 | 0.000349757 | 0.003497569 |
| Rpl13 | 64.75258 | -1.786319477 | 0.486019614 | -4.266622112 | 1.98E-05 | 0.000390295 | 0.003902946 |
| Lrrc17 | 19.136992 | -1.939309502 | 0.526393041 | -4.183227094 | 2.87E-05 | 0.000456367 | 0.004563665 |
| Rps8 | 56.816334 | -1.710058322 | 0.475732304 | -4.182516469 | 2.88E-05 | 0.000456367 | 0.004563665 |
| Cox8a | 63.387005 | -1.504397723 | 0.413013569 | -4.181130373 | 2.90E-05 | 0.000456367 | 0.004563665 |
| Rplp1 | 96.82634 | -1.191273798 | 0.332290896 | -4.057448336 | 4.96E-05 | 0.000731774 | 0.007317739 |
| Gm10062 | 48.74776 | -0.144269504 | 0.620659898 | 4.000452749 | 6.32E-05 | 0.000877662 | 0.008776619 |
| Ndufa4 | 16.210327 | -1.9083542 | 0.551707468 | -3.971784227 | 7.13E-05 | 0.000935298 | 0.009352984 |
| Rpl18 | 21.894825 | -2.101764447 | 0.629341632 | -3.958083924 | 7.56E-05 | 0.000938453 | 0.009384529 |
| Rpl18a | 28.586264 | -1.601290404 | 0.487266031 | -3.865786943 | 0.000110732 | 0.001272787 | 0.012727871 |
| Rps3 | 25.716751 | -1.634780314 | 0.498169225 | -3.860281485 | 0.000113256 | 0.001272787 | 0.012727871 |
| Cfap74 | 14.551349 | -1.734729708 | 0.521269939 | -3.822393133 | 0.000132163 | 0.001417746 | 0.01417746 |
| Rps19 | 36.5045 | -0.144269504 | 0.620659876 | -3.668771672 | 0.000243719 | 0.002420533 | 0.024205326 |
| Rpl39 | 32.378006 | -1.383575656 | 0.45319818 | -3.635980444 | 0.000276925 | 0.002420533 | 0.024205326 |
| Fau | 47.95611 | -1.855603028 | 0.646575874 | -3.637817784 | 0.000274958 | 0.002420533 | 0.024205326 |
| Lrp1b | 173.90468 | 0.395849411 | 0.111839894 | 3.646099778 | 0.000266251 | 0.002420533 | 0.024205326 |
| Nrg3 | 11.854018 | -0.144269504 | 0.62065992 | 3.64934969 | 0.000262905 | 0.002420533 | 0.024205326 |
| Rps29 | 80.43468 | -0.859955617 | 0.266320196 | -3.605012275 | 0.000312138 | 0.002630879 | 0.026308789 |
| Rpl37a | 48.31256 | -1.123654835 | 0.37971481 | -3.486953904 | 0.000488556 | 0.003975831 | 0.039758314 |
| Cst3 | 242.28601 | -1.018877951 | 0.343427403 | -3.474401324 | 0.000511995 | 0.004010727 | 0.040107269 |
| Dop1b | 19.670872 | -0.144269504 | 0.620659908 | 3.458185214 | 0.000543827 | 0.004010727 | 0.040107269 |
| Acoxl | 17.486248 | -0.144269504 | 0.620659909 | 3.460271957 | 0.00053963 | 0.004010727 | 0.040107269 |
| Mt3 | 47.08135 | -1.070651859 | 0.366413146 | -3.435497532 | 0.000591467 | 0.004105476 | 0.041054758 |
| Cmpk1 | 41.880413 | -0.144269504 | 0.620659887 | 3.438293447 | 0.000585393 | 0.004105476 | 0.041054758 |
| Rpl23 | 32.235893 | -1.479795361 | 0.542720997 | -3.418604949 | 0.00062943 | 0.004244159 | 0.042441592 |
| Rps18 | 26.082087 | -1.40299208 | 0.507464793 | -3.408350066 | 0.00065357 | 0.004284514 | 0.042845138 |
| Rps28 | 26.850864 | -1.47139415 | 0.565856244 | -3.315406973 | 0.000915098 | 0.005821637 | 0.058216375 |
| Rpl36 | 22.518797 | -1.422794195 | 0.540452974 | -3.308676048 | 0.000937382 | 0.005821637 | 0.058216375 |
| Rplp2 | 13.500318 | -1.670431257 | 0.638001748 | -3.294870297 | 0.000984671 | 0.005958523 | 0.05958523 |
| Rpl26 | 44.179935 | -1.295395172 | 0.498211235 | -3.281372526 | 0.001033032 | 0.006094889 | 0.060948892 |
| Rps27a | 37.838913 | -1.218188703 | 0.470497278 | -3.234666527 | 0.001217849 | 0.007010056 | 0.070100564 |
| Rps2 | 19.85445 | -1.390062234 | 0.548839831 | -3.212729106 | 0.001314802 | 0.007387937 | 0.073879371 |
| Atp5j2 | 13.868503 | -1.548105496 | 0.619621982 | -3.192267198 | 0.001411607 | 0.00763532 | 0.076353203 |
| Cdk8 | 13.7301655 | 1.483037715 | 0.585311589 | 3.189836248 | 0.001423534 | 0.00763532 | 0.076353203 |
| Rps20 | 40.926395 | -1.065089088 | 0.409484075 | -3.174901918 | 0.001498872 | 0.007689863 | 0.076898628 |
| Rps14 | 46.925453 | -1.245820289 | 0.501971163 | -3.175388639 | 0.001496359 | 0.007689863 | 0.076898628 |
| Rpl17 | 40.623035 | -0.893418973 | 0.328854755 | -3.168441119 | 0.001532588 | 0.007695548 | 0.076955477 |
| Rpl38 | 41.65708 | -1.139947825 | 0.460948804 | -3.116318156 | 0.001831246 | 0.009003625 | 0.090036253 |
| Mt1 | 39.442078 | -1.035603653 | 0.409259213 | -3.104433839 | 0.001906436 | 0.009182016 | 0.091820164 |
| Rpl8 | 20.108957 | -1.365573132 | 0.591180303 | -3.057090635 | 0.002234967 | 0.010549044 | 0.105490436 |
| Rps24 | 66.27958 | -1.379232278 | 0.640257774 | -3.026031102 | 0.002477868 | 0.011466213 | 0.114662133 |
| Cox7b | 18.56032 | -1.188724239 | 0.505215805 | -3.003310864 | 0.002670595 | 0.011671489 | 0.116714887 |
| Atp5l | 14.397043 | -1.259853167 | 0.534779177 | -3.006873918 | 0.002639492 | 0.011671489 | 0.116714887 |
| Rpl32 | 29.868748 | -1.091142692 | 0.459000947 | -3.008144023 | 0.002628485 | 0.011671489 | 0.116714887 |
| Rpl13a | 24.009502 | -1.356133302 | 0.621993062 | -2.989835692 | 0.002791275 | 0.011977109 | 0.119771094 |
| Rpsa | 35.718075 | -1.350057972 | 0.648598266 | -2.949363071 | 0.003184296 | 0.013419535 | 0.13419535 |
| Rps16 | 27.02593 | -1.317870225 | 0.638851519 | -2.90869872 | 0.003629364 | 0.015026842 | 0.150268419 |
| Serf2 | 14.604772 | -1.236816608 | 0.578420369 | -2.861621561 | 0.004214798 | 0.017139516 | 0.171395156 |
| Apoe | 99.38509 | -0.661135981 | 0.263967552 | -2.856390415 | 0.004284879 | 0.017139516 | 0.171395156 |
| Camk1d | 42.539684 | 0.825020131 | 0.349339306 | 2.827624728 | 0.004689474 | 0.017850257 | 0.178502571 |
| Tns3 | 10.464528 | -0.144269504 | 0.620659915 | 2.833836512 | 0.004599286 | 0.017850257 | 0.178502571 |
| Ftl1 | 13.945372 | -1.254296325 | 0.60251018 | -2.830886048 | 0.004641926 | 0.017850257 | 0.178502571 |
| Atp5k | 13.025406 | -0.993017415 | 0.45116895 | -2.754121572 | 0.00588499 | 0.022045361 | 0.220453607 |
| Tpt1 | 32.712616 | -0.904187223 | 0.437388321 | -2.68030456 | 0.00735552 | 0.027123481 | 0.271234811 |
| AY036118 | 59.607933 | 0.779146179 | 0.358715444 | 2.644095351 | 0.008190958 | 0.029739478 | 0.297394777 |
| Pde1c | 45.888428 | -0.144269504 | 0.620659882 | 2.604421448 | 0.009202951 | 0.032907523 | 0.329075232 |
| Dbi | 22.779673 | -0.91393594 | 0.479773754 | -2.564457008 | 0.010333738 | 0.036399436 | 0.363994355 |
| Rpl19 | 32.816597 | -0.855656204 | 0.444589581 | -2.54713393 | 0.010861174 | 0.037694663 | 0.376946628 |
| Uqcrh | 18.319183 | -1.120251686 | 0.702907643 | -2.537278215 | 0.011171813 | 0.038210839 | 0.382108389 |

| | | | | | | | |
|---|---|---|---|---|---|---|---|
| Rpl28 | 26.411627 | -1.145666037 | 0.786235627 | -2.525127347 | 0.011565637 | 0.038992719 | 0.389927191 |
| Rps13 | 21.236837 | -0.929797908 | 0.538171327 | -2.471171141 | 0.013467135 | 0.044763999 | 0.447639987 |
| Rpl27a | 43.011143 | -0.862437386 | 0.49133204 | -2.456097017 | 0.014045523 | 0.046038104 | 0.460381042 |
| Fth1 | 187.56815 | -0.688077725 | 0.357070485 | -2.422563895 | 0.015411415 | 0.049823206 | 0.498232057 |
| Bc1 | 15.193476 | -1.101026156 | 0.929400556 | -2.396533105 | 0.016550998 | 0.052784265 | 0.527842649 |
| Rps23 | 28.651545 | -0.831470453 | 0.493243655 | -2.37625297 | 0.017489468 | 0.0543094 | 0.543094 |
| Sec61a1 | 151.10835 | -0.144269504 | 0.620659781 | -2.377309953 | 0.017439427 | 0.0543094 | 0.543094 |
| Cmss1 | 47.226562 | 0.651912188 | 0.346539484 | 2.322222601 | 0.020220953 | 0.061975907 | 0.619759067 |
| Pcsk1n | 22.0218 | -0.86096407 | 0.587896462 | -2.281789853 | 0.02250175 | 0.068082218 | 0.680822183 |
| Rps15 | 20.473446 | -0.767930437 | 0.478418341 | -2.251895655 | 0.024328867 | 0.07267864 | 0.726786399 |
| Selenow | 20.185394 | -0.88073699 | 0.682455346 | -2.23959358 | 0.02511732 | 0.074096094 | 0.740960937 |
| Rps15a | 14.6720915 | -0.864121218 | 0.635288203 | -2.221697617 | 0.026303748 | 0.075703469 | 0.75703469 |
| 4930441J1 | 12.73646 | -0.144269504 | 0.620659906 | 2.225427413 | 0.026052559 | 0.075703469 | 0.75703469 |
| Aldoc | 15.870669 | -0.833918818 | 0.604235962 | -2.202044548 | 0.027662162 | 0.078653858 | 0.786538578 |
| Rps7 | 29.40218 | -0.793030821 | 0.563975146 | -2.196164745 | 0.028080154 | 0.078891861 | 0.788918607 |
| Cox6c | 51.074463 | -0.758480402 | 0.532169315 | -2.179956639 | 0.029260676 | 0.081241406 | 0.81241406 |
| Uqcr11 | 14.993872 | -0.722822885 | 0.454425095 | -2.174424013 | 0.029673306 | 0.081429073 | 0.814290732 |
| 4930558K0 | 17.527044 | 0.713311873 | 0.457394948 | 2.159442886 | 0.030815824 | 0.08359235 | 0.835923496 |
| Cox7c | 23.877707 | -0.67631383 | 0.423398702 | -2.152174508 | 0.031383613 | 0.084165143 | 0.841651429 |
| mt-Nd5 | 19.64598 | 0.539701782 | 0.304425131 | 2.134447049 | 0.032806204 | 0.085079825 | 0.850798251 |
| Gm50306 | 26.048487 | -0.144269504 | 0.620659889 | 2.13503992 | 0.032757749 | 0.085079825 | 0.850798251 |
| Atp9b | 17.763279 | -0.144269504 | 0.620659904 | 2.138681195 | 0.032461497 | 0.085079825 | 0.850798251 |
| Sil1 | 22.671968 | 0.623885284 | 0.415106533 | 2.027232093 | 0.042638683 | 0.10937749 | 1.0937749 |
| Gm12216 | 86.75775 | 0.295345658 | 0.155990966 | 2.011072441 | 0.044317806 | 0.112462389 | 1.124623885 |
| Rps26 | 26.535107 | -0.708763834 | 0.706482124 | -1.991588949 | 0.046416177 | 0.116534231 | 1.165342314 |
| Rpl11 | 36.35638 | -0.680694543 | 0.5714122 | -1.981278096 | 0.047560095 | 0.11814929 | 1.181492896 |
| Kctd16 | 63.752888 | 0.301828197 | 0.162121959 | 1.952778675 | 0.050845826 | 0.12499599 | 1.249959899 |
| Svil | 10.612067 | 0.672033155 | 0.538412131 | 1.946467457 | 0.051598613 | 0.125538894 | 1.255388944 |
| Meg3 | 15.650882 | 0.700996795 | 0.648950693 | 1.931632794 | 0.053404845 | 0.128607586 | 1.286075865 |
| Gapdh | 16.667706 | -0.665525187 | 0.611453144 | -1.912792946 | 0.055774565 | 0.132957548 | 1.329575479 |
| Wdr72 | 21.71617 | 0.565327954 | 0.410747434 | 1.877878028 | 0.060397853 | 0.142538933 | 1.425389327 |
| Plp1 | 15.080953 | -0.631214605 | 0.562152854 | -1.861995034 | 0.06260379 | 0.146282124 | 1.462821237 |
| Xlr5a | 78.55026 | 0.418442429 | 0.271779842 | 1.810723255 | 0.070183701 | 0.162385818 | 1.623858184 |
| Rpl12 | 15.030688 | -0.593666609 | 0.527892895 | -1.799390854 | 0.071956875 | 0.164872064 | 1.648720639 |
| Rpl27 | 11.1473675 | -0.596066357 | 0.579933764 | -1.771732884 | 0.076438908 | 0.173457523 | 1.734575227 |
| Atp5e | 16.386784 | -0.507671873 | 0.395194263 | -1.73366613 | 0.082977339 | 0.186501449 | 1.865014485 |
| Nkain2 | 15.254354 | -0.144269504 | 0.620659906 | 1.723217211 | 0.084849252 | 0.188909656 | 1.889096564 |
| Wdyhv1 | 17.055882 | 0.530915042 | 0.450817074 | 1.718294206 | 0.085742966 | 0.189115326 | 1.891153263 |
| Il27 | 28.03912 | 0.478048503 | 0.37486375 | 1.692125421 | 0.090622062 | 0.194917054 | 1.949170539 |
| Nell1 | 19.526506 | 0.507222771 | 0.419609402 | 1.698264414 | 0.089457868 | 0.194917054 | 1.949170539 |
| Gm765 | 11.480239 | -0.144269504 | 0.6206599 | 1.690924784 | 0.090851169 | 0.194917054 | 1.949170539 |
| Vwc2l | 10.584623 | -0.144269504 | 0.620659901 | 1.659741031 | 0.096966561 | 0.204322397 | 2.043223967 |
| Tmsb4x | 58.230568 | -0.498892259 | 0.447652287 | -1.661410196 | 0.096631091 | 0.204322397 | 2.043223967 |
| Rpl9 | 36.284992 | -0.515627649 | 0.58851004 | -1.643111627 | 0.10035984 | 0.209601081 | 2.096010811 |
| Zmat4 | 21.90318 | 0.38742116 | 0.289357185 | 1.620353391 | 0.105156386 | 0.217692167 | 2.176921674 |
| 8430422H0 | 39.241966 | 0.419459862 | 0.328537782 | 1.612977069 | 0.10674949 | 0.219068518 | 2.190685184 |
| Pcca | 10.767588 | -0.144269504 | 0.620659872 | -1.594679359 | 0.110783956 | 0.225388049 | 2.253880488 |
| Malat1 | 52.137634 | 0.455274828 | 0.397872134 | 1.576977401 | 0.114800725 | 0.231563856 | 2.315638557 |
| Ntm | 43.572205 | 0.423846762 | 0.349598547 | 1.571507067 | 0.116064914 | 0.231745025 | 2.317450245 |
| Slc25a4 | 12.412752 | -0.414420385 | 0.690761849 | -1.568114202 | 0.116854483 | 0.231745025 | 2.317450245 |
| Rps9 | 13.941719 | -0.48079111 | 0.597551125 | -1.556191904 | 0.119662466 | 0.235336183 | 2.35336183 |
| Rps17 | 10.877604 | -0.484343394 | 0.533768885 | -1.543733256 | 0.122652966 | 0.239223967 | 2.392239674 |
| Tmem108 | 15.635063 | 0.460889046 | 0.45143097 | 1.528741091 | 0.126328641 | 0.242386661 | 2.423866613 |
| Fam169a | 29.780441 | -0.144269504 | 0.620659881 | 1.529098 | 0.126240151 | 0.242386661 | 2.423866613 |
| C1qa | 14.401668 | -0.144269504 | 0.620659877 | -1.503922097 | 0.132601427 | 0.248596579 | 2.48596579 |
| Rubcnl | 10.739467 | 0.465667679 | 0.516218283 | 1.503441265 | 0.132725292 | 0.248596579 | 2.48596579 |
| Rsrc1 | 34.800335 | 0.395050087 | 0.337111405 | 1.504177628 | 0.132535637 | 0.248596579 | 2.48596579 |
| Rapgef4 | 41.82113 | 0.387404876 | 0.332517067 | 1.486159104 | 0.137237042 | 0.253030796 | 2.53030796 |
| Atp5md | 16.34425 | -0.451758205 | 0.487259446 | -1.486435966 | 0.137163843 | 0.253030796 | 2.53030796 |
| Adk | 12.20822 | 0.423762405 | 0.462939142 | 1.419670557 | 0.155703614 | 0.284853123 | 2.848531228 |
| Prkag2 | 15.796461 | -0.144269504 | 0.620659893 | 1.414576465 | 0.157192713 | 0.285365233 | 2.853652331 |
| Mapk14 | 38.901752 | -0.144269504 | 0.620659877 | 1.402544527 | 0.160752704 | 0.287406349 | 2.874063489 |
| Rps5 | 29.869446 | -0.416232774 | 0.49674472 | -1.402802959 | 0.160675604 | 0.287406349 | 2.874063489 |
| Eef1a1 | 23.16148 | -0.393899457 | 0.409827271 | -1.382814261 | 0.166721825 | 0.295837224 | 2.958372235 |
| mt-Atp6 | 199.6809 | 0.248665183 | 0.198528429 | 1.378202116 | 0.168140894 | 0.296128739 | 2.961287387 |
| Gm11337 | 11.763608 | 0.397227681 | 0.490095071 | 1.331554099 | 0.183006757 | 0.319922923 | 3.199229227 |
| Ptprd | 11.653818 | -0.144269504 | 0.620659908 | 1.31320054 | 0.189115382 | 0.32817081 | 3.2817081 |
| Ctnna3 | 20.84446 | -0.367806489 | 0.413086722 | -1.295861264 | 0.195023287 | 0.335952523 | 3.359525231 |
| Epha6 | 30.104229 | 0.354814709 | 0.393329801 | 1.271505311 | 0.203548939 | 0.348098186 | 3.480981857 |
| Galnt13 | 19.814262 | 0.351465514 | 0.396489971 | 1.258853119 | 0.20808339 | 0.353292662 | 3.532926617 |
| Scamp1 | 13.03109 | 0.362534043 | 0.481600837 | 1.236543593 | 0.216256572 | 0.364546793 | 3.645467934 |

| | | | | | | | |
|---|---|---|---|---|---|---|---|
| Ccdc60 | 11.796525 | 0.352626859 | 0.461316946 | 1.216686323 | 0.223723585 | 0.374459333 | 3.744593331 |
| Gnaz | 42.472843 | 0.310748882 | 0.335394662 | 1.202559813 | 0.229146707 | 0.380835372 | 3.808353723 |
| Ctnna2 | 10.995539 | 0.342337465 | 0.475745506 | 1.181929104 | 0.237233833 | 0.391518773 | 3.915187726 |
| Vit | 15.488907 | 0.333596696 | 0.427439231 | 1.174860967 | 0.240050344 | 0.391840827 | 3.918408269 |
| Gm42418 | 386.77994 | 0.292075991 | 0.311765295 | 1.173115078 | 0.240749661 | 0.391840827 | 3.918408269 |
| Rpl7 | 11.77445 | -0.189178759 | 0.605001071 | -1.163301747 | 0.244707097 | 0.395318398 | 3.953183984 |
| Ehbp1 | 35.2956 | -0.144269504 | 0.620659866 | 1.159332932 | 0.246320503 | 0.395318398 | 3.953183984 |
| 5830416I1 | 84.08612 | 0.255572367 | 0.26137868 | 1.153494019 | 0.248707671 | 0.395318398 | 3.953183984 |
| Lsamp | 22.104828 | 0.319578727 | 0.399214181 | 1.151354034 | 0.249586616 | 0.395318398 | 3.953183984 |
| Atp5h | 20.83939 | -0.265151373 | 0.597110485 | -1.124343578 | 0.260867298 | 0.405030805 | 4.050308049 |
| Pde6a | 55.96816 | 0.297301134 | 0.348471605 | 1.129994612 | 0.258478495 | 0.405030805 | 4.050308049 |
| Tcea1 | 13.930609 | -0.319646991 | 0.435099779 | -1.127638943 | 0.259472433 | 0.405030805 | 4.050308049 |
| Gm49497 | 10.412869 | 0.319792148 | 0.521688757 | 1.117239673 | 0.263891862 | 0.407048885 | 4.070488848 |
| mt-Co2 | 158.46565 | 0.166316602 | 0.159792635 | 1.1097823 | 0.267092848 | 0.407170567 | 4.071705673 |
| Fbxl7 | 13.382306 | 0.317589788 | 0.485337489 | 1.10902048 | 0.267421347 | 0.407170567 | 4.071705673 |
| Mc1r | 26.486118 | 0.292212209 | 0.361733236 | 1.097099264 | 0.272598002 | 0.412391849 | 4.123918489 |
| Ankrd12 | 20.563095 | 0.301640911 | 0.410140435 | 1.084198233 | 0.278276915 | 0.418301604 | 4.183016044 |
| Chsy3 | 21.024092 | -0.144269504 | 0.620659884 | 1.073679111 | 0.282966527 | 0.422658863 | 4.22658863 |
| Cntnap2 | 46.303696 | 0.259308576 | 0.306679486 | 1.059999278 | 0.289144928 | 0.428963855 | 4.289638545 |
| Igf2bp3 | 36.70198 | 0.233714105 | 0.261831358 | 1.056318006 | 0.290822952 | 0.428963855 | 4.289638545 |
| Ckb | 23.07366 | -0.291453717 | 0.432298774 | -1.044898955 | 0.296069673 | 0.433990328 | 4.339903276 |
| Rpl3 | 11.891058 | -0.144269504 | 0.620659891 | -1.034382782 | 0.300957251 | 0.43843155 | 4.384315502 |
| Camk2a | 17.791857 | 0.286703127 | 0.424637185 | 1.026094861 | 0.304846872 | 0.441373386 | 4.413733856 |
| Klkb1 | 11.750686 | 0.2793789 | 0.463127618 | 0.992694185 | 0.32085901 | 0.456161002 | 4.561610016 |
| Gas5 | 20.69724 | 0.276267903 | 0.41905594 | 0.993859069 | 0.320291484 | 0.456161002 | 4.561610016 |
| Cntn3 | 12.64693 | -0.144269504 | 0.620659898 | 0.993393891 | 0.320518038 | 0.456161002 | 4.561610016 |
| Gm29325 | 13.071094 | -0.144269504 | 0.620659888 | -0.957929262 | 0.33809843 | 0.461220979 | 4.612209791 |
| Dpyd | 11.061714 | 0.274524633 | 0.463259893 | 0.97869134 | 0.327732512 | 0.461220979 | 4.612209791 |
| Cox6b1 | 20.553802 | -0.249365421 | 0.545436932 | -0.969323343 | 0.332383886 | 0.461220979 | 4.612209791 |
| Fyb2 | 21.699911 | 0.257639333 | 0.376553136 | 0.961208702 | 0.336447242 | 0.461220979 | 4.612209791 |
| Zfhx3 | 12.507715 | 0.268910585 | 0.411764501 | 0.975549599 | 0.329287711 | 0.461220979 | 4.612209791 |
| Ubb | 14.642679 | -0.265979125 | 0.434196834 | -0.959811826 | 0.337149929 | 0.461220979 | 4.612209791 |
| Cpe | 20.28295 | 0.263686413 | 0.395503679 | 0.966714367 | 0.333686843 | 0.461220979 | 4.612209791 |
| Trim2 | 30.761068 | 0.248642364 | 0.353774532 | 0.948812139 | 0.342716167 | 0.464833422 | 4.648334219 |
| Itm2b | 12.290284 | 0.260516505 | 0.487065017 | 0.936599081 | 0.348964825 | 0.470603993 | 4.706039928 |
| Zbtb7a | 17.7495 | 0.250554698 | 0.417136672 | 0.909521004 | 0.363075176 | 0.478691293 | 4.786912931 |
| mt-Nd1 | 55.542908 | 0.236210824 | 0.340881732 | 0.915898013 | 0.359720389 | 0.478691293 | 4.786912931 |
| Gphn | 11.165733 | 0.253661212 | 0.479334891 | 0.913394268 | 0.361035219 | 0.478691293 | 4.786912931 |
| Pcgf6 | 11.465309 | -0.144269504 | 0.620659907 | 0.910128556 | 0.362754716 | 0.478691293 | 4.786912931 |
| Gm14582 | 10.845287 | 0.24256231 | 0.483010988 | 0.879301249 | 0.379237959 | 0.497223102 | 4.972231019 |
| Mbp | 57.92936 | 0.197320256 | 0.276679646 | 0.865182816 | 0.386938456 | 0.50451644 | 5.045164396 |
| Rps27 | 20.916819 | -0.223713664 | 0.375896458 | -0.841472334 | 0.400083378 | 0.518789435 | 5.187894347 |
| Pcdh15 | 24.104187 | -0.144269504 | 0.620659887 | 0.803042634 | 0.421950093 | 0.54189458 | 5.418945795 |
| Lrrc4c | 12.407509 | 0.219774731 | 0.446881371 | 0.802101791 | 0.422494079 | 0.54189458 | 5.418945795 |
| Gria4 | 20.558704 | 0.207544145 | 0.374716574 | 0.782568791 | 0.433880374 | 0.553490639 | 5.534906395 |
| Kcnip4 | 14.029852 | 0.204976934 | 0.400780056 | 0.762121376 | 0.445987566 | 0.565876697 | 5.658766967 |
| Rps3a1 | 41.11904 | -0.191631461 | 0.378560668 | -0.727421015 | 0.466968085 | 0.586193979 | 5.861939787 |
| D430042O | 51.79939 | 0.172691253 | 0.293068381 | 0.729206342 | 0.465875452 | 0.586193979 | 5.861939787 |
| Prkg1 | 27.762533 | 0.180756016 | 0.343482771 | 0.705696108 | 0.480377142 | 0.599836008 | 5.998360077 |
| Ttc28 | 25.026707 | 0.183545919 | 0.366123596 | 0.699144726 | 0.48446159 | 0.601752291 | 6.017522906 |
| Ms4a1 | 20.96246 | 0.176171954 | 0.377942043 | 0.666104968 | 0.505344009 | 0.621152012 | 6.211520116 |
| mt-Co3 | 273.03525 | 0.134130779 | 0.22902993 | 0.66898279 | 0.503506457 | 0.621152012 | 6.211520116 |
| Spock1 | 11.522049 | -0.144269504 | 0.620659897 | 0.647364356 | 0.517396159 | 0.632670951 | 6.326709508 |
| Calm1 | 21.18861 | 0.160700151 | 0.360963335 | 0.617706736 | 0.536768671 | 0.652976322 | 6.529763216 |
| Ctcflos | 14.221268 | 0.154904283 | 0.437935932 | 0.577584992 | 0.563544342 | 0.682033153 | 6.820331525 |
| Gm38973 | 18.559414 | 0.133812732 | 0.2888175 | 0.57083281 | 0.56811298 | 0.684054404 | 6.840544044 |
| Cdh23 | 29.209711 | 0.140785317 | 0.36548869 | 0.539536215 | 0.589516916 | 0.70622331 | 7.0622331 |
| Rab29 | 17.933344 | 0.136347164 | 0.404904342 | 0.511888287 | 0.608729196 | 0.725556012 | 7.255560118 |
| Ppp1r9a | 12.626955 | -0.126958757 | 0.435634945 | -0.479604841 | 0.631508403 | 0.748924539 | 7.489245388 |
| Asic2 | 15.394251 | -0.124130842 | 0.435120237 | -0.47081578 | 0.637772295 | 0.752571308 | 7.525713076 |
| AI314278 | 11.197256 | -0.144269504 | 0.620659897 | -0.460305674 | 0.645296829 | 0.753911147 | 7.539111468 |
| Ssh2 | 10.826203 | 0.121304027 | 0.478700929 | 0.461359447 | 0.644540741 | 0.753911147 | 7.539111468 |
| mt-Co1 | 133.49173 | -0.096685384 | 0.256845444 | -0.449033901 | 0.653407202 | 0.755902449 | 7.559024488 |
| Slx1b | 20.307749 | 0.119764387 | 0.388411485 | 0.452875867 | 0.65063813 | 0.755902449 | 7.559024488 |
| Gpx4 | 13.078108 | -0.101291571 | 0.513045749 | -0.416186444 | 0.677273578 | 0.779690559 | 7.796905585 |
| 4930511M | 30.31245 | -0.144269504 | 0.620659883 | 0.409665531 | 0.682051319 | 0.781379181 | 7.813791808 |
| Cspp1 | 12.829748 | -0.105446952 | 0.426048136 | -0.399965345 | 0.689182042 | 0.785734115 | 7.857341154 |
| Ahdc1 | 10.719946 | 0.098786474 | 0.455028859 | 0.373901647 | 0.708477491 | 0.803849461 | 8.038494608 |
| Chl1 | 12.945956 | -0.144269504 | 0.620659883 | -0.364073995 | 0.715802755 | 0.808274882 | 8.082748818 |
| Gm49767 | 12.271294 | -0.093743151 | 0.437592859 | -0.357573424 | 0.720662569 | 0.809887459 | 8.098874589 |
| Hexb | 15.148951 | 0.090926815 | 0.453912873 | 0.344356931 | 0.730577874 | 0.817139234 | 8.171392337 |

| | | | | | | | |
|---|---|---|---|---|---|---|---|
| 1700054A( | 135.83694 | -0.144269504 | 0.620659749 | -0.318989086 | 0.749734789 | 0.834610426 | 8.346104255 |
| mt-Nd2 | 84.98618 | -0.074690602 | 0.307988785 | -0.311769112 | 0.755216 | 0.836765146 | 8.367651455 |
| Rpl36a | 12.33492 | -0.077280794 | 0.472238114 | -0.302674481 | 0.762137947 | 0.839494218 | 8.39494218 |
| 1700126G( | 29.80312 | 0.077803906 | 0.365500745 | 0.29919212 | 0.764793461 | 0.839494218 | 8.39494218 |
| Tacc1 | 11.033001 | -0.075783046 | 0.435383586 | -0.289789471 | 0.771977303 | 0.843456683 | 8.434566833 |
| Ptpn9 | 15.482559 | -0.144269504 | 0.620659899 | 0.258613165 | 0.795933724 | 0.865623774 | 8.656237741 |
| Comt | 15.18931 | 0.066262068 | 0.403425657 | 0.250745895 | 0.802010575 | 0.868231632 | 8.682316316 |
| Sparcl1 | 18.419138 | -0.056195536 | 0.332724365 | -0.226138123 | 0.821093995 | 0.884831885 | 8.848318848 |
| Ppp1r16a | 13.233966 | -0.144269504 | 0.620659898 | -0.216706702 | 0.828436925 | 0.885247389 | 8.852473887 |
| Rpl35 | 11.955141 | -0.056242567 | 0.407131217 | -0.216007803 | 0.828981665 | 0.885247389 | 8.852473887 |
| Stxbp5l | 11.643013 | 0.055541188 | 0.43327809 | 0.209373247 | 0.834156875 | 0.886761363 | 8.867613631 |
| Ppia | 15.698862 | -0.044628361 | 0.415372119 | -0.172800106 | 0.862808554 | 0.913106811 | 9.131068106 |
| Csmd3 | 22.342834 | 0.037684668 | 0.377258257 | 0.143289621 | 0.886061452 | 0.93352903 | 9.335290299 |
| Shb | 48.784386 | -0.030017237 | 0.310261893 | -0.126904312 | 0.899016134 | 0.938795609 | 9.387956086 |
| Rpl23a | 13.855545 | -0.144269504 | 0.620659895 | -0.129668373 | 0.896828806 | 0.938795609 | 9.387956086 |
| Mrc2 | 14.887634 | -0.031541348 | 0.413784863 | -0.121829276 | 0.903034228 | 0.938837347 | 9.38837347 |
| 6030443J0 | 23.528912 | -0.028948924 | 0.355773402 | -0.114956306 | 0.908479755 | 0.940356238 | 9.403562379 |
| Zfp994 | 14.274862 | 0.023208827 | 0.417303648 | 0.086494255 | 0.931073523 | 0.955407054 | 9.55407054 |
| mt-Nd4 | 68.051285 | -0.018010693 | 0.279958761 | -0.081347777 | 0.935165379 | 0.955407054 | 9.55407054 |
| mt-Nd3 | 47.42681 | -0.020255384 | 0.329535276 | -0.083394361 | 0.933537972 | 0.955407054 | 9.55407054 |
| mt-Cytb | 73.80245 | 0.009580935 | 0.237544014 | 0.0449103 | 0.964178807 | 0.98080258 | 9.808025798 |
| Ninj2 | 14.1699705 | -0.009564069 | 0.400671824 | -0.037514844 | 0.970074505 | 0.982564734 | 9.825647345 |
| Phkg1 | 12.978579 | 0.00645345 | 0.317285147 | 0.031344736 | 0.974994614 | 0.983327901 | 9.833279008 |
| Kat7 | 11.734534 | -0.005299559 | 0.465721287 | -0.022548649 | 0.982010305 | 0.986189073 | 9.861890725 |
| Rps4x | 11.321524 | 0.003532222 | 0.544916235 | 0.012716848 | 0.989853697 | 0.989853697 | 9.898536968 |
| 4930402H: | 9.076421 | 0.198957747 | 0.472878872 | 0.732910537 | 0.463612996 | | |
| 4931429P1 | 6.352524 | -0.025371953 | 0.48464448 | -0.102133107 | 0.918651023 | | |
| A330008L1 | 8.747367 | -0.090482874 | 0.465269276 | -0.34981657 | 0.726476363 | | |
| A630012P( | 0.19104402 | -0.037093644 | 0.584478829 | -0.257845165 | 0.796526406 | | |
| A930016O | 6.780118 | 0.096158594 | 0.489572526 | 0.372417969 | 0.709581677 | | |
| Adamtsl1 | 1.1302043 | 0.150425816 | 0.719222389 | 0.91195009 | 0.361794992 | | |
| Alk | 4.9244676 | -0.29412319 | 0.564710298 | -1.089489756 | 0.27593797 | | |
| Alpl | 7.970038 | 0.110057119 | 0.483989766 | 0.42247477 | 0.672678512 | | |
| Anks1b | 8.026868 | 0.169582414 | 0.49431698 | 0.640055007 | 0.522136839 | | |
| Ano2 | 8.921904 | -0.045886838 | 0.45320672 | -0.178407544 | 0.858402925 | | |
| Ap3s1 | 9.49328 | 0.052123266 | 0.447061245 | 0.197825875 | 0.843181301 | | |
| Arvcf | 5.695173 | -0.221410822 | 0.532536007 | -0.852150842 | 0.394130376 | | |
| Asxl3 | 9.089721 | 0.245994921 | 0.449047538 | 0.890536563 | 0.373177845 | | |
| Atxn7l1 | 8.92539 | 0.065280848 | 0.46068747 | 0.249245059 | 0.803171227 | | |
| Brip1os | 5.968368 | -0.25704301 | 0.544068479 | -0.971273731 | 0.331411986 | | |
| C330002G( | 5.320138 | 0.202818804 | 0.542386115 | 0.780901591 | 0.434860376 | | |
| Cadm2 | 9.194128 | 0.159080667 | 0.502867497 | 0.606315695 | 0.544305138 | | |
| Cadps2 | 4.1596527 | 0.097003429 | 0.54441842 | 0.405134667 | 0.685378525 | | |
| Calb1 | 0.567519 | -0.022739965 | 0.63741543 | -0.200795443 | 0.840858527 | | |
| Car12 | 8.196756 | -0.389339653 | 0.623338763 | -1.393457999 | 0.163481337 | | |
| Cdh12 | 5.557151 | -0.120368781 | 0.512907512 | -0.484788563 | 0.627826336 | | |
| Cdh20 | 0.37721923 | 0.017830706 | 0.683221949 | 0.139837716 | 0.888788213 | | |
| Celf2 | 8.352298 | -0.144269504 | 0.620659906 | 1.263889282 | 0.206269769 | | |
| Cep128 | 9.558031 | 0.178761808 | 0.495379128 | 0.671247273 | 0.502063018 | | |
| Chchd6 | 5.7411637 | 0.061757502 | 0.500449459 | 0.244952217 | 0.806493429 | | |
| Cldn16 | 9.160386 | 0.261835116 | 0.394394803 | 0.965480059 | 0.334304419 | | |
| Col5a2 | 6.282319 | -0.403639057 | 0.531175104 | -1.33279275 | 0.182599826 | | |
| Cox11 | 5.257544 | -0.015309021 | 0.505540193 | -0.064060283 | 0.948922227 | | |
| Crhr2 | 4.843722 | -0.144269504 | 0.620659907 | 0.647042438 | 0.517604478 | | |
| Dctd | 6.7900023 | 0.101395436 | 0.507907064 | 0.40046952 | 0.688810729 | | |
| Dlg2 | 10.328302 | 0.268881927 | 0.607263863 | 1.03277544 | 0.301709 | | |
| Dlgap1 | 7.9720354 | 0.255128994 | 0.491893358 | 0.920765953 | 0.357172634 | | |
| Dmd | 9.512736 | 0.319708018 | 0.505883433 | 1.116264473 | 0.264308944 | | |
| Dppa2 | 10.103056 | -0.029085553 | 0.442451596 | -0.113033832 | 0.910003733 | | |
| Dtnb | 6.281887 | 0.185971251 | 0.513380491 | 0.705740942 | 0.480349255 | | |
| Elf1 | 6.25058 | -0.270368752 | 0.558155898 | -1.029045209 | 0.303458431 | | |
| Esf1 | 8.628525 | 0.267663689 | 0.51106544 | 0.965141369 | 0.334474009 | | |
| Fam155a | 7.3845167 | 0.349219334 | 0.548353965 | 1.208044905 | 0.22702999 | | |
| Fam189a2 | 6.49523 | -0.144269504 | 0.6206599 | 0.44835264 | 0.653898716 | | |
| Fbxo4 | 6.4332848 | 0.310998592 | 0.54711193 | 1.105499601 | 0.268943167 | | |
| Fgf12 | 8.296385 | 0.381103759 | 0.546007067 | 1.288244813 | 0.197660757 | | |
| Gm13219 | 6.0202317 | -0.144269504 | 0.620659901 | -0.689357674 | 0.490598212 | | |
| Gm17529 | 6.5060134 | -0.144269504 | 0.620659905 | -0.958853724 | 0.33763244 | | |
| Gm20342 | 7.6798315 | 0.122235296 | 0.521515768 | 0.48398035 | 0.628399815 | | |
| Gm26901 | 7.0512342 | -0.144269504 | 0.620659899 | -0.667104435 | 0.504705428 | | |
| Gm2895 | 5.542247 | 0.251908328 | 0.591212868 | 0.969905889 | 0.332093404 | | |

| | | | | | |
|---|---|---|---|---|---|
| Gm42791 | 4.404895 | 0.243130622 | 0.620650992 | 0.980616108 | 0.326782088 |
| Gm42889 | 8.830854 | -0.354154653 | 0.458536371 | -1.213335253 | 0.22500167 |
| Gm960 | 6.4016104 | -0.144269504 | 0.620659896 | -0.609463946 | 0.542216964 |
| Gnao1 | 8.653061 | 0.320406681 | 0.435945971 | 1.131808168 | 0.257715093 |
| Gria1 | 9.121605 | -0.125766468 | 0.456982561 | -0.478823074 | 0.6320645 |
| Grid2 | 9.691301 | -0.144269504 | 0.620659904 | 1.48059043 | 0.138715747 |
| Grik1 | 9.844908 | 0.236074942 | 0.501351306 | 0.861230366 | 0.389111179 |
| Hapln1 | 8.09209 | -0.130545457 | 0.46596912 | -0.497274381 | 0.618995576 |
| Hdac8 | 10.346247 | 0.272985959 | 0.464127512 | 0.974691352 | 0.329713384 |
| Hdac9 | 7.361722 | 0.202277955 | 0.500817013 | 0.752930265 | 0.45149182 |
| Hipk2 | 5.3017225 | -0.144269504 | 0.620659906 | -0.867315809 | 0.385768997 |
| Hspa14 | 5.6771154 | -0.077407096 | 0.492969976 | -0.308205991 | 0.757925594 |
| Il1rapl2 | 7.83866 | -0.144269504 | 0.620659908 | -0.205727129 | 0.837004066 |
| Il23a | 5.837542 | -0.030387293 | 0.490684452 | -0.122501266 | 0.902502044 |
| Kcnd2 | 6.987616 | 0.368832071 | 0.54439983 | 1.258051161 | 0.208373254 |
| Kcnq5 | 9.578302 | 0.019075872 | 0.475482987 | 0.072864934 | 0.941913598 |
| Klf10 | 6.2109737 | -0.144269504 | 0.620659913 | -0.11230478 | 0.910581752 |
| Klhl14 | 2.85936 | 0.361227804 | 0.741180851 | 1.412023831 | 0.157942944 |
| Lin28b | 7.461848 | 0.012299083 | 0.466126429 | 0.046228049 | 0.963128486 |
| Lingo2 | 7.3432455 | -0.144269504 | 0.620659908 | 1.686948447 | 0.091613265 |
| Lrrc51 | 3.743798 | -0.144269504 | 0.620659901 | 0.044362078 | 0.964615789 |
| Ly75 | 3.7452168 | -0.144269504 | 0.620659901 | -0.879595909 | 0.379078256 |
| Lyrm1 | 5.6628814 | -0.144269504 | 0.620659901 | 0.726764451 | 0.467370264 |
| Magi2 | 8.212161 | -0.144269504 | 0.620659906 | 0.477432712 | 0.633054025 |
| Micu1 | 5.3826747 | -0.253213584 | 0.549509848 | -0.965730821 | 0.334178893 |
| Naa11 | 5.254708 | -0.144269504 | 0.62065992 | 1.451621192 | 0.146606963 |
| Nrp2 | 6.3090897 | -0.144269504 | 0.620659911 | 0.603828563 | 0.545957627 |
| Nup210l | 7.297599 | -0.144269504 | 0.620659898 | -0.464246663 | 0.642471032 |
| Olfr57 | 6.0487924 | 0.15081933 | 0.50303358 | 0.578580216 | 0.562872457 |
| Pard3b | 5.9262924 | 0.247609582 | 0.546122386 | 0.923485498 | 0.355754254 |
| Pcsk2 | 7.305017 | 0.46486138 | 0.487334481 | 1.51249308 | 0.130408472 |
| Pecam1 | 5.728719 | 0.121476588 | 0.525548596 | 0.484874352 | 0.627765477 |
| Pkp2 | 8.365823 | -0.144269504 | 0.6206599 | 0.651872849 | 0.514483202 |
| Ppp3ca | 9.708731 | 0.304243492 | 0.478829833 | 1.0699047 | 0.284662207 |
| Ptprg | 8.17407 | 0.27180475 | 0.434345128 | 0.978135299 | 0.328007411 |
| Ralgps1 | 6.2349324 | 0.116956644 | 0.495428243 | 0.452234137 | 0.651100318 |
| Ralyl | 9.8957205 | -0.144269504 | 0.620659899 | 2.176338735 | 0.029529941 |
| Reps2 | 6.9960303 | -0.067705662 | 0.455033771 | -0.260755931 | 0.794280731 |
| Rgs6 | 9.6557255 | 0.166059875 | 0.579019246 | 0.68559954 | 0.492965671 |
| Riok3 | 6.514333 | -0.198253816 | 0.498708358 | -0.748513667 | 0.454150384 |
| Rtkn | 5.2543354 | 0.304405955 | 0.572754926 | 1.102592675 | 0.270204093 |
| Sdccag8 | 5.2209406 | -0.144269504 | 0.620659912 | 0.687911366 | 0.491508599 |
| Sema3c | 3.586898 | 0.240345223 | 0.639630481 | 0.998764165 | 0.317908949 |
| Sgcz | 5.369522 | -0.144269504 | 0.620659899 | -0.480673561 | 0.630748525 |
| Sgip1 | 4.315814 | -0.09132545 | 0.526815274 | -0.381949886 | 0.702498537 |
| Sh3bgrl2 | 5.074447 | 0.015415886 | 0.503436064 | 0.060887743 | 0.951448611 |
| Sh3glb1 | 7.251705 | 0.300509359 | 0.57597014 | 1.090040494 | 0.275695307 |
| Slc24a3 | 8.710463 | 0.167058189 | 0.406074352 | 0.62466131 | 0.532193372 |
| Slco2a1 | 7.080841 | 0.000995971 | 0.471064319 | 0.002287793 | 0.998174607 |
| Sorcs2 | 8.510727 | -0.144269504 | 0.620659911 | 0.821988102 | 0.411083668 |
| Srbd1 | 5.450283 | -0.144269504 | 0.620659909 | 1.272437622 | 0.203217674 |
| Srgn | 5.615288 | 0.281409069 | 0.573303489 | 1.039639381 | 0.298507474 |
| St18 | 7.3797083 | -0.144269504 | 0.620659895 | -0.266291444 | 0.790014762 |
| Stard7 | 7.2272997 | -0.144269504 | 0.620659901 | 0.202304922 | 0.839678352 |
| Sult2a2 | 7.9469604 | -0.144269504 | 0.620659906 | 0.614532041 | 0.538863807 |
| Syt1 | 9.239364 | 0.700173166 | 0.495129614 | 2.062481534 | 0.039161914 |
| Tmeff2 | 10.138145 | -0.144269504 | 0.620659912 | 1.498900313 | 0.133899495 |
| Tnni3k | 5.932917 | -0.144269504 | 0.620659907 | -0.178164896 | 0.858593477 |
| Trim33 | 7.46803 | -0.144269504 | 0.620659902 | -1.874493238 | 0.060862473 |
| Tshr | 7.4009957 | 0.094093391 | 0.419687735 | 0.353815148 | 0.72347742 |
| Tshz2 | 3.2275505 | 0.392666148 | 0.805959602 | 1.54467496 | 0.122424905 |
| Uba52 | 9.345032 | 0.11637957 | 0.479158652 | 0.445033256 | 0.656295721 |
| Ube2cbp | 7.5418496 | -0.351804969 | 0.530721723 | -1.215393499 | 0.224216046 |
| Unc5d | 1.8888023 | 0.184879425 | 0.667125497 | 0.899206212 | 0.368542832 |
| Usp3 | 8.490206 | 0.673845729 | 0.58583096 | 1.912001011 | 0.055876064 |
| Vmn2r103 | 7.5954895 | -0.144269504 | 0.620659902 | 0.575690071 | 0.564824688 |
| Wbscr25 | 5.105775 | -0.144269504 | 0.620659907 | -0.720279576 | 0.471352877 |
| Zfpm2 | 1.1305764 | 0.164138965 | 0.712908664 | 0.951593809 | 0.341303022 |
| mt-Atp8 | 6.522315 | -0.113044866 | 0.496093872 | -0.446871627 | 0.654967751 |
| mt-Nd4l | 8.752745 | -0.036479419 | 0.452399235 | -0.142020841 | 0.887063544 |

**table S4: Differential gene expression list quinine compared to elevated platform in GABA and Glut neurons**

| **GABA** | baseMean | log2FoldChange | lfcSE | stat | pvalue | padj | pglobadj |
|---|---|---|---|---|---|---|---|
| Gm42418 | 120.762505 | -2.414307526 | 0.281559464 | -8.842227871 | 9.38E-19 | 2.08E-16 | 2.08E-15 |
| mt-Co1 | 33.350822 | -1.612740115 | 0.299597766 | -5.709060465 | 1.14E-08 | 1.26E-06 | 1.26E-05 |
| Cox7c | 5.4592285 | -0.144269504 | 1.013834702 | -5.1666856 | 2.38E-07 | 1.76E-05 | 0.000176328 |
| Camk1d | 12.190769 | -2.005549356 | 0.501042035 | -4.491820479 | 7.06E-06 | 0.000313539 | 0.003135392 |
| mt-Atp6 | 61.073914 | -1.382406164 | 0.333325857 | -4.525735939 | 6.02E-06 | 0.000313539 | 0.003135392 |
| Lrp1b | 64.151474 | 0.876467178 | 0.211350012 | 4.334119781 | 1.46E-05 | 0.000541475 | 0.005414746 |
| mt-Nd2 | 19.671959 | -1.629016612 | 0.447084872 | -4.127745021 | 3.66E-05 | 0.001161815 | 0.011618148 |
| mt-Nd1 | 19.258715 | -1.573251971 | 0.446407058 | -4.02198223 | 5.77E-05 | 0.001601464 | 0.01601464 |
| Rtn1 | 4.818771 | -2.776540343 | 0.824236276 | -3.965436156 | 7.33E-05 | 0.001807126 | 0.018071259 |
| Kctd16 | 19.023716 | 1.629385602 | 0.485274821 | 3.692493077 | 0.000222066 | 0.004929874 | 0.049298738 |
| Mbp | 4.1102357 | -5.005476028 | 1.562792732 | -3.559378423 | 0.000371734 | 0.007502259 | 0.075022593 |
| mt-Co3 | 74.39738 | -1.006416997 | 0.322755674 | -3.496726201 | 0.000471005 | 0.008713592 | 0.087135922 |
| Ckb | 4.760432 | -2.412607507 | 0.895944655 | -3.444268331 | 0.000572607 | 0.009778368 | 0.097783676 |
| Hexb | 5.365507 | -1.860707343 | 0.787893479 | -3.145908506 | 0.001655718 | 0.026254956 | 0.262549557 |
| Psap | 4.012689 | -2.88371272 | 1.348121163 | -3.079539095 | 0.002073212 | 0.030683531 | 0.306835315 |
| mt-Nd3 | 14.217884 | -1.31478697 | 0.552213808 | -3.018461851 | 0.002540614 | 0.035251017 | 0.352510174 |
| Rpl9 | 2.8370523 | -2.356418929 | 1.196659045 | -2.920505548 | 0.00349464 | 0.043337214 | 0.433372138 |
| Calm1 | 6.5034804 | -1.783862885 | 0.859288288 | -2.91879891 | 0.003513828 | 0.043337214 | 0.433372138 |
| AY036118 | 19.411974 | -1.21448717 | 0.535026628 | -2.886748784 | 0.003892449 | 0.045480191 | 0.454801914 |
| Rpl38 | 8.163268 | -1.282643289 | 0.586401835 | -2.857670327 | 0.004267635 | 0.047370751 | 0.473707511 |
| Cst3 | 18.172358 | -1.15632179 | 0.521314637 | -2.833682438 | 0.004601503 | 0.048644464 | 0.486444644 |
| Ndrg4 | 2.455602 | -3.54983211 | 1.839517005 | -2.794175065 | 0.00520323 | 0.049558559 | 0.495585593 |
| mt-Co2 | 35.479572 | -0.909182527 | 0.387990545 | -2.784702707 | 0.005357682 | 0.049558559 | 0.495585593 |
| Tpt1 | 6.5563526 | -2.551155244 | 1.753861312 | -2.799798469 | 0.005113452 | 0.049558559 | 0.495585593 |
| Gm10062 | 14.307281 | 0.997451759 | 0.419539589 | 2.710094068 | 0.006726413 | 0.059730546 | 0.597305458 |
| Cox6b1 | 4.983884 | -0.144269504 | 1.013834754 | -2.677846886 | 0.007409708 | 0.060924267 | 0.609242673 |
| Camk2n1 | 4.055593 | -2.020447838 | 1.371784925 | -2.685848461 | 0.007234589 | 0.060924267 | 0.609242673 |
| Calm2 | 5.968031 | -1.048167725 | 2.367457158 | -2.555607582 | 0.010600259 | 0.084044909 | 0.840449088 |
| Igf2bp3 | 12.626369 | 1.101205963 | 0.555397503 | 2.494492416 | 0.012613742 | 0.096560371 | 0.965603713 |
| Nckap5 | 24.799383 | 0.776738583 | 0.390505062 | 2.370089008 | 0.017783803 | 0.131381765 | 1.313817655 |
| mt-Nd4 | 20.578793 | -0.882440102 | 0.487278842 | -2.358557803 | 0.018346102 | 0.131381765 | 1.313817655 |
| D430042O | 13.752109 | 0.981076694 | 0.549737775 | 2.313866616 | 0.02067504 | 0.141287002 | 1.412870025 |
| Rps20 | 3.7027924 | -0.929914836 | 1.633858729 | -2.307946326 | 0.021002122 | 0.141287002 | 1.412870025 |
| Cox8a | 4.676073 | -1.118628538 | 1.262966216 | -2.262244122 | 0.023682325 | 0.149866965 | 1.498669648 |
| mt-Nd5 | 5.0540237 | -1.037863867 | 0.774661879 | -2.24829423 | 0.024557434 | 0.149866965 | 1.498669648 |
| Rpl27a | 5.6623993 | -1.036877267 | 0.757392039 | -2.237603831 | 0.025246902 | 0.149866965 | 1.498669648 |
| 4930511M | 7.149277 | 1.22418466 | 0.773334427 | 2.246195767 | 0.024691471 | 0.149866965 | 1.498669648 |
| Rps29 | 13.822343 | -1.052075289 | 1.257211334 | -2.231426307 | 0.025652904 | 0.149866965 | 1.498669648 |
| Rpl37 | 5.5024624 | -1.073177909 | 1.004508162 | -2.20394798 | 0.027528003 | 0.156697861 | 1.566978615 |
| mt-Cytb | 19.874846 | -0.728686934 | 0.427910741 | -2.170435651 | 0.029973859 | 0.166354915 | 1.663549148 |
| Fam169a | 7.776216 | 0.941359802 | 0.557668215 | 2.135128971 | 0.032750477 | 0.17733185 | 1.773318499 |
| Gm48099 | 3.124489 | -0.941504823 | 1.171578391 | -2.110907707 | 0.034780245 | 0.182862973 | 1.828629727 |
| Rps21 | 5.7281804 | -0.967937442 | 0.860178903 | -2.1035307 | 0.035419405 | 0.182862973 | 1.828629727 |
| Gphn | 4.6438904 | -0.914863966 | 0.883178438 | -2.065058511 | 0.038917464 | 0.196356298 | 1.963562978 |
| Rpl21 | 3.3730946 | -0.90765273 | 1.078459997 | -2.044822143 | 0.040872396 | 0.201637153 | 2.016371529 |
| Fyb2 | 3.9299238 | 1.224824987 | 0.842617975 | 2.010918396 | 0.044334077 | 0.21396011 | 2.139601101 |
| Cox4i1 | 5.497971 | -0.827345408 | 0.970883374 | -1.970815458 | 0.048744988 | 0.225445569 | 2.254455692 |
| Rpl17 | 2.5901263 | -0.744376658 | 1.135153573 | -1.979297815 | 0.047782485 | 0.225445569 | 2.254455692 |
| Atp1b1 | 3.8904188 | -0.704001619 | 1.104658524 | -1.960877257 | 0.049893343 | 0.226047391 | 2.260473906 |
| Gria4 | 5.934157 | 0.960779213 | 0.804395302 | 1.936871576 | 0.052761044 | 0.229148859 | 2.291488588 |
| Nell1 | 6.0054364 | 0.938600987 | 0.779400448 | 1.925881065 | 0.054119226 | 0.229148859 | 2.291488588 |
| Adamtsl1 | 12.99348 | 0.688769783 | 0.458537094 | 1.921198323 | 0.05470671 | 0.229148859 | 2.291488588 |
| Rpl32 | 3.238497 | -0.734759736 | 1.079791915 | -1.927655743 | 0.05389796 | 0.229148859 | 2.291488588 |
| Mt3 | 5.347323 | -0.779330888 | 0.770401751 | -1.895928611 | 0.057969485 | 0.238318994 | 2.38318994 |
| Phactr1 | 7.1939235 | -0.572875857 | 0.999383546 | -1.84642879 | 0.06482997 | 0.257004524 | 2.570045245 |
| Prkg1 | 9.43711 | 0.760602562 | 0.591441762 | 1.849543507 | 0.064379372 | 0.257004524 | 2.570045245 |
| Malat1 | 8.29907 | -0.740452878 | 0.869598092 | -1.837493871 | 0.066137021 | 0.25758629 | 2.575862904 |
| AI314278 | 3.930977 | -0.739588047 | 0.813369238 | -1.827457538 | 0.067631005 | 0.258863502 | 2.588635019 |
| Fth1 | 7.0429077 | -0.730645228 | 0.723919868 | -1.79152293 | 0.073209418 | 0.275465947 | 2.754659472 |
| Pip5k1b | 3.6361911 | -0.684166839 | 0.893191141 | -1.778120139 | 0.075384126 | 0.278921265 | 2.789212646 |
| Ttc28 | 5.5298276 | 0.803088302 | 0.767694757 | 1.764536684 | 0.077641665 | 0.282564747 | 2.825647471 |
| Car12 | 3.8203118 | -0.589337998 | 0.97137019 | -1.73918766 | 0.082001759 | 0.28895858 | 2.889585804 |
| Rps27 | 2.5134103 | -0.144269504 | 1.013834799 | -1.746172568 | 0.08078097 | 0.28895858 | 2.889585804 |
| Uba52 | 2.4032862 | -0.597127489 | 0.985201907 | -1.713504165 | 0.086619828 | 0.291357602 | 2.913576024 |

| | | | | | | | |
|---|---|---|---|---|---|---|---|
| Tmeff2 | 5.0455685 | -0.658794942 | 0.806955773 | -1.716589366 | 0.086054227 | 0.291357602 | 2.913576024 |
| Gm19951 | 2.8695545 | -0.636101502 | 0.934553205 | -1.727776735 | 0.08402826 | 0.291357602 | 2.913576024 |
| Ntm | 17.162569 | 0.49544029 | 0.345276564 | 1.698073156 | 0.089493955 | 0.296532211 | 2.965322105 |
| Rpl23 | 5.883805 | -0.354846572 | 0.828357888 | -1.685531993 | 0.091885978 | 0.299980692 | 2.999806915 |
| Galnt13 | 7.401682 | 0.664190221 | 0.683494891 | 1.625849209 | 0.103981737 | 0.332059131 | 3.320591309 |
| Plp1 | 3.8196154 | -0.378104589 | 0.84895893 | -1.622467299 | 0.10470333 | 0.332059131 | 3.320591309 |
| Camk2a | 5.2652774 | 0.686824115 | 0.746448499 | 1.613523279 | 0.10663087 | 0.333409199 | 3.334091986 |
| Il1rapl2 | 2.5361302 | -0.144269504 | 1.01383476 | -1.60589718 | 0.108296503 | 0.333914219 | 3.339142187 |
| Dop1b | 5.258408 | 0.664334029 | 0.793682331 | 1.587596618 | 0.112377583 | 0.33713275 | 3.371327505 |
| Mc1r | 12.064563 | 0.579888973 | 0.512945627 | 1.58823759 | 0.112232624 | 0.33713275 | 3.371327505 |
| Xlr5a | 27.045874 | 0.481702705 | 0.38064784 | 1.580468811 | 0.113999544 | 0.33743865 | 3.374386503 |
| Shb | 15.035331 | 0.483353702 | 0.370443951 | 1.565477653 | 0.117470953 | 0.343138837 | 3.431388372 |
| Rab29 | 4.315765 | 0.63109613 | 0.903974624 | 1.556055292 | 0.119694945 | 0.345094516 | 3.450945163 |
| Acoxl | 5.559864 | 0.621853329 | 0.612445398 | 1.535964663 | 0.124547047 | 0.354480058 | 3.54480058 |
| Cep128 | 2.4896834 | 0.648564961 | 1.065861272 | 1.527585638 | 0.12661545 | 0.355805442 | 3.558054416 |
| Slx1b | 6.9713464 | 0.573723319 | 0.647003577 | 1.472685648 | 0.140835818 | 0.38225293 | 3.822529298 |
| Uqcrh | 2.2643454 | -0.437420604 | 0.92535462 | -1.471364666 | 0.141192524 | 0.38225293 | 3.822529298 |
| Lrrc4c | 2.9008563 | -0.144269504 | 1.013834812 | 1.473577583 | 0.140595361 | 0.38225293 | 3.822529298 |
| Klhl14 | 31.507303 | 0.38148638 | 0.304720256 | 1.462569399 | 0.143585242 | 0.384047273 | 3.84047273 |
| Mapk14 | 11.059541 | 0.517323899 | 0.504983985 | 1.456052461 | 0.145378118 | 0.384213597 | 3.842135973 |
| Snhg11 | 5.4747453 | -0.520841667 | 0.738888551 | -1.448145178 | 0.147576453 | 0.385434971 | 3.854349713 |
| 5830416I1 | 26.229902 | 0.425210484 | 0.372784798 | 1.426359933 | 0.153764473 | 0.394359312 | 3.943593124 |
| Rps7 | 2.3186011 | -0.333899469 | 0.861453749 | -1.423655515 | 0.154546217 | 0.394359312 | 3.943593124 |
| Cpe | 4.1240735 | -0.501523972 | 0.764283654 | -1.412149544 | 0.157905933 | 0.398353604 | 3.983536044 |
| Gm49497 | 2.7283678 | 0.589795474 | 0.817933407 | 1.39269575 | 0.163711814 | 0.403822475 | 4.038224755 |
| Rpl35a | 4.1580353 | -0.380847525 | 0.845337232 | -1.395558911 | 0.162847363 | 0.403822475 | 4.038224755 |
| Ppia | 2.6346543 | -0.369210499 | 0.882632945 | -1.319226394 | 0.187093437 | 0.456425747 | 4.564257468 |
| Cdk8 | 5.4957523 | -0.416134056 | 0.797829497 | -1.303338907 | 0.192459084 | 0.458838283 | 4.588382828 |
| Cspp1 | 3.1568568 | 0.452232939 | 0.923795892 | 1.298595585 | 0.194082754 | 0.458838283 | 4.588382828 |
| Gm765 | 2.9087982 | 0.47015365 | 0.909843232 | 1.298012975 | 0.194282877 | 0.458838283 | 4.588382828 |
| Kat7 | 4.6371884 | 0.480965695 | 0.744313338 | 1.27821 | 0.201175391 | 0.470115124 | 4.701151245 |
| Sema3c | 26.570084 | 0.3528113 | 0.337150398 | 1.2680577 | 0.204777346 | 0.473547612 | 4.735476124 |
| Ccdc60 | 3.6140106 | 0.466038793 | 0.812600564 | 1.255982465 | 0.209122328 | 0.478609863 | 4.78609863 |
| Ms4a1 | 7.180874 | 0.451472481 | 0.601794342 | 1.236823201 | 0.216152727 | 0.489652095 | 4.896520949 |
| Rps14 | 4.5782995 | -0.247161451 | 0.775579467 | -1.209097057 | 0.22662556 | 0.498127468 | 4.981274682 |
| Scamp1 | 4.529276 | 0.448927495 | 0.741090009 | 1.216154625 | 0.223926025 | 0.498127468 | 4.981274682 |
| Klkb1 | 2.677563 | 0.454341062 | 0.848170913 | 1.212303778 | 0.225396118 | 0.498127468 | 4.981274682 |
| 1700126G( | 10.648242 | 0.407218564 | 0.512791019 | 1.183422433 | 0.236641771 | 0.512475544 | 5.124755445 |
| Srgn | 3.0494854 | -0.396408384 | 0.740219246 | -1.18057856 | 0.237770185 | 0.512475544 | 5.124755445 |
| Pde1c | 10.393786 | 0.406538202 | 0.535666098 | 1.164897123 | 0.244060638 | 0.520975593 | 5.209755932 |
| mt-Nd4l | 2.6586802 | -0.207657785 | 0.773175514 | -1.156605046 | 0.24743376 | 0.523145663 | 5.231456632 |
| Tmem108 | 4.5202613 | 0.398661497 | 0.741662277 | 1.119355554 | 0.262988484 | 0.550787203 | 5.507872032 |
| Rps3a1 | 7.1764665 | -0.10327357 | 0.680015861 | -1.103801513 | 0.269679248 | 0.55540123 | 5.554012303 |
| Sil1 | 7.4833884 | 0.384671644 | 0.699982966 | 1.102613159 | 0.270195193 | 0.55540123 | 5.554012303 |
| Fau | 3.9611588 | -0.340927768 | 0.765794504 | -1.081266416 | 0.279578622 | 0.560253771 | 5.602537712 |
| Gm12216 | 27.56392 | 0.268393258 | 0.288958826 | 1.080034347 | 0.280126886 | 0.560253771 | 5.602537712 |
| Phkg1 | 4.9395423 | -0.144269504 | 1.013834789 | 1.085111398 | 0.27787232 | 0.560253771 | 5.602537712 |
| Epha6 | 16.498692 | -0.333514406 | 0.435931986 | -1.06348144 | 0.287563678 | 0.56999229 | 5.699922896 |
| Mt1 | 3.4968033 | -0.306299687 | 0.803130171 | -1.05179611 | 0.292893104 | 0.575418311 | 5.754183106 |
| Syt1 | 2.3429403 | -0.144269504 | 1.013834781 | 1.02534593 | 0.305199991 | 0.589168678 | 5.891686781 |
| 4930441J1 | 2.5527937 | 0.324571612 | 0.909680413 | 1.03058754 | 0.302734282 | 0.589168678 | 5.891686781 |
| Nrg1 | 2.8986304 | -0.287006462 | 0.825970132 | -1.011964594 | 0.311554983 | 0.596251777 | 5.962517772 |
| Cmpk1 | 15.688774 | 0.308716745 | 0.416379013 | 0.994339005 | 0.320057852 | 0.607289258 | 6.072892583 |
| Ctnna3 | 7.6658707 | 0.338762797 | 0.647905664 | 0.978204029 | 0.327973424 | 0.611849581 | 6.118495814 |
| Adk | 4.3268514 | -0.144269504 | 1.013834773 | -0.981849241 | 0.326174125 | 0.611849581 | 6.118495814 |
| Unc5d | 11.219402 | -0.307674093 | 0.477294906 | -0.965713587 | 0.334187519 | 0.61824691 | 6.182469097 |
| Rpl30 | 3.325701 | 0.319382479 | 0.805601019 | 0.959653077 | 0.337229846 | 0.618719222 | 6.187192221 |
| Hdac9 | 3.085551 | -0.144269504 | 1.01383482 | 0.945142846 | 0.344585954 | 0.627033457 | 6.270334566 |
| Rpl41 | 3.7210357 | -0.144269504 | 1.013834787 | -0.935783 | 0.349384926 | 0.630597183 | 6.305971834 |
| 6030443J0 | 7.5854096 | 0.314163428 | 0.605523789 | 0.915700995 | 0.359823743 | 0.640181368 | 6.401813678 |
| Ninj2 | 5.088945 | 0.307902389 | 0.720556696 | 0.914484189 | 0.360462482 | 0.640181368 | 6.401813678 |
| Rplp1 | 7.2411995 | -0.294529778 | 0.587323392 | -0.886733421 | 0.37522244 | 0.654649662 | 6.546496622 |
| Atp9b | 5.629104 | 0.302797124 | 0.652083558 | 0.880621516 | 0.378522709 | 0.654649662 | 6.546496622 |
| Gm38973 | 5.3755755 | -0.284426592 | 0.636963278 | -0.866342661 | 0.386302278 | 0.654649662 | 6.546496622 |
| Zbtb7a | 7.4976068 | 0.295531998 | 0.594532582 | 0.867159931 | 0.385854387 | 0.654649662 | 6.546496622 |
| 8430422H( | 9.972306 | -0.144269504 | 1.013834827 | 0.889229142 | 0.373879944 | 0.654649662 | 6.546496622 |
| St18 | 3.2548878 | 0.277284473 | 0.824856706 | 0.875249579 | 0.381438122 | 0.654649662 | 6.546496622 |

| | | | | | | | |
|---|---|---|---|---|---|---|---|
| Tmsb4x | 2.8991706 | -0.25653962 | 0.777282681 | -0.84702069 | 0.396983589 | 0.655240084 | 6.552400839 |
| Csmd3 | 6.9874845 | 0.282414198 | 0.584862642 | 0.827161173 | 0.408145715 | 0.655240084 | 6.552400839 |
| Tns3 | 3.2166498 | 0.26656476 | 0.822827654 | 0.846748979 | 0.397135052 | 0.655240084 | 6.552400839 |
| Bc1 | 3.4880683 | 0.236416686 | 0.864736281 | 0.81824997 | 0.413214467 | 0.655240084 | 6.552400839 |
| Ankrd12 | 6.749281 | -0.273161568 | 0.617628694 | -0.825282848 | 0.409211027 | 0.655240084 | 6.552400839 |
| Rpl26 | 8.074699 | -0.281904302 | 0.62362733 | -0.859377559 | 0.390132247 | 0.655240084 | 6.552400839 |
| 1700054A( | 51.351105 | 0.268681646 | 0.471129797 | 0.826247419 | 0.408663754 | 0.655240084 | 6.552400839 |
| Pcdh15 | 11.550101 | -0.260975885 | 0.488480697 | -0.819809803 | 0.412324542 | 0.655240084 | 6.552400839 |
| Ppp1r9a | 4.736625 | -0.144269504 | 1.013834805 | 0.841199406 | 0.400236233 | 0.655240084 | 6.552400839 |
| Cfap74 | 2.9782124 | 0.255410383 | 0.807459551 | 0.800798594 | 0.423248253 | 0.661817695 | 6.618176946 |
| Rpl12 | 3.1326845 | -0.223952237 | 0.797131176 | -0.800666344 | 0.423324832 | 0.661817695 | 6.618176946 |
| Rapgef4 | 15.866798 | -0.144269504 | 1.013834809 | 0.785078557 | 0.432407511 | 0.666628246 | 6.666282464 |
| Rpl13 | 5.3730884 | -0.238844353 | 0.715145263 | -0.78579672 | 0.431986589 | 0.666628246 | 6.666282464 |
| Hapln1 | 4.838067 | 0.240190151 | 0.687584059 | 0.726676295 | 0.467424278 | 0.687014051 | 6.870140511 |
| Cntnap2 | 19.890574 | -0.223062149 | 0.457704619 | -0.714726224 | 0.47477823 | 0.687014051 | 6.870140511 |
| Rps17 | 3.0598915 | -0.165765518 | 0.798931158 | -0.698474524 | 0.484880485 | 0.687014051 | 6.870140511 |
| Rps8 | 4.826319 | -0.191048128 | 0.767999883 | -0.751803108 | 0.452169473 | 0.687014051 | 6.870140511 |
| Pde6a | 18.263357 | 0.240174907 | 0.537141678 | 0.720141576 | 0.471437832 | 0.687014051 | 6.870140511 |
| Zfp994 | 4.4025946 | 0.210338446 | 0.842954042 | 0.741303883 | 0.458509208 | 0.687014051 | 6.870140511 |
| Ptgds | 7.3842645 | 0.22544472 | 0.712820409 | 0.709122038 | 0.478248748 | 0.687014051 | 6.870140511 |
| Ehbp1 | 10.079038 | 0.231738484 | 0.537905253 | 0.696906534 | 0.485861288 | 0.687014051 | 6.870140511 |
| Spock1 | 4.125073 | -0.242152889 | 0.680142007 | -0.756739912 | 0.449205698 | 0.687014051 | 6.870140511 |
| Fbxl7 | 5.428398 | 0.247448243 | 0.650368068 | 0.73689156 | 0.461188297 | 0.687014051 | 6.870140511 |
| Ahdc1 | 3.9941804 | 0.230601754 | 0.749751078 | 0.721063782 | 0.470870273 | 0.687014051 | 6.870140511 |
| Svil | 3.1932223 | 0.21015819 | 0.822119473 | 0.70059509 | 0.483555744 | 0.687014051 | 6.870140511 |
| Gnaz | 14.256076 | 0.233577079 | 0.473752094 | 0.724637011 | 0.468674747 | 0.687014051 | 6.870140511 |
| Kcnd2 | 3.1112175 | -0.192935691 | 0.76550099 | -0.679150839 | 0.497042293 | 0.69837588 | 6.983758801 |
| Nkain2 | 5.6990433 | 0.221336286 | 0.655733733 | 0.67250533 | 0.501262048 | 0.699875313 | 6.998753127 |
| Prkag2 | 5.047195 | -0.144269504 | 1.013834777 | 0.654904192 | 0.512529441 | 0.7111346 | 7.111345996 |
| Nrxn3 | 2.9000769 | -0.180156559 | 0.76300469 | -0.612039289 | 0.540511764 | 0.745301935 | 7.453019349 |
| Zfhx3 | 3.657378 | -0.144269504 | 1.013834773 | 0.597463241 | 0.550198142 | 0.753975231 | 7.539752315 |
| Nrxn1 | 3.2233655 | -0.150695135 | 0.789462947 | -0.571866863 | 0.567412175 | 0.77053305 | 7.705330503 |
| Apoe | 4.2958026 | -0.144269504 | 1.013834795 | 0.569196768 | 0.569222614 | 0.77053305 | 7.705330503 |
| Rps16 | 2.6041517 | -0.158278558 | 0.797544282 | -0.553861428 | 0.579673684 | 0.777822278 | 7.778222783 |
| Cdh23 | 8.561942 | -0.144269504 | 1.013834794 | -0.551027451 | 0.581614857 | 0.777822278 | 7.778222783 |
| Rpl37a | 5.688296 | -0.146481189 | 0.741910813 | -0.537849476 | 0.590680972 | 0.785216621 | 7.852166211 |
| Meg3 | 4.08225 | 0.130057744 | 0.849195678 | 0.500690146 | 0.616589209 | 0.801585258 | 8.015852583 |
| Vit | 2.6597064 | -0.144269504 | 1.013834801 | 0.49948565 | 0.617437294 | 0.801585258 | 8.015852583 |
| Ptpn9 | 6.567304 | 0.165723384 | 0.606950817 | 0.504550263 | 0.613874751 | 0.801585258 | 8.015852583 |
| Grm7 | 3.3865728 | -0.144269504 | 1.013834765 | -0.501920413 | 0.615723506 | 0.801585258 | 8.015852583 |
| Mrc2 | 5.057316 | 0.158440859 | 0.647729643 | 0.486564067 | 0.626567293 | 0.80403433 | 8.040343302 |
| Gnao1 | 2.5983818 | 0.14559005 | 0.808050019 | 0.487751913 | 0.625725576 | 0.80403433 | 8.040343302 |
| Eef1a1 | 2.634947 | -0.126909239 | 0.80384393 | -0.466239687 | 0.641043945 | 0.817883654 | 8.178836537 |
| Cox6c | 4.481963 | 0.140054652 | 0.693689682 | 0.443235817 | 0.657595174 | 0.817928479 | 8.179284786 |
| Il27 | 9.28567 | 0.145242895 | 0.523411185 | 0.445670806 | 0.655835056 | 0.817928479 | 8.179284786 |
| Dctd | 2.837946 | -0.125989389 | 0.763534927 | -0.440602365 | 0.65950089 | 0.817928479 | 8.179284786 |
| Kcnip4 | 2.6755905 | 0.138360044 | 0.785720063 | 0.450877869 | 0.652077574 | 0.817928479 | 8.179284786 |
| Lyrm1 | 2.6969721 | 0.129392335 | 0.819086932 | 0.446791249 | 0.65502579 | 0.817928479 | 8.179284786 |
| Dpyd | 5.592385 | -0.144269504 | 1.013834815 | 0.42939995 | 0.66763219 | 0.823413035 | 8.234130348 |
| Lrrc17 | 4.42784 | 0.12508561 | 0.695727568 | 0.398281726 | 0.690422529 | 0.846123469 | 8.461234692 |
| Sec61a1 | 52.64378 | 0.1188206 | 0.401966469 | 0.393881147 | 0.69366879 | 0.846123469 | 8.461234692 |
| Grik1 | 276.78488 | 0.040469456 | 0.110891211 | 0.374786148 | 0.707819517 | 0.853999635 | 8.539996348 |
| Hdac8 | 3.4485972 | 0.112414361 | 0.762342016 | 0.375253789 | 0.70747173 | 0.853999635 | 8.539996348 |
| Rps24 | 2.5770402 | -0.079878162 | 0.811494397 | -0.330515939 | 0.741010151 | 0.88681791 | 8.868179095 |
| Ppp1r16a | 5.2150073 | 0.101074318 | 0.704248783 | 0.327870473 | 0.7430096 | 0.88681791 | 8.868179095 |
| Dbi | 2.9943237 | -0.058628938 | 0.797861878 | -0.308326893 | 0.757833604 | 0.894888618 | 8.948886179 |
| Ctnna2 | 4.420046 | 0.096674266 | 0.705667609 | 0.310639542 | 0.756074663 | 0.894888618 | 8.948886179 |
| Trim2 | 11.130319 | -0.093035725 | 0.525705291 | -0.294730889 | 0.768199465 | 0.90232953 | 9.023295301 |
| Wdr72 | 8.42965 | 0.091030353 | 0.522996934 | 0.281315485 | 0.778468434 | 0.904816714 | 9.048167143 |
| Ptprg | 2.728707 | 0.079480528 | 0.819993952 | 0.283249236 | 0.776985788 | 0.904816714 | 9.048167143 |
| Wdyhv1 | 5.622881 | 0.088711233 | 0.626088408 | 0.275350058 | 0.783047309 | 0.905398451 | 9.053984514 |
| Zmat4 | 8.040158 | 0.084305676 | 0.542302848 | 0.261194834 | 0.793942261 | 0.908531866 | 9.085318662 |
| Sult2a2 | 3.2401161 | -0.144269504 | 1.01383476 | 0.263513358 | 0.792154929 | 0.908531866 | 9.085318662 |
| Kcnq5 | 2.5594702 | 0.059766357 | 0.842341304 | 0.227089888 | 0.820353847 | 0.919790677 | 9.197906772 |
| Cmss1 | 7.0652795 | -0.144269504 | 1.013834781 | -0.236358815 | 0.813154251 | 0.919790677 | 9.197906772 |
| Rabgap1l | 4.1028123 | -0.070222195 | 0.677806363 | -0.230387075 | 0.817791004 | 0.919790677 | 9.197906772 |
| Rpl11 | 2.9781857 | -0.040766887 | 0.824262793 | -0.239765884 | 0.810511756 | 0.919790677 | 9.197906772 |

| | | | | | | | |
|---|---|---|---|---|---|---|---|
| Lars2 | 2.271094 | -0.056109494 | 0.842521513 | -0.217762363 | 0.827614271 | 0.921355231 | 9.213552314 |
| Rsrc1 | 11.5518675 | 0.067506939 | 0.45731413 | 0.214637752 | 0.830049758 | 0.921355231 | 9.213552314 |
| Pcca | 3.9000213 | -0.144269504 | 1.013834806 | 0.195622119 | 0.844905946 | 0.923985819 | 9.239858188 |
| Gm11337 | 3.7218692 | -0.144269504 | 1.013834788 | 0.195622112 | 0.844905951 | 0.923985819 | 9.239858188 |
| Rps28 | 3.3896513 | 0.05695819 | 0.773224152 | 0.197046044 | 0.843791506 | 0.923985819 | 9.239858188 |
| Lsamp | 6.2088017 | 0.052272139 | 0.677457019 | 0.167664398 | 0.866847306 | 0.943333833 | 9.433338328 |
| Chsy3 | 6.558994 | -0.042799922 | 0.538872451 | -0.137738995 | 0.890446699 | 0.955102356 | 9.551023559 |
| Ctcflos | 4.9964113 | -0.043578911 | 0.670504532 | -0.144215711 | 0.885330135 | 0.955102356 | 9.551023559 |
| Rps15 | 2.5206728 | -0.029200757 | 0.827029859 | -0.132143644 | 0.894870676 | 0.955102356 | 9.551023559 |
| Alpl | 3.7230084 | 0.042741032 | 0.700067091 | 0.136163395 | 0.891692112 | 0.955102356 | 9.551023559 |
| Tacc1 | 2.534837 | 0.032991876 | 0.82380157 | 0.118893097 | 0.90536005 | 0.961674312 | 9.61674312 |
| Ptprd | 4.256398 | -0.026109951 | 0.742364727 | -0.095533128 | 0.923891378 | 0.96747116 | 9.674711604 |
| Gm49767 | 3.0391777 | 0.030062086 | 0.768444106 | 0.101594334 | 0.919078677 | 0.96747116 | 9.674711604 |
| Vwc2l | 5.2739124 | -0.029064877 | 0.645770054 | -0.096541521 | 0.923090499 | 0.96747116 | 9.674711604 |
| Gm29325 | 4.410524 | 0.022915334 | 0.675321283 | 0.071183658 | 0.943251588 | 0.967648255 | 9.676482548 |
| Pcgf6 | 4.2275147 | -0.144269504 | 1.013834795 | -0.057512431 | 0.954137004 | 0.967648255 | 9.676482548 |
| 4930558K0 | 7.5605907 | -0.144269504 | 1.013834783 | -0.074248328 | 0.940812792 | 0.967648255 | 9.676482548 |
| Cntn3 | 3.7232082 | -0.01774032 | 0.723535998 | -0.062387755 | 0.950254046 | 0.967648255 | 9.676482548 |
| 4930402H2 | 2.8789222 | -0.024046278 | 0.78297861 | -0.08927514 | 0.928863251 | 0.967648255 | 9.676482548 |
| Gm14582 | 4.4432654 | 0.023797474 | 0.65874642 | 0.073322144 | 0.941549771 | 0.967648255 | 9.676482548 |
| Chl1 | 3.8599033 | -0.144269504 | 1.013834817 | 0.056966442 | 0.954571927 | 0.967648255 | 9.676482548 |
| Gm50306 | 8.664798 | -0.012279509 | 0.576508881 | -0.041062319 | 0.967246215 | 0.975284892 | 9.752848917 |
| Tshz2 | 12.517045 | -0.144269504 | 1.013834793 | -0.036489919 | 0.970891716 | 0.975284892 | 9.752848917 |
| Ralyl | 3.6227722 | 0.007299317 | 0.730259878 | 0.022066823 | 0.982394652 | 0.982394652 | 9.823946518 |
| Aldoc | 0.9534373 | 0.054663947 | 1.031937517 | 0.226247259 | 0.821009116 | | |
| Anks1b | 1.8928432 | -0.144269504 | 1.013834769 | -0.412069579 | 0.680288426 | | |
| Arvcf | 1.2025291 | 0.431046629 | 1.32024377 | 1.157630342 | 0.247014921 | | |
| Atp6v0c | 0.93765974 | -0.175273185 | 0.75763061 | -1.288696337 | 0.197503676 | | |
| Atxn7l1 | 1.8535653 | -0.071954021 | 0.84456553 | -0.277250491 | 0.781587772 | | |
| Cadm2 | 1.4423465 | 0.373876736 | 1.403295967 | 1.20456604 | 0.228370873 | | |
| Csmd1 | 1.8290529 | 0.35813427 | 1.081174662 | 1.159057836 | 0.246432611 | | |
| Dlg2 | 1.6746006 | 0.082719365 | 0.899627468 | 0.313320836 | 0.754036921 | | |
| Dnm1 | 1.6511135 | -0.291463738 | 0.927896777 | -1.234057997 | 0.217181294 | | |
| Esf1 | 2.2412379 | -0.002455899 | 0.831967131 | -0.011451591 | 0.990863152 | | |
| Frmpd4 | 2.0246217 | -0.35157103 | 0.902590692 | -1.237119163 | 0.216042847 | | |
| Gas5 | 0.76831645 | 0.042887136 | 1.026553059 | 0.115295774 | 0.908210688 | | |
| Gm960 | 1.1630249 | 0.114839419 | 1.0327416 | 0.481205384 | 0.630370535 | | |
| Gnas | 1.4186062 | -0.144269504 | 1.01383474 | -1.888794722 | 0.05891934 | | |
| Gria1 | 1.5507308 | 0.247058201 | 1.035899837 | 0.898032503 | 0.369168222 | | |
| Grik2 | 1.1157385 | 0.129617796 | 1.055838613 | 0.536385316 | 0.591692277 | | |
| Hsp90ab1 | 1.6830295 | -0.293978094 | 0.833310665 | -1.498665499 | 0.133960431 | | |
| Kcnh7 | 1.598217 | 0.246098655 | 1.041320522 | 0.912380853 | 0.361568265 | | |
| Map2 | 1.5887762 | 0.029237739 | 0.919561253 | 0.13512543 | 0.892512704 | | |
| Mobp | 0.6598117 | -0.147470615 | 0.75911384 | -1.10993019 | 0.26702911 | | |
| Nlgn1 | 1.2868278 | -0.055102433 | 0.927094792 | -0.232888234 | 0.815848195 | | |
| Ppp3ca | 1.6195713 | 0.009043024 | 0.911990951 | 0.037049204 | 0.970445774 | | |
| Rpl13a | 2.187962 | -0.222936858 | 0.839717857 | -0.858682076 | 0.390515942 | | |
| Rpl24 | 0.5970339 | -0.010088571 | 0.984333998 | -0.171560383 | 0.863783156 | | |
| Rpl34 | 2.083867 | -0.212230718 | 0.830102686 | -0.928084544 | 0.353363716 | | |
| Rpl36 | 0.41904178 | 0.055950164 | 1.741083361 | 0.237755896 | 0.812070428 | | |
| Rpl39 | 1.1329759 | -0.019663536 | 0.885010274 | -0.117349777 | 0.906582881 | | |
| Rpl6 | 1.2310191 | 0.116491332 | 1.05005885 | 0.542021719 | 0.587803544 | | |
| Rps10 | 0.543171 | 0.172779122 | 1.344757683 | 0.548748629 | 0.583177969 | | |
| Rps12 | 1.5295702 | -0.095163128 | 0.843822889 | -0.421508594 | 0.673383737 | | |
| Rps23 | 1.4581437 | -0.044118603 | 0.873608571 | -0.260189897 | 0.794717297 | | |
| Rps25 | 0.8295014 | -0.144269504 | 1.013834785 | 1.036147219 | 0.300133466 | | |
| Rps27a | 1.8988329 | 0.066366306 | 0.918337919 | 0.284545337 | 0.775992496 | | |
| Rps3 | 2.0621865 | -0.377409486 | 0.929132844 | -1.379988755 | 0.167590107 | | |
| Selenow | 2.180446 | -0.049698299 | 0.835985919 | -0.18901031 | 0.850084736 | | |
| Sgip1 | 1.8380804 | -0.117037264 | 0.854294357 | -0.488695091 | 0.62505758 | | |
| Slc1a2 | 2.146905 | -0.311668237 | 0.874074761 | -1.353810575 | 0.175796819 | | |
| Snap25 | 1.882255 | -0.477869997 | 1.022921022 | -1.860619426 | 0.06279794 | | |
| Ttc3 | 2.0692492 | -0.111158015 | 0.855411152 | -0.484313467 | 0.62816342 | | |
| Tuba1a | 1.1539501 | -0.068458882 | 0.894017692 | -0.347584813 | 0.728152013 | | |
| Ubb | 0.76741 | -0.010357733 | 0.983352815 | -0.088526801 | 0.929457984 | | |

| **Glut** | baseMean | log2FoldChange | lfcSE | stat | pvalue | padj | pglobadj |
|---|---|---|---|---|---|---|---|

| | | | | | | | |
|---|---|---|---|---|---|---|---|
| mt-Nd4 | 99.5293 | -0.144269504 | 0.891945384 | -11.48267411 | 1.61E-30 | 4.24E-28 | 4.24E-27 |
| AY036118 | 115.76752 | -2.042013454 | 0.22819175 | -9.152206257 | 5.58E-20 | 7.34E-18 | 7.34E-17 |
| Camk1d | 86.15781 | -2.057596879 | 0.234726051 | -8.966734033 | 3.05E-19 | 2.68E-17 | 2.68E-16 |
| mt-Co3 | 321.70468 | -1.222299868 | 0.139653457 | -8.889949245 | 6.11E-19 | 4.02E-17 | 4.02E-16 |
| mt-Co1 | 172.39795 | -1.331149312 | 0.159880937 | -8.491984216 | 2.03E-17 | 1.07E-15 | 1.07E-14 |
| Rpl37 | 43.836258 | -0.144269504 | 0.891945536 | -7.217753126 | 5.29E-13 | 2.32E-11 | 2.32E-10 |
| mt-Nd5 | 25.289469 | -0.144269504 | 0.89194546 | -7.013216025 | 2.33E-12 | 8.75E-11 | 8.75E-10 |
| mt-Atp6 | 276.5879 | -1.152393284 | 0.169133456 | -6.975217917 | 3.05E-12 | 1.00E-10 | 1.00E-09 |
| Gm42418 | 533.4941 | -1.223408772 | 0.185889278 | -6.762517509 | 1.36E-11 | 3.96E-10 | 3.96E-09 |
| mt-Nd1 | 68.56083 | -1.038355427 | 0.15905101 | -6.679920834 | 2.39E-11 | 5.77E-10 | 5.77E-09 |
| Lrp1b | 220.60295 | 0.866387785 | 0.131749505 | 6.678615153 | 2.41E-11 | 5.77E-10 | 5.77E-09 |
| 1700054A( | 141.31175 | 0.895061055 | 0.139041507 | 6.555166783 | 5.56E-11 | 1.22E-09 | 1.22E-08 |
| Mbp | 64.30713 | -2.625254026 | 0.423452389 | -6.535964044 | 6.32E-11 | 1.28E-09 | 1.28E-08 |
| Atp1b1 | 13.367782 | -0.144269504 | 0.891945597 | -6.458741211 | 1.06E-10 | 1.98E-09 | 1.98E-08 |
| Gm12216 | 94.09649 | -0.144269504 | 0.891945591 | 6.396879725 | 1.59E-10 | 2.78E-09 | 2.78E-08 |
| Hsp90ab1 | 15.209404 | -2.7435117 | 0.453992189 | -6.327891765 | 2.49E-10 | 4.09E-09 | 4.09E-08 |
| Uqcrh | 13.775299 | -0.144269504 | 0.891945599 | -6.035781097 | 1.58E-09 | 2.45E-08 | 2.45E-07 |
| mt-Cytb | 112.09964 | -1.193617113 | 0.206249698 | -5.991378724 | 2.08E-09 | 3.04E-08 | 3.04E-07 |
| Xlr5a | 106.50605 | 1.005800245 | 0.17464977 | 5.908686377 | 3.45E-09 | 4.77E-08 | 4.77E-07 |
| 5830416I1 | 123.41928 | 0.972715859 | 0.169083824 | 5.898841839 | 3.66E-09 | 4.81E-08 | 4.81E-07 |
| mt-Co2 | 178.44832 | -1.040134994 | 0.187754659 | -5.721011729 | 1.06E-08 | 1.33E-07 | 1.33E-06 |
| 8430422H( | 44.64208 | -0.144269504 | 0.891945643 | 5.69813214 | 1.21E-08 | 1.45E-07 | 1.45E-06 |
| Cpe | 16.194384 | -0.144269504 | 0.891945594 | -5.606676682 | 2.06E-08 | 2.36E-07 | 2.36E-06 |
| Cox4i1 | 36.39977 | -1.652949358 | 0.319383433 | -5.480192085 | 4.25E-08 | 4.66E-07 | 4.66E-06 |
| Gnaz | 43.537 | -0.144269504 | 0.891945653 | 5.304093886 | 1.13E-07 | 1.19E-06 | 1.19E-05 |
| Rapgef4 | 53.225666 | -0.144269504 | 0.891945603 | 5.287650776 | 1.24E-07 | 1.25E-06 | 1.25E-05 |
| Rplp1 | 50.36301 | -1.078511026 | 0.213834809 | -5.244553134 | 1.57E-07 | 1.53E-06 | 1.53E-05 |
| Kctd16 | 82.1713 | 0.775286911 | 0.154983548 | 5.121284583 | 3.03E-07 | 2.85E-06 | 2.85E-05 |
| Fth1 | 122.870445 | -1.694112248 | 0.356098672 | -5.097208166 | 3.45E-07 | 3.13E-06 | 3.13E-05 |
| Mc1r | 43.232273 | -0.144269504 | 0.89194565 | 5.045752725 | 4.52E-07 | 3.96E-06 | 3.96E-05 |
| Rsrc1 | 47.870052 | -0.144269504 | 0.89194561 | 5.031937446 | 4.86E-07 | 4.12E-06 | 4.12E-05 |
| Rps14 | 26.258608 | -0.144269504 | 0.891945533 | -4.998881401 | 5.77E-07 | 4.74E-06 | 4.74E-05 |
| Gphn | 18.18605 | -1.600163523 | 0.351495321 | -4.878390166 | 1.07E-06 | 8.52E-06 | 8.52E-05 |
| Hexb | 12.893745 | -2.228591148 | 0.500595783 | -4.852711441 | 1.22E-06 | 9.42E-06 | 9.42E-05 |
| Apoe | 54.08286 | -0.982628035 | 0.212570666 | -4.826108628 | 1.39E-06 | 1.05E-05 | 0.000104619 |
| Car12 | 12.542942 | -2.113185317 | 0.478823004 | -4.803004877 | 1.56E-06 | 1.14E-05 | 0.000114187 |
| Tpt1 | 22.281645 | -1.584128917 | 0.355306863 | -4.789547637 | 1.67E-06 | 1.19E-05 | 0.000118818 |
| Sec61a1 | 169.02109 | 0.478586323 | 0.102670013 | 4.709273104 | 2.49E-06 | 1.72E-05 | 0.000172059 |
| mt-Nd4l | 10.935973 | -0.144269504 | 0.891945606 | -4.671013008 | 3.00E-06 | 2.02E-05 | 0.000202117 |
| Rps20 | 18.792002 | -0.144269504 | 0.891945574 | -4.661003376 | 3.15E-06 | 2.07E-05 | 0.000206897 |
| Adamtsl1 | 34.275814 | -0.144269504 | 0.891945652 | 4.627800908 | 3.70E-06 | 2.37E-05 | 0.000237065 |
| Rpl35a | 25.377203 | -0.144269504 | 0.891945577 | -4.559430891 | 5.13E-06 | 3.20E-05 | 0.000320138 |
| Calm2 | 13.950143 | -1.616887216 | 0.3851873 | -4.555174814 | 5.23E-06 | 3.20E-05 | 0.000320138 |
| Cox7c | 21.289083 | -1.465670227 | 0.350267879 | -4.515252322 | 6.32E-06 | 3.78E-05 | 0.000378011 |
| Shb | 62.057114 | -0.144269504 | 0.891945594 | 4.460837288 | 8.16E-06 | 4.77E-05 | 0.000477141 |
| Ttc3 | 11.944229 | -2.230393038 | 0.571856325 | -4.361849929 | 1.29E-05 | 7.37E-05 | 0.000737357 |
| Cox6b1 | 9.769847 | -2.035401755 | 0.521824353 | -4.322546563 | 1.54E-05 | 8.63E-05 | 0.000863079 |
| Pde1c | 48.71622 | 1.198608307 | 0.29752159 | 4.301763151 | 1.69E-05 | 9.28E-05 | 0.000928414 |
| Mobp | 13.428116 | -1.880075786 | 0.493224328 | -4.254555473 | 2.09E-05 | 0.000112427 | 0.00112427 |
| Rpl6 | 14.984661 | -0.144269504 | 0.891945621 | -4.243190797 | 2.20E-05 | 0.000115911 | 0.001159114 |
| Rps8 | 25.638395 | -0.144269504 | 0.891945503 | -4.203463628 | 2.63E-05 | 0.000135554 | 0.00135554 |
| Ms4a1 | 30.989298 | -0.144269504 | 0.891945649 | 4.164298014 | 3.12E-05 | 0.000157958 | 0.001579578 |
| 4930511M | 40.240997 | -0.144269504 | 0.891945629 | 4.159743013 | 3.19E-05 | 0.000158101 | 0.001581006 |
| Sema3c | 71.05895 | 0.806801401 | 0.202954021 | 4.134280669 | 3.56E-05 | 0.000173418 | 0.001734184 |
| Pde6a | 65.21776 | -0.144269504 | 0.891945574 | 4.06651906 | 4.77E-05 | 0.000224116 | 0.002241163 |
| Rpl39 | 22.424913 | -1.850695023 | 0.515243296 | -4.06806863 | 4.74E-05 | 0.000224116 | 0.002241163 |
| mt-Nd2 | 106.08404 | -0.886036125 | 0.232423428 | -4.023013534 | 5.75E-05 | 0.000265114 | 0.002651141 |
| Wdyhv1 | 16.211117 | 1.919284527 | 0.425653252 | 4.008899121 | 6.10E-05 | 0.000272079 | 0.002720788 |
| Cmss1 | 60.782124 | -1.296824649 | 0.354101217 | -4.008766706 | 6.10E-05 | 0.000272079 | 0.002720788 |
| Lars2 | 19.149841 | -1.450317926 | 0.403818869 | -3.975275254 | 7.03E-05 | 0.000308139 | 0.003081391 |
| Tuba1a | 8.8383875 | -0.144269504 | 0.891945628 | -3.946714462 | 7.92E-05 | 0.000341602 | 0.003416023 |
| Nckap5 | 102.8312 | 0.885512478 | 0.237335617 | 3.936966278 | 8.25E-05 | 0.000350037 | 0.003500369 |
| Fam169a | 25.7768 | -0.144269504 | 0.891945638 | 3.825267497 | 0.00013063 | 0.000545329 | 0.005453291 |
| Ppia | 16.155235 | -1.520143377 | 0.453359177 | -3.791116586 | 0.000149972 | 0.000616289 | 0.006162893 |
| Rpl9 | 16.844284 | -1.223319612 | 0.356245969 | -3.778353828 | 0.000157868 | 0.00063876 | 0.006387601 |
| Cox8a | 38.321594 | -1.800353626 | 0.561199488 | -3.736512418 | 0.00018659 | 0.000722728 | 0.007227281 |
| Slx1b | 29.005318 | -0.144269504 | 0.891945644 | 3.736142153 | 0.000186865 | 0.000722728 | 0.007227281 |

| | | | | | | | |
|---|---|---|---|---|---|---|---|
| Gm10062 | 67.16049 | 0.85745273 | 0.242536054 | 3.738178707 | 0.000185358 | 0.000722728 | 0.007227281 |
| Rpl23 | 18.408915 | -1.340934939 | 0.409612868 | -3.676409649 | 0.00023654 | 0.000901593 | 0.009015929 |
| Ctcflos | 17.103666 | 1.180064174 | 0.344290186 | 3.659618485 | 0.000252591 | 0.000949021 | 0.009490206 |
| Gm38973 | 14.194267 | -0.144269504 | 0.89194568 | 3.65026491 | 0.00026197 | 0.000970396 | 0.009703959 |
| Rps27a | 23.433489 | -1.597478381 | 0.50781038 | -3.636561386 | 0.000276302 | 0.001009269 | 0.010092692 |
| Nrxn3 | 15.641308 | -1.61420841 | 0.514470802 | -3.630033881 | 0.000283384 | 0.001020959 | 0.010209589 |
| Ehbp1 | 37.673763 | 0.909778193 | 0.278224719 | 3.499943805 | 0.000465356 | 0.001653901 | 0.016539013 |
| Gm48099 | 14.693597 | -0.144269504 | 0.891945519 | -3.420148949 | 0.000625869 | 0.002194712 | 0.021947124 |
| mt-Nd3 | 53.351776 | -1.009592416 | 0.325249544 | -3.413089624 | 0.000642308 | 0.002222724 | 0.022227243 |
| Ttc28 | 28.257631 | -0.144269504 | 0.891945598 | 3.365379564 | 0.000764384 | 0.002610819 | 0.026108191 |
| 4930558K0 | 20.962511 | -0.144269504 | 0.891945643 | 3.330643367 | 0.000866455 | 0.002884528 | 0.028845285 |
| Rpl12 | 7.4294677 | -0.144269504 | 0.891945627 | -3.332238785 | 0.000861503 | 0.002884528 | 0.028845285 |
| Chsy3 | 24.266058 | -0.144269504 | 0.891945628 | 3.310057046 | 0.00093277 | 0.00306648 | 0.0306648 |
| Mt3 | 22.449446 | -1.303936672 | 0.457093921 | -3.299182843 | 0.000969667 | 0.003148426 | 0.031484259 |
| 6030443J0 | 35.326588 | 0.699177578 | 0.22339533 | 3.273065574 | 0.001063878 | 0.003412194 | 0.034121938 |
| Ntm | 72.91777 | 0.487859304 | 0.153831448 | 3.255441988 | 0.00113216 | 0.00354474 | 0.035447398 |
| Rpl26 | 22.922571 | -0.965640885 | 0.327845932 | -3.258181544 | 0.001121286 | 0.00354474 | 0.035447398 |
| Svil | 12.984544 | -0.144269504 | 0.891945659 | 3.250461845 | 0.001152177 | 0.003564972 | 0.035649721 |
| Rpl21 | 29.280321 | -1.317407826 | 0.476965327 | -3.229345276 | 0.00124074 | 0.003733226 | 0.03733226 |
| Uba52 | 9.639106 | -1.449169837 | 0.534247806 | -3.223695991 | 0.001265477 | 0.003733226 | 0.03733226 |
| Rps25 | 15.280683 | -1.406388631 | 0.516313013 | -3.228252639 | 0.001245489 | 0.003733226 | 0.03733226 |
| Prkg1 | 36.00238 | 0.821088803 | 0.273346743 | 3.221532402 | 0.00127507 | 0.003733226 | 0.03733226 |
| Rps29 | 83.45581 | -1.161764643 | 0.411788024 | -3.220980188 | 0.00127753 | 0.003733226 | 0.03733226 |
| Pcca | 11.026051 | -0.144269504 | 0.891945657 | 3.187381484 | 0.001435673 | 0.004104152 | 0.041041516 |
| Gm19951 | 11.873416 | -1.175828963 | 0.422859049 | -3.18896354 | 0.001427839 | 0.004104152 | 0.041041516 |
| Ubb | 11.435586 | -1.135869605 | 0.410338658 | -3.16847739 | 0.001532397 | 0.00428745 | 0.042874503 |
| Camk2n1 | 8.961582 | -0.144269504 | 0.891945632 | -3.171411879 | 0.001516999 | 0.00428745 | 0.042874503 |
| Cdk8 | 20.925909 | -0.931230227 | 0.32770643 | -3.14899814 | 0.001638312 | 0.004535538 | 0.045355385 |
| Rps24 | 30.843369 | -1.284757141 | 0.48379064 | -3.136696027 | 0.001708632 | 0.004680939 | 0.04680939 |
| Spock1 | 13.501007 | 1.123478532 | 0.394701413 | 3.131307141 | 0.0017403 | 0.004718547 | 0.047185467 |
| Snap25 | 13.494748 | -1.583110177 | 0.634171554 | -3.105040023 | 0.001902533 | 0.005105777 | 0.051057768 |
| Acoxl | 23.2717 | -0.144269504 | 0.89194565 | 3.098784781 | 0.001943161 | 0.005162135 | 0.051621353 |
| Scamp1 | 19.733292 | -0.144269504 | 0.891945652 | 3.073482982 | 0.002115758 | 0.005564442 | 0.055644424 |
| Vit | 14.803189 | 1.152476701 | 0.422083533 | 3.06307871 | 0.002190724 | 0.005704559 | 0.057045593 |
| Il27 | 32.86697 | 0.782929514 | 0.275467979 | 3.058949763 | 0.002221144 | 0.005727067 | 0.057270672 |
| Hapln1 | 15.937945 | 1.085780092 | 0.395974166 | 3.055519551 | 0.00224671 | 0.005736744 | 0.057367438 |
| Rps10 | 17.40676 | -0.882757147 | 0.320959867 | -3.051316897 | 0.0022784 | 0.005761722 | 0.057617222 |
| Calm1 | 23.249405 | -1.233356512 | 0.483984449 | -3.023250262 | 0.002500753 | 0.006263792 | 0.062637919 |
| Rpl36 | 15.970733 | -1.236115769 | 0.49053453 | -3.00539333 | 0.002652376 | 0.006578559 | 0.065785588 |
| Sil1 | 30.168123 | -0.144269504 | 0.891945645 | 3.002644604 | 0.002676448 | 0.006578559 | 0.065785588 |
| Aldoc | 11.45493 | -1.195309886 | 0.471719017 | -2.988907112 | 0.002799772 | 0.006694001 | 0.066940011 |
| Rpl13 | 28.522684 | -1.173012025 | 0.461912119 | -2.993771551 | 0.002755522 | 0.006694001 | 0.066940011 |
| Epha6 | 42.808353 | 0.737301721 | 0.264924868 | 2.989387282 | 0.002795376 | 0.006694001 | 0.066940011 |
| Hdac8 | 15.433358 | 1.112926503 | 0.421615247 | 2.981103802 | 0.002872114 | 0.006805099 | 0.068050995 |
| Mapk14 | 43.059185 | 0.750816865 | 0.271655871 | 2.977487949 | 0.002906211 | 0.006824405 | 0.068244055 |
| Prkag2 | 19.312868 | -0.144269504 | 0.891945648 | 2.954420731 | 0.003132566 | 0.00729084 | 0.072908399 |
| Zmat4 | 23.255938 | 0.934905791 | 0.354239928 | 2.937464591 | 0.00330908 | 0.007634105 | 0.076341047 |
| 1700126G0 | 34.913403 | 0.632148083 | 0.229946393 | 2.909610024 | 0.0036188 | 0.008276038 | 0.08276038 |
| Psap | 9.908793 | -1.351992256 | 0.577494571 | -2.90408508 | 0.003683281 | 0.008350886 | 0.08350886 |
| Eef1a1 | 20.31284 | -1.069399737 | 0.43461205 | -2.882858859 | 0.00394084 | 0.008858469 | 0.088584688 |
| Csmd3 | 27.648584 | 0.768709746 | 0.297988304 | 2.809197864 | 0.004966511 | 0.011069428 | 0.110694275 |
| Nell1 | 25.19159 | -0.144269504 | 0.891945643 | 2.773402779 | 0.005547341 | 0.01226009 | 0.122600905 |
| Gm49497 | 15.45173 | -0.144269504 | 0.891945651 | 2.764697564 | 0.005697559 | 0.012282444 | 0.122824435 |
| 4930441J1 | 14.683736 | 0.912403285 | 0.366200877 | 2.764973009 | 0.005692751 | 0.012282444 | 0.122824435 |
| Grik1 | 5.603626 | 2.077191617 | 0.902970322 | 2.766396053 | 0.005667965 | 0.012282444 | 0.122824435 |
| Rps23 | 20.041182 | -0.962068375 | 0.404906221 | -2.759895081 | 0.005781993 | 0.012363123 | 0.123631227 |
| Atp6v0c | 12.940946 | -1.008638538 | 0.43052226 | -2.750306416 | 0.005953956 | 0.012597589 | 0.125975886 |
| Gm11337 | 14.409234 | 0.895170145 | 0.358921577 | 2.748468047 | 0.005987447 | 0.012597589 | 0.125975886 |
| Ctnna3 | 25.755857 | 0.825254275 | 0.334450273 | 2.742777074 | 0.006092203 | 0.012716265 | 0.127162648 |
| Ninj2 | 19.79367 | 0.862152164 | 0.350544847 | 2.740135864 | 0.006141379 | 0.012717974 | 0.127179742 |
| Fbxl7 | 17.824291 | 1.0169746 | 0.434344531 | 2.709824985 | 0.006731872 | 0.013831893 | 0.138318932 |
| Zbtb7a | 27.59445 | 0.742921506 | 0.300984577 | 2.702374526 | 0.006884616 | 0.014036078 | 0.140360784 |
| Rpl32 | 21.320887 | -0.888365808 | 0.378502645 | -2.697945077 | 0.006976895 | 0.014114795 | 0.141147954 |
| Wdr72 | 24.990583 | 0.842240518 | 0.351886534 | 2.68459577 | 0.007261758 | 0.014578948 | 0.145789483 |
| Rpl27a | 24.33323 | -0.951546339 | 0.41477329 | -2.677873038 | 0.00740913 | 0.01476213 | 0.147621296 |
| Lrrc17 | 21.825884 | 0.647707455 | 0.274624817 | 2.670850417 | 0.007565935 | 0.014961211 | 0.149612107 |
| Gm29325 | 15.722427 | 0.928653144 | 0.396519703 | 2.664873892 | 0.00770172 | 0.015116062 | 0.15116062 |

| | | | | | | | |
|---|---|---|---|---|---|---|---|
| Ptgds | 74.75537 | 0.905495613 | 0.393790669 | 2.656104942 | 0.007904901 | 0.015399918 | 0.153999184 |
| Rpl11 | 31.186632 | -1.213310185 | 0.588032536 | -2.641707471 | 0.008248927 | 0.015951969 | 0.159519687 |
| Ppp1r16a | 15.081142 | -0.144269504 | 0.891945638 | 2.630754065 | 0.008519566 | 0.01635508 | 0.163550799 |
| Pcdh15 | 29.29712 | 0.587261791 | 0.237757708 | 2.6282401 | 0.008582791 | 0.016357059 | 0.163570588 |
| Rps17 | 11.7198 | -1.068558408 | 0.507155574 | -2.589173779 | 0.009620653 | 0.018203106 | 0.18203106 |
| Phkg1 | 13.678538 | 0.978343294 | 0.445206368 | 2.562077728 | 0.010404803 | 0.019546166 | 0.195461655 |
| Rps15 | 16.84303 | -0.938378533 | 0.43715906 | -2.552660908 | 0.010690352 | 0.019902647 | 0.199026469 |
| Cmpk1 | 54.895275 | 0.620151972 | 0.262283034 | 2.550854563 | 0.010745916 | 0.019902647 | 0.199026469 |
| Gria4 | 31.359358 | 0.668879563 | 0.290517594 | 2.519953001 | 0.01173705 | 0.021586324 | 0.215863236 |
| Cdh23 | 31.59858 | 0.65907498 | 0.285996253 | 2.514008307 | 0.011936762 | 0.021781625 | 0.217816254 |
| Fau | 21.339561 | -1.099230598 | 0.558680719 | -2.511883255 | 0.012008881 | 0.021781625 | 0.217816254 |
| Tns3 | 12.934249 | 0.938224438 | 0.440247108 | 2.491748287 | 0.012711609 | 0.022898309 | 0.228983089 |
| St18 | 4.6962943 | 1.731587181 | 0.886190397 | 2.487528495 | 0.012863415 | 0.023014137 | 0.230141367 |
| AI314278 | 10.076811 | 1.017587739 | 0.488240231 | 2.469631382 | 0.013525234 | 0.023867956 | 0.238679558 |
| Plp1 | 7.740564 | -1.337646557 | 0.776163083 | -2.471210614 | 0.013465649 | 0.023867956 | 0.238679558 |
| Ppp1r9a | 17.912962 | 0.8388833 | 0.390846774 | 2.46493907 | 0.013703655 | 0.023867956 | 0.238679558 |
| Ptpn9 | 15.364464 | 0.895427309 | 0.42248737 | 2.466428577 | 0.013646794 | 0.023867956 | 0.238679558 |
| Dpyd | 18.939875 | 0.793621073 | 0.366556326 | 2.458876585 | 0.013937252 | 0.023957499 | 0.239574985 |
| Rpl13a | 19.485212 | -1.102251171 | 0.580967768 | -2.460541742 | 0.013872743 | 0.023957499 | 0.239574985 |
| Mrc2 | 15.145778 | 0.872599675 | 0.41538329 | 2.436338573 | 0.01483679 | 0.025338155 | 0.253381545 |
| Tmem108 | 22.477928 | -0.144269504 | 0.891945623 | 2.428526141 | 0.015160332 | 0.02572366 | 0.257236604 |
| Klkb1 | 12.750455 | 0.830746798 | 0.390567129 | 2.42459785 | 0.015325353 | 0.025836973 | 0.258369729 |
| Phactr1 | 9.087869 | -0.922545442 | 0.465997064 | -2.41313806 | 0.015815827 | 0.026494029 | 0.264940294 |
| Dlg2 | 17.788498 | -0.929038657 | 0.472075291 | -2.405306982 | 0.016158886 | 0.026897385 | 0.268973854 |
| Galnt13 | 35.5822 | -0.144269504 | 0.891945625 | 2.392846203 | 0.016718247 | 0.027653453 | 0.276534531 |
| Ctnna2 | 16.06405 | 0.756595146 | 0.357115202 | 2.384282517 | 0.017112459 | 0.028128605 | 0.281286052 |
| Ppp3ca | 13.0466585 | -1.015608564 | 0.556984907 | -2.349981317 | 0.018774353 | 0.030668664 | 0.306686642 |
| Gas5 | 24.85064 | -1.278483481 | 0.870083465 | -2.327956837 | 0.019914396 | 0.03220341 | 0.322034104 |
| Rab29 | 16.764362 | -0.144269504 | 0.89194563 | 2.327122098 | 0.019958768 | 0.03220341 | 0.322034104 |
| Trim2 | 50.253483 | 0.419903518 | 0.190052652 | 2.318993808 | 0.020395372 | 0.032707213 | 0.327072125 |
| Tshz2 | 5.361759 | 1.256399967 | 0.697787426 | 2.307943669 | 0.02100227 | 0.033476345 | 0.334763451 |
| Rpl37a | 30.679293 | -1.086546512 | 0.672319371 | -2.269754726 | 0.023222469 | 0.036792225 | 0.367922249 |
| Cox6c | 20.9289 | -0.977922186 | 0.567781926 | -2.261381875 | 0.02373562 | 0.037380049 | 0.373800485 |
| Anks1b | 9.369207 | -0.739896001 | 0.39404025 | -2.223444396 | 0.026185849 | 0.040993323 | 0.409933229 |
| Dctd | 8.537931 | -0.144269504 | 0.891945649 | 2.21483304 | 0.026771526 | 0.041662198 | 0.41662198 |
| Nlgn1 | 8.871985 | -0.974131063 | 0.60704465 | -2.169432581 | 0.030049858 | 0.046488897 | 0.464888974 |
| Cspp1 | 17.172728 | 0.761839128 | 0.417467866 | 2.158028822 | 0.030925591 | 0.047563921 | 0.475639207 |
| Syt1 | 14.075573 | -0.876829274 | 0.525156125 | -2.139970621 | 0.032357141 | 0.049476326 | 0.494763262 |
| Dop1b | 24.097761 | 0.664554743 | 0.363968221 | 2.098212754 | 0.035886355 | 0.054555558 | 0.545555577 |
| D430042O | 61.552265 | 0.346398453 | 0.173233285 | 2.084426136 | 0.037121423 | 0.056108817 | 0.561088168 |
| Vwc2l | 9.925723 | 0.820886801 | 0.483183731 | 2.07970589 | 0.037552517 | 0.056436069 | 0.564360691 |
| Rps12 | 15.100746 | -0.721004519 | 0.425677623 | -2.046427525 | 0.040714326 | 0.060840158 | 0.608401581 |
| Zfhx3 | 15.366595 | 0.693580815 | 0.403079825 | 2.02569262 | 0.042796293 | 0.063232726 | 0.632327257 |
| Rpl24 | 14.56541 | -0.980568499 | 0.744319084 | -2.025948176 | 0.042770096 | 0.063232726 | 0.632327257 |
| Rpl30 | 24.25163 | -0.876275309 | 0.592764932 | -2.019107479 | 0.04347605 | 0.063878218 | 0.638782183 |
| Rps27 | 25.235474 | -0.739403635 | 0.45403204 | -2.013224469 | 0.04409102 | 0.064421879 | 0.644218794 |
| Gm50306 | 36.891846 | -0.144269504 | 0.891945612 | 2.005008319 | 0.044962158 | 0.065331754 | 0.65331754 |
| Gm49767 | 10.397022 | -0.144269504 | 0.891945644 | 1.987552835 | 0.046861167 | 0.067346923 | 0.673469231 |
| Ckb | 24.891195 | -0.806976445 | 0.528848248 | -1.989645223 | 0.046630031 | 0.067346923 | 0.673469231 |
| Rpl38 | 27.752401 | -0.653643477 | 0.399427351 | -1.951269177 | 0.051025034 | 0.072932521 | 0.729325209 |
| Chl1 | 14.471017 | 0.589061434 | 0.352504148 | 1.90931001 | 0.056222112 | 0.079496857 | 0.794968569 |
| Tmsb4x | 25.429573 | -0.763271174 | 0.521893082 | -1.910000573 | 0.05613314 | 0.079496857 | 0.794968569 |
| Rps7 | 13.783592 | -0.773996002 | 0.53911058 | -1.90608746 | 0.056638859 | 0.079657861 | 0.796578611 |
| Gm14582 | 14.14446 | 0.678359331 | 0.440521883 | 1.878649337 | 0.06029239 | 0.084317728 | 0.843177281 |
| Cntnap2 | 63.37012 | 0.4024131 | 0.229795044 | 1.87645121 | 0.060593348 | 0.084317728 | 0.843177281 |
| Zfp994 | 14.614243 | 0.624403245 | 0.398196093 | 1.85010322 | 0.064298674 | 0.089002901 | 0.890029013 |
| Adk | 18.05157 | 0.494094758 | 0.302610546 | 1.81163532 | 0.070042564 | 0.096446043 | 0.964460431 |
| Esf1 | 10.646127 | 0.609939449 | 0.407154703 | 1.776009779 | 0.075731305 | 0.103736111 | 1.037361109 |
| Cst3 | 61.025986 | -0.440348791 | 0.273082367 | -1.772098354 | 0.076378232 | 0.104080182 | 1.040801816 |
| Rpl41 | 30.727098 | -0.651455335 | 0.477539018 | -1.738769086 | 0.082075388 | 0.11126715 | 1.112671501 |
| Rpl17 | 13.341359 | -0.697864318 | 0.544699746 | -1.728918799 | 0.083823629 | 0.113054433 | 1.130544331 |
| Dnm1 | 6.8677077 | -0.683591564 | 0.532750063 | -1.72613599 | 0.08432295 | 0.113147633 | 1.131476325 |
| Igf2bp3 | 52.856205 | 0.621565906 | 0.456837276 | 1.705135346 | 0.088169185 | 0.1177081 | 1.177080996 |
| Dbi | 14.339963 | -0.529111907 | 0.37753547 | -1.664480218 | 0.096016501 | 0.127537069 | 1.275370695 |
| Cntn3 | 12.479238 | -0.144269504 | 0.891945638 | 1.632108691 | 0.102656591 | 0.135671776 | 1.356717765 |
| Atp9b | 26.291191 | 0.585529917 | 0.456860704 | 1.613525606 | 0.106630365 | 0.139843745 | 1.39843745 |
| Map2 | 8.605354 | -0.568165626 | 0.445343346 | -1.612391476 | 0.106876779 | 0.139843745 | 1.39843745 |

| | | | | | | | |
|---|---|---|---|---|---|---|---|
| Selenow | 12.75893 | -0.660120646 | 0.576810332 | -1.605946848 | 0.108285589 | 0.140985692 | 1.409856924 |
| Rps21 | 54.943985 | -0.594774582 | 0.493490246 | -1.572159317 | 0.115913608 | 0.150173787 | 1.501737874 |
| Rtn1 | 11.511458 | -0.648770608 | 0.812432881 | -1.490533373 | 0.136084048 | 0.17544169 | 1.754416895 |
| Mt1 | 28.332375 | -0.602436688 | 0.585546772 | -1.478418886 | 0.139295693 | 0.178706182 | 1.787061823 |
| Slc1a2 | 12.561978 | -0.624404766 | 0.683992766 | -1.460163431 | 0.144245163 | 0.18415766 | 1.841576596 |
| Ndrg4 | 12.235044 | -0.579644097 | 0.558976319 | -1.449322622 | 0.147247506 | 0.186636238 | 1.866362377 |
| Cadm2 | 13.261956 | -0.51729154 | 0.455988287 | -1.448040065 | 0.147605846 | 0.186636238 | 1.866362377 |
| Grik2 | 6.7209067 | -0.558168643 | 0.542923854 | -1.430309073 | 0.152628332 | 0.192063403 | 1.920634033 |
| Tacc1 | 10.839199 | 0.497372738 | 0.444532678 | 1.400768547 | 0.161283297 | 0.201988129 | 2.019881291 |
| Ptprd | 15.735741 | 0.52227039 | 0.490510781 | 1.394989553 | 0.16301899 | 0.203194286 | 2.03194286 |
| Atxn7l1 | 14.766819 | 0.449715248 | 0.390349486 | 1.381987894 | 0.166975418 | 0.207144033 | 2.071440332 |
| Ahdc1 | 12.1700945 | 0.485398867 | 0.441955887 | 1.376973655 | 0.168520393 | 0.208079171 | 2.080791707 |
| Gm765 | 14.451148 | 0.412698237 | 0.35820877 | 1.348003862 | 0.17765714 | 0.218335644 | 2.183356445 |
| Frmpd4 | 9.483613 | -0.516838105 | 0.53914265 | -1.332441029 | 0.182715307 | 0.223507562 | 2.235075619 |
| Pcgf6 | 13.5450325 | -0.144269504 | 0.891945621 | 1.310054143 | 0.19017752 | 0.23155874 | 2.315587397 |
| Pip5k1b | 10.7171335 | 0.453312736 | 0.444104114 | 1.285401446 | 0.198652036 | 0.240762606 | 2.407626062 |
| Unc5d | 1.757701 | 0.597398564 | 0.863912873 | 1.277429639 | 0.201450606 | 0.242273996 | 2.42273996 |
| Cfap74 | 11.383468 | -0.144269504 | 0.891945647 | 1.276605768 | 0.201741464 | 0.242273996 | 2.42273996 |
| Meg3 | 32.51984 | -0.445186258 | 0.467366537 | -1.231936731 | 0.217972719 | 0.259461931 | 2.594619313 |
| Rps3a1 | 17.534512 | -0.454045785 | 0.486847177 | -1.231791589 | 0.218026946 | 0.259461931 | 2.594619313 |
| Sult2a2 | 11.900178 | 0.432058107 | 0.453882955 | 1.216112858 | 0.223941933 | 0.265300578 | 2.653005783 |
| Alpl | 12.127792 | 0.402635297 | 0.426352534 | 1.177213581 | 0.239110269 | 0.282000004 | 2.820000037 |
| Kcnip4 | 19.734201 | 0.383767076 | 0.402292994 | 1.164852898 | 0.244078542 | 0.28657436 | 2.865743598 |
| Gnas | 6.961285 | -0.416944512 | 0.537510724 | -1.098986711 | 0.271773862 | 0.317673448 | 3.176734478 |
| Kat7 | 15.104942 | 0.347211912 | 0.386100353 | 1.084142458 | 0.27830164 | 0.323864298 | 3.23864298 |
| Rpl34 | 18.58343 | -0.418021429 | 0.562516232 | -1.080413224 | 0.27995821 | 0.324356869 | 3.243568687 |
| Ralyl | 17.852365 | -0.364923055 | 0.445527318 | -1.054421713 | 0.291689886 | 0.334084482 | 3.340844819 |
| Lrrc4c | 22.398071 | -0.393452986 | 0.508730394 | -1.053383777 | 0.292165136 | 0.334084482 | 3.340844819 |
| Gm960 | 9.572825 | -0.144269504 | 0.89194565 | 1.054134806 | 0.291821203 | 0.334084482 | 3.340844819 |
| Rps16 | 18.085295 | -0.380404593 | 0.486517571 | -1.045100406 | 0.295976566 | 0.336977649 | 3.369776492 |
| Tmeff2 | 11.272396 | -0.144269504 | 0.891945652 | 1.038905037 | 0.298848904 | 0.3387813 | 3.387813002 |
| Lsamp | 30.734985 | 0.261673993 | 0.282349272 | 1.032690898 | 0.301748574 | 0.340600322 | 3.40600322 |
| Malat1 | 80.67501 | -0.340199499 | 0.409918413 | -1.023501108 | 0.306070976 | 0.34400285 | 3.440028495 |
| Arvcf | 7.2156377 | -0.357393186 | 0.496217093 | -0.985534066 | 0.324361807 | 0.363009171 | 3.630091711 |
| Fyb2 | 21.724798 | 0.310057445 | 0.37964394 | 0.980543232 | 0.32681804 | 0.36420824 | 3.642082397 |
| Ankrd12 | 19.460518 | 0.332331868 | 0.434983247 | 0.970794019 | 0.331650862 | 0.367385751 | 3.673857506 |
| Bc1 | 16.305922 | 0.338649241 | 0.447521193 | 0.969164439 | 0.332463151 | 0.367385751 | 3.673857506 |
| Rps3 | 11.808124 | -0.310296886 | 0.411847106 | -0.942488602 | 0.345942544 | 0.380681544 | 3.806815444 |
| Srgn | 7.3214526 | 0.333159431 | 0.528504936 | 0.886243125 | 0.37548653 | 0.411470655 | 4.114706555 |
| Klhl14 | 3.2298203 | 0.328192626 | 0.728232535 | 0.820218884 | 0.412091338 | 0.449709635 | 4.497096347 |
| Lyrm1 | 9.521898 | -0.144269504 | 0.891945661 | 0.791102384 | 0.428884248 | 0.466101477 | 4.661014765 |
| Grm7 | 7.474356 | -0.244524321 | 0.438202673 | -0.725388927 | 0.468213464 | 0.506749552 | 5.067495518 |
| Kcnq5 | 15.415363 | 0.229904807 | 0.405662246 | 0.701375226 | 0.483068879 | 0.520684898 | 5.206848978 |
| Kcnh7 | 9.575889 | 0.229439326 | 0.424517723 | 0.685537665 | 0.493004701 | 0.529225455 | 5.292254549 |
| Hdac9 | 10.943138 | 0.226447134 | 0.424039243 | 0.673997944 | 0.500312621 | 0.53488707 | 5.3488707 |
| Nrg1 | 10.063369 | -0.226962679 | 0.759196367 | -0.627649213 | 0.530233762 | 0.564580889 | 5.645808887 |
| Rabgap1l | 10.037972 | 0.2173435 | 0.494538292 | 0.600952127 | 0.547871873 | 0.581009284 | 5.810092843 |
| Gria1 | 11.72491 | -0.166387212 | 0.393653786 | -0.52569578 | 0.599099596 | 0.632783911 | 6.32783911 |
| Ptprg | 10.404542 | 0.151616578 | 0.429599995 | 0.451330656 | 0.651751253 | 0.685642319 | 6.856423186 |
| Snhg11 | 9.6927 | 0.153245159 | 0.457997813 | 0.440221355 | 0.659776794 | 0.688576575 | 6.885765751 |
| Nkain2 | 23.461985 | 0.120402265 | 0.308113222 | 0.44316566 | 0.657645916 | 0.688576575 | 6.885765751 |
| Kcnd2 | 9.405159 | -0.150152319 | 0.463976374 | -0.43496517 | 0.66358771 | 0.689816473 | 6.898164727 |
| 4930402H1 | 12.072494 | -0.144269504 | 0.891945635 | 0.387594161 | 0.698316391 | 0.723059885 | 7.23059885 |
| Nrxn1 | 13.530051 | -0.139929077 | 0.580513195 | -0.374988207 | 0.707669237 | 0.729870625 | 7.298706248 |
| Sgip1 | 7.5403495 | -0.116755505 | 0.53712142 | -0.321227981 | 0.748037629 | 0.765501543 | 7.655015425 |
| Ccdc60 | 18.026014 | 0.109981681 | 0.437665434 | 0.32346818 | 0.746340702 | 0.765501543 | 7.655015425 |
| Camk2a | 21.779188 | 0.077203651 | 0.323312673 | 0.274506728 | 0.783695234 | 0.798883126 | 7.988831259 |
| Il1rapl2 | 11.235707 | 0.069744242 | 0.430014734 | 0.206491164 | 0.836407266 | 0.846058119 | 8.460581191 |
| Rps28 | 22.09065 | -0.073207827 | 0.488318577 | -0.2085042 | 0.834835303 | 0.846058119 | 8.460581191 |
| Cep128 | 11.021495 | 0.045321389 | 0.40909659 | 0.137665474 | 0.890504807 | 0.893903681 | 8.939036807 |
| Csmd1 | 15.056169 | -0.044782147 | 0.416016376 | -0.138412209 | 0.889914648 | 0.893903681 | 8.939036807 |
| Gnao1 | 11.105964 | -0.037653162 | 0.429934391 | -0.115495499 | 0.908052388 | 0.908052388 | 9.080523881 |

**table S5: Differential gene expression list social compared to elevated platform in GABA and Glut neurons**

| **GABA** | baseMean | log2FoldChange | lfcSE | stat | pvalue | padj | pglobadj |
|---|---|---|---|---|---|---|---|
| Kctd16 | 10.054747 | -0.144269504 | 0.516808996 | 3.752918821 | 0.000174787 | 0.050863154 | 0.508631537 |
| Gm42418 | 100.87583 | -0.860213576 | 0.339271818 | -3.285215832 | 0.001019043 | 0.148270724 | 1.482707241 |
| Calm1 | 4.159139 | -0.144269504 | 0.516808934 | -3.115078353 | 0.001838961 | 0.178379173 | 1.783791726 |
| mt-Nd4 | 13.449779 | -0.940915083 | 0.455004539 | -2.938563309 | 0.003297373 | 0.23988389 | 2.398838905 |
| mt-Atp6 | 49.76098 | -0.144269504 | 0.516808979 | -2.67548486 | 0.007462124 | 0.361913031 | 3.619130306 |
| Lrp1b | 33.654198 | 0.553664723 | 0.255691263 | 2.714643781 | 0.006634709 | 0.361913031 | 3.619130306 |
| Mt3 | 3.100317 | -0.21501584 | 0.478784881 | -2.376422987 | 0.01748141 | 0.530705802 | 5.307058018 |
| Igf2bp3 | 6.8507752 | -0.144269504 | 0.516808984 | 2.454559846 | 0.014105718 | 0.530705802 | 5.307058018 |
| Hexb | 3.7118638 | -0.283832328 | 0.587100792 | -2.383194464 | 0.017163125 | 0.530705802 | 5.307058018 |
| mt-Nd1 | 13.19425 | -0.637578788 | 0.678372694 | -2.360764399 | 0.018237313 | 0.530705802 | 5.307058018 |
| Ckb | 3.4032974 | -0.188950614 | 0.448419633 | -2.143398508 | 0.032081113 | 0.777966986 | 7.779669857 |
| 5830416I1! | 17.28555 | 0.519593557 | 0.439831234 | 2.14432853 | 0.032006572 | 0.777966986 | 7.779669857 |
| Mbp | 3.633173 | -0.140646372 | 0.40963556 | -1.56241077 | 0.118191252 | 0.854747824 | 8.547478238 |
| Camk2n1 | 3.0341349 | -0.177399221 | 0.443336036 | -1.998161016 | 0.045699206 | 0.854747824 | 8.547478238 |
| Camk1d | 9.677498 | -0.232065893 | 0.454898682 | -1.695240844 | 0.090029742 | 0.854747824 | 8.547478238 |
| Gphn | 3.0994344 | -0.151062795 | 0.428724093 | -1.732147013 | 0.08324739 | 0.854747824 | 8.547478238 |
| Mapk14 | 7.4354506 | 0.304685906 | 0.522490308 | 1.775479389 | 0.075818766 | 0.854747824 | 8.547478238 |
| Malat1 | 5.594701 | -0.107601132 | 0.375018163 | -1.529396585 | 0.126166159 | 0.854747824 | 8.547478238 |
| Gm10062 | 8.332621 | 0.203801562 | 0.509180136 | 1.599146765 | 0.109787996 | 0.854747824 | 8.547478238 |
| Rps20 | 2.9428258 | -0.130246758 | 0.408407965 | -1.617768221 | 0.105712558 | 0.854747824 | 8.547478238 |
| Slx1b | 4.696141 | 0.189917005 | 0.518437731 | 1.59753673 | 0.110146117 | 0.854747824 | 8.547478238 |
| Rpl32 | 2.327905 | -0.110582824 | 0.403174591 | -1.505570374 | 0.132177499 | 0.854747824 | 8.547478238 |
| Cmpk1 | 11.141671 | 0.273961505 | 0.465701444 | 1.617867433 | 0.10569117 | 0.854747824 | 8.547478238 |
| Ldhb | 1.6373371 | -0.054400922 | 0.327881449 | -1.523005916 | 0.127757225 | 0.854747824 | 8.547478238 |
| Fyb2 | 2.1210632 | 0.133341925 | 0.53576399 | 1.551812248 | 0.120707155 | 0.854747824 | 8.547478238 |
| Ccdc60 | 2.8408208 | 0.170616424 | 0.549704466 | 1.689547707 | 0.091114517 | 0.854747824 | 8.547478238 |
| Cox7c | 4.787345 | -0.144269504 | 0.51680895 | -2.023674393 | 0.043003665 | 0.854747824 | 8.547478238 |
| Calm2 | 4.7257137 | -0.083697253 | 0.356290424 | -1.530106101 | 0.125990468 | 0.854747824 | 8.547478238 |
| Adamtsl1 | 7.7476234 | 0.228844356 | 0.468608122 | 1.51334471 | 0.130192121 | 0.854747824 | 8.547478238 |
| D430042O( | 7.439981 | 0.281308343 | 0.537216287 | 1.777134928 | 0.075546043 | 0.854747824 | 8.547478238 |
| Phactr1 | 4.6497188 | -0.054825319 | 0.322921866 | -1.94251792 | 0.052074433 | 0.854747824 | 8.547478238 |
| 4930511M | 3.7473068 | 0.185063125 | 0.552786777 | 1.7235486 | 0.084789364 | 0.854747824 | 8.547478238 |
| mt-Nd2 | 15.759436 | -0.313708962 | 0.427579547 | -1.717206817 | 0.08594139 | 0.854747824 | 8.547478238 |
| Rpl38 | 5.6481957 | -0.278334526 | 0.504560043 | -1.804266139 | 0.071189595 | 0.854747824 | 8.547478238 |
| Rps8 | 2.7214947 | -0.08199715 | 0.360828967 | -1.576897488 | 0.114819115 | 0.854747824 | 8.547478238 |
| mt-Co3 | 56.971947 | -0.263348759 | 0.256237997 | -1.528163434 | 0.126471965 | 0.854747824 | 8.547478238 |
| Rpl26 | 4.4618654 | -0.13779972 | 0.403584522 | -1.602621873 | 0.109018162 | 0.854747824 | 8.547478238 |
| Fau | 1.9818405 | -0.050712906 | 0.323652338 | -1.579906008 | 0.114128393 | 0.854747824 | 8.547478238 |
| Rtn1 | 3.9032948 | -0.140880604 | 0.410065376 | -1.676826068 | 0.093576499 | 0.854747824 | 8.547478238 |
| Nckap5 | 14.870321 | 0.39273963 | 0.512600424 | 1.919446399 | 0.054927864 | 0.854747824 | 8.547478238 |
| Fam169a | 4.5740232 | 0.186139597 | 0.508473869 | 1.552149166 | 0.120626536 | 0.854747824 | 8.547478238 |
| Gm19951 | 1.9425768 | -0.144269504 | 0.516808951 | -1.689357124 | 0.091151012 | 0.854747824 | 8.547478238 |
| Rpl27a | 3.789782 | -0.153893208 | 0.420796699 | -1.642591772 | 0.100467424 | 0.854747824 | 8.547478238 |
| Atp5k | 2.0002773 | -0.083602537 | 0.35088423 | -1.720910937 | 0.085266987 | 0.854747824 | 8.547478238 |
| Grik1 | 184.36032 | 0.249225746 | 0.139803638 | 2.029868003 | 0.042369958 | 0.854747824 | 8.547478238 |
| Mc1r | 7.4474297 | 0.197668701 | 0.477435942 | 1.469259685 | 0.141762367 | 0.891539724 | 8.915397236 |
| Syt1 | 2.108912 | 0.115910827 | 0.525650135 | 1.461076719 | 0.143994388 | 0.891539724 | 8.915397236 |
| Pde1c | 6.013044 | -0.144269504 | 0.516808973 | 1.321611265 | 0.18629763 | 0.902574768 | 9.025747681 |
| Eef1a1 | 3.0431514 | 0.093733622 | 0.471252587 | 1.106312574 | 0.268591251 | 0.902574768 | 9.025747681 |
| Rps29 | 10.679612 | -0.071212607 | 0.354100234 | -1.186764153 | 0.235320662 | 0.902574768 | 9.025747681 |
| mt-Nd5 | 3.7092254 | -0.115425708 | 0.410085502 | -1.196539019 | 0.231486281 | 0.902574768 | 9.025747681 |
| Rps19 | 1.2547617 | 0.069029549 | 0.52735895 | 1.161758584 | 0.245333541 | 0.902574768 | 9.025747681 |
| Gm12216 | 18.365543 | 0.218095311 | 0.387176155 | 1.318506127 | 0.187334275 | 0.902574768 | 9.025747681 |
| Rpl34 | 1.2356588 | -0.050565545 | 0.353776775 | -1.211545328 | 0.225686473 | 0.902574768 | 9.025747681 |
| Grm7 | 1.4026153 | -0.065537908 | 0.400405329 | -1.097635249 | 0.272363795 | 0.902574768 | 9.025747681 |
| Rpl35 | 1.6030182 | -0.046092044 | 0.344707599 | -1.273526412 | 0.202831306 | 0.902574768 | 9.025747681 |
| Rplp1 | 4.1578293 | -0.153494407 | 0.425626164 | -1.44005186 | 0.149852727 | 0.902574768 | 9.025747681 |
| Rpl6 | 1.2873253 | 0.075677933 | 0.535067042 | 1.221853837 | 0.221762905 | 0.902574768 | 9.025747681 |
| Rpl35a | 2.8133197 | -0.088533415 | 0.374535557 | -1.367958121 | 0.171325185 | 0.902574768 | 9.025747681 |
| Gm765 | 1.4331813 | -0.144269504 | 0.516808956 | 1.119302216 | 0.263011231 | 0.902574768 | 9.025747681 |
| Rpl41 | 1.9971235 | -0.066400441 | 0.376160272 | -1.134137607 | 0.256736831 | 0.902574768 | 9.025747681 |
| Gnas | 2.272737 | 0.091054078 | 0.476340377 | 1.096308999 | 0.272943572 | 0.902574768 | 9.025747681 |

| | | | | | | | |
|---|---|---|---|---|---|---|---|
| Gnaz | 9.3656435 | 0.181746923 | 0.386750988 | 1.165344832 | 0.243879439 | 0.902574768 | 9.025747681 |
| Dop1b | 3.0111034 | 0.11156019 | 0.490905593 | 1.283849968 | 0.199194456 | 0.902574768 | 9.025747681 |
| Rps3 | 1.296896 | -0.050221817 | 0.352525047 | -1.225472482 | 0.220397252 | 0.902574768 | 9.025747681 |
| Rpl23 | 4.472335 | -0.049799771 | 0.344884681 | -1.146128376 | 0.251742038 | 0.902574768 | 9.025747681 |
| Rpl37 | 4.3471746 | -0.091315413 | 0.387332811 | -1.111202391 | 0.266481242 | 0.902574768 | 9.025747681 |
| Ppia | 1.696819 | -0.075326466 | 0.38066833 | -1.324767778 | 0.185248185 | 0.902574768 | 9.025747681 |
| Atp1b1 | 3.0105686 | -0.098624238 | 0.398325067 | -1.271152996 | 0.203674225 | 0.902574768 | 9.025747681 |
| Srgn | 1.4749106 | -0.059868036 | 0.345717919 | -1.399259723 | 0.161735113 | 0.902574768 | 9.025747681 |
| mt-Co1 | 28.27414 | -0.235483398 | 0.30480917 | -1.35397998 | 0.175742765 | 0.902574768 | 9.025747681 |
| mt-Co2 | 26.223597 | -0.222112012 | 0.340501511 | -1.317001901 | 0.187837987 | 0.902574768 | 9.025747681 |
| Rab29 | 1.9150372 | -0.144269504 | 0.516808956 | 1.224976048 | 0.220584245 | 0.902574768 | 9.025747681 |
| Lrrc4c | 1.8728139 | 0.093069763 | 0.505176771 | 1.260337824 | 0.207547521 | 0.902574768 | 9.025747681 |
| Acoxl | 3.309197 | 0.110481736 | 0.471033973 | 1.159851955 | 0.246109087 | 0.902574768 | 9.025747681 |
| AY036118 | 15.501163 | -0.183838051 | 0.363919583 | -1.195745777 | 0.231795782 | 0.902574768 | 9.025747681 |
| Klkb1 | 1.5227352 | -0.144269504 | 0.516808972 | 1.282618794 | 0.199625663 | 0.902574768 | 9.025747681 |
| Rpl17 | 2.0687087 | -0.144269504 | 0.51680896 | -1.404922721 | 0.160044259 | 0.902574768 | 9.025747681 |
| Kcnq5 | 2.2035956 | 0.119393705 | 0.476153171 | 1.164792972 | 0.244102804 | 0.902574768 | 9.025747681 |
| Cpe | 2.7406192 | -0.093403166 | 0.398406954 | -1.160994877 | 0.245643983 | 0.902574768 | 9.025747681 |
| Nell1 | 3.1006572 | 0.124258715 | 0.499318244 | 1.370271309 | 0.170602226 | 0.902574768 | 9.025747681 |
| Cspp1 | 2.0211616 | 0.06898142 | 0.509691163 | 1.107664266 | 0.268006838 | 0.902574768 | 9.025747681 |
| Rsrc1 | 8.455423 | 0.156473671 | 0.402912257 | 1.096734992 | 0.272757255 | 0.902574768 | 9.025747681 |
| mt-Nd3 | 10.922144 | -0.204441588 | 0.408250606 | -1.391441968 | 0.164091446 | 0.902574768 | 9.025747681 |
| Rpsa | 1.696819 | -0.075326466 | 0.38066833 | -1.324767778 | 0.185248185 | 0.902574768 | 9.025747681 |
| Rps5 | 2.2747567 | -0.061986717 | 0.354045843 | -1.250978711 | 0.210942245 | 0.902574768 | 9.025747681 |
| Cox4i1 | 4.2313905 | -0.106598689 | 0.397603119 | -1.179178643 | 0.238327048 | 0.902574768 | 9.025747681 |
| Xlr5a | 16.316275 | 0.156500633 | 0.404086906 | 1.124929483 | 0.260618917 | 0.902574768 | 9.025747681 |
| mt-Nd4l | 1.8330154 | -0.022301051 | 0.33449312 | -0.858885542 | 0.390403667 | 0.90274242 | 9.0274242 |
| Phkg1 | 2.830667 | -0.144269504 | 0.516808958 | 0.862928505 | 0.388176777 | 0.90274242 | 9.0274242 |
| Gria4 | 2.6592212 | 0.06236281 | 0.484411913 | 0.956042619 | 0.339050703 | 0.90274242 | 9.0274242 |
| Kcnd2 | 1.7110088 | -0.055267719 | 0.371600279 | -0.980145507 | 0.327014299 | 0.90274242 | 9.0274242 |
| Ms4a1 | 4.2882752 | 0.105649571 | 0.438210757 | 0.989637201 | 0.32235148 | 0.90274242 | 9.0274242 |
| Mrc2 | 3.652254 | 0.088066036 | 0.439965722 | 0.907666251 | 0.36405458 | 0.90274242 | 9.0274242 |
| Kat7 | 2.4633076 | 0.076551604 | 0.45402364 | 0.871926283 | 0.383248594 | 0.90274242 | 9.0274242 |
| Rpl13a | 1.2706659 | -0.045429719 | 0.370716573 | -0.955156472 | 0.339498575 | 0.90274242 | 9.0274242 |
| Rpl30 | 1.0352559 | -0.042462748 | 0.39907892 | -0.807569317 | 0.419338556 | 0.90274242 | 9.0274242 |
| Kcnip4 | 1.9796486 | 0.064914122 | 0.459598593 | 0.823661329 | 0.410132018 | 0.90274242 | 9.0274242 |
| Hsp90ab1 | 1.3602197 | -0.048558933 | 0.397704127 | -0.809443075 | 0.418260334 | 0.90274242 | 9.0274242 |
| Il27 | 6.2832284 | 0.09786324 | 0.408309405 | 0.836507761 | 0.402869312 | 0.90274242 | 9.0274242 |
| Plp1 | 2.6336343 | -0.023465932 | 0.330638379 | -0.895776956 | 0.3703719 | 0.90274242 | 9.0274242 |
| Psap | 3.582861 | -0.053513439 | 0.364248029 | -0.919461593 | 0.357854186 | 0.90274242 | 9.0274242 |
| Fbxl7 | 3.5138466 | 0.086453586 | 0.440217361 | 0.874273875 | 0.381969123 | 0.90274242 | 9.0274242 |
| Nrxn1 | 1.932917 | -0.056383683 | 0.397974348 | -0.853614916 | 0.393318391 | 0.90274242 | 9.0274242 |
| Cox6b1 | 3.715559 | -0.070942121 | 0.388593363 | -0.828593391 | 0.407334529 | 0.90274242 | 9.0274242 |
| 6030443J0 | 4.7003493 | 0.097118051 | 0.428238465 | 0.898318279 | 0.36901589 | 0.90274242 | 9.0274242 |
| Sema3c | 15.617407 | 0.154208512 | 0.372735428 | 1.028377608 | 0.303772237 | 0.90274242 | 9.0274242 |
| Rpl36a | 1.0454094 | 0.061863242 | 0.531416003 | 1.040212611 | 0.298241134 | 0.90274242 | 9.0274242 |
| Shb | 9.115747 | 0.129996692 | 0.410204586 | 1.013723706 | 0.310714609 | 0.90274242 | 9.0274242 |
| Sil1 | 4.2721095 | 0.072057812 | 0.435927789 | 0.802746855 | 0.422121065 | 0.90274242 | 9.0274242 |
| Cep128 | 1.1042031 | 0.045670951 | 0.52984222 | 0.927816733 | 0.353502643 | 0.90274242 | 9.0274242 |
| AI314278 | 2.8831494 | -0.08253838 | 0.403466094 | -0.92747628 | 0.353679302 | 0.90274242 | 9.0274242 |
| Cdk8 | 3.8737385 | -0.080048748 | 0.38909769 | -0.958546251 | 0.337787381 | 0.90274242 | 9.0274242 |
| Car12 | 2.8213964 | -0.050076951 | 0.360463054 | -0.973174459 | 0.330466603 | 0.90274242 | 9.0274242 |
| Camk2a | 2.4877346 | 0.071854408 | 0.449576011 | 0.839357953 | 0.401268471 | 0.90274242 | 9.0274242 |
| Cadm2 | 0.59431374 | 0.035751677 | 0.723697001 | 0.795917143 | 0.426080196 | 0.90274242 | 9.0274242 |
| Zbtb7a | 4.4359345 | -0.144269504 | 0.516808971 | 0.862681533 | 0.388312589 | 0.90274242 | 9.0274242 |
| Ubb | 0.99423677 | 0.041531429 | 0.519305539 | 0.841331276 | 0.400162374 | 0.90274242 | 9.0274242 |
| Atxn7l1 | 1.806874 | -0.144269504 | 0.516808966 | 1.067271907 | 0.28584907 | 0.90274242 | 9.0274242 |
| Cox6c | 1.8433632 | -0.069831383 | 0.396636134 | -1.084349752 | 0.278209754 | 0.90274242 | 9.0274242 |
| Ttc28 | 2.799432 | 0.070049522 | 0.474531924 | 0.987621841 | 0.323337886 | 0.90274242 | 9.0274242 |
| Tpt1 | 5.9276953 | -0.057844406 | 0.360598826 | -0.883491758 | 0.376970634 | 0.90274242 | 9.0274242 |
| Rps4x | 1.5752174 | -0.041775407 | 0.367166824 | -0.856573267 | 0.391680765 | 0.90274242 | 9.0274242 |
| Rpl9 | 2.4686317 | -0.086222267 | 0.404375649 | -1.066102473 | 0.286377322 | 0.90274242 | 9.0274242 |
| Gm48099 | 2.6234372 | -0.0560135 | 0.384220713 | -0.821427294 | 0.411402922 | 0.90274242 | 9.0274242 |
| Rps10 | 0.56175023 | 0.037752667 | 0.684378708 | 0.792439161 | 0.428104653 | 0.90274242 | 9.0274242 |
| Rps17 | 1.7309535 | -0.022037647 | 0.335647273 | -0.833816112 | 0.404384614 | 0.90274242 | 9.0274242 |

| | | | | | | | |
|---|---|---|---|---|---|---|---|
| Rps18 | 0.8503305 | -0.042779739 | 0.368751283 | -0.981943613 | 0.326127628 | 0.90274242 | 9.0274242 |
| Rps21 | 4.558441 | -0.103114981 | 0.397196248 | -0.991672051 | 0.321357529 | 0.90274242 | 9.0274242 |
| Csmd1 | 1.0128459 | 0.060370606 | 0.519772419 | 1.008734134 | 0.313102157 | 0.90274242 | 9.0274242 |
| Galnt13 | 3.9169014 | 0.102762448 | 0.458354123 | 1.070151486 | 0.284551127 | 0.90274242 | 9.0274242 |
| Frmpd4 | 2.6953914 | 0.072704308 | 0.468802461 | 0.93304338 | 0.350797571 | 0.90274242 | 9.0274242 |
| Rps27 | 1.8113155 | -0.144269504 | 0.516808951 | -0.792796583 | 0.427896348 | 0.90274242 | 9.0274242 |
| Epha6 | 10.879962 | -0.144269504 | 0.51680897 | -1.002927143 | 0.315896015 | 0.90274242 | 9.0274242 |
| Rps27a | 1.6181068 | 0.061014549 | 0.482618567 | 0.910984509 | 0.362303536 | 0.90274242 | 9.0274242 |
| Ehbp1 | 7.3124123 | 0.084604541 | 0.440719465 | 0.929399492 | 0.352682091 | 0.90274242 | 9.0274242 |
| 4930441J1 | 1.5797255 | 0.071145569 | 0.484846876 | 1.012158866 | 0.3114621 | 0.90274242 | 9.0274242 |
| Fth1 | 4.9377313 | -0.090095135 | 0.380024525 | -1.025849307 | 0.304962621 | 0.90274242 | 9.0274242 |
| Rps3a1 | 5.56022 | -0.017489676 | 0.329090967 | -0.786208466 | 0.431745367 | 0.903869798 | 9.038697978 |
| Cox8a | 3.9048557 | -0.049162092 | 0.370503328 | -0.778315863 | 0.436382826 | 0.907052874 | 9.070528743 |
| Snhg11 | 3.8826401 | -0.056887554 | 0.378439586 | -0.769955613 | 0.441326223 | 0.910822205 | 9.108222045 |
| Gm50306 | 6.528987 | 0.082754864 | 0.407534004 | 0.758137049 | 0.448368942 | 0.918840578 | 9.188405784 |
| Tmeff2 | 3.6059976 | -0.033785697 | 0.363888949 | -0.663140174 | 0.507240783 | 0.934221948 | 9.342219484 |
| Spock1 | 2.4953773 | -0.052278208 | 0.394063831 | -0.672840175 | 0.501048975 | 0.934221948 | 9.342219484 |
| Pip5k1b | 3.0078826 | -0.071076748 | 0.422219385 | -0.674764765 | 0.49982523 | 0.934221948 | 9.342219484 |
| Ptprg | 2.0954328 | 0.047744046 | 0.457105316 | 0.688771775 | 0.4909669 | 0.934221948 | 9.342219484 |
| Rpl12 | 1.8963392 | -0.027763499 | 0.361919268 | -0.669634822 | 0.50309061 | 0.934221948 | 9.342219484 |
| Sec61a1 | 34.732437 | 0.087078186 | 0.359079753 | 0.663939463 | 0.506729057 | 0.934221948 | 9.342219484 |
| Rpl23a | 0.64499253 | -0.012070807 | 0.315741024 | -0.693103734 | 0.488244455 | 0.934221948 | 9.342219484 |
| Tmem108 | 2.4935522 | 0.054496616 | 0.442329841 | 0.681365387 | 0.495640319 | 0.934221948 | 9.342219484 |
| Sgip1 | 1.0718338 | -0.035345577 | 0.393557101 | -0.705210682 | 0.480679135 | 0.934221948 | 9.342219484 |
| 1700054A0 | 30.814533 | 0.092429664 | 0.370083186 | 0.723887268 | 0.469134947 | 0.934221948 | 9.342219484 |
| Ndufa4 | 1.0065343 | -0.018648635 | 0.342197798 | -0.718903025 | 0.472200673 | 0.934221948 | 9.342219484 |
| Il1rapl2 | 1.7379271 | -0.042533839 | 0.391747572 | -0.667034597 | 0.504750035 | 0.934221948 | 9.342219484 |
| Cntnap2 | 12.196445 | -0.080130415 | 0.36566659 | -0.735547566 | 0.462006082 | 0.934221948 | 9.342219484 |
| Ndrg2 | 1.6221211 | 0.038597921 | 0.480268302 | 0.731267454 | 0.464615806 | 0.934221948 | 9.342219484 |
| Ahdc1 | 2.4895377 | 0.0603049 | 0.44660063 | 0.721925216 | 0.470340457 | 0.934221948 | 9.342219484 |
| Negr1 | 0.97646224 | -0.034145814 | 0.394973218 | -0.666001052 | 0.505410428 | 0.934221948 | 9.342219484 |
| Wdr72 | 5.6548285 | 0.076863653 | 0.398262207 | 0.656612832 | 0.511429896 | 0.936013205 | 9.360132055 |
| Gria2 | 0.429995 | -0.011590555 | 0.318660715 | -0.579527165 | 0.562233522 | 0.951220669 | 9.512206687 |
| Gm960 | 0.8599682 | 0.025262735 | 0.484349903 | 0.589249347 | 0.555694019 | 0.951220669 | 9.512206687 |
| Uba52 | 1.9636774 | -0.044254815 | 0.407074041 | -0.602557305 | 0.54680323 | 0.951220669 | 9.512206687 |
| Tspan7 | 0.7915368 | -0.01538764 | 0.351260704 | -0.580302164 | 0.561710869 | 0.951220669 | 9.512206687 |
| Apoe | 2.826653 | 0.054048284 | 0.436023275 | 0.625685869 | 0.531521004 | 0.951220669 | 9.512206687 |
| Esf1 | 1.8126919 | 0.042897146 | 0.451424303 | 0.611674319 | 0.540753256 | 0.951220669 | 9.512206687 |
| Csmd3 | 4.069974 | 0.062375756 | 0.414347299 | 0.613319938 | 0.539664812 | 0.951220669 | 9.512206687 |
| Sparcl1 | 2.2664254 | -0.055812807 | 0.412159274 | -0.635432999 | 0.525146052 | 0.951220669 | 9.512206687 |
| Ttc3 | 1.1714523 | -0.016871376 | 0.351650051 | -0.595394015 | 0.551580125 | 0.951220669 | 9.512206687 |
| Aldoc | 1.0430903 | 0.01854143 | 0.530562447 | 0.59459198 | 0.552116242 | 0.951220669 | 9.512206687 |
| Rpl36 | 0.41520596 | 0.027669651 | 0.69468906 | 0.591786592 | 0.553993503 | 0.951220669 | 9.512206687 |
| Prkag2 | 2.3373704 | -0.144269504 | 0.516808958 | -0.624173611 | 0.532513575 | 0.951220669 | 9.512206687 |
| Pcgf6 | 2.1524448 | -0.044151296 | 0.398578489 | -0.579922028 | 0.5619672 | 0.951220669 | 9.512206687 |
| Cntn3 | 1.9270995 | -0.034945726 | 0.391231584 | -0.569043276 | 0.569326772 | 0.95532507 | 9.553250705 |
| Zmat4 | 4.238412 | -0.056363537 | 0.390947755 | -0.566247859 | 0.5712253 | 0.95532507 | 9.553250705 |
| Atp6v0c | 1.4594114 | 0.028799908 | 0.465125451 | 0.548581605 | 0.583292612 | 0.969932287 | 9.699322867 |
| Pcdh15 | 7.6551676 | -0.052676242 | 0.370366923 | -0.484483855 | 0.62804252 | 0.971207213 | 9.712072135 |
| Arvcf | 0.5373234 | 0.015768709 | 0.958386267 | 0.515423663 | 0.606257 | 0.971207213 | 9.712072135 |
| Rps23 | 1.2558768 | 0.013125123 | 0.448251412 | 0.283887541 | 0.776496565 | 0.971207213 | 9.712072135 |
| Gapdh | 0.6564336 | 0.013375967 | 0.465395676 | 0.310628579 | 0.756082999 | 0.971207213 | 9.712072135 |
| Tmsb4x | 2.6881373 | 0.031667275 | 0.424647612 | 0.419272655 | 0.675016879 | 0.971207213 | 9.712072135 |
| Prkg1 | 4.030952 | 0.042875495 | 0.413647323 | 0.467494783 | 0.640145924 | 0.971207213 | 9.712072135 |
| Atp5j2 | 0.33078152 | -0.010953738 | 0.324577133 | -0.450675936 | 0.652223128 | 0.971207213 | 9.712072135 |
| Rps14 | 3.6888611 | -0.010273885 | 0.358479101 | -0.352417687 | 0.72452504 | 0.971207213 | 9.712072135 |
| Nkain2 | 2.7724729 | -0.022782305 | 0.401991792 | -0.293974166 | 0.768777644 | 0.971207213 | 9.712072135 |
| Gm29325 | 3.1352184 | 0.0501419 | 0.421197642 | 0.531582926 | 0.595014893 | 0.971207213 | 9.712072135 |
| Rplp2 | 1.6708024 | 0.006683617 | 0.48555919 | 0.284923166 | 0.775703007 | 0.971207213 | 9.712072135 |
| Gm15564 | 1.2029009 | 0.018354229 | 0.448825343 | 0.32331521 | 0.746456536 | 0.971207213 | 9.712072135 |
| Pcsk1n | 1.2183781 | -0.018721126 | 0.401794428 | -0.360105063 | 0.718768567 | 0.971207213 | 9.712072135 |
| Dlg2 | 1.0871168 | 0.022773755 | 0.44941087 | 0.387117742 | 0.698669043 | 0.971207213 | 9.712072135 |
| Dctd | 2.3264232 | 0.02679187 | 0.423958921 | 0.362799712 | 0.716754507 | 0.971207213 | 9.712072135 |
| Dbi | 1.6482006 | -0.013661871 | 0.354843467 | -0.446709341 | 0.655084937 | 0.971207213 | 9.712072135 |
| Ctnna3 | 3.839693 | 0.029870415 | 0.403037052 | 0.301863496 | 0.762756124 | 0.971207213 | 9.712072135 |

| | | | | | | | |
|---|---|---|---|---|---|---|---|
| Svil | 1.7819316 | 0.022857341 | 0.441512118 | 0.36067466 | 0.71834267 | 0.971207213 | 9.712072135 |
| St18 | 1.1818001 | -0.029317154 | 0.41587527 | -0.495708025 | 0.620100425 | 0.971207213 | 9.712072135 |
| Ctcflos | 3.6227195 | 0.028314994 | 0.416323686 | 0.380044944 | 0.703912053 | 0.971207213 | 9.712072135 |
| Cst3 | 15.340955 | -0.038253288 | 0.357374226 | -0.347698312 | 0.728066765 | 0.971207213 | 9.712072135 |
| Bc1 | 1.1102152 | -0.009674325 | 0.346318402 | -0.452703498 | 0.65076226 | 0.971207213 | 9.712072135 |
| Selenow | 1.6499821 | 0.018894651 | 0.43997698 | 0.317312137 | 0.751006769 | 0.971207213 | 9.712072135 |
| Cmss1 | 4.3203454 | -0.144269504 | 0.516808966 | -0.398359062 | 0.69036553 | 0.971207213 | 9.712072135 |
| Nrxn3 | 2.4543583 | 0.027084264 | 0.42657145 | 0.369604333 | 0.711677322 | 0.971207213 | 9.712072135 |
| Chl1 | 2.0712001 | -0.031609312 | 0.401238621 | -0.464638121 | 0.642190628 | 0.971207213 | 9.712072135 |
| Cdh23 | 6.210955 | 0.052603943 | 0.384297645 | 0.453179655 | 0.650419381 | 0.971207213 | 9.712072135 |
| Ankrd12 | 4.59216 | -0.026842937 | 0.380529605 | -0.298805667 | 0.765088327 | 0.971207213 | 9.712072135 |
| Gm49767 | 2.0115237 | 0.019889048 | 0.428386638 | 0.282404627 | 0.777633267 | 0.971207213 | 9.712072135 |
| Pde6a | 10.6259 | 0.049100125 | 0.388629785 | 0.476962456 | 0.633388856 | 0.971207213 | 9.712072135 |
| Rpl39 | 1.0487356 | 0.022857286 | 0.450595477 | 0.449475186 | 0.653088905 | 0.971207213 | 9.712072135 |
| Mobp | 0.5765392 | -0.010025308 | 0.367131225 | -0.38695541 | 0.698789219 | 0.971207213 | 9.712072135 |
| Zfp994 | 2.3308425 | 0.028809073 | 0.434002267 | 0.415382854 | 0.677861657 | 0.971207213 | 9.712072135 |
| Ptprd | 2.5817294 | -0.144269504 | 0.516808972 | -0.395744562 | 0.692293478 | 0.971207213 | 9.712072135 |
| Meg3 | 2.4789577 | 0.016509115 | 0.435222868 | 0.326716996 | 0.743881944 | 0.971207213 | 9.712072135 |
| Lyrm1 | 1.8249676 | 0.028063454 | 0.44050925 | 0.467950534 | 0.639819964 | 0.971207213 | 9.712072135 |
| Alpl | 2.5561879 | 0.038591183 | 0.424126957 | 0.452116459 | 0.651185087 | 0.971207213 | 9.712072135 |
| Lsamp | 4.3221893 | 0.03548925 | 0.422152596 | 0.486672752 | 0.626490258 | 0.971207213 | 9.712072135 |
| Rpl13 | 3.516727 | -0.034709494 | 0.378953899 | -0.52130584 | 0.602153734 | 0.971207213 | 9.712072135 |
| Lrrc17 | 2.2260056 | -0.026994188 | 0.403600325 | -0.384616137 | 0.700521837 | 0.971207213 | 9.712072135 |
| Rpl21 | 3.0930173 | -0.034022583 | 0.39381225 | -0.387165469 | 0.698633713 | 0.971207213 | 9.712072135 |
| Vit | 1.2143638 | -0.016340437 | 0.409055889 | -0.302064998 | 0.762602514 | 0.971207213 | 9.712072135 |
| Adk | 2.9246624 | -0.032741213 | 0.39961951 | -0.381753752 | 0.702644026 | 0.971207213 | 9.712072135 |
| Ppp1r16a | 3.338581 | 0.035393489 | 0.422313954 | 0.462669679 | 0.643601155 | 0.971207213 | 9.712072135 |
| Uqcrh | 1.9008694 | -0.023553618 | 0.39685393 | -0.367890347 | 0.712954998 | 0.971207213 | 9.712072135 |
| Hdac9 | 1.5400158 | 0.040249691 | 0.450774527 | 0.452913435 | 0.650611076 | 0.971207213 | 9.712072135 |
| Jarid2 | 1.8519077 | -0.013127729 | 0.378392371 | -0.303822365 | 0.761263227 | 0.971207213 | 9.712072135 |
| Rpl3 | 1.6328958 | -0.020652148 | 0.413857936 | -0.296913377 | 0.766532643 | 0.971207213 | 9.712072135 |
| Rpl37a | 3.7412033 | -0.016107666 | 0.373874222 | -0.347251739 | 0.728402204 | 0.971207213 | 9.712072135 |
| Gpx4 | 0.3826424 | 0.024251087 | 0.685800584 | 0.523323977 | 0.600748818 | 0.971207213 | 9.712072135 |
| Gria1 | 0.63602114 | 0.013822424 | 0.465155931 | 0.309491774 | 0.756947468 | 0.971207213 | 9.712072135 |
| Mt1 | 3.395228 | 0.024539249 | 0.416398362 | 0.332569745 | 0.739459085 | 0.971207213 | 9.712072135 |
| Rpl28 | 1.4102774 | 0.017383797 | 0.434351544 | 0.337040198 | 0.736086597 | 0.971207213 | 9.712072135 |
| Grik2 | 0.5976399 | 0.029737971 | 0.470849026 | 0.452374884 | 0.650998938 | 0.971207213 | 9.712072135 |
| Cfap74 | 1.0864286 | -0.022926139 | 0.406297129 | -0.414510466 | 0.678500304 | 0.971207213 | 9.712072135 |
| Hbb-bs | 2.4392 | -0.024979278 | 0.390326734 | -0.378215193 | 0.70527074 | 0.971207213 | 9.712072135 |
| Uqcr11 | 0.23609814 | 0.016841084 | 0.665388286 | 0.296483557 | 0.766860822 | 0.971207213 | 9.712072135 |
| Hapln1 | 2.6859262 | 0.02518408 | 0.424920417 | 0.32086498 | 0.748312714 | 0.971207213 | 9.712072135 |
| Vwc2l | 3.2805588 | -0.022754541 | 0.404992038 | -0.224636966 | 0.822261705 | 0.984134304 | 9.841343043 |
| Anks1b | 1.3550904 | 0.00864886 | 0.4284784 | 0.142086874 | 0.887011386 | 0.984134304 | 9.841343043 |
| Atp9b | 2.5734684 | -0.011269557 | 0.405959562 | -0.134726805 | 0.892827879 | 0.984134304 | 9.841343043 |
| Trim2 | 7.6706095 | 0.022491846 | 0.376616411 | 0.204351918 | 0.838078501 | 0.984134304 | 9.841343043 |
| Wdyhv1 | 3.10377 | -0.010701891 | 0.401325777 | -0.123884246 | 0.901406927 | 0.984134304 | 9.841343043 |
| 4930558K0 | 4.527486 | -0.023292842 | 0.387915941 | -0.230485392 | 0.817714613 | 0.984134304 | 9.841343043 |
| 4930402H2 | 1.7428621 | -0.144269504 | 0.516808958 | -0.14863568 | 0.881841117 | 0.984134304 | 9.841343043 |
| Snap25 | 1.9701834 | -0.007607131 | 0.407626572 | -0.129163573 | 0.897228219 | 0.984134304 | 9.841343043 |
| Tns3 | 1.5644426 | 0.015642181 | 0.43313648 | 0.230502485 | 0.817701332 | 0.984134304 | 9.841343043 |
| Tacc1 | 1.6923778 | 0.015088326 | 0.435108807 | 0.241845911 | 0.808899563 | 0.984134304 | 9.841343043 |
| Zfhx3 | 2.3324132 | 0.01151634 | 0.425945615 | 0.229168642 | 0.818737844 | 0.984134304 | 9.841343043 |
| Unc5d | 7.921019 | 0.028193009 | 0.369908918 | 0.234621856 | 0.814502244 | 0.984134304 | 9.841343043 |
| Aldoa | 0.960985 | 0.007088203 | 0.438961254 | 0.130762513 | 0.895963178 | 0.984134304 | 9.841343043 |
| Nlgn1 | 0.9975629 | 0.011406003 | 0.439120007 | 0.206604581 | 0.836318682 | 0.984134304 | 9.841343043 |
| Slc1a2 | 1.8604496 | -0.006823225 | 0.376346458 | -0.223777085 | 0.822930761 | 0.984134304 | 9.841343043 |
| Ntm | 7.8656616 | 0.01842101 | 0.374930643 | 0.171105549 | 0.864140773 | 0.984134304 | 9.841343043 |
| Pcca | 2.358471 | -0.144269504 | 0.516808968 | 0.154862365 | 0.876929824 | 0.984134304 | 9.841343043 |
| Ppp1r9a | 2.7022164 | 0.014244423 | 0.4148031 | 0.204906965 | 0.837644813 | 0.984134304 | 9.841343043 |
| Ptgds | 3.6931489 | 0.015978157 | 0.40072708 | 0.166237649 | 0.867969932 | 0.984134304 | 9.841343043 |
| Rabgap1l | 2.5734684 | -0.011269557 | 0.405959562 | -0.134726805 | 0.892827879 | 0.984134304 | 9.841343043 |
| Klhl14 | 16.04287 | -0.029302775 | 0.304498073 | -0.200606335 | 0.841006404 | 0.984134304 | 9.841343043 |
| Hdac8 | 1.9765167 | 0.007946729 | 0.420512135 | 0.121912156 | 0.902968588 | 0.984134304 | 9.841343043 |
| Slc25a4 | 0.59944326 | 0.004306626 | 0.447346047 | 0.15356275 | 0.87795451 | 0.984134304 | 9.841343043 |
| Gm38973 | 3.883844 | -0.01358751 | 0.390509812 | -0.145755798 | 0.884114171 | 0.984134304 | 9.841343043 |

| | | | | | | | |
|---|---|---|---|---|---|---|---|
| Rps12 | 1.0601985 | -0.008932084 | 0.416172784 | -0.148503671 | 0.881945289 | 0.984134304 | 9.841343043 |
| Rps15 | 1.480534 | -0.015760594 | 0.403049541 | -0.261136319 | 0.793987385 | 0.984134304 | 9.841343043 |
| Rps24 | 1.6858717 | -0.009971281 | 0.409460852 | -0.160668682 | 0.872354357 | 0.984134304 | 9.841343043 |
| Gas5 | 0.4836592 | 0.008237448 | 0.460687139 | 0.158567582 | 0.874009571 | 0.984134304 | 9.841343043 |
| Nrg1 | 2.3236961 | -0.007024547 | 0.399235036 | -0.132726588 | 0.894409615 | 0.984134304 | 9.841343043 |
| Eif1 | 0.5790308 | 0.00515458 | 0.450246401 | 0.158567582 | 0.874009571 | 0.984134304 | 9.841343043 |
| Rps28 | 1.8586462 | -0.009480688 | 0.408604813 | -0.15664239 | 0.875526692 | 0.984134304 | 9.841343043 |
| Rps6 | 0.508086 | 0.00668192 | 0.451006615 | 0.14546718 | 0.884342027 | 0.984134304 | 9.841343043 |
| Ctnna2 | 2.272759 | -0.008864545 | 0.397452061 | -0.130863197 | 0.895883529 | 0.984134304 | 9.841343043 |
| Rps2 | 0.9342391 | -0.009209019 | 0.396714159 | -0.245222041 | 0.806284511 | 0.984134304 | 9.841343043 |
| Gm11337 | 2.116555 | -0.004083906 | 0.41168212 | -0.05416215 | 0.956805976 | 0.986531172 | 9.865311723 |
| Chsy3 | 4.1233163 | -0.007777014 | 0.391828258 | -0.085900298 | 0.931545675 | 0.986531172 | 9.865311723 |
| Scamp1 | 1.8120037 | -0.144269504 | 0.516808965 | 0.092866585 | 0.926009552 | 0.986531172 | 9.865311723 |
| 8430422H( | 5.263237 | 0.014127484 | 0.370907227 | 0.109816527 | 0.912554883 | 0.986531172 | 9.865311723 |
| mt-Cytb | 15.885096 | -0.144269504 | 0.516808978 | -0.06382208 | 0.949111897 | 0.986531172 | 9.865311723 |
| Ninj2 | 2.3014805 | 0.003816413 | 0.415392574 | 0.046639962 | 0.962800182 | 0.986531172 | 9.865311723 |
| Ndrg4 | 2.9026191 | -0.00474645 | 0.396851942 | -0.081130825 | 0.935337912 | 0.986531172 | 9.865311723 |
| Ppp3ca | 1.0359442 | 0.002227849 | 0.427284367 | 0.049598753 | 0.960442141 | 0.986531172 | 9.865311723 |
| Ptpn9 | 3.6076093 | 0.004322653 | 0.39777981 | 0.050184272 | 0.959975545 | 0.986531172 | 9.865311723 |
| Ralyl | 2.1369674 | -0.003864653 | 0.408700551 | -0.055832581 | 0.955475179 | 0.986531172 | 9.865311723 |
| Rapgef4 | 8.921907 | -0.144269504 | 0.516808968 | 0.068669585 | 0.945252629 | 0.986531172 | 9.865311723 |
| Lars2 | 1.6476631 | 0.003202782 | 0.419205189 | 0.057759079 | 0.953940534 | 0.986531172 | 9.865311723 |
| Dpyd | 3.1711082 | 0.010014588 | 0.402934606 | 0.107820398 | 0.914138162 | 0.986531172 | 9.865311723 |
| Gm49497 | 0.8510186 | -0.0038867 | 0.4283237 | -0.074823405 | 0.940355219 | 0.986531172 | 9.865311723 |
| Tshz2 | 7.7823305 | 0.008729854 | 0.358429328 | 0.067278237 | 0.946360202 | 0.986531172 | 9.865311723 |
| Gm14582 | 2.5490417 | -0.006700277 | 0.404637083 | -0.090404901 | 0.927965462 | 0.986531172 | 9.865311723 |
| Gnao1 | 1.3225267 | 0.002952304 | 0.423734411 | 0.046829524 | 0.962649098 | 0.986531172 | 9.865311723 |
| Rps7 | 2.285315 | -0.002498066 | 0.407186174 | -0.041988 | 0.966508265 | 0.986632881 | 9.866328807 |
| 1700126G( | 5.133263 | 0.004091633 | 0.387270737 | 0.038009101 | 0.969680426 | 0.986632881 | 9.866328807 |
| Actb | 1.7671645 | 0.001346432 | 0.410008863 | 0.028052976 | 0.977619899 | 0.991245264 | 9.912452637 |
| Rpl27 | 0.08955391 | 0.008597471 | 0.634686674 | 0.011620666 | 0.990728259 | 0.995704464 | 9.957044644 |
| Rps16 | 2.0092049 | 0.001147601 | 0.411068528 | 0.017372683 | 0.986139302 | 0.995704464 | 9.957044644 |
| Sult2a2 | 1.9073753 | 0.000791316 | 0.421130024 | 0.009672227 | 0.9922828 | 0.995704464 | 9.957044644 |
| Rpl11 | 2.1779647 | 0.000152254 | 0.406683846 | 0.001213916 | 0.999031435 | 0.999031435 | 9.990314351 |

| **Glut** | baseMean | log2FoldChange | lfcSE | stat | pvalue | padj | pglobadj |
|---|---|---|---|---|---|---|---|
| mt-Co1 | 161.67729 | -0.144269504 | 0.515390726 | -7.044658394 | 1.86E-12 | 3.63E-10 | 3.63E-09 |
| mt-Atp6 | 259.72635 | -0.522239028 | 0.127572751 | -4.345670675 | 1.39E-05 | 0.001353795 | 0.013537949 |
| 8430422H( | 29.454706 | -0.144269504 | 0.515390841 | 4.207464563 | 2.58E-05 | 0.001678637 | 0.016786373 |
| Camk1d | 86.7444 | -0.929789498 | 0.266463518 | -3.99879307 | 6.37E-05 | 0.003103733 | 0.031037327 |
| Car12 | 11.724478 | -0.144269504 | 0.515390807 | -3.765747912 | 0.000166051 | 0.006475999 | 0.064759994 |
| Rpl6 | 12.117716 | -0.144269504 | 0.5153908 | -3.719791324 | 0.000199387 | 0.006480092 | 0.06480092 |
| AY036118 | 118.59495 | -0.803600277 | 0.265778655 | -3.559467668 | 0.000371607 | 0.010351917 | 0.10351917 |
| Rpl21 | 27.682667 | -0.779100789 | 0.268017582 | -3.41050534 | 0.000648426 | 0.014323697 | 0.143236974 |
| Cst3 | 81.87364 | 0.946611178 | 0.339890657 | 3.37634833 | 0.000734549 | 0.014323697 | 0.143236974 |
| mt-Nd2 | 94.23299 | -0.525967367 | 0.172179099 | -3.395412271 | 0.000685253 | 0.014323697 | 0.143236974 |
| Xlr5a | 68.85628 | 0.639570919 | 0.223074785 | 3.298330886 | 0.000972614 | 0.017241802 | 0.172418021 |
| Ehbp1 | 25.875452 | -0.144269504 | 0.515390832 | 3.250670961 | 0.00115133 | 0.018709116 | 0.187091162 |
| Il27 | 24.08543 | -0.144269504 | 0.515390836 | 3.20697142 | 0.001341403 | 0.020121048 | 0.201210482 |
| 4930558K( | 15.388913 | -0.144269504 | 0.515390839 | 3.066954552 | 0.002162517 | 0.029887674 | 0.298876745 |
| mt-Cytb | 100.34639 | -0.720626821 | 0.294492715 | -3.048606785 | 0.002299052 | 0.029887674 | 0.298876745 |
| mt-Co2 | 174.3826 | -0.144269504 | 0.515390777 | -3.01700242 | 0.002552878 | 0.031113197 | 0.311131974 |
| Ndufa4 | 11.315106 | -0.144269504 | 0.515390807 | -2.977873619 | 0.002902556 | 0.033294029 | 0.332940294 |
| Pde1c | 33.343422 | 0.913460149 | 0.409782393 | 2.948189946 | 0.003196406 | 0.034627732 | 0.346277316 |
| Rapgef4 | 35.786278 | 0.635548548 | 0.27130833 | 2.861765297 | 0.004212888 | 0.042285752 | 0.422857517 |
| Gm10062 | 45.72148 | 0.590233169 | 0.248731759 | 2.852549906 | 0.004337 | 0.042285752 | 0.422857517 |
| Gnaz | 31.083302 | 0.824726453 | 0.401090436 | 2.7822376 | 0.00539855 | 0.049689119 | 0.496891193 |
| Lrp1b | 150.28911 | 0.567086202 | 0.247334044 | 2.769796528 | 0.005609132 | 0.049689119 | 0.496891193 |
| mt-Nd4 | 93.62785 | -0.634756779 | 0.295371922 | -2.755471115 | 0.005860768 | 0.049689119 | 0.496891193 |
| Wdyhv1 | 12.175273 | 1.050541002 | 0.654775347 | 2.61625009 | 0.008890142 | 0.064206578 | 0.642065783 |
| Wdr72 | 18.82549 | 0.760357478 | 0.404608942 | 2.619525877 | 0.008805209 | 0.064206578 | 0.642065783 |
| Gphn | 17.866236 | -0.144269504 | 0.515390807 | -2.626715443 | 0.00862134 | 0.064206578 | 0.642065783 |
| Cox4i1 | 34.3311 | -0.785311621 | 0.437418899 | -2.646869825 | 0.008124062 | 0.064206578 | 0.642065783 |
| Fam169a | 20.200886 | 0.831248576 | 0.495901077 | 2.5543712 | 0.010637978 | 0.074085918 | 0.740859176 |
| Rpl41 | 24.108028 | -0.790375271 | 0.506216434 | -2.524791353 | 0.0115767 | 0.076556698 | 0.765566982 |

| | | | | | | | |
|---|---|---|---|---|---|---|---|
| Atp5k | 10.8501835 | -0.144269504 | 0.515390805 | -2.518728198 | 0.011777954 | 0.076556698 | 0.765566982 |
| Rpl39 | 21.842203 | -0.780871336 | 0.52588309 | -2.483736491 | 0.013001197 | 0.079226047 | 0.79226047 |
| Calm2 | 12.349148 | -0.9012379 | 0.698761189 | -2.492220059 | 0.012694736 | 0.079226047 | 0.79226047 |
| Nckap5 | 70.68766 | 0.583457305 | 0.321787663 | 2.431243795 | 0.015047086 | 0.088914599 | 0.889145992 |
| mt-Nd3 | 50.742096 | -0.144269504 | 0.515390792 | -2.3625239 | 0.018150971 | 0.090653196 | 0.90653196 |
| Gm12216 | 65.97405 | 0.565356289 | 0.326052765 | 2.35866913 | 0.0183406 | 0.090653196 | 0.90653196 |
| Rps14 | 24.104664 | -0.558682163 | 0.266609065 | -2.392032096 | 0.016755377 | 0.090653196 | 0.90653196 |
| Rpl13a | 17.019503 | -0.721823056 | 0.546954768 | -2.358109811 | 0.018368259 | 0.090653196 | 0.90653196 |
| Rps21 | 42.12771 | -0.719072985 | 0.613816605 | -2.359426103 | 0.018303226 | 0.090653196 | 0.90653196 |
| Rplp2 | 10.934576 | -0.775237594 | 0.794971939 | -2.353541593 | 0.018595527 | 0.090653196 | 0.90653196 |
| Rpl35a | 21.166391 | -0.698817997 | 0.48587185 | -2.379671416 | 0.017328082 | 0.090653196 | 0.90653196 |
| Cdh23 | 22.590399 | 0.490137067 | 0.267811959 | 2.325636169 | 0.020037968 | 0.095302532 | 0.953025323 |
| 1700054A0 | 94.9814 | 0.518292182 | 0.295804881 | 2.314040765 | 0.020665486 | 0.095932771 | 0.95932771 |
| Rpl3 | 13.013179 | -0.144269504 | 0.515390803 | -2.305217309 | 0.021154406 | 0.095932771 | 0.95932771 |
| Uqcrh | 13.08399 | -0.67782227 | 0.741769172 | -2.265798572 | 0.023463719 | 0.103986936 | 1.039869358 |
| Adamtsl1 | 21.643253 | 0.532569425 | 0.332465914 | 2.233983843 | 0.025484136 | 0.109820042 | 1.098200416 |
| Rpl37 | 39.904488 | -0.570918528 | 0.39869311 | -2.227614001 | 0.025906266 | 0.109820042 | 1.098200416 |
| Rpl38 | 23.999256 | -0.144269504 | 0.515390806 | -2.215239549 | 0.026743627 | 0.1109576 | 1.109575999 |
| Hbb-bs | 28.182812 | 0.437629157 | 0.846217352 | 2.195825479 | 0.028104437 | 0.11193286 | 1.119328602 |
| Lars2 | 17.637733 | -0.623456797 | 0.554579944 | -2.195514405 | 0.028126719 | 0.11193286 | 1.119328602 |
| Mobp | 13.288434 | -0.606383084 | 0.492810989 | -2.176118414 | 0.029546407 | 0.115230989 | 1.152309887 |
| Negr1 | 14.903694 | -0.487609101 | 0.725189419 | -2.136712974 | 0.032621344 | 0.124728669 | 1.247286687 |
| Pde6a | 49.259518 | 0.483100547 | 0.315757125 | 2.126719968 | 0.033443357 | 0.12541259 | 1.254125901 |
| Sema3c | 51.47796 | 0.518252256 | 0.363382071 | 2.116357262 | 0.034314434 | 0.125768711 | 1.257687107 |
| Rps28 | 14.59144 | -0.540754452 | 0.403994441 | -2.110349552 | 0.034828258 | 0.125768711 | 1.257687107 |
| Rpl36 | 13.913916 | -0.554158874 | 0.611185034 | -2.092683826 | 0.036377388 | 0.125972863 | 1.259728632 |
| Rsrc1 | 30.191051 | 0.444198305 | 0.284116599 | 2.087722911 | 0.036822837 | 0.125972863 | 1.259728632 |
| Cox8a | 37.7302 | -0.551335192 | 0.520252382 | -2.101109331 | 0.03563137 | 0.125972863 | 1.259728632 |
| Cox7c | 20.10826 | -0.529881427 | 0.463622932 | -2.050891823 | 0.040277483 | 0.135415674 | 1.354156745 |
| mt-Nd5 | 26.785528 | -0.144269504 | 0.515390811 | -2.020120977 | 0.04337084 | 0.143344303 | 1.433443031 |
| Rpl23 | 16.993267 | -0.49864007 | 0.464485734 | -1.975370798 | 0.048226084 | 0.149595416 | 1.495954164 |
| Rps29 | 74.03566 | -0.469129154 | 0.561894576 | -1.974447981 | 0.048330827 | 0.149595416 | 1.495954164 |
| Zmat4 | 16.809929 | 0.521692452 | 0.516207563 | 1.994996601 | 0.046043251 | 0.149595416 | 1.495954164 |
| Hsp90ab1 | 17.179193 | -0.494443256 | 0.416652381 | -1.985324372 | 0.047108394 | 0.149595416 | 1.495954164 |
| Chsy3 | 15.987341 | -0.144269504 | 0.515390823 | 1.918128628 | 0.055094703 | 0.167866674 | 1.678666741 |
| Snap25 | 13.115135 | -0.45469839 | 0.525219881 | -1.890045533 | 0.058751871 | 0.176255612 | 1.762556122 |
| Nell1 | 18.427603 | 0.425605907 | 0.356328203 | 1.849298454 | 0.064414729 | 0.186734384 | 1.867343841 |
| Tpt1 | 21.913692 | -0.440827365 | 0.444097372 | -1.849192017 | 0.064430091 | 0.186734384 | 1.867343841 |
| Mapk14 | 31.587137 | 0.445784751 | 0.422126185 | 1.84444969 | 0.065117631 | 0.186734384 | 1.867343841 |
| Rps15 | 14.688202 | -0.432156259 | 0.444690587 | -1.817130155 | 0.069197186 | 0.193752453 | 1.937524528 |
| Sec61a1 | 116.780624 | -0.144269504 | 0.515390836 | 1.814816183 | 0.069552163 | 0.193752453 | 1.937524528 |
| Rpl11 | 28.021149 | -0.37465135 | 0.529624973 | -1.782476522 | 0.074671557 | 0.205056337 | 2.050563372 |
| Ttc3 | 13.601254 | -0.144269504 | 0.515390808 | -1.77612019 | 0.075713109 | 0.205056337 | 2.050563372 |
| Mbp | 73.51145 | -0.394041378 | 0.362497583 | -1.767493971 | 0.077145538 | 0.206073697 | 2.060736968 |
| Aldoc | 10.9872055 | -0.144269504 | 0.515390809 | -1.699663159 | 0.089194303 | 0.235039041 | 2.350390413 |
| 1700126G0 | 25.044415 | 0.358760442 | 0.318245153 | 1.67738862 | 0.093466513 | 0.239815396 | 2.39815396 |
| Acoxl | 16.777653 | 0.382808742 | 0.441048808 | 1.678057131 | 0.093335947 | 0.239815396 | 2.39815396 |
| Ttc28 | 20.06641 | 0.379961663 | 0.412603412 | 1.668708626 | 0.095175139 | 0.241027949 | 2.41027949 |
| mt-Co3 | 318.66357 | -0.295403109 | 0.228504023 | -1.644183919 | 0.100138218 | 0.250345544 | 2.503455439 |
| Fbxl7 | 11.705378 | 0.339689657 | 0.507509931 | 1.612173301 | 0.106924233 | 0.253212636 | 2.532126364 |
| Gm42418 | 525.09143 | -0.313783303 | 0.270107609 | -1.607457098 | 0.107954133 | 0.253212636 | 2.532126364 |
| Rpl32 | 18.241766 | -0.348953708 | 0.448505724 | -1.61601352 | 0.10609139 | 0.253212636 | 2.532126364 |
| Kctd16 | 54.999382 | 0.334404111 | 0.304539802 | 1.60235915 | 0.109076213 | 0.253212636 | 2.532126364 |
| 5830416I1 | 75.34862 | 0.331338663 | 0.296761284 | 1.602634916 | 0.10901528 | 0.253212636 | 2.532126364 |
| Eef1a1 | 19.555952 | -0.144269504 | 0.515390807 | -1.62617829 | 0.103911733 | 0.253212636 | 2.532126364 |
| Rps3a1 | 21.603226 | 0.347523648 | 0.441452341 | 1.576953511 | 0.114806222 | 0.263378981 | 2.63378981 |
| Rab29 | 13.005945 | 0.328105999 | 0.471836626 | 1.549974546 | 0.121147626 | 0.269335626 | 2.693356259 |
| D430042O0 | 46.858917 | 0.276666516 | 0.227216215 | 1.548315586 | 0.121546334 | 0.269335626 | 2.693356259 |
| Rpl13 | 26.57371 | -0.332717262 | 0.424241288 | -1.559850554 | 0.118795202 | 0.269335626 | 2.693356259 |
| Prkag2 | 13.786876 | 0.326263515 | 0.467116379 | 1.539067721 | 0.123787764 | 0.271220383 | 2.712203826 |
| Fth1 | 129.05562 | -0.317838388 | 0.322821726 | -1.529663623 | 0.126100012 | 0.273216693 | 2.73216693 |
| Rps23 | 18.492157 | -0.320013744 | 0.332614236 | -1.514327121 | 0.129942893 | 0.278449057 | 2.784490569 |
| Dop1b | 19.192867 | 0.287491945 | 0.511266043 | 1.498589606 | 0.133980131 | 0.283979625 | 2.839796254 |
| Rps24 | 29.452856 | -0.291179385 | 0.41968831 | -1.436841686 | 0.150762983 | 0.316115931 | 3.161159314 |
| Sil1 | 21.662725 | 0.298084343 | 0.391499069 | 1.411728009 | 0.158030063 | 0.327828322 | 3.27828322 |

| | | | | | | | |
|---|---|---|---|---|---|---|---|
| Cmpk1 | 38.822376 | 0.291500411 | 0.34224749 | 1.392669952 | 0.163719619 | 0.33605606 | 3.360560603 |
| Ankrd12 | 17.152811 | 0.247764133 | 0.4863706 | 1.356724569 | 0.174868743 | 0.355202134 | 3.552021342 |
| Rplp1 | 49.186565 | -0.248685321 | 0.251440839 | -1.333949212 | 0.182220502 | 0.366319565 | 3.663195651 |
| Rps27a | 24.053555 | -0.261571167 | 0.406200734 | -1.326222419 | 0.184766036 | 0.367646703 | 3.676467034 |
| Gria4 | 20.414885 | -0.144269504 | 0.515390827 | 1.315986867 | 0.188178451 | 0.370654524 | 3.706545241 |
| Rps16 | 14.018872 | -0.230105848 | 0.434761367 | -1.274748977 | 0.202398105 | 0.394071074 | 3.940710739 |
| Epha6 | 29.302063 | 0.257489032 | 0.389085895 | 1.267658721 | 0.204919852 | 0.394071074 | 3.940710739 |
| Rpl37a | 30.26386 | -0.246212883 | 0.387218712 | -1.253664347 | 0.209964032 | 0.394071074 | 3.940710739 |
| mt-Nd4l | 11.158327 | -0.235339661 | 0.425960081 | -1.260177981 | 0.207605164 | 0.394071074 | 3.940710739 |
| Ckb | 22.974964 | -0.247813248 | 0.383974292 | -1.253094714 | 0.210171239 | 0.394071074 | 3.940710739 |
| Fyb2 | 17.778055 | 0.240705016 | 0.408580585 | 1.219653686 | 0.222596186 | 0.413392916 | 4.13392916 |
| Rps27 | 22.39879 | -0.240422168 | 0.367904031 | -1.21160321 | 0.225664305 | 0.415137165 | 4.151371651 |
| Prkg1 | 23.337463 | 0.242338421 | 0.376000426 | 1.20534776 | 0.228069079 | 0.415639911 | 4.156399112 |
| Ms4a1 | 18.933796 | 0.235223927 | 0.386188401 | 1.182939019 | 0.236833316 | 0.42761571 | 4.276157098 |
| Shb | 38.882122 | 0.189925846 | 0.205193262 | 1.155982012 | 0.247688515 | 0.44311248 | 4.431124804 |
| Zbtb7a | 17.260128 | -0.144269504 | 0.51539082 | 1.140673801 | 0.254005693 | 0.450282819 | 4.502828192 |
| Ppia | 16.427113 | -0.20692797 | 0.401325243 | -1.122500828 | 0.261649556 | 0.455550565 | 4.555505655 |
| Scamp1 | 12.732742 | 0.214028782 | 0.41230597 | 1.12608206 | 0.260130787 | 0.455550565 | 4.555505655 |
| Selenow | 11.286031 | -0.190808482 | 0.420161419 | -1.104788016 | 0.269251455 | 0.456556814 | 4.565568143 |
| Rps10 | 15.577658 | -0.211191999 | 0.389131328 | -1.115512997 | 0.264630652 | 0.456556814 | 4.565568143 |
| 4930511M | 24.077318 | -0.144269504 | 0.51539082 | 1.108618193 | 0.267594926 | 0.456556814 | 4.565568143 |
| Mt1 | 25.606833 | -0.210607752 | 0.343704925 | -1.079579311 | 0.280329558 | 0.471243654 | 4.712436543 |
| Tmem108 | 15.926537 | 0.202918062 | 0.39051699 | 1.060300062 | 0.289008111 | 0.481680185 | 4.816801851 |
| Cmss1 | 62.80669 | -0.203876157 | 0.312280347 | -1.052926433 | 0.29237471 | 0.483161598 | 4.831615978 |
| Gm48099 | 16.995985 | -0.144269504 | 0.515390828 | 1.043903233 | 0.296530162 | 0.48591077 | 4.859107702 |
| Lrrc4c | 17.602245 | -0.146078184 | 0.426509765 | -1.030357278 | 0.30284232 | 0.488051673 | 4.880516734 |
| mt-Nd1 | 67.17599 | -0.196690667 | 0.284300487 | -1.032291318 | 0.301935667 | 0.488051673 | 4.880516734 |
| Mc1r | 25.007929 | 0.199571014 | 0.347811042 | 1.02247224 | 0.306557445 | 0.489989359 | 4.899893588 |
| Pcdh15 | 20.537975 | 0.197739483 | 0.357845454 | 1.016223283 | 0.309523071 | 0.490707308 | 4.907073083 |
| Fau | 19.209433 | -0.121612409 | 0.42652128 | -1.010603517 | 0.312206232 | 0.490969478 | 4.909694775 |
| Ppp3ca | 12.878865 | -0.183202491 | 0.375860022 | -0.982469576 | 0.325868564 | 0.50034937 | 5.003493703 |
| Actb | 15.762492 | -0.144269504 | 0.515390817 | 0.986263417 | 0.324003867 | 0.50034937 | 5.003493703 |
| Slx1b | 17.608353 | 0.192145177 | 0.348023792 | 0.9877404 | 0.323279803 | 0.50034937 | 5.003493703 |
| Rpl30 | 23.899248 | -0.18134221 | 0.348748162 | -0.954647833 | 0.339755819 | 0.516667759 | 5.166677592 |
| Tmsb4x | 30.03501 | 0.182770606 | 0.349717446 | 0.946499273 | 0.343893993 | 0.516667759 | 5.166677592 |
| Csmd3 | 17.87035 | 0.176099442 | 0.386827616 | 0.945418673 | 0.344445173 | 0.516667759 | 5.166677592 |
| Dlg2 | 16.44134 | -0.148199791 | 0.409549055 | -0.927933425 | 0.353442104 | 0.52611611 | 5.261161097 |
| Rps8 | 23.86433 | -0.168665573 | 0.352569402 | -0.899457314 | 0.368409123 | 0.543786417 | 5.437864166 |
| Rtn1 | 11.306291 | -0.144076559 | 0.409381743 | -0.894807084 | 0.370890223 | 0.543786417 | 5.437864166 |
| Rpl17 | 16.527699 | 0.143204765 | 0.431489111 | 0.874547124 | 0.381820369 | 0.547463029 | 5.474630293 |
| Syt1 | 13.347616 | -0.151120717 | 0.394231945 | -0.882019437 | 0.377766296 | 0.547463029 | 5.474630293 |
| Malat1 | 66.34335 | -0.166000777 | 0.314982595 | -0.87463642 | 0.381771764 | 0.547463029 | 5.474630293 |
| Gas5 | 25.984365 | -0.131686822 | 0.404870854 | -0.84648457 | 0.397282478 | 0.56547506 | 5.654750598 |
| Atp6v0c | 12.498774 | -0.1464398 | 0.382884035 | -0.833138902 | 0.404766396 | 0.568392316 | 5.683923155 |
| Adk | 13.152185 | 0.147374266 | 0.397466088 | 0.832438104 | 0.405161702 | 0.568392316 | 5.683923155 |
| Rpl27a | 23.546612 | -0.149676921 | 0.357118913 | -0.817226684 | 0.413798896 | 0.576362748 | 5.76362748 |
| Rps5 | 13.64657 | 0.139577389 | 0.403397451 | 0.803337472 | 0.421779705 | 0.580138373 | 5.801383733 |
| Nrxn3 | 17.17096 | -0.146285581 | 0.359299209 | -0.802161163 | 0.422459738 | 0.580138373 | 5.801383733 |
| Rpl34 | 15.662382 | -0.12788635 | 0.394702574 | -0.782317844 | 0.434027802 | 0.591856094 | 5.918560941 |
| Atp9b | 17.144903 | 0.137393748 | 0.387331606 | 0.773201305 | 0.439403254 | 0.59502524 | 5.950252396 |
| Rpl28 | 11.991038 | -0.128835167 | 0.38679862 | -0.758614097 | 0.448083438 | 0.602594968 | 6.025949677 |
| Atp1b1 | 16.694134 | -0.144269504 | 0.515390819 | -0.750395568 | 0.453016499 | 0.605056284 | 6.050562836 |
| Meg3 | 26.511145 | -0.092653632 | 0.415551518 | -0.739274535 | 0.459740309 | 0.609859594 | 6.098595937 |
| Rps18 | 12.561462 | -0.125203123 | 0.377606269 | -0.723425676 | 0.469418401 | 0.618490461 | 6.18490461 |
| Ubb | 11.475844 | -0.129990883 | 0.358791788 | -0.717830132 | 0.472862032 | 0.618846284 | 6.188462837 |
| Camk2n1 | 11.194849 | 0.116654562 | 0.401158028 | 0.692036285 | 0.488914542 | 0.619080102 | 6.190801024 |
| Rps4x | 11.205172 | -0.112164682 | 0.395151795 | -0.696847967 | 0.485897944 | 0.619080102 | 6.190801024 |
| Pcsk1n | 11.937058 | -0.110214919 | 0.40150873 | -0.71226996 | 0.476297622 | 0.619080102 | 6.190801024 |
| Ralyl | 14.434893 | -0.114306637 | 0.391458124 | -0.704135546 | 0.481348365 | 0.619080102 | 6.190801024 |
| Ptgds | 34.964783 | -0.130152116 | 0.319608819 | -0.70036098 | 0.483701898 | 0.619080102 | 6.190801024 |
| Igf2bp3 | 33.135895 | 0.123974988 | 0.365222526 | 0.685708595 | 0.492896886 | 0.620096082 | 6.200960819 |
| Rpl26 | 22.106962 | -0.121043753 | 0.348280105 | -0.670499035 | 0.502539721 | 0.628174652 | 6.281746517 |
| Calm1 | 24.714283 | -0.118035956 | 0.321873584 | -0.63887173 | 0.52290638 | 0.64535914 | 6.453591402 |
| Dbi | 12.409876 | -0.108156239 | 0.380688933 | -0.643114607 | 0.520149737 | 0.64535914 | 6.453591402 |
| Cadm2 | 11.268212 | -0.087508203 | 0.411506553 | -0.63356462 | 0.526364996 | 0.645541976 | 6.455419764 |

| | | | | | | | |
|---|---|---|---|---|---|---|---|
| Lsamp | 22.810705 | 0.113798103 | 0.299535681 | 0.610301016 | 0.541662424 | 0.660151079 | 6.601510793 |
| Ninj2 | 11.383703 | 0.095494033 | 0.384163077 | 0.558469988 | 0.576523493 | 0.698273796 | 6.982737958 |
| Cntnap2 | 44.181908 | 0.102246058 | 0.281789779 | 0.553081904 | 0.580207328 | 0.698397709 | 6.983977091 |
| Dpyd | 11.289407 | 0.089355794 | 0.391895307 | 0.537339873 | 0.591032868 | 0.702904321 | 7.029043211 |
| Mt3 | 24.288797 | -0.097775607 | 0.326947378 | -0.537154993 | 0.591160557 | 0.702904321 | 7.029043211 |
| Rps20 | 19.523159 | -0.08595201 | 0.345979602 | -0.486703285 | 0.626468617 | 0.740372002 | 7.403720017 |
| Camk2a | 17.494576 | 0.081601843 | 0.347922135 | 0.455089676 | 0.649044725 | 0.757866595 | 7.578665949 |
| Ndrg4 | 11.332241 | -0.059127343 | 0.412530519 | -0.455736495 | 0.648579475 | 0.757866595 | 7.578665949 |
| Ctnna3 | 14.824598 | 0.076646179 | 0.367488049 | 0.442789766 | 0.657917806 | 0.763654597 | 7.636545966 |
| Gm50306 | 25.903702 | 0.075156449 | 0.320357682 | 0.409610874 | 0.682091419 | 0.78702856 | 7.870285601 |
| Apoe | 57.67766 | 0.062376703 | 0.209560925 | 0.376376027 | 0.706637366 | 0.810554625 | 8.105546252 |
| Csmd1 | 12.766144 | 0.054823071 | 0.385057592 | 0.334817911 | 0.737762448 | 0.83158195 | 8.315819503 |
| Rpl35 | 11.108496 | -0.056229978 | 0.375246805 | -0.344325869 | 0.730601231 | 0.83158195 | 8.315819503 |
| Trim2 | 34.314507 | 0.061962293 | 0.283708386 | 0.33724833 | 0.735929707 | 0.83158195 | 8.315819503 |
| Rps6 | 11.909627 | -0.052766973 | 0.374287702 | -0.323588648 | 0.746249484 | 0.836314077 | 8.363140767 |
| Gm19951 | 13.942896 | 0.052418753 | 0.373874811 | 0.31131985 | 0.755557478 | 0.840986833 | 8.409868328 |
| Cox6c | 23.76232 | 0.053594892 | 0.35322194 | 0.306735737 | 0.759044526 | 0.840986833 | 8.409868328 |
| Ccdc60 | 13.773865 | 0.048051663 | 0.367238784 | 0.282249106 | 0.777752507 | 0.856845982 | 8.568459824 |
| Hexb | 16.373718 | -0.046220537 | 0.34618426 | -0.266289973 | 0.790015895 | 0.865466851 | 8.654668512 |
| Kcnip4 | 13.339096 | 0.040020412 | 0.366432281 | 0.234375782 | 0.814693258 | 0.873647428 | 8.736474275 |
| Galnt13 | 23.35115 | -0.144269504 | 0.515390824 | 0.238061937 | 0.811833056 | 0.873647428 | 8.736474275 |
| Rps12 | 14.798416 | -0.041995193 | 0.354724428 | -0.246593799 | 0.805222604 | 0.873647428 | 8.736474275 |
| Rpsa | 14.230893 | -0.033940719 | 0.395984302 | -0.23345995 | 0.815404266 | 0.873647428 | 8.736474275 |
| Cox6b1 | 12.227582 | -0.034194664 | 0.368965184 | -0.208043828 | 0.835194746 | 0.889961615 | 8.899616146 |
| Ntm | 48.500732 | 0.035965032 | 0.279435314 | 0.195949092 | 0.844650013 | 0.895006745 | 8.950067445 |
| Rps7 | 13.868416 | -0.030980067 | 0.370035541 | -0.190255449 | 0.849108963 | 0.895006745 | 8.950067445 |
| Sparcl1 | 14.376535 | 0.030717878 | 0.364781255 | 0.180459762 | 0.856791641 | 0.898249301 | 8.982493011 |
| 6030443J0 | 21.044775 | -0.144269504 | 0.515390817 | 0.170070334 | 0.864954824 | 0.901958239 | 9.019582387 |
| Cdk8 | 21.388523 | -0.027522941 | 0.328877794 | -0.155742409 | 0.876236068 | 0.908861879 | 9.088618787 |
| Psap | 11.310385 | -0.019352142 | 0.388272045 | -0.127379314 | 0.898640188 | 0.927168448 | 9.271684477 |
| Slc1a2 | 13.1157255 | -0.144269504 | 0.515390823 | -0.092637258 | 0.926191744 | 0.94066349 | 9.406634895 |
| Nkain2 | 17.19099 | -0.0175836 | 0.342703187 | -0.102618022 | 0.918266139 | 0.94066349 | 9.406634895 |
| Rpl23a | 11.890071 | -0.015313438 | 0.37846114 | -0.097453947 | 0.922365905 | 0.94066349 | 9.406634895 |
| Cpe | 19.586956 | 0.008678341 | 0.329744381 | 0.046977945 | 0.962530806 | 0.972505218 | 9.725052182 |
| Rpl9 | 18.87185 | 0.004742646 | 0.374393609 | 0.027544058 | 0.9780258 | 0.98306717 | 9.830671703 |
| Rps19 | 22.630238 | 0.000964211 | 0.328261058 | 0.005329168 | 0.995747959 | 0.995747959 | 9.95747959 |
| 4930402H2 | 9.076544 | -0.144269504 | 0.515390816 | 0.068855617 | 0.945104547 | | |
| 4930441J1 | 10.652756 | 0.373927387 | 0.525631222 | 1.706813944 | 0.08785664 | | |
| AI314278 | 6.1418185 | 0.189320512 | 0.478924769 | 1.158329558 | 0.246729573 | | |
| Ahdc1 | 8.7918 | 0.139793942 | 0.424580209 | 0.842070977 | 0.399748225 | | |
| Aldoa | 7.526311 | -0.355193299 | 0.583822088 | -1.843717368 | 0.065224341 | | |
| Alpl | 7.753427 | -0.00132107 | 0.39857568 | -0.010811304 | 0.991373995 | | |
| Anks1b | 8.893751 | -0.144269504 | 0.515390816 | -0.522073193 | 0.60161937 | | |
| Arvcf | 5.629838 | -0.144269504 | 0.515390817 | -1.18305288 | 0.236788191 | | |
| Atp5j2 | 8.170194 | -0.144269504 | 0.515390806 | -2.40210883 | 0.016300858 | | |
| Atxn7l1 | 9.241308 | -0.009343457 | 0.381617175 | -0.060204152 | 0.951993039 | | |
| Bc1 | 8.642321 | -0.154467482 | 0.440656662 | -1.144873145 | 0.25226171 | | |
| Cep128 | 9.068796 | 0.061100091 | 0.401767821 | 0.388567875 | 0.697595835 | | |
| Cfap74 | 6.1150217 | -0.144269504 | 0.515390816 | -1.501874188 | 0.133129604 | | |
| Chl1 | 9.726554 | 0.118268988 | 0.400820667 | 0.696998757 | 0.485803572 | | |
| Cntn3 | 9.957409 | 0.24326507 | 0.459088294 | 1.290550735 | 0.196859508 | | |
| Cspp1 | 9.700636 | 0.05660018 | 0.347015571 | 0.316968437 | 0.751267552 | | |
| Ctcflos | 10.201791 | 0.192391646 | 0.4801542 | 1.173043366 | 0.240778416 | | |
| Ctnna2 | 9.889812 | 0.101862432 | 0.400975039 | 0.61475461 | 0.538716789 | | |
| Dctd | 5.54887 | 0.145111382 | 0.46614346 | 0.961288894 | 0.336406931 | | |
| Eif1 | 8.936033 | -0.208675734 | 0.441848223 | -1.224280505 | 0.220846429 | | |
| Esf1 | 7.3313704 | -0.144269504 | 0.515390823 | 1.186673771 | 0.235356325 | | |
| Frmpd4 | 8.980852 | -0.144269504 | 0.515390813 | -0.370910914 | 0.710703886 | | |
| Gapdh | 10.714477 | -0.095160591 | 0.38412629 | -0.579698368 | 0.562118044 | | |
| Gm11337 | 9.823441 | 0.279396394 | 0.476549652 | 1.423504231 | 0.154590037 | | |
| Gm14582 | 9.58585 | -0.144269504 | 0.515390825 | 1.570059436 | 0.116401284 | | |
| Gm15564 | 5.3903313 | -0.497584635 | 0.844463149 | -2.20674779 | 0.027331684 | | |
| Gm29325 | 9.221433 | 0.134447835 | 0.417599943 | 0.804155148 | 0.421307378 | | |
| Gm38973 | 10.311735 | 0.878007167 | 0.649507061 | 2.436612559 | 0.014825555 | | |
| Gm49497 | 10.13219 | 0.163265614 | 0.437979705 | 0.972766927 | 0.330669154 | | |

| | | | | | |
|---|---|---|---|---|---|
| Gm49767 | 7.511084 | -0.144269504 | 0.515390819 | 1.627149577 | 0.103705334 |
| Gm765 | 10.8140335 | 0.175542196 | 0.390728744 | 0.945175628 | 0.34456922 |
| Gm960 | 6.197568 | -0.02373437 | 0.412069612 | -0.17278699 | 0.862818864 |
| Gnao1 | 9.669408 | 0.066313923 | 0.412744864 | 0.438561512 | 0.66097929 |
| Gnas | 6.836722 | 0.025873281 | 0.404725522 | 0.17105662 | 0.864179245 |
| Gpx4 | 9.617338 | -0.005119572 | 0.385663995 | -0.034585385 | 0.972410355 |
| Gria1 | 8.624296 | -0.144269504 | 0.515390817 | -1.135388113 | 0.256212738 |
| Gria2 | 8.166512 | -0.144269504 | 0.515390816 | -0.850312462 | 0.39515139 |
| Grik1 | 2.8808987 | 0.192517028 | 0.642301724 | 1.685332853 | 0.091924371 |
| Grik2 | 6.774379 | -0.144269504 | 0.515390824 | 0.102574301 | 0.91830084 |
| Grm7 | 5.1874037 | -0.237974591 | 0.503010688 | -1.571234776 | 0.116128125 |
| Hapln1 | 7.3785295 | -0.005584944 | 0.396266559 | -0.03856332 | 0.969238547 |
| Hdac8 | 9.471822 | 0.279308649 | 0.487114292 | 1.439506931 | 0.150006948 |
| Hdac9 | 7.803376 | 0.015356994 | 0.398324723 | 0.099534711 | 0.92071373 |
| Il1rapl2 | 7.6280694 | -0.087939065 | 0.401436544 | -0.572585322 | 0.5669255 |
| Jarid2 | 10.196609 | -0.153932273 | 0.405605973 | -0.915555271 | 0.359900201 |
| Kat7 | 10.592988 | 0.068976929 | 0.354250397 | 0.389104239 | 0.697199038 |
| Kcnd2 | 7.665193 | -0.025749107 | 0.394281563 | -0.170618828 | 0.864523492 |
| Kcnq5 | 10.671002 | -0.017427713 | 0.377114954 | -0.110028228 | 0.912386988 |
| Klhl14 | 2.5764966 | 0.077389556 | 0.49329248 | 0.727654759 | 0.466824951 |
| Klkb1 | 9.189222 | 0.254266387 | 0.496751041 | 1.393528589 | 0.163460006 |
| Ldhb | 8.447196 | -0.144269504 | 0.515390804 | -1.298280774 | 0.194190871 |
| Lrrc17 | 9.677987 | -0.403946849 | 0.579995355 | -1.864978314 | 0.062184444 |
| Lyrm1 | 6.137168 | -0.100425301 | 0.376197846 | -0.586609209 | 0.557466196 |
| Mrc2 | 10.618246 | 0.309919354 | 0.504520064 | 1.540037781 | 0.123551144 |
| Ndrg2 | 8.470592 | -0.043409968 | 0.415993989 | -0.339996779 | 0.733858953 |
| Nlgn1 | 9.592201 | -0.057322534 | 0.339932098 | -0.32220217 | 0.747299541 |
| Nrg1 | 7.5292664 | -0.157026983 | 0.444634971 | -1.182730129 | 0.236916119 |
| Nrxn1 | 9.978237 | -0.144269504 | 0.515390803 | -1.482403079 | 0.138233075 |
| Pcca | 6.166953 | 0.187068949 | 0.477128893 | 1.146457521 | 0.251605894 |
| Pcgf6 | 10.155868 | 0.123081383 | 0.408527758 | 0.733304643 | 0.463372644 |
| Phactr1 | 8.675378 | -0.144269504 | 0.515390806 | -0.975522861 | 0.329300968 |
| Phkg1 | 9.142755 | 0.296161993 | 0.51392013 | 1.521804333 | 0.128058116 |
| Pip5k1b | 7.6645293 | -0.144269504 | 0.515390828 | 1.016942407 | 0.309180828 |
| Plp1 | 8.522659 | -0.082720377 | 0.405148675 | -0.557768885 | 0.577002212 |
| Ppp1r16a | 9.776506 | 0.172874888 | 0.424119828 | 0.982558939 | 0.325824562 |
| Ppp1r9a | 9.570783 | 0.019314351 | 0.382006505 | 0.118017244 | 0.906053994 |
| Ptpn9 | 10.572884 | -0.144269504 | 0.515390831 | 2.346520509 | 0.018949623 |
| Ptprd | 10.320229 | 0.093256667 | 0.39451219 | 0.559824387 | 0.575599228 |
| Ptprg | 7.5578933 | 0.005166021 | 0.397441386 | 0.03227209 | 0.974255066 |
| Rabgap1l | 8.6605 | 0.182145954 | 0.441586333 | 1.051872887 | 0.292857873 |
| Rpl12 | 8.568083 | -0.00478262 | 0.391540132 | -0.032680205 | 0.973929609 |
| Rpl27 | 10.503557 | -0.233039863 | 0.448353314 | -1.312225481 | 0.189444067 |
| Rpl36a | 9.166282 | -0.012763381 | 0.392279777 | -0.084850259 | 0.932380437 |
| Rps17 | 10.353416 | -0.314030966 | 0.496696583 | -1.599412328 | 0.109729015 |
| Rps2 | 9.411554 | -0.127465606 | 0.398172044 | -0.773999648 | 0.438931 |
| Rps3 | 9.962103 | -0.049332963 | 0.381171178 | -0.308649948 | 0.757587821 |
| Sgip1 | 5.437327 | -0.104017483 | 0.422592393 | -0.728583135 | 0.466256698 |
| Slc25a4 | 9.770193 | -0.558095889 | 0.673636391 | -2.108969623 | 0.034947204 |
| Snhg11 | 9.443001 | 0.389592129 | 0.486766413 | 1.702646357 | 0.088634269 |
| Spock1 | 8.31042 | 0.240289317 | 0.488757267 | 1.33680717 | 0.181285588 |
| Srgn | 5.6584344 | 0.094392825 | 0.449510267 | 0.676297707 | 0.498851649 |
| St18 | 2.819745 | 0.262142555 | 0.672207835 | 1.852835462 | 0.063905946 |
| Sult2a2 | 7.8269706 | 0.042532181 | 0.400839588 | 0.274933165 | 0.783367586 |
| Svil | 8.817462 | 0.435566784 | 0.589043948 | 1.868897529 | 0.061637072 |
| Tacc1 | 7.43867 | 0.096459039 | 0.416643444 | 0.610763329 | 0.541356274 |
| Tmeff2 | 10.088047 | 0.260888381 | 0.473189885 | 1.36596197 | 0.1719509 |
| Tns3 | 9.338982 | 0.370113612 | 0.558774628 | 1.735296751 | 0.082688257 |
| Tshz2 | 3.0121968 | 0.14876937 | 0.527386323 | 1.186086086 | 0.235588305 |
| Tspan7 | 9.426498 | -0.214250367 | 0.443064682 | -1.24483993 | 0.213190595 |
| Uba52 | 10.173332 | -0.193057636 | 0.353237765 | -0.998830451 | 0.317876832 |
| Unc5d | 1.4916295 | 0.044017764 | 0.494217463 | 0.501520021 | 0.616005191 |
| Uqcr11 | 8.346692 | -0.144269504 | 0.515390821 | -0.226130543 | 0.82109989 |
| Vit | 9.474493 | 0.371044898 | 0.539641636 | 1.712441454 | 0.086815345 |
| Vwc2l | 6.578592 | -0.144269504 | 0.515390816 | 1.579523024 | 0.11421614 |

| | | | | | |
|---|---|---|---|---|---|
| Zfhx3 | 10.775464 | 0.234588481 | 0.441867393 | 1.23672154 | 0.216190479 |
| Zfp994 | 10.36792 | 0.148107992 | 0.439829105 | 0.913613542 | 0.360919948 |

**table S6: Differential gene expression list social compared to quinine in GABA and Glut neurons**

| **GABA** | baseMean | log2FoldChange | lfcSE | stat | pvalue | padj | pglobadj |
|---|---|---|---|---|---|---|---|
| Gm42418 | 197.87868 | 1.197690371 | 0.2243554 | 5.791100831 | 6.99E-09 | 3.59E-06 | 3.59E-05 |
| Cst3 | 38.42689 | 0.985627888 | 0.385962329 | 3.424868147 | 0.000615098 | 0.157772601 | 1.577726008 |
| Lrrc17 | 12.543546 | -0.144269504 | 0.366866616 | -3.139086352 | 0.001694755 | 0.289803102 | 2.898031018 |
| Rpl30 | 8.055436 | -2.384224174 | 0.952493846 | -2.979672291 | 0.002885569 | 0.370074231 | 3.70074231 |
| 2900097C1 | 8.523247 | 1.368672247 | 0.891404093 | 2.879420556 | 0.003984067 | 0.408765233 | 4.087652329 |
| mt-Co1 | 62.78899 | 0.579017104 | 0.398585912 | 2.762813162 | 0.005730556 | 0.489962532 | 4.899625323 |
| Ndrg4 | 3.9226623 | 0.03911124 | 0.217019253 | 2.483772079 | 0.012999898 | 0.654163557 | 6.541635571 |
| Ralgps1 | 6.0168247 | 0.055845582 | 0.199826142 | 2.425149793 | 0.015302072 | 0.654163557 | 6.541635571 |
| Frmpd4 | 6.773378 | 0.069697863 | 0.212136824 | 2.541796808 | 0.011028427 | 0.654163557 | 6.541635571 |
| Psap | 7.451934 | 0.035452896 | 0.257267162 | 2.438447019 | 0.014750521 | 0.654163557 | 6.541635571 |
| Vit | 7.636275 | -0.064632515 | 0.172880697 | -2.455087209 | 0.014085041 | 0.654163557 | 6.541635571 |
| Cfap74 | 6.8458595 | -0.074567698 | 0.18247769 | -2.539400544 | 0.011104261 | 0.654163557 | 6.541635571 |
| Gnas | 6.1879773 | 0.051704691 | 0.195891186 | 2.273371089 | 0.023003827 | 0.657646378 | 6.576463783 |
| Scamp1 | 11.683549 | -0.144269504 | 0.366866619 | -2.346856786 | 0.01893253 | 0.657646378 | 6.576463783 |
| Unc5d | 14.692937 | 0.0888368 | 0.221228176 | 2.274512828 | 0.022935171 | 0.657646378 | 6.576463783 |
| Nlgn1 | 3.4264915 | -0.144269504 | 0.366866632 | 2.30266744 | 0.021297561 | 0.657646378 | 6.576463783 |
| Cox7c | 6.407233 | 0.056693097 | 0.198812688 | 2.272185453 | 0.023075312 | 0.657646378 | 6.576463783 |
| Bc1 | 8.238236 | -0.062549543 | 0.171682801 | -2.371951726 | 0.017694402 | 0.657646378 | 6.576463783 |
| Rps27 | 7.522174 | 0.060458496 | 0.192857336 | 2.160746231 | 0.030714948 | 0.805755251 | 8.057552514 |
| mt-Co3 | 140.95633 | 0.125657773 | 0.256687856 | 2.132276374 | 0.032984133 | 0.805755251 | 8.057552514 |
| Fsip1 | 7.0084763 | -0.144269504 | 0.366866631 | 2.137376624 | 0.032567371 | 0.805755251 | 8.057552514 |
| Camk1d | 18.575476 | 0.089344095 | 0.212087001 | 2.105300997 | 0.035265115 | 0.822318372 | 8.223183724 |
| Fau | 4.5767965 | -0.041886126 | 0.157094573 | -2.068565728 | 0.038586857 | 0.860654675 | 8.606546752 |
| Ctnna2 | 10.305824 | -0.087083607 | 0.190764773 | -1.992222058 | 0.046346699 | 0.906023765 | 9.060237647 |
| mt-Atp6 | 116.925674 | 0.112777025 | 0.233435095 | 1.980160816 | 0.047685461 | 0.906023765 | 9.060237647 |
| Dlg2 | 8.813414 | 0.054859746 | 0.195812859 | 2.002786016 | 0.045200261 | 0.906023765 | 9.060237647 |
| Zeb1 | 4.422812 | -0.144269504 | 0.366866628 | -1.997037629 | 0.045821095 | 0.906023765 | 9.060237647 |
| Ntm | 48.24848 | -0.139682496 | 0.228591879 | -1.96230234 | 0.049727295 | 0.911075083 | 9.110750827 |
| Apoe | 17.863935 | -0.144269504 | 0.366866624 | 1.869302885 | 0.061580687 | 0.974260533 | 9.742605327 |
| Prnp | 3.9422488 | 0.034523893 | 0.182217825 | 1.86691656 | 0.061913241 | 0.974260533 | 9.742605327 |
| Atp6v1a | 4.2723627 | 0.042646281 | 0.169783285 | 1.885567521 | 0.059353258 | 0.974260533 | 9.742605327 |
| Nrxn3 | 4.7698565 | 0.040775963 | 0.184507746 | 1.861513258 | 0.06267173 | 0.974260533 | 9.742605327 |
| mt-Cytb | 44.338604 | 0.131777267 | 0.21841716 | 1.888957583 | 0.058897513 | 0.974260533 | 9.742605327 |
| Meis2 | 2.2039726 | 0.006257936 | 0.162678676 | 0.43161777 | 0.666019241 | 0.981446689 | 9.814466887 |
| Lingo2 | 5.8015375 | -0.02592222 | 0.153525667 | -1.405374293 | 0.159910007 | 0.981446689 | 9.814466887 |
| Lrmda | 2.7361138 | 0.010202067 | 0.163959704 | 0.649847148 | 0.515790961 | 0.981446689 | 9.814466887 |
| Prkg1 | 26.798965 | -0.144269504 | 0.366866614 | -1.675176805 | 0.093899547 | 0.981446689 | 9.814466887 |
| Prkag2 | 14.606763 | -0.020430175 | 0.152579427 | -0.61182021 | 0.540656717 | 0.981446689 | 9.814466887 |
| Ppp3ca | 3.36855 | -0.144269504 | 0.36686663 | 0.890757105 | 0.373059492 | 0.981446689 | 9.814466887 |
| Ppp1r9a | 13.191389 | -0.019217887 | 0.155367057 | -0.856440333 | 0.391754263 | 0.981446689 | 9.814466887 |
| Ppp1r16a | 14.036166 | -0.144269504 | 0.366866627 | 0.705636548 | 0.480414189 | 0.981446689 | 9.814466887 |
| Ppia | 1.793348 | -0.003874184 | 0.155711807 | -0.417853448 | 0.67605427 | 0.981446689 | 9.814466887 |
| Ppfia3 | 3.1646302 | -0.144269504 | 0.366866631 | 1.060009455 | 0.289140298 | 0.981446689 | 9.814466887 |
| Lrp1b | 193.3402 | -0.148823715 | 0.182679133 | -1.784944004 | 0.0742704 | 0.981446689 | 9.814466887 |
| Ppfia2 | 3.8350978 | 0.024032643 | 0.191602398 | 1.471240392 | 0.141226117 | 0.981446689 | 9.814466887 |
| Lrrc51 | 2.8095875 | -0.144269504 | 0.366866628 | -1.270889724 | 0.203767883 | 0.981446689 | 9.814466887 |
| Lrrc58 | 2.569153 | 0.011919554 | 0.166495055 | 0.787587685 | 0.43093792 | 0.981446689 | 9.814466887 |
| Ly75 | 3.9941173 | 0.013483565 | 0.162851057 | 0.716205061 | 0.473864738 | 0.981446689 | 9.814466887 |
| Pou6f2 | 1.2359202 | 0.014790062 | 0.174027585 | 1.370591434 | 0.170502355 | 0.981446689 | 9.814466887 |
| Plp1 | 2.1562288 | -0.010063353 | 0.154092025 | -0.739875842 | 0.459375334 | 0.981446689 | 9.814466887 |
| Plekhg1 | 3.2453194 | -0.144269504 | 0.366866626 | -1.528145856 | 0.126476328 | 0.981446689 | 9.814466887 |
| Pip5k1b | 7.5735574 | 0.01697151 | 0.170770266 | 0.79620786 | 0.42591123 | 0.981446689 | 9.814466887 |
| Phkg1 | 13.211901 | 0.011194853 | 0.153430596 | 0.3372723 | 0.735911639 | 0.981446689 | 9.814466887 |
| Phactr1 | 4.014428 | -0.014888957 | 0.154457992 | -0.87954701 | 0.379104756 | 0.981446689 | 9.814466887 |
| Lin28b | 7.330146 | 0.01028969 | 0.158177707 | 0.42874132 | 0.668111487 | 0.981446689 | 9.814466887 |
| Kmt2c | 3.5601394 | -0.144269504 | 0.366866631 | -0.468329376 | 0.639549064 | 0.981446689 | 9.814466887 |
| Ptgds | 18.82497 | 0.006270757 | 0.160148328 | 0.214306916 | 0.830307725 | 0.981446689 | 9.814466887 |
| Ptger3 | 8.53652 | 0.008850367 | 0.158411896 | 0.36654604 | 0.713957664 | 0.981446689 | 9.814466887 |

| | | | | | | | |
|---|---|---|---|---|---|---|---|
| Kirrel3 | 3.1277974 | -0.002239318 | 0.159785555 | -0.218864151 | 0.826755872 | 0.981446689 | 9.814466887 |
| Rpl17 | 3.1845405 | 0.00842538 | 0.169754077 | 0.627080561 | 0.530606428 | 0.981446689 | 9.814466887 |
| Klf10 | 5.660973 | -0.144269504 | 0.36686663 | 0.41546885 | 0.677798714 | 0.981446689 | 9.814466887 |
| Rpl13a | 4.6201034 | -0.011495203 | 0.155487262 | -0.666892276 | 0.504840945 | 0.981446689 | 9.814466887 |
| Rpl13 | 9.596256 | -0.018430915 | 0.156333155 | -0.785293029 | 0.432281782 | 0.981446689 | 9.814466887 |
| Robo2 | 2.420084 | 0.004275957 | 0.160221581 | 0.278398231 | 0.780706677 | 0.981446689 | 9.814466887 |
| Rnf180 | 3.4050643 | -0.015818433 | 0.153921004 | -0.913196476 | 0.361139217 | 0.981446689 | 9.814466887 |
| Riok3 | 3.8580315 | 0.018174813 | 0.180048086 | 1.157868509 | 0.2469177 | 0.981446689 | 9.814466887 |
| Klhl14 | 14.641721 | -0.144269504 | 0.366866627 | -1.096497432 | 0.272861146 | 0.981446689 | 9.814466887 |
| Pecam1 | 4.417114 | 0.020790958 | 0.166963163 | 1.009056666 | 0.312947459 | 0.981446689 | 9.814466887 |
| Rgs6 | 22.064264 | -0.032698153 | 0.157843978 | -0.995530447 | 0.319478344 | 0.981446689 | 9.814466887 |
| Rbfox1 | 2.495619 | 0.008404145 | 0.175948595 | 0.679180135 | 0.497023733 | 0.981446689 | 9.814466887 |
| Rapgef4 | 39.855263 | -0.035522346 | 0.14513914 | -0.696043919 | 0.486401326 | 0.981446689 | 9.814466887 |
| Rap1gap2 | 3.9017208 | 0.009250046 | 0.160887823 | 0.524701752 | 0.599790539 | 0.981446689 | 9.814466887 |
| Ralyl | 11.024073 | -0.013142307 | 0.153831938 | -0.476004832 | 0.634070936 | 0.981446689 | 9.814466887 |
| Klhl32 | 2.5686674 | 0.005220835 | 0.159712247 | 0.341918769 | 0.732412024 | 0.981446689 | 9.814466887 |
| Rabgap1l | 8.62533 | -0.010397129 | 0.153723252 | -0.38949256 | 0.696911813 | 0.981446689 | 9.814466887 |
| Ptprg | 6.1265078 | -0.144269504 | 0.366866626 | -0.596442185 | 0.550879866 | 0.981446689 | 9.814466887 |
| Ptprd | 9.341485 | -0.020633313 | 0.154405513 | -1.051063121 | 0.293229601 | 0.981446689 | 9.814466887 |
| Ptpn9 | 15.398825 | -0.023854618 | 0.154088248 | -0.758685597 | 0.448040655 | 0.981446689 | 9.814466887 |
| Reps2 | 5.085646 | -0.144269504 | 0.366866631 | 1.598897273 | 0.10984343 | 0.981446689 | 9.814466887 |
| Meg3 | 10.497679 | -0.027394428 | 0.155725327 | -1.072820409 | 0.283351702 | 0.981446689 | 9.814466887 |
| Pde4d | 3.9666913 | -0.019448653 | 0.153004288 | -1.15783833 | 0.246930018 | 0.981446689 | 9.814466887 |
| Pcgf6 | 8.9621725 | -0.029970308 | 0.155942221 | -1.084911589 | 0.277960815 | 0.981446689 | 9.814466887 |
| Ninj2 | 14.324038 | -0.050548325 | 0.163744806 | -1.472347346 | 0.140927104 | 0.981446689 | 9.814466887 |
| Nin | 4.3187513 | -0.003425162 | 0.159146051 | -0.214530653 | 0.830133266 | 0.981446689 | 9.814466887 |
| Nell1 | 18.728252 | -0.016033026 | 0.147961775 | -0.40827196 | 0.683074023 | 0.981446689 | 9.814466887 |
| Ndst3 | 3.9188282 | -0.008203231 | 0.154582609 | -0.482336223 | 0.629567119 | 0.981446689 | 9.814466887 |
| Mar-03 | 2.4135928 | -0.006649034 | 0.154997032 | -0.395220608 | 0.692680085 | 0.981446689 | 9.814466887 |
| Nckap5 | 79.63872 | -0.049483048 | 0.129539323 | -0.763099264 | 0.445404199 | 0.981446689 | 9.814466887 |
| Ncam2 | 3.0140529 | -0.144269504 | 0.366866631 | 1.238500459 | 0.215530552 | 0.981446689 | 9.814466887 |
| Naa11 | 6.8538857 | -0.007234692 | 0.15511474 | -0.333079324 | 0.739074406 | 0.981446689 | 9.814466887 |
| Mbp | 8.120255 | 0.023198368 | 0.17321741 | 1.083110277 | 0.278759478 | 0.981446689 | 9.814466887 |
| Nkain2 | 12.778161 | -0.031120158 | 0.155401743 | -0.96616824 | 0.333960002 | 0.981446689 | 9.814466887 |
| Myo1d | 3.1124558 | 0.021738801 | 0.172716944 | 1.268990067 | 0.204444607 | 0.981446689 | 9.814466887 |
| Mt1 | 12.3967495 | 0.029767811 | 0.19852136 | 1.52755452 | 0.126623181 | 0.981446689 | 9.814466887 |
| Mrpl15 | 3.7977905 | -0.004894472 | 0.154336494 | -0.240036848 | 0.810301691 | 0.981446689 | 9.814466887 |
| Mc1r | 37.432 | -0.07195733 | 0.156828788 | -1.212955514 | 0.225146829 | 0.981446689 | 9.814466887 |
| Mrc2 | 16.814459 | 0.020343546 | 0.154354598 | 0.56074339 | 0.574972483 | 0.981446689 | 9.814466887 |
| Mobp | 2.2887347 | 0.003105663 | 0.159411206 | 0.243125222 | 0.807908393 | 0.981446689 | 9.814466887 |
| Mir99ahg | 2.030341 | 0.003733506 | 0.160665525 | 0.291915056 | 0.770351572 | 0.981446689 | 9.814466887 |
| Micu1 | 3.0098612 | -0.144269504 | 0.36686663 | 1.238500443 | 0.215530558 | 0.981446689 | 9.814466887 |
| Mia2 | 3.6693451 | -0.016985515 | 0.15314591 | -1.028550362 | 0.303691014 | 0.981446689 | 9.814466887 |
| Meis3 | 3.5685163 | -0.011097955 | 0.153464121 | -0.822730587 | 0.410661217 | 0.981446689 | 9.814466887 |
| Mt3 | 3.8989177 | -0.144269504 | 0.366866631 | -0.440574964 | 0.659520731 | 0.981446689 | 9.814466887 |
| Mapk6 | 3.7598083 | -0.144269504 | 0.366866629 | 0.936787934 | 0.348867653 | 0.981446689 | 9.814466887 |
| Mapk14 | 35.734413 | 0.04999337 | 0.155549578 | 0.987219324 | 0.32353513 | 0.981446689 | 9.814466887 |
| Nqo2 | 3.6422973 | 0.013506752 | 0.168439101 | 0.787150004 | 0.43119406 | 0.981446689 | 9.814466887 |
| Macrod1 | 3.5645714 | -0.144269504 | 0.36686663 | 1.253846598 | 0.209897768 | 0.981446689 | 9.814466887 |
| Pcdh9 | 3.8833435 | 0.008031268 | 0.172176579 | 0.594872989 | 0.551928374 | 0.981446689 | 9.814466887 |
| Pcdh17 | 3.282865 | -0.004261721 | 0.157728048 | -0.332744903 | 0.739326851 | 0.981446689 | 9.814466887 |
| Pcdh15 | 20.877182 | -0.021566894 | 0.151002667 | -0.583395631 | 0.559626998 | 0.981446689 | 9.814466887 |
| Pard3b | 4.22236 | -0.01560584 | 0.154075507 | -0.819349067 | 0.412587285 | 0.981446689 | 9.814466887 |
| Palld | 3.6743817 | -0.144269504 | 0.366866631 | 1.091718365 | 0.274956911 | 0.981446689 | 9.814466887 |
| Olfr905 | 2.8664994 | 0.007993811 | 0.163557616 | 0.513585723 | 0.607541664 | 0.981446689 | 9.814466887 |
| Olfr729 | 4.568175 | -0.020193567 | 0.154073404 | -1.019793604 | 0.307826357 | 0.981446689 | 9.814466887 |
| Olfr694 | 3.366053 | -0.144269504 | 0.366866631 | 0.577496617 | 0.563604023 | 0.981446689 | 9.814466887 |
| Ogfrl1 | 4.3952684 | 0.005507068 | 0.158419672 | 0.271566954 | 0.785955012 | 0.981446689 | 9.814466887 |
| Nup210l | 5.6336856 | 0.021836721 | 0.165587283 | 1.007060278 | 0.313905808 | 0.981446689 | 9.814466887 |
| Magi2 | 5.7723165 | 0.005877478 | 0.162438491 | 0.326392659 | 0.744127291 | 0.981446689 | 9.814466887 |
| Nsf | 3.0547614 | 0.022418574 | 0.202441807 | 1.561296467 | 0.118453817 | 0.981446689 | 9.814466887 |

| | | | | | | | |
|---|---|---|---|---|---|---|---|
| Rpl18a | 6.301051 | 0.01131383 | 0.17750349 | 0.67417772 | 0.500198332 | 0.981446689 | 9.814466887 |
| Nrxn1 | 5.7600465 | -0.006623854 | 0.156417672 | -0.477864204 | 0.632746861 | 0.981446689 | 9.814466887 |
| Maml3 | 2.465767 | -0.013475254 | 0.153349397 | -0.93425475 | 0.3501725 | 0.981446689 | 9.814466887 |
| Nrp2 | 5.550371 | -0.009967995 | 0.155976954 | -0.472556087 | 0.636529918 | 0.981446689 | 9.814466887 |
| Nrgn | 3.1027417 | -0.020739261 | 0.153057632 | -1.236192549 | 0.216387 | 0.981446689 | 9.814466887 |
| Nrg3 | 7.9420066 | 0.011422815 | 0.157151556 | 0.444140046 | 0.656941335 | 0.981446689 | 9.814466887 |
| Mapk10 | 4.4292445 | -0.005287695 | 0.156602282 | -0.274891931 | 0.783399267 | 0.981446689 | 9.814466887 |
| Nrg1 | 5.9068103 | 0.007975396 | 0.166905785 | 0.412975003 | 0.679624928 | 0.981446689 | 9.814466887 |
| Pde1c | 34.822823 | -0.030076348 | 0.145728478 | -0.62107598 | 0.534549633 | 0.981446689 | 9.814466887 |
| Rpl19 | 6.514001 | 0.013284306 | 0.167292766 | 0.638025038 | 0.523457381 | 0.981446689 | 9.814466887 |
| 1700030K( | 4.864253 | 0.006525969 | 0.159742383 | 0.31575406 | 0.75218919 | 0.981446689 | 9.814466887 |
| Rpl27a | 9.139463 | -0.019805855 | 0.156222691 | -0.863399192 | 0.387918024 | 0.981446689 | 9.814466887 |
| Tor1aip1 | 5.202605 | -0.021265867 | 0.154996665 | -1.013250907 | 0.31094033 | 0.981446689 | 9.814466887 |
| Tns3 | 9.678324 | -0.144269504 | 0.366866622 | -1.109950735 | 0.267020256 | 0.981446689 | 9.814466887 |
| Tnni3k | 3.2726853 | -0.010823744 | 0.153963236 | -0.679281219 | 0.496959694 | 0.981446689 | 9.814466887 |
| Tmsb4x | 9.689733 | 0.03005028 | 0.180315732 | 1.299494078 | 0.193774424 | 0.981446689 | 9.814466887 |
| Tmem108 | 17.578232 | -0.012296708 | 0.152738468 | -0.385431156 | 0.699918003 | 0.981446689 | 9.814466887 |
| Tmeff2 | 8.183288 | 0.013382476 | 0.169063368 | 0.603423133 | 0.546227237 | 0.981446689 | 9.814466887 |
| Tmed9 | 3.1894383 | -0.144269504 | 0.366866631 | -0.468329372 | 0.639549067 | 0.981446689 | 9.814466887 |
| Tlr4 | 4.3207636 | -0.008245531 | 0.155337738 | -0.441690922 | 0.65871288 | 0.981446689 | 9.814466887 |
| Tcea1 | 6.314889 | -0.009032693 | 0.157470519 | -0.538395779 | 0.59030384 | 0.981446689 | 9.814466887 |
| Tacc1 | 7.2924743 | -0.018192644 | 0.155478701 | -0.796925822 | 0.425494113 | 0.981446689 | 9.814466887 |
| Syt11 | 2.5023417 | -0.006141273 | 0.150434631 | -0.711552884 | 0.476741692 | 0.981446689 | 9.814466887 |
| Tpt1 | 2.7894385 | 0.003032255 | 0.17880335 | 0.308214944 | 0.757918782 | 0.981446689 | 9.814466887 |
| Syt1 | 8.153926 | 0.008601428 | 0.157651298 | 0.343772818 | 0.731017144 | 0.981446689 | 9.814466887 |
| Susd4 | 3.0219922 | -0.144269504 | 0.366866629 | -0.871564439 | 0.383446038 | 0.981446689 | 9.814466887 |
| Sult2a2 | 8.987222 | -0.026023334 | 0.155779196 | -0.960878046 | 0.336613491 | 0.981446689 | 9.814466887 |
| Stxbp5l | 11.460402 | -0.05448289 | 0.167034741 | -1.732576353 | 0.083170995 | 0.981446689 | 9.814466887 |
| Stx8 | 3.011309 | -0.019799781 | 0.152859078 | -1.242928723 | 0.2138941 | 0.981446689 | 9.814466887 |
| Strbp | 2.4765105 | 0.018570795 | 0.183852596 | 1.341847249 | 0.179645529 | 0.981446689 | 9.814466887 |
| Stard7 | 9.268483 | -0.029934407 | 0.15649491 | -1.062352953 | 0.288075484 | 0.981446689 | 9.814466887 |
| Stard13 | 4.7596574 | -0.00795985 | 0.156426876 | -0.410811658 | 0.681210644 | 0.981446689 | 9.814466887 |
| St18 | 33.618313 | -0.065479155 | 0.150425552 | -1.099631623 | 0.271492658 | 0.981446689 | 9.814466887 |
| Ssh2 | 5.726728 | -0.011091476 | 0.156356757 | -0.62586653 | 0.53140249 | 0.981446689 | 9.814466887 |
| Srgn | 5.1956344 | -0.026899044 | 0.154999759 | -1.293707176 | 0.195766586 | 0.981446689 | 9.814466887 |
| Srbd1 | 5.352723 | 0.024419606 | 0.169385578 | 1.174421725 | 0.240226147 | 0.981446689 | 9.814466887 |
| Svil | 10.100415 | -0.035521661 | 0.157748023 | -1.323750084 | 0.185586059 | 0.981446689 | 9.814466887 |
| Sptlc3 | 3.1596541 | 0.004957109 | 0.159385608 | 0.291967141 | 0.770311748 | 0.981446689 | 9.814466887 |
| Trim2 | 32.228504 | -0.144269504 | 0.366866614 | 0.973325548 | 0.330391529 | 0.981446689 | 9.814466887 |
| Tshr | 5.839035 | -0.014141735 | 0.155198042 | -0.640773537 | 0.521669829 | 0.981446689 | 9.814466887 |
| mt-Nd4 | 37.65131 | 0.036398213 | 0.14963744 | 0.731900885 | 0.464229066 | 0.981446689 | 9.814466887 |
| mt-Nd3 | 21.990744 | 0.045744071 | 0.169634688 | 1.182871879 | 0.236859928 | 0.981446689 | 9.814466887 |
| mt-Nd2 | 38.638184 | 0.074762467 | 0.178798605 | 1.457460034 | 0.144989436 | 0.981446689 | 9.814466887 |
| mt-Nd1 | 24.586586 | 0.048769138 | 0.161264486 | 1.093313057 | 0.274256372 | 0.981446689 | 9.814466887 |
| mt-Co2 | 71.412186 | 0.049729502 | 0.16243318 | 1.003484498 | 0.315627153 | 0.981446689 | 9.814466887 |
| mt-Atp8 | 4.24499 | 0.006179492 | 0.16366887 | 0.351112426 | 0.725504006 | 0.981446689 | 9.814466887 |
| Zmat4 | 17.611155 | -0.029206879 | 0.154637066 | -0.833622335 | 0.404493836 | 0.981446689 | 9.814466887 |
| Zfpm2 | 23.544079 | -0.023347259 | 0.149658973 | -0.571191969 | 0.567869521 | 0.981446689 | 9.814466887 |
| Zfp994 | 11.950861 | -0.009196385 | 0.156564327 | -0.376208933 | 0.706761575 | 0.981446689 | 9.814466887 |
| Zbtb7a | 23.660015 | -0.053973222 | 0.161015971 | -1.233487687 | 0.217393868 | 0.981446689 | 9.814466887 |
| Xlr5a | 83.52263 | -0.030915457 | 0.148764204 | -0.628501291 | 0.529675601 | 0.981446689 | 9.814466887 |
| Trim33 | 6.189843 | -0.012204028 | 0.154845513 | -0.554610674 | 0.579160986 | 0.981446689 | 9.814466887 |
| Xirp2 | 6.82658 | -0.144269504 | 0.366866627 | 0.275999894 | 0.782548148 | 0.981446689 | 9.814466887 |
| Wdr86 | 2.3811314 | 0.006983558 | 0.162387526 | 0.486232538 | 0.626802304 | 0.981446689 | 9.814466887 |
| Wdr72 | 23.593578 | 0.011720249 | 0.148558422 | 0.282362918 | 0.777665246 | 0.981446689 | 9.814466887 |
| Wbscr25 | 2.1560493 | -0.004172076 | 0.15518894 | -0.361057566 | 0.718056414 | 0.981446689 | 9.814466887 |
| Vmn1r13 | 2.0559335 | -0.006813536 | 0.154188931 | -0.532045347 | 0.594694589 | 0.981446689 | 9.814466887 |
| Usp3 | 12.444709 | 0.014841518 | 0.164368855 | 0.551401581 | 0.581358417 | 0.981446689 | 9.814466887 |
| Uqcrh | 2.3927786 | 0.016277744 | 0.18050886 | 1.146226146 | 0.251701592 | 0.981446689 | 9.814466887 |
| Ube2cbp | 8.1268 | 0.01627297 | 0.162433097 | 0.646697033 | 0.517828044 | 0.981446689 | 9.814466887 |
| Ubb | 3.9243503 | 0.020488686 | 0.193142338 | 1.391794788 | 0.163984549 | 0.981446689 | 9.814466887 |

| | | | | | | | |
|---|---|---|---|---|---|---|---|
| Uba52 | 2.5813446 | 0.012397861 | 0.165349712 | 0.800307475 | 0.423532674 | 0.981446689 | 9.814466887 |
| Ttc3 | 2.3023152 | -0.012182868 | 0.15303757 | -0.866526973 | 0.386201241 | 0.981446689 | 9.814466887 |
| Ttc28 | 20.672173 | -0.042089831 | 0.156276225 | -1.061633161 | 0.288402253 | 0.981446689 | 9.814466887 |
| Wdyhv1 | 13.513174 | -0.021334091 | 0.153949961 | -0.664451566 | 0.506401339 | 0.981446689 | 9.814466887 |
| Rpl21 | 7.0366287 | 0.02434227 | 0.210487179 | 1.506218789 | 0.132011018 | 0.981446689 | 9.814466887 |
| Spock1 | 8.498045 | -0.044242513 | 0.158155122 | -1.260994219 | 0.20731093 | 0.981446689 | 9.814466887 |
| Spata16 | 3.1708157 | 0.015079741 | 0.157403513 | 0.749930719 | 0.453296432 | 0.981446689 | 9.814466887 |
| Rps3 | 2.4814267 | 0.003273204 | 0.162001159 | 0.225681872 | 0.821448858 | 0.981446689 | 9.814466887 |
| Rps29 | 19.558094 | -0.014640829 | 0.144153203 | -0.345599554 | 0.729643681 | 0.981446689 | 9.814466887 |
| Rps28 | 7.925555 | -0.022534882 | 0.155485452 | -0.951018195 | 0.34159514 | 0.981446689 | 9.814466887 |
| Rps27a | 6.605666 | 0.010339346 | 0.17862711 | 0.62948433 | 0.529032024 | 0.981446689 | 9.814466887 |
| Rps24 | 6.1713 | -0.003661753 | 0.160150204 | -0.305612245 | 0.759899896 | 0.981446689 | 9.814466887 |
| Rps23 | 3.547158 | 0.014100591 | 0.178481234 | 0.903236887 | 0.366400184 | 0.981446689 | 9.814466887 |
| Rps17 | 3.297554 | -0.003529607 | 0.15687499 | -0.227447839 | 0.820075524 | 0.981446689 | 9.814466887 |
| Rps15a | 3.5761046 | 0.017009398 | 0.189407059 | 1.272628068 | 0.203150054 | 0.981446689 | 9.814466887 |
| Rps15 | 6.495318 | 0.005645451 | 0.163952672 | 0.281373369 | 0.778424041 | 0.981446689 | 9.814466887 |
| Rps13 | 2.6396441 | -0.022201212 | 0.150637761 | -1.487737756 | 0.136820066 | 0.981446689 | 9.814466887 |
| Rps12 | 4.9291825 | 0.027794043 | 0.176488545 | 1.43057883 | 0.152550958 | 0.981446689 | 9.814466887 |
| Rps3a1 | 3.969914 | 0.009726652 | 0.158379795 | 0.464914632 | 0.641992592 | 0.981446689 | 9.814466887 |
| Rps10 | 4.5890503 | 0.019095094 | 0.176973026 | 1.055794685 | 0.291062027 | 0.981446689 | 9.814466887 |
| Rpl8 | 2.486145 | -0.006134001 | 0.156028116 | -0.465801264 | 0.64135776 | 0.981446689 | 9.814466887 |
| Rpl6 | 7.629247 | 0.039421123 | 0.16263484 | 1.306814341 | 0.191275774 | 0.981446689 | 9.814466887 |
| Rpl41 | 7.334943 | 0.021601559 | 0.165587871 | 0.94862895 | 0.342809362 | 0.981446689 | 9.814466887 |
| Rpl39 | 3.9424877 | 0.009563234 | 0.159393665 | 0.520886233 | 0.602446027 | 0.981446689 | 9.814466887 |
| Rpl37a | 8.407156 | -0.045453889 | 0.161967907 | -1.612368835 | 0.106881702 | 0.981446689 | 9.814466887 |
| Kctd16 | 68.58101 | -0.081403967 | 0.14977619 | -1.218320443 | 0.223102225 | 0.981446689 | 9.814466887 |
| Rpl37 | 12.240199 | 0.008439567 | 0.18699495 | 0.624838724 | 0.532076912 | 0.981446689 | 9.814466887 |
| Rpl36 | 2.4158921 | 0.003247298 | 0.16333679 | 0.235265325 | 0.814002806 | 0.981446689 | 9.814466887 |
| Rpl35a | 5.751913 | -0.007251069 | 0.157463444 | -0.402972299 | 0.686968603 | 0.981446689 | 9.814466887 |
| Rpl34 | 4.8314853 | 0.006518096 | 0.160157083 | 0.319662841 | 0.749223931 | 0.981446689 | 9.814466887 |
| Rpl32 | 2.143354 | -0.006654859 | 0.152766254 | -0.646161766 | 0.518174598 | 0.981446689 | 9.814466887 |
| Rplp1 | 13.382118 | -0.032798716 | 0.15522788 | -1.61315424 | 0.106711002 | 0.981446689 | 9.814466887 |
| Spdye4a | 4.774047 | -0.00487194 | 0.157653758 | -0.275260748 | 0.783115919 | 0.981446689 | 9.814466887 |
| Rps7 | 2.8617632 | 0.018156608 | 0.177158815 | 1.139730883 | 0.254398437 | 0.981446689 | 9.814466887 |
| Rsrc1 | 33.569794 | 0.013290256 | 0.14529975 | 0.282948985 | 0.777215944 | 0.981446689 | 9.814466887 |
| Sorcs2 | 10.033137 | -0.021320285 | 0.155686184 | -0.902840201 | 0.366610712 | 0.981446689 | 9.814466887 |
| Sntg1 | 1.8139235 | -0.006724471 | 0.155214011 | -0.51263213 | 0.608208674 | 0.981446689 | 9.814466887 |
| Snrpn | 1.9011647 | -0.005304756 | 0.154763345 | -0.419630822 | 0.674755168 | 0.981446689 | 9.814466887 |
| Snhg11 | 9.128835 | 0.004649514 | 0.162133648 | 0.227687098 | 0.819889503 | 0.981446689 | 9.814466887 |
| Snap25 | 3.0424485 | 0.029041216 | 0.161188396 | 1.543049959 | 0.122818655 | 0.981446689 | 9.814466887 |
| Smim13 | 1.2482914 | 0.005577689 | 0.176128675 | 0.549217465 | 0.58285622 | 0.981446689 | 9.814466887 |
| Slco2a1 | 5.0572495 | -0.032371008 | 0.156535331 | -1.352556307 | 0.176197419 | 0.981446689 | 9.814466887 |
| Slc4a10 | 2.2548785 | 0.015615075 | 0.21683363 | 1.401623499 | 0.161027706 | 0.981446689 | 9.814466887 |
| Slc31a2 | 2.889853 | 0.012118604 | 0.169391165 | 0.764017737 | 0.444856673 | 0.981446689 | 9.814466887 |
| Slc1a2 | 3.3066635 | 0.02169091 | 0.17621622 | 1.239538862 | 0.215146006 | 0.981446689 | 9.814466887 |
| Skap1 | 2.6202974 | 0.009262614 | 0.16383106 | 0.58077387 | 0.561392868 | 0.981446689 | 9.814466887 |
| Rps8 | 7.8138857 | 0.002214906 | 0.178453396 | 0.237478392 | 0.812285681 | 0.981446689 | 9.814466887 |
| Sil1 | 23.10908 | -0.144269504 | 0.366866615 | -0.803281543 | 0.421812024 | 0.981446689 | 9.814466887 |
| Sh3glb1 | 3.237924 | -0.144269504 | 0.36686663 | -0.757978704 | 0.448463732 | 0.981446689 | 9.814466887 |
| Sh3bgrl2 | 4.35456 | -0.013388176 | 0.153848256 | -0.742598761 | 0.457724639 | 0.981446689 | 9.814466887 |
| Sgip1 | 5.1121607 | -0.01047863 | 0.154612286 | -0.48516045 | 0.627562534 | 0.981446689 | 9.814466887 |
| Sgcz | 3.8964386 | -0.011784414 | 0.154188157 | -0.641072419 | 0.521475633 | 0.981446689 | 9.814466887 |
| Senp2 | 3.14891 | -0.031441818 | 0.153900492 | -1.725140379 | 0.084502177 | 0.981446689 | 9.814466887 |
| Sema3c | 152.0299 | -0.144269504 | 0.366866578 | 1.303515911 | 0.192398688 | 0.981446689 | 9.814466887 |
| Selenow | 5.3289976 | 0.006999659 | 0.170919689 | 0.419878726 | 0.67457405 | 0.981446689 | 9.814466887 |
| Sdccag8 | 8.908551 | 0.013619269 | 0.158384637 | 0.509994469 | 0.610055337 | 0.981446689 | 9.814466887 |
| Samd12 | 4.4635863 | -0.01557662 | 0.152033564 | -1.04569337 | 0.295702625 | 0.981446689 | 9.814466887 |
| Rubcnl | 9.297504 | 0.031905642 | 0.17253886 | 1.180611238 | 0.237757197 | 0.981446689 | 9.814466887 |
| Rtn1 | 3.8922057 | 0.004984742 | 0.166470301 | 0.337900686 | 0.735438031 | 0.981446689 | 9.814466887 |
| Shb | 44.75136 | -0.144269504 | 0.366866608 | -1.352491026 | 0.176218288 | 0.981446689 | 9.814466887 |
| Kcnq5 | 13.127025 | -0.144269504 | 0.366866624 | 0.336279373 | 0.736660203 | 0.981446689 | 9.814466887 |

| | | | | | | | |
|---|---|---|---|---|---|---|---|
| Lrrc4c | 11.133396 | -0.021466162 | 0.154262098 | -0.76397558 | 0.444881795 | 0.981446689 | 9.814466887 |
| mt-Nd5 | 7.1740417 | -0.144269504 | 0.366866629 | 0.34819663 | 0.727692518 | 0.981446689 | 9.814466887 |
| Crhr2 | 3.1307728 | -0.007151325 | 0.156381286 | -0.432679782 | 0.66524742 | 0.981446689 | 9.814466887 |
| AI314278 | 5.7923813 | -0.010640376 | 0.155252614 | -0.492600997 | 0.622294543 | 0.981446689 | 9.814466887 |
| Creb5 | 5.9548693 | -0.021394321 | 0.152974346 | -0.824064492 | 0.409902915 | 0.981446689 | 9.814466887 |
| Cox8a | 8.8485365 | 0.011946647 | 0.151473484 | 0.391490383 | 0.695434796 | 0.981446689 | 9.814466887 |
| Cox6c | 8.175585 | -0.025313439 | 0.15149398 | -0.771682046 | 0.440302767 | 0.981446689 | 9.814466887 |
| Cox6b1 | 4.8922834 | 0.01401261 | 0.169650369 | 0.725793111 | 0.467965611 | 0.981446689 | 9.814466887 |
| Cox4i1 | 7.1629577 | 0.014786492 | 0.173175615 | 0.79783796 | 0.424964529 | 0.981446689 | 9.814466887 |
| Cox11 | 3.43364 | -0.024521312 | 0.154139362 | -1.065947746 | 0.286447264 | 0.981446689 | 9.814466887 |
| AY036118 | 35.385612 | 0.070264072 | 0.200800771 | 1.805162391 | 0.071049274 | 0.981446689 | 9.814466887 |
| Comt | 7.2996964 | 0.013141833 | 0.170181057 | 0.715568776 | 0.474257659 | 0.981446689 | 9.814466887 |
| Csmd1 | 8.979083 | -0.032971157 | 0.157093554 | -1.337351772 | 0.181107839 | 0.981446689 | 9.814466887 |
| Col25a1 | 3.1188798 | -0.004488742 | 0.158480426 | -0.362183968 | 0.717214559 | 0.981446689 | 9.814466887 |
| Cntn3 | 7.9885926 | -0.029978736 | 0.15656984 | -1.136828686 | 0.255609909 | 0.981446689 | 9.814466887 |
| Cmss1 | 17.470535 | -0.144269504 | 0.366866617 | -1.711058106 | 0.087070387 | 0.981446689 | 9.814466887 |
| Cmpk1 | 45.31142 | 0.030611562 | 0.130937011 | 0.513152646 | 0.607844548 | 0.981446689 | 9.814466887 |
| Cmc1 | 3.3882625 | 0.018189078 | 0.160338665 | 0.982875861 | 0.32566854 | 0.981446689 | 9.814466887 |
| Ckb | 5.6728783 | 0.024430822 | 0.186177073 | 1.319411988 | 0.187031416 | 0.981446689 | 9.814466887 |
| Chsy3 | 15.00914 | -0.038134573 | 0.15756153 | -1.12688204 | 0.259792358 | 0.981446689 | 9.814466887 |
| Chl1 | 9.093343 | -0.024033252 | 0.154502891 | -0.856609739 | 0.391660602 | 0.981446689 | 9.814466887 |
| Chchd6 | 5.5926538 | -0.008775261 | 0.155301101 | -0.408183558 | 0.683138918 | 0.981446689 | 9.814466887 |
| Cep128 | 8.595963 | -0.144269504 | 0.366866622 | -1.312857621 | 0.189230929 | 0.981446689 | 9.814466887 |
| Cenpi | 5.506442 | -0.144269504 | 0.366866627 | -0.901115371 | 0.367526981 | 0.981446689 | 9.814466887 |
| Cntnap2 | 41.337933 | -0.071181369 | 0.145444686 | -1.112563652 | 0.265895878 | 0.981446689 | 9.814466887 |
| Cdk8 | 7.617289 | 0.019151604 | 0.179180278 | 0.966744248 | 0.333671902 | 0.981446689 | 9.814466887 |
| Csmd3 | 21.43251 | -0.144269504 | 0.366866618 | -0.505532243 | 0.613185058 | 0.981446689 | 9.814466887 |
| Kcnq2 | 3.3470104 | 0.009665872 | 0.15547643 | 0.520183521 | 0.60293567 | 0.981446689 | 9.814466887 |
| Eif2b3 | 3.8721156 | 0.015431576 | 0.164020166 | 0.828487161 | 0.407394663 | 0.981446689 | 9.814466887 |
| Ehbp1 | 31.22829 | 0.023328473 | 0.142670303 | 0.474299081 | 0.635286646 | 0.981446689 | 9.814466887 |
| Eef1a1 | 13.013707 | 0.061552233 | 0.183994837 | 1.755832257 | 0.079117055 | 0.981446689 | 9.814466887 |
| Dtnb | 6.7634826 | -0.013078828 | 0.154320779 | -0.607768372 | 0.543341109 | 0.981446689 | 9.814466887 |
| Dtd1 | 2.656812 | -0.016703043 | 0.152957017 | -1.080562632 | 0.279891712 | 0.981446689 | 9.814466887 |
| Dpyd | 15.116226 | -0.033299635 | 0.155304334 | -0.9575522 | 0.338288613 | 0.981446689 | 9.814466887 |
| Dppa2 | 7.833646 | 0.022223153 | 0.162755323 | 0.845931374 | 0.397591029 | 0.981446689 | 9.814466887 |
| Dpp10 | 5.468886 | -0.00935486 | 0.155687862 | -0.575579183 | 0.564899655 | 0.981446689 | 9.814466887 |
| Dop1b | 19.52095 | -0.144269504 | 0.366866619 | -0.237378107 | 0.812363472 | 0.981446689 | 9.814466887 |
| Dmd | 5.525551 | -0.020821552 | 0.154268103 | -1.04391893 | 0.296522899 | 0.981446689 | 9.814466887 |
| Cspp1 | 11.101294 | -0.022196937 | 0.153658332 | -0.729635658 | 0.46561292 | 0.981446689 | 9.814466887 |
| Disc1 | 5.831886 | -0.01492732 | 0.153763818 | -0.689064304 | 0.490782802 | 0.981446689 | 9.814466887 |
| Dclk1 | 3.4222398 | 0.003992121 | 0.160127459 | 0.248082431 | 0.804070628 | 0.981446689 | 9.814466887 |
| A930016O | 7.457064 | -0.144269504 | 0.366866628 | -0.557367759 | 0.577276189 | 0.981446689 | 9.814466887 |
| Dcc | 3.7580352 | -0.005023957 | 0.15812984 | -0.370946483 | 0.710677393 | 0.981446689 | 9.814466887 |
| Dbi | 4.200456 | -0.009958505 | 0.155276908 | -0.527162592 | 0.598080684 | 0.981446689 | 9.814466887 |
| Dach1 | 8.38423 | -0.144269504 | 0.366866628 | 0.668468391 | 0.503834653 | 0.981446689 | 9.814466887 |
| Dab1 | 2.1689057 | -0.003424469 | 0.157552256 | -0.241334651 | 0.809295757 | 0.981446689 | 9.814466887 |
| D430042O | 46.90667 | -0.02005274 | 0.129949311 | -0.348119709 | 0.727750283 | 0.981446689 | 9.814466887 |
| D030024E( | 3.0913346 | -0.011033967 | 0.154304862 | -0.703229501 | 0.481912737 | 0.981446689 | 9.814466887 |
| Ctnna3 | 20.55162 | -0.026890956 | 0.152977682 | -0.729547451 | 0.465666853 | 0.981446689 | 9.814466887 |
| Ctcflos | 14.059766 | -0.144269504 | 0.366866626 | 0.371910955 | 0.709959149 | 0.981446689 | 9.814466887 |
| Dctd | 8.773735 | 0.038024913 | 0.173324409 | 1.406949981 | 0.159442221 | 0.981446689 | 9.814466887 |
| Cdh8 | 4.101449 | 0.006948624 | 0.160256782 | 0.392970527 | 0.69434125 | 0.981446689 | 9.814466887 |
| Cdh23 | 23.583685 | 0.020405991 | 0.151004009 | 0.493946423 | 0.621344018 | 0.981446689 | 9.814466887 |
| Abat | 4.4407716 | -0.144269504 | 0.366866629 | -0.550686418 | 0.581848657 | 0.981446689 | 9.814466887 |
| Atp5e | 1.7737614 | 0.005253564 | 0.171863055 | 0.513496948 | 0.607603745 | 0.981446689 | 9.814466887 |
| Atp2b4 | 3.7729695 | -0.013926938 | 0.152688574 | -0.914378109 | 0.3605182 | 0.981446689 | 9.814466887 |
| Atp2b2 | 4.5780697 | 0.01739264 | 0.247240236 | 1.549351469 | 0.121297253 | 0.981446689 | 9.814466887 |
| Atp1b1 | 4.54629 | 0.024949964 | 0.177519682 | 1.315951148 | 0.18819044 | 0.981446689 | 9.814466887 |
| Actg1 | 3.5254388 | 0.014822292 | 0.188551273 | 1.133757549 | 0.256896262 | 0.981446689 | 9.814466887 |
| Atg13 | 1.7677553 | 0.004511849 | 0.162531478 | 0.355883714 | 0.721927653 | 0.981446689 | 9.814466887 |
| Asxl3 | 11.009017 | -0.014744538 | 0.154511322 | -0.573605628 | 0.566234703 | 0.981446689 | 9.814466887 |

| | | | | | | | |
|---|---|---|---|---|---|---|---|
| Astn2 | 3.277967 | 0.008249877 | 0.165519871 | 0.568196155 | 0.569901782 | 0.981446689 | 9.814466887 |
| Asic2 | 5.5682707 | -0.144269504 | 0.366866626 | -0.502208775 | 0.615520671 | 0.981446689 | 9.814466887 |
| Arvcf | 5.850197 | -0.007592244 | 0.158128481 | -0.405583209 | 0.685048868 | 0.981446689 | 9.814466887 |
| Atp5md | 7.0844955 | 0.005003513 | 0.173273681 | 0.36691044 | 0.713685824 | 0.981446689 | 9.814466887 |
| Arsb | 2.3654716 | -0.00920023 | 0.153178571 | -0.717247177 | 0.473221595 | 0.981446689 | 9.814466887 |
| Ano2 | 6.575887 | 0.00571601 | 0.158699545 | 0.253812472 | 0.799640437 | 0.981446689 | 9.814466887 |
| Adamtsl1 | 76.03911 | -0.022574999 | 0.132727939 | -0.390697394 | 0.696020926 | 0.981446689 | 9.814466887 |
| Ankrd12 | 14.751108 | 0.043643732 | 0.158136726 | 1.070935757 | 0.284198317 | 0.981446689 | 9.814466887 |
| Adgrg6 | 3.079143 | -0.144269504 | 0.366866629 | -0.871564492 | 0.383446009 | 0.981446689 | 9.814466887 |
| Alpl | 9.188238 | -0.015349965 | 0.154108938 | -0.566995635 | 0.570717147 | 0.981446689 | 9.814466887 |
| Alk | 6.4438066 | -0.144269504 | 0.366866628 | 0.594804152 | 0.551974392 | 0.981446689 | 9.814466887 |
| Aldoc | 4.788664 | 0.016244972 | 0.174465452 | 1.005980144 | 0.314425122 | 0.981446689 | 9.814466887 |
| Akap10 | 0.9909276 | 0.002142314 | 0.16769242 | 0.234285643 | 0.814763231 | 0.981446689 | 9.814466887 |
| Adk | 11.742018 | -0.008381326 | 0.151768671 | -0.265172512 | 0.79087657 | 0.981446689 | 9.814466887 |
| Ahcyl2 | 4.4735384 | -0.007555084 | 0.154914303 | -0.402046344 | 0.68764992 | 0.981446689 | 9.814466887 |
| Ap3s1 | 6.3816214 | 0.01745722 | 0.169304988 | 0.836036179 | 0.40313455 | 0.981446689 | 9.814466887 |
| Atp6v0c | 4.9662495 | 0.026024867 | 0.214731442 | 1.774730654 | 0.075942371 | 0.981446689 | 9.814466887 |
| Atp8a2 | 3.7063644 | 0.002708629 | 0.16906303 | 0.215848411 | 0.829105911 | 0.981446689 | 9.814466887 |
| Atp9b | 15.73347 | -0.144269504 | 0.366866621 | -1.405526418 | 0.159864799 | 0.981446689 | 9.814466887 |
| Cdh20 | 13.834691 | 0.010052644 | 0.144594042 | 0.251499816 | 0.801427704 | 0.981446689 | 9.814466887 |
| Cdh13 | 3.5069356 | 0.009188585 | 0.161749559 | 0.515600905 | 0.606133178 | 0.981446689 | 9.814466887 |
| Cdh12 | 5.969241 | -0.144269504 | 0.366866626 | -0.687779972 | 0.49159135 | 0.981446689 | 9.814466887 |
| Cdc14a | 2.7962465 | 0.016721414 | 0.181050359 | 1.207788855 | 0.227128489 | 0.981446689 | 9.814466887 |
| Ccdc60 | 15.320256 | -0.144269504 | 0.366866624 | 0.975628353 | 0.329248669 | 0.981446689 | 9.814466887 |
| Car12 | 4.1280303 | 0.007357637 | 0.167062902 | 0.49735047 | 0.618941927 | 0.981446689 | 9.814466887 |
| Camk2n1 | 1.9965439 | 0.009312544 | 0.167113905 | 0.684847882 | 0.493439917 | 0.981446689 | 9.814466887 |
| Camk2a | 15.446192 | -0.02525808 | 0.155620899 | -0.863332375 | 0.387954749 | 0.981446689 | 9.814466887 |
| Calm2 | 2.5644755 | 0.004295444 | 0.163014548 | 0.311646707 | 0.755309034 | 0.981446689 | 9.814466887 |
| Calm1 | 6.505156 | 0.013349538 | 0.174548599 | 0.778015525 | 0.436559861 | 0.981446689 | 9.814466887 |
| Calb1 | 3.5795054 | 0.005135102 | 0.162994727 | 0.346308706 | 0.729110727 | 0.981446689 | 9.814466887 |
| Abcg2 | 2.8091018 | -0.023108195 | 0.151667175 | -1.501629145 | 0.133192912 | 0.981446689 | 9.814466887 |
| Cadm2 | 5.785207 | -0.011969346 | 0.154952214 | -0.752540154 | 0.451726292 | 0.981446689 | 9.814466887 |
| Cacnb4 | 1.9863459 | 0.003855984 | 0.1709309 | 0.329767147 | 0.741575913 | 0.981446689 | 9.814466887 |
| C330002G( | 4.8138323 | -0.014398168 | 0.150287174 | -0.518598738 | 0.604040589 | 0.981446689 | 9.814466887 |
| C130073E2 | 3.6393151 | 0.004257268 | 0.160479924 | 0.234841445 | 0.814331799 | 0.981446689 | 9.814466887 |
| Brip1os | 4.872756 | 0.012938178 | 0.162095677 | 0.647586078 | 0.517252704 | 0.981446689 | 9.814466887 |
| Abhd6 | 2.1923254 | -0.005292153 | 0.154113583 | -0.43729512 | 0.661897334 | 0.981446689 | 9.814466887 |
| Bmt2 | 1.591141 | -0.008460111 | 0.151694659 | -0.791913473 | 0.42841113 | 0.981446689 | 9.814466887 |
| Bmpr1b | 1.2644954 | -0.144269504 | 0.36686663 | 1.60337848 | 0.108851119 | 0.981446689 | 9.814466887 |
| Bicc1 | 2.792174 | -0.010477635 | 0.154943083 | -0.66531745 | 0.505847472 | 0.981446689 | 9.814466887 |
| Bckdhb | 1.1911409 | 0.003332625 | 0.187550373 | 0.43568588 | 0.663064652 | 0.981446689 | 9.814466887 |
| Atxn7l1 | 8.993247 | -0.144269504 | 0.366866629 | 0.960404889 | 0.336851479 | 0.981446689 | 9.814466887 |
| Elf1 | 5.91902 | 0.014090323 | 0.159418334 | 0.595598923 | 0.551443196 | 0.981446689 | 9.814466887 |
| A630012P( | 11.478003 | -0.144269504 | 0.366866627 | 1.126145429 | 0.260103968 | 0.981446689 | 9.814466887 |
| Dnah1 | 2.6858728 | -0.009011369 | 0.155009151 | -0.607326814 | 0.543634047 | 0.981446689 | 9.814466887 |
| Gm42791 | 3.690221 | 0.023967831 | 0.177469242 | 1.303798768 | 0.192302203 | 0.981446689 | 9.814466887 |
| Gria1 | 4.716452 | -0.144269504 | 0.366866622 | -1.806104977 | 0.070901944 | 0.981446689 | 9.814466887 |
| Gpm6a | 5.923379 | 0.010528547 | 0.162675376 | 0.533093593 | 0.593968794 | 0.981446689 | 9.814466887 |
| Gpc5 | 1.6015186 | -0.011472328 | 0.151832908 | -0.936445043 | 0.349044096 | 0.981446689 | 9.814466887 |
| Gnaz | 38.746597 | 0.012984983 | 0.143978852 | 0.269803207 | 0.787311655 | 0.981446689 | 9.814466887 |
| 4930511M | 23.202368 | -0.022175757 | 0.154086193 | -0.627298864 | 0.530463348 | 0.981446689 | 9.814466887 |
| Gmcl1 | 4.490162 | 0.009898964 | 0.166138629 | 0.543966634 | 0.586464434 | 0.981446689 | 9.814466887 |
| Gm960 | 5.4595504 | 0.013256522 | 0.160517717 | 0.609669432 | 0.542080807 | 0.981446689 | 9.814466887 |
| Gm50306 | 26.180569 | 0.046001341 | 0.150009392 | 0.916092382 | 0.359618443 | 0.981446689 | 9.814466887 |
| Gm49797 | 5.4679165 | 0.009027775 | 0.166976796 | 0.480516688 | 0.63086004 | 0.981446689 | 9.814466887 |
| Gm49767 | 6.9007797 | -0.144269504 | 0.36686663 | 0.448064126 | 0.654106917 | 0.981446689 | 9.814466887 |
| Gria4 | 21.053968 | -0.144269504 | 0.366866616 | -0.92514306 | 0.354891497 | 0.981446689 | 9.814466887 |
| Gm49497 | 9.299484 | -0.030280174 | 0.156756836 | -1.237632218 | 0.215852463 | 0.981446689 | 9.814466887 |
| Gm48099 | 5.3664293 | 0.026797335 | 0.169630509 | 1.278587112 | 0.201042491 | 0.981446689 | 9.814466887 |
| 4930549G2 | 3.0712035 | -0.144269504 | 0.366866632 | 1.238500439 | 0.215530559 | 0.981446689 | 9.814466887 |
| Gm42889 | 7.961773 | -0.144269504 | 0.366866625 | -0.287105044 | 0.77403189 | 0.981446689 | 9.814466887 |

| | | | | | | | |
|---|---|---|---|---|---|---|---|
| Ephb2 | 4.2270985 | 0.018919586 | 0.16515236 | 0.9564546 | 0.338842611 | 0.981446689 | 9.814466887 |
| 4930558K0 | 17.679125 | -0.144269504 | 0.366866617 | -1.808629395 | 0.070508599 | 0.981446689 | 9.814466887 |
| Gm38304 | 3.9364216 | -0.018507768 | 0.153305023 | -1.065989339 | 0.286428461 | 0.981446689 | 9.814466887 |
| Gm37982 | 7.1370215 | -0.020579419 | 0.154767451 | -0.817165352 | 0.41383394 | 0.981446689 | 9.814466887 |
| Gm32647 | 2.3463705 | 0.012469188 | 0.157114618 | 0.780630536 | 0.435019827 | 0.981446689 | 9.814466887 |
| Gm29325 | 12.891184 | 0.00812042 | 0.157103739 | 0.273254253 | 0.784657779 | 0.981446689 | 9.814466887 |
| Gm2895 | 5.9506187 | -0.144269504 | 0.366866628 | 0.305058656 | 0.760321479 | 0.981446689 | 9.814466887 |
| Gm49171 | 2.0996242 | 0.023351799 | 0.184174645 | 1.609015595 | 0.107612932 | 0.981446689 | 9.814466887 |
| Grid2 | 7.6679535 | -0.005830755 | 0.154101821 | -0.229966808 | 0.818117562 | 0.981446689 | 9.814466887 |
| 4930441J1 | 10.034355 | 0.01545348 | 0.158223194 | 0.543405924 | 0.586850348 | 0.981446689 | 9.814466887 |
| Grik2 | 4.109635 | 0.010841403 | 0.167179646 | 0.655446772 | 0.512180146 | 0.981446689 | 9.814466887 |
| Kcnd2 | 4.998199 | -0.022634077 | 0.154281593 | -1.211572478 | 0.225676075 | 0.981446689 | 9.814466887 |
| Jakmip2 | 2.3758492 | -0.013598404 | 0.153325901 | -0.917133386 | 0.359072748 | 0.981446689 | 9.814466887 |
| Iqcb1 | 5.2411394 | -0.144269504 | 0.366866628 | 0.339257101 | 0.734416057 | 0.981446689 | 9.814466887 |
| 1700047F0 | 2.8115811 | -0.01801856 | 0.152805232 | -1.154500468 | 0.248295047 | 0.981446689 | 9.814466887 |
| 1700054A0 | 135.95944 | -0.05530318 | 0.130267095 | -0.839176003 | 0.401370551 | 0.981446689 | 9.814466887 |
| 1700102P0 | 3.330321 | -0.004367129 | 0.155767332 | -0.261088221 | 0.794024475 | 0.981446689 | 9.814466887 |
| Il23a | 5.4992266 | -0.007824651 | 0.155023955 | -0.355858692 | 0.721946392 | 0.981446689 | 9.814466887 |
| Il1rapl2 | 4.0735974 | -0.144269504 | 0.36686663 | -0.806600546 | 0.419896659 | 0.981446689 | 9.814466887 |
| Il1rapl1 | 4.5352893 | 0.005515254 | 0.180287518 | 0.442978226 | 0.657781484 | 0.981446689 | 9.814466887 |
| Igf2bp3 | 41.18624 | -0.10932327 | 0.170406967 | -1.526562524 | 0.126869832 | 0.981446689 | 9.814466887 |
| Igf2bp2 | 4.2253838 | -0.021074806 | 0.153314401 | -1.195421617 | 0.231922344 | 0.981446689 | 9.814466887 |
| 1700126G0 | 27.407665 | -0.020248706 | 0.152768697 | -0.531494675 | 0.59507603 | 0.981446689 | 9.814466887 |
| Hspa14 | 9.944516 | 0.034002299 | 0.173543091 | 1.302304193 | 0.192812419 | 0.981446689 | 9.814466887 |
| Hsp90ab1 | 2.9450097 | 0.028396543 | 0.157308199 | 1.407531888 | 0.159269729 | 0.981446689 | 9.814466887 |
| 2310002F0 | 4.5822587 | -0.028409696 | 0.154866262 | -1.464659968 | 0.143013712 | 0.981446689 | 9.814466887 |
| Hipk2 | 2.8712962 | 0.022309026 | 0.17469899 | 1.456457843 | 0.145266095 | 0.981446689 | 9.814466887 |
| Hexb | 5.7524843 | 0.027787978 | 0.17809757 | 1.421515524 | 0.155166944 | 0.981446689 | 9.814466887 |
| Hecw1 | 2.8604934 | 0.003469353 | 0.16723706 | 0.261165214 | 0.793965102 | 0.981446689 | 9.814466887 |
| 4930402H2 | 6.0613503 | -0.019177381 | 0.155159269 | -0.898795607 | 0.368761541 | 0.981446689 | 9.814466887 |
| Hdac9 | 8.86504 | -0.020742369 | 0.154219303 | -0.764465628 | 0.444589812 | 0.981446689 | 9.814466887 |
| Hapln1 | 12.379065 | -0.033450464 | 0.157592566 | -1.088137246 | 0.276534523 | 0.981446689 | 9.814466887 |
| Grm7 | 4.5105386 | -0.027013574 | 0.154471574 | -1.38358302 | 0.16648617 | 0.981446689 | 9.814466887 |
| Grin2b | 5.365261 | -0.009285081 | 0.15567361 | -0.609336605 | 0.542301349 | 0.981446689 | 9.814466887 |
| Gm28638 | 3.243386 | 0.007231271 | 0.160278801 | 0.437413894 | 0.66181121 | 0.981446689 | 9.814466887 |
| Gm28578 | 7.554862 | -0.014527994 | 0.154527786 | -0.592440746 | 0.553555488 | 0.981446689 | 9.814466887 |
| Agbl4 | 2.4272997 | -0.004289615 | 0.155792624 | -0.287032855 | 0.774087164 | 0.981446689 | 9.814466887 |
| Gm20342 | 8.248323 | 0.02935663 | 0.156348229 | 0.944999148 | 0.344659311 | 0.981446689 | 9.814466887 |
| Gas5 | 2.3400586 | -0.005257813 | 0.156736747 | -0.361932632 | 0.717402374 | 0.981446689 | 9.814466887 |
| Gm10062 | 54.468834 | -0.015231953 | 0.150753848 | -0.358945541 | 0.71963583 | 0.981446689 | 9.814466887 |
| Gm26901 | 5.5947323 | -0.144269504 | 0.366866631 | 1.459165746 | 0.144519494 | 0.981446689 | 9.814466887 |
| 8430422H0 | 31.612448 | -0.047872548 | 0.1546403 | -1.015731368 | 0.309757327 | 0.981446689 | 9.814466887 |
| Galnt17 | 4.862014 | -0.144269504 | 0.366866633 | 1.598897174 | 0.109843452 | 0.981446689 | 9.814466887 |
| Galnt13 | 22.919483 | -0.012247306 | 0.148919021 | -0.31103324 | 0.755775352 | 0.981446689 | 9.814466887 |
| Gm11337 | 9.638341 | -0.144269504 | 0.366866623 | -0.740231571 | 0.459159494 | 0.981446689 | 9.814466887 |
| Gabrb1 | 4.010501 | 0.013812042 | 0.16226126 | 0.723214457 | 0.469548138 | 0.981446689 | 9.814466887 |
| Gm11775 | 5.692351 | -0.144269504 | 0.366866629 | 1.32383579 | 0.185557587 | 0.981446689 | 9.814466887 |
| Fyb2 | 15.368901 | -0.037116214 | 0.149166821 | -0.851040506 | 0.394746852 | 0.981446689 | 9.814466887 |
| Fth1 | 20.526327 | 0.012087459 | 0.156361179 | 0.364243629 | 0.71567609 | 0.981446689 | 9.814466887 |
| Gm12216 | 78.83598 | 0.01698097 | 0.121251575 | 0.271027618 | 0.786369791 | 0.981446689 | 9.814466887 |
| Fstl4 | 4.5438514 | 0.006485491 | 0.158095418 | 0.319559849 | 0.749302016 | 0.981446689 | 9.814466887 |
| Foxn3 | 6.298606 | -0.005419982 | 0.159688352 | -0.318732156 | 0.749929628 | 0.981446689 | 9.814466887 |
| 5830416I1 | 88.99353 | 0.039278542 | 0.134871886 | 0.641379253 | 0.521276309 | 0.981446689 | 9.814466887 |
| Gm13219 | 2.3005614 | 0.008018063 | 0.170026585 | 0.565722645 | 0.571582339 | 0.981446689 | 9.814466887 |
| A230057D0 | 3.368232 | -0.020716881 | 0.147102396 | -1.619667767 | 0.105303664 | 0.981446689 | 9.814466887 |
| Gm19951 | 4.1404014 | 0.004058452 | 0.159263183 | 0.223114657 | 0.823446271 | 0.981446689 | 9.814466887 |
| Fam155a | 5.4177537 | 0.019600819 | 0.163345589 | 0.868810431 | 0.384950827 | 0.981446689 | 9.814466887 |
| Fam169a | 25.631237 | -0.020938422 | 0.145868095 | -0.467164962 | 0.64038186 | 0.981446689 | 9.814466887 |
| A330008L1 | 9.149897 | 0.01160861 | 0.157347561 | 0.450068319 | 0.65266118 | 0.981446689 | 9.814466887 |
| 4931429P1 | 4.9862547 | -0.144269504 | 0.366866631 | 0.339257115 | 0.734416046 | 0.981446689 | 9.814466887 |
| Fam53a | 3.7980964 | -0.144269504 | 0.366866631 | 0.577496454 | 0.563604134 | 0.981446689 | 9.814466887 |

| | | | | | | | |
|---|---|---|---|---|---|---|---|
| Gm17529 | 7.4276986 | 0.016471201 | 0.160865814 | 0.643180975 | 0.520106677 | 0.981446689 | 9.814466887 |
| Galntl6 | 6.7059746 | 0.03041902 | 0.183305256 | 1.525380221 | 0.127164289 | 0.981446689 | 9.814466887 |
| Gm14582 | 10.369181 | -0.010104813 | 0.153964521 | -0.351136242 | 0.72548614 | 0.981446689 | 9.814466887 |
| Fbxo4 | 5.469748 | -0.144269504 | 0.366866631 | 0.894222941 | 0.37120262 | 0.981446689 | 9.814466887 |
| Fgf12 | 5.036495 | 0.025292973 | 0.169194308 | 1.196450711 | 0.231520722 | 0.981446689 | 9.814466887 |
| Fgf13 | 2.5857751 | -0.013629296 | 0.153144032 | -0.909135429 | 0.363278643 | 0.981446689 | 9.814466887 |
| Gm14507 | 2.9591346 | -0.01163955 | 0.154633604 | -0.749271889 | 0.453693347 | 0.981446689 | 9.814466887 |
| Fgf14 | 4.405706 | 0.025866717 | 0.175815657 | 1.313923466 | 0.188871961 | 0.981446689 | 9.814466887 |
| 4932412D2 | 3.5255585 | -0.144269504 | 0.366866626 | -0.586891247 | 0.557276749 | 0.981446689 | 9.814466887 |
| Fgfr2 | 5.6097198 | -0.007263673 | 0.156660533 | -0.364462817 | 0.715512435 | 0.981446689 | 9.814466887 |
| Gphn | 5.080364 | 0.004443142 | 0.158740096 | 0.210672154 | 0.833143105 | 0.982534283 | 9.825342827 |
| Gria3 | 3.0391603 | -0.003267829 | 0.157453045 | -0.207685983 | 0.835474163 | 0.983023499 | 9.830234995 |
| Rps20 | 3.1889122 | 0.002225482 | 0.170323999 | 0.201629057 | 0.840206727 | 0.984077742 | 9.840777425 |
| Dlgap1 | 4.805951 | -0.144269504 | 0.366866627 | -0.201727149 | 0.840130037 | 0.984077742 | 9.840777425 |
| Snhg14 | 3.171061 | -0.002673598 | 0.156929911 | -0.174048427 | 0.861827404 | 0.986627254 | 9.866272544 |
| Wipf1 | 2.3992095 | 0.002479467 | 0.163648627 | 0.187810038 | 0.851025566 | 0.986627254 | 9.866272544 |
| Il27 | 26.21643 | 0.008054363 | 0.135683016 | 0.173645277 | 0.862144247 | 0.986627254 | 9.866272544 |
| Abcb4 | 2.1237674 | 0.002269934 | 0.159123289 | 0.171870608 | 0.863539254 | 0.986627254 | 9.866272544 |
| Celf2 | 7.3800097 | 0.002816977 | 0.167668537 | 0.183242551 | 0.854607709 | 0.986627254 | 9.866272544 |
| Arhgap18 | 2.9532552 | 0.002582673 | 0.158820728 | 0.172317824 | 0.863187673 | 0.986627254 | 9.866272544 |
| Tenm2 | 7.9103303 | -0.004002122 | 0.157310613 | -0.177156411 | 0.859385533 | 0.986627254 | 9.866272544 |
| Gm765 | 10.980562 | -0.0053696 | 0.152643771 | -0.178834967 | 0.858067288 | 0.986627254 | 9.866272544 |
| 6030443J0 | 22.903217 | -0.144269504 | 0.366866617 | -0.194802544 | 0.84554753 | 0.986627254 | 9.866272544 |
| Lyrm1 | 6.8762755 | -0.003794573 | 0.156575479 | -0.190372971 | 0.849016877 | 0.986627254 | 9.866272544 |
| Cpe | 5.8398447 | 0.003452187 | 0.159818894 | 0.178266635 | 0.85851358 | 0.986627254 | 9.866272544 |
| Rpl11 | 5.6096416 | 0.002901776 | 0.162587475 | 0.164845665 | 0.86906546 | 0.988537874 | 9.885378739 |
| Rab29 | 14.565226 | -0.144269504 | 0.366866622 | 0.166059498 | 0.868110127 | 0.988537874 | 9.885378739 |
| Tshz2 | 26.526703 | -0.144269504 | 0.366866615 | -0.161400258 | 0.871778164 | 0.989429641 | 9.894296415 |
| Col5a2 | 8.483255 | -0.004747567 | 0.149705401 | -0.157343902 | 0.874973823 | 0.990864395 | 9.908643952 |
| Ddx5 | 2.300076 | 0.002075387 | 0.163376134 | 0.148415578 | 0.882014807 | 0.992266658 | 9.92266658 |
| Kat7 | 13.281488 | -0.144269504 | 0.366866623 | -0.148439586 | 0.881995861 | 0.992266658 | 9.92266658 |
| Gm44593 | 5.707944 | 0.003070504 | 0.159219465 | 0.152137693 | 0.879078335 | 0.992266658 | 9.92266658 |
| Hdac8 | 11.37182 | -0.144269504 | 0.366866625 | -0.139950186 | 0.888699349 | 0.993712112 | 9.937121119 |
| Rpl12 | 3.47854 | 0.002372537 | 0.162771333 | 0.141507015 | 0.887469419 | 0.993712112 | 9.937121119 |
| Mast4 | 1.9890884 | -0.001227261 | 0.159370989 | -0.139429406 | 0.889110837 | 0.993712112 | 9.937121119 |
| Cldn16 | 7.197641 | -0.144269504 | 0.366866626 | -0.110640317 | 0.911901576 | 0.996008502 | 9.960085017 |
| Zfhx3 | 9.302843 | -0.003297737 | 0.155744591 | -0.133667674 | 0.893665368 | 0.996008502 | 9.960085017 |
| Vmn2r103 | 6.718278 | -0.144269504 | 0.366866627 | -0.104962603 | 0.916405483 | 0.996008502 | 9.960085017 |
| Ms4a1 | 24.781036 | -0.004154404 | 0.149434345 | -0.109524206 | 0.912786724 | 0.996008502 | 9.960085017 |
| Rps16 | 7.119069 | 0.002126031 | 0.159914303 | 0.108664355 | 0.913468715 | 0.996008502 | 9.960085017 |
| Sec61a1 | 145.17264 | -0.005821247 | 0.141780692 | -0.113755349 | 0.909431735 | 0.996008502 | 9.960085017 |
| Zfp236 | 8.02445 | -0.144269504 | 0.366866627 | -0.116040518 | 0.90762043 | 0.996008502 | 9.960085017 |
| Pkp2 | 8.761476 | 0.003587418 | 0.155029402 | 0.129547646 | 0.896924327 | 0.996008502 | 9.960085017 |
| Rpl26 | 12.747771 | 0.002773774 | 0.161374326 | 0.120113761 | 0.904393032 | 0.996008502 | 9.960085017 |
| Fmnl2 | 2.9049668 | -0.001647896 | 0.15831298 | -0.107350179 | 0.914511177 | 0.996008502 | 9.960085017 |
| Myt1l | 2.587045 | 0.001735035 | 0.160912891 | 0.117943246 | 0.906112627 | 0.996008502 | 9.960085017 |
| Slc24a3 | 7.1606812 | -0.144269504 | 0.366866629 | -0.110640314 | 0.911901578 | 0.996008502 | 9.960085017 |
| Rps14 | 5.149886 | -0.00192905 | 0.160510822 | -0.105221151 | 0.916200328 | 0.996008502 | 9.960085017 |
| Acoxl | 19.920992 | 0.001406782 | 0.153394921 | 0.039126207 | 0.968789767 | 0.998416399 | 9.984163993 |
| Vwc2l | 23.652805 | -0.144269504 | 0.36686662 | -0.042213443 | 0.966328547 | 0.998416399 | 9.984163993 |
| Usp37 | 4.724717 | -4.75E-05 | 0.157268435 | -0.003871391 | 0.996911084 | 0.998416399 | 9.984163993 |
| Kcnip4 | 8.848616 | -0.001499312 | 0.15752736 | -0.065097849 | 0.948096093 | 0.998416399 | 9.984163993 |
| Ank2 | 1.1421878 | 6.96E-05 | 0.171508797 | 0.015371444 | 0.987735845 | 0.998416399 | 9.984163993 |
| Pcsk2 | 6.9237137 | -0.001972948 | 0.15485897 | -0.085248087 | 0.932064162 | 0.998416399 | 9.984163993 |
| Gm12381 | 2.1627202 | -0.144269504 | 0.366866629 | -0.060600148 | 0.951677656 | 0.998416399 | 9.984163993 |
| Pcca | 10.118426 | 0.000604056 | 0.156782537 | 0.02196071 | 0.982479297 | 0.998416399 | 9.984163993 |
| Gm14164 | 5.9404206 | 0.001868038 | 0.157446793 | 0.081377027 | 0.935142118 | 0.998416399 | 9.984163993 |
| Olfr57 | 3.6490273 | 0.000222143 | 0.155653084 | 0.010565081 | 0.991570442 | 0.998416399 | 9.984163993 |
| Gm38973 | 12.814702 | -0.000716151 | 0.153911403 | -0.02631368 | 0.979007143 | 0.998416399 | 9.984163993 |
| Gm47371 | 5.4833465 | -0.000864958 | 0.158439029 | -0.04798263 | 0.961730086 | 0.998416399 | 9.984163993 |
| Gnao1 | 7.956073 | 0.002021692 | 0.156281332 | 0.077584831 | 0.938158309 | 0.998416399 | 9.984163993 |

| | | | | | | | |
|---|---|---|---|---|---|---|---|
| Grik1 | 504.84247 | 0.003171406 | 0.079819329 | 0.053311841 | 0.957483446 | 0.998416399 | 9.984163993 |
| Malat1 | 16.276428 | 0.002282616 | 0.160015073 | 0.088143408 | 0.929762696 | 0.998416399 | 9.984163993 |
| Lsamp | 17.453974 | -0.144269504 | 0.366866625 | -0.051700086 | 0.958767669 | 0.998416399 | 9.984163993 |
| mt-Nd4l | 2.988681 | 0.000623118 | 0.161919849 | 0.039512616 | 0.968481695 | 0.998416399 | 9.984163993 |
| Lars2 | 5.3278947 | -0.144269504 | 0.366866629 | -0.092568286 | 0.926246539 | 0.998416399 | 9.984163993 |
| Larp4 | 2.1627202 | -0.144269504 | 0.366866629 | -0.060600148 | 0.951677656 | 0.998416399 | 9.984163993 |
| Klkb1 | 10.227771 | -0.001503437 | 0.156737027 | -0.057359483 | 0.954258837 | 0.998416399 | 9.984163993 |
| Pde6a | 57.194782 | -0.003399989 | 0.140782842 | -0.067808572 | 0.94593802 | 0.998416399 | 9.984163993 |
| Ahdc1 | 11.586237 | -0.001985711 | 0.152444874 | -0.065631551 | 0.94767117 | 0.998416399 | 9.984163993 |
| Pgap1 | 3.0949821 | 0.000123368 | 0.161081805 | 0.006532681 | 0.994787712 | 0.998416399 | 9.984163993 |
| Fam189a2 | 5.841838 | -0.144269504 | 0.366866629 | 0.005261148 | 0.995802231 | 0.998416399 | 9.984163993 |
| Anks1b | 4.0464716 | -0.144269504 | 0.366866631 | 0.035291336 | 0.971847432 | 0.998416399 | 9.984163993 |
| Susd1 | 3.591877 | -0.144269504 | 0.366866627 | 0.054882257 | 0.956232267 | 0.998416399 | 9.984163993 |
| Bckdha | 3.9729373 | -0.000258948 | 0.158172046 | -0.015170079 | 0.987896493 | 0.998416399 | 9.984163993 |
| Spag17 | 2.8195398 | -0.000497384 | 0.160442918 | -0.036163115 | 0.971152297 | 0.998416399 | 9.984163993 |
| Cadps2 | 2.1912954 | -0.000935929 | 0.157280606 | -0.065632191 | 0.947670659 | 0.998416399 | 9.984163993 |
| Slx1b | 23.27221 | 0.003651607 | 0.150557818 | 0.092092417 | 0.926624613 | 0.998416399 | 9.984163993 |
| Slc39a2 | 3.634159 | -0.144269504 | 0.366866629 | 0.05488181 | 0.956232624 | 0.998416399 | 9.984163993 |
| Serinc1 | 2.093677 | 0.000518841 | 0.159836045 | 0.03848585 | 0.969300313 | 0.998416399 | 9.984163993 |
| Rtkn | 4.235541 | -0.000206793 | 0.161765489 | -0.017482554 | 0.986051651 | 0.998416399 | 9.984163993 |
| Rps26 | 4.4566765 | 0.000275426 | 0.163649815 | 0.020995837 | 0.983248976 | 0.998416399 | 9.984163993 |
| Rps21 | 11.421091 | -0.144269504 | 0.366866623 | 0.082929569 | 0.933907542 | 0.998416399 | 9.984163993 |
| Rpl9 | 4.063705 | -4.88E-05 | 0.157713421 | -0.004338432 | 0.996538443 | 0.998416399 | 9.984163993 |
| Rpl23 | 4.592517 | -0.00200323 | 0.149682272 | -0.085640574 | 0.931752143 | 0.998416399 | 9.984163993 |
| Epha6 | 35.940872 | 0.002407361 | 0.128728061 | -0.00198475 | 0.998416399 | 0.998416399 | 9.984163993 |
| Esf1 | 6.9082932 | 0.000460533 | 0.156860265 | 0.018246572 | 0.98544215 | 0.998416399 | 9.984163993 |
| Fbxl7 | 16.380646 | -0.000369165 | 0.145230743 | -0.008204766 | 0.993453617 | 0.998416399 | 9.984163993 |
| Rpl38 | 9.65513 | 0.001073254 | 0.16264574 | 0.052159944 | 0.958401249 | 0.998416399 | 9.984163993 |

| **Glut** | baseMean | log2FoldChange | lfcSE | stat | pvalue | padj | pglobadj |
|---|---|---|---|---|---|---|---|
| mt-Nd5 | 21.425013 | -0.144269504 | 0.416043185 | 5.027082516 | 4.98E-07 | 9.86E-05 | 0.000986097 |
| Hexb | 11.725084 | -0.144269504 | 0.416043188 | 4.914436708 | 8.90E-07 | 9.86E-05 | 0.000986097 |
| AY036118 | 74.46276 | 1.027500912 | 0.228672367 | 4.922157828 | 8.56E-07 | 9.86E-05 | 0.000986097 |
| mt-Co3 | 284.29404 | 0.672384944 | 0.145471885 | 4.943763596 | 7.66E-07 | 9.86E-05 | 0.000986097 |
| Ptgds | 95.20256 | -1.504625671 | 0.365220685 | -4.691165182 | 2.72E-06 | 0.000240685 | 0.00240685 |
| Mbp | 43.231262 | 2.129006466 | 0.546293973 | 4.444795746 | 8.80E-06 | 0.000649552 | 0.006495518 |
| Camk1d | 54.098747 | 0.95027269 | 0.243247769 | 4.384474017 | 1.16E-05 | 0.0007358 | 0.007358 |
| Atp1b1 | 11.707169 | 1.901636291 | 0.507085647 | 4.283547901 | 1.84E-05 | 0.001018548 | 0.010185482 |
| Cox6b1 | 10.22942 | 1.972558928 | 0.531880809 | 4.236838558 | 2.27E-05 | 0.001115814 | 0.011158137 |
| Cpe | 18.23968 | 1.570768983 | 0.47525144 | 3.93065272 | 8.47E-05 | 0.003752899 | 0.037528994 |
| Fth1 | 101.40413 | 1.146953095 | 0.356886782 | 3.816251581 | 0.000135494 | 0.005456727 | 0.054567273 |
| Hapln1 | 19.474379 | -0.144269504 | 0.416043164 | -3.671572023 | 0.000241063 | 0.008899247 | 0.088992469 |
| mt-Co1 | 131.00432 | 0.5265147 | 0.160883252 | 3.641134575 | 0.000271439 | 0.009249813 | 0.092498126 |
| Cst3 | 107.436134 | 1.395898809 | 0.480888149 | 3.59220723 | 0.000327889 | 0.009683653 | 0.096836532 |
| Rplp1 | 46.989296 | 0.856633665 | 0.284060531 | 3.59804269 | 0.000320621 | 0.009683653 | 0.096836532 |
| Cmss1 | 56.141346 | 1.007628223 | 0.348938539 | 3.530861159 | 0.000414209 | 0.011468415 | 0.114684147 |
| 6030443J0 | 42.34214 | -0.750774506 | 0.251230973 | -3.494655746 | 0.000474674 | 0.012369435 | 0.123694348 |
| Apoe | 61.643425 | 0.92950513 | 0.333561867 | 3.429802484 | 0.000604021 | 0.014865622 | 0.148656221 |
| Lrrc17 | 20.94026 | -1.495404641 | 0.568969604 | -3.410914627 | 0.000647454 | 0.015095892 | 0.15095892 |
| mt-Nd4 | 69.5036 | -0.144269504 | 0.416043162 | 3.32115321 | 0.000896463 | 0.018911103 | 0.189111029 |
| Xlr5a | 146.60124 | -0.144269504 | 0.416043107 | -3.329818923 | 0.000869025 | 0.018911103 | 0.189111029 |
| Bc1 | 16.422619 | -1.257745475 | 0.485712534 | -3.28076819 | 0.001035248 | 0.020846128 | 0.208461283 |
| Rpl12 | 7.536416 | -0.144269504 | 0.416043191 | 3.259738987 | 0.001115148 | 0.021478718 | 0.214787176 |
| mt-Co2 | 166.83102 | 0.67779233 | 0.256214866 | 3.201826853 | 0.001365591 | 0.025206525 | 0.252065253 |
| Rpl17 | 20.148624 | 1.173140672 | 0.497979701 | 3.174198032 | 0.001502511 | 0.026624502 | 0.266245021 |
| Galnt13 | 43.59496 | -0.144269504 | 0.416043154 | -3.098781189 | 0.001943185 | 0.0327695 | 0.327695002 |
| 5830416I1 | 163.03563 | -0.542535015 | 0.208298662 | -3.090642689 | 0.001997238 | 0.0327695 | 0.327695002 |
| mt-Atp6 | 232.6575 | 0.545184676 | 0.210965538 | 3.054655612 | 0.002253191 | 0.035648699 | 0.35648699 |
| Unc5d | 7.5450683 | -0.144269504 | 0.416043175 | -2.991522261 | 0.002775903 | 0.042404311 | 0.424043111 |
| Tpt1 | 17.57397 | -0.144269504 | 0.416043182 | 2.950742366 | 0.003170112 | 0.046198291 | 0.461982913 |
| Gm48099 | 17.89093 | 1.010319022 | 0.489210187 | 2.944684864 | 0.003232838 | 0.046198291 | 0.461982913 |

| | | | | | | | |
|---|---|---|---|---|---|---|---|
| Camk2n1 | 12.306559 | 1.197012205 | 0.607930581 | 2.928169176 | 0.003409645 | 0.046839036 | 0.468390359 |
| 4930511M | 53.437294 | -0.144269504 | 0.416043156 | -2.920996442 | 0.003489138 | 0.046839036 | 0.468390359 |
| Cox8a | 22.333979 | 1.025511954 | 0.515390913 | 2.896270288 | 0.00377627 | 0.049202576 | 0.492025763 |
| Ttc3 | 9.750542 | 1.198970366 | 0.690093072 | 2.79939042 | 0.005119919 | 0.063419362 | 0.634193619 |
| Hsp90ab1 | 9.8191185 | 1.263632542 | 0.737338803 | 2.797265404 | 0.005153718 | 0.063419362 | 0.634193619 |
| Gm42418 | 454.92178 | 0.562320146 | 0.252132989 | 2.787774289 | 0.005307151 | 0.063542371 | 0.635423706 |
| Tmsb4x | 33.171402 | 1.010136146 | 0.566766265 | 2.768524422 | 0.005631077 | 0.065646497 | 0.656464973 |
| Sec61a1 | 200.14557 | -0.28127733 | 0.111672174 | -2.697435527 | 0.006987581 | 0.079371758 | 0.793717582 |
| Mobp | 7.812506 | -0.144269504 | 0.416043186 | 2.674823876 | 0.007476852 | 0.081395758 | 0.813957582 |
| Rps3a1 | 28.6849 | 1.010304007 | 0.637591858 | 2.672303781 | 0.007533242 | 0.081395758 | 0.813957582 |
| Nrxn3 | 15.3793335 | 0.862805055 | 0.532822582 | 2.641791619 | 0.008246878 | 0.08496202 | 0.849620197 |
| Gm19951 | 13.628314 | 1.026194844 | 0.68207912 | 2.648091681 | 0.008094757 | 0.08496202 | 0.849620197 |
| Rps14 | 22.560736 | -0.144269504 | 0.41604318 | 2.619146756 | 0.008815002 | 0.088751038 | 0.887510384 |
| Ms4a1 | 40.452103 | -0.508597185 | 0.244465081 | -2.599882644 | 0.009325565 | 0.091805003 | 0.918050029 |
| Shb | 75.35348 | -0.144269504 | 0.416043139 | -2.559487998 | 0.010482648 | 0.100952454 | 1.009524535 |
| mt-Nd1 | 65.938736 | 0.522155717 | 0.272230411 | 2.530542966 | 0.011388614 | 0.107343741 | 1.073437405 |
| Actg1 | 8.641894 | 0.904841695 | 0.810626461 | 2.504016553 | 0.012279229 | 0.11332705 | 1.133270503 |
| Rsrc1 | 65.781166 | -0.381928646 | 0.183469286 | -2.450857734 | 0.014251627 | 0.128846341 | 1.288463409 |
| Gria4 | 39.796684 | -0.144269504 | 0.41604315 | -2.426107472 | 0.01526175 | 0.135219105 | 1.352191052 |
| Dab1 | 7.2935143 | 0.438755857 | 0.864972928 | 2.410351423 | 0.015937161 | 0.137432514 | 1.374325144 |
| Fgf14 | 8.012228 | 0.50762276 | 0.905598278 | 2.405914646 | 0.016132033 | 0.137432514 | 1.374325144 |
| Slx1b | 37.951138 | -0.144269504 | 0.416043148 | -2.327519113 | 0.019937653 | 0.159425766 | 1.594257659 |
| Rps24 | 24.425106 | 0.613150322 | 0.457644612 | 2.32348441 | 0.020153144 | 0.159425766 | 1.594257659 |
| Ube2cbp | 10.206972 | -0.144269504 | 0.416043179 | -2.335210065 | 0.01953245 | 0.159425766 | 1.594257659 |
| Rps20 | 19.378077 | 0.646943081 | 0.517880515 | 2.33330102 | 0.019632352 | 0.159425766 | 1.594257659 |
| Gas5 | 20.86644 | 0.654208454 | 0.661580238 | 2.308333266 | 0.020980608 | 0.16305981 | 1.630598098 |
| Ntm | 84.047455 | -0.391785472 | 0.215470142 | -2.271786398 | 0.023099415 | 0.176431737 | 1.764317374 |
| Cfap74 | 12.249493 | -0.144269504 | 0.416043175 | -2.246790484 | 0.02465342 | 0.180209074 | 1.802090742 |
| Mt3 | 22.130777 | 0.595821552 | 0.600896581 | 2.239909761 | 0.025096782 | 0.180209074 | 1.802090742 |
| Rapgef4 | 75.54937 | -0.144269504 | 0.416043123 | -2.237998782 | 0.025221135 | 0.180209074 | 1.802090742 |
| Gphn | 13.590273 | -0.144269504 | 0.416043182 | 2.252635455 | 0.024282144 | 0.180209074 | 1.802090742 |
| Hspa14 | 11.3665905 | -0.144269504 | 0.416043169 | -2.192456148 | 0.028346586 | 0.199325995 | 1.993259949 |
| Prkg1 | 49.208027 | -0.144269504 | 0.416043161 | -2.184350094 | 0.028936532 | 0.20029506 | 2.002950604 |
| Phactr1 | 8.39548 | -0.144269504 | 0.416043184 | 2.168440198 | 0.03012521 | 0.205314893 | 2.05314893 |
| Ctnna3 | 30.611032 | -0.487846511 | 0.430952363 | -2.091142985 | 0.036515248 | 0.245094774 | 2.450947737 |
| Arsb | 6.028934 | -0.144269504 | 0.416043188 | 2.036367517 | 0.041713465 | 0.275806938 | 2.758069383 |
| Atp6v1a | 7.590766 | -0.144269504 | 0.416043186 | 2.010439067 | 0.044384739 | 0.28880755 | 2.888075496 |
| Atp9b | 34.46911 | -0.144269504 | 0.41604316 | -2.004808078 | 0.044983569 | 0.28880755 | 2.888075496 |
| Lrp1b | 297.02847 | -0.210982773 | 0.11495446 | -1.997270052 | 0.045795855 | 0.289822339 | 2.898223386 |
| Syt1 | 13.465807 | 0.4413262 | 0.469192335 | 1.985142847 | 0.047128581 | 0.294055793 | 2.940557934 |
| Rpl27a | 21.897854 | 0.422647532 | 0.545785611 | 1.978807483 | 0.047837686 | 0.29433465 | 2.943346497 |
| Elf1 | 10.554319 | -0.391022856 | 0.563356345 | -1.956782255 | 0.050373074 | 0.300610494 | 3.006104935 |
| Mapk10 | 9.381102 | -0.415279384 | 0.567575269 | -1.958449792 | 0.050177257 | 0.300610494 | 3.006104935 |
| Mc1r | 56.35549 | -0.385158555 | 0.288290179 | -1.952377314 | 0.050893424 | 0.300610494 | 3.006104935 |
| A330008L1 | 13.030954 | -0.144269504 | 0.416043176 | -1.938642507 | 0.052544884 | 0.306281362 | 3.062813623 |
| Igf2bp2 | 6.105624 | -0.281829915 | 0.511697064 | -1.910589502 | 0.056057355 | 0.314346938 | 3.143469376 |
| Rpl9 | 18.211401 | 0.303384827 | 0.581919029 | 1.912778881 | 0.055776366 | 0.314346938 | 3.143469376 |
| Rps27a | 18.378384 | 0.409434165 | 0.481500273 | 1.917868149 | 0.055127732 | 0.314346938 | 3.143469376 |
| Cox6c | 25.040688 | 0.304326631 | 0.563678934 | 1.863516652 | 0.062389612 | 0.321379048 | 3.213790477 |
| mt-Nd4l | 7.357236 | 0.317920627 | 0.541352158 | 1.876883167 | 0.060534108 | 0.321379048 | 3.213790477 |
| Zbtb7a | 34.105064 | -0.144269504 | 0.416043167 | -1.876265945 | 0.060618771 | 0.321379048 | 3.213790477 |
| Gm10062 | 90.244774 | -0.250122632 | 0.153779497 | -1.864187096 | 0.062295435 | 0.321379048 | 3.213790477 |
| Nin | 7.5209975 | -0.144269504 | 0.416043174 | -1.892832417 | 0.058380159 | 0.321379048 | 3.213790477 |
| Fbxo4 | 7.84022 | 0.364756217 | 0.50477288 | 1.874895688 | 0.060807075 | 0.321379048 | 3.213790477 |
| Eef1a1 | 14.083366 | 0.359709592 | 0.52259201 | 1.869794479 | 0.061512363 | 0.321379048 | 3.213790477 |
| Cntnap2 | 76.74996 | -0.144269504 | 0.416043144 | -1.852918791 | 0.063893999 | 0.325345307 | 3.253453071 |
| Pcca | 14.824389 | -0.144269504 | 0.416043175 | -1.832267358 | 0.066911598 | 0.333054357 | 3.330543573 |
| Igf2bp3 | 67.86232 | -0.34015703 | 0.26133415 | -1.832792062 | 0.0668335 | 0.333054357 | 3.330543573 |
| mt-Nd3 | 47.639797 | 0.377712163 | 0.359488593 | 1.820253421 | 0.068720421 | 0.335938504 | 3.359385037 |
| Fstl4 | 7.3715515 | -0.144269504 | 0.416043174 | -1.818369455 | 0.069007684 | 0.335938504 | 3.359385037 |
| Ubb | 11.578455 | 0.358560632 | 0.432183824 | 1.788741497 | 0.073656457 | 0.354671851 | 3.546718506 |

| | | | | | | | |
|---|---|---|---|---|---|---|---|
| Nrg3 | 18.244083 | 0.360384849 | 0.420430502 | 1.776822943 | 0.075597376 | 0.360103629 | 3.601036286 |
| Rps26 | 12.015778 | 0.274506454 | 0.522638249 | 1.767811221 | 0.077092468 | 0.363318761 | 3.633187609 |
| Ddx5 | 6.023942 | 0.215891407 | 0.520729554 | 1.762221236 | 0.078031925 | 0.363875189 | 3.638751891 |
| Lars2 | 12.854558 | -0.144269504 | 0.416043181 | 1.753307675 | 0.079549209 | 0.36502579 | 3.650257898 |
| Slco2a1 | 12.440821 | -0.144269504 | 0.41604317 | -1.747833297 | 0.080492898 | 0.36502579 | 3.650257898 |
| Plp1 | 6.9241905 | 0.194985165 | 0.532254525 | 1.746347273 | 0.080750626 | 0.36502579 | 3.650257898 |
| Ppp1r9a | 21.5275 | -0.336840149 | 0.431586814 | -1.731965115 | 0.083279774 | 0.372655958 | 3.726559575 |
| Rpl19 | 17.258965 | 0.313379115 | 0.468776213 | 1.707416267 | 0.087744709 | 0.383216055 | 3.832160552 |
| Trim2 | 57.85163 | -0.323409223 | 0.282965503 | -1.711852162 | 0.086923916 | 0.383216055 | 3.832160552 |
| Stard7 | 15.240784 | -0.325640002 | 0.435530674 | -1.700164991 | 0.089099895 | 0.383216055 | 3.832160552 |
| Gm29325 | 19.812794 | -0.339782102 | 0.401489824 | -1.704023579 | 0.088376682 | 0.383216055 | 3.832160552 |
| Hdac8 | 21.701168 | -0.330157749 | 0.292612586 | -1.686908819 | 0.091620886 | 0.390269736 | 3.902697361 |
| Rps7 | 15.87695 | 0.308508936 | 0.447940019 | 1.670925566 | 0.094736381 | 0.399697303 | 3.99697303 |
| Rpl6 | 7.36718 | -0.221423327 | 0.439300767 | -1.646565982 | 0.099647287 | 0.416450456 | 4.164504561 |
| Uba52 | 8.7551985 | 0.240923191 | 0.484996757 | 1.640948072 | 0.100808195 | 0.417364769 | 4.173647694 |
| Cdk8 | 22.79811 | 0.304575158 | 0.401467798 | 1.605184105 | 0.108453294 | 0.444859347 | 4.448593469 |
| Anks1b | 9.436307 | 0.291756695 | 0.40712824 | 1.597100521 | 0.110243302 | 0.445439244 | 4.454392436 |
| Ctcflos | 24.750378 | -0.144269504 | 0.416043161 | -1.595476678 | 0.110605681 | 0.445439244 | 4.454392436 |
| Rpl39 | 14.220657 | 0.26567998 | 0.44209848 | 1.566051039 | 0.117336669 | 0.468289589 | 4.682895887 |
| Kctd16 | 108.661865 | -0.22966029 | 0.179454532 | -1.558817721 | 0.119039529 | 0.470843851 | 4.708438505 |
| Ank2 | 6.6865263 | 0.203275733 | 0.468101672 | 1.550224001 | 0.121087761 | 0.474706887 | 4.747068867 |
| Kmt2c | 7.6863174 | -0.144269504 | 0.416043181 | 1.543920044 | 0.122607704 | 0.476449234 | 4.76449234 |
| Slc4a10 | 9.207843 | -0.238103352 | 0.425226202 | -1.535804168 | 0.124586417 | 0.479928546 | 4.79928546 |
| Ninj2 | 24.909336 | -0.286902646 | 0.346965767 | -1.518019905 | 0.129009381 | 0.492682377 | 4.926823765 |
| Kat7 | 19.015747 | -0.144269504 | 0.41604317 | -1.511938665 | 0.130549467 | 0.494280784 | 4.94280784 |
| Gm17529 | 10.33267 | -0.261418564 | 0.408297584 | -1.507590204 | 0.131659441 | 0.494280784 | 4.94280784 |
| Il1rapl1 | 16.975739 | -0.186775637 | 0.409684723 | -1.48548102 | 0.137416447 | 0.506062059 | 5.060620592 |
| Cspp1 | 22.39606 | -0.27141802 | 0.371866706 | -1.488491475 | 0.136621329 | 0.506062059 | 5.060620592 |
| Psap | 7.0171747 | 0.211572603 | 0.443860681 | 1.482434854 | 0.138224626 | 0.506062059 | 5.060620592 |
| Rpl13 | 22.585205 | 0.244948032 | 0.426891329 | 1.468940846 | 0.141848835 | 0.515074047 | 5.150740468 |
| Ano2 | 13.782791 | -0.246021363 | 0.398413105 | -1.461383946 | 0.143910104 | 0.518310374 | 5.183103744 |
| Rps29 | 56.96026 | 0.260219714 | 0.269528435 | 1.44931084 | 0.147250795 | 0.526065341 | 5.260653409 |
| Scamp1 | 26.906717 | -0.144269504 | 0.416043162 | -1.417933394 | 0.156210212 | 0.55360899 | 5.536089901 |
| Pcsk2 | 15.964129 | -0.264781829 | 0.312332308 | -1.410029015 | 0.158531116 | 0.557375272 | 5.573752719 |
| Ppp3ca | 11.602921 | 0.212085602 | 0.420838239 | 1.385463701 | 0.165910724 | 0.570011812 | 5.700118122 |
| 1700126G( | 45.321327 | -0.232740475 | 0.224729778 | -1.39035685 | 0.164420543 | 0.570011812 | 5.700118122 |
| Calm1 | 21.541122 | 0.196126976 | 0.444954956 | 1.381022348 | 0.167272089 | 0.570011812 | 5.700118122 |
| Gm14164 | 9.191307 | -0.227744023 | 0.390203396 | -1.383431017 | 0.166532745 | 0.570011812 | 5.700118122 |
| Rpl41 | 23.7763 | -0.222894851 | 0.384259385 | -1.373868553 | 0.169482494 | 0.573135456 | 5.731354564 |
| mt-Cytb | 84.411514 | 0.226483365 | 0.225914955 | 1.358824654 | 0.174202159 | 0.584633003 | 5.846330033 |
| Rpl26 | 21.040993 | 0.230250473 | 0.379393178 | 1.343420779 | 0.179135762 | 0.596670246 | 5.96670246 |
| Il1rapl2 | 12.456991 | -0.238496668 | 0.345524453 | -1.326953348 | 0.184524115 | 0.609153302 | 6.091533017 |
| Atxn7l1 | 16.558521 | -0.231435102 | 0.353497677 | -1.323606923 | 0.185633625 | 0.609153302 | 6.091533017 |
| Ap3s1 | 15.414441 | -0.211322398 | 0.371262555 | -1.295929762 | 0.194999685 | 0.620583938 | 6.205839376 |
| Sdccag8 | 10.882109 | -0.200209282 | 0.382874112 | -1.30099619 | 0.193259759 | 0.620583938 | 6.205839376 |
| Rpl30 | 20.953156 | 0.167692365 | 0.444298231 | 1.294146951 | 0.195614666 | 0.620583938 | 6.205839376 |
| Riok3 | 9.238238 | -0.144269504 | 0.416043174 | -1.288643074 | 0.197522201 | 0.620583938 | 6.205839376 |
| Dpyd | 23.287388 | -0.219665558 | 0.354662647 | -1.292335412 | 0.196241014 | 0.620583938 | 6.205839376 |
| Tmed9 | 7.3592887 | 0.178126696 | 0.413792074 | 1.290882501 | 0.196744424 | 0.620583938 | 6.205839376 |
| Ppia | 14.577909 | 0.178375294 | 0.423177793 | 1.274220285 | 0.202585358 | 0.632009251 | 6.320092513 |
| Tor1aip1 | 9.50812 | 0.183960509 | 0.398346604 | 1.246170839 | 0.212701682 | 0.637796355 | 6.377963547 |
| Gm960 | 10.659881 | -0.18459301 | 0.375151734 | -1.22966577 | 0.21882229 | 0.637796355 | 6.377963547 |
| Nkain2 | 27.099146 | -0.144269504 | 0.416043169 | -1.238382318 | 0.215574334 | 0.637796355 | 6.377963547 |
| Rtn1 | 9.300041 | 0.139916447 | 0.449755411 | 1.257625005 | 0.208527406 | 0.637796355 | 6.377963547 |
| Stxbp5l | 19.681265 | -0.223588955 | 0.284908602 | -1.247715385 | 0.212135305 | 0.637796355 | 6.377963547 |
| Rps28 | 21.22325 | -0.178647808 | 0.379639205 | -1.238956939 | 0.215361446 | 0.637796355 | 6.377963547 |
| Atp5e | 12.293096 | 0.199374628 | 0.383491971 | 1.255550533 | 0.209278976 | 0.637796355 | 6.377963547 |
| Gm28578 | 14.677203 | -0.144269504 | 0.416043175 | -1.231453513 | 0.218153293 | 0.637796355 | 6.377963547 |
| Csmd3 | 35.537155 | -0.218567583 | 0.32179858 | -1.249165972 | 0.211604375 | 0.637796355 | 6.377963547 |
| 1700054A( | 195.7257 | -0.186142413 | 0.191912016 | -1.229624969 | 0.218837575 | 0.637796355 | 6.377963547 |
| Atp6v0c | 10.461972 | 0.170726317 | 0.397981066 | 1.190965012 | 0.233667321 | 0.651056833 | 6.510568327 |

| | | | | | | | |
|---|---|---|---|---|---|---|---|
| Gm49497 | 21.095951 | -0.210717139 | 0.312641308 | -1.201728611 | 0.229468692 | 0.651056833 | 6.510568327 |
| Mapk14 | 61.648254 | -0.144269504 | 0.416043134 | -1.201565642 | 0.22953186 | 0.651056833 | 6.510568327 |
| Cox4i1 | 23.908047 | 0.204818482 | 0.329191625 | 1.187210298 | 0.23514468 | 0.651056833 | 6.510568327 |
| Nlgn1 | 9.184643 | 0.156106593 | 0.41407251 | 1.19384498 | 0.232538613 | 0.651056833 | 6.510568327 |
| Ppp1r16a | 20.882874 | -0.20074756 | 0.338080384 | -1.189845809 | 0.234107001 | 0.651056833 | 6.510568327 |
| Snhg11 | 14.930077 | 0.18016391 | 0.393336525 | 1.198845833 | 0.230587897 | 0.651056833 | 6.510568327 |
| Snap25 | 9.105734 | 0.149799747 | 0.421572343 | 1.198042242 | 0.230900572 | 0.651056833 | 6.510568327 |
| Grm7 | 6.15634 | -0.128257826 | 0.380548339 | -1.167850458 | 0.242867091 | 0.657629036 | 6.576290358 |
| Col5a2 | 8.42301 | -0.154151012 | 0.378014223 | -1.171310287 | 0.241474078 | 0.657629036 | 6.576290358 |
| mt-Nd2 | 89.006584 | 0.198906558 | 0.242567259 | 1.173909136 | 0.240431423 | 0.657629036 | 6.576290358 |
| Slc24a3 | 14.504547 | -0.144269504 | 0.416043171 | -1.165235138 | 0.243923826 | 0.657629036 | 6.576290358 |
| Mapk6 | 5.9629583 | 0.142128027 | 0.409173641 | 1.162725635 | 0.244940837 | 0.657629036 | 6.576290358 |
| Epha6 | 55.580833 | -0.199342227 | 0.284508276 | -1.151957092 | 0.249338706 | 0.665403896 | 6.654038958 |
| Sh3glb1 | 10.047588 | 0.153191022 | 0.402341629 | 1.141180204 | 0.253794941 | 0.673240471 | 6.732404711 |
| Gnas | 7.3453293 | 0.120225547 | 0.434539855 | 1.133737968 | 0.256904478 | 0.677432642 | 6.774326416 |
| Gm50306 | 44.830368 | -0.192414859 | 0.267872288 | -1.116414813 | 0.264244616 | 0.692664881 | 6.926648807 |
| Gm47371 | 7.2979345 | -0.134203519 | 0.372677529 | -1.077324955 | 0.281335117 | 0.733126217 | 7.331262174 |
| Rpl18a | 11.955841 | 0.138329449 | 0.376851506 | 0.988248133 | 0.32303114 | 0.743492868 | 7.434928677 |
| Rpl37a | 24.871754 | 0.111460402 | 0.445269135 | 1.06273075 | 0.287904072 | 0.743492868 | 7.434928677 |
| Rbfox1 | 15.527408 | -0.117206114 | 0.367511542 | -0.976737898 | 0.328698928 | 0.743492868 | 7.434928677 |
| Pgap1 | 6.7447853 | -0.144269504 | 0.416043183 | 0.980880194 | 0.326651826 | 0.743492868 | 7.434928677 |
| Lingo2 | 10.584524 | 0.1565587 | 0.324118025 | 0.97136639 | 0.331365859 | 0.743492868 | 7.434928677 |
| Rps12 | 16.157665 | 0.156382539 | 0.348738413 | 1.000648861 | 0.316996599 | 0.743492868 | 7.434928677 |
| Myt1l | 6.136326 | 0.112631991 | 0.40111128 | 0.988676711 | 0.322821341 | 0.743492868 | 7.434928677 |
| Slc1a2 | 14.43218 | 0.102697833 | 0.444217062 | 1.032942647 | 0.301630739 | 0.743492868 | 7.434928677 |
| Ankrd12 | 27.62106 | 0.161081827 | 0.31451544 | 0.969479372 | 0.332306067 | 0.743492868 | 7.434928677 |
| Tlr4 | 8.745401 | -0.144269504 | 0.416043178 | -1.044652423 | 0.296183642 | 0.743492868 | 7.434928677 |
| Aldoc | 8.814409 | 0.130414424 | 0.384346182 | 0.989072348 | 0.322627746 | 0.743492868 | 7.434928677 |
| Chchd6 | 10.823148 | -0.137061745 | 0.357629807 | -0.989761932 | 0.322290496 | 0.743492868 | 7.434928677 |
| Cox11 | 8.621069 | -0.135566273 | 0.363401889 | -1.015997488 | 0.309630583 | 0.743492868 | 7.434928677 |
| Dbi | 14.679555 | -0.144269504 | 0.416043181 | 1.040754753 | 0.297989384 | 0.743492868 | 7.434928677 |
| 8430422H( | 65.08901 | -0.144269504 | 0.41604314 | -0.991999053 | 0.321197987 | 0.743492868 | 7.434928677 |
| Cox7c | 14.654309 | 0.179135242 | 0.288622243 | 1.03274814 | 0.301721779 | 0.743492868 | 7.434928677 |
| Cdh12 | 7.2508698 | -0.144269504 | 0.416043182 | -0.995220352 | 0.319629106 | 0.743492868 | 7.434928677 |
| Trim33 | 7.735228 | -0.127870089 | 0.362616726 | -0.980987238 | 0.326599035 | 0.743492868 | 7.434928677 |
| Csmd1 | 17.076962 | 0.139394458 | 0.386779621 | 1.006493001 | 0.314178477 | 0.743492868 | 7.434928677 |
| Alpl | 14.307622 | -0.171207911 | 0.29440498 | -1.002154367 | 0.316269044 | 0.743492868 | 7.434928677 |
| Gabrb1 | 8.30607 | 0.139400352 | 0.388786407 | 1.051695404 | 0.29293932 | 0.743492868 | 7.434928677 |
| Uqcrh | 7.830963 | 0.116518711 | 0.405882144 | 0.999766177 | 0.317423678 | 0.743492868 | 7.434928677 |
| Usp37 | 8.726714 | -0.144269504 | 0.416043174 | -1.044652392 | 0.296183656 | 0.743492868 | 7.434928677 |
| 4932412D2 | 7.161943 | -0.125468473 | 0.368640918 | -1.017106321 | 0.309102854 | 0.743492868 | 7.434928677 |
| Gm14582 | 19.499535 | -0.144269504 | 0.416043173 | -0.993638263 | 0.320399009 | 0.743492868 | 7.434928677 |
| Gm28638 | 7.3468695 | -0.144269504 | 0.416043184 | 0.971428487 | 0.331334949 | 0.743492868 | 7.434928677 |
| Gm49797 | 7.9473557 | -0.144269504 | 0.41604318 | -1.019991637 | 0.307732427 | 0.743492868 | 7.434928677 |
| Spock1 | 19.296722 | -0.162482774 | 0.324846898 | -1.001983334 | 0.316351642 | 0.743492868 | 7.434928677 |
| Gm12216 | 135.57246 | -0.113953519 | 0.133448313 | -0.9651798 | 0.334454763 | 0.744539999 | 7.445399989 |
| Rps21 | 42.87345 | -0.137729638 | 0.34957636 | -0.953510237 | 0.340331612 | 0.75383452 | 7.538345202 |
| Phkg1 | 19.854284 | -0.149862308 | 0.320846549 | -0.936961967 | 0.348778122 | 0.76870004 | 7.687000404 |
| Pip5k1b | 13.284409 | -0.144269504 | 0.416043173 | -0.931756093 | 0.35146259 | 0.77078182 | 7.707818196 |
| Gm20342 | 13.577959 | -0.1323808 | 0.345528966 | -0.917124418 | 0.359077447 | 0.783602507 | 7.836025074 |
| Rpl8 | 8.272719 | 0.095479273 | 0.393205037 | 0.835363047 | 0.403513329 | 0.790647788 | 7.906477878 |
| Zfp236 | 8.841368 | -0.111647313 | 0.350139625 | -0.838166085 | 0.401937432 | 0.790647788 | 7.906477878 |
| Gpc5 | 5.7927704 | 0.097476208 | 0.388558543 | 0.857554187 | 0.391138682 | 0.790647788 | 7.906477878 |
| D030024E( | 7.7510514 | 0.103867516 | 0.364096782 | 0.796499783 | 0.425741602 | 0.790647788 | 7.906477878 |
| Meis3 | 6.8671646 | -0.144269504 | 0.416043181 | -0.807063644 | 0.419629817 | 0.790647788 | 7.906477878 |
| Mrpl15 | 7.4101195 | -0.09860978 | 0.355935368 | -0.804494114 | 0.421111667 | 0.790647788 | 7.906477878 |
| Naa11 | 10.836784 | -0.109109037 | 0.340684564 | -0.782882371 | 0.433696191 | 0.790647788 | 7.906477878 |
| Rps8 | 24.841814 | 0.126767947 | 0.303382667 | 0.783547831 | 0.433305477 | 0.790647788 | 7.906477878 |
| Chl1 | 18.524576 | -0.130271787 | 0.270906097 | -0.786912174 | 0.431333281 | 0.790647788 | 7.906477878 |
| Cep128 | 13.595762 | 0.117449422 | 0.351618672 | 0.815231538 | 0.414939788 | 0.790647788 | 7.906477878 |
| Rpl37 | 30.822403 | 0.135036717 | 0.351642878 | 0.890611196 | 0.37313779 | 0.790647788 | 7.906477878 |

| | | | | | | | |
|---|---|---|---|---|---|---|---|
| Tmeff2 | 15.771518 | -0.144269504 | 0.41604318 | 0.803170605 | 0.421876133 | 0.790647788 | 7.906477878 |
| Cdh23 | 43.743217 | -0.134613029 | 0.204816389 | -0.861079766 | 0.389194112 | 0.790647788 | 7.906477878 |
| Nsf | 5.768386 | -0.144269504 | 0.41604318 | 0.870782054 | 0.383873166 | 0.790647788 | 7.906477878 |
| Rpl23 | 14.544383 | 0.124701309 | 0.353778834 | 0.856269781 | 0.391848572 | 0.790647788 | 7.906477878 |
| Atp2b2 | 8.060129 | -0.060167823 | 0.366064892 | -0.822243741 | 0.410938189 | 0.790647788 | 7.906477878 |
| Pcdh17 | 6.5773635 | 0.098265526 | 0.372252307 | 0.797876403 | 0.424942218 | 0.790647788 | 7.906477878 |
| Wbscr25 | 7.0775514 | -0.144269504 | 0.416043181 | -0.902435968 | 0.366825321 | 0.790647788 | 7.906477878 |
| Pde4d | 5.9735985 | 0.0898054 | 0.382338127 | 0.788107784 | 0.430633662 | 0.790647788 | 7.906477878 |
| Nckap5 | 145.2572 | -0.128584381 | 0.209764437 | -0.82422815 | 0.409809935 | 0.790647788 | 7.906477878 |
| Tmem108 | 28.584448 | -0.144269504 | 0.416043171 | -0.880854176 | 0.378396753 | 0.790647788 | 7.906477878 |
| Rpl11 | 23.5805 | 0.136541664 | 0.343181039 | 0.885866282 | 0.375689587 | 0.790647788 | 7.906477878 |
| Fam189a2 | 8.9994755 | -0.144269504 | 0.416043177 | -0.818644235 | 0.412989421 | 0.790647788 | 7.906477878 |
| Fau | 16.124325 | 0.114796701 | 0.374960446 | 0.854242624 | 0.392970571 | 0.790647788 | 7.906477878 |
| Jakmip2 | 6.640921 | -0.097895878 | 0.358358296 | -0.816859845 | 0.414008528 | 0.790647788 | 7.906477878 |
| Il23a | 9.502472 | -0.107190365 | 0.348273945 | -0.804650557 | 0.421021359 | 0.790647788 | 7.906477878 |
| Fam155a | 9.914968 | 0.110355778 | 0.356601021 | 0.799928574 | 0.423752181 | 0.790647788 | 7.906477878 |
| Frmpd4 | 9.439089 | 0.101816492 | 0.387489951 | 0.846201208 | 0.397440508 | 0.790647788 | 7.906477878 |
| Fyb2 | 30.695557 | -0.144269504 | 0.416043168 | 0.843107716 | 0.399168204 | 0.790647788 | 7.906477878 |
| Kcnip4 | 25.411425 | -0.134608096 | 0.320227769 | -0.8598394 | 0.389877577 | 0.790647788 | 7.906477878 |
| Spag17 | 6.7585807 | -0.096667139 | 0.362414827 | -0.842339794 | 0.399597782 | 0.790647788 | 7.906477878 |
| Sorcs2 | 14.490852 | -0.144269504 | 0.41604317 | -0.852616123 | 0.393872219 | 0.790647788 | 7.906477878 |
| Dlgap1 | 10.49048 | 0.117147408 | 0.35375452 | 0.828593687 | 0.407334362 | 0.790647788 | 7.906477878 |
| Gm44593 | 7.5369215 | -0.107271191 | 0.359232922 | -0.869792068 | 0.384414047 | 0.790647788 | 7.906477878 |
| Rps10 | 15.820669 | 0.12012271 | 0.347336417 | 0.816730569 | 0.414082418 | 0.790647788 | 7.906477878 |
| Lyrm1 | 10.499059 | -0.111548948 | 0.352318098 | -0.846826346 | 0.397091921 | 0.790647788 | 7.906477878 |
| Sptlc3 | 7.487166 | -0.092143811 | 0.358257184 | -0.784690266 | 0.432635192 | 0.790647788 | 7.906477878 |
| Dctd | 11.168691 | -0.128929137 | 0.345234539 | -0.899986511 | 0.368127429 | 0.790647788 | 7.906477878 |
| Gm42889 | 11.820578 | -0.124866374 | 0.34442829 | -0.876702331 | 0.38064834 | 0.790647788 | 7.906477878 |
| Maml3 | 8.084639 | -0.144269504 | 0.416043183 | 0.850261954 | 0.395179464 | 0.790647788 | 7.906477878 |
| Grin2b | 7.271239 | -0.091430111 | 0.354096111 | -0.751686155 | 0.452239818 | 0.800425685 | 8.004256851 |
| Ptpn9 | 23.07427 | -0.118544984 | 0.301038612 | -0.744624909 | 0.456498505 | 0.800425685 | 8.004256851 |
| Foxn3 | 10.898032 | -0.101373404 | 0.344024036 | -0.75343284 | 0.451189855 | 0.800425685 | 8.004256851 |
| Vit | 28.113583 | -0.123467429 | 0.292709562 | -0.762786506 | 0.44559073 | 0.800425685 | 8.004256851 |
| Car12 | 5.946631 | -0.144269504 | 0.416043182 | 0.73689899 | 0.461183779 | 0.800425685 | 8.004256851 |
| Rps23 | 18.716457 | 0.108618431 | 0.343150077 | 0.742218496 | 0.457954964 | 0.800425685 | 8.004256851 |
| Zfp994 | 20.68027 | -0.116177855 | 0.314844609 | -0.753160973 | 0.451353188 | 0.800425685 | 8.004256851 |
| Rps15 | 12.40428 | 0.107307926 | 0.339574047 | 0.737987662 | 0.460521947 | 0.800425685 | 8.004256851 |
| Tenm2 | 10.901395 | -0.078122595 | 0.363641307 | -0.751713219 | 0.452223539 | 0.800425685 | 8.004256851 |
| Ndrg4 | 12.432465 | 0.106508716 | 0.340062245 | 0.734656897 | 0.462548477 | 0.800425685 | 8.004256851 |
| Cldn16 | 15.553372 | -0.114220237 | 0.321428379 | -0.754761873 | 0.450391875 | 0.800425685 | 8.004256851 |
| Pou6f2 | 6.514782 | -0.089079978 | 0.355725367 | -0.749885048 | 0.45332394 | 0.800425685 | 8.004256851 |
| Abat | 7.444665 | -0.144269504 | 0.416043181 | -0.747316523 | 0.454872521 | 0.800425685 | 8.004256851 |
| Srbd1 | 9.897157 | -0.097399743 | 0.33924877 | -0.713161357 | 0.475745916 | 0.807622939 | 8.076229387 |
| 2900097C1 | 6.721527 | 0.088205031 | 0.3666312 | 0.71765346 | 0.472970986 | 0.807622939 | 8.076229387 |
| 4930558K0 | 29.897451 | -0.144269504 | 0.41604317 | 0.71736267 | 0.473150347 | 0.807622939 | 8.076229387 |
| Fbxl7 | 26.032482 | -0.102886213 | 0.328638334 | -0.708538466 | 0.478610934 | 0.807622939 | 8.076229387 |
| Snrpn | 6.7307167 | 0.085955553 | 0.368392775 | 0.70918595 | 0.478209091 | 0.807622939 | 8.076229387 |
| Pkp2 | 15.413167 | -0.10637988 | 0.319200392 | -0.707156627 | 0.479469149 | 0.807622939 | 8.076229387 |
| Lin28b | 11.945574 | -0.101142319 | 0.33215892 | -0.709891304 | 0.477771543 | 0.807622939 | 8.076229387 |
| Lsamp | 39.143772 | -0.105335429 | 0.203571903 | -0.684881343 | 0.4934188 | 0.808264945 | 8.08264945 |
| Fam169a | 40.403946 | -0.144269504 | 0.416043171 | 0.700858202 | 0.483391511 | 0.808264945 | 8.08264945 |
| Dppa2 | 14.264458 | -0.109770114 | 0.293708742 | -0.68721253 | 0.49194881 | 0.808264945 | 8.08264945 |
| Usp3 | 17.644514 | -0.114013055 | 0.271777891 | -0.695416692 | 0.486794203 | 0.808264945 | 8.08264945 |
| Klf10 | 9.963331 | -0.089771225 | 0.343241733 | -0.68458845 | 0.493603658 | 0.808264945 | 8.08264945 |
| Nrgn | 7.1817718 | 0.07635791 | 0.382295465 | 0.68712051 | 0.492006791 | 0.808264945 | 8.08264945 |
| Ralyl | 14.376986 | 0.102005246 | 0.327307331 | 0.68325377 | 0.494446501 | 0.808264945 | 8.08264945 |
| Cmpk1 | 75.07277 | -0.102284947 | 0.188349781 | -0.685540984 | 0.493002608 | 0.808264945 | 8.08264945 |
| AI314278 | 14.637004 | -0.096796614 | 0.32419072 | -0.663388 | 0.50708209 | 0.816863148 | 8.168631479 |
| Mrc2 | 20.690413 | -0.144269504 | 0.41604317 | -0.664562087 | 0.506330625 | 0.816863148 | 8.168631479 |
| Ndst3 | 6.0131106 | -0.144269504 | 0.416043178 | -0.668420166 | 0.503865426 | 0.816863148 | 8.168631479 |
| Gm26901 | 6.7513933 | 0.079183199 | 0.369551038 | 0.667514636 | 0.504443464 | 0.816863148 | 8.168631479 |

| | | | | | | | |
|---|---|---|---|---|---|---|---|
| Spdye4a | 6.411588 | -0.069572801 | 0.36049146 | -0.659709454 | 0.509440298 | 0.817688595 | 8.17688595 |
| Ctnna2 | 21.734634 | -0.098545982 | 0.314452081 | -0.653071993 | 0.513709867 | 0.821564877 | 8.215648773 |
| Pcdh15 | 39.479504 | -0.106171769 | 0.264501975 | -0.650057275 | 0.515655226 | 0.821709587 | 8.21709587 |
| Iqcb1 | 9.742833 | -0.086314636 | 0.338302213 | -0.644770235 | 0.519076093 | 0.824196091 | 8.24196091 |
| Wdyhv1 | 28.000046 | 0.096848608 | 0.262730993 | 0.595602634 | 0.551440716 | 0.824476286 | 8.244762861 |
| Rtkn | 8.03187 | 0.077039322 | 0.356865258 | 0.608815026 | 0.542647053 | 0.824476286 | 8.244762861 |
| Rubcnl | 20.44625 | -0.144269504 | 0.416043177 | -0.604095783 | 0.545779963 | 0.824476286 | 8.244762861 |
| Sil1 | 40.758835 | -0.086042304 | 0.256548312 | -0.533485753 | 0.593697371 | 0.824476286 | 8.244762861 |
| Rps16 | 15.809079 | -0.081444803 | 0.32733636 | -0.577415732 | 0.563658649 | 0.824476286 | 8.244762861 |
| Stard13 | 10.583754 | 0.076875893 | 0.336373726 | 0.554085241 | 0.57952051 | 0.824476286 | 8.244762861 |
| Zfhx3 | 22.321194 | -0.087271384 | 0.297368238 | -0.562059372 | 0.574075568 | 0.824476286 | 8.244762861 |
| Sult2a2 | 13.528371 | -0.076408923 | 0.321299289 | -0.534383406 | 0.593076297 | 0.824476286 | 8.244762861 |
| Tcea1 | 14.393718 | -0.144269504 | 0.416043179 | -0.534389522 | 0.593072067 | 0.824476286 | 8.244762861 |
| Rpl21 | 22.089443 | 0.067273219 | 0.394616962 | 0.623170386 | 0.53317256 | 0.824476286 | 8.244762861 |
| Rpl13a | 12.340456 | 0.068117424 | 0.37157641 | 0.573576384 | 0.566254496 | 0.824476286 | 8.244762861 |
| Xirp2 | 12.11597 | -0.144269504 | 0.416043178 | 0.583878731 | 0.559301904 | 0.824476286 | 8.244762861 |
| Rabgap1l | 14.293994 | 0.085480392 | 0.344005138 | 0.615515321 | 0.538214454 | 0.824476286 | 8.244762861 |
| 1700030K0 | 9.7303295 | -0.068603632 | 0.34483234 | -0.553229172 | 0.580106494 | 0.824476286 | 8.244762861 |
| Gm38973 | 22.971754 | 0.086819862 | 0.297121052 | 0.552297984 | 0.580744212 | 0.824476286 | 8.244762861 |
| Gria1 | 11.842802 | -0.144269504 | 0.416043175 | -0.597871344 | 0.549925782 | 0.824476286 | 8.244762861 |
| Atp2b4 | 7.9582925 | -0.08889684 | 0.311599036 | -0.593532683 | 0.552824717 | 0.824476286 | 8.244762861 |
| Mar-03 | 7.669989 | 0.069085326 | 0.352071979 | 0.546382773 | 0.584802849 | 0.824476286 | 8.244762861 |
| Abcg2 | 6.1671276 | -0.065467383 | 0.352607633 | -0.574705711 | 0.565490344 | 0.824476286 | 8.244762861 |
| Nell1 | 37.07933 | -0.092274764 | 0.261065503 | -0.571648198 | 0.567560335 | 0.824476286 | 8.244762861 |
| Dtd1 | 6.049204 | -0.144269504 | 0.416043183 | 0.607104173 | 0.543781781 | 0.824476286 | 8.244762861 |
| Dpp10 | 7.456398 | -0.03199916 | 0.367423893 | -0.548404983 | 0.583413856 | 0.824476286 | 8.244762861 |
| A930016O | 9.818968 | -0.144269504 | 0.416043179 | -0.552704177 | 0.580465993 | 0.824476286 | 8.244762861 |
| Nrp2 | 10.334757 | -0.082684736 | 0.334064011 | -0.607322049 | 0.543637209 | 0.824476286 | 8.244762861 |
| Kcnq2 | 6.291892 | -0.061651921 | 0.358069807 | -0.581767729 | 0.560723143 | 0.824476286 | 8.244762861 |
| Dcc | 10.508539 | -0.144269504 | 0.416043173 | -0.595815058 | 0.551298783 | 0.824476286 | 8.244762861 |
| Alk | 6.1867847 | -0.06777635 | 0.352687812 | -0.591080415 | 0.554466542 | 0.824476286 | 8.244762861 |
| Cntn3 | 17.998539 | -0.144269504 | 0.41604318 | 0.602733313 | 0.546686116 | 0.824476286 | 8.244762861 |
| Ahcyl2 | 9.446116 | 0.07110845 | 0.345841237 | 0.538604022 | 0.590160112 | 0.824476286 | 8.244762861 |
| Ckb | 20.8185 | 0.065809756 | 0.380271392 | 0.569411005 | 0.56907725 | 0.824476286 | 8.244762861 |
| Ptprd | 21.2857 | -0.082812236 | 0.317916206 | -0.564945713 | 0.57211069 | 0.824476286 | 8.244762861 |
| Atp5md | 12.651631 | 0.083154325 | 0.337030381 | 0.589640049 | 0.555431995 | 0.824476286 | 8.244762861 |
| Calb1 | 6.1530104 | -0.069525902 | 0.35108644 | -0.595369317 | 0.55159663 | 0.824476286 | 8.244762861 |
| 4931429P1 | 5.7152615 | 0.071926666 | 0.370231623 | 0.630446179 | 0.528402705 | 0.824476286 | 8.244762861 |
| Camk2a | 26.517548 | -0.144269504 | 0.416043176 | 0.608014001 | 0.543178188 | 0.824476286 | 8.244762861 |
| 4930441J1 | 22.884426 | -0.144269504 | 0.416043169 | -0.562746159 | 0.573607748 | 0.824476286 | 8.244762861 |
| Prkag2 | 25.674173 | -0.144269504 | 0.416043173 | -0.582066315 | 0.560522013 | 0.824476286 | 8.244762861 |
| Cenpi | 13.821451 | -0.144269504 | 0.416043175 | -0.567124501 | 0.570629598 | 0.824476286 | 8.244762861 |
| Chsy3 | 32.73518 | -0.086619407 | 0.266887002 | -0.537194483 | 0.591133282 | 0.824476286 | 8.244762861 |
| Kcnq5 | 14.210515 | -0.086928726 | 0.337018213 | -0.640668307 | 0.521738209 | 0.824476286 | 8.244762861 |
| Dlg2 | 11.623713 | -0.06763069 | 0.336959913 | -0.517024973 | 0.605138726 | 0.825638088 | 8.256380877 |
| Sntg1 | 6.711516 | -0.061334212 | 0.348394174 | -0.521209938 | 0.602220532 | 0.825638088 | 8.256380877 |
| Rpl38 | 24.64436 | 0.079933461 | 0.307023854 | 0.518945559 | 0.603798706 | 0.825638088 | 8.256380877 |
| Ephb2 | 6.759665 | -0.059876597 | 0.351321439 | -0.522751956 | 0.601146877 | 0.825638088 | 8.256380877 |
| Sema3c | 6.1041503 | -0.055007234 | 0.356310504 | -0.515763854 | 0.606019351 | 0.825638088 | 8.256380877 |
| Pde1c | 75.51991 | -0.069900204 | 0.165041833 | -0.508207747 | 0.611307659 | 0.825638088 | 8.256380877 |
| D430042O | 78.718346 | -0.071611572 | 0.1705843 | -0.510505077 | 0.609697659 | 0.825638088 | 8.256380877 |
| Pcgf6 | 17.646658 | -0.144269504 | 0.416043175 | -0.523099102 | 0.60090529 | 0.825638088 | 8.256380877 |
| Hipk2 | 6.769567 | -0.058138548 | 0.35181889 | -0.512319227 | 0.608427611 | 0.825638088 | 8.256380877 |
| Rps13 | 13.523786 | 0.070945092 | 0.331336748 | 0.501894676 | 0.61574161 | 0.826586465 | 8.265864646 |
| Tnni3k | 7.789238 | -0.144269504 | 0.416043178 | -0.503776114 | 0.614418714 | 0.826586465 | 8.265864646 |
| Tshz2 | 8.23189 | -0.05647825 | 0.350805745 | -0.492798777 | 0.622154775 | 0.827479441 | 8.274794406 |
| Gpm6a | 9.308586 | -0.064049776 | 0.33561737 | -0.490361091 | 0.623878404 | 0.827479441 | 8.274794406 |
| Rab29 | 24.73881 | -0.144269504 | 0.416043165 | -0.493649755 | 0.621553555 | 0.827479441 | 8.274794406 |
| Macrod1 | 9.057329 | 0.067960909 | 0.332453915 | 0.493732768 | 0.62149492 | 0.827479441 | 8.274794406 |
| Ssh2 | 14.778119 | 0.067381078 | 0.334360615 | 0.482748429 | 0.629274372 | 0.832144916 | 8.321449163 |
| Hdac9 | 12.908575 | -0.064917118 | 0.329941957 | -0.479823868 | 0.63135264 | 0.832408391 | 8.324083912 |

| | | | | | | | |
|---|---|---|---|---|---|---|---|
| Dmd | 13.929831 | 0.063101508 | 0.342448679 | 0.469191487 | 0.63893277 | 0.832491815 | 8.324918146 |
| Nrg1 | 7.284752 | -0.057575072 | 0.34489211 | -0.477079 | 0.633305868 | 0.832491815 | 8.324918146 |
| Arhgap18 | 5.685643 | -0.144269504 | 0.416043184 | 0.470657725 | 0.637885178 | 0.832491815 | 8.324918146 |
| Cadm2 | 12.924879 | 0.059385478 | 0.355635416 | 0.472452542 | 0.636603808 | 0.832491815 | 8.324918146 |
| Rpl36 | 11.369721 | 0.063647865 | 0.333902765 | 0.46139849 | 0.644512735 | 0.837299535 | 8.372995353 |
| Mt1 | 24.129139 | 0.051539675 | 0.37696619 | 0.455355862 | 0.648853243 | 0.838023285 | 8.380232853 |
| Adk | 21.201334 | -0.070659509 | 0.292217797 | -0.457124422 | 0.647581618 | 0.838023285 | 8.380232853 |
| Ppfia3 | 6.9764113 | -0.144269504 | 0.416043181 | -0.452532742 | 0.650885239 | 0.838203956 | 8.38203956 |
| Pcdh9 | 11.418508 | 0.063747161 | 0.324049756 | 0.447926981 | 0.654205895 | 0.839074519 | 8.390745185 |
| Vmn2r103 | 9.938954 | -0.144269504 | 0.416043176 | 0.446343141 | 0.655349398 | 0.839074519 | 8.390745185 |
| Gm11337 | 20.5174 | -0.066523552 | 0.295848955 | -0.434996659 | 0.663564853 | 0.845878731 | 8.458787314 |
| Selenow | 11.197987 | 0.055339314 | 0.349995576 | 0.433732621 | 0.664482615 | 0.845878731 | 8.458787314 |
| Susd4 | 6.0400133 | 0.048376459 | 0.360559442 | 0.423793977 | 0.671716071 | 0.852636732 | 8.526367315 |
| Sh3bgrl2 | 7.145208 | -0.0481054 | 0.345172416 | -0.405556591 | 0.68506843 | 0.852752862 | 8.527528624 |
| mt-Atp8 | 6.825624 | 0.054620175 | 0.333358359 | 0.410722077 | 0.681276337 | 0.852752862 | 8.527528624 |
| Gm37982 | 8.135304 | -0.144269504 | 0.416043178 | 0.405049837 | 0.685440878 | 0.852752862 | 8.527528624 |
| Celf2 | 16.3416 | -0.049032859 | 0.346599109 | -0.407279712 | 0.683802562 | 0.852752862 | 8.527528624 |
| Senp2 | 8.941299 | -0.144269504 | 0.416043177 | -0.416082056 | 0.67734996 | 0.852752862 | 8.527528624 |
| Kcnd2 | 10.868125 | 0.054295797 | 0.336637246 | 0.403886135 | 0.686296453 | 0.852752862 | 8.527528624 |
| Gm49767 | 15.366516 | -0.144269504 | 0.416043174 | -0.402648045 | 0.68720716 | 0.852752862 | 8.527528624 |
| Asxl3 | 15.20223 | -0.05986707 | 0.311959967 | -0.41264809 | 0.679864462 | 0.852752862 | 8.527528624 |
| Nrxn1 | 13.538358 | -0.045479265 | 0.351701677 | -0.398531188 | 0.690238673 | 0.854122157 | 8.541221574 |
| Vmn1r13 | 6.4503555 | -0.043666444 | 0.348111059 | -0.381005267 | 0.703199341 | 0.860832685 | 8.608326848 |
| Calm2 | 8.584985 | 0.048343613 | 0.345021378 | 0.381330659 | 0.702957906 | 0.860832685 | 8.608326848 |
| Ehbp1 | 54.4034 | -0.06066458 | 0.254556781 | -0.380985194 | 0.703214235 | 0.860832685 | 8.608326848 |
| Comt | 23.47635 | -0.058995439 | 0.285387798 | -0.380688526 | 0.703434383 | 0.860832685 | 8.608326848 |
| Samd12 | 8.580058 | 0.046500966 | 0.34785203 | 0.372983861 | 0.709160456 | 0.863071654 | 8.630716539 |
| Rpl32 | 19.181837 | 0.055519828 | 0.312801216 | 0.374606571 | 0.707953085 | 0.863071654 | 8.630716539 |
| Grik2 | 6.7997155 | 0.043834385 | 0.351874983 | 0.36631539 | 0.714129747 | 0.866738296 | 8.66738296 |
| Dclk1 | 6.2317123 | 0.042227491 | 0.352955376 | 0.358556027 | 0.719927248 | 0.871387352 | 8.713873517 |
| Gm13219 | 5.8314877 | -0.144269504 | 0.41604318 | 0.342329637 | 0.732102835 | 0.871832156 | 8.718321556 |
| 2310002F0 | 9.686885 | -0.044730483 | 0.331694557 | -0.343090026 | 0.731530734 | 0.871832156 | 8.718321556 |
| Cadps2 | 6.855507 | -0.048624975 | 0.311672123 | -0.347528205 | 0.728194533 | 0.871832156 | 8.718321556 |
| Agbl4 | 6.2274485 | 0.040507992 | 0.35409504 | 0.346884334 | 0.728678217 | 0.871832156 | 8.718321556 |
| Adgrg6 | 5.8655686 | -0.144269504 | 0.416043182 | 0.342329618 | 0.732102849 | 0.871832156 | 8.718321556 |
| Gm2895 | 9.839932 | -0.041927054 | 0.341357299 | -0.343729155 | 0.731049983 | 0.871832156 | 8.718321556 |
| Gm42791 | 6.778445 | 0.03484935 | 0.359733885 | 0.308463536 | 0.757729643 | 0.899930916 | 8.999309161 |
| Klhl14 | 6.4297285 | -0.03012411 | 0.358369053 | -0.297265493 | 0.766263824 | 0.907633353 | 9.076333531 |
| Wipf1 | 7.5803995 | 0.035832513 | 0.343089682 | 0.286488429 | 0.774504055 | 0.914947457 | 9.149474565 |
| Ccdc60 | 20.410414 | -0.043034229 | 0.270375198 | -0.273689257 | 0.784323434 | 0.919193866 | 9.191938661 |
| Gnaz | 65.72708 | -0.04401145 | 0.231587199 | -0.278665006 | 0.78050192 | 0.919193866 | 9.191938661 |
| Rpl35a | 18.569323 | 0.04088349 | 0.305179331 | 0.273807293 | 0.784232719 | 0.919193866 | 9.191938661 |
| Prnp | 6.7899947 | -0.029083139 | 0.349782589 | -0.259083178 | 0.795571064 | 0.927468372 | 9.274683717 |
| Ogfrl1 | 7.2132864 | -0.144269504 | 0.416043184 | 0.261289044 | 0.793869615 | 0.927468372 | 9.274683717 |
| Pecam1 | 8.383729 | 0.035009994 | 0.32199145 | 0.254659594 | 0.798986027 | 0.92900475 | 9.2900475 |
| Brip1os | 6.327769 | -0.024788491 | 0.360436795 | -0.250044215 | 0.802553156 | 0.930054172 | 9.300541716 |
| Crhr2 | 6.000898 | -0.144269504 | 0.416043184 | 0.218091779 | 0.827357603 | 0.930054172 | 9.300541716 |
| Olfr729 | 8.73949 | -0.027410928 | 0.341169039 | -0.226606054 | 0.820730084 | 0.930054172 | 9.300541716 |
| Kirrel3 | 5.8289046 | 0.02704133 | 0.353483127 | 0.237041065 | 0.812624933 | 0.930054172 | 9.300541716 |
| Atg13 | 6.0980806 | -0.024395993 | 0.347405278 | -0.21562405 | 0.829280808 | 0.930054172 | 9.300541716 |
| Rps17 | 8.93894 | 0.028661222 | 0.336737721 | 0.221275469 | 0.82487795 | 0.930054172 | 9.300541716 |
| Dtnb | 9.041158 | -0.144269504 | 0.416043178 | 0.217454767 | 0.827853954 | 0.930054172 | 9.300541716 |
| Ptprg | 13.163048 | 0.034793222 | 0.315676945 | 0.242249438 | 0.80858689 | 0.930054172 | 9.300541716 |
| Gm14507 | 6.5407157 | -0.144269504 | 0.416043184 | 0.24041087 | 0.810011752 | 0.930054172 | 9.300541716 |
| Sgcz | 7.5641694 | -0.025645593 | 0.356279495 | -0.243139649 | 0.807897217 | 0.930054172 | 9.300541716 |
| Serinc1 | 7.0369806 | 0.02496619 | 0.357111678 | 0.220930856 | 0.825146272 | 0.930054172 | 9.300541716 |
| Il27 | 48.842026 | -0.031928475 | 0.176766846 | -0.223457219 | 0.823179676 | 0.930054172 | 9.300541716 |
| Lrrc4c | 18.952847 | -0.032161995 | 0.300155248 | -0.217232152 | 0.828027429 | 0.930054172 | 9.300541716 |
| Gmcl1 | 9.497752 | 0.031644037 | 0.332577916 | 0.238850488 | 0.811221519 | 0.930054172 | 9.300541716 |
| Klkb1 | 18.450718 | -0.032154842 | 0.287866122 | -0.211397183 | 0.832577355 | 0.931393354 | 9.313933539 |
| Plekhg1 | 6.0983872 | -0.022005956 | 0.354638265 | -0.207099056 | 0.835932499 | 0.932791177 | 9.327911768 |

| | | | | | | | |
|---|---|---|---|---|---|---|---|
| Olfr57 | 9.326843 | -0.025484168 | 0.33568491 | -0.203318042 | 0.838886454 | 0.933735425 | 9.337354254 |
| Dop1b | 36.803204 | 0.031841378 | 0.26592855 | 0.199829912 | 0.841613607 | 0.934423127 | 9.344231272 |
| Disc1 | 7.1660576 | 0.023193824 | 0.345112733 | 0.191246 | 0.848332866 | 0.939528649 | 9.395286491 |
| A230057D( | 6.6212473 | -0.019533942 | 0.352760516 | -0.179288511 | 0.857711167 | 0.940141493 | 9.401414928 |
| Arvcf | 5.8929763 | -0.144269504 | 0.41604318 | -0.172074251 | 0.863379156 | 0.940141493 | 9.401414928 |
| Rap1gap2 | 5.939493 | -0.144269504 | 0.416043183 | -0.172074261 | 0.863379148 | 0.940141493 | 9.401414928 |
| Tshr | 10.289192 | 0.023961418 | 0.33862896 | 0.186708468 | 0.851889216 | 0.940141493 | 9.401414928 |
| Rps27 | 21.824091 | 0.025575692 | 0.320853638 | 0.17966566 | 0.857415054 | 0.940141493 | 9.401414928 |
| Meg3 | 31.199203 | 0.021932651 | 0.357075896 | 0.181469089 | 0.855999394 | 0.940141493 | 9.401414928 |
| Pde6a | 84.62703 | -0.027428977 | 0.231728973 | -0.171613069 | 0.863741733 | 0.940141493 | 9.401414928 |
| Gnao1 | 12.896826 | -0.144269504 | 0.416043182 | 0.158004278 | 0.874453428 | 0.9462182 | 9.462181996 |
| Gm38304 | 5.967835 | 0.01690225 | 0.359361256 | 0.156381372 | 0.87573242 | 0.9462182 | 9.462181996 |
| Acoxl | 32.851875 | -0.025310465 | 0.2594673 | -0.160869002 | 0.872196577 | 0.9462182 | 9.462181996 |
| Ttc28 | 40.03808 | -0.023093215 | 0.259621712 | -0.147360138 | 0.882847767 | 0.951585306 | 9.515853061 |
| Gria3 | 7.159691 | -0.01531355 | 0.349228258 | -0.136047989 | 0.891783344 | 0.958883547 | 9.588835468 |
| Wdr72 | 38.662872 | 0.020080024 | 0.231762569 | 0.12583492 | 0.899862599 | 0.965227921 | 9.65227921 |
| Rps3 | 12.916127 | 0.016944431 | 0.318874591 | 0.120197843 | 0.904326427 | 0.965320045 | 9.653200451 |
| Grid2 | 16.681768 | -0.144269504 | 0.416043176 | 0.117472493 | 0.906485641 | 0.965320045 | 9.653200451 |
| Creb5 | 9.735547 | -0.015836865 | 0.327974674 | -0.121675223 | 0.903156236 | 0.965320045 | 9.653200451 |
| Magi2 | 11.623744 | -0.014312068 | 0.323717862 | -0.107292404 | 0.91455701 | 0.97157975 | 9.715797495 |
| Cdh8 | 5.867067 | -0.011085779 | 0.353560756 | -0.103799299 | 0.917328623 | 0.972192775 | 9.721927748 |
| Nup210l | 8.683186 | -0.144269504 | 0.416043179 | 0.095654032 | 0.92379535 | 0.974384143 | 9.743841434 |
| Bckdha | 6.7999625 | 0.012231362 | 0.337365854 | 0.097712103 | 0.922160905 | 0.974384143 | 9.743841434 |
| Esf1 | 14.626984 | -0.011520434 | 0.307002511 | -0.080410934 | 0.935910431 | 0.984817865 | 9.848178646 |
| Bicc1 | 7.085659 | -0.009027029 | 0.3418231 | -0.076802771 | 0.938780446 | 0.985497009 | 9.854970087 |
| 4930402H2 | 14.7000065 | -0.008684289 | 0.311589552 | -0.06286583 | 0.949873345 | 0.988749263 | 9.887492629 |
| Reps2 | 9.043988 | -0.00738477 | 0.336965074 | -0.06032608 | 0.951895931 | 0.988749263 | 9.887492629 |
| Pard3b | 10.008259 | -0.144269504 | 0.416043179 | 0.049259523 | 0.960712477 | 0.988749263 | 9.887492629 |
| Rpl34 | 16.785885 | 0.007207218 | 0.322421739 | 0.050086752 | 0.960053257 | 0.988749263 | 9.887492629 |
| Malat1 | 85.781746 | 0.008657242 | 0.308292744 | 0.054308266 | 0.956689564 | 0.988749263 | 9.887492629 |
| Zmat4 | 35.043602 | -0.00889706 | 0.2643593 | -0.057617 | 0.954053708 | 0.988749263 | 9.887492629 |
| Slc39a2 | 7.616286 | -0.008197352 | 0.311967117 | -0.059120652 | 0.952856009 | 0.988749263 | 9.887492629 |
| Ahdc1 | 15.425038 | -0.006371823 | 0.318848409 | -0.047686633 | 0.961965987 | 0.988749263 | 9.887492629 |
| Svil | 19.824146 | -0.00800274 | 0.298486563 | -0.055098838 | 0.956059722 | 0.988749263 | 9.887492629 |
| Gm765 | 18.418118 | -0.00616744 | 0.263871908 | -0.041114795 | 0.96720438 | 0.989541664 | 9.895416636 |
| Fgf12 | 15.401117 | -0.005449157 | 0.322136425 | -0.041511522 | 0.966888107 | 0.989541664 | 9.895416636 |
| Rps15a | 9.871597 | 0.003730322 | 0.347764848 | 0.029661971 | 0.976336641 | 0.991345724 | 9.913457236 |
| Susd1 | 8.2491865 | -0.144269504 | 0.416043178 | -0.031673513 | 0.974732418 | 0.991345724 | 9.913457236 |
| C330002G( | 9.527081 | -0.004194983 | 0.334143319 | -0.034439893 | 0.972526373 | 0.991345724 | 9.913457236 |
| Srgn | 9.471736 | 0.003682524 | 0.337178069 | 0.027678064 | 0.977918919 | 0.991345724 | 9.913457236 |
| Tns3 | 19.613354 | -0.003330389 | 0.304006004 | -0.023250807 | 0.981450212 | 0.992653981 | 9.926539812 |
| Ralgps1 | 8.795305 | 0.00188892 | 0.34521535 | 0.013628594 | 0.989126292 | 0.998138832 | 9.981388321 |
| Zeb1 | 8.858541 | -0.144269504 | 0.416043178 | -0.002957569 | 0.997640205 | 0.99859102 | 9.985910201 |
| Mast4 | 6.266274 | 0.000779344 | 0.356274772 | 0.005610036 | 0.995523863 | 0.99859102 | 9.985910201 |
| Tacc1 | 13.222597 | -1.67E-05 | 0.316878405 | -0.001765895 | 0.99859102 | 0.99859102 | 9.985910201 |
| Asic2 | 18.165735 | 0.001027956 | 0.298649087 | 0.00527283 | 0.99579291 | 0.99859102 | 9.985910201 |
| 1700047F( | 4.6522107 | 0.024684648 | 0.360067984 | 0.233137996 | 0.815654251 | | |
| 1700102P( | 4.542301 | -0.007708385 | 0.360887558 | -0.079917181 | 0.936303125 | | |
| 4930549G2 | 3.049757 | 0.00405692 | 0.370279723 | 0.045380607 | 0.963803938 | | |
| A630012P( | 0.75610065 | -0.008436678 | 0.379091004 | -0.181347889 | 0.856094519 | | |
| Abcb4 | 4.661708 | 0.02459957 | 0.359551704 | 0.231456736 | 0.816959992 | | |
| Abhd6 | 4.056553 | 0.067160393 | 0.419106605 | 0.799099749 | 0.424232575 | | |
| Adamtsl1 | 3.7572446 | -0.035728905 | 0.361848144 | -0.384168419 | 0.700853624 | | |
| Akap10 | 5.0634646 | 0.246885969 | 0.511239891 | 1.784098378 | 0.074407681 | | |
| Astn2 | 4.9827747 | -0.144269504 | 0.416043186 | 1.029436651 | 0.303274534 | | |
| Atp8a2 | 3.6725814 | -0.07645732 | 0.366491309 | -0.796602381 | 0.425681994 | | |
| Bckdhb | 4.332469 | -0.032967168 | 0.361731106 | -0.345897262 | 0.729419926 | | |
| Bmpr1b | 2.8447673 | 0.065961703 | 0.393328708 | 0.737742647 | 0.460670851 | | |
| Bmt2 | 3.6812673 | 0.094394547 | 0.399904292 | 0.939840716 | 0.347299271 | | |
| C130073E2 | 4.8236737 | -0.144269504 | 0.416043183 | -0.812593878 | 0.416450943 | | |
| Cacnb4 | 4.398444 | -0.000974723 | 0.359640727 | -0.011047857 | 0.991185265 | | |

| | | | | | |
|---|---|---|---|---|---|
| Cdc14a | 3.9893844 | -0.014378757 | 0.359904288 | -0.148625849 | 0.881848875 |
| Cdh13 | 4.852521 | -0.06472512 | 0.359597072 | -0.622279436 | 0.533758141 |
| Cdh20 | 1.0013392 | -0.144269504 | 0.416043187 | 0.460377975 | 0.645244941 |
| Cmc1 | 5.4820776 | -0.144269504 | 0.416043179 | -2.022948745 | 0.043078432 |
| Col25a1 | 4.212558 | 0.084399843 | 0.396732153 | 0.83793072 | 0.402069614 |
| Dach1 | 1.1488812 | -0.072572613 | 0.359380772 | -1.11026402 | 0.266885273 |
| Dnah1 | 4.584845 | 0.058577097 | 0.382237674 | 0.583237734 | 0.559733273 |
| Eif2b3 | 4.970553 | -0.15641656 | 0.398311578 | -1.410052101 | 0.158524299 |
| Fam53a | 3.9098713 | 0.033187216 | 0.368558913 | 0.335116056 | 0.737537541 |
| Fgf13 | 5.3616858 | -0.016944038 | 0.356060322 | -0.163770074 | 0.869912151 |
| Fgfr2 | 5.105867 | 0.025750985 | 0.372412172 | 0.262216166 | 0.793154791 |
| Fmnl2 | 4.31436 | 0.046379 | 0.369689174 | 0.449427319 | 0.653123428 |
| Fsip1 | 5.1837482 | -0.008315394 | 0.357502158 | -0.081854815 | 0.934762167 |
| Galnt17 | 4.9463263 | -0.028630673 | 0.356730264 | -0.280977291 | 0.778727818 |
| Galntl6 | 4.9721513 | 0.043380389 | 0.371141671 | 0.421249494 | 0.673572906 |
| Gm11775 | 3.8012867 | -0.003042818 | 0.362606215 | -0.033139031 | 0.973563718 |
| Gm12381 | 4.6018014 | -0.046199573 | 0.361092461 | -0.477800798 | 0.632791993 |
| Gm32647 | 3.8484664 | -0.003660316 | 0.361520625 | -0.039141389 | 0.968777663 |
| Gm49171 | 2.9792106 | -0.026711932 | 0.36996989 | -0.342451371 | 0.732011234 |
| Grik1 | 5.2223997 | -0.008397428 | 0.369668743 | -0.094907488 | 0.924388309 |
| Hecw1 | 4.9448857 | -0.144269504 | 0.416043182 | -0.922494692 | 0.356270598 |
| Klhl32 | 3.5486538 | 0.015240098 | 0.375109298 | 0.173288253 | 0.862424856 |
| Larp4 | 4.4198475 | -0.144269504 | 0.416043182 | -1.013221628 | 0.310954312 |
| Lrmda | 4.5780797 | -0.144269504 | 0.41604318 | -1.121126966 | 0.262233822 |
| Lrrc51 | 5.1778035 | -0.007638153 | 0.35409248 | -0.073019951 | 0.941790241 |
| Lrrc58 | 3.8690581 | 0.071480163 | 0.395407657 | 0.753071482 | 0.45140696 |
| Ly75 | 5.1721644 | -0.129719783 | 0.380252018 | -1.163004829 | 0.244827543 |
| Meis2 | 5.0457025 | -0.026810166 | 0.366909445 | -0.305163718 | 0.760241464 |
| Mia2 | 4.81705 | -0.144269504 | 0.416043181 | 0.167835986 | 0.866712311 |
| Micu1 | 5.5047164 | -0.06334478 | 0.359338677 | -0.604600173 | 0.545444691 |
| Mir99ahg | 4.6764874 | 0.022331994 | 0.363341527 | 0.216836537 | 0.828335737 |
| Myo1d | 5.327755 | -0.038244784 | 0.364321608 | -0.415936524 | 0.677456452 |
| Ncam2 | 5.199977 | -0.050283831 | 0.355019236 | -0.469708371 | 0.638563386 |
| Nqo2 | 4.649644 | -0.05586777 | 0.357318368 | -0.534939487 | 0.592691702 |
| Olfr694 | 4.5218077 | 0.033185625 | 0.365676914 | 0.321802936 | 0.74760199 |
| Olfr905 | 5.253632 | 0.027028705 | 0.356614636 | 0.245058123 | 0.806411427 |
| Palld | 3.1472957 | -0.144269504 | 0.416043182 | -0.617246352 | 0.537072247 |
| Ppfia2 | 4.588239 | -0.089191161 | 0.367167984 | -0.868541235 | 0.385098109 |
| Ptger3 | 0.41824993 | 0.013280082 | 0.430706106 | 0.364455355 | 0.715518006 |
| Rgs6 | 0.58242667 | 0.003706581 | 0.393134112 | 0.090394319 | 0.927973871 |
| Rnf180 | 2.8745008 | 0.15595803 | 0.48387728 | 1.653510868 | 0.098226936 |
| Robo2 | 5.292184 | -0.144269504 | 0.416043184 | 2.26262169 | 0.02365902 |
| Sgip1 | 5.421542 | -0.053231407 | 0.359524129 | -0.523535595 | 0.600601587 |
| Skap1 | 5.1535754 | -0.005876191 | 0.354373137 | -0.056704916 | 0.954780258 |
| Slc31a2 | 5.344804 | -0.098624221 | 0.336355567 | -0.723488355 | 0.469379906 |
| Smim13 | 4.3185587 | 0.160412548 | 0.421524611 | 1.321524616 | 0.1863265 |
| Snhg14 | 5.0788894 | -0.039722704 | 0.368456732 | -0.524199853 | 0.600139543 |
| Spata16 | 4.664937 | -0.144269504 | 0.416043184 | 1.339364743 | 0.180451961 |
| St18 | 1.5516726 | 0.014013575 | 0.40339601 | 0.250269315 | 0.802379085 |
| Strbp | 4.8617706 | -0.028108229 | 0.357957991 | -0.279024261 | 0.780226206 |
| Stx8 | 2.8512423 | 0.167893375 | 0.486818335 | 1.704770335 | 0.088237266 |
| Syt11 | 5.28325 | -0.144269504 | 0.416043184 | 0.740483949 | 0.459006397 |
| Vwc2l | 1.9856879 | -0.047395345 | 0.368418567 | -0.672547095 | 0.501235469 |
| Wdr86 | 3.9860065 | 0.068451919 | 0.38283633 | 0.677234267 | 0.498257329 |
| Zfpm2 | 2.991291 | -0.028965058 | 0.365644938 | -0.341526389 | 0.732707343 |

**table S7: Differential gene expression list condensed milk compared to rest (3) conditions in GABA and Glut neurons**

| GABA | baseMean | log2FoldChange | lfcSE | stat | pvalue | padj | pglobadj |
|---|---|---|---|---|---|---|---|
| Grik1 | 283.93326 | 0.415954792 | 0.110693097 | 4.080496983 | 4.49E-05 | 0.010201269 | 0.102012694 |
| Rps15 | 4.6742244 | 2.004885756 | 0.694771651 | 3.657457374 | 0.00025473 | 0.02891181 | 0.289118102 |
| Rps12 | 3.9755926 | 2.300482299 | 0.991505885 | 3.181869285 | 0.001463278 | 0.110721402 | 1.10721402 |
| Fth1 | 10.972276 | 1.579365326 | 0.734105411 | 3.092064937 | 0.001987694 | 0.112801613 | 1.12801613 |
| Mrc2 | 6.650044 | -0.144269504 | 0.32517569 | 2.858463605 | 0.004256979 | 0.184876928 | 1.848769276 |
| Rps10 | 1.7781105 | 1.372568107 | 0.96980549 | 2.814414674 | 0.004886615 | 0.184876928 | 1.848769276 |
| Rpl41 | 4.4877243 | 1.119794213 | 0.812406007 | 2.733567829 | 0.006265222 | 0.203172191 | 2.031721909 |
| Rps28 | 4.243435 | 0.550309235 | 0.892796637 | 2.486617568 | 0.012896395 | 0.252703112 | 2.527031124 |
| Chsy3 | 7.38024 | 0.573556314 | 0.515636527 | 2.465223189 | 0.013692793 | 0.252703112 | 2.527031124 |
| Atp9b | 6.026926 | 0.556750599 | 0.501567889 | 2.445329533 | 0.014471984 | 0.252703112 | 2.527031124 |
| Gria4 | 7.6431046 | 0.63866895 | 0.524021319 | 2.488014723 | 0.012845841 | 0.252703112 | 2.527031124 |
| 4930402H24Rik | 3.5811813 | 0.682235903 | 0.819985824 | 2.611144812 | 0.009023967 | 0.252703112 | 2.527031124 |
| Tmsb4x | 4.6209717 | 0.474770331 | 0.873328434 | 2.502494228 | 0.012332164 | 0.252703112 | 2.527031124 |
| Gm42418 | 88.84172 | -0.191122891 | 0.375592636 | -2.412546334 | 0.015841524 | 0.256858993 | 2.568589928 |
| Rpl30 | 4.705174 | 0.196268769 | 0.454872011 | 2.236204392 | 0.025338386 | 0.383454237 | 3.834542365 |
| AY036118 | 14.886724 | -0.203413815 | 0.381781468 | -2.10115155 | 0.035627665 | 0.44930444 | 4.493044403 |
| Ctnna2 | 5.1470156 | 0.312291348 | 0.489323226 | 2.126484051 | 0.033462976 | 0.44930444 | 4.493044403 |
| Nkain2 | 3.9198008 | -0.144269504 | 0.325175688 | -2.106268939 | 0.035180998 | 0.44930444 | 4.493044403 |
| Sec61a1 | 55.388393 | 0.3403631 | 0.248593673 | 2.053560593 | 0.040018239 | 0.47811265 | 4.781126505 |
| Rpl39 | 2.3683107 | 0.142671682 | 0.389169964 | 2.01043198 | 0.044385489 | 0.503775296 | 5.037752958 |
| Gm11337 | 4.2377167 | 0.243809896 | 0.407318885 | 1.976578379 | 0.048089308 | 0.519822521 | 5.198225211 |
| Rps21 | 6.852249 | 0.19654383 | 0.4035426 | 1.934366297 | 0.053068107 | 0.547566379 | 5.475663788 |
| 1700054A03Rik | 52.824097 | 0.261536736 | 0.275564355 | 1.660799631 | 0.096753695 | 0.563156124 | 5.631561236 |
| Cox8a | 6.496518 | 0.113112307 | 0.379215707 | 1.693300666 | 0.090398251 | 0.563156124 | 5.631561236 |
| Lrrc4c | 2.4291487 | -0.154076835 | 0.325645587 | -1.686710574 | 0.091659018 | 0.563156124 | 5.631561236 |
| Ptprd | 2.9578784 | -0.126803392 | 0.309129418 | -1.666676461 | 0.095578756 | 0.563156124 | 5.631561236 |
| D430042O09Rik | 15.700211 | 0.25305849 | 0.339934992 | 1.688757544 | 0.091265903 | 0.563156124 | 5.631561236 |
| Hdac9 | 2.0922823 | -0.154825247 | 0.330013102 | -1.721239753 | 0.085207327 | 0.563156124 | 5.631561236 |
| Adamtsl1 | 11.004572 | -0.247656944 | 0.336797595 | -1.665844045 | 0.095744481 | 0.563156124 | 5.631561236 |
| Dop1b | 4.380478 | -0.186660648 | 0.34726631 | -1.741672638 | 0.081565743 | 0.563156124 | 5.631561236 |
| Wdr72 | 7.101959 | -0.248488226 | 0.364046322 | -1.719782185 | 0.085472042 | 0.563156124 | 5.631561236 |
| Chl1 | 4.161993 | 0.194916913 | 0.364419695 | 1.83607851 | 0.066346046 | 0.563156124 | 5.631561236 |
| Rps24 | 3.1835344 | 0.147099604 | 0.361480987 | 1.790541518 | 0.073366899 | 0.563156124 | 5.631561236 |
| Tpt1 | 4.37262 | -0.052940727 | 0.262502667 | -1.850943301 | 0.06417771 | 0.563156124 | 5.631561236 |
| Il1rapl2 | 2.7269056 | 0.13968471 | 0.332669813 | 1.717620676 | 0.085865827 | 0.563156124 | 5.631561236 |
| Calm1 | 3.654584 | -0.061883795 | 0.272803241 | -1.697989781 | 0.089509691 | 0.563156124 | 5.631561236 |
| 4930441J16Rik | 3.6492968 | 0.167810502 | 0.351953207 | 1.757902444 | 0.07876411 | 0.563156124 | 5.631561236 |
| Ankrd12 | 7.1624546 | 0.24650352 | 0.375063401 | 1.802978398 | 0.071391607 | 0.563156124 | 5.631561236 |
| Rpl11 | 4.7606373 | 0.109606751 | 0.377284085 | 1.721457946 | 0.085167757 | 0.563156124 | 5.631561236 |
| Rplp1 | 7.5951495 | 0.153456858 | 0.359931146 | 1.621728052 | 0.10486159 | 0.566751925 | 5.667519248 |
| Csmd3 | 7.3274617 | -0.144269504 | 0.325175689 | 1.623477413 | 0.104487388 | 0.566751925 | 5.667519248 |
| Ptpn9 | 6.340883 | -0.144269504 | 0.325175688 | 1.626611001 | 0.103819741 | 0.566751925 | 5.667519248 |
| Lyrm1 | 2.0649328 | -0.106922372 | 0.297332706 | -1.562110625 | 0.11826193 | 0.624312981 | 6.243129808 |
| Mt1 | 3.035079 | -0.072828842 | 0.282490684 | -1.428940261 | 0.153021404 | 0.640428195 | 6.404281949 |
| Fam169a | 6.7369084 | -0.18340204 | 0.319235729 | -1.513134013 | 0.130245622 | 0.640428195 | 6.404281949 |
| mt-Nd1 | 13.160506 | -0.116934416 | 0.300101372 | -1.469456434 | 0.14170903 | 0.640428195 | 6.404281949 |
| Epha6 | 12.959585 | -0.197251177 | 0.304780365 | -1.474898807 | 0.140239751 | 0.640428195 | 6.404281949 |
| Cst3 | 20.57817 | 0.159508532 | 0.333286368 | 1.443185572 | 0.148968203 | 0.640428195 | 6.404281949 |
| Ppp1r16a | 5.960048 | 0.162739497 | 0.312080006 | 1.44147094 | 0.149451681 | 0.640428195 | 6.404281949 |
| Apoe | 4.915247 | 0.168137651 | 0.323071493 | 1.515112425 | 0.129743936 | 0.640428195 | 6.404281949 |
| Gm14582 | 4.4432197 | -0.144269504 | 0.325175691 | 1.506361389 | 0.131974427 | 0.640428195 | 6.404281949 |
| Hexb | 3.3334806 | -0.070734684 | 0.283679208 | -1.423219334 | 0.154672583 | 0.640428195 | 6.404281949 |
| Vit | 1.6428826 | -0.104812619 | 0.293341135 | -1.421505599 | 0.155169827 | 0.640428195 | 6.404281949 |
| Alpl | 3.0174866 | -0.140277684 | 0.306789004 | -1.481063629 | 0.138589619 | 0.640428195 | 6.404281949 |
| Tacc1 | 1.8357242 | -0.144269504 | 0.325175692 | -1.501173511 | 0.133310689 | 0.640428195 | 6.404281949 |
| Rpl35a | 4.942784 | 0.079615002 | 0.353369283 | 1.375895816 | 0.168853891 | 0.64965819 | 6.496581898 |
| Pde1c | 8.5778265 | -0.185324725 | 0.29802617 | -1.402720332 | 0.160700252 | 0.64965819 | 6.496581898 |
| Ntm | 12.9065 | -0.188815693 | 0.290463432 | -1.393815711 | 0.163373263 | 0.64965819 | 6.496581898 |
| Rpl23 | 3.8133543 | -0.046198235 | 0.27070535 | -1.380576633 | 0.167409172 | 0.64965819 | 6.496581898 |
| Rpl13 | 5.65667 | 0.113485612 | 0.316393903 | 1.34990126 | 0.177047657 | 0.650383675 | 6.503836754 |
| Gm50306 | 7.646293 | -0.13767914 | 0.300112689 | -1.353078876 | 0.176030433 | 0.650383675 | 6.503836754 |
| Cpe | 2.8087764 | -0.086315764 | 0.287001879 | -1.348063897 | 0.177637832 | 0.650383675 | 6.503836754 |
| mt-Cytb | 17.050423 | -0.149010167 | 0.2941164 | -1.31159177 | 0.189657911 | 0.676105448 | 6.761054478 |
| Rps7 | 1.746049 | -0.050939669 | 0.268529654 | -1.30874708 | 0.190620038 | 0.676105448 | 6.761054478 |
| Acoxl | 4.884706 | -0.151670275 | 0.290069682 | -1.293574371 | 0.19581248 | 0.683837429 | 6.838374291 |
| Zfhx3 | 3.9178238 | -0.144269504 | 0.325175691 | 1.27483507 | 0.202367625 | 0.696021982 | 6.960219824 |
| Gphn | 2.9946876 | -0.068421919 | 0.282712928 | -1.260825286 | 0.207371801 | 0.702588043 | 7.025880432 |
| 4930558K02Rik | 5.919937 | -0.132525398 | 0.282603996 | -1.210432005 | 0.226113167 | 0.744924917 | 7.449249171 |
| Fbxl7 | 4.7657743 | -0.127052022 | 0.28377821 | -1.209603897 | 0.226430922 | 0.744924917 | 7.449249171 |
| Mc1r | 10.825235 | -0.144485613 | 0.2754687 | -1.183315184 | 0.236684257 | 0.748825722 | 7.488257219 |
| Phkg1 | 4.360975 | -0.144269504 | 0.325175689 | -1.075975182 | 0.281938356 | 0.748825722 | 7.488257219 |
| Rab29 | 4.6720915 | 0.092935635 | 0.291279726 | 1.033307415 | 0.301460058 | 0.748825722 | 7.488257219 |
| Lrrc17 | 4.172238 | -0.144269504 | 0.325175689 | 1.086127144 | 0.277422745 | 0.748825722 | 7.488257219 |
| Rps14 | 3.1900764 | -0.025318107 | 0.26480946 | -1.038338018 | 0.299112715 | 0.748825722 | 7.488257219 |
| Klkb1 | 2.2612617 | -0.082141371 | 0.277670762 | -1.092517724 | 0.274605604 | 0.748825722 | 7.488257219 |
| Tshz2 | 10.283613 | -0.14364658 | 0.257058162 | -1.113952074 | 0.265299748 | 0.748825722 | 7.488257219 |
| Unc5d | 11.152448 | 0.120127075 | 0.267316144 | 1.035677792 | 0.300352486 | 0.748825722 | 7.488257219 |

| | | | | | | | |
|---|---|---|---|---|---|---|---|
| Xlr5a | 27.32727 | 0.14038274 | 0.228098145 | 1.046212485 | 0.29546294 | 0.748825722 | 7.488257219 |
| Zbtb7a | 6.3876896 | -0.131928425 | 0.260215716 | -1.076413748 | 0.281742256 | 0.748825722 | 7.488257219 |
| mt-Atp6 | 52.440006 | -0.104050192 | 0.275944329 | -1.028980499 | 0.303488839 | 0.748825722 | 7.488257219 |
| Hapln1 | 3.8963327 | -0.144269504 | 0.325175692 | -1.136321744 | 0.255821934 | 0.748825722 | 7.488257219 |
| mt-Nd4 | 15.204892 | -0.102436257 | 0.283778613 | -1.114940938 | 0.264875732 | 0.748825722 | 7.488257219 |
| Rpl32 | 3.3037553 | 0.066369241 | 0.32801883 | 1.153151608 | 0.248848162 | 0.748825722 | 7.488257219 |
| mt-Nd5 | 4.970977 | 0.090726347 | 0.305349636 | 1.177534327 | 0.238982305 | 0.748825722 | 7.488257219 |
| Galnt13 | 6.1900806 | -0.109214386 | 0.273011608 | -1.035961177 | 0.300220255 | 0.748825722 | 7.488257219 |
| Dpyd | 4.496455 | -0.144269504 | 0.32517569 | -1.075975213 | 0.281938342 | 0.748825722 | 7.488257219 |
| Ctnna3 | 7.525647 | 0.116417999 | 0.278270182 | 1.057845426 | 0.290125919 | 0.748825722 | 7.488257219 |
| Eef1a1 | 4.1184964 | 0.078032559 | 0.302427574 | 1.05288792 | 0.292392363 | 0.748825722 | 7.488257219 |
| Cox6b1 | 3.6377096 | -0.072211825 | 0.280756301 | -1.053655417 | 0.292040707 | 0.748825722 | 7.488257219 |
| Gm49497 | 2.7023165 | 0.087057828 | 0.285457673 | 1.076708537 | 0.281610497 | 0.748825722 | 7.488257219 |
| Cfap74 | 2.7922401 | 0.091807542 | 0.294243867 | 1.131594546 | 0.257804934 | 0.748825722 | 7.488257219 |
| Cadm2 | 1.128733 | -0.049109114 | 0.272086738 | -1.047392794 | 0.294918454 | 0.748825722 | 7.488257219 |
| Ninj2 | 4.790965 | 0.09601654 | 0.275638481 | 0.979056211 | 0.327552206 | 0.750414487 | 7.504144867 |
| Cntnap2 | 15.640811 | -0.11977317 | 0.258474464 | -1.001738067 | 0.316470116 | 0.750414487 | 7.504144867 |
| Fyb2 | 4.7031064 | 0.091257882 | 0.282255546 | 0.978370883 | 0.327890924 | 0.750414487 | 7.504144867 |
| Camk1d | 9.3028755 | -0.069061892 | 0.280703015 | -1.008674754 | 0.313130643 | 0.750414487 | 7.504144867 |
| Shb | 15.204553 | -0.144269504 | 0.325175683 | 0.990194213 | 0.322079201 | 0.750414487 | 7.504144867 |
| Spock1 | 3.860509 | 0.083644551 | 0.278455418 | 0.975747905 | 0.329189406 | 0.750414487 | 7.504144867 |
| Ms4a1 | 6.423043 | -0.107726758 | 0.261772711 | -0.966318444 | 0.33388486 | 0.750414487 | 7.504144867 |
| Hdac8 | 3.5206144 | 0.086380811 | 0.275547572 | 0.973400155 | 0.330354462 | 0.750414487 | 7.504144867 |
| 4930511M06Rik | 7.9351397 | 0.107495058 | 0.268488173 | 0.966449315 | 0.333819397 | 0.750414487 | 7.504144867 |
| Trim2 | 9.635662 | -0.115345236 | 0.247128607 | -0.938547583 | 0.347963076 | 0.76687008 | 7.668700801 |
| Rpl12 | 2.0700948 | -0.051411782 | 0.276250951 | -0.939140487 | 0.34765862 | 0.76687008 | 7.668700801 |
| Ppp1r9a | 4.709364 | 0.082379014 | 0.281211308 | 0.909799899 | 0.362928048 | 0.784092116 | 7.84092116 |
| Rps20 | 2.5459812 | -0.030998388 | 0.274385395 | -0.891240844 | 0.372799977 | 0.784092116 | 7.84092116 |
| Plp1 | 2.3849928 | -0.025619624 | 0.269104434 | -0.884359123 | 0.376502382 | 0.784092116 | 7.84092116 |
| Cox6c | 3.8020508 | 0.070636592 | 0.284250481 | 0.885465914 | 0.375905396 | 0.784092116 | 7.84092116 |
| Dlg2 | 1.8535248 | 0.062142206 | 0.278734595 | 0.893676521 | 0.371494991 | 0.784092116 | 7.84092116 |
| Rpl9 | 3.0056663 | 0.048831028 | 0.312604645 | 0.90894579 | 0.363378741 | 0.784092116 | 7.84092116 |
| Vwc2l | 4.158543 | -0.079236819 | 0.267503585 | -0.876323363 | 0.380854268 | 0.785944716 | 7.859447162 |
| mt-Co3 | 64.31273 | -0.083497409 | 0.254951607 | -0.760497136 | 0.446957481 | 0.803152885 | 8.031528847 |
| mt-Nd3 | 11.394203 | -0.065447363 | 0.273451889 | -0.793441706 | 0.42752052 | 0.803152885 | 8.031528847 |
| Prkg1 | 7.6053395 | -0.068930566 | 0.256909116 | -0.681845609 | 0.495336581 | 0.803152885 | 8.031528847 |
| 5830416I19Rik | 24.880907 | -0.097975075 | 0.202493001 | -0.75287459 | 0.451525279 | 0.803152885 | 8.031528847 |
| Rpl34 | 1.8285625 | 0.030865893 | 0.303035065 | 0.685451362 | 0.493059143 | 0.803152885 | 8.031528847 |
| mt-Nd2 | 16.228426 | -0.053170589 | 0.273357441 | -0.693766572 | 0.487828609 | 0.803152885 | 8.031528847 |
| Aldoc | 1.6210175 | 0.030938377 | 0.310619826 | 0.70100135 | 0.483302173 | 0.803152885 | 8.031528847 |
| Cdk8 | 4.057161 | -0.050264498 | 0.276944999 | -0.821684182 | 0.411256664 | 0.803152885 | 8.031528847 |
| Rps17 | 1.9367694 | -0.028147181 | 0.275133307 | -0.746550852 | 0.455334725 | 0.803152885 | 8.031528847 |
| Wdyhv1 | 5.2000837 | 0.067553491 | 0.257973728 | 0.690415027 | 0.48993323 | 0.803152885 | 8.031528847 |
| Rps27 | 2.3981893 | 0.04319335 | 0.283232745 | 0.698243428 | 0.485024971 | 0.803152885 | 8.031528847 |
| Rps3 | 1.8366036 | 0.032617785 | 0.315273426 | 0.76770084 | 0.442664937 | 0.803152885 | 8.031528847 |
| Rps8 | 4.412556 | 0.039203894 | 0.311631617 | 0.742805572 | 0.457599401 | 0.803152885 | 8.031528847 |
| Rsrc1 | 12.455451 | 0.086436245 | 0.247556806 | 0.751689755 | 0.452237653 | 0.803152885 | 8.031528847 |
| Scamp1 | 4.1156707 | 0.068081352 | 0.266826967 | 0.740363454 | 0.459079488 | 0.803152885 | 8.031528847 |
| Ahdc1 | 3.5728967 | -0.144269504 | 0.325175687 | -0.841559953 | 0.400034314 | 0.803152885 | 8.031528847 |
| Sil1 | 7.6066775 | 0.079962286 | 0.251980227 | 0.732376054 | 0.46393907 | 0.803152885 | 8.031528847 |
| AI314278 | 3.58537 | 0.060918113 | 0.281900009 | 0.80539429 | 0.420592186 | 0.803152885 | 8.031528847 |
| Cep128 | 2.7124422 | 0.064844386 | 0.275612728 | 0.807938647 | 0.419125902 | 0.803152885 | 8.031528847 |
| Gnao1 | 2.0144234 | -0.057811355 | 0.268161259 | -0.804729232 | 0.420975947 | 0.803152885 | 8.031528847 |
| Lsamp | 5.6331697 | -0.065491312 | 0.262828975 | -0.706744963 | 0.479724982 | 0.803152885 | 8.031528847 |
| Kctd16 | 20.133091 | 0.090127104 | 0.259986983 | 0.790265722 | 0.429372601 | 0.803152885 | 8.031528847 |
| Gm10062 | 13.044163 | -0.092537378 | 0.256702011 | -0.831361272 | 0.405769571 | 0.803152885 | 8.031528847 |
| Dctd | 2.5801604 | -0.051003254 | 0.269734956 | -0.717403764 | 0.473124998 | 0.803152885 | 8.031528847 |
| Meg3 | 3.3725424 | -0.049964658 | 0.274421008 | -0.801893435 | 0.422614604 | 0.803152885 | 8.031528847 |
| Mt3 | 4.3413577 | 0.042696499 | 0.299697659 | 0.697275913 | 0.485630139 | 0.803152885 | 8.031528847 |
| Nckap5 | 23.137186 | -0.090323525 | 0.224958217 | -0.720182722 | 0.471412501 | 0.803152885 | 8.031528847 |
| Nell1 | 5.491928 | -0.066527026 | 0.262695739 | -0.72785343 | 0.466703313 | 0.803152885 | 8.031528847 |
| Pcca | 3.6836677 | 0.061343675 | 0.26462478 | 0.691395253 | 0.489317186 | 0.803152885 | 8.031528847 |
| Gria1 | 1.568388 | -0.144269504 | 0.325175692 | 0.765567483 | 0.443933698 | 0.803152885 | 8.031528847 |
| Gnaz | 13.247394 | -0.08488706 | 0.218488065 | -0.67497225 | 0.499693396 | 0.804470928 | 8.04470928 |
| Tmeff2 | 3.7606382 | -0.037821803 | 0.275693293 | -0.640436359 | 0.521888951 | 0.809067265 | 8.090672646 |
| Fau | 2.4518764 | -0.02292522 | 0.276727379 | -0.568573272 | 0.569645768 | 0.809067265 | 8.090672646 |
| Sgip1 | 1.58918 | 0.030117702 | 0.280350193 | 0.557162738 | 0.577416246 | 0.809067265 | 8.090672646 |
| Cmss1 | 5.8559604 | -0.061361049 | 0.250594615 | -0.608515875 | 0.542845381 | 0.809067265 | 8.090672646 |
| mt-Co1 | 29.908287 | -0.053367891 | 0.264740746 | -0.592423798 | 0.553566834 | 0.809067265 | 8.090672646 |
| Klhl14 | 26.143715 | -0.079788262 | 0.19349247 | -0.618039232 | 0.536549478 | 0.809067265 | 8.090672646 |
| Kcnq5 | 2.647503 | -0.039358879 | 0.268669341 | -0.594945078 | 0.551880184 | 0.809067265 | 8.090672646 |
| Rpl36 | 0.8958757 | 0.020396489 | 0.321162203 | 0.563917901 | 0.572810007 | 0.809067265 | 8.090672646 |
| Ptprg | 2.6142354 | -0.045670873 | 0.263823795 | -0.614747426 | 0.538721534 | 0.809067265 | 8.090672646 |
| Ckb | 3.2405775 | -0.025927105 | 0.279826006 | -0.554920453 | 0.578949072 | 0.809067265 | 8.090672646 |
| Gm38973 | 4.666378 | -0.04851334 | 0.263565331 | -0.595715128 | 0.55136555 | 0.809067265 | 8.090672646 |
| Cdh23 | 7.7059603 | -0.067907089 | 0.249676346 | -0.637874309 | 0.523555502 | 0.809067265 | 8.090672646 |
| Ubb | 1.2982892 | 0.025797643 | 0.292741315 | 0.567200924 | 0.57057768 | 0.809067265 | 8.090672646 |
| Pde6a | 17.578203 | 0.070570967 | 0.231208601 | 0.584808871 | 0.558676239 | 0.809067265 | 8.090672646 |
| Bc1 | 2.8435912 | 0.038811929 | 0.279602361 | 0.578003412 | 0.563261815 | 0.809067265 | 8.090672646 |
| Camk2a | 5.2381573 | 0.059581195 | 0.257324944 | 0.614073112 | 0.539167015 | 0.809067265 | 8.090672646 |

| | | | | | | | |
|---|---|---|---|---|---|---|---|
| Rpl17 | 2.507469 | 0.032177912 | 0.295780459 | 0.628274548 | 0.529824102 | 0.809067265 | 8.090672646 |
| Car12 | 3.4988654 | 0.029134474 | 0.301561664 | 0.551982717 | 0.580960194 | 0.809067265 | 8.090672646 |
| Cox7c | 4.358693 | -0.029501603 | 0.277875025 | -0.595976452 | 0.551190958 | 0.809067265 | 8.090672646 |
| 6030443J06Rik | 7.6030917 | 0.063576813 | 0.24287249 | 0.578493834 | 0.562930759 | 0.809067265 | 8.090672646 |
| Cox4i1 | 4.267222 | -0.030134066 | 0.27773729 | -0.564835044 | 0.572185969 | 0.809067265 | 8.090672646 |
| Uba52 | 1.9512028 | -0.0298059 | 0.27412602 | -0.615935776 | 0.537936909 | 0.809067265 | 8.090672646 |
| Gm12216 | 27.355824 | 0.06748177 | 0.19122808 | 0.531143054 | 0.595319651 | 0.821684868 | 8.21684868 |
| Tmem108 | 3.9934008 | -0.144269504 | 0.325175691 | -0.528344968 | 0.597259926 | 0.821684868 | 8.21684868 |
| Rps16 | 2.7770374 | 0.028103901 | 0.294175746 | 0.513800132 | 0.607391736 | 0.825616312 | 8.256163117 |
| mt-Nd4l | 1.7701625 | -0.014452585 | 0.276604788 | -0.517921221 | 0.604513235 | 0.825616312 | 8.256163117 |
| Slx1b | 6.8200607 | -0.049143541 | 0.252225264 | -0.496126445 | 0.619805202 | 0.837474886 | 8.374748862 |
| Kat7 | 4.219699 | -0.050317713 | 0.241301178 | -0.485749417 | 0.627144842 | 0.842377983 | 8.423779828 |
| Cmpk1 | 15.532138 | -0.052695045 | 0.200454179 | -0.432430793 | 0.665428342 | 0.863818223 | 8.638182233 |
| Dbi | 2.738859 | 0.018590742 | 0.301609568 | 0.43707579 | 0.662056384 | 0.863818223 | 8.638182233 |
| Rpl37 | 5.0523815 | 0.027316758 | 0.290169888 | 0.442184126 | 0.658355972 | 0.863818223 | 8.638182233 |
| Lrp1b | 56.844112 | -0.054597822 | 0.187053612 | -0.431727939 | 0.665939159 | 0.863818223 | 8.638182233 |
| Rpl26 | 6.793167 | 0.030073053 | 0.285309677 | 0.449134265 | 0.653334804 | 0.863818223 | 8.638182233 |
| Rpl13a | 1.4650565 | -0.021898981 | 0.27408584 | -0.452768472 | 0.650715469 | 0.863818223 | 8.638182233 |
| Ppp3ca | 1.3173085 | -0.022005466 | 0.266971317 | -0.40326017 | 0.68675684 | 0.885760242 | 8.857602421 |
| Mbp | 3.1925125 | -0.013916181 | 0.279665754 | -0.387489681 | 0.698393723 | 0.890648175 | 8.906481745 |
| Rps27a | 2.27826 | 0.023354561 | 0.279157355 | 0.388140309 | 0.697912204 | 0.890648175 | 8.906481745 |
| Rpl21 | 3.447512 | 0.018283348 | 0.290732465 | 0.33562308 | 0.737155118 | 0.899646299 | 8.996462994 |
| Kcnip4 | 2.905939 | 0.026364942 | 0.262740725 | 0.344455868 | 0.73050348 | 0.899646299 | 8.996462994 |
| Ttc28 | 5.5357037 | 0.031715466 | 0.263658606 | 0.361762165 | 0.717529768 | 0.899646299 | 8.996462994 |
| Rpl27a | 4.768087 | 0.023442639 | 0.282332266 | 0.366973953 | 0.713638447 | 0.899646299 | 8.996462994 |
| Kcnd2 | 2.514149 | 0.023039772 | 0.274430089 | 0.356224093 | 0.72167275 | 0.899646299 | 8.996462994 |
| Rpl38 | 7.104176 | 0.02443584 | 0.281823468 | 0.368491442 | 0.712506825 | 0.899646299 | 8.996462994 |
| Igf2bp3 | 12.022297 | -0.040527887 | 0.2256576 | -0.349693262 | 0.726568911 | 0.899646299 | 8.996462994 |
| Uqcrh | 1.8911818 | -0.016606154 | 0.277152961 | -0.339897798 | 0.733933495 | 0.899646299 | 8.996462994 |
| Gm49767 | 2.9797692 | 0.022575997 | 0.259354442 | 0.286546698 | 0.774459432 | 0.920430844 | 9.204308437 |
| Anks1b | 1.6476322 | -0.144269504 | 0.325175691 | -0.286597305 | 0.774420678 | 0.920430844 | 9.204308437 |
| Il27 | 8.782036 | -0.144269504 | 0.325175688 | -0.287506005 | 0.773724905 | 0.920430844 | 9.204308437 |
| Esf1 | 2.2181897 | -0.020562664 | 0.263379321 | -0.301130064 | 0.763315317 | 0.920430844 | 9.204308437 |
| mt-Co2 | 31.080164 | -0.03022148 | 0.249492677 | -0.296619466 | 0.766757047 | 0.920430844 | 9.204308437 |
| Ehbp1 | 10.738027 | 0.030243757 | 0.234573671 | 0.274574797 | 0.783642932 | 0.926494508 | 9.264945077 |
| 1700126G02Rik | 9.276643 | 0.03139287 | 0.216010043 | 0.265712645 | 0.790460524 | 0.929712637 | 9.29712637 |
| 8430422H06Rik | 8.173612 | -0.023156215 | 0.240426051 | -0.218725954 | 0.82686353 | 0.936528355 | 9.365283551 |
| Atxn7l1 | 2.1083577 | -0.015485012 | 0.263796199 | -0.231538242 | 0.81689668 | 0.936528355 | 9.365283551 |
| Zmat4 | 6.7431383 | 0.022529386 | 0.246711128 | 0.22684464 | 0.820544551 | 0.936528355 | 9.365283551 |
| Gas5 | 0.86245275 | 0.009330518 | 0.273485417 | 0.199798957 | 0.841637817 | 0.936528355 | 9.365283551 |
| Rps3a1 | 6.1928663 | 0.007860253 | 0.297455268 | 0.20320128 | 0.838977713 | 0.936528355 | 9.365283551 |
| St18 | 2.3145986 | -0.013232787 | 0.267155578 | -0.200992569 | 0.840704385 | 0.936528355 | 9.365283551 |
| Srgn | 2.2238126 | 0.013866806 | 0.275903826 | 0.23431069 | 0.814743788 | 0.936528355 | 9.365283551 |
| Prkag2 | 4.1486835 | 0.018246375 | 0.25427631 | 0.204517681 | 0.837948977 | 0.936528355 | 9.365283551 |
| Rapgef4 | 13.728402 | -0.144269504 | 0.325175686 | -0.24745277 | 0.804557839 | 0.936528355 | 9.365283551 |
| Mapk14 | 11.374496 | 0.026125042 | 0.218565173 | 0.218902552 | 0.826725959 | 0.936528355 | 9.365283551 |
| Rps23 | 1.558236 | 0.008382039 | 0.290411303 | 0.202850854 | 0.839251608 | 0.936528355 | 9.365283551 |
| Ccdc60 | 3.9716513 | -0.01651348 | 0.252042935 | -0.185699416 | 0.852680486 | 0.94418766 | 9.4418766 |
| Selenow | 2.1228044 | 0.010666727 | 0.271834341 | 0.169706388 | 0.86524105 | 0.953445235 | 9.534452349 |
| Arvcf | 1.1981798 | -0.008236574 | 0.27181469 | -0.161465207 | 0.871727013 | 0.95413146 | 9.541314598 |
| Sema3c | 24.274586 | -0.019797044 | 0.192818218 | -0.158236763 | 0.874270236 | 0.95413146 | 9.541314598 |
| Tns3 | 2.8131933 | 0.009492171 | 0.262328255 | 0.124347089 | 0.901040465 | 0.964793328 | 9.647933278 |
| Gm960 | 1.248124 | 0.006599501 | 0.273399828 | 0.126734773 | 0.899150323 | 0.964793328 | 9.647933278 |
| Syt1 | 2.76881 | -0.144269504 | 0.32517569 | -0.127154613 | 0.898818027 | 0.964793328 | 9.647933278 |
| Ctcflos | 5.029918 | 0.011848715 | 0.250348254 | 0.129090462 | 0.897286069 | 0.964793328 | 9.647933278 |
| Svil | 2.9297297 | -0.144269504 | 0.325175691 | 0.118298633 | 0.90583104 | 0.965369231 | 9.653692309 |
| Ralyl | 3.207875 | 0.008547868 | 0.256506071 | 0.105882814 | 0.91567533 | 0.967270116 | 9.672701157 |
| Pcdh15 | 10.199923 | 0.010982523 | 0.241623588 | 0.105301095 | 0.916136894 | 0.967270116 | 9.672701157 |
| Gm765 | 2.7776783 | 0.007397864 | 0.258850476 | 0.093708169 | 0.925340981 | 0.972464827 | 9.724648273 |
| Pcgf6 | 3.4580526 | -0.005435331 | 0.253983553 | -0.065553943 | 0.947732958 | 0.988492345 | 9.884923448 |
| Cspp1 | 3.2586973 | -0.144269504 | 0.32517569 | 0.06358457 | 0.949301018 | 0.988492345 | 9.884923448 |
| Malat1 | 6.5285845 | -0.002568976 | 0.284969702 | -0.054126926 | 0.95683404 | 0.991786882 | 9.917868817 |
| Ppia | 1.9710991 | 0.001253321 | 0.288209213 | 0.02916322 | 0.976734415 | 0.994348678 | 9.943486785 |
| Adk | 3.7155418 | -0.00219536 | 0.255652754 | -0.026765123 | 0.978647071 | 0.994348678 | 9.943486785 |
| Gm29325 | 4.275987 | -0.001717868 | 0.239048636 | -0.033712984 | 0.973106025 | 0.994348678 | 9.943486785 |
| Sult2a2 | 2.8649654 | 0.001757449 | 0.257371301 | 0.022404405 | 0.982125366 | 0.994348678 | 9.943486785 |
| Rps29 | 11.573063 | -0.000293598 | 0.282949937 | -0.007447561 | 0.994057761 | 0.994348678 | 9.943486785 |
| Cntn3 | 2.9678428 | -0.144269504 | 0.325175693 | -0.00708294 | 0.994348678 | 0.994348678 | 9.943486785 |
| Zfp994 | 3.931868 | 0.000607642 | 0.267142038 | 0.007521999 | 0.993998369 | 0.994348678 | 9.943486785 |
| Rpl37a | 4.900374 | -0.002148273 | 0.278221984 | -0.038038945 | 0.969656631 | 0.994348678 | 9.943486785 |

| **Glut** | baseMean | log2FoldChange | lfcSE | stat | pvalue | padj | pglobadj |
|---|---|---|---|---|---|---|---|
| Rps12 | 35.98471 | 2.462618387 | 0.406953734 | 6.397346936 | 1.58E-10 | 2.00E-08 | 2.00E-07 |
| Rps3 | 19.521986 | 1.821425842 | 0.306300449 | 6.319397497 | 2.63E-10 | 2.00E-08 | 2.00E-07 |
| Cst3 | 153.94177 | 1.856793114 | 0.464086646 | 4.443558516 | 8.85E-06 | 0.000268989 | 0.002689886 |
| Rps10 | 27.12138 | 1.603959273 | 0.401840125 | 4.477810031 | 7.54E-06 | 0.000268989 | 0.002689886 |
| Rpl35a | 37.172855 | 1.60762834 | 0.401135716 | 4.487610983 | 7.20E-06 | 0.000268989 | 0.002689886 |
| Rps8 | 42.907177 | 1.664560564 | 0.424147964 | 4.404133431 | 1.06E-05 | 0.000269059 | 0.002690592 |
| Galnt13 | 29.213446 | -0.144269504 | 0.471400307 | -3.858133993 | 0.000114256 | 0.002443891 | 0.02443891 |
| Dbi | 18.818296 | 1.113930842 | 0.337434072 | 3.829074932 | 0.000128626 | 0.002443891 | 0.02443891 |

| | | | | | | | |
|---|---|---|---|---|---|---|---|
| Rpl13 | 48.121593 | 1.650362231 | 0.534090385 | 3.668606979 | 0.000243876 | 0.00370691 | 0.037069098 |
| Rplp1 | 76.570724 | 1.314431554 | 0.417330582 | 3.681440735 | 0.00023192 | 0.00370691 | 0.037069098 |
| Rps20 | 30.235338 | 1.381924585 | 0.457630817 | 3.602590946 | 0.000315061 | 0.004353572 | 0.043535718 |
| Rpl17 | 26.852716 | 1.531550737 | 0.51637146 | 3.56347007 | 0.000365984 | 0.004635803 | 0.046358029 |
| Rpl26 | 34.769566 | 1.297250781 | 0.459194139 | 3.426798786 | 0.000610741 | 0.007140977 | 0.071409769 |
| Rps24 | 50.57764 | 1.503141605 | 0.565610503 | 3.304882432 | 0.000950162 | 0.009628313 | 0.096283132 |
| Rps14 | 37.294437 | 1.244713847 | 0.459021308 | 3.32265831 | 0.000891641 | 0.009628313 | 0.096283132 |
| Rps16 | 24.292126 | 1.217056806 | 0.457917771 | 3.278733742 | 0.00104274 | 0.009906027 | 0.09906027 |
| Fau | 36.68445 | 1.630510592 | 0.640071173 | 3.233767028 | 0.001221691 | 0.010794028 | 0.107940281 |
| Ntm | 60.91025 | -0.581516333 | 0.199212735 | -3.220820866 | 0.00127824 | 0.010794028 | 0.107940281 |
| Trim2 | 42.692062 | -0.538604908 | 0.186625775 | -3.139556489 | 0.001692038 | 0.013536303 | 0.135363031 |
| Rpl38 | 36.311028 | 1.037057315 | 0.418288474 | 3.085191768 | 0.002034209 | 0.014845266 | 0.148452657 |
| Apoe | 80.2238 | 1.008340739 | 0.401340801 | 3.082747549 | 0.002050991 | 0.014845266 | 0.148452657 |
| Cox4i1 | 55.03479 | 1.34039575 | 0.569909768 | 3.048594145 | 0.002299149 | 0.015885027 | 0.158850267 |
| Acoxl | 21.082848 | -0.144269504 | 0.471400317 | -2.996555908 | 0.002730482 | 0.018044922 | 0.180449222 |
| Cdk8 | 18.143866 | -1.190074525 | 0.576439192 | -2.962951615 | 0.003047045 | 0.019297949 | 0.192979492 |
| Rpl23 | 26.072792 | 1.197977174 | 0.554634361 | 2.897383028 | 0.0037629 | 0.022878434 | 0.228784338 |
| Mt3 | 35.23202 | 1.143254956 | 0.533883748 | 2.860125886 | 0.004234729 | 0.02330034 | 0.233003398 |
| Rpl37 | 58.321743 | 1.035065567 | 0.46873541 | 2.868185785 | 0.00412833 | 0.02330034 | 0.233003398 |
| Rps7 | 21.615023 | 1.174941211 | 0.557227003 | 2.855850795 | 0.004292168 | 0.02330034 | 0.233003398 |
| Tmem108 | 19.81278 | -0.144269504 | 0.471400324 | -2.773301939 | 0.005549061 | 0.028880434 | 0.288804341 |
| Cox6c | 36.238724 | 1.172125192 | 0.586739812 | 2.76455294 | 0.005700086 | 0.028880434 | 0.288804341 |
| Rpl27a | 34.65671 | 1.01403714 | 0.499879395 | 2.737655009 | 0.006187895 | 0.030340647 | 0.303406469 |
| Sil1 | 27.170454 | -0.534987033 | 0.220861632 | -2.71830927 | 0.006561647 | 0.031167825 | 0.311678251 |
| Rps21 | 69.29485 | 1.015503966 | 0.50834811 | 2.702966401 | 0.006872369 | 0.031654549 | 0.316545494 |
| Rpl32 | 26.072998 | 0.876854995 | 0.443177614 | 2.651588399 | 0.008011414 | 0.034792426 | 0.347924259 |
| Gm50306 | 32.413666 | -0.144269504 | 0.471400281 | -2.658191093 | 0.007856133 | 0.034792426 | 0.347924259 |
| Rpl41 | 37.511967 | 0.938206077 | 0.484150186 | 2.640226216 | 0.00828507 | 0.034981408 | 0.349814085 |
| Tmsb4x | 42.59606 | 0.979091562 | 0.528754261 | 2.595740703 | 0.009438727 | 0.038775309 | 0.387753091 |
| Cntnap2 | 56.53396 | -0.144269504 | 0.471400165 | -2.556411985 | 0.010575782 | 0.042303129 | 0.423031288 |
| Meg3 | 25.358404 | -0.95098049 | 0.675196912 | -2.537671449 | 0.011159269 | 0.043492537 | 0.434925367 |
| Rpl30 | 34.068516 | 0.923331074 | 0.53072163 | 2.505167461 | 0.012239343 | 0.046509503 | 0.465095032 |
| Rps28 | 25.47918 | 0.831677198 | 0.466486216 | 2.481505974 | 0.013082852 | 0.04850228 | 0.485022795 |
| Rps23 | 25.459724 | 0.811531577 | 0.459090876 | 2.468754667 | 0.013558414 | 0.049068545 | 0.490685452 |
| Gm10062 | 59.852055 | -0.500146634 | 0.238447827 | -2.447380975 | 0.014389864 | 0.050866498 | 0.508664977 |
| Rpl9 | 27.202162 | 0.950706031 | 0.603869283 | 2.420834647 | 0.015484919 | 0.052710071 | 0.527100706 |
| Selenow | 17.175367 | 0.897834732 | 0.566125162 | 2.418026155 | 0.015604955 | 0.052710071 | 0.527100706 |
| Sec61a1 | 171.56808 | 0.299522177 | 0.131148462 | 2.409660887 | 0.015967354 | 0.052761692 | 0.527616922 |
| Rpl36 | 19.816837 | 0.857646615 | 0.559604287 | 2.367819402 | 0.017893268 | 0.05786759 | 0.578675895 |
| Rps27a | 33.156513 | 0.893813516 | 0.604540317 | 2.341282708 | 0.019217609 | 0.060855762 | 0.608557616 |
| Cmpk1 | 49.48254 | -0.430534899 | 0.214269074 | -2.284494954 | 0.022342459 | 0.067786482 | 0.677864817 |
| Rpl21 | 37.689503 | 0.799199454 | 0.532433725 | 2.28544022 | 0.022287028 | 0.067786482 | 0.677864817 |
| Rpl34 | 22.53576 | 0.716962347 | 0.452022088 | 2.277705187 | 0.022744148 | 0.067786482 | 0.677864817 |
| 4930558K02Rik | 19.8751 | -0.478714883 | 0.244460161 | -2.268510369 | 0.023298118 | 0.06810219 | 0.681021903 |
| Rps15 | 19.44779 | 0.682642259 | 0.44416836 | 2.227606939 | 0.025906738 | 0.074298568 | 0.742985681 |
| Rps3a1 | 29.006617 | 0.738384516 | 0.50680707 | 2.207930272 | 0.027249134 | 0.076701267 | 0.767012666 |
| Mt1 | 35.063984 | 0.68504884 | 0.460390696 | 2.180569293 | 0.029215288 | 0.080740433 | 0.807404335 |
| Rsrc1 | 41.97339 | -0.419865312 | 0.222994929 | -2.164402947 | 0.030433438 | 0.082605046 | 0.826050456 |
| Cox8a | 54.110287 | 0.756080482 | 0.685511711 | 2.083843961 | 0.037174363 | 0.099131635 | 0.99131635 |
| Scamp1 | 17.284771 | -0.455108912 | 0.26078734 | -2.070135187 | 0.038439687 | 0.099581705 | 0.995817045 |
| Atp9b | 22.188927 | -0.525313054 | 0.334766338 | -2.067857501 | 0.038653425 | 0.099581705 | 0.995817045 |
| Gria4 | 27.528088 | -0.386890133 | 0.213145128 | -2.053174007 | 0.040055704 | 0.101474451 | 1.014744511 |
| Tpt1 | 28.210365 | 0.657818662 | 0.52600162 | 2.038923606 | 0.041457652 | 0.103304314 | 1.033043144 |
| Sema3c | 66.135925 | -0.372128431 | 0.211859971 | -2.010198214 | 0.044410214 | 0.108876655 | 1.088766547 |
| Zbtb7a | 23.56369 | -0.429910494 | 0.256382706 | -1.994897083 | 0.046054106 | 0.111114668 | 1.111146678 |
| Camk1d | 70.181915 | -0.54616219 | 0.664707449 | -1.985107629 | 0.047132498 | 0.111939683 | 1.119396833 |
| Rpl37a | 40.960964 | 0.650726603 | 0.58884927 | 1.970672787 | 0.048761315 | 0.11402646 | 1.140264601 |
| AY036118 | 96.25058 | -0.529288282 | 0.640076101 | -1.945657223 | 0.051695928 | 0.11905729 | 1.190572895 |
| Lsamp | 27.76111 | -0.3416601 | 0.201229266 | -1.924167005 | 0.054333654 | 0.12086902 | 1.208690202 |
| Cox6b1 | 15.940308 | 0.619847269 | 0.676979237 | 1.927152177 | 0.053960668 | 0.12086902 | 1.208690202 |
| Xlr5a | 95.355095 | -0.362482682 | 0.219968937 | -1.913661945 | 0.055663365 | 0.12086902 | 1.208690202 |
| Adk | 15.99909 | -0.457323708 | 0.299445511 | -1.916334486 | 0.055322534 | 0.12086902 | 1.208690202 |
| Rpl13a | 23.156572 | 0.584065347 | 0.562282692 | 1.878181732 | 0.060356308 | 0.129213505 | 1.292135051 |
| Nell1 | 23.617365 | -0.343469585 | 0.214337158 | -1.836560651 | 0.066274781 | 0.139913426 | 1.399134261 |
| Rpl39 | 28.428679 | 0.557925983 | 0.596342595 | 1.8195246 | 0.068831433 | 0.141383485 | 1.413834846 |
| Mc1r | 36.4037 | -0.393356274 | 0.262301576 | -1.825363719 | 0.06794616 | 0.141383485 | 1.413834846 |
| Nkain2 | 20.22035 | -0.407846581 | 0.279447003 | -1.808607109 | 0.070512063 | 0.142904448 | 1.429044483 |
| Wdr72 | 23.826147 | -0.404915526 | 0.279929993 | -1.796877501 | 0.072355069 | 0.144710138 | 1.447101385 |
| Cox7c | 24.603357 | 0.512842068 | 0.471724302 | 1.774033975 | 0.076057531 | 0.150139541 | 1.501395411 |
| Dop1b | 23.20367 | -0.397237459 | 0.280285355 | -1.762431939 | 0.077996346 | 0.15157461 | 1.5157461 |
| Fth1 | 159.25063 | 0.529099239 | 0.51035342 | 1.757815488 | 0.078778909 | 0.15157461 | 1.5157461 |
| Nckap5 | 94.96033 | -0.322520871 | 0.215722465 | -1.733381922 | 0.083027809 | 0.157752836 | 1.577528363 |
| Uqcrh | 17.226807 | 0.48491182 | 0.61753923 | 1.710774734 | 0.087122705 | 0.163489521 | 1.634895214 |
| Malat1 | 69.80103 | -0.419803778 | 0.366440549 | -1.679589995 | 0.093037116 | 0.172459044 | 1.724590436 |
| Lrrc17 | 20.887321 | 0.459684457 | 0.437616058 | 1.656716192 | 0.097576868 | 0.178694987 | 1.786949868 |
| Epha6 | 38.557938 | -0.345755051 | 0.259996374 | -1.63387796 | 0.102284478 | 0.185086199 | 1.85086199 |
| Rapgef4 | 48.983097 | -0.303637228 | 0.231885906 | -1.545987345 | 0.122107624 | 0.218357163 | 2.183571632 |
| Prkg1 | 32.408478 | -0.283027377 | 0.211608787 | -1.530344455 | 0.12593149 | 0.222576587 | 2.225765871 |
| Bc1 | 16.326283 | 0.40168989 | 0.465577603 | 1.495258672 | 0.134846947 | 0.235594665 | 2.355946652 |
| Fbxl7 | 16.057865 | -0.352451437 | 0.356950635 | -1.426665379 | 0.153676369 | 0.262639679 | 2.626396788 |

| | | | | | | | |
|---|---|---|---|---|---|---|---|
| Lrp1b | 204.00333 | -0.245707479 | 0.195373594 | -1.426297647 | 0.153782443 | 0.262639679 | 2.626396788 |
| Eef1a1 | 22.886635 | 0.346189002 | 0.4086505 | 1.355733942 | 0.175183836 | 0.295866034 | 2.95866034 |
| Cmss1 | 56.48602 | -0.323521907 | 0.436832086 | -1.334578919 | 0.182014201 | 0.300719115 | 3.007191155 |
| 5830416I19Rik | 109.243774 | -0.249609568 | 0.217718037 | -1.340258512 | 0.180161315 | 0.300719115 | 3.007191155 |
| Rpl11 | 34.706364 | 0.330163253 | 0.486327679 | 1.305951605 | 0.191569016 | 0.313102047 | 3.131020474 |
| Slx1b | 25.369318 | -0.267319455 | 0.253961125 | -1.282922739 | 0.199519146 | 0.319230633 | 3.19230633 |
| Lrrc4c | 18.606462 | -0.313766778 | 0.413268527 | -1.284052207 | 0.199123689 | 0.319230633 | 3.19230633 |
| Mapk14 | 41.72899 | -0.212832843 | 0.191631995 | -1.24485731 | 0.213184205 | 0.330653053 | 3.306530534 |
| Ms4a1 | 27.41386 | -0.266993374 | 0.267573531 | -1.2472639 | 0.21230075 | 0.330653053 | 3.306530534 |
| 8430422H06Rik | 42.030968 | -0.261935979 | 0.25854107 | -1.255165569 | 0.209418662 | 0.330653053 | 3.306530534 |
| Igf2bp3 | 45.83929 | -0.254859451 | 0.285604297 | -1.1615915 | 0.245401436 | 0.376777963 | 3.76777963 |
| Gm42418 | 495.07285 | -0.273048324 | 0.382779041 | -1.149878678 | 0.250193844 | 0.380294643 | 3.802946427 |
| Kctd16 | 76.70161 | -0.157525832 | 0.155930158 | -1.094773786 | 0.273615754 | 0.411778164 | 4.117781638 |
| Pde6a | 63.212955 | -0.18201959 | 0.1982004 | -1.043698824 | 0.296624754 | 0.442029046 | 4.420290456 |
| Il27 | 31.86366 | -0.202385938 | 0.231121398 | -1.036256042 | 0.300082708 | 0.442840501 | 4.428405006 |
| Rps29 | 86.63813 | 0.233242995 | 0.404466944 | 0.970419941 | 0.331837213 | 0.48499285 | 4.849928496 |
| Pcdh15 | 27.633442 | -0.144269504 | 0.471400313 | -0.922723252 | 0.356151446 | 0.515571616 | 5.155716163 |
| Prkag2 | 18.591381 | -0.190032669 | 0.260317512 | -0.905928307 | 0.364973801 | 0.523358657 | 5.233586574 |
| mt-Atp6 | 255.12491 | -0.201566541 | 0.345882835 | -0.883165552 | 0.377146831 | 0.535759984 | 5.357599836 |
| 6030443J06Rik | 30.79876 | -0.161366468 | 0.219446562 | -0.858464538 | 0.390636004 | 0.549784005 | 5.497840053 |
| Gm38973 | 16.28398 | 0.191466476 | 0.326608249 | 0.832417102 | 0.405173552 | 0.56501266 | 5.650126596 |
| Ckb | 26.128765 | 0.192387129 | 0.382869287 | 0.821339629 | 0.411452841 | 0.568553016 | 5.68553016 |
| Kcnip4 | 17.83586 | -0.177384764 | 0.288012575 | -0.811286945 | 0.417200905 | 0.571302141 | 5.713021408 |
| D430042O09Rik | 59.267593 | -0.116957833 | 0.164801013 | -0.778424307 | 0.436318914 | 0.592147097 | 5.921470969 |
| mt-Nd5 | 23.92398 | -0.164373749 | 0.414059522 | -0.756135582 | 0.44956791 | 0.604728517 | 6.047285166 |
| Wdyhv1 | 16.539324 | -0.168367931 | 0.335036111 | -0.737837096 | 0.460613448 | 0.614151264 | 6.14151264 |
| 1700054A03Rik | 141.57812 | 0.124013995 | 0.190961827 | 0.728119797 | 0.466540256 | 0.616644513 | 6.166445126 |
| Adamtsl1 | 32.24669 | -0.13611542 | 0.223981141 | -0.714515152 | 0.47490869 | 0.617960816 | 6.17960816 |
| Ctnna3 | 24.259981 | 0.145369894 | 0.250069865 | 0.713288573 | 0.475667207 | 0.617960816 | 6.17960816 |
| mt-Cytb | 99.99927 | -0.158418041 | 0.362448369 | -0.703390205 | 0.481812609 | 0.618158388 | 6.181583882 |
| Camk2a | 20.71487 | -0.145774698 | 0.26051503 | -0.699958938 | 0.483952949 | 0.618158388 | 6.181583882 |
| 4930511M06Rik | 35.85655 | -0.130360109 | 0.233249529 | -0.668502586 | 0.503812832 | 0.63816292 | 6.3816292 |
| Chsy3 | 23.052742 | -0.120304872 | 0.21425147 | -0.65055024 | 0.515336861 | 0.647365313 | 6.473653126 |
| mt-Nd1 | 66.879105 | -0.144047857 | 0.332930262 | -0.643701728 | 0.519768869 | 0.647580886 | 6.475808858 |
| mt-Nd4 | 90.86455 | -0.134897727 | 0.366468745 | -0.605061849 | 0.545137902 | 0.673666351 | 6.736663508 |
| Ppia | 17.00175 | 0.110464105 | 0.398817055 | 0.502322356 | 0.615440786 | 0.742436503 | 7.424365033 |
| Zmat4 | 23.312378 | -0.0996604 | 0.237142738 | -0.505769858 | 0.613018221 | 0.742436503 | 7.424365033 |
| Pde1c | 48.006958 | -0.108921914 | 0.279722779 | -0.511844251 | 0.608760017 | 0.742436503 | 7.424365033 |
| Ptpn9 | 16.17321 | 0.086680718 | 0.265809612 | 0.413629593 | 0.679145397 | 0.812835436 | 8.128354363 |
| Gas5 | 23.922663 | -0.076177499 | 0.43514323 | -0.401551617 | 0.688014043 | 0.817016676 | 8.170166763 |
| Gm12216 | 92.376305 | -0.064373536 | 0.193389428 | -0.377249147 | 0.70598846 | 0.831862371 | 8.318623715 |
| Rab29 | 17.81273 | 0.071985648 | 0.242828936 | 0.356745714 | 0.721282179 | 0.843345317 | 8.433453173 |
| mt-Co3 | 317.89905 | -0.073703977 | 0.330340951 | -0.333983509 | 0.738392003 | 0.852321207 | 8.523212066 |
| Ttc28 | 27.770508 | -0.064196961 | 0.231146865 | -0.331623358 | 0.740173679 | 0.852321207 | 8.523212066 |
| Cpe | 18.960005 | 0.067154706 | 0.394995065 | 0.309886427 | 0.756647325 | 0.858286518 | 8.582865181 |
| Rps27 | 24.597618 | 0.070037752 | 0.333381987 | 0.31333924 | 0.75402294 | 0.858286518 | 8.582865181 |
| Ninj2 | 17.927513 | -0.05683817 | 0.243655315 | -0.283083896 | 0.777112526 | 0.874971141 | 8.749711408 |
| Ehbp1 | 37.233177 | -0.049379749 | 0.21453733 | -0.268227784 | 0.78852399 | 0.881291518 | 8.812915183 |
| Fyb2 | 22.179024 | -0.048900631 | 0.248768313 | -0.247590975 | 0.804450895 | 0.89252946 | 8.925294597 |
| mt-Nd2 | 100.98584 | -0.038219281 | 0.310946337 | -0.174398045 | 0.861552651 | 0.895863725 | 8.958637248 |
| mt-Co1 | 163.52351 | -0.041180337 | 0.345819393 | -0.187693449 | 0.851116966 | 0.895863725 | 8.958637248 |
| mt-Co2 | 179.1176 | -0.042420786 | 0.311987106 | -0.192896509 | 0.847040019 | 0.895863725 | 8.958637248 |
| Gnaz | 44.17836 | -0.039319953 | 0.254955607 | -0.193227622 | 0.846780707 | 0.895863725 | 8.958637248 |
| Mbp | 64.762535 | -0.03944711 | 0.434603204 | -0.214328085 | 0.830291218 | 0.895863725 | 8.958637248 |
| Cdh23 | 31.393366 | -0.032937462 | 0.186464648 | -0.197801696 | 0.843200219 | 0.895863725 | 8.958637248 |
| Ctcflos | 16.329626 | -0.037145233 | 0.342714587 | -0.168239937 | 0.866394523 | 0.895863725 | 8.958637248 |
| Shb | 57.12811 | -0.034674262 | 0.187087426 | -0.207866275 | 0.835333382 | 0.895863725 | 8.958637248 |
| Fam169a | 27.947054 | 0.042088385 | 0.249547733 | 0.206747708 | 0.836206897 | 0.895863725 | 8.958637248 |
| Calm1 | 23.933855 | -0.037179617 | 0.382541539 | -0.17444318 | 0.861517182 | 0.895863725 | 8.958637248 |
| Ankrd12 | 21.009596 | 0.029307708 | 0.271498784 | 0.138864318 | 0.889557368 | 0.913599459 | 9.135994592 |
| Ppp1r9a | 15.939527 | 0.015958868 | 0.290379244 | 0.072773726 | 0.941986179 | 0.960952344 | 9.609523436 |
| Csmd3 | 26.341698 | 0.00876747 | 0.248349011 | 0.04333168 | 0.965437138 | 0.978309633 | 9.783096327 |
| mt-Nd3 | 53.05994 | 0.004745362 | 0.336688532 | 0.020820317 | 0.98338899 | 0.9899015 | 9.899015002 |
| 1700126G02Rik | 34.365047 | 0.000199725 | 0.197777465 | -0.005574329 | 0.995552352 | 0.995552352 | 9.955523521 |
| 4930402H24Rik | 10.878186 | -0.144269504 | 0.471400323 | -0.897202009 | 0.369611134 | | |
| 4930441J16Rik | 14.087247 | -0.319809897 | 0.293431402 | -1.397566125 | 0.162243397 | | |
| AI314278 | 10.483916 | 0.193812031 | 0.30343342 | 0.866128756 | 0.386419557 | | |
| Ahdc1 | 11.812328 | -0.028112985 | 0.286769023 | -0.130610909 | 0.896083112 | | |
| Aldoc | 15.485122 | 0.957114771 | 0.54294827 | 2.56603686 | 0.01028679 | | |
| Alpl | 10.492561 | -0.186688967 | 0.327314118 | -0.814224023 | 0.415516626 | | |
| Anks1b | 9.166362 | -0.038649372 | 0.3588414 | -0.177224544 | 0.859332017 | | |
| Arvcf | 7.0802913 | -0.144269504 | 0.47140034 | 1.33537944 | 0.18175219 | | |
| Atxn7l1 | 12.299738 | -0.356266815 | 0.333802519 | -1.45818805 | 0.144788717 | | |
| Cadm2 | 11.891537 | -0.10276444 | 0.36966792 | -0.465416706 | 0.641633073 | | |
| Car12 | 13.269348 | 0.221472991 | 0.486063697 | 1.011744797 | 0.311660091 | | |
| Ccdc60 | 15.782949 | -0.48991017 | 0.285650355 | -2.084168121 | 0.037144877 | | |
| Cep128 | 10.783486 | -0.115477868 | 0.299170455 | -0.523519099 | 0.600613064 | | |
| Cfap74 | 13.209162 | 1.08517473 | 0.340496702 | 3.713096103 | 0.000204739 | | |
| Chl1 | 13.870078 | -0.014945714 | 0.276530227 | -0.071176167 | 0.943257549 | | |
| Cntn3 | 12.88803 | -0.032471034 | 0.269499143 | -0.154342943 | 0.877339339 | | |

| | | | | | |
|---|---|---|---|---|---|
| Cspp1 | 15.578925 | 0.023188508 | 0.295129754 | 0.105598252 | 0.915901112 |
| Ctnna2 | 13.414483 | -0.483125348 | 0.353906646 | -1.851786584 | 0.064056475 |
| Dctd | 7.827331 | -0.2211418 | 0.352101396 | -0.935993438 | 0.349276566 |
| Dlg2 | 14.809554 | -0.336273164 | 0.527369311 | -1.473425785 | 0.140636262 |
| Dpyd | 15.7993355 | -0.425094077 | 0.300819092 | -1.790502185 | 0.073373216 |
| Esf1 | 9.794302 | -0.227122679 | 0.332129992 | -0.96908491 | 0.332502826 |
| Gm11337 | 13.186041 | -0.351411022 | 0.33343846 | -1.440305976 | 0.149780851 |
| Gm14582 | 12.835498 | -0.323846373 | 0.327820322 | -1.352846067 | 0.176104812 |
| Gm29325 | 14.84829 | 0.048349108 | 0.272657934 | 0.227432781 | 0.820087233 |
| Gm49497 | 13.54824 | -0.353159226 | 0.348710249 | -1.430322577 | 0.152624458 |
| Gm49767 | 11.525636 | 0.299640703 | 0.308657992 | 1.304135683 | 0.192187325 |
| Gm765 | 13.4760685 | -0.144269504 | 0.471400333 | -1.254229375 | 0.209758646 |
| Gm960 | 8.686207 | -0.144269504 | 0.47140034 | 0.111884828 | 0.910914726 |
| Gnao1 | 10.703454 | -0.134258826 | 0.326092681 | -0.594263929 | 0.5523356 |
| Gphn | 15.688767 | -0.273381372 | 0.465022288 | -1.204643648 | 0.228340899 |
| Gria1 | 10.882296 | 0.085771158 | 0.294409399 | 0.391555225 | 0.695386877 |
| Grik1 | 5.4989185 | -0.02157633 | 0.399064883 | -0.103773811 | 0.91734885 |
| Hapln1 | 12.383218 | -0.322619187 | 0.362766199 | -1.307305191 | 0.191109082 |
| Hdac8 | 13.568848 | -0.424850217 | 0.328933771 | -1.711577472 | 0.086974563 |
| Hdac9 | 9.57025 | -0.144269504 | 0.471400331 | -1.389159562 | 0.164784235 |
| Hexb | 15.088464 | 0.028952826 | 0.439054137 | 0.150910664 | 0.880046188 |
| Il1rapl2 | 9.701231 | -0.130217302 | 0.312567199 | -0.580762205 | 0.561400731 |
| Kat7 | 14.177729 | -0.02971243 | 0.23438756 | -0.159762971 | 0.873067794 |
| Kcnd2 | 8.189679 | -0.331902477 | 0.379881967 | -1.31330904 | 0.189078834 |
| Kcnq5 | 13.298319 | -0.247206244 | 0.313027144 | -1.069500319 | 0.284844285 |
| Klhl14 | 2.6490629 | -1.086343565 | 1.004610199 | -2.093300996 | 0.036322294 |
| Klkb1 | 12.379919 | -0.142619026 | 0.304562078 | -0.639153945 | 0.522722789 |
| Lyrm1 | 7.9197645 | -0.144269504 | 0.471400336 | -1.119344218 | 0.262993319 |
| Mrc2 | 15.52196 | 0.065190366 | 0.296772858 | 0.296748763 | 0.766658325 |
| Pcca | 11.302964 | 0.225275059 | 0.320580771 | 0.976737363 | 0.328699193 |
| Pcgf6 | 13.125739 | -0.144269504 | 0.471400332 | -0.473493779 | 0.635860938 |
| Phkg1 | 13.941239 | 0.08425616 | 0.286556065 | 0.387585884 | 0.698322517 |
| Plp1 | 11.947595 | 0.756112754 | 0.694148199 | 2.11901619 | 0.034089099 |
| Ppp1r16a | 14.606436 | -0.031012429 | 0.230739681 | -0.170764803 | 0.864408706 |
| Ppp3ca | 11.9548855 | -0.219792519 | 0.417015242 | -0.964936608 | 0.334576564 |
| Ptprd | 14.081265 | -0.203615771 | 0.315554819 | -0.891407247 | 0.372710731 |
| Ptprg | 9.615968 | -0.068397235 | 0.301128041 | -0.311524876 | 0.755401635 |
| Ralyl | 14.507205 | -0.604461841 | 0.462599823 | -2.044440829 | 0.040910017 |
| Rpl12 | 12.270738 | 1.272597042 | 0.531098799 | 3.120336522 | 0.001806445 |
| Rps17 | 12.002829 | 0.24806296 | 0.417655033 | 1.041566058 | 0.297612912 |
| Sgip1 | 6.526274 | -0.082698991 | 0.357774227 | -0.371133466 | 0.710538126 |
| Spock1 | 12.781663 | -0.095012551 | 0.287792577 | -0.437067542 | 0.662062366 |
| Srgn | 6.927038 | -0.144269504 | 0.471400336 | -0.964761958 | 0.334664054 |
| St18 | 5.592503 | 0.174415593 | 0.405533996 | 0.763534871 | 0.445144474 |
| Sult2a2 | 10.396637 | -0.213375019 | 0.313411926 | -0.930177007 | 0.352279445 |
| Svil | 12.132911 | -0.336355536 | 0.351073588 | -1.361970922 | 0.173207062 |
| Syt1 | 12.191012 | -0.33292979 | 0.466681891 | -1.366251251 | 0.171860116 |
| Tacc1 | 11.41664 | 0.275481578 | 0.293558574 | 1.234361305 | 0.217068301 |
| Tmeff2 | 11.866141 | -0.145968032 | 0.256374828 | -0.727131345 | 0.467145499 |
| Tns3 | 11.983182 | -0.489137485 | 0.351818192 | -1.869944738 | 0.061491492 |
| Tshz2 | 4.0018873 | -1.700220181 | 0.876346344 | -2.675977576 | 0.007451163 |
| Uba52 | 9.810962 | -0.009344602 | 0.404870867 | -0.046735009 | 0.962724428 |
| Ubb | 13.352551 | 0.309454593 | 0.435202235 | 1.242000286 | 0.214236457 |
| Unc5d | 1.6227521 | -0.144269504 | 0.471400343 | -0.888902692 | 0.374055379 |
| Vit | 14.305894 | -0.117942806 | 0.272538183 | -0.551421375 | 0.581344852 |
| Vwc2l | 9.199624 | -0.232529064 | 0.2866929 | -1.08372527 | 0.278486628 |
| Zfhx3 | 14.494242 | -0.218687104 | 0.274049801 | -1.006115619 | 0.314359956 |
| Zfp994 | 14.944137 | 0.09522881 | 0.270373538 | 0.451247843 | 0.651810931 |
| mt-Nd4l | 10.59298 | -0.008964983 | 0.387441333 | -0.043246843 | 0.965504765 |

**table S8: Differential gene expression list elevated platform compared to rest (3) conditions in GABA and Glut neurons**

| **GABA** | baseMean | log2FoldChange | lfcSE | stat | pvalue | padj | pglobadj |
|---|---|---|---|---|---|---|---|
| mt-Nd1 | 13.160506 | 1.757960472 | 0.339405196 | 5.654674494 | 1.56E-08 | 3.54E-06 | 3.54E-05 |
| Gm42418 | 88.84172 | 2.07301591 | 0.428931831 | 5.202024691 | 1.97E-07 | 2.24E-05 | 0.000223741 |
| Hexb | 3.3334806 | 2.504139466 | 0.670926981 | 4.37713659 | 1.20E-05 | 0.000682411 | 0.006824109 |
| mt-Nd4 | 15.204892 | 1.357991156 | 0.347811362 | 4.383592645 | 1.17E-05 | 0.000682411 | 0.006824109 |
| Camk1d | 9.3028755 | 1.820358355 | 0.487776667 | 4.321371626 | 1.55E-05 | 0.000703983 | 0.007039826 |
| Ckb | 3.2405775 | 2.544086873 | 0.734811361 | 4.172143604 | 3.02E-05 | 0.001141611 | 0.011416108 |
| mt-Atp6 | 52.440006 | 1.145434974 | 0.330814014 | 3.902924382 | 9.50E-05 | 0.003081927 | 0.030819272 |
| Calm1 | 3.654584 | 2.813077531 | 0.887777663 | 3.747526447 | 0.000178587 | 0.00479683 | 0.047968303 |
| mt-Co3 | 64.31273 | 0.935464776 | 0.281072244 | 3.731712214 | 0.000190183 | 0.00479683 | 0.047968303 |
| mt-Nd3 | 11.394203 | 1.336006373 | 0.448133013 | 3.629072771 | 0.000284441 | 0.006456812 | 0.06456812 |
| Kctd16 | 20.133091 | -0.144269504 | 0.5137751 | -3.579508384 | 0.000344241 | 0.007103887 | 0.071038867 |
| mt-Nd2 | 16.228426 | 1.347944254 | 0.46076916 | 3.544566261 | 0.00039326 | 0.007439162 | 0.074391616 |
| AY036118 | 14.886724 | 1.321753311 | 0.496781041 | 3.294671363 | 0.000985368 | 0.017206049 | 0.172060489 |
| mt-Co1 | 29.908287 | 1.076164752 | 0.404180811 | 3.210754403 | 0.00132387 | 0.021465611 | 0.214656108 |
| Gphn | 2.9946876 | 1.704865959 | 0.798538689 | 3.140342677 | 0.001687503 | 0.025537548 | 0.255375479 |
| mt-Co2 | 31.080164 | 0.866484255 | 0.349037935 | 2.97713914 | 0.002909519 | 0.039749729 | 0.397497292 |
| Cox6b1 | 3.6377096 | 1.236927016 | 0.624937052 | 2.970118188 | 0.002976852 | 0.039749729 | 0.397497292 |
| Rps20 | 2.5459812 | 2.109196446 | 1.104120675 | 2.934697666 | 0.003338728 | 0.039889016 | 0.398890162 |
| Cox7c | 4.358693 | 1.703888952 | 0.849566216 | 2.946956877 | 0.00320918 | 0.039889016 | 0.398890162 |
| Rpl23 | 3.8133543 | 2.078712132 | 1.110233262 | 2.874295159 | 0.004049303 | 0.045959594 | 0.45959594 |
| Cpe | 2.8087764 | 1.109962183 | 0.748030847 | 2.63335718 | 0.008454538 | 0.091389535 | 0.913895346 |
| mt-Cytb | 17.050423 | 0.74061086 | 0.382952989 | 2.4881002 | 0.012842754 | 0.132513875 | 1.325138747 |
| Rpl27a | 4.768087 | 0.996699326 | 0.73432565 | 2.397989775 | 0.016485325 | 0.162702995 | 1.627029945 |
| Mbp | 3.1925125 | 1.02909517 | 1.33146421 | 2.302545747 | 0.021304415 | 0.201504256 | 2.015042557 |
| Epha6 | 12.959585 | 0.631988792 | 0.381946547 | 2.196140742 | 0.028081871 | 0.205989569 | 2.05989569 |
| Igf2bp3 | 12.022297 | -0.779436522 | 0.458548039 | -2.207215382 | 0.027299016 | 0.205989569 | 2.05989569 |
| Cox4i1 | 4.267222 | 0.899511476 | 0.893713203 | 2.221342214 | 0.026327792 | 0.205989569 | 2.05989569 |
| Cntnap2 | 15.640811 | 0.558976077 | 0.319864325 | 2.195458369 | 0.028130734 | 0.205989569 | 2.05989569 |
| 4930511M | 7.9351397 | -1.034579713 | 0.711174618 | -2.198456041 | 0.027916624 | 0.205989569 | 2.05989569 |
| D430042O | 15.700211 | -0.805761193 | 0.517785506 | -2.207849628 | 0.027254757 | 0.205989569 | 2.05989569 |
| Tpt1 | 4.37262 | 0.931082722 | 1.440915663 | 2.255637189 | 0.024093362 | 0.205989569 | 2.05989569 |
| Plp1 | 2.3849928 | 0.755616512 | 1.253923351 | 2.165287515 | 0.030365674 | 0.215406499 | 2.154064991 |
| Tmeff2 | 3.7606382 | 0.756670883 | 0.800567185 | 2.108716327 | 0.034969075 | 0.238746306 | 2.38746306 |
| Rps29 | 11.573063 | 0.842852747 | 0.849604001 | 2.087852917 | 0.036811104 | 0.238746306 | 2.38746306 |
| Cdk8 | 4.057161 | 0.762718042 | 0.786512417 | 2.092003911 | 0.036438166 | 0.238746306 | 2.38746306 |
| Lrp1b | 56.844112 | -0.436221791 | 0.236071581 | -2.0738302 | 0.038095083 | 0.240210665 | 2.402106648 |
| Gria4 | 7.6431046 | -0.853270992 | 0.767993933 | -1.990426265 | 0.046543999 | 0.285553724 | 2.855537238 |
| Uba52 | 1.9512028 | 0.53528106 | 0.81065492 | 1.953737595 | 0.050732258 | 0.303058487 | 3.030584871 |
| AI314278 | 3.58537 | 0.578883482 | 0.6721646 | 1.855068027 | 0.063586513 | 0.366402637 | 3.664026372 |
| Fyb2 | 4.7031064 | -0.858099244 | 0.837249578 | -1.848262752 | 0.064564341 | 0.366402637 | 3.664026372 |
| Rpl13a | 1.4650565 | 0.432207358 | 0.706183223 | 1.804111058 | 0.071213899 | 0.394281828 | 3.942818284 |
| Rpl38 | 7.104176 | 0.574255458 | 0.656095361 | 1.771595446 | 0.076461736 | 0.413257479 | 4.132574793 |
| Fau | 2.4518764 | 0.445432233 | 0.766537901 | 1.73390501 | 0.082934939 | 0.427868889 | 4.278688891 |
| Rpl37 | 5.0523815 | 0.517130894 | 0.709154147 | 1.742508965 | 0.081419424 | 0.427868889 | 4.278688891 |
| Mt3 | 4.3413577 | 0.494907878 | 0.714989635 | 1.714233758 | 0.086485803 | 0.433927406 | 4.33927406 |
| Nckap5 | 23.137186 | -0.455535076 | 0.332029008 | -1.706406475 | 0.087932426 | 0.433927406 | 4.33927406 |
| mt-Nd5 | 4.970977 | 0.476917841 | 0.63007114 | 1.640232922 | 0.100956746 | 0.458343629 | 4.58343629 |
| Nell1 | 5.491928 | -0.591145597 | 0.57620906 | -1.640712777 | 0.100857051 | 0.458343629 | 4.58343629 |
| Gm10062 | 13.044163 | -0.508311669 | 0.437565912 | -1.642335355 | 0.100520524 | 0.458343629 | 4.58343629 |
| Ccdc60 | 3.9716513 | -0.144269504 | 0.513775112 | -1.648880659 | 0.099172086 | 0.458343629 | 4.58343629 |
| mt-Nd4l | 1.7701625 | 0.292249853 | 0.744528968 | 1.610282612 | 0.107336174 | 0.477751204 | 4.777512037 |
| Malat1 | 6.5285845 | 0.425001993 | 0.722012087 | 1.590369246 | 0.111751598 | 0.482054575 | 4.820545753 |
| Srgn | 2.2238126 | 0.422352879 | 0.659920906 | 1.58683426 | 0.112550187 | 0.482054575 | 4.820545753 |
| Ppia | 1.9710991 | 0.316731445 | 0.669091382 | 1.513175333 | 0.130235128 | 0.525769953 | 5.257699526 |
| Rpl12 | 2.0700948 | 0.352004501 | 0.634794722 | 1.506177827 | 0.13202153 | 0.525769953 | 5.257699526 |
| Rpl17 | 2.507469 | 0.349336458 | 0.665004902 | 1.514765511 | 0.129831798 | 0.525769953 | 5.257699526 |
| Uqcrh | 1.8911818 | 0.355524859 | 0.65378753 | 1.527655584 | 0.126598073 | 0.525769953 | 5.257699526 |
| Cep128 | 2.7124422 | -0.144269504 | 0.513775113 | -1.486663945 | 0.13710359 | 0.536595084 | 5.365950844 |
| Rab29 | 4.6720915 | -0.441527976 | 0.618000443 | -1.450319901 | 0.146969332 | 0.565458277 | 5.654582771 |

| | | | | | | | |
|---|---|---|---|---|---|---|---|
| Ttc28 | 5.5357037 | -0.436191853 | 0.576024 | -1.420552111 | 0.155447009 | 0.572480086 | 5.724800863 |
| Rps3a1 | 6.1928663 | 0.280924669 | 0.743266939 | 1.412118537 | 0.157915061 | 0.572480086 | 5.724800863 |
| Slx1b | 6.8200607 | -0.144269504 | 0.513775113 | -1.413605 | 0.157477913 | 0.572480086 | 5.724800863 |
| Pcdh15 | 10.199923 | 0.380358348 | 0.398876111 | 1.400364894 | 0.161404077 | 0.572480086 | 5.724800863 |
| Rps14 | 3.1900764 | 0.263695113 | 0.73723434 | 1.408118686 | 0.159095929 | 0.572480086 | 5.724800863 |
| Fam169a | 6.7369084 | -0.419458833 | 0.485964967 | -1.361074238 | 0.17349023 | 0.591406703 | 5.914067027 |
| Rps17 | 1.9367694 | 0.263225142 | 0.647670782 | 1.349546549 | 0.177161479 | 0.591406703 | 5.914067027 |
| Xlr5a | 27.32727 | -0.32885369 | 0.289880075 | -1.363438386 | 0.17274439 | 0.591406703 | 5.914067027 |
| Adk | 3.7155418 | -0.144269504 | 0.513775115 | 1.351246109 | 0.176616609 | 0.591406703 | 5.914067027 |
| Mapk14 | 11.374496 | -0.144269504 | 0.513775111 | -1.324952285 | 0.185186978 | 0.598890374 | 5.988903743 |
| Rpl9 | 3.0056663 | 0.288906241 | 0.635749484 | 1.325790273 | 0.184909176 | 0.598890374 | 5.988903743 |
| Gm38973 | 4.666378 | 0.361336523 | 0.504518477 | 1.318554377 | 0.187318135 | 0.598890374 | 5.988903743 |
| Kcnd2 | 2.514149 | 0.322079856 | 0.578396231 | 1.295908605 | 0.195006974 | 0.609973806 | 6.099738062 |
| Rps7 | 1.746049 | 0.229516237 | 0.655030777 | 1.274362867 | 0.202534846 | 0.609973806 | 6.099738062 |
| Car12 | 3.4988654 | 0.286393879 | 0.629268693 | 1.288277753 | 0.197649294 | 0.609973806 | 6.099738062 |
| Camk2a | 5.2381573 | -0.387247366 | 0.521302203 | -1.283398646 | 0.199352448 | 0.609973806 | 6.099738062 |
| Cspp1 | 3.2586973 | -0.359887192 | 0.56819087 | -1.269619205 | 0.204220305 | 0.609973806 | 6.099738062 |
| 4930441J1 | 3.6492968 | -0.307913061 | 0.569194761 | -1.221982775 | 0.221714141 | 0.653624805 | 6.536248051 |
| Rps27 | 2.3981893 | 0.271376219 | 0.567507729 | 1.200492386 | 0.229948168 | 0.66920813 | 6.692081298 |
| Rps10 | 1.7781105 | -0.302338098 | 0.629997453 | -1.130270804 | 0.258362133 | 0.733102553 | 7.331025525 |
| 5830416I1 | 24.880907 | -0.273604191 | 0.293614339 | -1.130506426 | 0.258262892 | 0.733102553 | 7.331025525 |
| Dop1b | 4.380478 | -0.309176347 | 0.507451882 | -1.110276677 | 0.26687982 | 0.747922459 | 7.479224586 |
| Rpl26 | 6.793167 | 0.256029528 | 0.542060354 | 1.035453593 | 0.300457128 | 0.748896174 | 7.488961744 |
| Sil1 | 7.6066775 | -0.289542916 | 0.449791405 | -1.038921111 | 0.298841427 | 0.748896174 | 7.488961744 |
| Mc1r | 10.825235 | -0.270603714 | 0.399923137 | -0.980962759 | 0.326611107 | 0.748896174 | 7.488961744 |
| Trim2 | 9.635662 | 0.249565819 | 0.344830262 | 1.005618474 | 0.314599134 | 0.748896174 | 7.488961744 |
| Syt1 | 2.76881 | -0.139480055 | 0.456577227 | -1.001974444 | 0.316355936 | 0.748896174 | 7.488961744 |
| Shb | 15.204553 | -0.144269504 | 0.513775107 | -1.036643321 | 0.299902116 | 0.748896174 | 7.488961744 |
| Arvcf | 1.1981798 | -0.299659217 | 0.616552695 | -1.043619738 | 0.296661357 | 0.748896174 | 7.488961744 |
| Spock1 | 3.860509 | 0.258432311 | 0.493452038 | 1.027188395 | 0.304331762 | 0.748896174 | 7.488961744 |
| Cst3 | 20.57817 | 0.270409795 | 0.463194516 | 0.999514313 | 0.317545609 | 0.748896174 | 7.488961744 |
| Galnt13 | 6.1900806 | -0.308053614 | 0.468169048 | -1.089585151 | 0.275895927 | 0.748896174 | 7.488961744 |
| Gm765 | 2.7776783 | -0.279885411 | 0.531347488 | -1.081468277 | 0.279488865 | 0.748896174 | 7.488961744 |
| Rpl34 | 1.8285625 | 0.186096091 | 0.590591224 | 1.044781551 | 0.296123944 | 0.748896174 | 7.488961744 |
| Cadm2 | 1.128733 | -0.246304529 | 0.588776212 | -0.989553651 | 0.322392334 | 0.748896174 | 7.488961744 |
| Acoxl | 4.884706 | -0.287297344 | 0.455867823 | -1.014618703 | 0.31028762 | 0.748896174 | 7.488961744 |
| Rpl32 | 3.3037553 | 0.218117194 | 0.593388106 | 1.065086641 | 0.286836724 | 0.748896174 | 7.488961744 |
| Prkg1 | 7.6053395 | -0.272711273 | 0.440796081 | -0.984559637 | 0.324840424 | 0.748896174 | 7.488961744 |
| Rps3 | 1.8366036 | 0.169533342 | 0.597726152 | 1.006398076 | 0.314224119 | 0.748896174 | 7.488961744 |
| Rpl21 | 3.447512 | 0.239347992 | 0.58603526 | 1.082521108 | 0.279021042 | 0.748896174 | 7.488961744 |
| Mt1 | 3.035079 | 0.202219101 | 0.562045242 | 0.973213328 | 0.330447288 | 0.750115344 | 7.501153437 |
| Cdh23 | 7.7059603 | -0.144269504 | 0.513775105 | 0.962258838 | 0.3359196 | 0.754987616 | 7.54987616 |
| Rpl30 | 4.705174 | -0.158249034 | 0.542725513 | -0.913253258 | 0.361109359 | 0.758998376 | 7.589983757 |
| Gm49497 | 2.7023165 | -0.215368448 | 0.518089711 | -0.925063814 | 0.354932714 | 0.758998376 | 7.589983757 |
| Cmss1 | 5.8559604 | -0.144269504 | 0.513775113 | 0.93717367 | 0.348669231 | 0.758998376 | 7.589983757 |
| Lrrc4c | 2.4291487 | -0.21399901 | 0.517974838 | -0.924053602 | 0.355458408 | 0.758998376 | 7.589983757 |
| Adamtsl1 | 11.004572 | -0.247245167 | 0.391638002 | -0.915147859 | 0.360114013 | 0.758998376 | 7.589983757 |
| Il1rapl2 | 2.7269056 | 0.207101888 | 0.525840894 | 0.933237675 | 0.350697267 | 0.758998376 | 7.589983757 |
| Kat7 | 4.219699 | -0.255395043 | 0.473255009 | -0.943764579 | 0.345289965 | 0.758998376 | 7.589983757 |
| Rps21 | 6.852249 | 0.213967735 | 0.516286079 | 0.885085002 | 0.376110788 | 0.783276595 | 7.832765954 |
| Rpl37a | 4.900374 | 0.188057432 | 0.533458888 | 0.856427775 | 0.391761207 | 0.794016017 | 7.940160173 |
| 1700054A( | 52.824097 | -0.216920253 | 0.32737894 | -0.866461793 | 0.38623697 | 0.794016017 | 7.940160173 |
| 4930558K( | 5.919937 | 0.221790959 | 0.42063169 | 0.858916983 | 0.390386319 | 0.794016017 | 7.940160173 |
| Ms4a1 | 6.423043 | -0.144269504 | 0.513775112 | -0.833600893 | 0.404505923 | 0.794439339 | 7.944393388 |
| Tmem108 | 3.9934008 | -0.144269504 | 0.513775115 | -0.832741841 | 0.404990341 | 0.794439339 | 7.944393388 |
| Tshz2 | 10.283613 | 0.214520695 | 0.345504025 | 0.848101994 | 0.396381171 | 0.794439339 | 7.944393388 |
| Mrc2 | 6.650044 | -0.216807851 | 0.457846149 | -0.831008192 | 0.405969001 | 0.794439339 | 7.944393388 |
| Klkb1 | 2.2612617 | -0.179551463 | 0.503237492 | -0.796179043 | 0.425927976 | 0.800767664 | 8.007676641 |
| Sema3c | 24.274586 | -0.144269504 | 0.513775097 | -0.788561186 | 0.430368524 | 0.800767664 | 8.007676641 |
| Scamp1 | 4.1156707 | -0.202253648 | 0.467135116 | -0.790626996 | 0.429161689 | 0.800767664 | 8.007676641 |
| Ptprd | 2.9578784 | 0.174937429 | 0.522063047 | 0.79648131 | 0.425752335 | 0.800767664 | 8.007676641 |
| Sgip1 | 1.58918 | 0.145855951 | 0.531885967 | 0.795039214 | 0.426590692 | 0.800767664 | 8.007676641 |

| | | | | | | | |
|---|---|---|---|---|---|---|---|
| Rps12 | 3.9755926 | -0.116204684 | 0.542143734 | -0.802640695 | 0.42218244 | 0.800767664 | 8.007676641 |
| Gria1 | 1.568388 | -0.166344605 | 0.518784744 | -0.772724437 | 0.439685481 | 0.811452067 | 8.11452067 |
| Vwc2l | 4.158543 | 0.189312624 | 0.443712111 | 0.763307375 | 0.445280105 | 0.815149869 | 8.15149869 |
| 6030443J0 | 7.6030917 | -0.193493617 | 0.404115025 | -0.726140026 | 0.467752934 | 0.828619064 | 8.286190636 |
| Rps8 | 4.412556 | 0.138380025 | 0.566644471 | 0.731208537 | 0.464651786 | 0.828619064 | 8.286190636 |
| Ctnna3 | 7.525647 | -0.190718247 | 0.426862967 | -0.726243772 | 0.467689342 | 0.828619064 | 8.286190636 |
| Cmpk1 | 15.532138 | -0.18614949 | 0.324476729 | -0.7402858 | 0.459126596 | 0.828619064 | 8.286190636 |
| Atp9b | 6.026926 | -0.187179419 | 0.440862213 | -0.721032941 | 0.470889248 | 0.828619064 | 8.286190636 |
| Gm12216 | 27.355824 | -0.16485284 | 0.276426114 | -0.713785767 | 0.475359662 | 0.830051102 | 8.300511017 |
| Pcgf6 | 3.4580526 | -0.144269504 | 0.513775111 | 0.706177363 | 0.480077846 | 0.831890618 | 8.318906183 |
| Rps15 | 4.6742244 | -0.122180135 | 0.531299036 | -0.692708543 | 0.488492477 | 0.833742799 | 8.337427992 |
| Ankrd12 | 7.1624546 | 0.179294206 | 0.432086488 | 0.694108619 | 0.487614094 | 0.833742799 | 8.337427992 |
| 1700126G( | 9.276643 | -0.144269504 | 0.51377511 | -0.616750599 | 0.537399241 | 0.86459497 | 8.645949695 |
| Ninj2 | 4.790965 | -0.16009547 | 0.433409237 | -0.62286903 | 0.533370591 | 0.86459497 | 8.645949695 |
| Rpl13 | 5.65667 | 0.14780551 | 0.491372593 | 0.637822117 | 0.523589481 | 0.86459497 | 8.645949695 |
| Csmd3 | 7.3274617 | -0.144269504 | 0.513775114 | -0.656191429 | 0.511700963 | 0.86459497 | 8.645949695 |
| Rpl39 | 2.3683107 | -0.09634193 | 0.534891928 | -0.619255753 | 0.535747888 | 0.86459497 | 8.645949695 |
| Cntn3 | 2.9678428 | 0.146092302 | 0.470728305 | 0.637786522 | 0.523612654 | 0.86459497 | 8.645949695 |
| Svil | 2.9297297 | -0.144269504 | 0.513775115 | -0.611531223 | 0.540847955 | 0.86459497 | 8.645949695 |
| Ehbp1 | 10.738027 | -0.164675458 | 0.377593533 | -0.628346654 | 0.529776875 | 0.86459497 | 8.645949695 |
| Dctd | 2.5801604 | 0.142014128 | 0.485562678 | 0.627779821 | 0.530148187 | 0.86459497 | 8.645949695 |
| Aldoc | 1.6210175 | -0.076100282 | 0.536588456 | -0.595547394 | 0.551477628 | 0.875422529 | 8.754225291 |
| Apoe | 4.915247 | -0.144073235 | 0.440796464 | -0.575312672 | 0.565079853 | 0.883217023 | 8.832170225 |
| Phkg1 | 4.360975 | -0.144269504 | 0.513775113 | -0.575501243 | 0.56495235 | 0.883217023 | 8.832170225 |
| Rpl36 | 0.8958757 | -0.090210803 | 0.534177943 | -0.570910537 | 0.568060288 | 0.883217023 | 8.832170225 |
| Zfp994 | 3.931868 | -0.120459667 | 0.477723747 | -0.538604619 | 0.5901597 | 0.911335046 | 9.113350463 |
| Rps27a | 2.27826 | -0.095577083 | 0.504610793 | -0.510934253 | 0.609397095 | 0.915960535 | 9.159605349 |
| Rpl35a | 4.942784 | 0.093277215 | 0.558803351 | 0.517578334 | 0.604752501 | 0.915960535 | 9.159605349 |
| Tns3 | 2.8131933 | -0.112828152 | 0.474921041 | -0.505325262 | 0.613330402 | 0.915960535 | 9.159605349 |
| Sec61a1 | 55.388393 | -0.121182337 | 0.298664134 | -0.513301777 | 0.607740241 | 0.915960535 | 9.159605349 |
| Ptpn9 | 6.340883 | -0.131229093 | 0.416979928 | -0.512432361 | 0.608348447 | 0.915960535 | 9.159605349 |
| Gm50306 | 7.646293 | 0.104499314 | 0.421325796 | 0.409798111 | 0.681954052 | 0.925898644 | 9.258986438 |
| Ahdc1 | 3.5728967 | -0.111539245 | 0.446952081 | -0.463243886 | 0.643189563 | 0.925898644 | 9.258986438 |
| Alpl | 3.0174866 | 0.080151273 | 0.449686292 | 0.346066998 | 0.729292365 | 0.925898644 | 9.258986438 |
| Ctcflos | 5.029918 | 0.097145287 | 0.42074837 | 0.389250945 | 0.697090521 | 0.925898644 | 9.258986438 |
| Ctnna2 | 5.1470156 | -0.082806251 | 0.452340834 | -0.346677671 | 0.728833487 | 0.925898644 | 9.258986438 |
| Kcnip4 | 2.905939 | -0.092712285 | 0.470603824 | -0.420545355 | 0.674087104 | 0.925898644 | 9.258986438 |
| Dbi | 2.738859 | 0.067173486 | 0.565110249 | 0.411680865 | 0.680573352 | 0.925898644 | 9.258986438 |
| Rps28 | 4.243435 | -0.095080868 | 0.490639223 | -0.457277272 | 0.647471764 | 0.925898644 | 9.258986438 |
| Pde1c | 8.5778265 | -0.115422499 | 0.37860283 | -0.445691283 | 0.655820262 | 0.925898644 | 9.258986438 |
| Pde6a | 17.578203 | -0.096782011 | 0.326979804 | -0.389023127 | 0.697259038 | 0.925898644 | 9.258986438 |
| Zmat4 | 6.7431383 | 0.105328292 | 0.41280153 | 0.421920788 | 0.673082836 | 0.925898644 | 9.258986438 |
| Ppp1r16a | 5.960048 | -0.093112612 | 0.420513177 | -0.369568738 | 0.711703848 | 0.925898644 | 9.258986438 |
| Cox8a | 6.496518 | 0.071205991 | 0.54873846 | 0.397021406 | 0.691351676 | 0.925898644 | 9.258986438 |
| Ppp1r9a | 4.709364 | -0.092156407 | 0.465743254 | -0.406913909 | 0.68407122 | 0.925898644 | 9.258986438 |
| Eef1a1 | 4.1184964 | -0.084528621 | 0.497329608 | -0.421883892 | 0.673109768 | 0.925898644 | 9.258986438 |
| Zfhx3 | 3.9178238 | -0.094518263 | 0.433198861 | -0.390969827 | 0.695819539 | 0.925898644 | 9.258986438 |
| Anks1b | 1.6476322 | 0.093557143 | 0.496230234 | 0.47733972 | 0.633120231 | 0.925898644 | 9.258986438 |
| Wdr72 | 7.101959 | 0.08721472 | 0.382831648 | 0.344041322 | 0.730815211 | 0.925898644 | 9.258986438 |
| Rplp1 | 7.5951495 | 0.079976245 | 0.495721747 | 0.362531468 | 0.716954912 | 0.925898644 | 9.258986438 |
| Tacc1 | 1.8357242 | 0.066478834 | 0.487532441 | 0.340200626 | 0.733705446 | 0.925898644 | 9.258986438 |
| Ralyl | 3.207875 | 0.082009715 | 0.451525518 | 0.355427807 | 0.72226912 | 0.925898644 | 9.258986438 |
| Vit | 1.6428826 | 0.069858914 | 0.499488594 | 0.368811809 | 0.712268002 | 0.925898644 | 9.258986438 |
| Rpl11 | 4.7606373 | -0.057310026 | 0.530923176 | -0.374018139 | 0.708390821 | 0.925898644 | 9.258986438 |
| Cfap74 | 2.7922401 | -0.102255904 | 0.483220592 | -0.472857578 | 0.636314792 | 0.925898644 | 9.258986438 |
| Unc5d | 11.152448 | 0.09965469 | 0.367305612 | 0.39672163 | 0.691572748 | 0.925898644 | 9.258986438 |
| Ubb | 1.2982892 | -0.064856974 | 0.527830232 | -0.431945867 | 0.665780757 | 0.925898644 | 9.258986438 |
| Bc1 | 2.8435912 | -0.067494663 | 0.490031544 | -0.339553558 | 0.734192757 | 0.925898644 | 9.258986438 |
| Gm960 | 1.248124 | -0.081982014 | 0.513043968 | -0.455820878 | 0.648518789 | 0.925898644 | 9.258986438 |
| Fbxl7 | 4.7657743 | -0.082852676 | 0.420831955 | -0.332695619 | 0.739364057 | 0.927268734 | 9.272687339 |
| Zbtb7a | 6.3876896 | -0.079033081 | 0.400844154 | -0.312683507 | 0.754521127 | 0.941078549 | 9.410785488 |
| Dlg2 | 1.8535248 | -0.144269504 | 0.513775119 | -0.301916255 | 0.762715904 | 0.941237474 | 9.412374737 |

| | | | | | | | |
|---|---|---|---|---|---|---|---|
| Nkain2 | 3.9198008 | 0.069946526 | 0.448747172 | 0.301620487 | 0.762941388 | 0.941237474 | 9.412374737 |
| Ppp3ca | 1.3173085 | -0.144269504 | 0.513775115 | 0.272009781 | 0.785614498 | 0.945490445 | 9.454904455 |
| Grik1 | 283.93326 | -0.047553668 | 0.181012427 | -0.286325407 | 0.7746289 | 0.945490445 | 9.454904455 |
| 8430422H( | 8.173612 | -0.068917919 | 0.360799429 | -0.269929474 | 0.787214512 | 0.945490445 | 9.454904455 |
| Tmsb4x | 4.6209717 | -0.057582213 | 0.48862692 | -0.277326708 | 0.781529254 | 0.945490445 | 9.454904455 |
| Hdac8 | 3.5206144 | -0.144269504 | 0.513775114 | -0.271903818 | 0.785695975 | 0.945490445 | 9.454904455 |
| Rps16 | 2.7770374 | 0.046567835 | 0.521508685 | 0.261362631 | 0.793812872 | 0.948397484 | 9.483974839 |
| Dpyd | 4.496455 | -0.144269504 | 0.513775113 | 0.250712103 | 0.802036703 | 0.953205924 | 9.532059241 |
| Gnaz | 13.247394 | -0.053363376 | 0.312048486 | -0.217977704 | 0.827446484 | 0.956473015 | 9.564730146 |
| Rsrc1 | 12.455451 | -0.056080061 | 0.351261749 | -0.223277782 | 0.823319319 | 0.956473015 | 9.564730146 |
| Ntm | 12.9065 | -0.062092735 | 0.365628773 | -0.241689133 | 0.80902105 | 0.956473015 | 9.564730146 |
| Kcnq5 | 2.647503 | -0.047243145 | 0.479565735 | -0.230178595 | 0.817952994 | 0.956473015 | 9.564730146 |
| Lsamp | 5.6331697 | 0.058371198 | 0.419538993 | 0.23296623 | 0.815787629 | 0.956473015 | 9.564730146 |
| Gm14582 | 4.4432197 | -0.144269504 | 0.513775112 | 0.214615781 | 0.830066889 | 0.956473015 | 9.564730146 |
| Ptprg | 2.6142354 | -0.040838282 | 0.464462162 | -0.191802202 | 0.847897146 | 0.967199257 | 9.671992572 |
| Hapln1 | 3.8963327 | -0.045776998 | 0.435886452 | -0.194351922 | 0.845900333 | 0.967199257 | 9.671992572 |
| Rapgef4 | 13.728402 | -0.144269504 | 0.513775111 | 0.169642838 | 0.865291031 | 0.975080689 | 9.750806892 |
| 4930402H2 | 3.5811813 | -0.03462812 | 0.477944478 | -0.164849463 | 0.869062471 | 0.975080689 | 9.750806892 |
| Selenow | 2.1228044 | 0.0295143 | 0.489751713 | 0.152374775 | 0.878891351 | 0.975080689 | 9.750806892 |
| Meg3 | 3.3725424 | -0.031101726 | 0.490478948 | -0.157733425 | 0.874666861 | 0.975080689 | 9.750806892 |
| St18 | 2.3145986 | -0.032591036 | 0.482263914 | -0.162059762 | 0.871258793 | 0.975080689 | 9.750806892 |
| Sult2a2 | 2.8649654 | 0.033375624 | 0.455458578 | 0.150234666 | 0.880579477 | 0.975080689 | 9.750806892 |
| Prkag2 | 4.1486835 | 0.019369414 | 0.430892938 | 0.081305237 | 0.93519921 | 0.976744347 | 9.767443473 |
| Lyrm1 | 2.0649328 | 0.014940914 | 0.496696078 | 0.079083358 | 0.93696632 | 0.976744347 | 9.767443473 |
| Cox6c | 3.8020508 | 0.024501149 | 0.468587844 | 0.112700593 | 0.910267931 | 0.976744347 | 9.767443473 |
| Pcca | 3.6836677 | -0.144269504 | 0.513775112 | 0.087024944 | 0.930651685 | 0.976744347 | 9.767443473 |
| Atxn7l1 | 2.1083577 | 0.017233569 | 0.475217817 | 0.083541603 | 0.933420899 | 0.976744347 | 9.767443473 |
| Klhl14 | 26.143715 | -0.022619807 | 0.270132161 | -0.100714482 | 0.919777115 | 0.976744347 | 9.767443473 |
| Rps23 | 1.558236 | 0.016926999 | 0.535590659 | 0.120635023 | 0.903980127 | 0.976744347 | 9.767443473 |
| Wdyhv1 | 5.2000837 | 0.026774193 | 0.413920265 | 0.109083725 | 0.913136083 | 0.976744347 | 9.767443473 |
| Rpl41 | 4.4877243 | 0.020296332 | 0.50247697 | 0.101500823 | 0.919152904 | 0.976744347 | 9.767443473 |
| Chl1 | 4.161993 | 0.017784439 | 0.448545414 | 0.077760207 | 0.938018801 | 0.976744347 | 9.767443473 |
| Fth1 | 10.972276 | -0.023427552 | 0.50060585 | -0.115656075 | 0.90792512 | 0.976744347 | 9.767443473 |
| Gm11337 | 4.2377167 | -0.030573927 | 0.444988406 | -0.131404907 | 0.895455007 | 0.976744347 | 9.767443473 |
| Gm29325 | 4.275987 | 0.02614007 | 0.427201587 | 0.108368005 | 0.91370378 | 0.976744347 | 9.767443473 |
| Esf1 | 2.2181897 | 0.012756026 | 0.471217962 | 0.061409519 | 0.951033071 | 0.976852974 | 9.768529736 |
| Il27 | 8.782036 | -0.016565391 | 0.356086827 | -0.06475961 | 0.948365401 | 0.976852974 | 9.768529736 |
| Gas5 | 0.86245275 | -0.02566649 | 0.515881756 | -0.06711698 | 0.946488577 | 0.976852974 | 9.768529736 |
| Hdac9 | 2.0922823 | -0.010012625 | 0.48341308 | -0.050757326 | 0.959518896 | 0.981129682 | 9.811296821 |
| Rps24 | 3.1835344 | -0.006126829 | 0.501523064 | -0.032823815 | 0.973815087 | 0.984111854 | 9.841118542 |
| Gm49767 | 2.9797692 | -0.144269504 | 0.513775114 | 0.030784671 | 0.975441265 | 0.984111854 | 9.841118542 |
| Lrrc17 | 4.172238 | -0.007184896 | 0.439161471 | -0.031205473 | 0.975105675 | 0.984111854 | 9.841118542 |
| Chsy3 | 7.38024 | -0.003179017 | 0.410252917 | -0.013537719 | 0.989198793 | 0.989198793 | 9.891987927 |
| Gnao1 | 2.0144234 | 0.003233266 | 0.469726007 | 0.015153354 | 0.987909836 | 0.989198793 | 9.891987927 |

| **Glut** | baseMean | log2FoldChange | lfcSE | stat | pvalue | padj | pglobadj |
|---|---|---|---|---|---|---|---|
| mt-Cytb | 99.99927 | 1.141406854 | 0.231830091 | 5.30713948 | 1.11E-07 | 1.26E-05 | 0.000126392 |
| Camk1d | 70.181915 | 1.807268701 | 0.366694036 | 5.329986324 | 9.82E-08 | 1.26E-05 | 0.000126392 |
| mt-Nd4 | 90.86455 | 1.16966317 | 0.244458729 | 5.185046467 | 2.16E-07 | 1.63E-05 | 0.000163411 |
| Gphn | 15.688767 | 1.476196532 | 0.327633536 | 5.012875821 | 5.36E-07 | 3.04E-05 | 0.000304308 |
| AY036118 | 96.25058 | 1.709825754 | 0.38181176 | 4.90868955 | 9.17E-07 | 4.16E-05 | 0.000416259 |
| mt-Atp6 | 255.12491 | 0.997950344 | 0.227409737 | 4.781843395 | 1.74E-06 | 6.57E-05 | 0.000657146 |
| 1700054A( | 141.57812 | -0.539049354 | 0.121393985 | -4.663797361 | 3.10E-06 | 0.000100667 | 0.001006671 |
| mt-Co1 | 163.52351 | 0.979109622 | 0.283125074 | 3.931289295 | 8.45E-05 | 0.002397447 | 0.023974471 |
| Gm42418 | 495.07285 | 0.981567016 | 0.296576039 | 3.800782445 | 0.00014424 | 0.003638051 | 0.036380509 |
| mt-Nd2 | 100.98584 | 0.761468147 | 0.262929865 | 3.392312729 | 0.000693053 | 0.0114044 | 0.114043996 |
| mt-Co3 | 317.89905 | 0.80687565 | 0.279733626 | 3.388267997 | 0.000703355 | 0.0114044 | 0.114043996 |
| Pde1c | 48.006958 | -0.743757823 | 0.253003506 | -3.438088162 | 0.000585837 | 0.0114044 | 0.114043996 |
| Gnaz | 44.17836 | -0.67050468 | 0.224453665 | -3.404209871 | 0.000663557 | 0.0114044 | 0.114043996 |
| Gm12216 | 92.376305 | -0.457566487 | 0.143169698 | -3.441893766 | 0.000577657 | 0.0114044 | 0.114043996 |
| mt-Nd1 | 66.879105 | 0.794575058 | 0.288078121 | 3.300618229 | 0.000964721 | 0.014057807 | 0.140578072 |
| mt-Nd5 | 23.92398 | 1.084855779 | 0.417505599 | 3.293109531 | 0.000990859 | 0.014057807 | 0.140578072 |

| | | | | | | | |
|---|---|---|---|---|---|---|---|
| mt-Nd4l | 10.59298 | 1.067714491 | 0.440243925 | 3.221097989 | 0.001277005 | 0.01705177 | 0.170517698 |
| Cmss1 | 56.48602 | 0.990334263 | 0.392467925 | 3.179421164 | 0.001475695 | 0.018548538 | 0.185485384 |
| Gm38973 | 16.28398 | -0.144269504 | 0.40719626 | -3.164682837 | 0.001552521 | 0.018548538 | 0.185485384 |
| mt-Co2 | 179.1176 | 0.680158079 | 0.274650631 | 2.988661041 | 0.002802028 | 0.031803017 | 0.318030171 |
| Ppp3ca | 11.9548855 | 0.934309275 | 0.441678627 | 2.941920987 | 0.003261831 | 0.035258844 | 0.352588443 |
| Syt1 | 12.191012 | 0.944369731 | 0.446875783 | 2.917432403 | 0.003529261 | 0.036415561 | 0.364155606 |
| 8430422H( | 42.030968 | -0.57631884 | 0.247996726 | -2.794297301 | 0.005201264 | 0.047584741 | 0.47584741 |
| Wdyhv1 | 16.539324 | -0.853684122 | 0.404612061 | -2.794637735 | 0.00519579 | 0.047584741 | 0.47584741 |
| Fam169a | 27.947054 | -0.643788386 | 0.283056649 | -2.791859523 | 0.00524061 | 0.047584741 | 0.47584741 |
| Lrrc4c | 18.606462 | 0.764572219 | 0.394736782 | 2.689202701 | 0.007162291 | 0.062532313 | 0.62532313 |
| Adamtsl1 | 32.24669 | -0.529445287 | 0.23792832 | -2.64327387 | 0.008210859 | 0.069032038 | 0.690320378 |
| Ehbp1 | 37.233177 | -0.497586556 | 0.224470332 | -2.616586898 | 0.008881375 | 0.07200258 | 0.720025796 |
| Zmat4 | 23.312378 | -0.144269504 | 0.407196252 | -2.58030096 | 0.009871424 | 0.077269425 | 0.772694248 |
| St18 | 5.592503 | -1.445234022 | 0.826120855 | -2.526826281 | 0.011509842 | 0.086850853 | 0.868508532 |
| Dlg2 | 14.809554 | 0.877363386 | 0.574738389 | 2.51626225 | 0.011860689 | 0.086850853 | 0.868508532 |
| mt-Nd3 | 53.05994 | 0.637978405 | 0.350714198 | 2.487058276 | 0.01288043 | 0.091370549 | 0.913705488 |
| Vit | 14.305894 | -0.144269504 | 0.407196257 | -2.428260547 | 0.01517144 | 0.104361116 | 1.043611162 |
| Pcca | 11.302964 | -0.144269504 | 0.407196264 | -2.414141752 | 0.015772325 | 0.105303463 | 1.053034631 |
| Ralyl | 14.507205 | 0.622511485 | 0.407905704 | 2.345557625 | 0.018998642 | 0.118834143 | 1.188341431 |
| Mbp | 64.762535 | 0.842885089 | 0.650566295 | 2.355107991 | 0.018517323 | 0.118834143 | 1.188341431 |
| Rps27 | 24.597618 | 0.577129819 | 0.351635387 | 2.33834339 | 0.019369442 | 0.118834143 | 1.188341431 |
| Cdk8 | 18.143866 | 0.703997218 | 0.514141578 | 2.312868503 | 0.02072987 | 0.123833697 | 1.238336972 |
| Lrp1b | 204.00333 | -0.340956079 | 0.173441393 | -2.241730242 | 0.024978812 | 0.145389496 | 1.453894959 |
| Kctd16 | 76.70161 | -0.309453805 | 0.158399576 | -2.172537896 | 0.029815114 | 0.165073926 | 1.650739265 |
| Malat1 | 69.80103 | 0.529293577 | 0.343819146 | 2.178241191 | 0.029388085 | 0.165073926 | 1.650739265 |
| Mapk14 | 41.72899 | -0.35934542 | 0.201079632 | -2.090967508 | 0.036530977 | 0.179082888 | 1.790828883 |
| Uba52 | 9.810962 | 0.574039748 | 0.557633832 | 2.105790558 | 0.035222549 | 0.179082888 | 1.790828883 |
| Anks1b | 9.166362 | 0.504319918 | 0.411864286 | 2.076283404 | 0.037867747 | 0.179082888 | 1.790828883 |
| AI314278 | 10.483916 | -0.144269504 | 0.407196256 | -2.084184601 | 0.037143379 | 0.179082888 | 1.790828883 |
| Calm1 | 23.933855 | 0.557377836 | 0.456323421 | 2.085332069 | 0.037039168 | 0.179082888 | 1.790828883 |
| Meg3 | 25.358404 | 0.618570408 | 0.560906431 | 2.115879697 | 0.034355041 | 0.179082888 | 1.790828883 |
| Grik1 | 5.4989185 | -0.690242814 | 0.829611557 | -2.098925555 | 0.035823464 | 0.179082888 | 1.790828883 |
| Spock1 | 12.781663 | -0.525353983 | 0.3668622 | -2.063480808 | 0.039066969 | 0.179625002 | 1.79625002 |
| Phkg1 | 13.941239 | -0.523746895 | 0.375736925 | -2.058262119 | 0.039564978 | 0.179625002 | 1.79625002 |
| Mrc2 | 15.52196 | -0.483291418 | 0.328097862 | -2.031606287 | 0.04219353 | 0.187802573 | 1.878025733 |
| Cadm2 | 11.891537 | 0.499012226 | 0.430765432 | 2.011074896 | 0.044317546 | 0.193463135 | 1.93463135 |
| Ppia | 17.00175 | 0.519847253 | 0.501814117 | 1.99395848 | 0.046156592 | 0.197689555 | 1.976895551 |
| Gas5 | 23.922663 | 0.514762947 | 0.623317291 | 1.982393156 | 0.047435255 | 0.199403757 | 1.994037571 |
| Rapgef4 | 48.983097 | -0.371751543 | 0.240487209 | -1.94882899 | 0.051315851 | 0.21179451 | 2.117945104 |
| Svil | 12.132911 | -0.488716225 | 0.394664901 | -1.933492609 | 0.053175543 | 0.215550861 | 2.155508606 |
| Ptpn9 | 16.17321 | -0.438043713 | 0.33207297 | -1.891548137 | 0.058551212 | 0.220346645 | 2.203466454 |
| Gm29325 | 14.84829 | -0.144269504 | 0.407196258 | -1.900275046 | 0.057397034 | 0.220346645 | 2.203466454 |
| Nckap5 | 94.96033 | -0.344053576 | 0.224352392 | -1.905373902 | 0.056731485 | 0.220346645 | 2.203466454 |
| Rps29 | 86.63813 | 0.476493955 | 0.443831001 | 1.879452129 | 0.060182784 | 0.220346645 | 2.203466454 |
| Car12 | 13.269348 | 0.442946816 | 0.585711972 | 1.882624455 | 0.05975128 | 0.220346645 | 2.203466454 |
| Xlr5a | 95.355095 | -0.354186132 | 0.241388746 | -1.883913922 | 0.05957662 | 0.220346645 | 2.203466454 |
| Rps17 | 12.002829 | 0.440275233 | 0.466423351 | 1.836463372 | 0.066289155 | 0.238851398 | 2.388513981 |
| 5830416I1 | 109.243774 | -0.316660665 | 0.215657155 | -1.807555889 | 0.070675649 | 0.249469867 | 2.494698671 |
| Cdh23 | 31.393366 | -0.144269504 | 0.407196254 | -1.802707887 | 0.071434103 | 0.249469867 | 2.494698671 |
| Il27 | 31.86366 | -0.346090083 | 0.247731988 | -1.783345932 | 0.074530009 | 0.25633806 | 2.563380604 |
| Ttc28 | 27.770508 | -0.345891494 | 0.250491491 | -1.763821352 | 0.077762061 | 0.263462507 | 2.634625067 |
| Ctcflos | 16.329626 | -0.394835199 | 0.413412472 | -1.721110545 | 0.085230766 | 0.284520352 | 2.845203518 |
| Hdac8 | 13.568848 | -0.38096067 | 0.361534928 | -1.670428412 | 0.094834632 | 0.311992196 | 3.119921958 |
| Ctnna3 | 24.259981 | -0.334311437 | 0.268542504 | -1.660508462 | 0.096812207 | 0.313948157 | 3.139481572 |
| Cst3 | 153.94177 | -0.230203132 | 0.458647537 | -1.643221256 | 0.100337164 | 0.314683947 | 3.146839471 |
| Sec61a1 | 171.56808 | -0.206320534 | 0.138594768 | -1.632573625 | 0.102558702 | 0.314683947 | 3.146839471 |
| 4930441J1 | 14.087247 | -0.362460214 | 0.330589883 | -1.632452514 | 0.102584194 | 0.314683947 | 3.146839471 |
| Rab29 | 17.81273 | -0.335489757 | 0.275252996 | -1.637524498 | 0.101520924 | 0.314683947 | 3.146839471 |
| 4930558K( | 19.8751 | -0.342951582 | 0.308844205 | -1.609854146 | 0.107429701 | 0.323934945 | 3.239349454 |
| Chsy3 | 23.052742 | -0.320901863 | 0.266709209 | -1.598728164 | 0.109881017 | 0.323934945 | 3.239349454 |
| Gm49767 | 11.525636 | -0.144269504 | 0.407196256 | -1.603272674 | 0.108874467 | 0.323934945 | 3.239349454 |
| Prkag2 | 18.591381 | -0.326226925 | 0.287566353 | -1.567286548 | 0.117047728 | 0.328911715 | 3.289117154 |

| | | | | | | | |
|---|---|---|---|---|---|---|---|
| Ninj2 | 17.927513 | -0.316433549 | 0.269460419 | -1.568799349 | 0.1166947 | 0.328911715 | 3.289117154 |
| Arvcf | 7.0802913 | -0.144269504 | 0.407196268 | 1.565930138 | 0.117364973 | 0.328911715 | 3.289117154 |
| Ckb | 26.128765 | 0.351100292 | 0.4027356 | 1.582129783 | 0.113619946 | 0.328911715 | 3.289117154 |
| 1700126G0 | 34.365047 | -0.265611159 | 0.207599998 | -1.538074965 | 0.124030287 | 0.343352135 | 3.433521347 |
| Pde6a | 63.212955 | -0.239605084 | 0.180849347 | -1.529700054 | 0.12609099 | 0.344851263 | 3.448512626 |
| Wdr72 | 23.826147 | -0.314089063 | 0.313442425 | -1.502953966 | 0.132850915 | 0.359013783 | 3.59013783 |
| Tns3 | 11.983182 | -0.309330348 | 0.362339813 | -1.442723859 | 0.149098275 | 0.365125804 | 3.651258039 |
| Rsrc1 | 41.97339 | -0.27301014 | 0.256401386 | -1.428553909 | 0.15313249 | 0.365125804 | 3.651258039 |
| Vwc2l | 9.199624 | -0.144269504 | 0.407196265 | -1.418575131 | 0.156022921 | 0.365125804 | 3.651258039 |
| Shb | 57.12811 | -0.230366885 | 0.188463211 | -1.435480867 | 0.151150117 | 0.365125804 | 3.651258039 |
| Prkg1 | 32.408478 | -0.267351235 | 0.229753063 | -1.462079435 | 0.143719444 | 0.365125804 | 3.651258039 |
| Kcnd2 | 8.189679 | 0.29360461 | 0.340310074 | 1.453247166 | 0.146155141 | 0.365125804 | 3.651258039 |
| Sema3c | 66.135925 | -0.261573802 | 0.231595697 | -1.444794035 | 0.148515747 | 0.365125804 | 3.651258039 |
| Ppp1r16a | 14.606436 | -0.297838104 | 0.304973812 | -1.421759847 | 0.15509598 | 0.365125804 | 3.651258039 |
| Ms4a1 | 27.41386 | -0.289924597 | 0.278926838 | -1.449676876 | 0.147148647 | 0.365125804 | 3.651258039 |
| Cpe | 18.960005 | 0.30563006 | 0.423539953 | 1.46839183 | 0.14199782 | 0.365125804 | 3.651258039 |
| Fbxl7 | 16.057865 | -0.300780025 | 0.363787125 | -1.427944473 | 0.153307842 | 0.365125804 | 3.651258039 |
| Gria1 | 10.882296 | -0.144269504 | 0.407196263 | 1.486788786 | 0.137070604 | 0.365125804 | 3.651258039 |
| Csmd3 | 26.341698 | -0.285524592 | 0.260489562 | -1.462182738 | 0.143691141 | 0.365125804 | 3.651258039 |
| 4930511M | 35.85655 | -0.251936886 | 0.22427465 | -1.402215372 | 0.160850943 | 0.372583306 | 3.72583306 |
| Hexb | 15.088464 | 0.251024159 | 0.462932525 | 1.390616381 | 0.164341786 | 0.376824096 | 3.768240956 |
| Zfp994 | 14.944137 | -0.144269504 | 0.407196252 | -1.380661119 | 0.167383181 | 0.379959822 | 3.799598219 |
| Gm11337 | 13.186041 | -0.279885287 | 0.322315851 | -1.342679806 | 0.179375676 | 0.403151273 | 4.031512727 |
| Eef1a1 | 22.886635 | 0.267491325 | 0.38684069 | 1.323445187 | 0.185687373 | 0.405298401 | 4.052984009 |
| Cox7c | 24.603357 | 0.263203241 | 0.410503973 | 1.327201321 | 0.184442096 | 0.405298401 | 4.052984009 |
| Klkb1 | 12.379919 | -0.274951217 | 0.34523734 | -1.327343847 | 0.184394966 | 0.405298401 | 4.052984009 |
| Sgip1 | 6.526274 | 0.251433759 | 0.363245664 | 1.293463514 | 0.195850795 | 0.423410767 | 4.234107671 |
| Dctd | 7.827331 | -0.144269504 | 0.407196259 | -1.244505306 | 0.213313648 | 0.456813189 | 4.568131894 |
| Gm10062 | 59.852055 | -0.22939487 | 0.256840996 | -1.222661066 | 0.221457738 | 0.469821557 | 4.698215567 |
| Acoxl | 21.082848 | -0.22889951 | 0.264576925 | -1.186372313 | 0.235475301 | 0.48857322 | 4.885732199 |
| Zfhx3 | 14.494242 | -0.237446543 | 0.289451525 | -1.188113478 | 0.234788705 | 0.48857322 | 4.885732199 |
| Mc1r | 36.4037 | -0.227828079 | 0.277810601 | -1.183140315 | 0.236753543 | 0.48857322 | 4.885732199 |
| Scamp1 | 17.284771 | -0.231522483 | 0.27812664 | -1.176881821 | 0.239242679 | 0.489262055 | 4.892620549 |
| Dop1b | 23.20367 | -0.227298646 | 0.291266689 | -1.15545995 | 0.247902124 | 0.502444484 | 5.024444842 |
| Rpl11 | 34.706364 | 0.198786046 | 0.415962996 | 1.13230092 | 0.257507944 | 0.517294719 | 5.172947186 |
| Slx1b | 25.369318 | -0.215465869 | 0.27393258 | -1.117325752 | 0.263855069 | 0.525395619 | 5.253956191 |
| Nkain2 | 20.22035 | 0.207908532 | 0.260764333 | 1.109421216 | 0.267248514 | 0.527525328 | 5.27525328 |
| Rps12 | 35.98471 | -0.128960897 | 0.400644648 | -1.09805351 | 0.272181124 | 0.529198145 | 5.291981452 |
| Pcdh15 | 27.633442 | -0.144269504 | 0.407196254 | -1.09673211 | 0.272758515 | 0.529198145 | 5.291981452 |
| Ubb | 13.352551 | 0.198633248 | 0.382924667 | 1.08899582 | 0.276155729 | 0.531248733 | 5.312487332 |
| Ppp1r9a | 15.939527 | -0.207560867 | 0.310413965 | -1.061961668 | 0.288253087 | 0.549860932 | 5.498609315 |
| Il1rapl2 | 9.701231 | -0.144269504 | 0.407196264 | 1.034141669 | 0.301069939 | 0.563688088 | 5.636880879 |
| Nell1 | 23.617365 | -0.192684655 | 0.25239502 | -1.030125051 | 0.302951307 | 0.563688088 | 5.636880879 |
| Cspp1 | 15.578925 | -0.202356529 | 0.304512079 | -1.035766507 | 0.300311087 | 0.563688088 | 5.636880879 |
| Cntn3 | 12.88803 | -0.144269504 | 0.407196256 | -1.019162792 | 0.308125688 | 0.5640688 | 5.640687997 |
| Tacc1 | 11.41664 | -0.200408232 | 0.305859434 | -1.023015194 | 0.306300662 | 0.5640688 | 5.640687997 |
| Ankrd12 | 21.009596 | -0.186125697 | 0.27998973 | -0.979897041 | 0.327136944 | 0.59408069 | 5.940806895 |
| Rps15 | 19.44779 | 0.172992018 | 0.366669071 | 0.954569602 | 0.339795396 | 0.61217107 | 6.1217107 |
| Uqcrh | 17.226807 | 0.138235094 | 0.418828691 | 0.937918209 | 0.348286445 | 0.617664242 | 6.17664242 |
| Rpl39 | 28.428679 | 0.148245635 | 0.409656357 | 0.940740464 | 0.346837877 | 0.617664242 | 6.17664242 |
| Epha6 | 38.557938 | -0.172710078 | 0.257962256 | -0.931469981 | 0.351610506 | 0.618182956 | 6.181829557 |
| Tshz2 | 4.0018873 | -0.125352153 | 0.386721052 | -0.926809443 | 0.354025481 | 0.618182956 | 6.181829557 |
| Tpt1 | 28.210365 | 0.156791059 | 0.387682118 | 0.920655162 | 0.357230493 | 0.619017724 | 6.190177242 |
| Hapln1 | 12.383218 | -0.16376613 | 0.33024488 | -0.884396335 | 0.376482301 | 0.647435472 | 6.474354716 |
| Ccdc60 | 15.782949 | 0.161871985 | 0.289324079 | 0.863218638 | 0.388017268 | 0.661067812 | 6.610678119 |
| Rpl17 | 26.852716 | -0.115788647 | 0.385610124 | -0.85919333 | 0.390233863 | 0.661067812 | 6.610678119 |
| Chl1 | 13.870078 | -0.160549872 | 0.283204722 | -0.847000967 | 0.396994582 | 0.662630663 | 6.62630663 |
| Rpl13a | 23.156572 | 0.135754761 | 0.392542077 | 0.847006636 | 0.396991423 | 0.662630663 | 6.62630663 |
| Fth1 | 159.25063 | 0.137607707 | 0.380220128 | 0.81886519 | 0.412863333 | 0.68408742 | 6.840874196 |
| Gm14582 | 12.835498 | -0.149509518 | 0.31522487 | -0.804625458 | 0.421035847 | 0.687590915 | 6.875909148 |
| Gria4 | 27.528088 | -0.143332008 | 0.227206916 | -0.808853915 | 0.41859918 | 0.687590915 | 6.875909148 |
| Rps3 | 19.521986 | -0.116893822 | 0.375829138 | -0.798044454 | 0.424844693 | 0.688855324 | 6.888553242 |

| | | | | | | | |
|---|---|---|---|---|---|---|---|
| Gm49497 | 13.54824 | -0.144794883 | 0.319879874 | -0.789551152 | 0.429789946 | 0.691931331 | 6.91931331 |
| Rpl36 | 19.816837 | 0.120660217 | 0.38454177 | 0.763501443 | 0.445164402 | 0.711636051 | 7.116360514 |
| Cfap74 | 13.209162 | -0.120488857 | 0.356814354 | -0.743137461 | 0.457398462 | 0.726080076 | 7.260800759 |
| Ahdc1 | 11.812328 | -0.13635831 | 0.288782837 | -0.727143588 | 0.467138 | 0.731312593 | 7.313125925 |
| Dpyd | 15.7993355 | -0.136312456 | 0.284536154 | -0.730987515 | 0.464786779 | 0.731312593 | 7.313125925 |
| Sil1 | 27.170454 | -0.131043319 | 0.258625013 | -0.712291958 | 0.476284003 | 0.740523758 | 7.405237578 |
| Cmpk1 | 49.48254 | -0.122125052 | 0.230044625 | -0.693275975 | 0.488136377 | 0.753788828 | 7.537888277 |
| Rps3a1 | 29.006617 | -0.105648526 | 0.363989291 | -0.68723941 | 0.491931873 | 0.75451713 | 7.545171301 |
| Rpl21 | 37.689503 | 0.108113629 | 0.376984955 | 0.676212761 | 0.498905572 | 0.755533479 | 7.555334785 |
| Zbtb7a | 23.56369 | -0.123248021 | 0.267673033 | -0.668842029 | 0.503596253 | 0.755533479 | 7.555334785 |
| 6030443J0 | 30.79876 | -0.116288811 | 0.224573478 | -0.665222987 | 0.50590788 | 0.755533479 | 7.555334785 |
| Igf2bp3 | 45.83929 | -0.123876644 | 0.277458376 | -0.671723937 | 0.501759459 | 0.755533479 | 7.555334785 |
| Esf1 | 9.794302 | -0.119798893 | 0.305221904 | -0.651417575 | 0.514776969 | 0.763433522 | 7.634335223 |
| 4930402H2 | 10.878186 | 0.118402097 | 0.277224076 | 0.646548708 | 0.517924064 | 0.763433522 | 7.634335223 |
| Lrrc17 | 20.887321 | -0.099041963 | 0.348510705 | -0.607933013 | 0.543231903 | 0.78543721 | 7.854372102 |
| Camk2a | 20.71487 | 0.106374276 | 0.229524327 | 0.608320566 | 0.542974883 | 0.78543721 | 7.854372102 |
| Fyb2 | 22.179024 | -0.107564704 | 0.220714415 | -0.617290366 | 0.53704322 | 0.78543721 | 7.854372102 |
| Rpl32 | 26.072998 | 0.100526698 | 0.355860726 | 0.601662841 | 0.547398589 | 0.786452404 | 7.864524036 |
| Ctnna2 | 13.414483 | -0.108657621 | 0.292285508 | -0.590011193 | 0.555183146 | 0.789997377 | 7.899973766 |
| Gnao1 | 10.703454 | 0.105952611 | 0.301523518 | 0.587561968 | 0.556826345 | 0.789997377 | 7.899973766 |
| Tmem108 | 19.81278 | -0.103931379 | 0.237659817 | -0.579476933 | 0.562267407 | 0.79276212 | 7.9276212 |
| Tmeff2 | 11.866141 | -0.099978079 | 0.262993891 | -0.555194496 | 0.578761633 | 0.806005465 | 8.060054649 |
| Rps23 | 25.459724 | 0.092634447 | 0.353113963 | 0.555744723 | 0.578385378 | 0.806005465 | 8.060054649 |
| Cox8a | 54.110287 | 0.075502195 | 0.38993469 | 0.535517602 | 0.592291989 | 0.81981879 | 8.198187896 |
| Lyrm1 | 7.9197645 | -0.144269504 | 0.407196261 | 0.52753611 | 0.597821347 | 0.82123105 | 8.212310497 |
| Unc5d | 1.6227521 | -0.144269504 | 0.407196266 | 0.523612887 | 0.600547816 | 0.82123105 | 8.212310497 |
| Kcnq5 | 13.298319 | 0.091641045 | 0.295967029 | 0.504507466 | 0.613904817 | 0.834469422 | 8.344694221 |
| Rpl37a | 40.960964 | 0.072661234 | 0.374616177 | 0.482758693 | 0.629267084 | 0.83590678 | 8.3590678 |
| Ptprd | 14.081265 | -0.088335664 | 0.290760915 | -0.486047845 | 0.626933244 | 0.83590678 | 8.3590678 |
| Ntm | 60.91025 | 0.084104275 | 0.229428545 | 0.48216056 | 0.629691892 | 0.83590678 | 8.3590678 |
| Aldoc | 15.485122 | 0.073059948 | 0.370853073 | 0.486245942 | 0.626792802 | 0.83590678 | 8.3590678 |
| Rpl37 | 58.321743 | 0.077007253 | 0.359156735 | 0.476739541 | 0.633547603 | 0.836135499 | 8.361354991 |
| Rpl41 | 37.511967 | 0.0742768 | 0.360549797 | 0.465314579 | 0.641706196 | 0.840255067 | 8.40255067 |
| Mt1 | 35.063984 | 0.076599405 | 0.348582207 | 0.462012766 | 0.644072166 | 0.840255067 | 8.40255067 |
| Rps27a | 33.156513 | 0.065808895 | 0.377721349 | 0.451301549 | 0.651772228 | 0.845441691 | 8.454416906 |
| Hdac9 | 9.57025 | -0.144269504 | 0.407196255 | 0.444329346 | 0.656804487 | 0.847128515 | 8.471285146 |
| Trim2 | 42.692062 | 0.069839948 | 0.234662888 | 0.394609703 | 0.693130951 | 0.856581493 | 8.565814926 |
| Tmsb4x | 42.59606 | -0.060761127 | 0.362470571 | -0.42227623 | 0.672823406 | 0.856581493 | 8.565814926 |
| Cep128 | 10.783486 | 0.074296709 | 0.293882898 | 0.413125004 | 0.679515031 | 0.856581493 | 8.565814926 |
| D430042O | 59.267593 | -0.144269504 | 0.407196239 | -0.383245534 | 0.701537719 | 0.856581493 | 8.565814926 |
| Lsamp | 27.76111 | 0.070463979 | 0.236850577 | 0.39903779 | 0.68986536 | 0.856581493 | 8.565814926 |
| Gm765 | 13.4760685 | -0.068263605 | 0.25119173 | -0.383117497 | 0.701632647 | 0.856581493 | 8.565814926 |
| Atp9b | 22.188927 | -0.072704474 | 0.289352306 | -0.402613433 | 0.687232626 | 0.856581493 | 8.565814926 |
| Pcgf6 | 13.125739 | -0.073312488 | 0.265336155 | -0.401883916 | 0.68776946 | 0.856581493 | 8.565814926 |
| Galnt13 | 29.213446 | 0.071140237 | 0.249977952 | 0.39515813 | 0.692726191 | 0.856581493 | 8.565814926 |
| Rps8 | 42.907177 | -0.052132286 | 0.368419485 | -0.38279936 | 0.701868536 | 0.856581493 | 8.565814926 |
| Rpl34 | 22.53576 | 0.056719176 | 0.348383877 | 0.350546286 | 0.725928759 | 0.870078153 | 8.700781531 |
| Klhl14 | 2.6490629 | 0.047025497 | 0.364628043 | 0.35172423 | 0.725045087 | 0.870078153 | 8.700781531 |
| Ptprg | 9.615968 | -0.144269504 | 0.407196264 | 0.348231222 | 0.727666542 | 0.870078153 | 8.700781531 |
| Srgn | 6.927038 | -0.144269504 | 0.407196261 | -0.34744205 | 0.728259247 | 0.870078153 | 8.700781531 |
| Adk | 15.99909 | -0.057974992 | 0.244087809 | -0.323772197 | 0.746110507 | 0.880264192 | 8.80264192 |
| Cox6c | 36.238724 | -0.043074743 | 0.368827452 | -0.320725457 | 0.748418454 | 0.880264192 | 8.80264192 |
| Rpl23 | 26.072792 | 0.046330823 | 0.372125018 | 0.321866715 | 0.74755367 | 0.880264192 | 8.80264192 |
| Rps21 | 69.29485 | 0.045471482 | 0.363323006 | 0.300080816 | 0.764115512 | 0.894093923 | 8.940939233 |
| Sult2a2 | 10.396637 | -0.053570248 | 0.267365284 | -0.294897809 | 0.768071946 | 0.894114522 | 8.941145218 |
| Gm50306 | 32.413666 | 0.045623139 | 0.196725431 | 0.280934093 | 0.778760951 | 0.901932326 | 9.019323258 |
| Plp1 | 11.947595 | 0.035541751 | 0.382653347 | 0.274572855 | 0.783644424 | 0.902981138 | 9.029811378 |
| Cox4i1 | 55.03479 | 0.036463729 | 0.375594622 | 0.261042973 | 0.794059368 | 0.910360993 | 9.10360993 |
| Bc1 | 16.326283 | -0.037749982 | 0.347166906 | -0.245925169 | 0.80574016 | 0.919110635 | 9.191106346 |
| Cox6b1 | 15.940308 | 0.028286172 | 0.38235901 | 0.218340405 | 0.827163895 | 0.924691745 | 9.246917451 |
| Kat7 | 14.177729 | -0.037599023 | 0.264953748 | -0.208198419 | 0.835074043 | 0.924691745 | 9.246917451 |
| Selenow | 17.175367 | 0.033402195 | 0.363445869 | 0.226513334 | 0.82080219 | 0.924691745 | 9.246917451 |

| | | | | | | | |
|---|---|---|---|---|---|---|---|
| Rpl38 | 36.311028 | 0.033693879 | 0.345869762 | 0.210279859 | 0.833449254 | 0.924691745 | 9.246917451 |
| Rpl30 | 34.068516 | 0.034583118 | 0.361438881 | 0.230424485 | 0.817761936 | 0.924691745 | 9.246917451 |
| Rps10 | 27.12138 | -0.031508924 | 0.361107973 | -0.221364988 | 0.824808252 | 0.924691745 | 9.246917451 |
| Rps7 | 21.615023 | -0.027012297 | 0.365006664 | -0.197058836 | 0.843781496 | 0.929798056 | 9.297980562 |
| Rpl13 | 48.121593 | -0.024355528 | 0.373184657 | -0.190000037 | 0.849309102 | 0.931367952 | 9.31367952 |
| Fau | 36.68445 | -0.021036099 | 0.381851144 | -0.183205608 | 0.854636694 | 0.932704469 | 9.327044695 |
| Rps14 | 37.294437 | 0.026004943 | 0.357540357 | 0.170921731 | 0.864285309 | 0.938721365 | 9.387213648 |
| Rps20 | 30.235338 | -0.019514023 | 0.359741271 | -0.137538021 | 0.890605542 | 0.958139611 | 9.581396114 |
| Atxn7l1 | 12.299738 | 0.025198186 | 0.283719673 | 0.138091704 | 0.890167942 | 0.958139611 | 9.581396114 |
| Rpl9 | 27.202162 | -0.015730972 | 0.367597507 | -0.116668422 | 0.907122815 | 0.966745911 | 9.667459108 |
| Rpl27a | 34.65671 | 0.018395768 | 0.357426508 | 0.121176334 | 0.903551369 | 0.966745911 | 9.667459108 |
| Rps28 | 25.47918 | -0.015699043 | 0.346515256 | -0.102340677 | 0.91848627 | 0.969750619 | 9.697506195 |
| Rpl26 | 34.769566 | -0.015002163 | 0.357485776 | -0.10427362 | 0.916952212 | 0.969750619 | 9.697506195 |
| Dbi | 18.818296 | 0.014484084 | 0.33572053 | 0.087890759 | 0.929963502 | 0.97023563 | 9.702356302 |
| Rps24 | 50.57764 | -0.008662013 | 0.375263795 | -0.069501253 | 0.944590636 | 0.97023563 | 9.702356302 |
| Rps16 | 24.292126 | -0.010616225 | 0.356253251 | -0.075392055 | 0.939902781 | 0.97023563 | 9.702356302 |
| Gm960 | 8.686207 | -0.144269504 | 0.407196266 | -0.081692203 | 0.934891479 | 0.97023563 | 9.702356302 |
| Apoe | 80.2238 | -0.015064111 | 0.339946363 | -0.095566939 | 0.923864524 | 0.97023563 | 9.702356302 |
| Cntnap2 | 56.53396 | 0.010957104 | 0.176737414 | 0.071157295 | 0.943272568 | 0.97023563 | 9.702356302 |
| Rplp1 | 76.570724 | -0.008334937 | 0.354587338 | -0.05798821 | 0.953758019 | 0.975239056 | 9.75239056 |
| Rpl12 | 12.270738 | -0.003384483 | 0.365094875 | -0.026157022 | 0.979132096 | 0.984828481 | 9.848284813 |
| Mt3 | 35.23202 | 0.004726951 | 0.365627923 | 0.031827695 | 0.97460946 | 0.984828481 | 9.848284813 |
| Kcnip4 | 17.83586 | -0.005041769 | 0.276934061 | -0.02873093 | 0.977079188 | 0.984828481 | 9.848284813 |
| Alpl | 10.492561 | 0.004568775 | 0.29255811 | 0.024454559 | 0.98049003 | 0.984828481 | 9.848284813 |
| Rpl35a | 37.172855 | 0.001274284 | 0.361116162 | 0.007080097 | 0.994350947 | 0.994350947 | 9.943509473 |

**table S9: Differential gene expression list quinine compared to rest (3) conditions in GABA and Glut neurons**

| **GABA** | baseMean | log2FoldChange | lfcSE | stat | pvalue | padj | pglobadj |
|---|---|---|---|---|---|---|---|
| Cst3 | 20.57817 | -1.40531633 | 0.436610172 | -4.029334519 | 5.59E-05 | 0.002551156 | 0.02551156 |
| mt-Co1 | 29.908287 | -1.208045081 | 0.372061266 | -4.028770326 | 5.61E-05 | 0.002551156 | 0.02551156 |
| mt-Atp6 | 52.440006 | -0.941372072 | 0.403186228 | -3.321871014 | 0.00089416 | 0.024074178 | 0.240741777 |
| Ntm | 12.9065 | 0.56368479 | 0.208240516 | 3.210996162 | 0.001322757 | 0.024074178 | 0.240741777 |
| Rps21 | 6.852249 | -1.410357488 | 0.664547055 | -3.228219253 | 0.001245635 | 0.024074178 | 0.240741777 |
| Cox8a | 6.496518 | -0.149649504 | 0.330804148 | -2.944854569 | 0.003231065 | 0.044282677 | 0.442826774 |
| Gm42418 | 88.84172 | -0.304686311 | 0.646908122 | -2.873926815 | 0.004054029 | 0.044282677 | 0.442826774 |
| mt-Co3 | 64.31273 | -0.652673061 | 0.346027828 | -2.879324759 | 0.003985277 | 0.044282677 | 0.442826774 |
| Lrp1b | 56.844112 | 0.334231247 | 0.134589199 | 2.849441533 | 0.004379605 | 0.044282677 | 0.442826774 |
| Grik1 | 283.93326 | -0.310143812 | 0.128650308 | -2.779867145 | 0.005438114 | 0.048583724 | 0.485837243 |
| mt-Nd2 | 16.228426 | -0.830714896 | 0.658368991 | -2.754802472 | 0.005872758 | 0.048583724 | 0.485837243 |
| Prkg1 | 7.6053395 | -0.144269504 | 0.401982134 | 2.724988597 | 0.006430371 | 0.048763645 | 0.487636451 |
| Camk1d | 9.3028755 | -0.465842326 | 1.013363589 | -2.609179881 | 0.009075952 | 0.063531664 | 0.635316635 |
| Unc5d | 11.152448 | -0.144269504 | 0.401982115 | -2.555294578 | 0.010609797 | 0.066781965 | 0.667819655 |
| Fth1 | 10.972276 | -0.172860379 | 0.356732174 | -2.542444243 | 0.011008016 | 0.066781965 | 0.667819655 |
| mt-Co2 | 31.080164 | -0.49983989 | 0.41217816 | -2.410348125 | 0.015937305 | 0.090643424 | 0.906434236 |
| mt-Nd3 | 11.394203 | -0.479525012 | 0.592570983 | -2.384492781 | 0.017102684 | 0.091549659 | 0.91549659 |
| mt-Nd5 | 4.970977 | -0.303231411 | 0.556599836 | -2.35035052 | 0.018755739 | 0.094820682 | 0.948206823 |
| Rps29 | 11.573063 | -0.129487483 | 0.311840414 | -2.293756233 | 0.021804505 | 0.104432101 | 1.044321014 |
| Rpl37 | 5.0523815 | -0.181699047 | 0.376428939 | -2.267217102 | 0.023376967 | 0.106365198 | 1.063651982 |
| Klhl14 | 26.143715 | 0.295857685 | 0.196619059 | 2.042006467 | 0.04115089 | 0.16281439 | 1.628143903 |
| Rps3a1 | 6.1928663 | -0.056915202 | 0.264069558 | -2.067473224 | 0.038689585 | 0.16281439 | 1.628143903 |
| Ankrd12 | 7.1624546 | -0.360559889 | 0.39977318 | -2.058932403 | 0.039500715 | 0.16281439 | 1.628143903 |
| mt-Cytb | 17.050423 | -0.336101048 | 0.363836381 | -1.999599951 | 0.045543479 | 0.172685692 | 1.726856923 |
| Igf2bp3 | 12.022297 | 0.275202591 | 0.216674545 | 1.844511743 | 0.065108596 | 0.236995289 | 2.369952894 |
| Rpl38 | 7.104176 | -0.130333692 | 0.318158187 | -1.722466384 | 0.084985067 | 0.297447735 | 2.974477351 |
| Galnt13 | 6.1900806 | 0.258970902 | 0.275714547 | 1.67765 | 0.093415446 | 0.313712257 | 3.137122573 |
| AY036118 | 14.886724 | -0.152902226 | 0.324590734 | -1.661929812 | 0.096526848 | 0.313712257 | 3.137122573 |
| mt-Nd1 | 13.160506 | -0.146832818 | 0.321797372 | -1.627473906 | 0.103636486 | 0.325204146 | 3.252041464 |
| Rsrc1 | 12.455451 | -0.217480618 | 0.227536152 | -1.526779423 | 0.12681587 | 0.384674806 | 3.846748058 |
| Nell1 | 5.491928 | 0.180186636 | 0.331126044 | 1.4142975 | 0.157274571 | 0.458221319 | 4.582213188 |
| Mrc2 | 6.650044 | -0.174313778 | 0.314354927 | -1.401271931 | 0.161132771 | 0.458221319 | 4.582213188 |
| Rpl13 | 5.65667 | -0.104998394 | 0.3033684 | -1.323836167 | 0.185557462 | 0.494854335 | 4.948543351 |
| Fam169a | 6.7369084 | 0.166647052 | 0.308794624 | 1.296001492 | 0.194974971 | 0.494854335 | 4.948543351 |
| Chsy3 | 7.38024 | -0.151359693 | 0.305651125 | -1.296710587 | 0.194730785 | 0.494854335 | 4.948543351 |
| Camk2a | 5.2381573 | -0.144269504 | 0.401982141 | 1.293707279 | 0.19576655 | 0.494854335 | 4.948543351 |
| Adamtsl1 | 11.004572 | 0.161596689 | 0.272231767 | 1.203465444 | 0.228796256 | 0.562715116 | 5.627151157 |
| Cdh23 | 7.7059603 | -0.144269504 | 0.401982119 | -1.130369494 | 0.258320563 | 0.581803437 | 5.818034368 |
| 8430422H( | 8.173612 | 0.149192983 | 0.237207996 | 1.106466484 | 0.268524663 | 0.581803437 | 5.818034368 |
| Gm10062 | 13.044163 | 0.145108559 | 0.271792305 | 1.115934072 | 0.264450356 | 0.581803437 | 5.818034368 |
| mt-Nd4 | 15.204892 | -0.1083093 | 0.286422345 | -1.115595581 | 0.264595284 | 0.581803437 | 5.818034368 |
| Sec61a1 | 55.388393 | -0.150121195 | 0.235558284 | -1.135257341 | 0.25626751 | 0.581803437 | 5.818034368 |
| Malat1 | 6.5285845 | -0.049282832 | 0.275207738 | -1.068142653 | 0.285456168 | 0.589746721 | 5.897467213 |
| Nckap5 | 23.137186 | 0.139380644 | 0.211294139 | 1.040486943 | 0.298113727 | 0.589746721 | 5.897467213 |
| Rplp1 | 7.5951495 | -0.071780306 | 0.283432429 | -1.047127789 | 0.295040644 | 0.589746721 | 5.897467213 |
| 1700126G( | 9.276643 | 0.140593281 | 0.258741595 | 1.070405467 | 0.284436839 | 0.589746721 | 5.897467213 |
| Pde1c | 8.5778265 | 0.130094789 | 0.267940761 | 1.019191367 | 0.308112124 | 0.596557518 | 5.965575175 |
| Kctd16 | 20.133091 | 0.122180351 | 0.266118139 | 0.987208824 | 0.323540277 | 0.613378442 | 6.13378442 |
| Ctcflos | 5.029918 | -0.103520918 | 0.28779016 | -0.941058642 | 0.346674808 | 0.633788606 | 6.33788606 |
| Ms4a1 | 6.423043 | 0.115514053 | 0.274312967 | 0.938017342 | 0.348235498 | 0.633788606 | 6.33788606 |
| Rapgef4 | 13.728402 | -0.144269504 | 0.401982132 | 0.909748936 | 0.36295493 | 0.647625463 | 6.476254634 |
| Ttc28 | 5.5357037 | 0.094589655 | 0.291082402 | 0.869008269 | 0.384842608 | 0.673474564 | 6.73474564 |
| Ppp1r16a | 5.960048 | -0.089919064 | 0.281341759 | -0.853255201 | 0.393517797 | 0.675662633 | 6.756626333 |
| Zbtb7a | 6.3876896 | 0.092884613 | 0.274511793 | 0.792826115 | 0.427879139 | 0.687582855 | 6.875828546 |
| Mc1r | 10.825235 | 0.102062203 | 0.25288393 | 0.820285689 | 0.412053263 | 0.687582855 | 6.875828546 |
| Pcdh15 | 10.199923 | -0.097650798 | 0.244458621 | -0.788022122 | 0.430683766 | 0.687582855 | 6.875828546 |
| Ehbp1 | 10.738027 | -0.103911741 | 0.209233641 | -0.802720579 | 0.422136255 | 0.687582855 | 6.875828546 |
| 4930558K( | 5.919937 | -0.144269504 | 0.401982134 | 0.647249193 | 0.517470678 | 0.81189365 | 8.118936504 |
| Ctnna2 | 5.1470156 | -0.060679112 | 0.281290231 | -0.618228005 | 0.536425052 | 0.827367454 | 8.273674536 |
| 4930511M | 7.9351397 | 0.067587006 | 0.262191954 | 0.592102145 | 0.553782191 | 0.839902989 | 8.39902989 |
| Gm12216 | 27.355824 | -0.072794682 | 0.19368727 | -0.577297445 | 0.563738539 | 0.840987001 | 8.409870013 |
| Trim2 | 9.635662 | -0.069193589 | 0.220051601 | -0.549643283 | 0.582564066 | 0.855053709 | 8.550537092 |

| | | | | | | | |
|---|---|---|---|---|---|---|---|
| Il27 | 8.782036 | -0.144269504 | 0.401982126 | -0.516606163 | 0.605431113 | 0.868121716 | 8.681217159 |
| Cmpk1 | 15.532138 | -0.063991485 | 0.212514423 | -0.509292605 | 0.610547141 | 0.868121716 | 8.681217159 |
| Rpl26 | 6.793167 | -0.029616365 | 0.279377337 | -0.481407813 | 0.630226685 | 0.882317358 | 8.823173585 |
| 1700054A( | 52.824097 | -0.048627073 | 0.227691726 | -0.414488248 | 0.678516572 | 0.896521339 | 8.965213394 |
| Mapk14 | 11.374496 | -0.144269504 | 0.401982126 | -0.412763472 | 0.679779917 | 0.896521339 | 8.965213394 |
| Epha6 | 12.959585 | -0.04981001 | 0.244715108 | -0.436241185 | 0.66266175 | 0.896521339 | 8.965213394 |
| Csmd3 | 7.3274617 | -0.144269504 | 0.401982122 | -0.453190818 | 0.650411344 | 0.896521339 | 8.965213394 |
| Zmat4 | 6.7431383 | 0.039745482 | 0.271656254 | 0.385323538 | 0.699997725 | 0.909997042 | 9.099970421 |
| Ctnna3 | 7.525647 | 0.040576802 | 0.259523969 | 0.365232261 | 0.714938038 | 0.916329035 | 9.16329035 |
| Sil1 | 7.6066775 | 0.035400185 | 0.251033782 | 0.306585496 | 0.759158895 | 0.94634876 | 9.463487595 |
| Slx1b | 6.8200607 | 0.033498978 | 0.264498802 | 0.31802202 | 0.750468234 | 0.94634876 | 9.463487595 |
| Lsamp | 5.6331697 | -0.026759347 | 0.262513454 | -0.249027123 | 0.803339801 | 0.950269552 | 9.502695517 |
| Wdr72 | 7.101959 | 0.032641713 | 0.253956936 | 0.284460094 | 0.776057812 | 0.950269552 | 9.502695517 |
| Sema3c | 24.274586 | 0.02758308 | 0.150896057 | 0.248077769 | 0.804074236 | 0.950269552 | 9.502695517 |
| Tshz2 | 10.283613 | -0.144269504 | 0.401982134 | 0.261179427 | 0.793954143 | 0.950269552 | 9.502695517 |
| 6030443J0 | 7.6030917 | -0.144269504 | 0.401982127 | -0.216762914 | 0.828393115 | 0.954224981 | 9.542249807 |
| Gm50306 | 7.646293 | -0.02036379 | 0.267355825 | -0.220335115 | 0.825610175 | 0.954224981 | 9.542249807 |
| Shb | 15.204553 | -0.017988833 | 0.208495742 | -0.146717541 | 0.883354973 | 0.960111279 | 9.601112787 |
| Cntnap2 | 15.640811 | -0.016787907 | 0.241043155 | -0.150731037 | 0.880187889 | 0.960111279 | 9.601112787 |
| Cmss1 | 5.8559604 | 0.015833963 | 0.2628239 | 0.143042565 | 0.886256565 | 0.960111279 | 9.601112787 |
| D430042O( | 15.700211 | 0.02056946 | 0.249241384 | 0.190008009 | 0.849302855 | 0.960111279 | 9.601112787 |
| 5830416I1! | 24.880907 | 0.018966294 | 0.184452914 | 0.151794978 | 0.879348644 | 0.960111279 | 9.601112787 |
| Atp9b | 6.026926 | -0.01192279 | 0.270075183 | -0.123486616 | 0.901721772 | 0.965372721 | 9.653727207 |
| Pde6a | 17.578203 | 0.002417136 | 0.210763278 | 0.018497939 | 0.985241622 | 0.985750061 | 9.857500614 |
| Gria4 | 7.6431046 | 0.002168722 | 0.267345468 | 0.022136582 | 0.982339006 | 0.985750061 | 9.857500614 |
| Wdyhv1 | 5.2000837 | -0.001715426 | 0.270396028 | -0.017860599 | 0.985750061 | 0.985750061 | 9.857500614 |
| Xlr5a | 27.32727 | -0.007494302 | 0.204528025 | -0.06500493 | 0.948170075 | 0.985750061 | 9.857500614 |
| Ptpn9 | 6.340883 | 0.006901203 | 0.237477318 | 0.049304066 | 0.96067698 | 0.985750061 | 9.857500614 |
| Gnaz | 13.247394 | 0.006512036 | 0.20213355 | 0.050356926 | 0.959837961 | 0.985750061 | 9.857500614 |
| 4930402H2 | 3.5811813 | -0.119019949 | 0.321386543 | -1.282586903 | 0.199636841 | | |
| 4930441J1 | 3.6492968 | -0.045404204 | 0.288123677 | -0.576982603 | 0.563951208 | | |
| AI314278 | 3.58537 | -0.235564998 | 0.432676873 | -1.988816424 | 0.046721469 | | |
| Acoxl | 4.884706 | 0.101700697 | 0.293768573 | 0.914799113 | 0.3602971 | | |
| Adk | 3.7155418 | -0.073210081 | 0.296392958 | -0.74187631 | 0.458162281 | | |
| Ahdc1 | 3.5728967 | -0.144269504 | 0.401982122 | 0.562986214 | 0.573444272 | | |
| Aldoc | 1.6210175 | -0.029263333 | 0.277475277 | -0.784046662 | 0.433012731 | | |
| Alpl | 3.0174866 | 0.025336777 | 0.292278234 | 0.268647569 | 0.788200902 | | |
| Anks1b | 1.6476322 | -0.144269504 | 0.40198212 | -0.720081583 | 0.471474766 | | |
| Apoe | 4.915247 | -0.085567752 | 0.287053115 | -0.849646568 | 0.395521613 | | |
| Arvcf | 1.1981798 | 0.045634929 | 0.324437572 | 0.731094717 | 0.464721301 | | |
| Atxn7l1 | 2.1083577 | -0.097883942 | 0.320526223 | -1.134931184 | 0.256404153 | | |
| Bc1 | 2.8435912 | 0.102063827 | 0.339893892 | 1.160604836 | 0.245802638 | | |
| Cadm2 | 1.128733 | 0.077326969 | 0.360062174 | 1.236231166 | 0.216372649 | | |
| Calm1 | 3.654584 | -0.02832227 | 0.272184252 | -0.836783145 | 0.402714474 | | |
| Car12 | 3.4988654 | -0.084546038 | 0.289781852 | -1.689124316 | 0.091195609 | | |
| Ccdc60 | 3.9716513 | -0.005864442 | 0.273758535 | -0.059580236 | 0.95248996 | | |
| Cdk8 | 4.057161 | -0.064035647 | 0.292163093 | -0.921735787 | 0.356666409 | | |
| Cep128 | 2.7124422 | 0.047734722 | 0.300162713 | 0.528352665 | 0.597254585 | | |
| Cfap74 | 2.7922401 | 0.034331393 | 0.302493037 | 0.43543225 | 0.663248706 | | |
| Chl1 | 4.161993 | -0.05236023 | 0.29055723 | -0.614731146 | 0.538732287 | | |
| Ckb | 3.2405775 | -0.126937894 | 0.326010805 | -2.100793316 | 0.035659113 | | |
| Cntn3 | 2.9678428 | -0.144269504 | 0.40198213 | 0.491107992 | 0.623350069 | | |
| Cox4i1 | 4.267222 | -0.093756983 | 0.298985209 | -1.538106271 | 0.124022633 | | |
| Cox6b1 | 3.6377096 | -0.108127464 | 0.314476355 | -1.27284212 | 0.203074071 | | |
| Cox6c | 3.8020508 | 0.032760686 | 0.29829222 | 0.414923276 | 0.67819807 | | |
| Cox7c | 4.358693 | -1.910695345 | 0.960989383 | -3.164997467 | 0.001550843 | | |
| Cpe | 2.8087764 | -0.028223932 | 0.297744366 | -0.406320922 | 0.684506815 | | |
| Cspp1 | 3.2586973 | -0.144269504 | 0.40198212 | 0.244911718 | 0.806524788 | | |
| Dbi | 2.738859 | -0.014415472 | 0.280170676 | -0.376705613 | 0.706392391 | | |
| Dctd | 2.5801604 | -0.104021591 | 0.315988996 | -1.00858696 | 0.313172764 | | |
| Dlg2 | 1.8535248 | -0.033394277 | 0.304177404 | -0.451722344 | 0.65146902 | | |
| Dop1b | 4.380478 | 0.227749145 | 0.35960811 | 1.621158294 | 0.104983695 | | |
| Dpyd | 4.496455 | 0.083124124 | 0.287506634 | 0.748848303 | 0.453948642 | | |
| Eef1a1 | 4.1184964 | -0.171949711 | 0.361713187 | -1.980283478 | 0.047671684 | | |
| Esf1 | 2.2181897 | -0.062952417 | 0.307588653 | -0.733981659 | 0.462959916 | | |

| | | | | | |
|---|---|---|---|---|---|
| Fau | 2.4518764 | -0.001634928 | 0.297634522 | -0.0374779 | 0.970103961 |
| Fbxl7 | 4.7657743 | -0.144269504 | 0.40198213 | 0.480007157 | 0.631222304 |
| Fyb2 | 4.7031064 | 0.034083815 | 0.279857522 | 0.353598395 | 0.723639877 |
| Gas5 | 0.86245275 | -0.010090368 | 0.312951761 | -0.225880252 | 0.821294558 |
| Gm11337 | 4.2377167 | -0.085252913 | 0.295570933 | -0.863276942 | 0.387985218 |
| Gm14582 | 4.4432197 | -0.144269504 | 0.401982121 | -0.827781397 | 0.407794312 |
| Gm29325 | 4.275987 | -0.06949 | 0.271580701 | -0.598466153 | 0.549528937 |
| Gm38973 | 4.666378 | -0.077490936 | 0.290346724 | -0.796659785 | 0.425648646 |
| Gm49497 | 2.7023165 | 0.075684989 | 0.321541088 | 0.869683659 | 0.384473305 |
| Gm49767 | 2.9797692 | -0.035279304 | 0.292397098 | -0.390545072 | 0.696133534 |
| Gm765 | 2.7776783 | 0.074596951 | 0.310005602 | 0.779967369 | 0.435410082 |
| Gm960 | 1.248124 | -0.005209323 | 0.309472797 | -0.089906365 | 0.928361623 |
| Gnao1 | 2.0144234 | -0.144269504 | 0.401982125 | 1.131122496 | 0.258003537 |
| Gphn | 2.9946876 | -0.052305969 | 0.297613377 | -0.847347402 | 0.396801513 |
| Gria1 | 1.568388 | -0.144269504 | 0.401982129 | 0.649350791 | 0.516111663 |
| Hapln1 | 3.8963327 | -0.144269504 | 0.401982124 | 0.996401732 | 0.319054992 |
| Hdac8 | 3.5206144 | -0.144269504 | 0.401982122 | -0.207170715 | 0.835876537 |
| Hdac9 | 2.0922823 | 0.123867089 | 0.361963064 | 1.368612046 | 0.171120577 |
| Hexb | 3.3334806 | -0.07356898 | 0.294387595 | -1.328733874 | 0.183935785 |
| Il1rapl2 | 2.7269056 | -0.1513207 | 0.358341009 | -1.709712128 | 0.087319119 |
| Kat7 | 4.219699 | -0.144269504 | 0.401982124 | 1.093916002 | 0.27399182 |
| Kcnd2 | 2.514149 | -0.030650505 | 0.296704871 | -0.441880627 | 0.65857559 |
| Kcnip4 | 2.905939 | -0.045256307 | 0.293968399 | -0.519413324 | 0.603472542 |
| Kcnq5 | 2.647503 | -0.031515364 | 0.293386874 | -0.41570276 | 0.677627521 |
| Klkb1 | 2.2612617 | 0.076273545 | 0.320696335 | 0.876657314 | 0.380672799 |
| Lrrc17 | 4.172238 | 0.017328843 | 0.286888626 | 0.184433991 | 0.853673006 |
| Lrrc4c | 2.4291487 | 0.087903795 | 0.326083394 | 1.001526153 | 0.316572503 |
| Lyrm1 | 2.0649328 | 0.04194384 | 0.317721309 | 0.620583154 | 0.534873926 |
| Mbp | 3.1925125 | -0.216609953 | 0.441122757 | -3.164116203 | 0.001555547 |
| Meg3 | 3.3725424 | 0.037787698 | 0.303790456 | 0.536777274 | 0.591421472 |
| Mt1 | 3.035079 | -0.052317974 | 0.285582805 | -0.958135015 | 0.337994681 |
| Mt3 | 4.3413577 | -0.063469682 | 0.287089464 | -1.099952103 | 0.271352991 |
| Ninj2 | 4.790965 | 0.048022598 | 0.281976596 | 0.476333671 | 0.633836681 |
| Nkain2 | 3.9198008 | -0.144269504 | 0.401982127 | 2.297197576 | 0.021607501 |
| Pcca | 3.6836677 | -0.029022393 | 0.283958627 | -0.306997931 | 0.758844947 |
| Pcgf6 | 3.4580526 | -0.144269504 | 0.401982127 | 0.638100935 | 0.523407978 |
| Phkg1 | 4.360975 | -0.144269504 | 0.401982119 | 0.915435041 | 0.35996329 |
| Plp1 | 2.3849928 | -0.02967227 | 0.267686089 | -1.0076854 | 0.313605516 |
| Ppia | 1.9710991 | -0.034164357 | 0.285787381 | -0.796812754 | 0.425559787 |
| Ppp1r9a | 4.709364 | -0.004378392 | 0.281770177 | -0.054546134 | 0.956500054 |
| Ppp3ca | 1.3173085 | -0.004583246 | 0.317585949 | -0.071162709 | 0.94326826 |
| Prkag2 | 4.1486835 | -0.144269504 | 0.401982124 | 1.181573624 | 0.237374924 |
| Ptprd | 2.9578784 | 0.033343678 | 0.305694278 | 0.479349028 | 0.631690349 |
| Ptprg | 2.6142354 | -0.020610986 | 0.296127166 | -0.239778394 | 0.810502057 |
| Rab29 | 4.6720915 | 0.038705993 | 0.28650184 | 0.442768178 | 0.657933423 |
| Ralyl | 3.207875 | -0.144269504 | 0.401982125 | -0.108733148 | 0.913414149 |
| Rpl11 | 4.7606373 | -0.051730688 | 0.273163012 | -1.261974033 | 0.20695813 |
| Rpl12 | 2.0700948 | -0.003113712 | 0.304121065 | -0.05192631 | 0.95858741 |
| Rpl13a | 1.4650565 | -0.013193672 | 0.30869673 | -0.238375573 | 0.811589811 |
| Rpl17 | 2.507469 | -0.175729444 | 0.387761561 | -2.302693619 | 0.021296087 |
| Rpl21 | 3.447512 | -0.157123908 | 0.348390674 | -2.299118816 | 0.021498193 |
| Rpl23 | 3.8133543 | -0.052797833 | 0.273187254 | -1.4441956 | 0.148683962 |
| Rpl27a | 4.768087 | -0.116497605 | 0.314327818 | -1.562963394 | 0.118061205 |
| Rpl30 | 4.705174 | -0.019477988 | 0.281798752 | -0.382872792 | 0.701814086 |
| Rpl32 | 3.3037553 | -0.106804537 | 0.302435192 | -2.026048724 | 0.042759792 |
| Rpl34 | 1.8285625 | -0.033646369 | 0.284876026 | -0.819424665 | 0.412544167 |
| Rpl35a | 4.942784 | -0.070557535 | 0.275782045 | -1.752521031 | 0.079684257 |
| Rpl36 | 0.8958757 | -0.00816249 | 0.282098335 | -0.308404547 | 0.757774523 |
| Rpl37a | 4.900374 | -0.027974154 | 0.283810001 | -0.460160376 | 0.64540111 |
| Rpl39 | 2.3683107 | -0.075302968 | 0.285662603 | -1.584309299 | 0.113123351 |
| Rpl41 | 4.4877243 | -0.082407799 | 0.292100419 | -1.284244285 | 0.199056494 |
| Rpl9 | 3.0056663 | -2.282100577 | 1.047588828 | -3.172111957 | 0.001513347 |
| Rps10 | 1.7781105 | -0.101896029 | 0.316444112 | -1.545320807 | 0.122268685 |
| Rps12 | 3.9755926 | -0.066728106 | 0.271797774 | -1.852001615 | 0.064025591 |
| Rps14 | 3.1900764 | -0.023876734 | 0.268765167 | -0.821141579 | 0.411565629 |

| | | | | | |
|---|---|---|---|---|---|
| Rps15 | 4.6742244 | -0.078364992 | 0.284098828 | -1.525158405 | 0.127219593 |
| Rps16 | 2.7770374 | -0.042076303 | 0.283776642 | -0.814090284 | 0.415593231 |
| Rps17 | 1.9367694 | -0.002264434 | 0.298763451 | -0.055080572 | 0.956074274 |
| Rps20 | 2.5459812 | -0.071487078 | 0.280669476 | -1.830923711 | 0.067111931 |
| Rps23 | 1.558236 | -0.027025796 | 0.281939238 | -0.677207259 | 0.498274463 |
| Rps24 | 3.1835344 | -0.081112486 | 0.295168685 | -1.278026806 | 0.201239975 |
| Rps27 | 2.3981893 | -0.109433149 | 0.326608718 | -1.446496498 | 0.148037994 |
| Rps27a | 2.27826 | -0.043247087 | 0.29320298 | -0.648413677 | 0.516717427 |
| Rps28 | 4.243435 | -0.066642629 | 0.29029014 | -0.903955203 | 0.366019155 |
| Rps3 | 1.8366036 | -0.042346954 | 0.279619542 | -1.102204659 | 0.270372707 |
| Rps7 | 1.746049 | -0.031915545 | 0.271240634 | -1.018369739 | 0.308502278 |
| Rps8 | 4.412556 | -0.032138484 | 0.276937672 | -0.721894809 | 0.470359153 |
| Scamp1 | 4.1156707 | -0.144269504 | 0.401982125 | 1.361054985 | 0.173496314 |
| Selenow | 2.1228044 | -0.052192174 | 0.29835974 | -0.73929132 | 0.459730119 |
| Sgip1 | 1.58918 | -0.02754913 | 0.306732397 | -0.474418159 | 0.635201747 |
| Spock1 | 3.860509 | -0.144269504 | 0.401982122 | -1.018349138 | 0.308512064 |
| Srgn | 2.2238126 | -0.01253748 | 0.300111863 | -0.199617632 | 0.841779637 |
| St18 | 2.3145986 | 0.156173025 | 0.379149245 | 1.504930809 | 0.132341867 |
| Sult2a2 | 2.8649654 | -0.144269504 | 0.401982125 | 0.181391926 | 0.856059956 |
| Svil | 2.9297297 | -0.144269504 | 0.401982126 | 0.478025175 | 0.632632287 |
| Syt1 | 2.76881 | -0.035342887 | 0.289706383 | -0.418017684 | 0.675934188 |
| Tacc1 | 1.8357242 | 0.032747705 | 0.313710961 | 0.428819373 | 0.668054679 |
| Tmeff2 | 3.7606382 | -0.07426627 | 0.296965094 | -1.063634328 | 0.287494385 |
| Tmem108 | 3.9934008 | 0.095522483 | 0.302111567 | 0.891489007 | 0.372666887 |
| Tmsb4x | 4.6209717 | -0.89304218 | 0.859107653 | -2.672477172 | 0.00752935 |
| Tns3 | 2.8131933 | 0.065163948 | 0.311282623 | 0.728422088 | 0.466355246 |
| Tpt1 | 4.37262 | -0.047621375 | 0.260565615 | -1.931898729 | 0.053372007 |
| Uba52 | 1.9512028 | -0.092858149 | 0.323991347 | -1.387318579 | 0.16534464 |
| Ubb | 1.2982892 | -0.058494176 | 0.287929686 | -1.271654457 | 0.203495919 |
| Uqcrh | 1.8911818 | -0.093511442 | 0.313578872 | -1.47515932 | 0.140169715 |
| Vit | 1.6428826 | 0.106730787 | 0.362252785 | 1.305143395 | 0.191844025 |
| Vwc2l | 4.158543 | 0.019439266 | 0.287145756 | 0.216752222 | 0.828401448 |
| Zfhx3 | 3.9178238 | -0.036017681 | 0.286116561 | -0.397187422 | 0.691229257 |
| Zfp994 | 3.931868 | 0.048526399 | 0.297754268 | 0.569252529 | 0.569184777 |
| mt-Nd4l | 1.7701625 | -0.021892343 | 0.273709143 | -0.777157424 | 0.437065898 |

| **Glut** | baseMean | log2FoldChange | lfcSE | stat | pvalue | padj | pglobadj |
|---|---|---|---|---|---|---|---|
| Cpe | 18.960005 | -1.775914494 | 0.266985338 | -6.858679744 | 6.95E-12 | 1.58E-09 | 1.58E-08 |
| Mbp | 64.762535 | -2.353230573 | 0.442840443 | -5.817058327 | 5.99E-09 | 5.35E-07 | 5.35E-06 |
| Fth1 | 159.25063 | -1.943232652 | 0.364716513 | -5.789088916 | 7.08E-09 | 5.35E-07 | 5.35E-06 |
| Mc1r | 36.4037 | -0.144269504 | 0.59534174 | 5.505139413 | 3.69E-08 | 2.09E-06 | 2.09E-05 |
| Apoe | 80.2238 | -1.546311713 | 0.329109342 | -5.123833412 | 2.99E-07 | 1.36E-05 | 0.000135921 |
| 5830416I1 | 109.243774 | 0.566466813 | 0.117734137 | 4.913538734 | 8.94E-07 | 3.38E-05 | 0.000338408 |
| Hexb | 15.088464 | -2.212126058 | 0.505333313 | -4.873755614 | 1.09E-06 | 3.55E-05 | 0.000355081 |
| mt-Co3 | 317.89905 | -1.079382138 | 0.238768558 | -4.835361746 | 1.33E-06 | 3.77E-05 | 0.000377114 |
| Rsrc1 | 41.97339 | -0.144269504 | 0.595341713 | 4.761458641 | 1.92E-06 | 4.36E-05 | 0.000436291 |
| mt-Co2 | 179.1176 | -0.975562238 | 0.216803496 | -4.776038515 | 1.79E-06 | 4.36E-05 | 0.000436291 |
| Cox7c | 24.603357 | -1.553778156 | 0.362751108 | -4.70141795 | 2.58E-06 | 5.33E-05 | 0.000533163 |
| Hapln1 | 12.383218 | -0.144269504 | 0.595341777 | 4.664828947 | 3.09E-06 | 5.84E-05 | 0.000584286 |
| Cox6b1 | 15.940308 | -2.521586243 | 0.628035125 | -4.60411558 | 4.14E-06 | 7.23E-05 | 0.000723296 |
| Tpt1 | 28.210365 | -1.755713479 | 0.431831195 | -4.578332379 | 4.69E-06 | 7.60E-05 | 0.00075996 |
| Mt3 | 35.23202 | -1.90194458 | 0.490081286 | -4.47171087 | 7.76E-06 | 0.000117429 | 0.001174291 |
| Rpl12 | 12.270738 | -2.011576633 | 0.510758905 | -4.432649394 | 9.31E-06 | 0.00013206 | 0.001320603 |
| Rps20 | 30.235338 | -1.794499991 | 0.474260972 | -4.349186653 | 1.37E-05 | 0.000182459 | 0.001824591 |
| mt-Nd4l | 10.59298 | -1.382815956 | 0.349236633 | -4.289416925 | 1.79E-05 | 0.000214029 | 0.002140285 |
| Rps27a | 33.156513 | -1.977743932 | 0.542129203 | -4.294931172 | 1.75E-05 | 0.000214029 | 0.002140285 |
| Ppia | 17.00175 | -1.470577863 | 0.384884147 | -4.260719205 | 2.04E-05 | 0.000231279 | 0.002312791 |
| Eef1a1 | 22.886635 | -1.273082844 | 0.335392794 | -4.203513587 | 2.63E-05 | 0.000259375 | 0.002593755 |
| Cox4i1 | 55.03479 | -2.009401569 | 0.572506845 | -4.205356052 | 2.61E-05 | 0.000259375 | 0.002593755 |
| Cox8a | 54.110287 | -2.149150087 | 0.618205278 | -4.208593043 | 2.57E-05 | 0.000259375 | 0.002593755 |
| Rplp1 | 76.570724 | -1.596063677 | 0.444498378 | -4.167040358 | 3.09E-05 | 0.000291865 | 0.002918651 |
| Rpl39 | 28.428679 | -1.976949829 | 0.567781774 | -4.15600116 | 3.24E-05 | 0.00029407 | 0.002940705 |
| Cox6c | 36.238724 | -1.844309215 | 0.563518915 | -3.964527475 | 7.35E-05 | 0.000638635 | 0.006386346 |
| Rpl37 | 58.321743 | -1.505857884 | 0.446979996 | -3.949226986 | 7.84E-05 | 0.000638635 | 0.006386346 |
| mt-Nd5 | 23.92398 | -1.35204553 | 0.391252686 | -3.939687914 | 8.16E-05 | 0.000638635 | 0.006386346 |

| | | | | | | | |
|---|---|---|---|---|---|---|---|
| mt-Co1 | 163.52351 | -1.070775207 | 0.301298412 | -3.945997249 | 7.95E-05 | 0.000638635 | 0.006386346 |
| Rpl21 | 37.689503 | -1.594996878 | 0.483001239 | -3.9185885 | 8.91E-05 | 0.000673956 | 0.006739556 |
| Tmsb4x | 42.59606 | -1.618957759 | 0.504798153 | -3.851569972 | 0.000117363 | 0.0008594 | 0.008593999 |
| Rpl17 | 26.852716 | -1.821574666 | 0.596862251 | -3.772007814 | 0.000161939 | 0.001148756 | 0.011487561 |
| Uqcrh | 17.226807 | -2.01444825 | 0.674245636 | -3.746721088 | 0.000179161 | 0.001232411 | 0.012324112 |
| Xlr5a | 95.355095 | 0.546542716 | 0.151810277 | 3.710101507 | 0.000207176 | 0.001383206 | 0.013832056 |
| Car12 | 13.269348 | -1.814461759 | 0.606835567 | -3.678467604 | 0.000234639 | 0.001439545 | 0.01439545 |
| Rps24 | 50.57764 | -1.862548786 | 0.647313688 | -3.683095246 | 0.000230419 | 0.001439545 | 0.01439545 |
| Calm1 | 23.933855 | -1.215862466 | 0.378365962 | -3.689241838 | 0.000224923 | 0.001439545 | 0.01439545 |
| Aldoc | 15.485122 | -1.570094588 | 0.511680435 | -3.664563894 | 0.00024776 | 0.001480042 | 0.014800423 |
| Ubb | 13.352551 | -1.295483413 | 0.411426701 | -3.624197075 | 0.000289861 | 0.001687137 | 0.016871369 |
| Rpl23 | 26.072792 | -1.688741778 | 0.583082052 | -3.611780725 | 0.000304102 | 0.001725777 | 0.017257769 |
| Rps14 | 37.294437 | -1.486707194 | 0.503787895 | -3.596934058 | 0.00032199 | 0.001782726 | 0.017827255 |
| Plp1 | 11.947595 | -1.996243089 | 0.713997765 | -3.578596374 | 0.000345444 | 0.001867045 | 0.018670452 |
| Rpl9 | 27.202162 | -1.715525686 | 0.612758911 | -3.552520362 | 0.000381559 | 0.002014279 | 0.020142792 |
| Rps23 | 25.459724 | -1.284124722 | 0.427841841 | -3.541801684 | 0.000397404 | 0.002050244 | 0.02050244 |
| Cst3 | 153.94177 | -1.770866161 | 0.671842032 | -3.498880761 | 0.000467215 | 0.002356842 | 0.023568425 |
| 6030443J0 | 30.79876 | -0.144269504 | 0.595341702 | 3.484249156 | 0.00049352 | 0.002383596 | 0.023835958 |
| Rpl26 | 34.769566 | -1.480451693 | 0.526811028 | -3.485144962 | 0.00049187 | 0.002383596 | 0.023835958 |
| Adamtsl1 | 32.24669 | -0.144269504 | 0.595341746 | 3.464215679 | 0.00053178 | 0.002514876 | 0.025148761 |
| mt-Nd4 | 90.86455 | -1.030302467 | 0.341879868 | -3.456507315 | 0.000547224 | 0.002535101 | 0.025351006 |
| Rps3a1 | 29.006617 | -1.338055263 | 0.4698055 | -3.442311225 | 0.000576766 | 0.002618519 | 0.026185188 |
| Rpl37a | 40.960964 | -1.547820947 | 0.584889288 | -3.402997396 | 0.000666509 | 0.002789039 | 0.027890388 |
| mt-Nd1 | 66.879105 | -0.865056969 | 0.282567981 | -3.39672203 | 0.000681982 | 0.002789039 | 0.027890388 |
| Rpl30 | 34.068516 | -1.443648474 | 0.527697122 | -3.417585197 | 0.000631793 | 0.002789039 | 0.027890388 |
| Rpl27a | 34.65671 | -1.400948186 | 0.509294354 | -3.407168971 | 0.000656405 | 0.002789039 | 0.027890388 |
| Rapgef4 | 48.983097 | 0.449432859 | 0.135656251 | 3.396641475 | 0.000682183 | 0.002789039 | 0.027890388 |
| Zbtb7a | 23.56369 | 0.540259052 | 0.163854073 | 3.394299185 | 0.000688045 | 0.002789039 | 0.027890388 |
| Dpyd | 15.7993355 | -0.144269504 | 0.59534175 | 3.385679672 | 0.000710022 | 0.002827632 | 0.028276319 |
| Prkg1 | 32.408478 | -0.144269504 | 0.595341743 | 3.365322438 | 0.000764543 | 0.002941545 | 0.029415452 |
| Rpl13 | 48.121593 | -1.739300377 | 0.701177959 | -3.365470013 | 0.000764134 | 0.002941545 | 0.029415452 |
| mt-Nd3 | 53.05994 | -0.923558787 | 0.313082642 | -3.333166183 | 0.000858636 | 0.003195253 | 0.031952535 |
| Rps8 | 42.907177 | -1.515333677 | 0.588199643 | -3.33577978 | 0.000850605 | 0.003195253 | 0.031952535 |
| Galnt13 | 29.213446 | 0.500227212 | 0.156245917 | 3.310857097 | 0.000930107 | 0.003405392 | 0.034053915 |
| Uba52 | 9.810962 | -1.302528435 | 0.486356303 | -3.234290696 | 0.001219453 | 0.004393901 | 0.043939009 |
| Rps12 | 35.98471 | -1.800145893 | 0.820133495 | -3.203430797 | 0.001358007 | 0.004796096 | 0.04796096 |
| Rps10 | 27.12138 | -1.398911005 | 0.566129133 | -3.194248241 | 0.001401955 | 0.004796096 | 0.04796096 |
| Rps15 | 19.44779 | -1.124112757 | 0.418032182 | -3.198278716 | 0.001382506 | 0.004796096 | 0.04796096 |
| Rpl36 | 19.816837 | -1.408905449 | 0.568863735 | -3.191453736 | 0.001415588 | 0.004796096 | 0.04796096 |
| Rps29 | 86.63813 | -1.107704316 | 0.421421911 | -3.180009372 | 0.001472703 | 0.004916229 | 0.049162288 |
| mt-Atp6 | 255.12491 | -0.8678506 | 0.310947716 | -3.17193214 | 0.001514284 | 0.004941086 | 0.04941086 |
| Shb | 57.12811 | 0.349715972 | 0.112354461 | 3.170134688 | 0.001523683 | 0.004941086 | 0.04941086 |
| Rpl35a | 37.172855 | -1.378399092 | 0.573440075 | -3.147098919 | 0.001648992 | 0.005272128 | 0.052721282 |
| Rps7 | 21.615023 | -1.432674725 | 0.624522402 | -3.079067692 | 0.002076495 | 0.006546727 | 0.065467271 |
| Rpl32 | 26.072998 | -1.129711992 | 0.452596319 | -3.066494438 | 0.002165848 | 0.006734898 | 0.067348984 |
| Lrp1b | 204.00333 | 0.407658423 | 0.137805243 | 3.036323762 | 0.002394821 | 0.007346274 | 0.073462741 |
| 4930511M | 35.85655 | 0.479828052 | 0.166172797 | 3.0108768 | 0.002604945 | 0.007875162 | 0.07875162 |
| Gm42418 | 495.07285 | -0.921359088 | 0.357996054 | -3.007205095 | 0.002636618 | 0.007875162 | 0.07875162 |
| Ms4a1 | 27.41386 | 0.556227817 | 0.196669509 | 2.993722349 | 0.002755966 | 0.008124731 | 0.081247314 |
| Gria4 | 27.528088 | 0.432246629 | 0.151180023 | 2.97218351 | 0.002956899 | 0.008605334 | 0.08605334 |
| mt-Nd2 | 100.98584 | -0.732392454 | 0.283296891 | -2.899867667 | 0.003733202 | 0.01072705 | 0.107270498 |
| Fau | 36.68445 | -1.575844484 | 0.905642648 | -2.874884835 | 0.004041749 | 0.011468463 | 0.114684625 |
| Ntm | 60.91025 | 0.427339914 | 0.15614548 | 2.847609161 | 0.004404898 | 0.012344592 | 0.12344592 |
| Tshz2 | 4.0018873 | 1.101808747 | 0.465936715 | 2.837170876 | 0.004551525 | 0.012599952 | 0.125999522 |
| Dbi | 18.818296 | -0.979446638 | 0.419498618 | -2.833016291 | 0.004611104 | 0.012611091 | 0.126110906 |
| Rps17 | 12.002829 | -1.017570769 | 0.439700598 | -2.826990597 | 0.004698771 | 0.012697869 | 0.126978686 |
| Rpl11 | 34.706364 | -1.240334601 | 0.612216643 | -2.818583929 | 0.0048236 | 0.012775165 | 0.127751652 |
| Ckb | 26.128765 | -0.934897669 | 0.400908623 | -2.813784186 | 0.004896209 | 0.012775165 | 0.127751652 |
| Mt1 | 35.063984 | -1.044540801 | 0.471280722 | -2.816469005 | 0.004855473 | 0.012775165 | 0.127751652 |
| Rpl13a | 23.156572 | -1.240535703 | 0.614675545 | -2.804719004 | 0.005036046 | 0.01299071 | 0.129907102 |
| Slx1b | 25.369318 | 0.535105135 | 0.203369072 | 2.79378959 | 0.005209436 | 0.013286989 | 0.13286989 |
| Cmss1 | 56.48602 | -0.973564682 | 0.436911563 | -2.779262395 | 0.00544825 | 0.013741696 | 0.137416961 |
| AY036118 | 96.25058 | -1.221750811 | 0.632579192 | -2.767033018 | 0.005656903 | 0.014111175 | 0.141111745 |
| Ninj2 | 17.927513 | -0.144269504 | 0.595341715 | 2.736064169 | 0.00621789 | 0.015341968 | 0.153419676 |
| Rpl38 | 36.311028 | -1.027452061 | 0.484798011 | -2.732315747 | 0.006289084 | 0.015350774 | 0.153507743 |

| | | | | | | | |
|---|---|---|---|---|---|---|---|
| Rps27 | 24.597618 | -0.746925873 | 0.318352047 | -2.6886532 | 0.007174091 | 0.017324666 | 0.173246661 |
| Gas5 | 23.922663 | -1.248182874 | 0.699855648 | -2.678893404 | 0.00738659 | 0.017650064 | 0.176500636 |
| Camk1d | 70.181915 | -1.174990314 | 0.662064898 | -2.640710483 | 0.008273238 | 0.019361084 | 0.193610837 |
| Hdac8 | 13.568848 | 0.639925371 | 0.266213405 | 2.642305559 | 0.008234373 | 0.019361084 | 0.193610837 |
| Kctd16 | 76.70161 | 0.330842693 | 0.129555784 | 2.618205281 | 0.008839361 | 0.020474847 | 0.204748467 |
| Gphn | 15.688767 | -1.00574733 | 0.503163789 | -2.61004908 | 0.009052923 | 0.020563378 | 0.205633784 |
| 8430422H( | 42.030968 | 0.493428664 | 0.19981204 | 2.609828706 | 0.009058757 | 0.020563378 | 0.205633784 |
| mt-Cytb | 99.99927 | -0.801359808 | 0.371731195 | -2.593066855 | 0.009512428 | 0.021379417 | 0.213794168 |
| Selenow | 17.175367 | -1.050312116 | 0.609329486 | -2.481394011 | 0.013086962 | 0.029124906 | 0.291249064 |
| Cntnap2 | 56.53396 | -0.144269504 | 0.595341708 | 2.469886783 | 0.013515582 | 0.029786768 | 0.297867681 |
| Rps3 | 19.521986 | -1.026707495 | 0.607126251 | -2.443514874 | 0.01454497 | 0.03144484 | 0.3144484 |
| Gm10062 | 59.852055 | 0.484623199 | 0.210546548 | 2.446395261 | 0.014429272 | 0.03144484 | 0.3144484 |
| Nckap5 | 94.96033 | 0.429174253 | 0.186064193 | 2.417997767 | 0.015606173 | 0.033420766 | 0.334207658 |
| Spock1 | 12.781663 | -0.144269504 | 0.595341762 | 2.401106435 | 0.016345581 | 0.034677075 | 0.346770746 |
| Ctnna2 | 13.414483 | 0.588800979 | 0.275221588 | 2.37779282 | 0.017416608 | 0.036607131 | 0.366071308 |
| Anks1b | 9.166362 | -0.685511918 | 0.337696842 | -2.340280119 | 0.019269281 | 0.040129604 | 0.401296039 |
| Rpl41 | 37.511967 | -0.897807616 | 0.54446644 | -2.315208686 | 0.020601514 | 0.042514034 | 0.425140337 |
| Scamp1 | 17.284771 | 0.498170001 | 0.23396896 | 2.308242694 | 0.020985642 | 0.042916583 | 0.429165828 |
| Vit | 14.305894 | -0.144269504 | 0.595341786 | 2.279854819 | 0.0226163 | 0.045838395 | 0.458383947 |
| Rpl34 | 22.53576 | -0.793967554 | 0.48556312 | -2.190592637 | 0.028481284 | 0.057214616 | 0.572146156 |
| Cmpk1 | 49.48254 | 0.349307274 | 0.16770299 | 2.174272389 | 0.029684685 | 0.059108978 | 0.591089775 |
| Gm12216 | 92.376305 | 0.276142044 | 0.131353094 | 2.155774235 | 0.031101298 | 0.061391258 | 0.613912583 |
| Ppp3ca | 11.9548855 | -0.752653358 | 0.456821883 | -2.138437886 | 0.03248122 | 0.063562388 | 0.635623882 |
| Gm50306 | 32.413666 | -0.144269504 | 0.595341754 | 2.128834072 | 0.033267991 | 0.064545589 | 0.645455892 |
| Rps21 | 69.29485 | -0.827885883 | 0.598030253 | -2.108067025 | 0.035025192 | 0.067378972 | 0.673789716 |
| Epha6 | 38.557938 | 0.408854611 | 0.208173659 | 2.092177984 | 0.036422597 | 0.0694784 | 0.694783996 |
| Trim2 | 42.692062 | 0.355919528 | 0.180343244 | 2.075152724 | 0.037972383 | 0.07183109 | 0.718310905 |
| Igf2bp3 | 45.83929 | 0.469015742 | 0.24710392 | 2.065570964 | 0.038869008 | 0.072919545 | 0.729195447 |
| Svil | 12.132911 | 0.492091603 | 0.27218217 | 2.015196552 | 0.043884059 | 0.081653126 | 0.816531255 |
| Ppp1r9a | 15.939527 | 0.48954594 | 0.272466253 | 2.003283469 | 0.045146871 | 0.083319835 | 0.833198347 |
| Rps16 | 24.292126 | -0.760042479 | 0.583842482 | -1.971788427 | 0.048633764 | 0.089031165 | 0.890311649 |
| Gm11337 | 13.186041 | 0.447387606 | 0.252823463 | 1.951195521 | 0.051033792 | 0.092677366 | 0.926773655 |
| Atp9b | 22.188927 | 0.48373726 | 0.289372954 | 1.881295046 | 0.059931795 | 0.10797236 | 1.0797236 |
| Fbxl7 | 16.057865 | 0.525478397 | 0.324115908 | 1.870687948 | 0.061388346 | 0.109725627 | 1.097256267 |
| Sil1 | 27.170454 | 0.357009505 | 0.205030989 | 1.858071874 | 0.063158808 | 0.112008199 | 1.12008199 |
| Dctd | 7.827331 | -0.144269504 | 0.595341797 | 1.819298347 | 0.068865925 | 0.11963167 | 1.1963167 |
| Gm49497 | 13.54824 | 0.480696492 | 0.300942701 | 1.820182089 | 0.06873128 | 0.11963167 | 1.1963167 |
| Gm29325 | 14.84829 | 0.399411065 | 0.240260664 | 1.818167466 | 0.069038541 | 0.11963167 | 1.1963167 |
| Acoxl | 21.082848 | 0.393155993 | 0.235636159 | 1.814510838 | 0.069599116 | 0.119689388 | 1.196893883 |
| 4930558K( | 19.8751 | 0.358795482 | 0.213855892 | 1.808834102 | 0.070476781 | 0.120097012 | 1.200970118 |
| Tmem108 | 19.81278 | 0.373402101 | 0.223506533 | 1.806154107 | 0.070894271 | 0.120097012 | 1.200970118 |
| Cspp1 | 15.578925 | 0.433046301 | 0.269285983 | 1.793505795 | 0.072892085 | 0.122566691 | 1.225666909 |
| Pde1c | 48.006958 | 0.410469952 | 0.255427794 | 1.760063479 | 0.078397044 | 0.129371944 | 1.293719443 |
| Sema3c | 66.135925 | 0.356451923 | 0.216534866 | 1.758579112 | 0.078649023 | 0.129371944 | 1.293719443 |
| Vwc2l | 9.199624 | -0.144269504 | 0.595341803 | 1.762720812 | 0.077947589 | 0.129371944 | 1.293719443 |
| Chsy3 | 23.052742 | 0.331646876 | 0.204841999 | 1.729646442 | 0.083693463 | 0.136679253 | 1.366792526 |
| 1700054A( | 141.57812 | 0.251022394 | 0.152054244 | 1.707080696 | 0.087807055 | 0.141789697 | 1.417896966 |
| Atxn7l1 | 12.299738 | 0.425542841 | 0.281949142 | 1.70565671 | 0.088072014 | 0.141789697 | 1.417896966 |
| Ehbp1 | 37.233177 | 0.28678133 | 0.177303038 | 1.698025832 | 0.089502887 | 0.143078558 | 1.43078558 |
| Prkag2 | 18.591381 | -0.144269504 | 0.595341774 | 1.664340053 | 0.096044492 | 0.152462236 | 1.524622362 |
| Ctnna3 | 24.259981 | 0.354980111 | 0.235639471 | 1.64061978 | 0.100876366 | 0.159020383 | 1.590203832 |
| Adk | 15.99909 | -0.144269504 | 0.595341786 | 1.60753086 | 0.107937965 | 0.168978746 | 1.689787458 |
| Ctcflos | 16.329626 | 0.454121363 | 0.347182911 | 1.551300325 | 0.120829729 | 0.184082876 | 1.840828756 |
| Ttc28 | 27.770508 | 0.225521349 | 0.152063064 | 1.557294063 | 0.119400684 | 0.184082876 | 1.840828756 |
| Ppp1r16a | 14.606436 | -0.144269504 | 0.595341764 | 1.555556761 | 0.119813527 | 0.184082876 | 1.840828756 |
| Arvcf | 7.0802913 | -0.144269504 | 0.595341801 | -1.551448548 | 0.120794229 | 0.184082876 | 1.840828756 |
| Syt1 | 12.191012 | -0.533004752 | 0.499894529 | -1.522381923 | 0.127913412 | 0.19357563 | 1.935756296 |
| Csmd3 | 26.341698 | 0.302679691 | 0.215881742 | 1.506763129 | 0.131871383 | 0.198243735 | 1.982437355 |
| Grik1 | 5.4989185 | 0.534153497 | 0.491535025 | 1.488881105 | 0.13651868 | 0.203879871 | 2.038798706 |
| Tns3 | 11.983182 | 0.408625622 | 0.329422894 | 1.455082764 | 0.14564635 | 0.216089683 | 2.160896831 |
| Pcdh15 | 27.633442 | -0.144269504 | 0.595341752 | 1.447539117 | 0.147745989 | 0.217781425 | 2.177814251 |
| Ptprd | 14.081265 | 0.353433724 | 0.278471399 | 1.4290403 | 0.152992651 | 0.224060205 | 2.240602045 |
| Alpl | 10.492561 | 0.345764033 | 0.276429974 | 1.416103079 | 0.156745327 | 0.226631778 | 2.266317781 |
| Zmat4 | 23.312378 | 0.26196878 | 0.196288105 | 1.418684889 | 0.155990905 | 0.226631778 | 2.266317781 |
| Gm14582 | 12.835498 | 0.381079863 | 0.314528912 | 1.408108299 | 0.159099004 | 0.228578949 | 2.285789487 |

| | | | | | | | |
|---|---|---|---|---|---|---|---|
| Gnaz | 44.17836 | 0.289107101 | 0.222262287 | 1.399146247 | 0.161769132 | 0.230953415 | 2.309534148 |
| Mapk14 | 41.72899 | 0.253787666 | 0.199467246 | 1.34819417 | 0.177595939 | 0.251964239 | 2.519642389 |
| Lyrm1 | 7.9197645 | -0.144269504 | 0.595341797 | 1.321612454 | 0.186297234 | 0.261192252 | 2.611922518 |
| Nell1 | 23.617365 | 0.270226207 | 0.220930334 | 1.321299505 | 0.186401519 | 0.261192252 | 2.611922518 |
| Cdk8 | 18.143866 | -0.44907874 | 0.508884166 | -1.306804577 | 0.19127909 | 0.266382537 | 2.66382537 |
| Wdr72 | 23.826147 | 0.323991178 | 0.281252256 | 1.296464428 | 0.194815528 | 0.2696532 | 2.696531998 |
| Pcca | 11.302964 | 0.333616509 | 0.295461849 | 1.292678341 | 0.196122332 | 0.269816784 | 2.698167835 |
| Il27 | 31.86366 | 0.256358345 | 0.215291188 | 1.278617564 | 0.201031762 | 0.274904879 | 2.74904879 |
| Sult2a2 | 10.396637 | 0.317491453 | 0.284904399 | 1.268192148 | 0.20472934 | 0.278284792 | 2.782847921 |
| Phkg1 | 13.941239 | 0.257887806 | 0.22144636 | 1.262758702 | 0.206675911 | 0.279258522 | 2.79258522 |
| 4930441J1 | 14.087247 | 0.317834471 | 0.292329609 | 1.240250938 | 0.214882593 | 0.288629282 | 2.886292816 |
| Wdyhv1 | 16.539324 | 0.34609347 | 0.336400648 | 1.213804003 | 0.224822577 | 0.300204265 | 3.002042648 |
| AI314278 | 10.483916 | -0.144269504 | 0.595341794 | 1.205125515 | 0.228154851 | 0.302872229 | 3.028722292 |
| Zfhx3 | 14.494242 | 0.285747885 | 0.279669142 | 1.155298793 | 0.24796809 | 0.325478204 | 3.254782035 |
| Dlg2 | 14.809554 | -0.394910198 | 0.571569569 | -1.151599966 | 0.249485495 | 0.325478204 | 3.254782035 |
| Cadm2 | 11.891537 | -0.363654686 | 0.411497223 | -1.152742396 | 0.249016134 | 0.325478204 | 3.254782035 |
| Pde6a | 63.212955 | 0.178892461 | 0.162697662 | 1.144500155 | 0.252416274 | 0.327419967 | 3.274199665 |
| Esf1 | 9.794302 | 0.295067056 | 0.309151925 | 1.111108558 | 0.266521624 | 0.343752322 | 3.437523225 |
| Cdh23 | 31.393366 | 0.160397804 | 0.150744424 | 1.100019867 | 0.271323466 | 0.347968513 | 3.479685126 |
| Klkb1 | 12.379919 | 0.269443016 | 0.298610549 | 1.037417891 | 0.299541144 | 0.381999099 | 3.819990994 |
| Rps28 | 25.47918 | -0.341941238 | 0.48862034 | -1.027487432 | 0.304191001 | 0.385761772 | 3.857617717 |
| St18 | 5.592503 | 0.318257226 | 0.422422353 | 0.987084202 | 0.323601362 | 0.408097273 | 4.080972731 |
| Hdac9 | 9.57025 | -0.144269504 | 0.595341776 | 0.943774189 | 0.345285053 | 0.433037056 | 4.330370557 |
| Kcnip4 | 17.83586 | 0.236456858 | 0.292781474 | 0.925257506 | 0.354831977 | 0.442565158 | 4.425651583 |
| Dop1b | 23.20367 | 0.221445191 | 0.287388467 | 0.872561853 | 0.382901941 | 0.470288739 | 4.702887386 |
| Chl1 | 13.870078 | 0.205546799 | 0.263956146 | 0.871877951 | 0.383274963 | 0.470288739 | 4.702887386 |
| 1700126G0 | 34.365047 | 0.147209082 | 0.176941801 | 0.872207173 | 0.383095367 | 0.470288739 | 4.702887386 |
| Nkain2 | 20.22035 | 0.167022329 | 0.215114154 | 0.845883795 | 0.397617573 | 0.485264458 | 4.852644577 |
| Kcnq5 | 13.298319 | 0.208174862 | 0.301907279 | 0.797044633 | 0.42542511 | 0.516425134 | 5.16425134 |
| Ptpn9 | 16.17321 | 0.177892212 | 0.249792715 | 0.786059533 | 0.431832611 | 0.521414908 | 5.214149076 |
| Gm960 | 8.686207 | -0.144269504 | 0.59534174 | 0.775648375 | 0.437956629 | 0.526011401 | 5.260114014 |
| Gm765 | 13.4760685 | 0.160356989 | 0.226348846 | 0.769524796 | 0.44158183 | 0.527574081 | 5.275740812 |
| Mrc2 | 15.52196 | 0.185218059 | 0.276615698 | 0.755902276 | 0.449707789 | 0.534469467 | 5.344694665 |
| Malat1 | 69.80103 | -0.202220725 | 0.370274525 | -0.685772091 | 0.492856838 | 0.580737574 | 5.807375735 |
| Unc5d | 1.6227521 | -0.144269504 | 0.59534181 | 0.684348935 | 0.493754853 | 0.580737574 | 5.807375735 |
| Gria1 | 10.882296 | -0.166970526 | 0.288306108 | -0.665499066 | 0.505731341 | 0.591757806 | 5.917578062 |
| Gnao1 | 10.703454 | -0.179236782 | 0.342598283 | -0.634151432 | 0.525982 | 0.612296995 | 6.12296995 |
| Kat7 | 14.177729 | 0.122612165 | 0.212061257 | 0.622627479 | 0.533529349 | 0.617914093 | 6.179140935 |
| Lsamp | 27.76111 | 0.128655122 | 0.232919549 | 0.604341 | 0.545616953 | 0.628705829 | 6.287058291 |
| Lrrc17 | 20.887321 | 0.191872206 | 0.45284784 | 0.585812883 | 0.558001266 | 0.639728724 | 6.397287237 |
| Lrrc4c | 18.606462 | -0.180722175 | 0.417240885 | -0.580409002 | 0.561638836 | 0.640663396 | 6.406633962 |
| Il1rapl2 | 9.701231 | -0.144269504 | 0.595341778 | 0.566839916 | 0.570822949 | 0.647884047 | 6.478840467 |
| Fam169a | 27.947054 | 0.112395151 | 0.214727907 | 0.561868295 | 0.574205757 | 0.648481128 | 6.48481128 |
| Zfp994 | 14.944137 | 0.118954534 | 0.255625781 | 0.516119852 | 0.605770703 | 0.670780242 | 6.707802424 |
| Srgn | 6.927038 | -0.144269504 | 0.595341796 | 0.528211695 | 0.597352413 | 0.670780242 | 6.707802424 |
| Ankrd12 | 21.009596 | -0.131933303 | 0.289041354 | -0.523264866 | 0.600789947 | 0.670780242 | 6.707802424 |
| Tmeff2 | 11.866141 | -0.113801531 | 0.241416105 | -0.519823517 | 0.603186587 | 0.670780242 | 6.707802424 |
| Ahdc1 | 11.812328 | 0.111298332 | 0.241410998 | 0.507457332 | 0.611833967 | 0.674205391 | 6.742053908 |
| Klhl14 | 2.6490629 | 0.161925833 | 0.561618008 | 0.492556041 | 0.622326315 | 0.682454461 | 6.824544612 |
| Gm38973 | 16.28398 | 0.13491041 | 0.335321253 | 0.479917997 | 0.631285704 | 0.688951225 | 6.889512249 |
| Cep128 | 10.783486 | -0.120027778 | 0.295694162 | -0.469864144 | 0.638452083 | 0.693438387 | 6.934383872 |
| Cfap74 | 13.209162 | -0.140686415 | 0.448962903 | -0.440543709 | 0.659543362 | 0.712934968 | 7.12934968 |
| D430042O0 | 59.267593 | 0.04882702 | 0.106321006 | 0.396034792 | 0.692079362 | 0.744559314 | 7.445593139 |
| Ralyl | 14.507205 | -0.118530664 | 0.408691771 | -0.385675756 | 0.69973682 | 0.749246501 | 7.49246501 |
| Ccdc60 | 15.782949 | 0.096336785 | 0.296922898 | 0.374556571 | 0.707990277 | 0.754524849 | 7.545248488 |
| Camk2a | 20.71487 | -0.070163155 | 0.215306829 | -0.347963274 | 0.727867765 | 0.772084031 | 7.720840313 |
| Meg3 | 25.358404 | -0.101279107 | 0.499642034 | -0.315515783 | 0.752370069 | 0.794362818 | 7.943628175 |
| Pcgf6 | 13.125739 | 0.065566338 | 0.256651942 | 0.284422971 | 0.776086257 | 0.815609169 | 8.156091686 |
| Fyb2 | 22.179024 | -0.053497595 | 0.236521327 | -0.248462741 | 0.803776395 | 0.837057051 | 8.370570508 |
| Kcnd2 | 8.189679 | -0.069066395 | 0.338292289 | -0.24834205 | 0.803869767 | 0.837057051 | 8.370570508 |
| 4930402H2 | 10.878186 | -0.144269504 | 0.595341793 | 0.233796982 | 0.815142593 | 0.844919491 | 8.449194912 |
| Ptprg | 9.615968 | -0.144269504 | 0.595341772 | 0.182944219 | 0.854841786 | 0.882041298 | 8.820412977 |
| Sec61a1 | 171.56808 | 0.020685938 | 0.141880425 | 0.149653004 | 0.881038388 | 0.904957982 | 9.049579818 |
| Rab29 | 17.81273 | 0.030782029 | 0.265047892 | 0.129429339 | 0.897017934 | 0.91722104 | 9.172210403 |
| Sgip1 | 6.526274 | -0.034183323 | 0.368893748 | -0.117914941 | 0.906135054 | 0.922388598 | 9.223885979 |

| | | | | | | | |
|---|---|---|---|---|---|---|---|
| Bc1 | 16.326283 | 0.0310921 | 0.464208831 | 0.095381747 | 0.924011614 | 0.936386769 | 9.363867694 |
| Gm49767 | 11.525636 | -0.022417805 | 0.297357665 | -0.088071537 | 0.929819819 | 0.938084884 | 9.380848839 |
| Cntn3 | 12.88803 | -0.144269504 | 0.595341795 | -0.019746719 | 0.984245421 | 0.984245421 | 9.842454213 |
| Tacc1 | 11.41664 | -0.144269504 | 0.595341778 | -0.023570442 | 0.981195249 | 0.984245421 | 9.842454213 |

**table S10: Differential gene expression list social compared to rest (3) conditions in GABA and Glut neurons**

| **GABA** | baseMean | log2FoldChange | lfcSE | stat | pvalue | padj | pglobadj |
|---|---|---|---|---|---|---|---|
| 1700054A( | 52.824097 | -1.03E-06 | 0.001124862 | -0.360947417 | 0.718138756 | 0.997423128 | 9.974231283 |
| Rpl13 | 5.65667 | -8.97E-07 | 0.001113357 | -0.825558395 | 0.409054644 | 0.997423128 | 9.974231283 |
| Rpl13a | 1.4650565 | -7.40E-07 | 0.001128108 | -0.798832219 | 0.424387706 | 0.997423128 | 9.974231283 |
| Rpl17 | 2.507469 | -3.28E-07 | 0.001125614 | -0.456739257 | 0.647858471 | 0.997423128 | 9.974231283 |
| Rpl21 | 3.447512 | 9.13E-08 | 0.001156609 | 0.106620901 | 0.915089736 | 0.997423128 | 9.974231283 |
| Rpl23 | 3.8133543 | -2.99E-07 | 0.001099821 | -0.543821843 | 0.586564076 | 0.997423128 | 9.974231283 |
| Rpl26 | 6.793167 | -1.38E-06 | 0.001080426 | -1.544185065 | 0.122543506 | 0.997423128 | 9.974231283 |
| Rpl27a | 4.768087 | -1.06E-06 | 0.001100929 | -1.130227151 | 0.258380522 | 0.997423128 | 9.974231283 |
| Rpl30 | 4.705174 | -1.41E-06 | 0.001054536 | -2.077490406 | 0.03775632 | 0.997423128 | 9.974231283 |
| Rpl32 | 3.3037553 | -7.55E-07 | 0.001079106 | -1.193893502 | 0.23251963 | 0.997423128 | 9.974231283 |
| Rpl34 | 1.8285625 | -7.24E-07 | 0.001077649 | -1.26950214 | 0.204262028 | 0.997423128 | 9.974231283 |
| Rpl35a | 4.942784 | -7.04E-07 | 0.001060769 | -1.331260544 | 0.183103296 | 0.997423128 | 9.974231283 |
| Rpl12 | 2.0700948 | -5.33E-07 | 0.001135382 | -0.515994914 | 0.605857961 | 0.997423128 | 9.974231283 |
| Rpl36 | 0.8958757 | 1.41E-07 | 0.001190017 | 0.258488618 | 0.796029832 | 0.997423128 | 9.974231283 |
| Rpl37a | 4.900374 | -3.68E-07 | 0.001119454 | -0.378630091 | 0.704962573 | 0.997423128 | 9.974231283 |
| Rpl38 | 7.104176 | -6.80E-07 | 0.001111074 | -0.684669883 | 0.493552258 | 0.997423128 | 9.974231283 |
| Rpl39 | 2.3683107 | -1.29E-07 | 0.001125588 | -0.187489305 | 0.851277008 | 0.997423128 | 9.974231283 |
| Rpl41 | 4.4877243 | -1.66E-06 | 0.001075313 | -1.904770488 | 0.056809912 | 0.997423128 | 9.974231283 |
| Rpl9 | 3.0056663 | -3.38E-07 | 0.001117621 | -0.480755084 | 0.630690577 | 0.997423128 | 9.974231283 |
| Rplp1 | 7.5951495 | -1.46E-06 | 0.001078676 | -1.559289878 | 0.118927786 | 0.997423128 | 9.974231283 |
| Rps10 | 1.7781105 | 4.42E-08 | 0.00115199 | 0.050398065 | 0.959805179 | 0.997423128 | 9.974231283 |
| Rps12 | 3.9755926 | -5.19E-07 | 0.001064194 | -1.056653211 | 0.290669886 | 0.997423128 | 9.974231283 |
| Rps14 | 3.1900764 | -4.28E-09 | 0.00113748 | -0.042611899 | 0.96601091 | 0.997423128 | 9.974231283 |
| Rps15 | 4.6742244 | -8.45E-07 | 0.001072727 | -1.259973643 | 0.20767887 | 0.997423128 | 9.974231283 |
| Rps17 | 1.9367694 | -5.51E-07 | 0.001084096 | -0.968774378 | 0.332657773 | 0.997423128 | 9.974231283 |
| Rpl37 | 5.0523815 | -4.79E-07 | 0.001118244 | -0.538086756 | 0.590517156 | 0.997423128 | 9.974231283 |
| Rps20 | 2.5459812 | -2.76E-07 | 0.001108349 | -0.497713343 | 0.618686102 | 0.997423128 | 9.974231283 |
| Rpl11 | 4.7606373 | -3.34E-07 | 0.001094495 | -0.557748872 | 0.57701588 | 0.997423128 | 9.974231283 |
| Ralyl | 3.207875 | -4.16E-07 | 0.001139846 | -0.280303352 | 0.779244779 | 0.997423128 | 9.974231283 |
| Mt1 | 3.035079 | 1.17E-06 | 0.001240223 | 1.137242561 | 0.255436899 | 0.997423128 | 9.974231283 |
| Mt3 | 4.3413577 | -2.04E-06 | 0.001069912 | -2.234514625 | 0.025449231 | 0.997423128 | 9.974231283 |
| Nckap5 | 23.137186 | 2.61E-06 | 0.001158707 | 0.862052829 | 0.388658447 | 0.997423128 | 9.974231283 |
| Nell1 | 5.491928 | 5.71E-07 | 0.001146495 | 0.354703335 | 0.722811852 | 0.997423128 | 9.974231283 |
| Ninj2 | 4.790965 | -1.98E-06 | 0.001123206 | -1.163024864 | 0.244819414 | 0.997423128 | 9.974231283 |
| Nkain2 | 3.9198008 | -5.80E-07 | 0.001125653 | -0.429148794 | 0.667814945 | 0.997423128 | 9.974231283 |
| Ntm | 12.9065 | -1.93E-06 | 0.001121684 | -0.846755208 | 0.397131579 | 0.997423128 | 9.974231283 |
| Pcca | 3.6836677 | -5.14E-07 | 0.001138287 | -0.331574439 | 0.740210623 | 0.997423128 | 9.974231283 |
| Pcdh15 | 10.199923 | -8.18E-07 | 0.001138393 | -0.412874021 | 0.679698916 | 0.997423128 | 9.974231283 |
| Pcgf6 | 3.4580526 | -1.67E-06 | 0.001125626 | -1.067851351 | 0.28558757 | 0.997423128 | 9.974231283 |
| Pde1c | 8.5778265 | 1.09E-06 | 0.001145992 | 0.411363153 | 0.680806268 | 0.997423128 | 9.974231283 |
| Rapgef4 | 13.728402 | -0.144269504 | 0.112592779 | -1.061228486 | 0.288586077 | 0.997423128 | 9.974231283 |
| Pde6a | 17.578203 | -6.60E-07 | 0.001136953 | -0.262154861 | 0.793202052 | 0.997423128 | 9.974231283 |
| Plp1 | 2.3849928 | -3.19E-07 | 0.001071896 | -0.763103059 | 0.445401936 | 0.997423128 | 9.974231283 |
| Ppia | 1.9710991 | -5.79E-07 | 0.001089692 | -0.974019257 | 0.330046978 | 0.997423128 | 9.974231283 |
| Ppp1r16a | 5.960048 | -1.91E-07 | 0.001145417 | -0.112209148 | 0.910657576 | 0.997423128 | 9.974231283 |
| Ppp1r9a | 4.709364 | -6.21E-07 | 0.00112072 | -0.50319426 | 0.6148277 | 0.997423128 | 9.974231283 |
| Ppp3ca | 1.3173085 | 2.15E-07 | 0.001151977 | 0.215701845 | 0.829220163 | 0.997423128 | 9.974231283 |
| Prkag2 | 4.1486835 | -2.31E-06 | 0.001119511 | -1.350591638 | 0.176826281 | 0.997423128 | 9.974231283 |
| Prkg1 | 7.6053395 | -1.15E-06 | 0.001123603 | -0.628561352 | 0.529636269 | 0.997423128 | 9.974231283 |
| Ptpn9 | 6.340883 | -0.144269504 | 0.11259278 | -1.164447413 | 0.244242743 | 0.997423128 | 9.974231283 |
| Ptprd | 2.9578784 | 5.88E-08 | 0.001141965 | 0.057259896 | 0.954338167 | 0.997423128 | 9.974231283 |
| Ptprg | 2.6142354 | -0.144269504 | 0.112592781 | 1.204869733 | 0.228253594 | 0.997423128 | 9.974231283 |
| Rab29 | 4.6720915 | -6.13E-07 | 0.001121255 | -0.468587464 | 0.63936454 | 0.997423128 | 9.974231283 |
| Phkg1 | 4.360975 | 5.31E-07 | 0.001146238 | 0.29388253 | 0.768847668 | 0.997423128 | 9.974231283 |
| mt-Nd4l | 1.7701625 | -1.15E-07 | 0.0010804 | -0.698139716 | 0.485089822 | 0.997423128 | 9.974231283 |
| Rps21 | 6.852249 | -7.72E-07 | 0.001108388 | -0.775782949 | 0.437877153 | 0.997423128 | 9.974231283 |
| Rps24 | 3.1835344 | -5.68E-07 | 0.001114508 | -0.643578037 | 0.519849096 | 0.997423128 | 9.974231283 |
| Trim2 | 9.635662 | 1.32E-06 | 0.001156833 | 0.538684538 | 0.590104544 | 0.997423128 | 9.974231283 |

| | | | | | | | |
|---|---|---|---|---|---|---|---|
| Tshz2 | 10.283613 | -0.144269504 | 0.112592779 | 0.06833061 | 0.945522458 | 0.997423128 | 9.974231283 |
| Ttc28 | 5.5357037 | -1.11E-07 | 0.001130179 | -0.079182225 | 0.936887683 | 0.997423128 | 9.974231283 |
| Uba52 | 1.9512028 | 2.98E-07 | 0.001173598 | 0.31482713 | 0.752892911 | 0.997423128 | 9.974231283 |
| Ubb | 1.2982892 | 6.94E-07 | 0.001248504 | 0.847661653 | 0.396626428 | 0.997423128 | 9.974231283 |
| Unc5d | 11.152448 | 1.21E-06 | 0.001162859 | 0.478225067 | 0.632490023 | 0.997423128 | 9.974231283 |
| Uqcrh | 1.8911818 | 2.56E-07 | 0.001178822 | 0.257343453 | 0.796913651 | 0.997423128 | 9.974231283 |
| Vit | 1.6428826 | -5.18E-07 | 0.001124386 | -0.503811686 | 0.614393715 | 0.997423128 | 9.974231283 |
| Vwc2l | 4.158543 | -2.92E-07 | 0.001135968 | -0.168150675 | 0.866464744 | 0.997423128 | 9.974231283 |
| Wdr72 | 7.101959 | 1.89E-06 | 0.001156683 | 0.858238164 | 0.390760966 | 0.997423128 | 9.974231283 |
| Wdyhv1 | 5.2000837 | -1.37E-06 | 0.001133034 | -0.756325086 | 0.449454311 | 0.997423128 | 9.974231283 |
| Tpt1 | 4.37262 | 4.75E-08 | 0.001148417 | 0.086910702 | 0.930742493 | 0.997423128 | 9.974231283 |
| Xlr5a | 27.32727 | 4.44E-07 | 0.001147755 | 0.121232605 | 0.903506799 | 0.997423128 | 9.974231283 |
| Zfhx3 | 3.9178238 | -3.16E-07 | 0.001137908 | -0.220416068 | 0.825547132 | 0.997423128 | 9.974231283 |
| Zfp994 | 3.931868 | -2.70E-07 | 0.001130212 | -0.222925354 | 0.823593603 | 0.997423128 | 9.974231283 |
| Zmat4 | 6.7431383 | -2.11E-06 | 0.001120757 | -1.181979029 | 0.237214022 | 0.997423128 | 9.974231283 |
| mt-Atp6 | 52.440006 | 3.97E-08 | 0.001139977 | 0.132882752 | 0.894286108 | 0.997423128 | 9.974231283 |
| mt-Co1 | 29.908287 | 3.80E-07 | 0.001147247 | 0.223285812 | 0.823313069 | 0.997423128 | 9.974231283 |
| mt-Co2 | 31.080164 | -5.13E-07 | 0.001129403 | -0.271258855 | 0.78619195 | 0.997423128 | 9.974231283 |
| mt-Cytb | 17.050423 | 1.96E-06 | 0.001176426 | 0.880741986 | 0.378457486 | 0.997423128 | 9.974231283 |
| mt-Nd1 | 13.160506 | -8.37E-07 | 0.001112122 | -0.693526761 | 0.487979038 | 0.997423128 | 9.974231283 |
| mt-Nd2 | 16.228426 | -2.86E-07 | 0.001130566 | -0.217311562 | 0.827965548 | 0.997423128 | 9.974231283 |
| mt-Nd3 | 11.394203 | -3.20E-07 | 0.00113884 | -0.226414446 | 0.820879093 | 0.997423128 | 9.974231283 |
| mt-Nd4 | 15.204892 | -1.49E-06 | 0.001108461 | -1.00099249 | 0.316830439 | 0.997423128 | 9.974231283 |
| Zbtb7a | 6.3876896 | 1.29E-07 | 0.001136664 | 0.052603048 | 0.958048188 | 0.997423128 | 9.974231283 |
| Rps23 | 1.558236 | 2.19E-07 | 0.001187402 | 0.31778266 | 0.750649805 | 0.997423128 | 9.974231283 |
| Tns3 | 2.8131933 | -8.11E-07 | 0.001121942 | -0.634292608 | 0.52588988 | 0.997423128 | 9.974231283 |
| Tmeff2 | 3.7606382 | -2.98E-07 | 0.001142632 | -0.296540304 | 0.766817492 | 0.997423128 | 9.974231283 |
| Rps27 | 2.3981893 | -3.23E-07 | 0.001142471 | -0.336159717 | 0.736750428 | 0.997423128 | 9.974231283 |
| Rps27a | 2.27826 | 7.95E-07 | 0.001185404 | 0.707949962 | 0.478976332 | 0.997423128 | 9.974231283 |
| Rps28 | 4.243435 | -1.18E-06 | 0.001104901 | -1.118309792 | 0.263434706 | 0.997423128 | 9.974231283 |
| Rps29 | 11.573063 | -6.23E-07 | 0.001087741 | -0.821581127 | 0.411315334 | 0.997423128 | 9.974231283 |
| Rps3 | 1.8366036 | -7.04E-07 | 0.001070469 | -1.225355501 | 0.220441305 | 0.997423128 | 9.974231283 |
| Rps3a1 | 6.1928663 | -4.10E-07 | 0.001059823 | -0.994600534 | 0.319930588 | 0.997423128 | 9.974231283 |
| Rps7 | 1.746049 | 4.59E-07 | 0.001264322 | 0.680998867 | 0.495872208 | 0.997423128 | 9.974231283 |
| Rps8 | 4.412556 | -9.01E-07 | 0.001058092 | -1.653225719 | 0.098284934 | 0.997423128 | 9.974231283 |
| Rsrc1 | 12.455451 | 1.63E-06 | 0.001162714 | 0.675039619 | 0.499650594 | 0.997423128 | 9.974231283 |
| Scamp1 | 4.1156707 | -2.38E-06 | 0.001118247 | -1.412614819 | 0.15776901 | 0.997423128 | 9.974231283 |
| Sec61a1 | 55.388393 | -3.85E-07 | 0.001132416 | -0.142517746 | 0.886671063 | 0.997423128 | 9.974231283 |
| Tmem108 | 3.9934008 | -1.19E-07 | 0.001135498 | -0.074676483 | 0.940472119 | 0.997423128 | 9.974231283 |
| Selenow | 2.1228044 | 8.20E-07 | 0.001173216 | 0.637209289 | 0.523988528 | 0.997423128 | 9.974231283 |
| Sgip1 | 1.58918 | -6.56E-07 | 0.001118701 | -0.803986163 | 0.421404966 | 0.997423128 | 9.974231283 |
| Shb | 15.204553 | 2.73E-07 | 0.001146799 | 0.087716651 | 0.930101885 | 0.997423128 | 9.974231283 |
| Sil1 | 7.6066775 | -5.06E-07 | 0.001137256 | -0.236572571 | 0.812988401 | 0.997423128 | 9.974231283 |
| Slx1b | 6.8200607 | 2.09E-06 | 0.001170191 | 1.016667109 | 0.309311818 | 0.997423128 | 9.974231283 |
| Spock1 | 3.860509 | -1.15E-06 | 0.001131188 | -0.837903736 | 0.40208477 | 0.997423128 | 9.974231283 |
| Srgn | 2.2238126 | -1.86E-06 | 0.00109801 | -1.739163598 | 0.08200599 | 0.997423128 | 9.974231283 |
| St18 | 2.3145986 | -1.46E-06 | 0.001101489 | -1.330104955 | 0.183483693 | 0.997423128 | 9.974231283 |
| Sult2a2 | 2.8649654 | -3.92E-07 | 0.001138226 | -0.275593391 | 0.782860386 | 0.997423128 | 9.974231283 |
| Svil | 2.9297297 | -2.81E-07 | 0.001138767 | -0.191059727 | 0.8484788 | 0.997423128 | 9.974231283 |
| Syt1 | 2.76881 | 2.18E-06 | 0.001179422 | 1.451095042 | 0.146753397 | 0.997423128 | 9.974231283 |
| Tacc1 | 1.8357242 | -0.144269504 | 0.11259278 | 0.734306836 | 0.462761753 | 0.997423128 | 9.974231283 |
| Sema3c | 24.274586 | 1.26E-06 | 0.001149265 | 0.317586537 | 0.750798588 | 0.997423128 | 9.974231283 |
| Mrc2 | 6.650044 | 1.38E-07 | 0.001147586 | 0.061728514 | 0.950779032 | 0.997423128 | 9.974231283 |
| Ms4a1 | 6.423043 | 5.26E-07 | 0.001133148 | 0.198702737 | 0.842495283 | 0.997423128 | 9.974231283 |
| Mc1r | 10.825235 | 2.48E-06 | 0.00116212 | 0.967659088 | 0.333214657 | 0.997423128 | 9.974231283 |
| Ccdc60 | 3.9716513 | -0.144269504 | 0.112592781 | 1.931633021 | 0.053404817 | 0.997423128 | 9.974231283 |
| Cdh23 | 7.7059603 | 1.94E-06 | 0.001159816 | 0.850205717 | 0.395210724 | 0.997423128 | 9.974231283 |
| Cdk8 | 4.057161 | -2.99E-07 | 0.00113881 | -0.292902451 | 0.769596715 | 0.997423128 | 9.974231283 |
| Cep128 | 2.7124422 | -0.144269504 | 0.11259278 | -0.33005911 | 0.741355299 | 0.997423128 | 9.974231283 |
| Cfap74 | 2.7922401 | -2.13E-06 | 0.001105529 | -1.665347032 | 0.09584354 | 0.997423128 | 9.974231283 |
| Chl1 | 4.161993 | -1.91E-06 | 0.001126392 | -1.25702126 | 0.208745937 | 0.997423128 | 9.974231283 |

| | | | | | | | |
|---|---|---|---|---|---|---|---|
| Chsy3 | 7.38024 | -1.58E-06 | 0.001131717 | -0.862678113 | 0.38831447 | 0.997423128 | 9.974231283 |
| Ckb | 3.2405775 | -8.67E-07 | 0.001090961 | -1.226820526 | 0.219890054 | 0.997423128 | 9.974231283 |
| Cmpk1 | 15.532138 | 5.74E-06 | 0.001162735 | 1.641455113 | 0.100702978 | 0.997423128 | 9.974231283 |
| Cmss1 | 5.8559604 | -6.24E-07 | 0.001136613 | -0.310715614 | 0.756016826 | 0.997423128 | 9.974231283 |
| Cntn3 | 2.9678428 | -0.144269504 | 0.11259278 | -1.096404971 | 0.27290159 | 0.997423128 | 9.974231283 |
| Car12 | 3.4988654 | -5.09E-07 | 0.001094046 | -0.797418411 | 0.42520807 | 0.997423128 | 9.974231283 |
| Cntnap2 | 15.640811 | -1.80E-06 | 0.001134329 | -0.726957932 | 0.467251726 | 0.997423128 | 9.974231283 |
| Cox6c | 3.8020508 | -2.82E-06 | 0.001101338 | -2.044877019 | 0.040866984 | 0.997423128 | 9.974231283 |
| Cox7c | 4.358693 | -7.88E-08 | 0.001139842 | -0.104498875 | 0.916773461 | 0.997423128 | 9.974231283 |
| Cox8a | 6.496518 | -5.36E-07 | 0.001081292 | -0.858556293 | 0.390585361 | 0.997423128 | 9.974231283 |
| Cpe | 2.8087764 | -4.42E-07 | 0.00113048 | -0.434747837 | 0.663745471 | 0.997423128 | 9.974231283 |
| Csmd3 | 7.3274617 | -0.144269504 | 0.11259278 | -0.409365708 | 0.682271302 | 0.997423128 | 9.974231283 |
| Cspp1 | 3.2586973 | 8.83E-07 | 0.001155593 | 0.580287575 | 0.561720705 | 0.997423128 | 9.974231283 |
| Cst3 | 20.57817 | -4.07E-07 | 0.001124624 | -0.305718995 | 0.759818609 | 0.997423128 | 9.974231283 |
| Ctcflos | 5.029918 | 1.15E-06 | 0.001161408 | 0.620654242 | 0.534827143 | 0.997423128 | 9.974231283 |
| Ctnna2 | 5.1470156 | -1.86E-06 | 0.001115975 | -1.275567845 | 0.202108327 | 0.997423128 | 9.974231283 |
| Ctnna3 | 7.525647 | -2.21E-06 | 0.001125607 | -1.060100122 | 0.289099052 | 0.997423128 | 9.974231283 |
| D430042O | 15.700211 | -3.36E-07 | 0.001131983 | -0.190621146 | 0.848822422 | 0.997423128 | 9.974231283 |
| Cox4i1 | 4.267222 | -2.06E-07 | 0.001132831 | -0.24067973 | 0.80980335 | 0.997423128 | 9.974231283 |
| Meg3 | 3.3725424 | 2.76E-07 | 0.001150654 | 0.264075018 | 0.791722113 | 0.997423128 | 9.974231283 |
| Camk2a | 5.2381573 | -1.24E-06 | 0.001127458 | -0.675672862 | 0.49924837 | 0.997423128 | 9.974231283 |
| Calm1 | 3.654584 | -8.54E-07 | 0.001047388 | -1.767253269 | 0.077185822 | 0.997423128 | 9.974231283 |
| 1700126G( | 9.276643 | -3.16E-06 | 0.00112862 | -1.25367432 | 0.209960405 | 0.997423128 | 9.974231283 |
| 4930402H: | 3.5811813 | -9.84E-07 | 0.001127107 | -0.817774815 | 0.413485781 | 0.997423128 | 9.974231283 |
| 4930441J1 | 3.6492968 | -1.80E-07 | 0.001141556 | -0.139750712 | 0.888856957 | 0.997423128 | 9.974231283 |
| 4930511M | 7.9351397 | 1.46E-07 | 0.001143753 | 0.039587318 | 0.968422138 | 0.997423128 | 9.974231283 |
| 4930558K( | 5.919937 | -4.08E-07 | 0.001140106 | -0.204397171 | 0.83804314 | 0.997423128 | 9.974231283 |
| 5830416I1 | 24.880907 | 6.62E-06 | 0.00115751 | 1.550433021 | 0.121037618 | 0.997423128 | 9.974231283 |
| 6030443J0 | 7.6030917 | -0.144269504 | 0.11259278 | 0.184802644 | 0.853383834 | 0.997423128 | 9.974231283 |
| 8430422H( | 8.173612 | -1.27E-06 | 0.001131583 | -0.59919523 | 0.549042704 | 0.997423128 | 9.974231283 |
| AI314278 | 3.58537 | -5.69E-07 | 0.001143558 | -0.44835774 | 0.653895036 | 0.997423128 | 9.974231283 |
| AY036118 | 14.886724 | 3.76E-07 | 0.001155603 | 0.280265056 | 0.779274158 | 0.997423128 | 9.974231283 |
| Acoxl | 4.884706 | 1.14E-06 | 0.001157491 | 0.638103423 | 0.523406358 | 0.997423128 | 9.974231283 |
| Camk1d | 9.3028755 | -3.91E-07 | 0.001127368 | -0.351371585 | 0.725309597 | 0.997423128 | 9.974231283 |
| Adamtsl1 | 11.004572 | 2.04E-06 | 0.001155775 | 0.795597071 | 0.42626627 | 0.997423128 | 9.974231283 |
| Ahdc1 | 3.5728967 | 6.92E-07 | 0.001148063 | 0.404992553 | 0.685482985 | 0.997423128 | 9.974231283 |
| Aldoc | 1.6210175 | 2.97E-07 | 0.001188676 | 0.455699402 | 0.648606152 | 0.997423128 | 9.974231283 |
| Alpl | 3.0174866 | 1.14E-06 | 0.001153287 | 0.726585234 | 0.467480077 | 0.997423128 | 9.974231283 |
| Ankrd12 | 7.1624546 | -7.07E-07 | 0.001133694 | -0.458275746 | 0.646754347 | 0.997423128 | 9.974231283 |
| Anks1b | 1.6476322 | -0.144269504 | 0.112592781 | 0.755680274 | 0.449840914 | 0.997423128 | 9.974231283 |
| Apoe | 4.915247 | -3.68E-08 | 0.001148793 | -0.038689141 | 0.969138231 | 0.997423128 | 9.974231283 |
| Arvcf | 1.1981798 | 3.11E-07 | 0.001164533 | 0.328239838 | 0.742730328 | 0.997423128 | 9.974231283 |
| Atp9b | 6.026926 | -4.48E-06 | 0.001126303 | -1.90693668 | 0.056528788 | 0.997423128 | 9.974231283 |
| Atxn7l1 | 2.1083577 | 1.99E-06 | 0.001180823 | 1.434830357 | 0.151335445 | 0.997423128 | 9.974231283 |
| Bc1 | 2.8435912 | -2.40E-06 | 0.001094643 | -1.9480071 | 0.051414114 | 0.997423128 | 9.974231283 |
| Cadm2 | 1.128733 | 2.90E-07 | 0.001171626 | 0.417800451 | 0.676093021 | 0.997423128 | 9.974231283 |
| Adk | 3.7155418 | 8.24E-08 | 0.001128394 | 0.036279468 | 0.971059521 | 0.997423128 | 9.974231283 |
| Dctd | 2.5801604 | 1.34E-06 | 0.001179777 | 0.962408366 | 0.335844513 | 0.997423128 | 9.974231283 |
| Dbi | 2.738859 | -3.98E-07 | 0.001077049 | -0.782195414 | 0.434099739 | 0.997423128 | 9.974231283 |
| Dop1b | 4.380478 | 1.13E-06 | 0.001159628 | 0.740338047 | 0.4590949 | 0.997423128 | 9.974231283 |
| Gphn | 2.9946876 | -7.30E-07 | 0.001118371 | -0.824820826 | 0.409473321 | 0.997423128 | 9.974231283 |
| Gria1 | 1.568388 | -0.144269504 | 0.11259278 | -0.863929074 | 0.387626854 | 0.997423128 | 9.974231283 |
| Gria4 | 7.6431046 | -1.32E-06 | 0.001113635 | -0.866038966 | 0.386468793 | 0.997423128 | 9.974231283 |
| Grik1 | 283.93326 | -1.41E-06 | 0.001129859 | -0.285905271 | 0.774950676 | 0.997423128 | 9.974231283 |
| Hdac8 | 3.5206144 | -0.144269504 | 0.11259278 | -0.515642649 | 0.606104017 | 0.997423128 | 9.974231283 |
| Hexb | 3.3334806 | -8.10E-07 | 0.001096657 | -1.10821742 | 0.267767929 | 0.997423128 | 9.974231283 |
| Igf2bp3 | 12.022297 | 7.38E-07 | 0.001144023 | 0.305734631 | 0.759806703 | 0.997423128 | 9.974231283 |
| Il1rapl2 | 2.7269056 | -8.77E-07 | 0.001130658 | -0.799695527 | 0.423887225 | 0.997423128 | 9.974231283 |
| Il27 | 8.782036 | 1.93E-06 | 0.001151354 | 0.746164009 | 0.455568347 | 0.997423128 | 9.974231283 |
| Kat7 | 4.219699 | -3.67E-07 | 0.001131048 | -0.146351579 | 0.88364385 | 0.997423128 | 9.974231283 |
| Kcnd2 | 2.514149 | -1.47E-06 | 0.001118461 | -1.257324205 | 0.208636263 | 0.997423128 | 9.974231283 |

| | | | | | | | |
|---|---|---|---|---|---|---|---|
| Dlg2 | 1.8535248 | -5.34E-08 | 0.001142813 | -0.056528046 | 0.954921154 | 0.997423128 | 9.974231283 |
| Kcnip4 | 2.905939 | 8.91E-07 | 0.001160812 | 0.622708777 | 0.533475914 | 0.997423128 | 9.974231283 |
| Kctd16 | 20.133091 | 5.82E-07 | 0.001143534 | 0.277587015 | 0.781329401 | 0.997423128 | 9.974231283 |
| Klhl14 | 26.143715 | -1.20E-05 | 0.001129004 | -2.34029732 | 0.019268394 | 0.997423128 | 9.974231283 |
| Klkb1 | 2.2612617 | 6.60E-07 | 0.001156825 | 0.511728717 | 0.608840885 | 0.997423128 | 9.974231283 |
| Lrp1b | 56.844112 | -2.03E-07 | 0.001135204 | -0.162191836 | 0.871154788 | 0.997423128 | 9.974231283 |
| Lrrc17 | 4.172238 | -0.144269504 | 0.11259278 | -1.514351185 | 0.129936793 | 0.997423128 | 9.974231283 |
| Lrrc4c | 2.4291487 | 1.14E-06 | 0.00117582 | 0.921876812 | 0.356592836 | 0.997423128 | 9.974231283 |
| Lsamp | 5.6331697 | 1.15E-06 | 0.001160424 | 0.672890124 | 0.501017195 | 0.997423128 | 9.974231283 |
| Lyrm1 | 2.0649328 | 6.40E-07 | 0.001159989 | 0.628257964 | 0.529834964 | 0.997423128 | 9.974231283 |
| Malat1 | 6.5285845 | -6.41E-07 | 0.00107647 | -1.015966788 | 0.309645202 | 0.997423128 | 9.974231283 |
| Mapk14 | 11.374496 | -0.144269504 | 0.11259278 | 1.262380552 | 0.206811884 | 0.997423128 | 9.974231283 |
| Mbp | 3.1925125 | -2.07E-07 | 0.001110287 | -0.331118228 | 0.740555183 | 0.997423128 | 9.974231283 |
| Kcnq5 | 2.647503 | 2.04E-06 | 0.001165949 | 1.370668606 | 0.170478286 | 0.997423128 | 9.974231283 |
| Gnaz | 13.247394 | 1.84E-06 | 0.001152549 | 0.695186758 | 0.48693827 | 0.997423128 | 9.974231283 |
| mt-Nd5 | 4.970977 | -8.72E-07 | 0.001119492 | -0.821460303 | 0.411384127 | 0.997423128 | 9.974231283 |
| Gm960 | 1.248124 | 3.41E-07 | 0.001155582 | 0.402113107 | 0.687600787 | 0.997423128 | 9.974231283 |
| Eef1a1 | 4.1184964 | 1.10E-06 | 0.00120194 | 0.916846332 | 0.35922317 | 0.997423128 | 9.974231283 |
| Esf1 | 2.2181897 | 1.13E-06 | 0.001180502 | 0.897952546 | 0.369210849 | 0.997423128 | 9.974231283 |
| Fam169a | 6.7369084 | 1.46E-06 | 0.001160245 | 0.758034359 | 0.448430414 | 0.997423128 | 9.974231283 |
| Fau | 2.4518764 | -1.46E-06 | 0.001063078 | -1.696508734 | 0.089789578 | 0.997423128 | 9.974231283 |
| Fbxl7 | 4.7657743 | 1.51E-06 | 0.001157404 | 0.760638821 | 0.446872826 | 0.997423128 | 9.974231283 |
| Fth1 | 10.972276 | -1.11E-06 | 0.001067002 | -1.477706886 | 0.139486251 | 0.997423128 | 9.974231283 |
| Fyb2 | 4.7031064 | 6.89E-08 | 0.001140522 | 0.043972713 | 0.964926155 | 0.997423128 | 9.974231283 |
| Galnt13 | 6.1900806 | 5.15E-07 | 0.001144935 | 0.264081836 | 0.79171686 | 0.997423128 | 9.974231283 |
| Gnao1 | 2.0144234 | -4.60E-07 | 0.001129852 | -0.374170717 | 0.708277309 | 0.997423128 | 9.974231283 |
| Gas5 | 0.86245275 | -0.144269504 | 0.112592781 | 0.444282476 | 0.656838369 | 0.997423128 | 9.974231283 |
| Gm10062 | 13.044163 | 1.85E-06 | 0.001161799 | 0.779549825 | 0.435655899 | 0.997423128 | 9.974231283 |
| Gm11337 | 4.2377167 | -1.19E-06 | 0.00113188 | -0.816707523 | 0.414095591 | 0.997423128 | 9.974231283 |
| Gm12216 | 27.355824 | 4.72E-06 | 0.001153138 | 1.037385584 | 0.299556194 | 0.997423128 | 9.974231283 |
| Gm14582 | 4.4432197 | -1.06E-06 | 0.001133321 | -0.507188726 | 0.612022405 | 0.997423128 | 9.974231283 |
| Gm29325 | 4.275987 | 1.54E-06 | 0.001129575 | 0.699937829 | 0.483966131 | 0.997423128 | 9.974231283 |
| Dpyd | 4.496455 | -0.144269504 | 0.112592781 | -0.378117973 | 0.705342957 | 0.997423128 | 9.974231283 |
| Gm38973 | 4.666378 | 5.35E-07 | 0.001157763 | 0.347268821 | 0.728389372 | 0.997423128 | 9.974231283 |
| Gm42418 | 88.84172 | -1.61E-07 | 0.001119985 | -0.162903797 | 0.870594181 | 0.997423128 | 9.974231283 |
| Gm49497 | 2.7023165 | -2.14E-06 | 0.001106873 | -1.636795542 | 0.101673198 | 0.997423128 | 9.974231283 |
| Gm49767 | 2.9797692 | 2.53E-07 | 0.001150512 | 0.173203656 | 0.86249135 | 0.997423128 | 9.974231283 |
| Gm50306 | 7.646293 | 2.00E-06 | 0.001181424 | 1.091118476 | 0.275220754 | 0.997423128 | 9.974231283 |
| Gm765 | 2.7776783 | -1.54E-06 | 0.001129239 | -0.362101257 | 0.717276365 | 0.997423128 | 9.974231283 |
| Epha6 | 12.959585 | -1.54E-07 | 0.001144736 | -0.069915693 | 0.944260763 | 0.997423128 | 9.974231283 |
| Ehbp1 | 10.738027 | 2.31E-06 | 0.001160214 | 0.946773149 | 0.343754387 | 0.997423128 | 9.974231283 |
| Rps16 | 2.7770374 | -1.00E-07 | 0.001139998 | 0.002698274 | 0.997847092 | 0.997847092 | 9.978470917 |
| Cox6b1 | 3.6377096 | 1.73E-08 | 0.001159983 | 0.013972316 | 0.988852067 | 0.997847092 | 9.978470917 |
| Hdac9 | 2.0922823 | 2.12E-08 | 0.001141755 | 0.019026311 | 0.984820116 | 0.997847092 | 9.978470917 |
| Hapln1 | 3.8963327 | 9.18E-09 | 0.001144333 | 0.008003891 | 0.993613887 | 0.997847092 | 9.978470917 |
| mt-Co3 | 64.31273 | -5.75E-08 | 0.001134032 | -0.029523566 | 0.976447024 | 0.997847092 | 9.978470917 |
| Tmsb4x | 4.6209717 | -9.85E-09 | 0.001146521 | -0.009917822 | 0.992086852 | 0.997847092 | 9.978470917 |

| **Glut** | baseMean | log2FoldChange | lfcSE | stat | pvalue | padj | pglobadj |
|---|---|---|---|---|---|---|---|
| Lrrc17 | 20.887321 | -1.640094203 | 0.536162375 | -3.856297822 | 0.000115117 | 0.02613161 | 0.2613161 |
| 1700054A( | 141.57812 | 7.58E-07 | 0.001131599 | 0.168657949 | 0.866065696 | 0.990729506 | 9.907295057 |
| Rpl17 | 26.852716 | -1.84E-07 | 0.001119409 | -0.166368429 | 0.867867017 | 0.990729506 | 9.907295057 |
| Rpl21 | 37.689503 | -1.45E-06 | 0.00108314 | -1.221328414 | 0.221961699 | 0.990729506 | 9.907295057 |
| Rpl23 | 26.072792 | -1.37E-06 | 0.001076595 | -1.334795205 | 0.181943383 | 0.990729506 | 9.907295057 |
| Rpl26 | 34.769566 | -1.04E-06 | 0.001093289 | -0.864537046 | 0.387292939 | 0.990729506 | 9.907295057 |
| Rpl27a | 34.65671 | -9.34E-07 | 0.001096418 | -0.77116099 | 0.440611513 | 0.990729506 | 9.907295057 |
| Rpl30 | 34.068516 | -9.19E-07 | 0.001095096 | -0.768758933 | 0.442036432 | 0.990729506 | 9.907295057 |
| Rpl32 | 26.072998 | -1.85E-06 | 0.001087119 | -1.351429704 | 0.176557824 | 0.990729506 | 9.907295057 |
| Rpl34 | 22.53576 | -1.39E-06 | 0.001096531 | -0.98242344 | 0.325891283 | 0.990729506 | 9.907295057 |
| Rpl35a | 37.172855 | -2.23E-06 | 0.001064926 | -2.01587883 | 0.043812648 | 0.990729506 | 9.907295057 |
| Rpl36 | 19.816837 | -1.77E-06 | 0.001075139 | -1.61258235 | 0.106835276 | 0.990729506 | 9.907295057 |

| | | | | | | | |
|---|---|---|---|---|---|---|---|
| Rpl37 | 58.321743 | -1.78E-06 | 0.001079807 | -1.413757326 | 0.157433167 | 0.990729506 | 9.907295057 |
| Rpl37a | 40.960964 | -9.58E-07 | 0.001091258 | -0.840463846 | 0.400648365 | 0.990729506 | 9.907295057 |
| Rpl38 | 36.311028 | -1.81E-06 | 0.001086751 | -1.303399963 | 0.192438249 | 0.990729506 | 9.907295057 |
| Rpl39 | 28.428679 | -1.17E-06 | 0.001076014 | -1.234138449 | 0.217151318 | 0.990729506 | 9.907295057 |
| Rpl41 | 37.511967 | -2.58E-06 | 0.001069691 | -2.055956345 | 0.039786724 | 0.990729506 | 9.907295057 |
| Rpl9 | 27.202162 | -2.95E-07 | 0.001114022 | -0.269337407 | 0.787670049 | 0.990729506 | 9.907295057 |
| Rplp1 | 76.570724 | -1.14E-06 | 0.00109004 | -0.934566883 | 0.350011553 | 0.990729506 | 9.907295057 |
| Rps10 | 27.12138 | -1.47E-06 | 0.001079948 | -1.310560685 | 0.190006229 | 0.990729506 | 9.907295057 |
| Rps12 | 35.98471 | -9.94E-07 | 0.001069242 | -1.246516092 | 0.212574985 | 0.990729506 | 9.907295057 |
| Rps14 | 37.294437 | -1.48E-06 | 0.001082828 | -1.240605976 | 0.214751343 | 0.990729506 | 9.907295057 |
| Rps15 | 19.44779 | -1.84E-06 | 0.001093717 | -1.258277183 | 0.20829153 | 0.990729506 | 9.907295057 |
| Rps16 | 24.292126 | -1.88E-06 | 0.001076182 | -1.603841109 | 0.108749081 | 0.990729506 | 9.907295057 |
| Rps17 | 12.002829 | -1.49E-06 | 0.001103348 | -1.019421242 | 0.308003026 | 0.990729506 | 9.907295057 |
| Rpl13a | 23.156572 | -1.58E-06 | 0.001076333 | -1.464152837 | 0.143152193 | 0.990729506 | 9.907295057 |
| Rpl13 | 48.121593 | -1.24E-06 | 0.001070328 | -1.36559604 | 0.17206579 | 0.990729506 | 9.907295057 |
| Rpl12 | 12.270738 | -4.45E-07 | 0.001112948 | -0.3876517 | 0.698273804 | 0.990729506 | 9.907295057 |
| Rpl11 | 34.706364 | -1.22E-06 | 0.001085327 | -1.082471319 | 0.279043153 | 0.990729506 | 9.907295057 |
| Mt3 | 35.23202 | -6.37E-07 | 0.001101163 | -0.56916852 | 0.569241782 | 0.990729506 | 9.907295057 |
| Nckap5 | 94.96033 | 3.88E-06 | 0.001147267 | 0.994375289 | 0.320040194 | 0.990729506 | 9.907295057 |
| Nell1 | 23.617365 | 3.88E-06 | 0.001135808 | 0.999744259 | 0.317434287 | 0.990729506 | 9.907295057 |
| Ninj2 | 17.927513 | -4.31E-06 | 0.001121697 | -1.186500605 | 0.235424663 | 0.990729506 | 9.907295057 |
| Nkain2 | 20.22035 | -5.14E-07 | 0.001129842 | -0.143685523 | 0.885748803 | 0.990729506 | 9.907295057 |
| Ntm | 60.91025 | -7.20E-07 | 0.001126451 | -0.209571208 | 0.834002353 | 0.990729506 | 9.907295057 |
| Pcca | 11.302964 | -1.53E-06 | 0.001119832 | -0.647684528 | 0.517189013 | 0.990729506 | 9.907295057 |
| Pcdh15 | 27.633442 | -3.95E-06 | 0.001124067 | 0.148617356 | 0.881855577 | 0.990729506 | 9.907295057 |
| Pcgf6 | 13.125739 | -0.144269504 | 0.112106272 | 0.519636666 | 0.603316838 | 0.990729506 | 9.907295057 |
| Pde1c | 48.006958 | 3.01E-06 | 0.001151382 | 0.973149792 | 0.33047886 | 0.990729506 | 9.907295057 |
| Pde6a | 63.212955 | 5.85E-06 | 0.001144701 | 1.23482867 | 0.216894275 | 0.990729506 | 9.907295057 |
| Rps20 | 30.235338 | -8.39E-07 | 0.0010978 | -0.717206438 | 0.473246728 | 0.990729506 | 9.907295057 |
| Phkg1 | 13.941239 | 3.46E-07 | 0.001130998 | 0.127075179 | 0.898880896 | 0.990729506 | 9.907295057 |
| Ppia | 17.00175 | -2.85E-07 | 0.001124639 | -0.203400714 | 0.838821841 | 0.990729506 | 9.907295057 |
| Ppp1r16a | 14.606436 | -6.06E-07 | 0.001127301 | -0.201947313 | 0.839957914 | 0.990729506 | 9.907295057 |
| Ppp1r9a | 15.939527 | -3.47E-06 | 0.001111203 | -1.355812963 | 0.175158686 | 0.990729506 | 9.907295057 |
| Ppp3ca | 11.9548855 | 3.30E-07 | 0.001143205 | 0.202529227 | 0.839503012 | 0.990729506 | 9.907295057 |
| Prkag2 | 18.591381 | 3.42E-06 | 0.001130876 | 0.88670174 | 0.375239502 | 0.990729506 | 9.907295057 |
| Prkg1 | 32.408478 | 9.17E-08 | 0.001129878 | 0.036374578 | 0.970983685 | 0.990729506 | 9.907295057 |
| Ptpn9 | 16.17321 | 9.92E-07 | 0.001132796 | 0.320060089 | 0.74892278 | 0.990729506 | 9.907295057 |
| Ptprd | 14.081265 | -3.49E-07 | 0.001128297 | -0.078115427 | 0.937736236 | 0.990729506 | 9.907295057 |
| Ptprg | 9.615968 | -0.144269504 | 0.112106273 | -0.193703192 | 0.846408292 | 0.990729506 | 9.907295057 |
| Rab29 | 17.81273 | 3.38E-06 | 0.001133138 | 0.922102136 | 0.356475303 | 0.990729506 | 9.907295057 |
| Rapgef4 | 48.983097 | 3.00E-06 | 0.001141179 | 0.800990452 | 0.423137173 | 0.990729506 | 9.907295057 |
| Plp1 | 11.947595 | -4.80E-07 | 0.001103444 | -0.501220949 | 0.616215633 | 0.990729506 | 9.907295057 |
| Rps21 | 69.29485 | -2.41E-06 | 0.001061963 | -2.185407805 | 0.028858959 | 0.990729506 | 9.907295057 |
| Rps23 | 25.459724 | -1.33E-06 | 0.001097979 | -0.928147112 | 0.353331264 | 0.990729506 | 9.907295057 |
| Rps24 | 50.57764 | -1.16E-06 | 0.0010735 | -1.25256516 | 0.210364001 | 0.990729506 | 9.907295057 |
| Trim2 | 42.692062 | -1.71E-07 | 0.00112949 | -0.053532702 | 0.957307476 | 0.990729506 | 9.907295057 |
| Tshz2 | 4.0018873 | 5.94E-07 | 0.001150461 | 0.487830846 | 0.625669661 | 0.990729506 | 9.907295057 |
| Ttc28 | 27.770508 | 2.69E-06 | 0.001132305 | 0.596715003 | 0.550697674 | 0.990729506 | 9.907295057 |
| Uba52 | 9.810962 | 2.77E-07 | 0.001144621 | 0.195195357 | 0.845240013 | 0.990729506 | 9.907295057 |
| Ubb | 13.352551 | -3.11E-07 | 0.001125768 | -0.206766505 | 0.836192216 | 0.990729506 | 9.907295057 |
| Unc5d | 1.6227521 | -3.61E-07 | 0.001122797 | -0.334409818 | 0.738070332 | 0.990729506 | 9.907295057 |
| Uqcrh | 17.226807 | -1.13E-06 | 0.001072448 | -1.317268293 | 0.187748709 | 0.990729506 | 9.907295057 |
| Vit | 14.305894 | 1.10E-06 | 0.001132563 | 0.377568302 | 0.705751316 | 0.990729506 | 9.907295057 |
| Vwc2l | 9.199624 | 6.31E-07 | 0.001127658 | 0.276764277 | 0.781961113 | 0.990729506 | 9.907295057 |
| Wdr72 | 23.826147 | 4.48E-06 | 0.001156512 | 1.428801235 | 0.153061371 | 0.990729506 | 9.907295057 |
| Wdyhv1 | 16.539324 | 3.93E-06 | 0.001161339 | 1.473780993 | 0.140540567 | 0.990729506 | 9.907295057 |
| Tpt1 | 28.210365 | -1.24E-06 | 0.001090559 | -1.018342282 | 0.308515321 | 0.990729506 | 9.907295057 |
| Xlr5a | 95.355095 | 2.32E-06 | 0.001141415 | 0.649263794 | 0.516167884 | 0.990729506 | 9.907295057 |
| Zfhx3 | 14.494242 | 1.55E-06 | 0.00113817 | 0.568789602 | 0.569498932 | 0.990729506 | 9.907295057 |
| Zmat4 | 23.312378 | 3.95E-06 | 0.001139529 | 1.068252173 | 0.285406776 | 0.990729506 | 9.907295057 |
| mt-Atp6 | 255.12491 | -2.89E-07 | 0.00112365 | -0.145762997 | 0.884108488 | 0.990729506 | 9.907295057 |

| | | | | | | | |
|---|---|---|---|---|---|---|---|
| mt-Co1 | 163.52351 | -1.05E-06 | 0.001110614 | -0.557081026 | 0.57747207 | 0.990729506 | 9.907295057 |
| mt-Co2 | 179.1176 | 3.32E-07 | 0.001133108 | 0.13004509 | 0.896530753 | 0.990729506 | 9.907295057 |
| mt-Co3 | 317.89905 | -2.22E-09 | 0.001130152 | 0.049676956 | 0.96037982 | 0.990729506 | 9.907295057 |
| mt-Cytb | 99.99927 | -1.22E-06 | 0.00110717 | -0.680234265 | 0.49635614 | 0.990729506 | 9.907295057 |
| mt-Nd1 | 66.879105 | 3.89E-07 | 0.001136319 | 0.176283805 | 0.860070985 | 0.990729506 | 9.907295057 |
| mt-Nd2 | 100.98584 | -1.22E-06 | 0.001115056 | -0.53683213 | 0.591383576 | 0.990729506 | 9.907295057 |
| mt-Nd3 | 53.05994 | -2.75E-07 | 0.001126287 | -0.138253529 | 0.890040051 | 0.990729506 | 9.907295057 |
| mt-Nd4 | 90.86455 | -8.64E-07 | 0.001113113 | -0.476036572 | 0.634048323 | 0.990729506 | 9.907295057 |
| Zbtb7a | 23.56369 | -2.42E-07 | 0.001127413 | -0.136634748 | 0.891319509 | 0.990729506 | 9.907295057 |
| Tns3 | 11.983182 | 2.62E-06 | 0.001153802 | 1.114540662 | 0.26504731 | 0.990729506 | 9.907295057 |
| Tmsb4x | 42.59606 | 6.19E-08 | 0.001131588 | 0.044166852 | 0.964771404 | 0.990729506 | 9.907295057 |
| Tmem108 | 19.81278 | 2.28E-06 | 0.001138146 | 0.705846834 | 0.480283393 | 0.990729506 | 9.907295057 |
| Rps27 | 24.597618 | -9.31E-07 | 0.001120802 | -0.459994272 | 0.645520332 | 0.990729506 | 9.907295057 |
| Rps27a | 33.156513 | -9.63E-07 | 0.001087839 | -0.91158167 | 0.361988975 | 0.990729506 | 9.907295057 |
| Rps28 | 25.47918 | -2.70E-06 | 0.00107573 | -1.967969297 | 0.04907157 | 0.990729506 | 9.907295057 |
| Rps29 | 86.63813 | -2.04E-06 | 0.001085257 | -1.37605134 | 0.168805739 | 0.990729506 | 9.907295057 |
| Rps3 | 19.521986 | -1.40E-06 | 0.001083309 | -1.235777847 | 0.216541152 | 0.990729506 | 9.907295057 |
| Rps3a1 | 29.006617 | 7.99E-07 | 0.001156719 | 0.514698254 | 0.606763894 | 0.990729506 | 9.907295057 |
| Rps7 | 21.615023 | -6.37E-07 | 0.001102037 | -0.583257406 | 0.559720031 | 0.990729506 | 9.907295057 |
| Rps8 | 42.907177 | -1.25E-06 | 0.001080637 | -1.168690455 | 0.242528369 | 0.990729506 | 9.907295057 |
| Rsrc1 | 41.97339 | 8.75E-07 | 0.001133967 | 0.268792075 | 0.788089692 | 0.990729506 | 9.907295057 |
| Scamp1 | 17.284771 | 9.17E-07 | 0.001132648 | 0.313804995 | 0.753669151 | 0.990729506 | 9.907295057 |
| Sec61a1 | 171.56808 | -6.98E-06 | 0.001119463 | -1.206339548 | 0.227686595 | 0.990729506 | 9.907295057 |
| Selenow | 17.175367 | -1.25E-06 | 0.001087619 | -1.100262739 | 0.271217662 | 0.990729506 | 9.907295057 |
| Sema3c | 66.135925 | 4.61E-06 | 0.001149312 | 1.171740983 | 0.241301064 | 0.990729506 | 9.907295057 |
| Sgip1 | 6.526274 | -1.23E-06 | 0.001121914 | -0.677430856 | 0.498132626 | 0.990729506 | 9.907295057 |
| Shb | 57.12811 | -3.78E-06 | 0.00112056 | -0.890734786 | 0.373071468 | 0.990729506 | 9.907295057 |
| Sil1 | 27.170454 | 2.92E-06 | 0.001144417 | 0.897758018 | 0.369314571 | 0.990729506 | 9.907295057 |
| Slx1b | 25.369318 | -9.14E-07 | 0.001124531 | -0.311084334 | 0.755736511 | 0.990729506 | 9.907295057 |
| Spock1 | 12.781663 | 7.18E-08 | 0.001129571 | 0.055258545 | 0.955932488 | 0.990729506 | 9.907295057 |
| Srgn | 6.927038 | 1.30E-06 | 0.001135822 | 0.593723007 | 0.552697392 | 0.990729506 | 9.907295057 |
| St18 | 5.592503 | 2.62E-07 | 0.001131854 | 0.16088298 | 0.872185568 | 0.990729506 | 9.907295057 |
| Sult2a2 | 10.396637 | -1.20E-06 | 0.001124803 | -0.397375468 | 0.691090603 | 0.990729506 | 9.907295057 |
| Svil | 12.132911 | 2.11E-06 | 0.00114744 | 0.892570207 | 0.37208738 | 0.990729506 | 9.907295057 |
| Syt1 | 12.191012 | 3.14E-07 | 0.001142222 | 0.201650379 | 0.840190057 | 0.990729506 | 9.907295057 |
| Tacc1 | 11.41664 | -0.144269504 | 0.112106272 | -0.557887155 | 0.576921444 | 0.990729506 | 9.907295057 |
| Tmeff2 | 11.866141 | 4.80E-06 | 0.001144321 | 1.644022409 | 0.100171574 | 0.990729506 | 9.907295057 |
| Mt1 | 35.063984 | -1.20E-06 | 0.001099176 | -0.844661171 | 0.398300045 | 0.990729506 | 9.907295057 |
| mt-Nd4l | 10.59298 | -1.91E-07 | 0.001131783 | -0.126606191 | 0.899252096 | 0.990729506 | 9.907295057 |
| Ms4a1 | 27.41386 | -2.79E-07 | 0.001127817 | -0.091993692 | 0.926703051 | 0.990729506 | 9.907295057 |
| Meg3 | 25.358404 | -8.03E-08 | 0.001127308 | -0.055452168 | 0.955778236 | 0.990729506 | 9.907295057 |
| Cdh23 | 31.393366 | 2.63E-06 | 0.001131099 | 0.548977427 | 0.583020941 | 0.990729506 | 9.907295057 |
| Cdk8 | 18.143866 | 1.65E-06 | 0.001183002 | 0.973148346 | 0.330479579 | 0.990729506 | 9.907295057 |
| Cep128 | 10.783486 | 1.94E-06 | 0.001135273 | 0.711517205 | 0.476763793 | 0.990729506 | 9.907295057 |
| Cfap74 | 13.209162 | -4.36E-06 | 0.001082908 | -2.565854521 | 0.010292198 | 0.990729506 | 9.907295057 |
| Chl1 | 13.870078 | -7.78E-07 | 0.001126174 | -0.22832003 | 0.819397454 | 0.990729506 | 9.907295057 |
| Chsy3 | 23.052742 | 8.65E-08 | 0.001128475 | 0.033668959 | 0.973141132 | 0.990729506 | 9.907295057 |
| Ckb | 26.128765 | -9.96E-07 | 0.001111098 | -0.603348294 | 0.546277012 | 0.990729506 | 9.907295057 |
| Cmpk1 | 49.48254 | 2.87E-06 | 0.001142086 | 0.768797608 | 0.442013469 | 0.990729506 | 9.907295057 |
| Cmss1 | 56.48602 | 6.74E-07 | 0.001148161 | 0.389049239 | 0.697239723 | 0.990729506 | 9.907295057 |
| Cntn3 | 12.88803 | -0.144269504 | 0.112106272 | 1.205273816 | 0.228097614 | 0.990729506 | 9.907295057 |
| Cntnap2 | 56.53396 | -1.78E-07 | 0.001130652 | -0.039343771 | 0.968616309 | 0.990729506 | 9.907295057 |
| Ccdc60 | 15.782949 | 1.57E-06 | 0.001136819 | 0.528927025 | 0.596856074 | 0.990729506 | 9.907295057 |
| Cox4i1 | 55.03479 | -1.45E-06 | 0.001067343 | -1.552986014 | 0.120426476 | 0.990729506 | 9.907295057 |
| Cox6c | 36.238724 | -3.51E-07 | 0.001112302 | -0.310547867 | 0.756144365 | 0.990729506 | 9.907295057 |
| Cox7c | 24.603357 | -1.52E-06 | 0.001092577 | -1.116768558 | 0.264093296 | 0.990729506 | 9.907295057 |
| Cox8a | 54.110287 | -1.09E-06 | 0.001074343 | -1.199030094 | 0.230516244 | 0.990729506 | 9.907295057 |
| Cpe | 18.960005 | 1.20E-06 | 0.001166701 | 0.707463781 | 0.479278313 | 0.990729506 | 9.907295057 |
| Csmd3 | 26.341698 | -1.70E-06 | 0.001125692 | -0.451581434 | 0.651570548 | 0.990729506 | 9.907295057 |
| Cspp1 | 15.578925 | -2.70E-06 | 0.001113257 | -1.098222404 | 0.272107385 | 0.990729506 | 9.907295057 |
| Cst3 | 153.94177 | -8.34E-08 | 0.001123477 | -0.074331957 | 0.940746249 | 0.990729506 | 9.907295057 |

| | | | | | | | |
|---|---|---|---|---|---|---|---|
| Ctcflos | 16.329626 | 9.27E-08 | 0.00113019 | 0.045416241 | 0.963775536 | 0.990729506 | 9.907295057 |
| Ctnna2 | 13.414483 | -2.63E-07 | 0.001127264 | -0.10515543 | 0.916252476 | 0.990729506 | 9.907295057 |
| Ctnna3 | 24.259981 | -4.32E-06 | 0.001114709 | -1.371357394 | 0.170263572 | 0.990729506 | 9.907295057 |
| D430042O | 59.267593 | 4.57E-06 | 0.001135167 | 0.792959534 | 0.427801399 | 0.990729506 | 9.907295057 |
| Cox6b1 | 15.940308 | -1.65E-07 | 0.001119205 | -0.170452866 | 0.864653999 | 0.990729506 | 9.907295057 |
| Car12 | 13.269348 | -1.43E-06 | 0.001076647 | -1.45617687 | 0.145343732 | 0.990729506 | 9.907295057 |
| Camk2a | 20.71487 | 2.76E-06 | 0.001132427 | 0.69406162 | 0.487643566 | 0.990729506 | 9.907295057 |
| Camk1d | 70.181915 | -2.09E-07 | 0.001119474 | -0.168405468 | 0.866264306 | 0.990729506 | 9.907295057 |
| 1700126G( | 34.365047 | 1.46E-06 | 0.001132531 | 0.334881212 | 0.737714695 | 0.990729506 | 9.907295057 |
| 4930402H2 | 10.878186 | -3.16E-07 | 0.001129054 | 0.04067825 | 0.967552401 | 0.990729506 | 9.907295057 |
| 4930441J1 | 14.087247 | 2.77E-06 | 0.00114742 | 1.053576998 | 0.292076625 | 0.990729506 | 9.907295057 |
| 4930511M | 35.85655 | -3.61E-06 | 0.001121159 | -0.898461646 | 0.368939484 | 0.990729506 | 9.907295057 |
| 4930558K( | 19.8751 | 4.25E-06 | 0.00114949 | 1.352564574 | 0.176194776 | 0.990729506 | 9.907295057 |
| 5830416I1 | 109.243774 | -3.65E-07 | 0.00112708 | -0.097873458 | 0.922032776 | 0.990729506 | 9.907295057 |
| 6030443J0 | 30.79876 | -4.52E-06 | 0.001117502 | -1.252769938 | 0.210289445 | 0.990729506 | 9.907295057 |
| 8430422H( | 42.030968 | 3.01E-06 | 0.001145887 | 0.901850504 | 0.367136287 | 0.990729506 | 9.907295057 |
| AI314278 | 10.483916 | -0.144269504 | 0.112106272 | -0.593967572 | 0.552533802 | 0.990729506 | 9.907295057 |
| AY036118 | 96.25058 | -6.30E-09 | 0.001125308 | -0.064768222 | 0.948358543 | 0.990729506 | 9.907295057 |
| Acoxl | 21.082848 | 3.70E-06 | 0.001146516 | 1.163791138 | 0.244508664 | 0.990729506 | 9.907295057 |
| Adamtsl1 | 32.24669 | 4.49E-07 | 0.001131009 | 0.130705538 | 0.896008251 | 0.990729506 | 9.907295057 |
| Adk | 15.99909 | 2.65E-06 | 0.001131689 | 0.759296708 | 0.447675086 | 0.990729506 | 9.907295057 |
| Ahdc1 | 11.812328 | 6.88E-07 | 0.00113255 | 0.258656381 | 0.795900378 | 0.990729506 | 9.907295057 |
| Aldoc | 15.485122 | -1.15E-06 | 0.001092759 | -0.985074826 | 0.324587318 | 0.990729506 | 9.907295057 |
| Alpl | 10.492561 | -1.40E-06 | 0.001121976 | -0.608335793 | 0.542964786 | 0.990729506 | 9.907295057 |
| Ankrd12 | 21.009596 | 4.41E-06 | 0.001153677 | 1.4006431 | 0.161320826 | 0.990729506 | 9.907295057 |
| Anks1b | 9.166362 | 7.45E-07 | 0.001144402 | 0.381929553 | 0.702513619 | 0.990729506 | 9.907295057 |
| Apoe | 80.2238 | -3.90E-07 | 0.001117663 | -0.258530864 | 0.795997232 | 0.990729506 | 9.907295057 |
| Arvcf | 7.0802913 | -1.72E-06 | 0.00112239 | -0.871342094 | 0.383567393 | 0.990729506 | 9.907295057 |
| Atp9b | 22.188927 | 6.45E-07 | 0.001136096 | 0.270530769 | 0.786751949 | 0.990729506 | 9.907295057 |
| Atxn7l1 | 12.299738 | -1.34E-06 | 0.001122516 | -0.540459736 | 0.588880022 | 0.990729506 | 9.907295057 |
| Bc1 | 16.326283 | -3.64E-06 | 0.001075111 | -2.44549861 | 0.014465201 | 0.990729506 | 9.907295057 |
| Cadm2 | 11.891537 | -5.28E-07 | 0.001126817 | -0.288257722 | 0.773149467 | 0.990729506 | 9.907295057 |
| Calm1 | 23.933855 | 6.25E-07 | 0.001147684 | 0.359697119 | 0.719073647 | 0.990729506 | 9.907295057 |
| Dbi | 18.818296 | -1.53E-06 | 0.001100891 | -0.997041495 | 0.318744369 | 0.990729506 | 9.907295057 |
| Mrc2 | 15.52196 | 1.11E-06 | 0.001137475 | 0.432135848 | 0.665642682 | 0.990729506 | 9.907295057 |
| Dctd | 7.827331 | 5.43E-07 | 0.001129426 | 0.226502856 | 0.820810338 | 0.990729506 | 9.907295057 |
| Dop1b | 23.20367 | 5.62E-06 | 0.001155915 | 1.686740408 | 0.091653278 | 0.990729506 | 9.907295057 |
| Gria4 | 27.528088 | -1.24E-07 | 0.001129059 | -0.033505203 | 0.973271717 | 0.990729506 | 9.907295057 |
| Grik1 | 5.4989185 | 2.34E-07 | 0.001134123 | 0.170748925 | 0.864421191 | 0.990729506 | 9.907295057 |
| Hapln1 | 12.383218 | -2.89E-06 | 0.001104422 | -1.382910767 | 0.166692228 | 0.990729506 | 9.907295057 |
| Hdac8 | 13.568848 | 7.72E-07 | 0.001135378 | 0.345097003 | 0.730021447 | 0.990729506 | 9.907295057 |
| Hdac9 | 9.57025 | -2.68E-07 | 0.001128651 | -0.074310443 | 0.940763367 | 0.990729506 | 9.907295057 |
| Hexb | 15.088464 | 6.37E-07 | 0.001164819 | 0.501030922 | 0.616349361 | 0.990729506 | 9.907295057 |
| Igf2bp3 | 45.83929 | -3.49E-07 | 0.001125674 | -0.142674801 | 0.886547019 | 0.990729506 | 9.907295057 |
| Il1rapl2 | 9.701231 | -0.144269504 | 0.112106272 | -0.992443346 | 0.320981303 | 0.990729506 | 9.907295057 |
| Il27 | 31.86366 | 3.82E-06 | 0.001146024 | 1.080510493 | 0.279914917 | 0.990729506 | 9.907295057 |
| Kat7 | 14.177729 | -7.95E-07 | 0.001127274 | -0.277392278 | 0.78147891 | 0.990729506 | 9.907295057 |
| Kcnd2 | 8.189679 | 1.03E-06 | 0.001130637 | 0.402448927 | 0.687353668 | 0.990729506 | 9.907295057 |
| Kcnip4 | 17.83586 | -7.47E-07 | 0.001126784 | -0.273128842 | 0.784754176 | 0.990729506 | 9.907295057 |
| Kcnq5 | 13.298319 | -8.01E-07 | 0.001124678 | -0.34993765 | 0.726385491 | 0.990729506 | 9.907295057 |
| Kctd16 | 76.70161 | 1.24E-06 | 0.001132023 | 0.22963879 | 0.818372463 | 0.990729506 | 9.907295057 |
| Klhl14 | 2.6490629 | 1.03E-06 | 0.001165053 | 0.846531872 | 0.397256102 | 0.990729506 | 9.907295057 |
| Klkb1 | 12.379919 | 1.60E-06 | 0.001141979 | 0.655397361 | 0.51221195 | 0.990729506 | 9.907295057 |
| Lrp1b | 204.00333 | 4.08E-06 | 0.001142454 | 0.870975734 | 0.383767403 | 0.990729506 | 9.907295057 |
| Lrrc4c | 18.606462 | -1.10E-06 | 0.001112405 | -0.649910158 | 0.515750256 | 0.990729506 | 9.907295057 |
| Lsamp | 27.76111 | 1.97E-06 | 0.001134212 | 0.492386505 | 0.622446138 | 0.990729506 | 9.907295057 |
| Lyrm1 | 7.9197645 | -1.94E-06 | 0.001121921 | -0.838830867 | 0.401564227 | 0.990729506 | 9.907295057 |
| Malat1 | 69.80103 | 1.31E-07 | 0.001131612 | 0.059599014 | 0.952475004 | 0.990729506 | 9.907295057 |
| Mapk14 | 41.72899 | 5.67E-06 | 0.001141449 | 1.243125549 | 0.213821572 | 0.990729506 | 9.907295057 |
| Mc1r | 36.4037 | -9.04E-07 | 0.001123293 | -0.308496896 | 0.757704262 | 0.990729506 | 9.907295057 |
| Gphn | 15.688767 | -1.17E-07 | 0.00113027 | -0.081906289 | 0.934721233 | 0.990729506 | 9.907295057 |

| | | | | | | | |
|---|---|---|---|---|---|---|---|
| Gnaz | 44.17836 | 3.77E-06 | 0.001148626 | 1.071487053 | 0.283950489 | 0.990729506 | 9.907295057 |
| Gria1 | 10.882296 | -0.144269504 | 0.112106272 | -1.019056175 | 0.308176298 | 0.990729506 | 9.907295057 |
| Gm960 | 8.686207 | -2.22E-06 | 0.001121838 | -0.912550559 | 0.361478967 | 0.990729506 | 9.907295057 |
| Dpyd | 15.7993355 | -1.07E-06 | 0.001123558 | -0.400879377 | 0.688508934 | 0.990729506 | 9.907295057 |
| Eef1a1 | 22.886635 | -7.97E-07 | 0.001114819 | -0.499895197 | 0.617148875 | 0.990729506 | 9.907295057 |
| Ehbp1 | 37.233177 | 2.48E-06 | 0.001136601 | 0.616035953 | 0.537870791 | 0.990729506 | 9.907295057 |
| Epha6 | 38.557938 | 1.59E-06 | 0.001139052 | 0.497207397 | 0.619042806 | 0.990729506 | 9.907295057 |
| Esf1 | 9.794302 | 7.80E-07 | 0.001134066 | 0.333990471 | 0.73838675 | 0.990729506 | 9.907295057 |
| Fam169a | 27.947054 | 7.64E-06 | 0.001139686 | 1.735546717 | 0.082644014 | 0.990729506 | 9.907295057 |
| Fau | 36.68445 | -1.04E-06 | 0.00106394 | -1.38904136 | 0.164820174 | 0.990729506 | 9.907295057 |
| Fbxl7 | 16.057865 | 1.39E-06 | 0.001144408 | 0.609630617 | 0.542106525 | 0.990729506 | 9.907295057 |
| Fth1 | 159.25063 | -7.45E-07 | 0.001101716 | -0.58805617 | 0.556494591 | 0.990729506 | 9.907295057 |
| Fyb2 | 22.179024 | -0.144269504 | 0.112106273 | 1.538888441 | 0.123831534 | 0.990729506 | 9.907295057 |
| Galnt13 | 29.213446 | -4.94E-07 | 0.001127793 | -0.161783118 | 0.871476648 | 0.990729506 | 9.907295057 |
| Gnao1 | 10.703454 | 1.74E-06 | 0.001147074 | 0.757518507 | 0.448739283 | 0.990729506 | 9.907295057 |
| Gas5 | 23.922663 | 8.76E-08 | 0.001135753 | 0.073726812 | 0.941227764 | 0.990729506 | 9.907295057 |
| Gm11337 | 13.186041 | 1.61E-06 | 0.001140646 | 0.644140829 | 0.519484116 | 0.990729506 | 9.907295057 |
| Gm12216 | 92.376305 | 4.70E-06 | 0.001139701 | 0.898974156 | 0.368666427 | 0.990729506 | 9.907295057 |
| Gm14582 | 12.835498 | 1.02E-06 | 0.001137426 | 0.423949712 | 0.671602488 | 0.990729506 | 9.907295057 |
| Gm29325 | 14.84829 | -3.86E-06 | 0.001123841 | -1.039289071 | 0.298670316 | 0.990729506 | 9.907295057 |
| Gm38973 | 16.28398 | 2.04E-06 | 0.00114612 | 0.796079101 | 0.425986061 | 0.990729506 | 9.907295057 |
| Gm42418 | 495.07285 | 3.37E-07 | 0.001134862 | 0.171272313 | 0.86400965 | 0.990729506 | 9.907295057 |
| Gm49497 | 13.54824 | 5.93E-07 | 0.001135464 | 0.263373563 | 0.792262665 | 0.990729506 | 9.907295057 |
| Gm49767 | 11.525636 | 7.91E-08 | 0.001130974 | 0.037116549 | 0.970392077 | 0.990729506 | 9.907295057 |
| Gm50306 | 32.413666 | 2.57E-07 | 0.001129265 | 0.048374461 | 0.961417813 | 0.990729506 | 9.907295057 |
| Gm765 | 13.4760685 | 2.36E-06 | 0.001128507 | 0.667569335 | 0.504408538 | 0.990729506 | 9.907295057 |
| Gm10062 | 59.852055 | 2.96E-06 | 0.001146983 | 0.874319108 | 0.381944496 | 0.990729506 | 9.907295057 |
| mt-Nd5 | 23.92398 | 3.79E-07 | 0.001143395 | 0.239164622 | 0.810977934 | 0.990729506 | 9.907295057 |
| Ralyl | 14.507205 | 4.05E-08 | 0.001134361 | 0.020364759 | 0.983752396 | 0.996238136 | 9.962381356 |
| Mbp | 64.762535 | 2.17E-08 | 0.001130254 | 0.01571632 | 0.987460707 | 0.996238136 | 9.962381356 |
| Zfp994 | 14.944137 | -0.144269504 | 0.112106271 | -0.004427104 | 0.996467694 | 0.997053692 | 9.970536918 |
| Dlg2 | 14.809554 | 6.39E-09 | 0.001133226 | 0.003692658 | 0.997053692 | 0.997053692 | 9.970536918 |
